# Melanoma evolution in the lymph node shapes systemic outcomes

**DOI:** 10.64898/2026.09.15.751777

**Authors:** Katherine S. Ventre, Tara Muijlwijk, Robert Stagnitta, Shi Qiu, Ze Chen, Irineu Illa Bochaca, Ata S. Moshiri, Teresa Davoli, Iman Osman, Itai Yanai, Markus Schober, Amanda W. Lund

**Affiliations:** The Ronald O. Perelman Department of Dermatology, NYU Grossman School of Medicine, New York, New York; Department of Pathology, NYU Grossman School of Medicine, New York, New York 10016; Institute for Systems Genetics, NYU Grossman School of Medicine, New York, New York, USA 10016; Department of Biochemistry and Molecular Pharmacology, NYU Grossman School of Medicine, New York, New York, USA 10016; Laura and Isaac Perlmutter Cancer Center, NYU Grossman School of Medicine, New York, New York 10016

**Keywords:** melanoma, lymph node metastasis, tumor evolution, intratumoral heterogeneity, neural crest, immune evasion, chromosomal instability, aneuploidy

## Abstract

Lymph node (LN) metastasis predicts poor patient outcomes, but the mechanistic drivers that shape metastatic fitness, immune evasion, and clinical impact remain elusive. While preclinical models indicate that active tumor adaptation is necessary for LN metastasis, observations of clonal heterogeneity in human tumors has supported a stochastic model of passive and continuous seeding. Reconciling LN metastasis as a passive or active process is essential to understanding if LN metastasis is simply a marker of disease progression or a clinically informative therapeutic target. Here, we report evidence that LNs are active niches that facilitate ongoing melanoma evolution to progressively subvert immune surveillance and enable progression. To construct a spatial trajectory of LN metastasis, we examined paired primary melanomas and metastatic sentinel LNs through integrated genomic, phenotypic, and immunologic analyses. In contrast to a model of continuous seeding, we observe that early dissemination from the primary tumor is followed by extensive intra-nodal diversification, indicating that metastatic outgrowth requires ongoing adaptation within the LN. As clones evolve in the LN, they re-differentiate towards a melanocytic state and reprogram the microenvironment for immune exclusion. In further evolved clones, loss of inflammatory interferon signaling and induction of p53 and mitochondrial stress are associated with decreased overall survival. Collectively, these results implicate the LN as a critical battleground for melanoma progression, where tumor evolution drives adaptation and immune escape to biologically link regional metastasis to patient survival.

## INTRODUCTION

Lymph node (LN) metastasis is a strong adverse prognostic feature across many solid cancers, including melanoma^1^, where it defines stage III regionally advanced disease^2^. Yet, its causal contribution to systemic progression remains unclear. Despite evidence for sequential metastasis in preclinical studies^3–5^, the limited clonal overlap between LN and distant metastases in melanoma^6–8^, breast^9,10^, and colorectal^11,12^ cancer argues against LN involvement as a prerequisite for hematogenous dissemination. This has led to an alternative model, in which LN metastasis promotes systemic tolerance by suppressing anti-tumor immune surveillance^13^, thereby permitting the outgrowth of disseminated tumor cells at distant sites^14,15^. Distinguishing whether LN metastasis is simply a marker of aggressive disease or an active contributor to systemic progression will define its importance to regional disease management in the era of modern cancer therapy.

Refining the clinical utility of LN metastasis, however, depends on a mechanistic understanding of how tumor cells colonize LN tissue and how local tumor outgrowth affects LN function. Genomic studies in colorectal cancer^11,16^ report that LN metastases are more clonally heterogeneous than distant metastases, a finding interpreted as evidence for continuous, stochastic, and thereby passive, seeding from a heterogeneous primary tumor. Consistent with a low barrier to LN tumor formation, experimental tumors in mice are poorly surveilled by cytotoxic CD8^+^ T cells^17^, together indicating that LNs are intrinsically susceptible to tumor colonization. However, emerging preclinical data demonstrates that tumor cells undergo metabolic^4,18,19^ and immunologic^14,20,21^ adaptation within the LN to facilitate metastatic outgrowth and local and systemic immune suppression. These observations motivate a more active model of LN metastasis, in which multiple features of the LN environment present barriers to the establishment of LN lesions. If validated, these adaptive bottlenecks could serve as mechanistically informed biomarkers, linking regional melanoma progression to systemic outcomes, and as potential points for therapeutic interception. However, reconciling these disparate observations into a unifying model of LN metastasis has remained challenging, in part because preclinical models rarely capture the genomic diversity seen in patients, and high-resolution paired analyses of primary tumors and metastatic LNs from patients have remained limited.

Here, we reconstruct the genomic, transcriptomic, and immunologic adaptations that accompany LN colonization through an integrated multi-omic spatial analyses of paired primary melanomas and metastatic sentinel LNs. We show that melanoma dissemination occurs early, but successful LN colonization is accompanied by substantial intranodal genomic diversification and cell state adaptation. These changes are associated with the emergence of dense, immune-excluded LN lesions and with poor patient survival. Our findings reconcile nodal clonal diversity with an active model of LN adaptation, identify LN metastases as critical sites of early melanoma evolution, and define a biological framework integrating evolution, phenotype, and clinical outcomes, which may inform higher resolution prognostic or therapeutic use of LN metastasis.

## RESULTS

### Genomic diversification of LN metastases

To map the evolution and adaptation of melanoma cells between primary tumors and LN metastases, we collected a cohort of eight paired primary tumors and metastatic sentinel LNs (SLN), plus four unpaired SLNs, from patients with stage III cutaneous (n = 10) and acral (n = 2) melanoma **(Fig. 1a and Supplementary Table 1)**. The patients were predominantly male with a median age of 62 years, consistent with national melanoma patient demographics^22^, and were treatment-naïve at the time of surgery. Paired specimens were collected concurrently and profiled using Visium 10X spatial transcriptomics (ST) and Akoya Fusion high-dimensional whole-slide imaging^23^ on sequential sections. Primary and LN tumors were of varying sizes **(Fig. 1b-d and Supplementary Table 2)**. Tumor regions of interest for ST were selected using a tumor detection algorithm on adjacent slides^24,25^. Malignant ST spots were identified by reference-free SpatialInferCNV^26,27^, and annotations were validated with H&E and Fusion imaging **(Extended Data Fig. 1a,b)**.

**Fig. 1.**
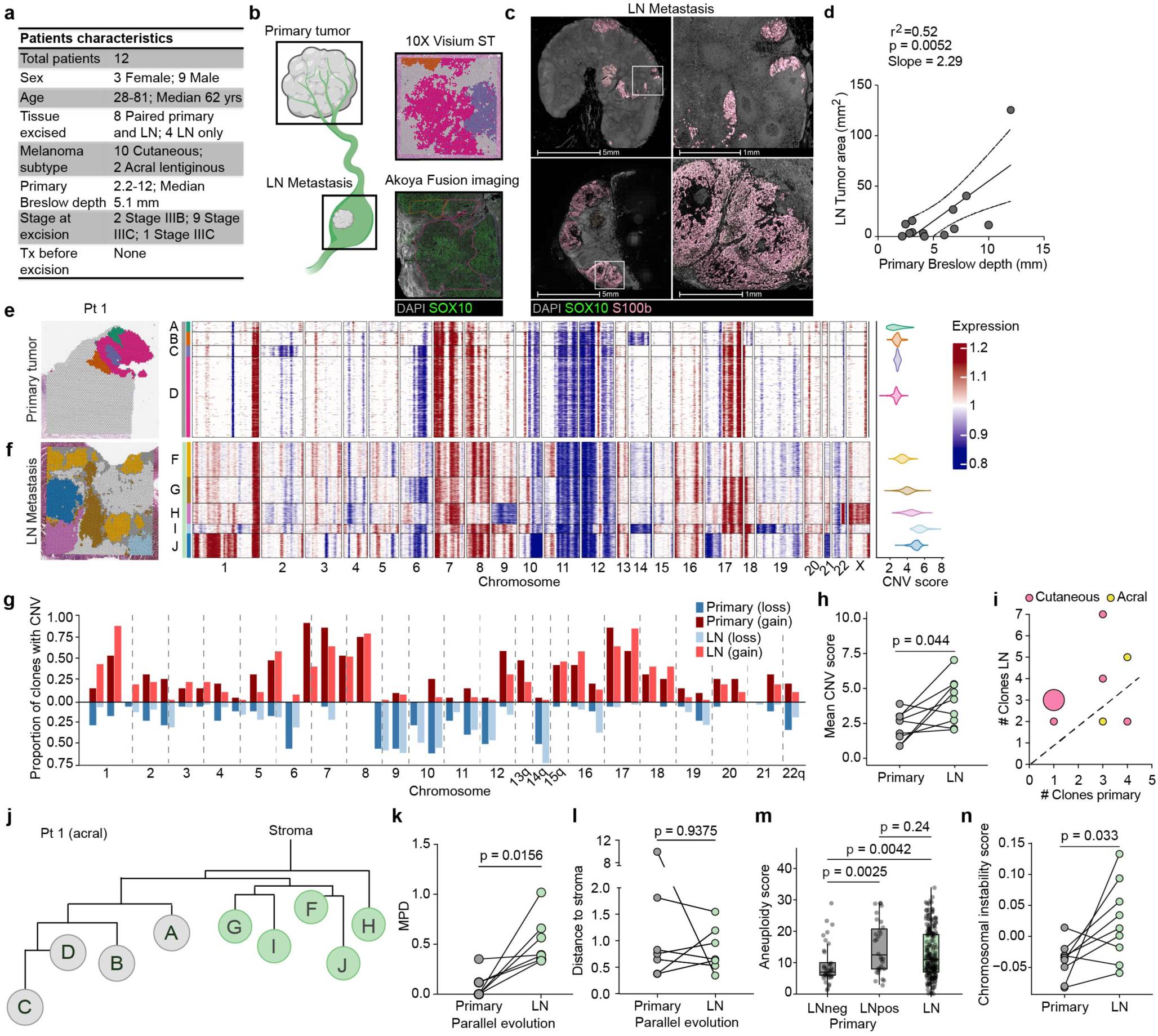
| Melanoma genomically diversifies during LN metastasis. **a**, Table of patient characteristics. **b,** Schematic depicting complementary ST and Akoya Fusion imaging on paired primary and LN melanomas. **c,** Representative images of small and large LN metastases. Scale bar, 5 mm (left), 1 mm (right) **d,** Linear regression of primary Breslow depth at excision and LN tumor area (n = 13 LN from 12 patients). **e-f,** Spatial map with CNV clone identities (left), spot-level gene expression heatmaps normalized to stroma and clustered by CNV clone (middle), violin plot of clone CNV score (right) for representative paired primary tumor **(e)** and metastatic LN **(f)**. **g,** Proportion of primary (n = 18) and LN (n = 33) clones with chromosome arm gains (red) or losses (blue). **h,** Mean CNV score within primary and LN tumors (n = 9 pairs). **i,** Number of clones identified in primary tumors (n = 8) and their paired metastatic LN (n = 9 from 8 patients). Spot size scaled to sample number. **j,** Maximum likelihood phylogenetic tree for a representative patient with parallel evolution. Branch lengths represent evolutionary distances between clones. **k-l,** Mean pairwise distance (MPD) between clones **(k)** and mean estimated distance to stroma per patient **(l)**, within primary and LN tumors from patients with parallel evolution (n = 7 pairs). **m,** Aneuploidy score in cutaneous melanoma primary tumors without LN metastasis (n = 38), primary tumors with LN metastasis (n = 30), and LN metastases (n = 201) from The Cancer Genome Atlas (TCGA). **n,** Chromosomal instability signature score within primary and LN tumors (n = 9 pairs). Statistics: paired two-tailed Wilcoxon test **(h, k, l, n)**; Kruskal-Wallis test (**m**, p = 0.0044) with pairwise two-tailed Mann-Whitney U test.

We first resolved the clonal architecture of each lesion by SpatialInferCNV^26,27^ **(Fig. 1e,f, Extended Data Fig.1c,d, and Supplementary Table 3 and 4)**. We observed substantial clonal heterogeneity **(Fig. 1e,f)**, with individual clones highly clustered in distinct spatial regions in both tumor sites **(Extended Data Fig. 1e,f)**, consistent with local outgrowth of unique populations. Comparison of arm-level copy number variation (CNV) distributions between primary and LN tumors revealed a broad distribution of CNVs in both sites **(Fig. 1g)**. While gains in chromosome 7p were enriched in primary tumors **(Extended Data Fig. 1g,h)**, no specific CNVs were exclusive to LN metastases. In most cases, CNV scores **(Fig. 1h)** and the number of unique clones **(Fig. 1i)** increased from primary tumor to LN metastasis, indicating greater clonal heterogeneity within the LN compared to its paired primary tumor independent of melanoma subtype **(Fig. 1i)**.

To determine the evolutionary relationships between clones in primary tumors and LN metastases, we constructed a maximum likelihood phylogenetic tree for each patient **(Fig. 1j and Extended Data Fig. 2, 3)**. In the six patients with more clonally diverse LN metastases than primary tumors, tumor progression followed a pattern of early dissemination relative to primary tumor clones and parallel genomic evolution in both sites. LN clones branched early from the stroma and were not directly descendent from observed primary clones **(Fig. 1j and Extended Data Fig. 2)**, though descent from unsampled ancestral populations cannot be excluded due to limits of detection and tissue sampling. Across these six patients, LN clones were more diverse from each other than the primary clones were (**Fig. 1k**), though clones in both sites had evolved to a similar extent from the diploid stroma (**Fig. 1l**). The two remaining patients exhibited diversification within the primary tumor, with LN clones emerging in a more linear fashion from common branches of existing primary clones (**Extended Data Fig. 3**). In these patients, LN clones were more evolved from stroma than their primary counterparts **(Extended Data Fig. 3c)** and relatively similar to each other genomically **(Extended Data Fig. 3d)**.

The high CNV score and marked clonal heterogeneity observed across this metastatic cohort prompted us to ask whether aneuploidy and chromosomal instability (CIN) are associated with regional dissemination. To test this, we analyzed primary melanomas in the TCGA SKCM cohort and found that tumors from patients presenting with regional LN metastasis had significantly higher aneuploidy scores than primaries without nodal involvement. This elevated aneuploidy persisted in LN metastases (**Fig. 1m**). Consistent with this, LN lesions in our paired samples exhibited increased expression of a CIN-associated score relative to paired primary tumors (**Fig. 1n**). These findings aligned with the dominant pattern of clonal diversification observed within the LN, linking nodal outgrowth to ongoing genomic instability. However, despite this increase in aneuploidy and CIN, we did not detect recurrent enrichment of specific CNVs in LN metastases relative to paired primary tumors. Together, these data suggest that successful LN colonization is associated less with the selection of a common nodal driver event than with general genomic plasticity, which may enable tumor cells to access adaptive phenotypic states in response to selective pressures from the LN microenvironment.

### Genotypic variation overlaps with but does not define transcriptional and immunological heterogeneity

Aneuploid cells diversify under selection pressure, where genomic instability offers the cell more flexibility in its search for new phenotypic states despite consequences for steady-state fitness^28^. Melanoma is a transcriptionally plastic tumor^29^, with heterogeneous cell states that emerge with progression and in response to therapy^30^, endowing the tumor with invasive and resistant properties. These cell states describe both continuums of differentiation (‘identity’ states) and functional environmental responses (‘adaptive’ states)^30^. However, it remains unclear to what extent these states emerge within LN metastases. To interrogate site-specific intratumoral heterogeneity and lineage plasticity in the spatial context, we performed non-negative matrix factorization (NMF) on primary and LN tumor spots and identified 25 factors that clustered into 12 metaprograms (MPs) **(Extended Data Fig. 4a)**. Given the resolution of the ST platform, these programs captured the colocalization of tumor intrinsic and extrinsic features and were defined by signatures of lipid metabolism (MP1, MP10), DNA damage and replicative stress (MP2, MP3, MP4, MP10), cytoskeleton remodeling (MP6, MP8), lysosomal activation (MP9, MP10, MP11), extracellular matrix production (MP3, MP7), cell adhesion (MP9, MP12), and interferon (IFN) response (MP1, MP12) **(Supplementary Table 5)**. All NMF factors and metaprograms were expressed in both primary and LN tumors **(Extended Data Fig. 4b)**, and only two metaprograms showed differential enrichment between sites: MP5, a signature of immune responses, and MP7, a metaprogram high for fibrotic and neural crest-like programs **(Extended Data Fig. 4c,d)**.

To map established identity and adaptive cell states onto the metaprograms, we generated consensus signatures based on prior literature for three identity states – melanocytic, mesenchymal, and neural crest-like^29,31–38^ **(Extended Data Fig. 4e and Supplementary Table 5)** – and six pan-cancer adaptive states, including proliferative^32^, p53 replicative stress^32^, hypoxic stress^32,33^, mitochondrial stress^32^, ferroptotic stress^39–41^, and IFN response^32,42^ **(Extended Data Fig. 4f and Supplementary Table 5)**. Using these consensus signatures, we deconvolved the relative fraction of tumor cells in each state per spot using the single-cell RNA-sequencing (scRNA-seq) reference dataset from Tirosh et al^34^ **(Extended Data Fig. 4g,h)**. Each metaprogram was enriched with a distinct combination of these cell states, indicating correlations between specific identity and adaptive states at the spot level (**Fig. 2a**). Direct evaluation of the colocalization of states across LN metastases revealed that melanocytic cells colocalized with signatures of mitosis and p53 replicative stress, while neural crest-like and mesenchymal cells colocalized with ferroptosis, hypoxic stress, and IFN response signatures **(Fig. 2b)**, consistent with established behavior of these differentiation states^30^. Importantly, this inferred intratumoral heterogeneity was readily captured at single cell resolution by immunofluorescence imaging **(Fig. 2c,d and Extended Data Fig. 4i,j)**, with differentiated melanocytic (GP100^+^) and de-differentiated neural crest-like (AQP1^+^ or NGFR^+^) tumor cells readily observed across LN metastases **(Fig. 2e)**. Consistent with the transcriptional data, GP100^+^ melanocytic cells were more likely to stain positive for the proliferative marker Ki-67 **(Fig. 2c and Extended Data Fig. 4k)**, while neural crest-like cells and remaining ‘unclassified’ tumor cells (S100b^+^GP100^−^ AQP1^−^NGFR^−^) expressed markers of immune activation (HLA-DR, PD-L1, Axl) and CSPG4 **(Fig. 2c)**, a melanoma-associated antigen that is highly expressed on early LN metastases^43^.

**Fig. 2.**
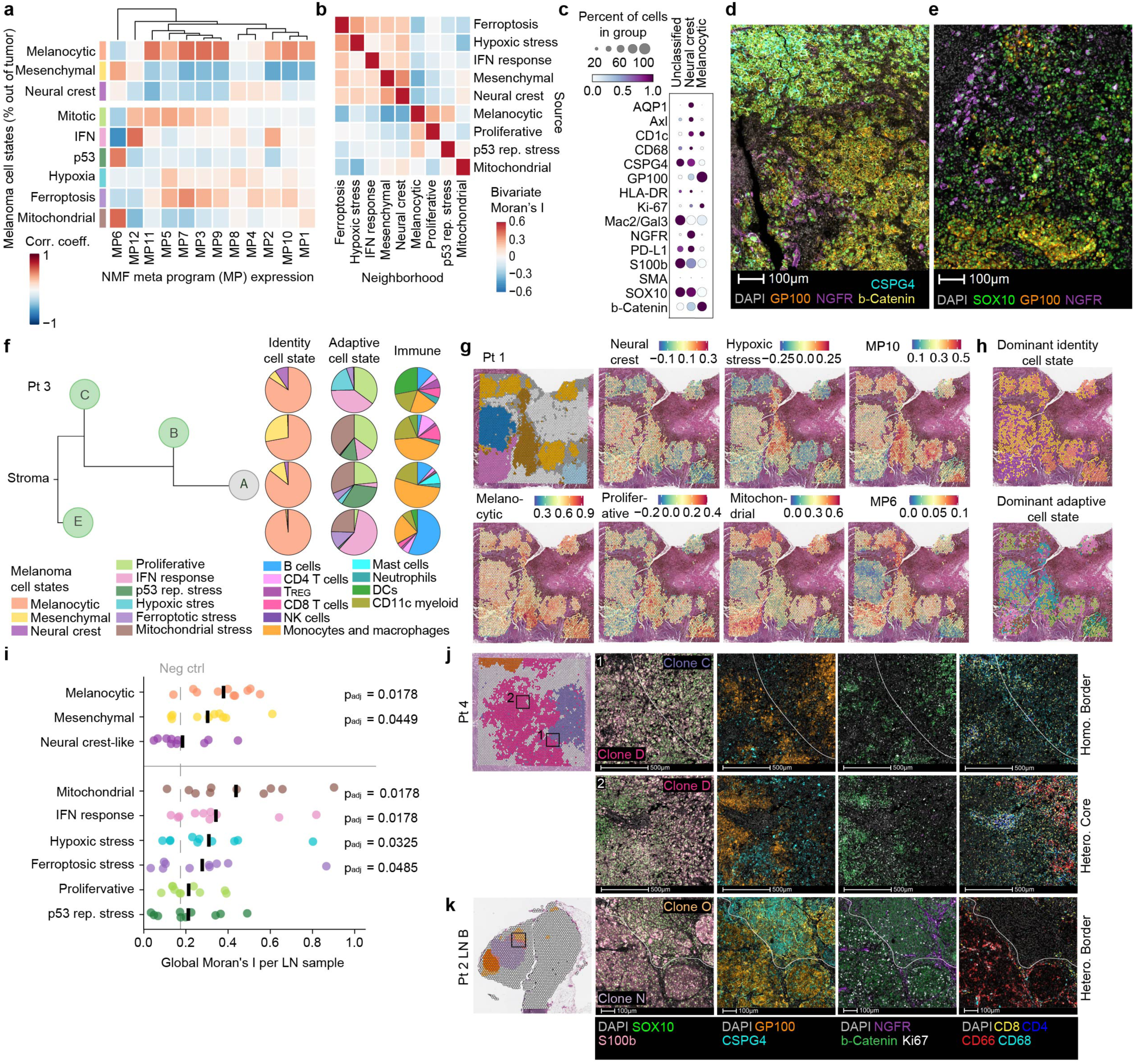
| Genomic variation overlaps with but does not define transcriptional and immunological heterogeneity. **a**, Spearman correlation between non-negative matrix factorization (NMF) metaprograms (MPs) and deconvolution-estimated cell state frequencies (% of tumor) across LN tumor spots (n = 14,158 spots from 10 LNs; 8 patients). Columns hierarchically clustered (Euclidean distance). Color scale reflects correlation coefficient. **b,** Mean global bivariate Moran’s I of identity and adaptive tumor cell states across LN samples (n = 10 LNs from 8 patients). Row axis (Source) indicates focal spot’s cell state expression; column axis (Neighborhood) indicates the mean expression of that cell state in the 6 nearest neighboring spots. Rows and columns are hierarchically clustered (Euclidean distance). **c,** Expression of tumor markers in LN tumors (n = 14 LNs from 12 patients) by imaging, scaled by marker. **d-e,** Representative images of LN tumor marker heterogeneity. Scale bar, 100 µm. **f,** Maximum likelihood tree of representative patient with identity/adaptive cell state proportions by deconvolution and immune cell frequencies by imaging in primary and LN tumor clones. **g,** Representative spatial maps of LN clones (upper left panel) and melanocytic, neural crest, proliferative, hypoxic stress, mitochondrial stress, and metaprogram 10 and 6 signature module scores. **h,** Representative spatial maps of dominant identity and adaptive cell state per spot, assigned as module score with the highest cohort-wide z-score. **i,** Global Moran’s I for clustering of consensus signatures. Negative control: RNA processing. Bar indicates mean across LN samples (n = 10). **j-k,** Spatial map of representative LN samples with clones (left) and images of homogeneous clonal borders and heterogeneous clonal core **(j)** and heterogeneous clonal border **(k).** Scale bar, 500 µm (upper and middle panels) and 100 µm (lower panels). Statistics: paired two-tailed Wilcoxon with BH-FDR correction against negative control; * = padj < 0.05 **(i)**.

Consistent with the concept of phenotypic adaptation, this transcriptional heterogeneity did not strictly depend on the underlying genomic structure. While each clone exhibited a different distribution of cell states, all were competent to acquire multiple identity and adaptive cell states with similar overall heterogeneity in the LN as seen in the primary tumor (**Fig. 2f and Extended Data Fig. 4l**). When comparing the immune microenvironment between sites, primary and LN melanomas had comparable levels of macrophage and CD4 and CD8 T cell infiltration, while LN metastases contained fewer neutrophils and more B cells and regulatory T cells (T_REG_) than primary melanomas **(Extended Data Fig. 4l)**. Within a single LN, analysis of the immune microenvironments of each LN clone highlighted differences across clones, with notable shifts in their association with B cells, dendritic cells, and monocytes/macrophages (**Fig. 2f**). These regional differences in immune composition could indicate altered microenvironments that shape the patterns of intratumoral heterogeneity across clonal boundaries. Indeed, we found several cell state and metaprogram signatures to be visibly clustered, generating distinct regions of biological activity (**Fig. 2g,h**). While melanocytic and mitochondrial states were highly spatially clustered, others, such as the neural crest-like and mitotic states, were dispersed within the tissue **(Fig. 2i)**.

These data indicate that genomic architecture and tumor phenotype are related but not strictly coupled within LN metastases. Transcriptional cell states frequently crossed clonal boundaries, showing that phenotypic heterogeneity could not be explained solely by clonal structure **(Fig. 2j)**. However, we also identified discrete regions in which a clonal interface coincided with an abrupt shift in tumor cell state (**Fig. 2k**). In several cases, the physical convergence of distinct clones was accompanied by a marked change in the local immune microenvironment **(Fig. 2k)**, suggesting that neighboring tumor populations had undergone partially independent programs of adaptation before merging into a single lesion. Thus, rather than representing uniform masses, LN metastases comprise the spatial coalescence of multiple parallel evolutionary trajectories, each shaped by its own combination of genomic diversification, phenotype plasticity, and microenvironmental interactions.

### Melanoma differentiation states occupy unique neighborhoods in the LN

The structured heterogeneity (**Fig. 2**) observed across LNs suggested that melanoma cell states may occupy reproducible spatial niches within the LN. We hypothesized that defining these organizational properties might inform a model of intranodal progression. In both primary tumors and LN metastases, GP100^+^ melanocytic melanoma cells appeared highly clustered in dense lesions, while NGFR^+^ neural crest-like cells were more likely to be seen at the sparse boundaries of lesions, even within the same clone (**Fig. 3a**). By deconvolution, tumor cells in the neural crest-like state were indeed more likely to be in sparse regions of the tumor **(Fig. 3b-d and Extended Data Fig. 5a-c, e-g)** of both the primary tumor and LN metastases, whereas melanocytic cells were enriched in tumor-dense regions (**Fig. 3e-g and Extended Data Fig. 5d, h-j**). To further define the state-specific cellular neighborhoods, we identified the 10 nearest neighbors of each tumor cell and compared them to a simulated control (neighbor enrichment score)^44^. Melanocytic (GP100^+^), neural crest-like (NGFR^+^ or AQP1^+^), and the remaining unclassified tumor cells each occupied distinct immune niches **(Fig. 3h,i and Extended Data Fig. 5k)**. Melanocytic tumor cells were most likely to be next to themselves, with a slight enrichment for MHC Class II (MHCII; HLA-DR)^−^ CD68^+^ macrophages **(Fig. 3h-j)**. Conversely, neural crest-like cells were more likely to be in immune rich neighborhoods enriched for dendritic cells, mast cells, and T cell populations including T_REG_ and effector and exhausted CD8 T cells **(Fig. 3h,i)**. No differences were observed between neural crest-like and unclassified tumor cells (data not shown). Notably, the neural crest-like niche showed clear evidence of local immune activation. Macrophages and monocytes proximal to neural crest-like cells were more likely to be HLA-DR^+^ (**Fig. 3h,j**) and the neural crest-like cells themselves more likely to express HLA-DR (**Fig. 3k-l**). These imaging observations were consistent with the association between the de-differentiated (mesenchymal and neural crest-like) and IFN response program identified in our earlier analyses **(Fig. 2b)**. Spatially, neural crest-like signatures exhibited preferential colocalization with both type I and II IFN and inflammatory TGFβ and TNFα programs relative to melanocytic cells within the same tissue (**Fig. 3m**). Together, these data indicate that established LN lesions acquire a recurrent organizational structure characterized by a dense, poorly inflamed, differentiated core surrounded by a more diffuse, neural crest-enriched peripheral zone of active inflammation.

**Fig. 3.**
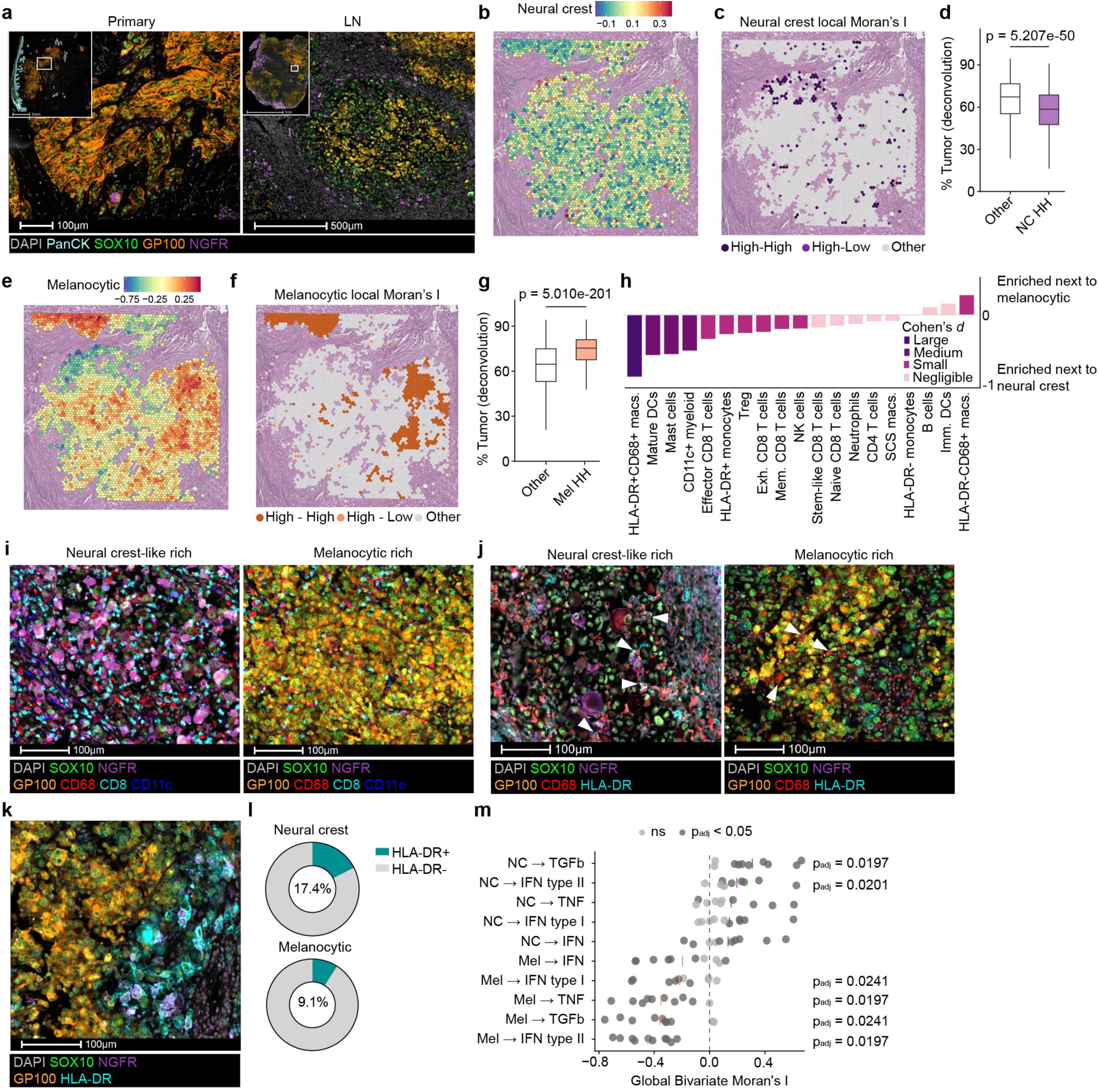
| De-differentiated and differentiated melanoma cells are spatially segregated in the LN. **a**, Representative image of dense melanocytic (GP100^+^) and sparse neural crest-like (NGFR^+^) regions in primary and LN melanoma. Scale bar, 100 µm. **b-c,** Representative spatial projection of neural crest (NC) signature **(b)** and hotspots **(c)**. **d,** Tumor density by deconvolution of NC hotspots (n = 873) versus other (n = 13,285) tumor spots (n = 10 LNs from 8 patients; Cliff’s Delta: – 0.30). **e-f,** Representative spatial projection of melanocytic (Mel) signature **(e)** and hotspots **(f)**. **g,** Tumor density by deconvolution of melanocytic hotspots (n = 2,012) versus other (n = 12,146) tumor spots (n = 10 LN; Cliff’s Delta: 0.42). **h,** Cohen’s *d* effect size of neighbor enrichment score between NC-like and melanocytic tumor cells (n = 14 LNs from 12 patients; large ≥0.8; medium 0.5-0.8; small 0.2-0.5; negligible <0.2). **i-k,** Representative image of NC-like tumor cells with infiltrating CD8^+^ T cells and CD11c^+^ dendritic cells (left) and dense melanocytic region with only CD68^+^ macrophages (right) **(i)**, NC-rich (left) and melanocytic-rich (right) LN tumor areas with adjacent HLA-DR^+^CD68^+^ or HLA-DR^−^CD68^+^ macrophages, respectively **(j)**, and NC-like cells expressing HLA-DR, while adjacent melanocytic cells do not **(k)**. Scale bars, 100 µm. **l,** Proportion of NC-like (upper, n = 391,876) and melanocytic (lower, n = 720,511) cells expressing HLA-DR (*X*^2^ = 1677, p = 2.2e-16, df =1). **m,** Global bivariate Moran’s I (n = 10 LNs) between melanocytic/NC module scores (source) and immune reactivity signatures in 6-spot neighborhood. Statistics: Mann-Whitney U test with Cliff’s Delta (negligible <0.11; small 0.11-0.28; medium 0.28-0.43; large >0.43). * = padj < 0.05 **(d, g)**. Pearson’s chi-squared test on tumor cells by imaging **(l)**. Global Moran’s I (499 permutations, BH-FDR corrected; dot color); one-sample Wilcoxon test vs. zero across LN samples, BH-FDR corrected **(m)**.

### Melanoma cells re-differentiate as they evolve towards dense, immune-excluded macro-metastases

The accumulation of CNV during clonal evolution provides a quantitative framework for placing clones along an inferred evolutionary trajectory **(Extended Fig. 5a)**. We, therefore, used this framework to reconstruct pseudotemporal changes in tumor cell state during LN progression and order the emergence of differentiated and de-differentiated states in their distinct LN niches. We first characterized the cellular composition of LN clones over evolutionary pseudotime. Progression along the inferred trajectory was associated with a marked increase in tumor cell density within LN clones **(Fig. 4a)**, a finding also supported by spot-level deconvolution **(Extended Data Fig. 6b)**. Notably, this relationship was specific to lesion density rather than lesion size, as evolutionary distance did not correlate with the area of the LN clone **(Fig. 4b and Extended Data Fig. 6c)**. Consistently, dense tumor spots had a higher CNV score than sparse spots in the LN **(Fig. 4c)**, placing them later in the evolutionary trajectory. By contrast, the opposite pattern was observed in the primary tumor, where sparse spots had higher CNV scores than dense regions **(Fig. 4c)**, supporting a continuous progression from dense-to-sparse organization in the primary tumor followed by sparse-to-dense organization in the LN.

**Fig. 4.**
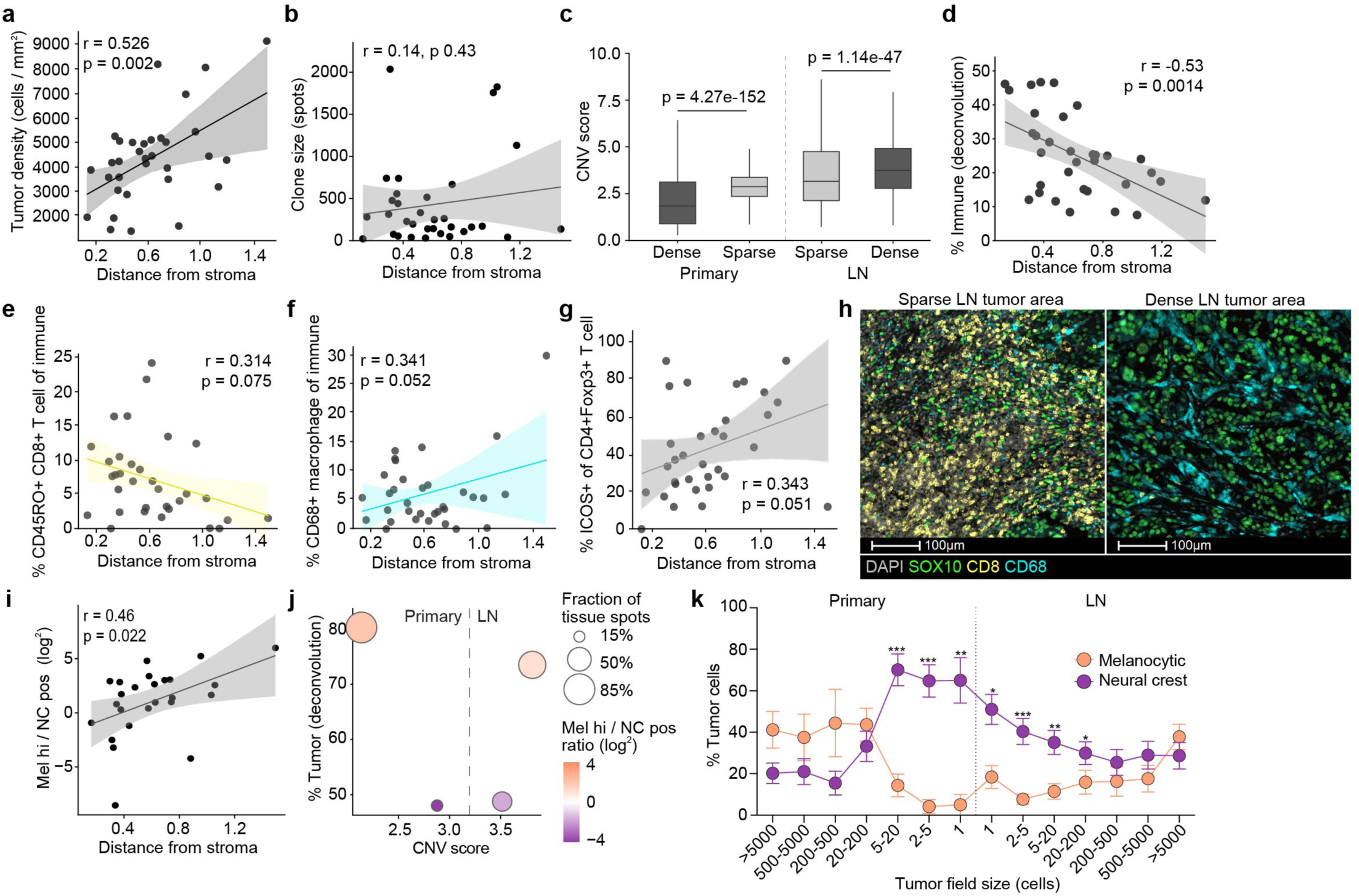
| LN melanomas evolve to be dense, differentiated, and immune excluded. **a**, LN clone (n = 33 from 10 LNs) tumor density (cells / mm^2^) by imaging as a function of evolutionary distance (EVD). **b,** Clone size by number of tumor spots over EVD. **c,** CNV score for dense (> 60% tumor) and sparse (< 60% tumor) spots in primary (dense n = 11,831, sparse n = 2,161; 8 tumors) and LN (dense n = 9,281, sparse n = 4,877; 10 LNs from 8 patients) tumors. Cliff’s Delta: primary –0.36; LNs 0.15. **d,** Mean deconvolved spot immune fraction per LN clone (n = 33 from 10 LNs) as a function of EVD. **e-f,** Percent antigen-experienced CD8 T cells **(e)** and CD68^+^ macrophages **(f)** of immune cells per LN clone over EVD. **g,** Percent regulatory T cells expressing ICOS per LN clone over EVD. **h,** Representative image of sparse SOX10^+^ tumor cells next to CD8^+^ T cells (left) and dense tumor region infiltrated by CD68^+^ macrophages (right). Scale bar, 100 µm. **i,** Log2 ratio of melanocytic-high to neural crest-positive proportion by deconvolution over EVD (n = 33). **j,** Tumor density by deconvolution and CNV score for dense and sparse spots in primary and LN melanomas, colored by log2 ratio of melanocytic-high to neural crest-positive proportion. Size scales for fraction of spots. **k,** Percent of melanocytic (GP100^+^ tumor) and neural crest-like (NGFR^+^ or AQP1^+^ tumor) cells in tumor fields in primary (n = 127 fields from 8 tumors) and LN (n = 231 fields from 14 LNs) melanoma. Statistics: Pearson correlation **(a, b, d-g, i)**, Mann-Whitney U test with Cliff’s Delta (negligible <0.11; small 0.11-0.28; medium 0.28-0.43; large >0.43) **(c)**, and paired two-tailed Wilcoxon with BH-FDR correction. *: padj < 0.05; **: padj < 0.01, ***: padj < 0.001 **(k)**.

We then examined how this shift in spatial organization related to the local microenvironment. Whereas the deconvolved fraction of fibroblasts **(Extended Data Fig. 6d)** and endothelial cells **(Extended Data Fig. 6e)** remained relatively stable across evolutionary pseudotime, the overall immune cell fraction declined significantly **(Fig. 4d and Extended Data Fig. 6f)**. Higher resolution analysis using imaging validated this result and demonstrated that B cells **(Extended Data Fig. 6g)** and antigen experienced CD45RO^+^ CD8 **(Fig. 4e)** and CD4 **(Extended Data Fig. 6h)** T cells specifically decreased in the tumor with evolutionary distance. In contrast, CD68^+^ **(Fig. 4f)** and Mac2^+^ monocytes **(Extended Data Fig. 6i)** increased, as did the proportion of activated, ICOS^+^, T_REG_ **(Fig. 4g and Extended Data Fig. 6j)**. Together, these data identify an immune phenotype tightly associated with LN evolution, in which sparse, immune-rich lesions progress toward dense, size-independent, myeloid-dominant, and immunosuppressed metastatic structures **(Fig. 4h).**

Consistent with our cell state analysis, which correlated differentiation with the formation of dense, immune-excluded lesions, the ratio of differentiated to de-differentiated tumor cells in a LN clone increased with evolution **(Fig. 4i)**. Using the spot-level metastatic trajectory of density and CNV established in **Fig. 4c**, we found that dense, low CNV spots in the primary were more melanocytic, then transitioned to sparse, neural crest-like spots, consistent with the hypothesis that invasive de-differentiated cells drive more aggressive tumor behavior^36,45^ **(Fig. 4j)**. Interestingly, the continued accumulation of CNVs associated with a reversal of de-differentiation in the LN, where sparse, neural crest-like lesions with lower CNV scores gave way to denser, differentiated lesions with higher CNV scores **(Fig. 4j)**, indicating that as tumors in the LN continue to evolve, they become denser and more differentiated.

While spatial transcriptomics provides both inferred genomic and transcriptional depth to our analyses, it is limited in the detection of the smallest – and possibly earliest – lesions within the LN. To increase our resolution of early metastatic lesions and the differentiation switches that occur within the LN, we defined distinct tumor fields in each specimen independent of its clonal architecture **(Extended Data Fig. 6k)**. Differentiation states plotted as a function of field size across anatomical compartments confirmed the predicted melanoma de-differentiation and re-differentiation axis during LN metastasis **(Fig. 4k)**. Taken together with the transcriptional analyses above, these data support a model in which phenotypic adaptation and immune exclusion occur in parallel to ongoing CNV accumulation and clonal diversification to establish metastatic lesions in the LN.

### Melanoma evolution in the LN is associated with a phenotype switch that correlates with poor outcomes

To define transcriptional programs associated with LN evolution, we compared tumor spots from the clones at the shortest (early, n = 7) and longest (late, n = 9) evolutionary distances from stroma **(Fig. 5a,b)**. Early clones were enriched for immune activation and de-differentiation programs, while highly evolved clones were enriched for signatures of RNA processing, mitochondrial metabolism, and cytoskeletal rearrangement **(Fig. 5b and Supplementary Table 6)**. Consistent with this shift, comparison of tumor-derived NMF programs showed depletion in late clones of factor 9, a signature of immune infiltration **(Extended Data Fig. 7a)**, and factor 11, a neural crest-like and extracellular matrix rich factor **(Fig. 5c)**. Conversely, factor 22, a signature of proliferation and mitochondrial activation, was enriched in late clones **(Fig. 5d)**. In parallel, the immune composition shifted toward myeloid and granulocytic dominance in late clones relative to early LN clones **(Fig. 5e)**, in agreement with the progressive microenvironmental remodeling observed across pseudotime **(Fig. 4)**.

**Fig. 5.**
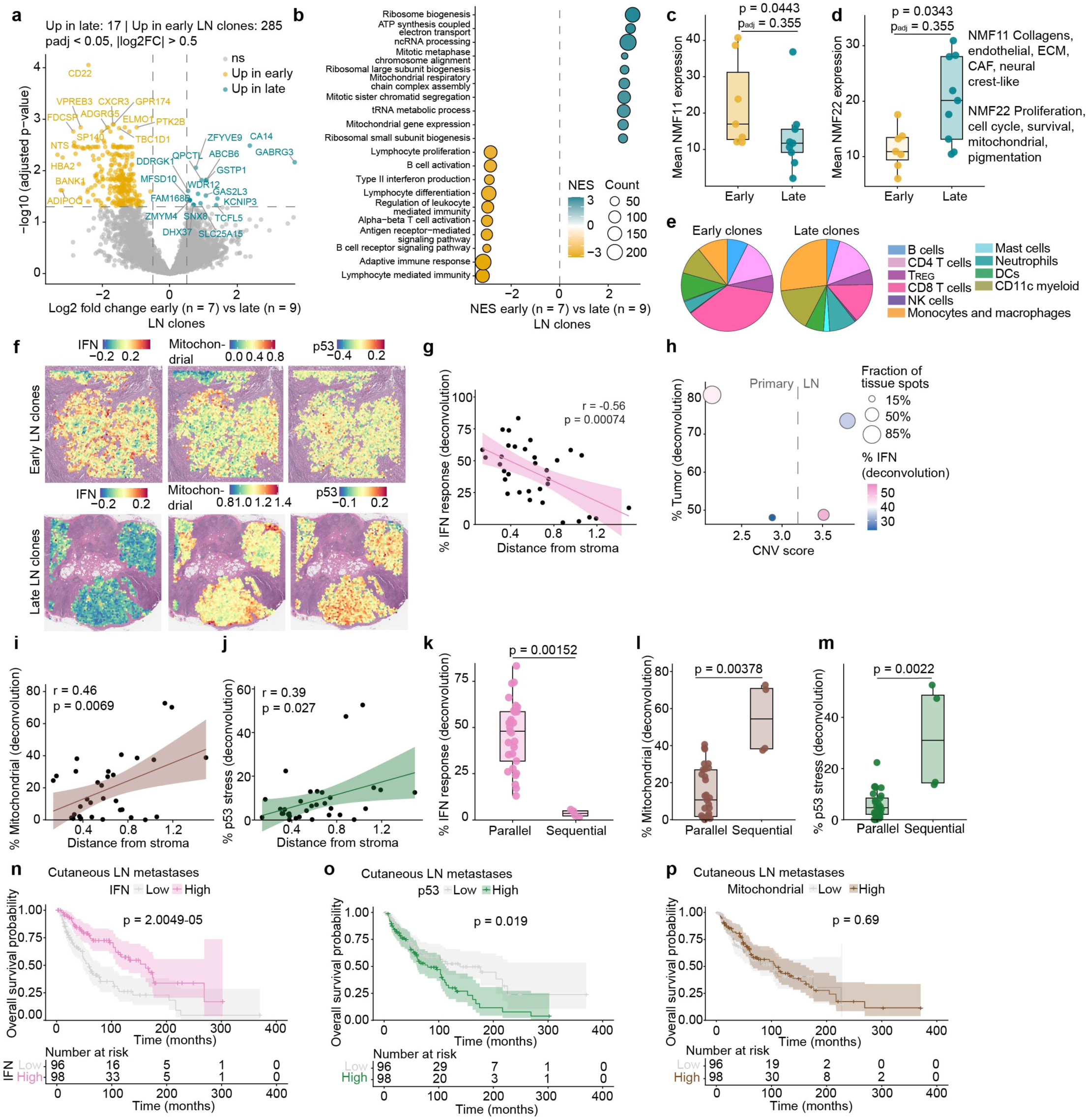
| A phenotype switch in the LN associates with disparate clinical outcomes. **a**, Differentially expressed genes between early (n = 7) and late (n = 9) LN clones, stratified by evolutionary distance. Pseudobulk DESeq2 (unpaired). Dashed lines indicate |log2FC|>0.5 and p_adj_ < 0.05. Teal = upregulated in late clones (n = 17); yellow = upregulated in early clones (n = 285). **b,** GSEA of GO Biological Process terms between early and late LN clones, ranked by log2fold change. Top 10 terms per direction shown after redundancy reduction. **c-d,** Mean expression of NMF program 11 **(c)** and 22 **(d)** in early and late LN clones. **e,** Immune microenvironment of early and late LN clones (*X*^2^ = 1171.1, p = 2.2e-16, df = 9). **f,** Representative spatial projection of IFN response, mitochondrial, and p53 stress signatures in tumor of early and late LN clones. **g,** Fraction of IFN response by deconvolution over evolutionary distance to stroma (n = 33 clones from 10 LNs). **h,** Tumor density by deconvolution and CNV score of primary and LN tumor spots (as in Fig. 4c). **i-j,** Fraction of mitochondrial **(i)** and p53 stress **(j)** by deconvolution over evolutionary distance. **k-m,** Mean deconvolution-estimated fraction of IFN-responsive **(k)**, mitochondrial **(l)**, and p53 stress **(m)** tumor cells in parallel (n = 29) and linear (n = 4) LN clones. **n-p,** Overall survival stratified by IFN responsiveness **(n)**, p53 **(o)**, and mitochondrial stress **(p)** in LN metastases (n = 194) from patients with cutaneous melanoma. Statistics: DESeq2 Wald test with BH-FDR correction **(a)**, Mann-Whitney U test with BH-FDR correction **(c-d)**, Pearson chi-squared test on clone-averaged immune cells by imaging **(e)**, Pearson correlation **(g, i-j)**, Mann-Whitney U test **(k-m)**, and Kaplan-Meier survival curve with log-rank Mantel-Cox test **(n-p)**.

Because neural crest-like cells preferentially localized to immune-reactive niches **(Fig. 3)** and were depleted as lesions progressed along the LN metastatic trajectory **(Fig. 4)**, we next asked whether inflammatory responsiveness also changed with LN clonal evolution. Module scores and deconvolution-based estimates both showed progressive reduction in TNFα, TGFβ, and IFN response programs with evolutionary distance to stroma **(Fig. 5f,g and Extended Data Fig. 7b-d)**. Among these, the decline of type II IFN response was more pronounced than type I IFN response **(Extended Data Fig. 7e,f)**, whereas cGAS-STING signaling increased with evolutionary pseudotime **(Extended Data Fig. 7g)**. Projecting these inflammatory programs onto the joint CNV-density metastatic trajectory further showed that TNFα, TGFβ, and IFN responses were enriched in tumor-sparse, low CNV-score regions of LN metastases **(Fig. 5h and Extended Data Fig. 7h, i)**. Notably, although TNFα and TGFβ responses were also enriched in sparse spots in the primary, the IFN response was not, suggesting that IFN may represent an environmental exposure encountered upon LN entry. In parallel with declining inflammatory reactivity, more evolved clones showed increased mitochondrial stress **(Fig. 5f,i)** and p53 replicative stress **(Fig. 5f,j)** programs, defining a phenotypic transition from an immune-reactive to stress-adapted metastatic state.

We next asked whether this shift was linked specifically to the site and timing of tumor evolution. Our data had identified two major evolutionary patterns: a predominant pattern of early dissemination and parallel evolution, where metastasis occurred early and genomic diversification took place in the LN itself, and a pattern of late dissemination and linear evolution, where genomic diversification was already advanced in the primary tumor before nodal spread. If attenuation of IFN responsiveness accompanies or enables metastatic outgrowth, then clones arriving in the LN after more extensive prior evolution should be less IFN responsive than those diversifying locally within the node. Consistent with this prediction, parallel LN clones were significantly more IFN responsive than linear LN clones **(Fig. 5k),** whereas TNFα and TGFβ did not differ detectably (data not shown). Linear clones also exhibited higher mitochondrial stress and p53 replicative stress programs **(Fig. 5l,m)**, further linking advanced genomic evolution to a stress-adapted, less inflammatory phenotype. Together these findings indicate there are tumor phenotypes more competent for LN outgrowth, and that in most cases these states are acquired locally, where environmental cues (e.g. IFN signaling) instruct adaptive reprogramming of genomically and phenotypically plastic melanoma cells.

Finally, we asked whether this phenotype switch had prognostic relevance for melanoma patients. We found that high IFN responsiveness in LN metastases was associated with improved overall survival in the TCGA SKCM cohort **(Fig. 5n).** Both type I and type II IFN responses stratified outcomes **(Extended Data Fig. 7j,k)**, although the association was stronger for type II IFN. By contrast, neither TNFα nor TGFβ response in the LN was prognostic **(Extended Data Fig. 7l,m)**. Conversely, the high p53 replicative stress score that was enriched in evolved clones in LN metastases was associated with worse overall survival **(Fig. 5o)**. Mitochondrial stress response did not stratify patient survival **(Fig. 5p)**. The evolutionary trajectory revealed through this work **(Extended Data Fig. 7n)**, therefore, highlights previously underappreciated biological heterogeneity within LN metastasis, driven by the extent of local evolution, that differentially stratifies patient survival outcomes. We therefore propose that the genomic and adaptive history of a LN lesion mechanistically informs patient prognosis beyond conventional staging, which may motivate future biomarker validation and novel targeting strategies.

## DISCUSSION

Despite increasing recognition that LN metastasis influences systemic cancer progression^14,15^ and therapeutic response^46,47^, the mechanisms that enable tumor outgrowth in the LN remain poorly defined. Using multimodal spatial analysis of paired primary melanoma tumors and metastatic sentinel LNs from patients, we delineate a trajectory of genomic, phenotypic, and immunologic adaptation during nodal metastasis. Our data support early dissemination from the primary tumor but indicate that successful LN colonization depends on specific tumor phenotypes that most often arise only after substantial local evolution and immune evasion in the LN. The resulting metastatic lesions present as composite ecosystems assembled through the spatial coalescence of multiple evolving populations rather than uniform tumor masses. Notably, the extent of this regional evolution correlates with patient survival, suggesting that LN metastases contain biological information that may refine prognostic stratification beyond conventional staging. Our work constructs an integrated framework for understanding the dynamic nature and clinical significance of LN metastasis which can be further mechanistically investigated.

Preclinical studies have suggested that LN metastasis is an active process shaped by local immunologic^14,20,21^ and metabolic^4,18,19^ constraints. In contrast, clonal heterogeneity in human metastatic LNs has been interpreted to support a stochastic model of continuous, passive LN seeding^16^. Our data offer an alternative explanation. The early phylogenetic divergence of primary and LN clones, together with the continued evolution of distinct CNV profiles, indicates that nodal heterogeneity may arise through diversification within the LN itself rather than from repeated seeding by multiple primary clones. This is consistent with the observation that early disseminated melanoma cells acquire driver mutations in the LN itself^48^. Beyond specific mutations, however, we found that general aneuploidy in the primary tumor was associated with increased risk of LN metastasis in cutaneous melanoma, an observation also made in head and neck cancer^49^. The increase in clonal diversity we observed after LN seeding suggests that selective pressures in the nodal microenvironment may favor more genomically plastic and adaptable cells. Genomic instability and aneuploidy are mechanistically linked to immune evasion^50–52^ and metastatic fitness^53,54^, consistent with the emerging model here that genomic instability facilitates the necessary adaptations for LN metastasis. Although canonical CD8^+^ T cells may not be the dominant constraint on LN metastases^17^, innate immune populations^19^, metabolic competition^4,18^, or structural features of the LN likely impose additional barriers to regional tumor fitness. Together, these findings support an active model of LN metastasis in which the nodal microenvironment sets up barriers to colonization that select for adaptive tumor states.

The progressive accumulation of CNVs provided a pseudotime framework for mapping non-genomic adaptations across anatomic sites and within the LN. As reported in pancreas^55,56^ and prostate cancer^57^, melanoma cell state transitions were only partly aligned with clonal architecture, highlighting substantial plasticity during metastasis. We infer that disseminating melanoma cells arise from de-differentiated states in the primary tumor^58^, including neural crest-like programs, whereas LN lesions acquire a more differentiated melanocytic identity at their core over time. This apparent re-differentiation resembles the mesenchymal-to-epithelial transitions described in squamous cell carcinoma distant metastases^59^ and parallels observations in breast cancer LN metastases^20,60^, where differentiated epithelial states occupy lesion cores and more mesenchymal states localize to the periphery^20^. Within this framework, single disseminated cells were less likely to exhibit a differentiated melanocytic program, consistent with recent reports that early metastatic seeds in melanoma adopt de-differentiated states^43^. Our data thus indicate that successful LN colonization may require sequential and dynamic use of distinct tumor programs, rather than fixation in a single stable phenotype.

Immune reactivity appears to be a component of this spatial organization. Across tumor types, inflammatory cues, including IFNs, have been linked to the induction and maintenance of de-differentiated states^61–63^, and in our data, de-differentiated melanoma cells colocalized with inflammatory infiltrates and IFN response signatures in the LN. Whether IFN signaling is required to sustain a de-differentiated state in vivo remains unclear, but these programs may nonetheless promote immune evasion at the lesion periphery while protecting a more immune-excluded melanocytic core. This interpretation is consistent with prior work showing that IFN-induced MHCII and PD-L1 expression can facilitate immune escape in LN metastasis^14,20^, including through engagement of T_REG_ populations^17,20^, and with broader evidence that chronic IFN signaling contributes to immune dysfunction^64^ and resistance to immunotherapy^65^. Interestingly, however, progression to more evolved LN clones is associated with attenuation of the IFN response program. One possibility is that inflammatory signaling is advantageous early during adaptation to the LN but becomes dispensable, or even deleterious over time. In renal cell carcinoma, loss of IFN sensing confers increased tolerance to CIN^50^ and is necessary for outgrowth. Our data indicate that melanoma may be analogous, with disseminated de-differentiated melanoma cells establishing an inflammatory periphery niche that supports the emergence and persistence of an immune-evasive differentiated core. Although this model requires functional testing, it provides a potential explanation for how local inflammatory signaling can be implicated in both early adaptive responses and later immune escape.

This new model implies that biologic heterogeneity within LN metastases captures features of tumor evolution that may provide a plausible basis for how nodal metastasis could contribute to systemic tolerance. In particular, the phenotype switch identified here tracked along the evolutionary trajectory in the LN and stratified prognosis. Melanoma cells may therefore reach the LN early, but clinically consequential nodal metastasis reflects substantial local evolution within the LN. The p53 and mitochondrial stress programs enriched in more evolved lesions, and the dynamic processes required to establish them, may therefore represent actionable vulnerabilities of established LN lesions, complementing previously described metabolic dependencies^19^. More broadly, because draining LNs are central to systemic immune surveillance, interventions that preserve LN structure and function by limiting metastatic outgrowth may have benefits that extend beyond local disease control.

Our cohort also suggests that LN evolution may proceed along more than one temporal route. Although early dissemination with parallel evolution predominated, two patients with evidence of linear metastasis showed relatively clonal but more evolved LN lesions enriched for the same biological programs associated with poor outcome. This observation raises the possibility that acquisition of these states is rate-limiting for nodal outgrowth and genomic instability facilitates their emergence. These findings will require validation in larger cohorts, and both our patient set and TCGA-based analyses are underpowered to establish the reported signatures as predictive biomarkers. Nonetheless, this study establishes LN metastasis as a dynamic process and provides a unifying framework for understanding how tumor evolution within the LN shapes systemic disease outcomes. This work also suggests that LN metastases are an underused clinical resource for evaluating tumor progression, prognosis, and therapeutic vulnerability.

## MATERIALS AND METHODS

### Melanoma tissue procurement, selection, and processing

Formalin-fixed paraffin-embedded (FFPE) cutaneous and acral melanoma specimens were obtained from the New York University Langone Medical Center under Institutional Review Board approval (IRB H10362). Written informed consent was obtained from all patients prior to tissue collection and analysis. FFPE blocks were maintained at 4°C until sectioning.

Blocks were assessed histologically (hematoxylin & eosin; 5 µm sections), digitally scanned, and evaluated using internally developed AI-based image analysis tools to confirm melanoma presence^25,24^. Only specimens with greater than 5 mm² of tumor area were advanced for downstream preparation and quality assessment. RNA integrity was assessed on a 10 µm scroll, and samples with a DV200 value exceeding 30% were deemed suitable for spatial transcriptomic (ST) profiling. Sequential 5 µm sections were collected for paired imaging and ST.

### Akoya Fusion Imaging

5 µm FFPE slides selected for imaging were baked vertically overnight at 65°C and stained according to the commercial protocol^23,66^ using a panel of 29 antibodies purchased directly from Akoya and 15 custom conjugated antibodies **(Supplementary Table 2)**. Slides were deparaffinized and rehydrated and antigen retrieval was performed at pH 9 in a pressure cooker for 20 minutes on high pressure. All primary antibodies were stained together with blocking oligonucleotides for 3 hours at room temperature. Tissues were fixed by sequential 1.6% PFA and 4°C methanol incubation before flow cell application and storage.

Antibodies were conjugated according to the Akoya commercial protocol. Briefly, 50 µg carrier-free antibody was filtered through a blocked 50kDa MWCO filter, its disulfide bonds were partially reduced, and the exposed sulfides tagged with maleimide-tagged oligonucleotides.

Imaging was performed on the Akoya Fusion PhenoCycler.

### Image analysis

#### Image post-processing and marker thresholding

Image post-processing was performed using HALO (Indica Labs). Nuclear segmentation was performed using the HALO AI Nuclei Seg V2 network (minimum nuclear area 8 µm^2^, maximum nuclear area 800 µm^2^) and the HALO AI DenseNet V2 network to distinguish tumor from non-tumor compartments. Briefly, representative tumor and non-tumor regions were manually annotated to train the tumor classifier, which integrates pixel-level features (including tissue architecture and cell morphology) together with the intensities of selected markers (DAPI, SOX10, S100B, GP100, and CSPG4). The minimum object size set at 50 mm^2^, and the analysis resolution was 0.65 mm per pixel. The trained models were then applied to segment individual cells and to classify tumor versus non-tumor regions across all tissues. Clones were manually annotated as detected on SpatialDimPlot by SpatialInferCNV; whole-tissue annotations encompassed the entire primary tumor section or the whole metastatic LN section. Tumor fields were defined as tumor classifier regions separated by > 100 mm. Staining artifacts and folded tissue were excluded.

Intensity-based thresholding was performed using HighPlex FL v4.3.2. For each marker, a positivity threshold was set on one representative section and all sections were normalized using min-max scaling. All thresholds were manually inspected and adjusted as needed. Finally, per-cell matrices containing cell IDs, coordinates, tumor classification, average marker intensities, and applied thresholds were exported from HALO.

#### Normalization and pre-processing

Image analysis was performed in Python (v3.9.21) and adapted from the SPACEc image analysis pipeline^67^. Individual ROI cellular matrices from HALO were as AnnData objects; cell intensity values were maintained as the data frame and all additional data was moved to metadata. Cell centroids were calculated by finding the mean of X and Y minimum and maximum values per segmented nuclei. The lowest 1% of cells per ROI by whole cell area (µm^2^) and DAPI Cell Intensity were removed. The cell intensity matrix of each ROI was Z scored and used for all downstream processing and analysis. The highest 1% of cells by either number of markers expressed or total intensity were removed. Individual ROI AnnData objects were merged into a single AnnData object for downstream analysis.

#### Cellular annotation

Initial cellular annotation was performed by Leiden clustering on cell-identifying markers (AQP1, b-catenin, CD11c, CD14, CD169, CD20, CD21, CD31, CD3e, CD4, CD40, CD57, CD66, CD68, CD8, Collagen IV, Foxp3, GP100, HLA-DR, mast cell tryptase, pan-cytokeratin, PNAd, podoplanin, S100b, SMA, and SOX10) with 20 nearest neighbors and a resolution of 0.1. Resulting clusters were annotated as tumor, immune, or stromal. Cells in the tumor cluster not annotated by the HALO classifier as tumor were moved to the stromal cluster for downstream annotation. Initial clusters were reclustered at a resolution of 0.1 and clearly identifiable populations were labelled. Remaining clusters were serially reclustered until defined populations could be clearly annotated. Ultimately, clusters strongly expressing no markers, all markers, or a biologically impossible combination of markers were removed from downstream analysis (∼10% cells per ROI).

HALO thresholds were used for manual annotation of subpopulations: mature DCs (HLA-DR^+^ CD40^+^ cDC1, cDC2, or CD11c^+^ myeloid); neural crest-like tumor cells (AQP1^+^ CD31^−^ or NGFR^+^ CD20^−^ CD21^−^ tumor cells); melanocytic tumor cells (GP100^+^ tumor cells); antigen experienced (CD45RO^+^) and naïve (CD45RO^−^) CD8 and CD4 T cells; effector CD8 T cells (Granzyme B^+^ or IFN*γ*^+^ CD45RO^+^ T cells); stem-like T cells (TCF1^+^ TOX^+^ CD45RO^+^ CD8 T cells); central memory CD8 T cells (TCF1^+^ TOX^−^ CD45RO^+^ CD8 T cells); exhausted CD8 T cells (TCF^−^ PD-1^+^ CD45RO^+^ CD8 T cells); resident memory CD8 T cells (CD103^+^ CD45RO^+^ CD8 T cells); effector memory CD8 T cells (CD45RO^+^ TOX^−^ TCF1^−^ PD-1^−^ Granzyme B^−^ IFN*γ*^−^ CD103^−^ CD8 T cells).

#### Neighborhood analysis and simulation

Neighborhood analysis was performed using a modified version of the SPACEc neighborhood function^67^ that exports the per cell neighbor matrix before k means clustering. Cellular neighborhoods were defined using k means clustering of 10 nearest neighbor distributions; 10 nearest neighbors approximate a radius of direct contact. A simulation function was generated that reads in the cell-level neighbor matrix from the neighborhood function and creates a simulated neighbor matrix weighted to the cellular composition of each ROI. This simulation was run at 1, 10, 50, 100, 300, 500, 800, 1000, and 1500 iterations; the point of convergence was identified as the number of iterations past which each cell’s distribution changed less than 1%. For each cell, the neighborhood enrichment score was defined:

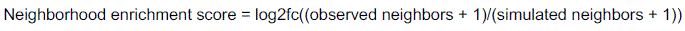

The neighborhood enrichment score was used to calculate the Cohen’s *d* effect size. This analysis was adapted from ESImap as described in Solis et al^44^.

### Spatial transcriptomics library construction and sequencing

Eligible specimens proceeded to spatial transcriptomic library preparation using the 10x Genomics Visium/CytAssist workflow optimized for FFPE tissue (Visium Human Transcriptome Probe Set v2.0). Tissue sections were methanol-fixed, stained with H&E, and digitally imaged. Either a 6.5 × 6.5 or 11 x 11 mm region of interest (ROI) **(Supplementary Table 3)** was delineated within OMERO Plus (OMERO.web v5.25.0). A board-certified dermatopathologist verified that melanoma was fully contained within the annotated ROI.

Library preparation was performed using the Visium Spatial Gene Expression Reagent Kit (10x Genomics), including tissue permeabilization, reverse transcription, second-strand synthesis, and amplification of cDNA. Final pooled libraries were sequenced on an Illumina NovaSeq X+ platform using a 10B 100-cycle flow cell.

### Spatial transcriptomics data processing and analysis

Sequencing data were aligned to the human reference transcriptome (GRCh38-2020-A, hg38) using Space Ranger (v2.1.1; 10x Genomics). The resulting filtered_feature_bc_matrix.h5 files were imported into Seurat (v5.3.0) within R (v4.4.1). Spots with fewer than 200 unique molecular identifiers (nCount_Spatial) were excluded to remove low-quality capture locations. Spots located outside tissue boundaries were manually removed using Loupe Browser (v7.0.1), and those barcodes were loaded using a csv file for removal from the object.

Data were normalized using the NormalizeData function, which performs global-scaling normalization. For each spatial spot, total transcript counts were scaled with a scale factor of 10,000 and log-transformed to correct for differences in sequencing depth and capture efficiency across spots.

Following preprocessing of individual samples, datasets were merged. The top 3,000 most variable features were identified using FindVariableFeatures with the variance-stabilizing transformation (vst) method, the data were scaled using ScaleData, and principal component analysis (PCA) was performed using RunPCA. Batch effects by sequencing batch were mitigated using Harmony (v1.2.4). Uniform Manifold Approximation and Projection (UMAP) embeddings were generated using RunUMAP, and a nearest neighbor graph was constructed using FindNeighbors based on the first 30 principal components, followed by unsupervised clustering across multiple resolutions using FindClusters. When subsetting samples of interest, or when restricting the analysis to tumor or stromal spots, FindVariableFeatures, ScaleData, RunPCA, RunHarmony, RunUMAP, FindNeighbors, and FindClusters were rerun on the subsetted data.

### Clone identification and analysis

#### Clone identification

ST clones were identified using sequential SpatialInferCNV^26,27^ (v.0.1.0). Briefly, SpatialInferCNV was run reference-free to define tumor and stromal regions. Tumor spots were verified against H&E. SpatialInferCNV was run on those tumor spots a second time, with the defined stroma serving as a sample-specific reference. Spot CNV predictions were hierarchically clustered and clone branching points were manually identified. SpatialInferCNV was then run again at the newly-identified clone level to predict clone-level CNV probabilities using a Hidden Markov Model (HMM) and Bayesian model. Clone CNVs that were determined to be more than 50% likely to be real by Bayesian posterior probability were kept as clone-defining CNVs and mapped to chromosome arms. Clones where fewer than 50% of spots contained clone-defining CNVs and which clustered spatially and on transcriptional UMAP between tumor and stromal regions were annotated as mixed tumor-stromal border and eliminated from downstream analysis.

#### Phylogenetic tree construction and analysis

A phylogenetic tree was constructed for each patient using a maximum likelihood model, manually rooted on a diploid stroma. Edge length distance is annotated as evolutionary distance to stroma. Mean pairwise distance (MPD) was calculated as the shortest distance between pairs of clones within the LN or within the primary tumor.

#### CNV score

The CNV score was calculated as the sum of absolute fold change expression deviations across all chromosomal arms per spot, normalized by the deconvolution-estimated fraction of tumor per spot, serving as a tumor content-corrected spot-level proxy for CNV.

#### CIN signature

The CIN high signature was derived in house from an isogenic mouse model of chromosomal instability in which KP lung adenocarcinoma cells were engineered with the murine adaptation of KaryoCreate, a doxycycline inducible AuroraB dCas9 fusion targeted to pericentromeric minor satellite repeats by sgMinSat5, with a non-targeting sgRNA (sgRosa26) as the isogenic near diploid control (adapted from Bosco et al^68^ and Cheng et al^69^). Bulk RNA sequencing of high aneuploidy (sgMinSat5) versus low aneuploidy (sgRosa26) cells was analyzed with DESeq2^70^, and genes ranked by the DESeq2 test statistic were used for GSEA against the Reactome collection of MSigDB. Three gene sets were enriched in high aneuploidy cells: extracellular matrix organization and collagen formation (ES = 0.82, q = 0.10), immunoregulatory interactions between a lymphoid and a non-lymphoid cell (ES = 0.71, q = 0.15), and initial triggering of complement (ES = 0.76, q = 0.10). From these programs we retained the most strongly regulated genes encoding secreted or cell surface proteins, that is, those able to act on the tumor microenvironment, yielding an 11 gene CIN high signature: *CCL2*, *C3*, *CXCL1*, *CXCL5*, *IL1RN*, *TFF1*, *TFF2*, *TFF3*, *CCL17*, *CD36*, and *CDCP1* (human orthologs used for scoring human data).

#### Spatial restriction of CNV clones

To map spatial restriction of CNV clones, the global Moran’s I statistic was calculated for each clone. Briefly, CNV clone status was treated as a binary spatial feature, with 1 = clone spot and 0 = other spot. CNV clones with fewer than 30 spots (1 LN clone) or samples with only 1 clone (4 primary tumors) were excluded from the analysis. The remaining clones were analyzed using the FindSpatiallyVariableFeatures() function in Seurat v5.3.0 with selection.method = moransi. Results were corrected across CNV clones on a per-sample basis for multiple comparison using BH-FDR (padj = 0.05).

### Consensus melanoma cell states

To generate consensus signatures from the literature for known melanoma cell identity states, overlapping genes were identified between signatures from the literature for melanocytic^29,31–38^, mesenchymal^29,32,33,35,36^, and neural crest^29,31–34,37,38^ cells. Any gene that appeared two or more times was added to the consensus signature. As adaptive states are less established, we used signatures directly from Pozniak et al^32^ for mitosis, p53 stress, and mitochondrial stress, a merged signature from Pozniak et al^32^ and Karras et al^33^ for hypoxia, and an IFN signature of combined type I IFN signaling (Pozniak et al^32^) and type II IFN signaling (Ayers et al^42^). For ferroptosis, we made a consensus signature of genes present in at least two of the following: Guo et al^41^; Liu et al^41^, and Pin et al^39^.

Module scores were calculated using AddModuleScore() in Seurat v5.3.0. Dominant identity and adaptive cell states were assigned per spot by z-scoring all module scores and selecting the cell state with the highest score within each category.

### Deconvolution of spatial transcriptomics

Visium 10X spatial transcriptomic data was deconvolved using Robust Cell Type Decomposition (RCTD) as included in spacexr v2.2.1. First, spots were deconvolved using Tirosh et al^34^ as a single cell RNA sequencing (scRNAseq) reference. scRNAseq data was downloaded from https://www.weizmann.ac.il/sites/3CA/skin and pre-processed in Seurat v5.3.0 according to the following steps and parameters: NormalizeData() with LogNormalize and scale.factor = 10,000; FindVariableFeatures() method vst, 2,000 variable genes; ScaleData(); RunPCA(); FindNeighbors() on pcs = 1:30; FindClusters(). Batch effect was corrected for using RunHarmony() by sample. The following cell types were pre-annotated and checked using FindAllMarkers(): T cells, B cells, NK cells, myeloid cells, fibroblasts, endothelial cells, and malignant cells. To annotate subclusters of malignant cells, module scores of consensus signatures were added, and dominant identity and adaptive cell states were identified as described above.

RCTD was performed three times: once on the whole tissue using the cell type references only (T cells, B cells, NK cells, myeloid cells, fibroblasts, endothelial cells, and malignant cells) to capture the broad cellular distribution of the data, once on tumor spots using the malignant cell reference annotated by identity state to identify the distribution of identity states per spot, and once on tumor spots using the malignant cell reference annotated by adaptive state to identify the distribution of adaptive states per spot. Deconvolution was run per sample using create.RCTD() and run.RCTD() with doublet_mode = full to estimate cell type fractions for all cell types simultaneously per spot.

#### Defining density by deconvolution

Tumor cells were defined as sparse if the RCTD-estimated malignant cell fraction per spot was in the bottom quartile of all tumor spots across all samples (n = 8 primaries and n = 10 LN), or < 59.51% tumor by RCTD. The remaining tumor spots were annotated as dense.

### Unbiased definition of transcriptional programs using non-negative matrix factorization (NMF)

To define transcriptional programs independent of reported cell states, we performed non-negative matrix factorization (NMF) on the paired primary (n = 8) and LN (n = 10) tumors. Samples were downsampled by random selection of 50% of tumor spots per sample. NMF v0.28 was used to run NMF with 25 ranks (defined by elbowplot) on the top 2,000 variable genes of the downsampled cohort. NMF() was run with seed = ica, method = non-smooth NMF, nrun = 5 iterations; 150 ranked genes define each NMF factor. Factors were annotated by gene ontology (GO) analysis (fgsea v 1.30.0, msigdbdf v24.1.0). Resulting pathways were filtered using Normalization Enrichment Score > 1.5 and p < 0.05 and ordered by median rank of pathway genes in NMF gene list. Redundant GO terms supported by overlapping genes were manually consolidated into single biological themes. Highly ranked and biologically interesting pathways were annotated. NNLS projection (nnls v1.6) was used to project factors on all spots; NNLS projection identifies the non-negative combination of factors that best reconstructs a spot’s expression and scores the spot appropriately for each factor. Metaprograms were defined by pairwise Jaccard similarity index (JSI) and manual annotation of the NMF factors and validated by pairwise hypergeometric testing of the original 2,000 variable genes. Metaprogram scores per spot were calculated by averaging the metaprogram’s constituent scaled (0-1 normalized) factor scores per spot.

### Spatial colocalization of gene signatures

To quantify directional spatial colocalization between pairs of module scores, we computed the global bivariate Moran’s I; this index describes if high expression of a module score in a spot predicts high expression of a second module score in k = 6 nearest neighbor spots, and vice versa. Global bivariate Moran’s I was computed as Pearson correlation between the z-scored expression of a module score and the spatially lagged z-scored expression of the second module score, using a k=6 nearest-neighbor spatial weights matrix (spdep v1.4-1) with 499 permutations per direction per pair per sample. Results were corrected using BH-FDR (padj = 0.05).

### Global spatial restriction of gene signatures and metaprograms

Global Moran’s I was used (FindSpatiallyVariableFeatures() Seurat v5.3.0, selection.method = moransi) to quantify the global spatial restriction of module scores and metaprograms on LN tumor spots. A RNA processing signature generated from Karras et al^33^ served as a negative control (Supplementary Table 5).

### Local spatial autocorrelation of neural crest-like and melanocytic programs

To identify local spatial clusters of spots (hotspots) of neural crest and melanocytic gene signatures, we calculated the local Moran’s I for each score per spot using localmoran() (spdep v 1.4-1) with k = 6 nearest neighbors and row-standardized weights. Raw local moran’s I values were Z-scored and p-values adjusted using BH-FDR (p_adj_ = 0.05). Spots were annotated as follows: High-high (padj < 0.05, spot z-score > 0, neighbor z-score > 0); high-low (p_adj_ < 0.05, spot z-score > 0, neighbor z-score < 0); low-low (padj < 0.05, spot z-score < 0, neighbor z-score < 0); low-high (padj < 0.05, spot z-score < 0, neighbor z-score > 0); non-significant (padj > 0.05). High-high spots therefore represent spots high for a signature surrounded by other spots high for that signature and were annotated as hotspots to be used for downstream analysis.

### Early and late LN CNV clones

#### Definition

To classify LN CNV clones along the evolutionary trajectory, clones below the median evolutionary distance from stroma were assigned as early and clones above the median as late. Per LN sample, the most extreme clone was selected (lowest evolutionary distance for early, highest for late).

#### Differential gene expression analysis

Pseudobulk gene expression profiles were generated for each of the early and late clones by aggregating raw counts across all spots assigned to that clone (AggregateExpression(), Seurat v5.3.0). Genes with fewer than 10 counts in at least 3 pseudobulk samples were removed prior to analysis. Differential expression between late and early clones was assessed using DESeq2 (v1.44) with an unpaired design. Genes which an absolute log2 fold change > 0.5 and BH-FDR p_adj_ < 0.05 were considered significantly differentially expressed.

#### Gene set enrichment analysis

Genes were ranked by log2 fold change (late vs. early) and Gene Set Enrichment Analysis (GSEA) was performed against Gene Ontology (GO) Biological Process gene sets (fgsea v.1.30, msigdbdf v24.1). To reduce redundancy among enriched terms, pairwise JSI of leading-edge gene sets was calculated, and terms with JSI > 0.5 were consolidated, retaining the term with the lowest p_adj_.

### The Cancer Genome Atlas (TCGA) Analysis

#### Data acquisition

Skin cutaneous melanoma TCGA clinical data was downloaded from https://www.cbioportal.org/study/summary?id=skcm_tcga_pan_can_atlas_2018. Samples were annotated using Tumor_Disease_Anatomic_Site to identify LN metastases and primary tumors with and without LN metastasis. Regional subcutaneous and distant metastatic samples were removed. To ensure only cutaneous samples were selected, glabrous skin samples as defined by the Firehose 2016 dataset (https://www.cbioportal.org/study/summary?id=skcm_tcga) were omitted from downstream analysis. Aneuploidy score (CNV score) was used as defined by Taylor et al^71^ and reported in the TCGA data.

#### Survival analysis

The TCGA FPKM expression matrix was downloaded via https://www.cbioportal.org/study/summary?id=skcm_tcga_pan_can_atlas_2018. The expression matrix was z-scored per gene across all samples, and a signature score per sample was calculated as the mean z-scored expression across all present signature genes. Survival analysis was performed with stratifying for low and high expression using the median across the tumors analyzed using surv() and survfit() in survival v3.8-3 and ggsurvplot() in survminer for visualization v0.5.1.

### Statistics and reproducibility

No statistical methods were used to predetermine sample size. Sample sizes reflect the number of patient specimens meeting the tissue and RNA quality criteria described above and are reported for each analysis in the corresponding figure legend. Non-parametric tests were used by default given the non-normal distribution of the biological data. Multiple comparisons were corrected using Benjamini-Hochberg false discovery (BH-FDR) correction. Plots show box plots (median and interquartile range) or individual data points, which display both the central tendency and spread of the data. Cliff’s Delta was calculated using the cliff.delta() function (effsize v0.8.1), defined as the probability that a randomly selected value from one group exceeds a randomly selected value from the other, minus the reverse probability. Cohen’s d was calculated as the difference in group means divided by the pooled standard deviation. Representative images shown (e.g. Fusion imaging, H&E, spatial projections) are representative of findings consistently observed across the samples analyzed in the relevant cohort for that panel. Where multiple spots, clones, or tumor fields were analyzed within the same tissue sample, sample size is reported in the figure legends to indicate both the total number of measurements and the number of independent samples/patients from which they were derived. For most analyses, hierarchical structure was addressed by aggregating measurements to the level of independent clones or samples prior to statistical testing. Density and CNV score comparisons between tumor spot categories were instead performed at the spot level, pooling spots across samples, to characterize within-tissue spatial density patterns rather than sample-level differences.

## Data availability

Raw and processed spatial transcriptomics data supporting the findings of this study have been deposited at the Gene Expression Omnibus (GEO) and will be made publicly available upon publication of the peer-reviewed article. Imaging data will be available upon request following publication.

## Code availability

This study did not involve the development of new software or algorithms. Analyses were performed using published tools and packages described in the Methods and Reporting Summary (Data Analysis). The custom code used for data analysis will be available from the corresponding author upon request following publication.

## Acknowledgements

The authors acknowledge technical assistance and feedback from Sabrina Solis, Joseph Daccache, Deborah Corcoran, Danielle Share, Yodit Girmay, and the rest of the Lund lab. We thank members of the Experimental Pathology Research Laboratory (RRID:SCR_017928) and NYU Langone’s Genome Technology Center (RRID: SCR_017929), which is partially supported by the Cancer Center Support Grant P30CA016087 at NYU Langone’s Laura and Isaac Perlmutter Cancer Center. This work was supported by grants from the Cancer Research Institute (Lloyd J. Old STAR Award to AWL), the American Association for Cancer Research (AACR-BMS Midcareer Female Investigator Award, AWL), the NIH (U54CA263001, AWL, MS, IY, IO; P50CA016087 to AWL, IY, and IO), and the DOD (W81XWH2110510 to AWL, MS, IY, IO). KSV was supported by the NIH (F30CA288142-01A). TM was supported by the CRI (CRI14497).

## Author contributions

Conceptualization: KSV, TM, IY, MS, AWL

Methodology: KSV, TM, RS, ZC, IIB,

Investigation: KSV, TM, RS

Funding acquisition: MS, AWL

Reagents and Biospecimens: SQ, ILB, IO, ASM

Supervision: TD, IO, MS, AWL

Writing – original draft: KSV, TM, IY, MS, AWL

Writing – review & editing: KSV, TM, RS, SQ, ZC, IIB, ASM, TD, IO, IY, MS, AWL

## Declaration of generative AI and AI-assisted technologies in the manuscript preparation process

During the preparation of this work the authors used NYU-proprietary generative AI to check grammar, clarity and consistency of text. NYU-proprietary generative AI and Claude Sonnet 4.6 were used for function generation and code clarity. After using these tools/services, the authors reviewed and edited the content as needed and take full responsibility for the content of the published article.

