## Extended Data Fig. 1-7 for "Melanoma evolution in the lymph node shapes systemic outcomes"

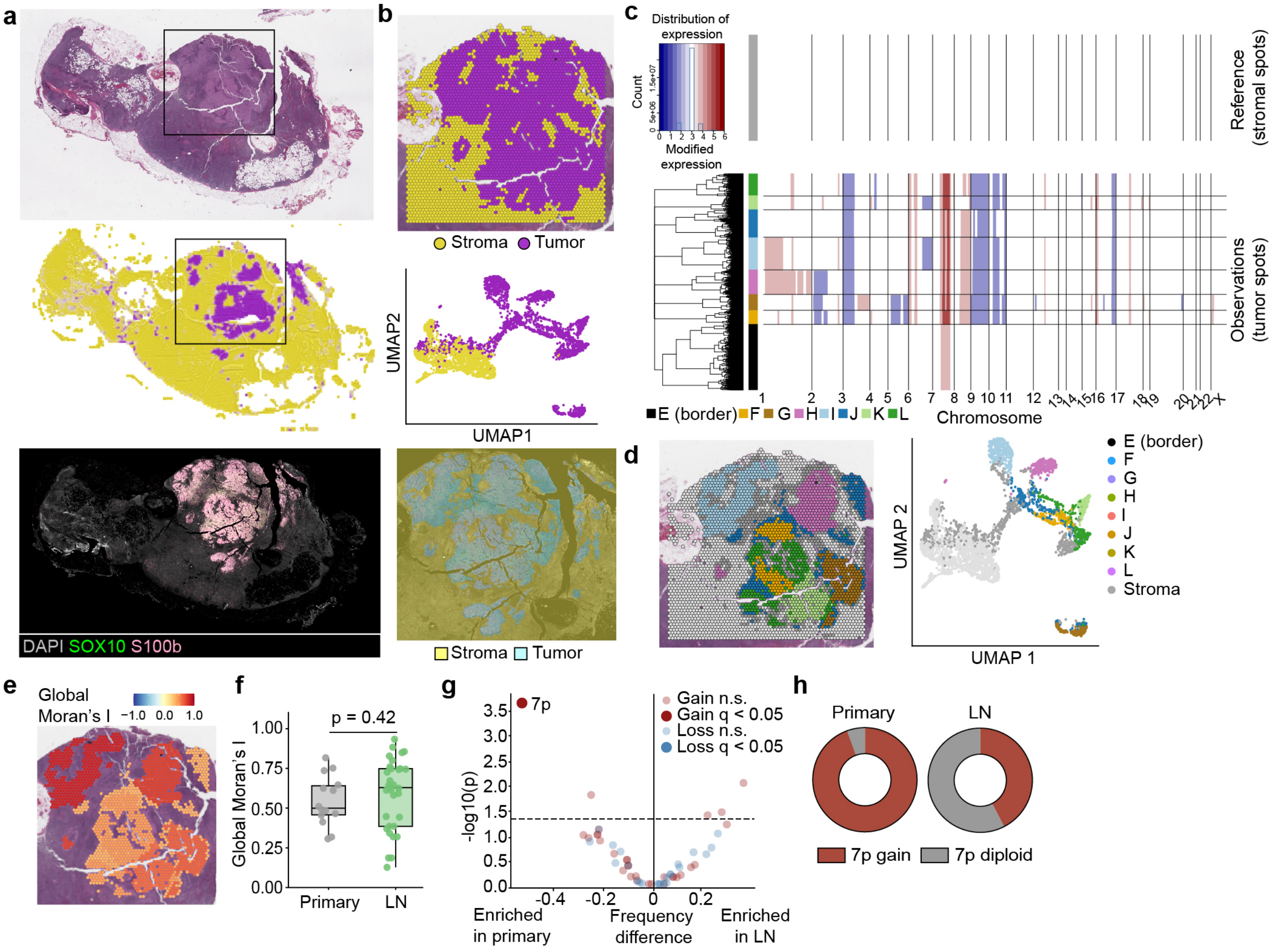


**Extended Data Fig. 1 | Clone identification in primary and LN metastatic melanomas. a,** Representative metastatic LN H&E (upper panel) with corresponding AI tumor detection used for ST placement (middle panel), and Quanterix Fusion whole-slide imaging (lower panel). **b,** Spatial map and Uniform Manifold Approximation and Projection (UMAP) of tumor and stromal spots (upper and middle panel); Fusion imaging of ST ROI highlighted by HALO tumor classifier (lower panel). **c,** Clone-defining CNV probability predictions by SpatialInferCNV Hidden Markov Model with Bayesian latent mixture model posterior probability corrections. **d,** Spatial map and UMAP of predicted clones (color), border clone (black), and stroma (grey). **e,** Representative spatial map of tumor spots colored by global Moran's I of clonal identity. **f,** Global Moran's I of primary ($n = 18$) and LN ($n = 33$) clones. **g-h,** Arm CNV enriched in primary ($n = 18$ from 8 tumors) and LN ($n = 33$ from 10 LNs) melanoma clones **(g)** and proportion of primary (left) and LN (right) tumor clones with gains in chromosome 7p **(h)**. Statistics: two-sided Mann-Whitney U test **(f)** and Fisher's exact test with Benjamini-Hochberg False Discovery Rate (BH-FDR) for multiple comparisons **(g)**. Significant = $q < 0.05$.

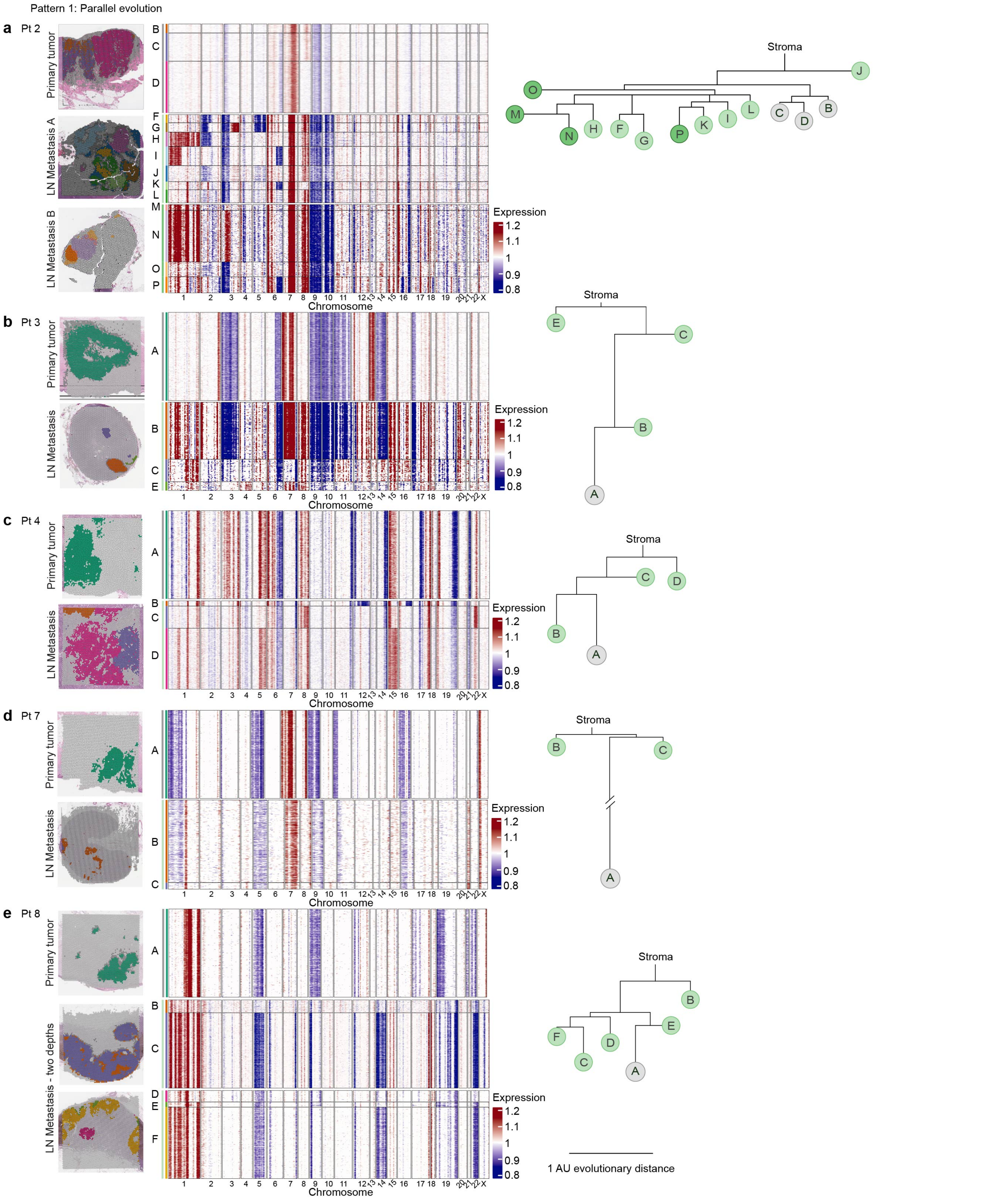


**Extended Data Fig. 2 | LN melanomas are clonally heterogeneous and follow two patterns of evolution. a-e,** Spatial map of paired primary and LN tumors colored by clone (left) and maximum likelihood trees with edge length distances (right) for melanomas with parallel evolutionary patterns. Scale bar represents 1 arbitrary unit (AU) of evolutionary distance.

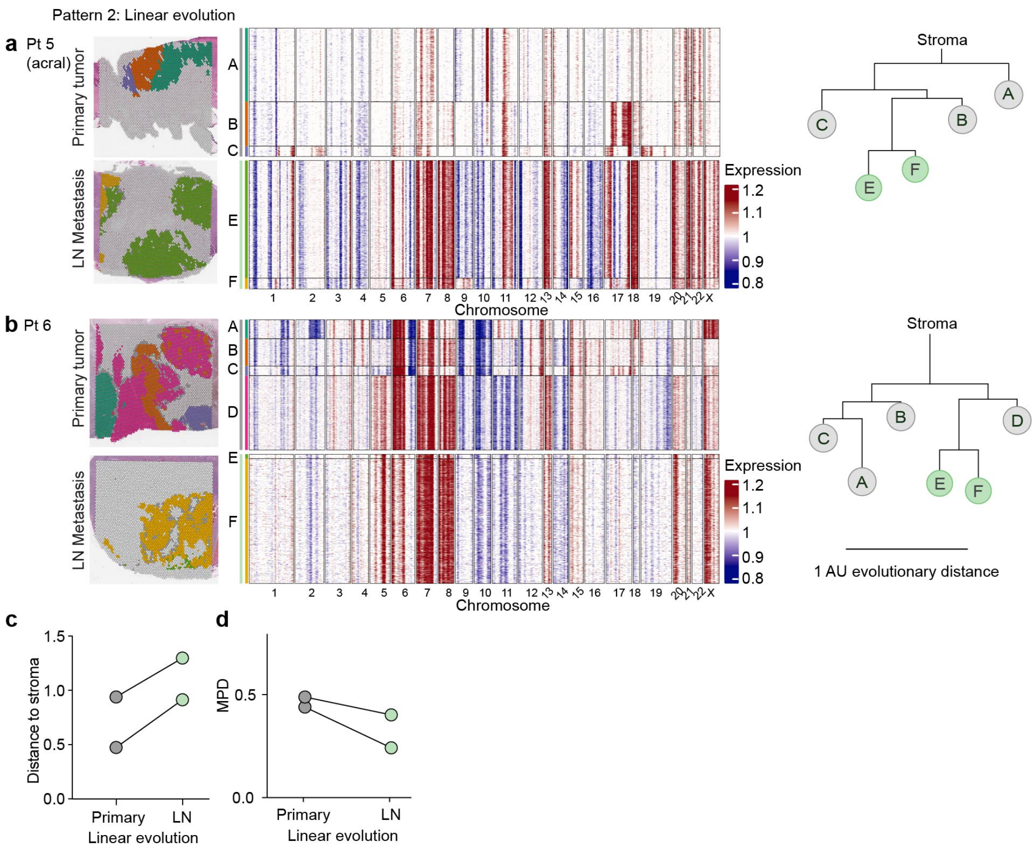


**Extended Data Fig. 3 | LN melanomas are clonally heterogeneous and follow two patterns of evolution. a-b,** Spatial map of paired primary and LN tumors colored by clone (left) and maximum likelihood trees with edge length distances (right) for melanomas with linear evolutionary patterns. **c,** Mean estimated distance to stroma per patient of primary and LN metastatic clones in patients with linear evolution (n = 2 pairs). **d,** Mean pairwise distance (MPD) between clones within primary and LN tumors from patients with linear evolution (n = 2 pairs). Scale bar represents 1 arbitrary unit (AU) of evolutionary distance.

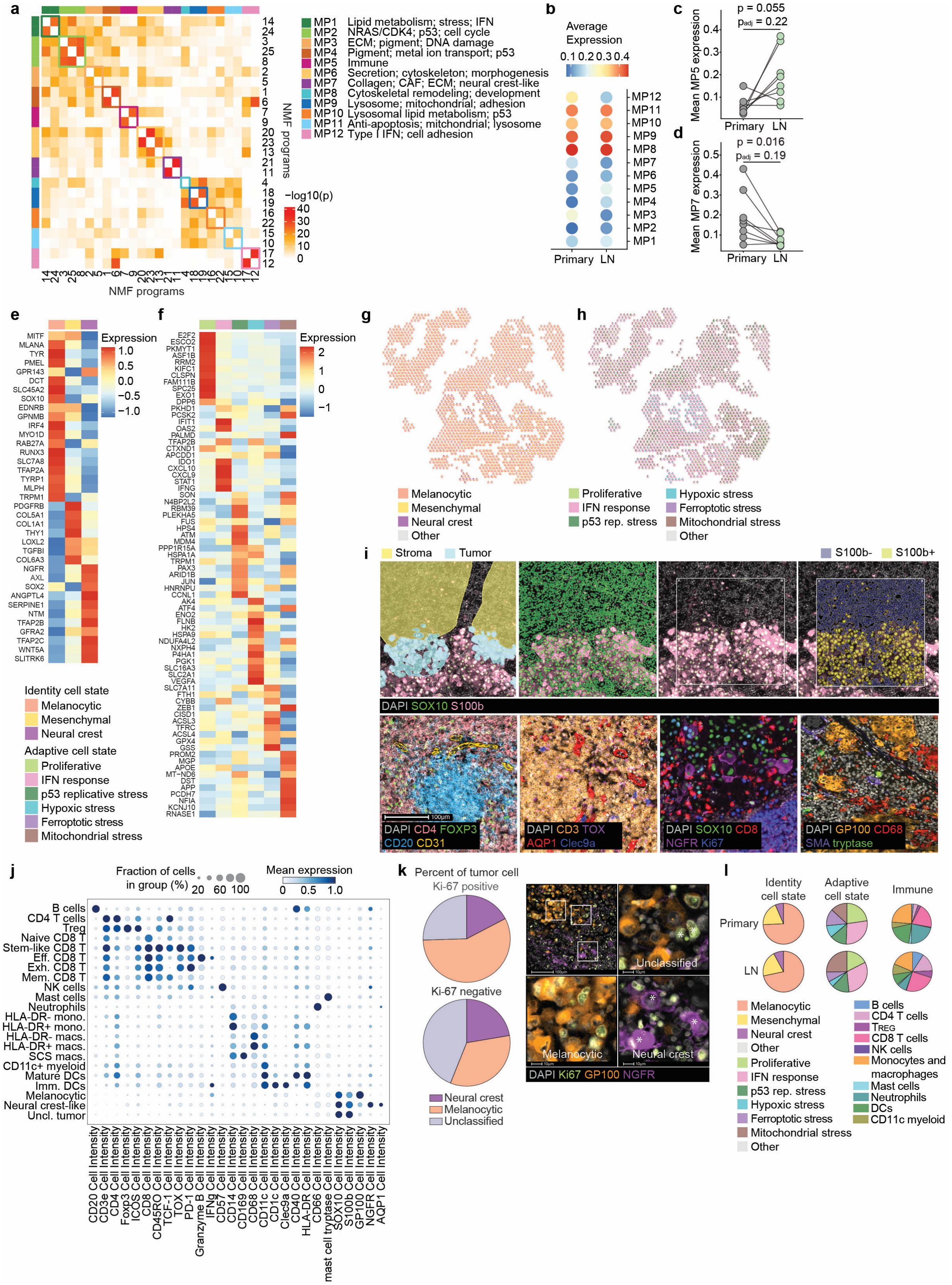

**Extended Data Fig. 4 | Phenotypic heterogeneity in treatment-naïve melanoma primary tumors and LN metastases.** **a,** Gene overlap significance between non-negative matrix factorization (NMF) factors (-log10 hypergeometric p-value), ordered by hierarchical clustering of the Jaccard Similarity Index (JSI) matrix (Euclidean distance); colored by metaprograms. **b,** Average expression of NMF metaprograms in paired primary and LN tumors (n = 8 pairs). **c-d,** Mean expression of metaprogram 5 **(c)** and metaprogram 7 **(d)** in primary and LN tumor spots (n = 8 pairs). **e-f,** Average gene expression of consensus identity **(e)** and adaptive **(f)** signatures, grouped by dominant state in tumor spots (n = 10 LNs from 8 patients). **g-h,** Representative spatial projection of identity **(g)** and adaptive **(h)** state deconvolution. **i,** Representative

ROI of imaging pipeline. Top: stroma/tumor annotations for tumor classifier training, nuclear segmentation, and S100b threshold detection. Bottom: images of B cell follicle, T cell zone, neural crest-like tumor, melanocytic tumor. All scale bars, 100 µm. **j,** Key imaging cell annotation markers, scaled by marker. **k,** Proportion of proliferative ($Ki\text{-}67^{+}$) and non-proliferative ($Ki\text{-}67^{-}$) LN tumor cells by imaging in each cell state ($X^2$ = 76938, p = 0.2.2e-16, df = 2, n = 391,897 neural crest, n = 720,511 melanocytic, n = 735,361 unclassified) and representative image. Scale bar, 100 µm and 10 µm. **l,** Deconvolution-estimated proportion of identity and adaptive tumor cell states and immune cell proportion by imaging in primary and LN tumors, averaged per sample and per patient (n = 8; identity $X^2$ = 0.17, p = 0.92, df = 2; adaptive $X^2$ = 6.04, p = 0.30, df = 5; immune $X^2$ = 3326, p = 2.2e-16, df = 9). Statistics: paired two-tailed Wilcoxon test with BH-FDR correction **(c-d)** and Pearson's chi-squared test on tumor cells by imaging **(k,l)** and sample-aggregated deconvolution-estimated fractions **(l)**.

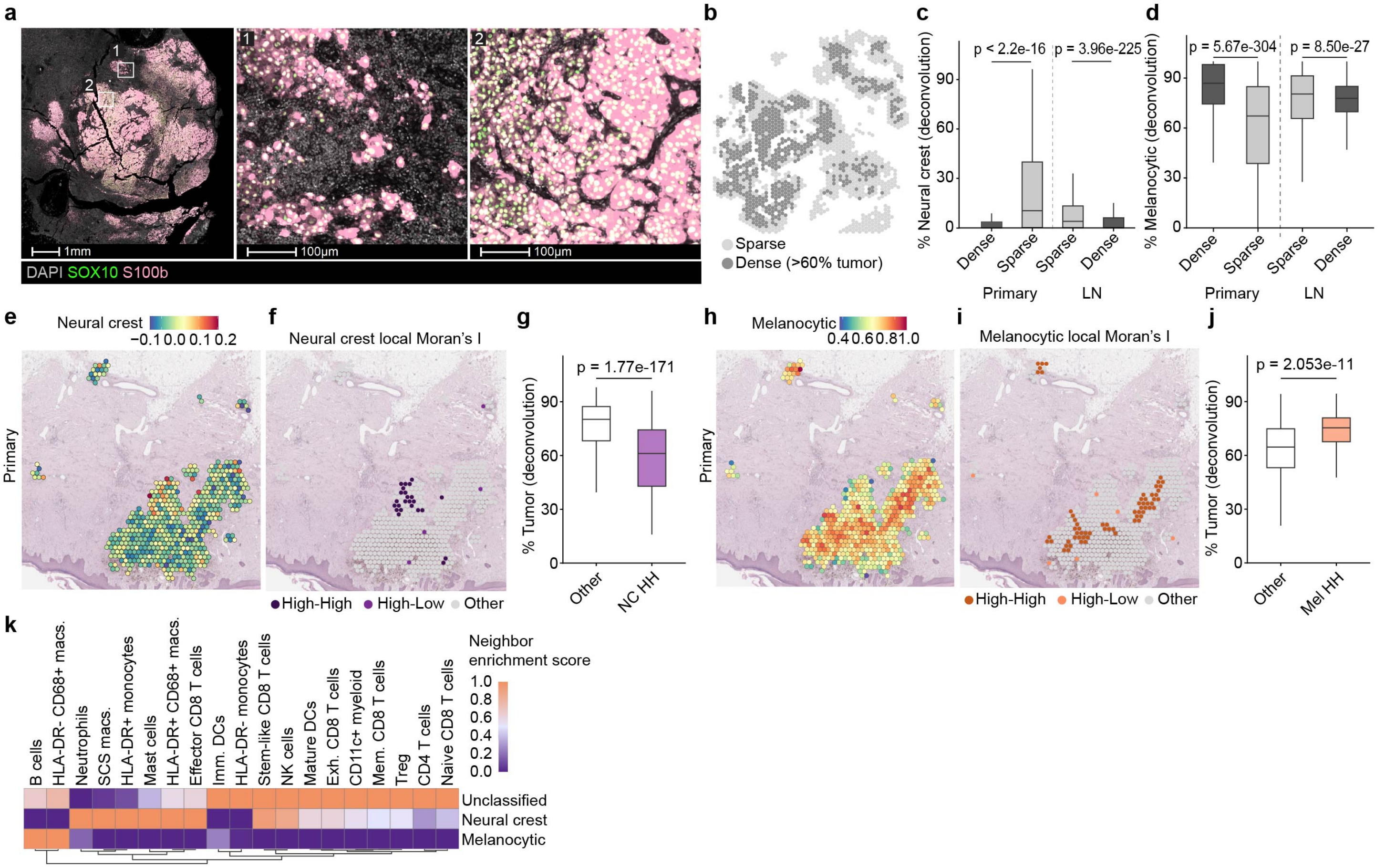


**Extended Data Fig. 5 | Melanoma cells de-differentiate in the primary tumor before metastasis. a,** Representative images of sparse (middle panel) and dense (right panel) tumor regions. Scale bar, 1 mm (left), 100 μm (middle, right). **b,** Representative annotation of sparse and dense tumor spots. Dense tumor spots are defined as > 60% tumor by deconvolution. **c,** Deconvolution-estimated fraction of neural crest-like tumor cells per spot in dense and sparse spots in primary (n = 11,831 dense; n = 2,161 sparse spots in n = 8 tumors from 8 patients) and LN (n = 9,281 dense; n = 4,877 sparse spots in n = 10 LNs from 8 patients) melanomas (Cliff's Delta: primary tumors 0.60; LNs -0.33) **d,** Same as **(c)** for melanocytic tumor cells (Cliff's Delta: primary tumors 0.50; LNs -0.11). **e-f,** Representative spatial projection of NC signature **(e)** and hotspots **(f)** in primary melanoma. **g,** Tumor density by deconvolution of primary NC hotspots (n = 1,010) versus other (n = 12,982) primary tumor spots (Cliff's Delta: -0.53). **h-i,** Representative spatial projection of melanocytic signature **(h)** and hotspots **(i)** in primary melanoma. **j,** Tumor density by deconvolution of melanocytic hotspots (n = 1,754) compared to other (n = 12,238) primary tumor spots (Cliff's Delta: 0.10). **k,** Heatmap (scaled by row) of neighbor enrichment score for melanocytic ($GP100^{+}$), neural crest-like ($NGFR^{+}$ or $AQP1^{+}$) or unclassified ($GP100^{-}NGFR^{-}AQP1^{-}$) tumor cells (n = 14 LNs from 12 patients). Statistics: Mann-Whitney U test with Cliff's Delta (negligible <0.11; small 0.11-0.28; medium 0.28-0.43; large >0.43 **(c, d, g, j)**.

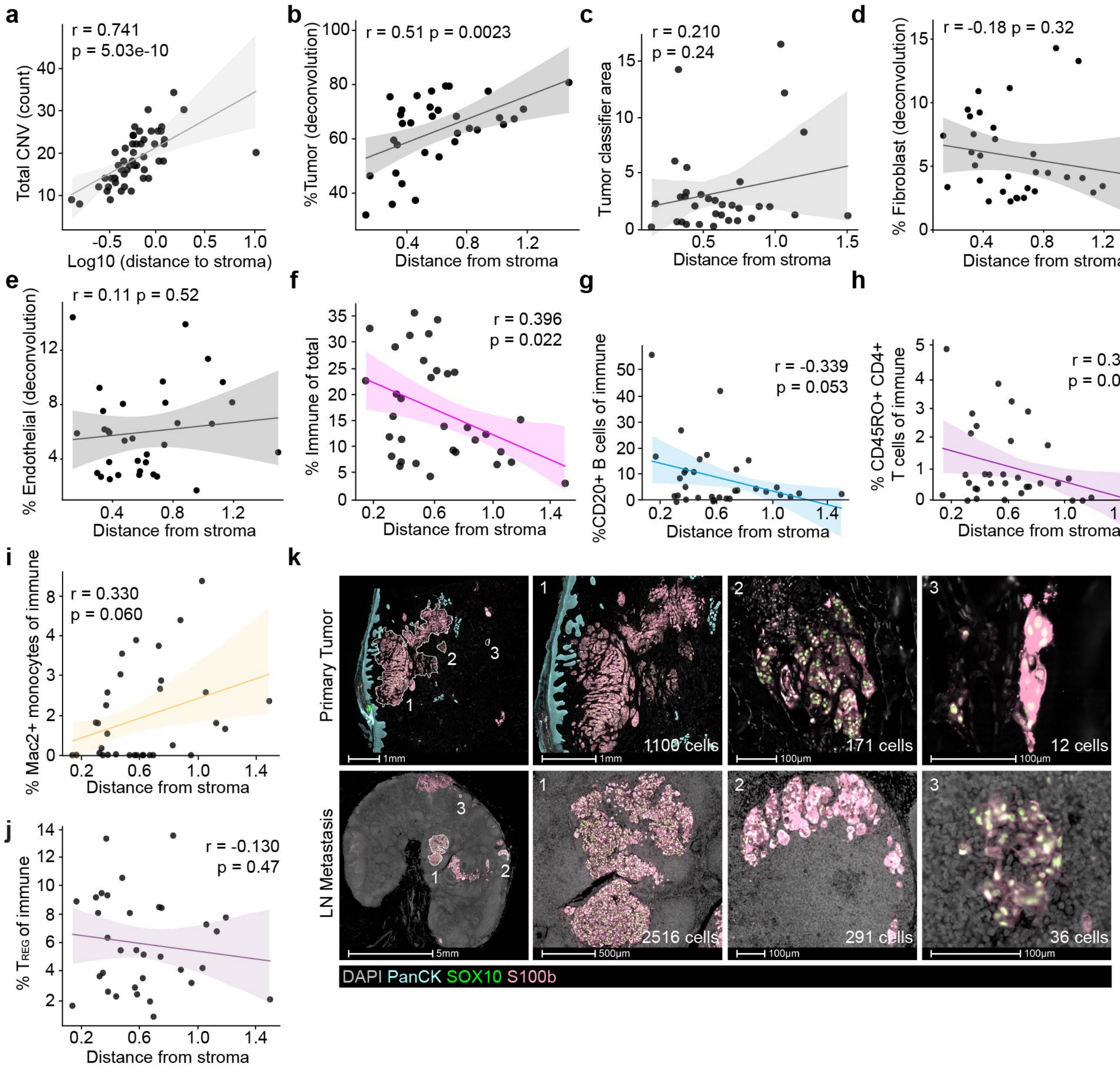


**Extended Data Fig. 6 | Microenvironment shifts as a function of LN tumor evolution. a,** Total CNV count over Log10 evolutionary distance (n = 51 clones from 8 primary tumors and 10 LNs). **b,** Mean spot tumor fraction by deconvolution per LN clone, over evolutionary distance (n = 33 clones from 10 LNs). **c,** Clone size by tumor classifier area over evolutionary distance. **d-e,** Mean spot fibroblast **(d)** and endothelial **(e)** fraction by deconvolution per LN clone, over evolutionary distance (n = 33 clones from 10 LNs). **f,** Percent immune cells of all cells in tumor classifier area per LN clone over evolutionary distance (n = 33 clones from 10 LNs). **g-j,** Percent CD20$^+$ B cells **(g)**, antigen-experienced (CD45RO$^+$) CD4$^+$ CD3$^+$ T cells **(h)**, Mac2$^+$ monocytes **(i)**, and CD4$^+$FOXP3$^+$ regulatory T cells ($T_{REG}$) **(j)** of all immune cells over evolutionary distance in LN clones (n = 33 clones from 10 LNs). **k,** Representative images of primary (top) and LN (bottom) tumor field sizes and locations. Scale bars top: 1 mm, 1 mm, 100 μm, 100 μm; bottom: 5 mm, 500 μm, 100 μm, 100 μm. Statistics: Pearson correlation **(a-j)**.

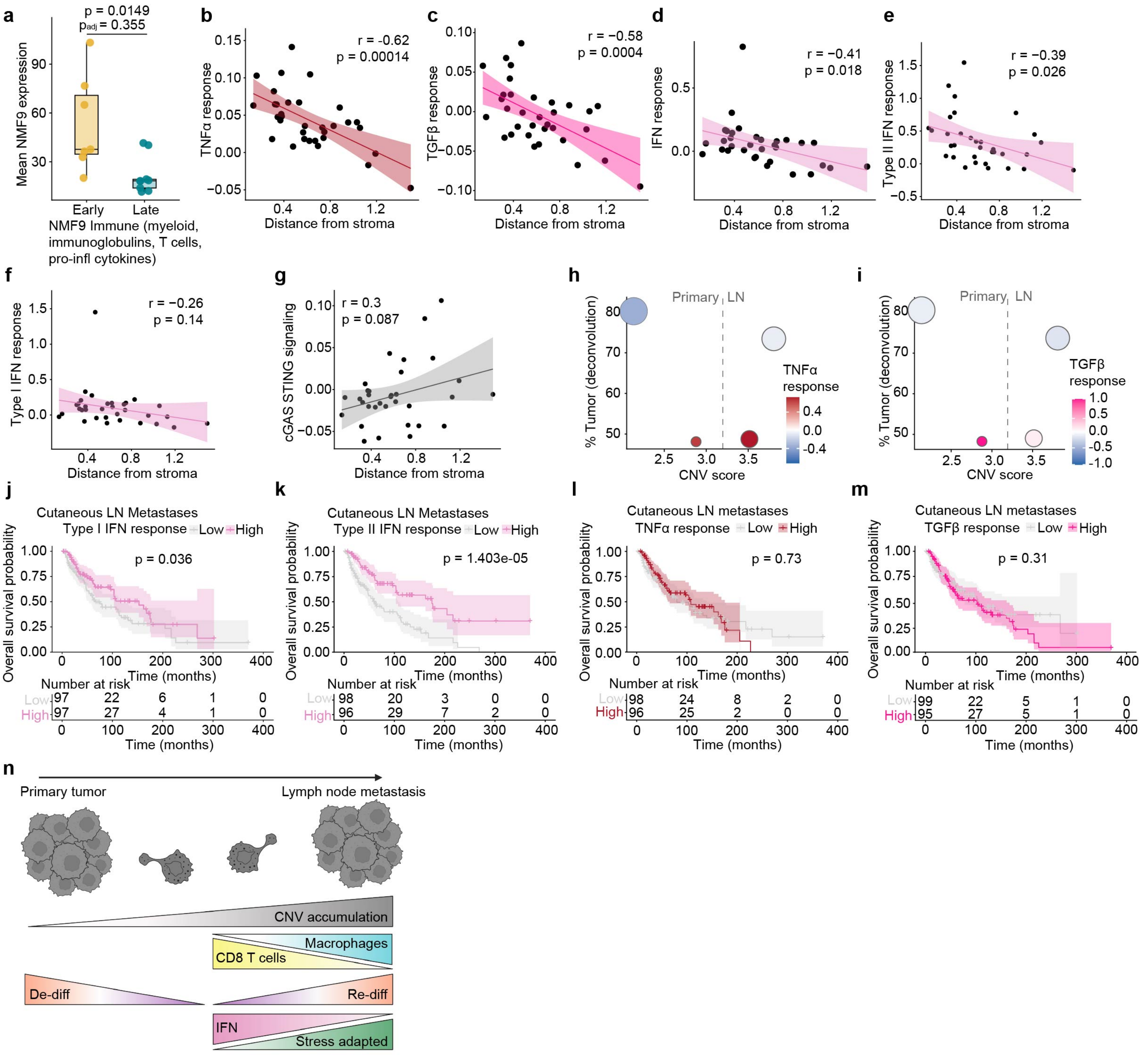


**Extended Data Fig. 7 | Evolution of LN metastasis correlates with decreased immune reactivity. a,** Mean expression of NMF program 9 in early (n = 7) and late (n = 9) LN clones. **b-g,** Mean expression of TNFα **(b)**, TGFβ **(c)**, IFN response **(d)**, type I IFN response **(e)**, type II IFN response **(f)**, and cGAS-STING activation **(g)** signatures per LN clone (n = 33 clones from 10 LNs) correlated with evolutionary distance (gene signatures: **Supplementary Table 5**). **h-i,** Tumor density by deconvolution and CNV score of primary (dense n = 11,831, sparse n = 2,161; 8 tumors) and LN (dense n = 9,281, sparse n = 4,877; 10 LNs from 8 patients) spots, colored by expression of TNFα **(h)** and TGFβ **(i)** signatures. **j-m,** Survival of patients with cutaneous melanoma (n = 194) stratified by LN expression of type I IFN **(j)**, type II IFN **(k)**, TNFα **(l)**, and TGFβ **(m)** response signatures. **n,** model of melanoma evolution and adaptation during LN metastasis. Statistics: Mann-Whitney U test with BH-FDR correction **(a)**, Pearson correlation **(b-g)**, and Kaplan-Meier survival curve with log-rank Mantel-Cox test **(j-m)**.
