## Supplementary Table 1 for "Melanoma evolution in the lymph node shapes systemic outcomes"

| Pt | Paired primary - LN? | Primary Melanoma Anatomic Location | Sex | Age at Excision | Stage at Excision | Type | Primary tumor thickness | Mutation status | Tx before excision |
|---|---|---|---|---|---|---|---|---|---|
| 1 | Paired | Left foot, digits 2 and 3 | Male | 74 | Stage IIIB | Acral lentiginous | 8 | No data | No |
| 2 | Paired | Left upper back | Male | 69 | Stage IIIC | Cutaneous | 3.1 | BRAF V600K, FGFR1 I234T, FGFR2 S372F | No |
| 3 | Paired | Right Upper Back | Female | 62 | Stage IIIC | Cutaneous | 6 | BRAF V600E, ALK T680I | No |
| 4 | Paired | Mid back skin | Male | 66 | Stage IIID | Cutaneous | 12 | No data | No |
| 5 | Paired | Left plantar forefoot | Female | 62 | Stage IIIC | Acral lentiginous | 6.8 | No data | No |
| 6 | Paired | Chest Wall | Male | 59 | Stage IIIC | Cutaneous | 10 | BRAF V600E | No |
| 7 | Paired | Left lower back | Male | 62 | Stage IIIC | Cutaneous | 4.1 | BRAF V600E, ALK T680I | No |
| 8 | Paired | Right Upper Arm | Male | 62 | Stage IIIC | Cutaneous | 2.5 | NRAS Q61K | No |
| 9 | LN only | Right Leg | Male | 56 | Stage IIIC | Cutaneous | 6.9 | BRAF V600E | No |
| 10 | LN only | Right posterior thigh | Female | 67 | Stage IIIC | Cutaneous | 2.85 | BRAF V600E, ALK G667R, SMO V270I | No |
| 11 | LN only | Left Upper Back | Male | 28 | Stage IIIB | Cutaneous | 2.2 | No data | No |
| 12 | LN only | Left upper back | Male | 81 | Stage IIIC | Cutaneous | 4.2 | N/A (BRAF negative) | No |

| Adjuvant Tx after excision | Any progression/recurrence after excision (Y/N) | Notes |
|---|---|---|
| N/A | Y | |
| nivo | N | Two LN resected |
| dab+tram | N | |
| Temodar | Y (muscle met) | |
| N/A | N/A | |
| ipi+nivo+toci (1 dose) | Y (liver met, bone met) | |
| nivo/nivo+rela, nivo (1 dose) | N | |
| No | Y (multiple cutaneous, LN mets, liver met, brain mets) | |
| Temodar + Thalidomide | Y (iliac, pelvic, popliteal fossa LNs) | Received ICI after further progression, and progressed again |
| No | Y (LN) | |
| Vaccine trial | N | |
| No | N | |
