## Supplementary Table 2 for "Melanoma evolution in the lymph node shapes systemic outcomes"

| Protein | Fluorescence channel | Clone | Barcode | Quanterix or Custom conjugation |
|---|---|---|---|---|
| CD20 | AF750 | L26 | BX007 | Quanterix |
| CD8 | Atto550 | CB/144B | BX026 | Quanterix |
| HLA-DR | AF647 | EPR3692 | BX033 | Quanterix |
| aSMA | AF750 | 1A4 | BX013 | Quanterix |
| Podoplanin | Atto550 | NC-08 | BX023 | Quanterix |
| CD4 | AF647 | EPR6855 | BX003 | Quanterix |
| Aquaporin1 | AF750 | EPR11588(B) | BX034 | Custom |
| Ki67 | Atto550 | B56 | BX047 | Quanterix |
| ICOS | AF647 | D1K2T | BX045 | Quanterix |
| CD31 | AF750 | WM59 | BX032 | Quanterix |
| Mac2/Gal3 | Atto550 | M3/38 | BX035 | Quanterix |
| CD68 | AF647 | KP1 | BX015 | Quanterix |
| B-catenin | Atto550 | 12F7 | BX096 | Quanterix |
| CD3e | AF647 | EP449E | BX045 | Quanterix |
| PanCK | AF750 | AE-1/AE-3 | BX019 | Quanterix |
| CD103 | Atto550 | EPR4166(2) | BX055 | Custom |
| FOXP3 | AF647 | 23RA/E7 | BX031 | Quanterix |
| SOX10 | AF750 | A-2 | BX004 | Custom |
| CD45RO | Atto550 | UCHL1 | BX017 | Quanterix |
| TCF-1 | AF647 | C63D9 | BX061 | Quanterix |
| IL-33 | AF750 | EPR20417 | BX022 | Custom |
| CD14 | Atto550 | EPR3653 | BX037 | Quanterix |
| PD-1 | AF647 | D4W2J | BX046 | Quanterix |
| S100b | AF750 | E7C3A | BX025 | Custom |
| Granzyme B | Atto550 | D6E9W | BX041 | Quanterix |
| PD-L1 | AF647 | 73-10 | BX043 | Quanterix |
| CSPG4 | AF750 | EPR9195 | BX040 | Custom |
| CD57 | Atto550 | HNK-1 | BX028 | Custom |
| Collagen IV | AF647 | EPR20966 | BX042 | Quanterix |
| IFNg | Atto550 | EPR21704 | BX020 | Quanterix |
| IDO1 | AF647 | V1NC3IDO | BX027 | Quanterix |
| CD21 | Atto550 | EP3093 | BX001 | Quanterix |
| CD11c | AF647 | D3V1E | BX024 | Custom |
| TOX | Atto550 | E6I3Q | BX060 | Quanterix |
| CD40 | AF647 | D8W3N | BX010 | Quanterix |
| CD1c | Atto550 | EPR23189-196 | BX094 | Custom |
| NGFR | AF647 | D4B3 | BX016 | Custom |
| Clec9a | Atto550 | EPR22324 | BX005 | Custom |
| Axl | AF647 | Polyclonal | BX030 | Custom |
| tryptase | Atto550 | AA1 | BX014 | Custom |

| | | | | |
|---|---|---|---|---|
| CD169 | AF647 | SP216 | BX006 | Custom |
| GP100 | Atto550 | HMB45 | BX094 | Quanterix |
| PNAd | Atto550 | MECA-79 | BX029 | Custom |
| CD66a/c/e | AF647 | ASL-32 | BX096 | Quanterix |
