## Supplementary Table 3 for "Melanoma evolution in the lymph node shapes systemic outcomes"

| Pt | Sample | Tissue | ST size | # total spots (ST) | total area (mm2) (imaging) | mean nCount (ST) | mean nFeature (ST) | # of tumor spots (ST) |
|---|---|---|---|---|---|---|---|---|
| 1 | 1_Primary | Primary | 6.5x6.5mm | 2,931 | 5.79 | 36,341 | 5,776 | 712 |
| 1 | 1_LN | LN | 6.5x6.5mm | 4,734 | 29.12 | 37,457 | 8,029 | 2,448 |
| 2 | 2_Primary | Primary | 11x11mm | 8,116 | 36.88 | 3,366 | 2,003 | 5,248 |
| 2 | 2_LN_A | LN | 6.5x6.5mm | 3,622 | 20.79 | 29,900 | 8,164 | 1,533 |
| 2 | 2_LN_B | LN | 6.5x6.5mm | 2,033 | 4.22 | 30,892 | 8,203 | 449 |
| 3 | 3_Primary | Primary | 6.5x6.5mm | 4,480 | 16.57 | 9,056 | 3,429 | 1,731 |
| 3 | 3_LN | LN | 6.5x6.5mm | 3,069 | 2.17 | 47,229 | 9,092 | 212 |
| 4 | 4_Primary | Primary | 6.5x6.5mm | 4,303 | 15.34 | 12,444 | 4,414 | 1,282 |
| 4 | 4_LN | LN | 6.5x6.5mm | 4,990 | 35.38 | 13,259 | 5,120 | 2,948 |
| 5 | 5_Primary | Primary | 6.5x6.5mm | 3,713 | 14.13 | 20,512 | 5,955 | 1,032 |
| 5 | 5_LN | LN | 6.5x6.5mm | 4,750 | 24.08 | 32,690 | 7,824 | 1,919 |
| 6 | 6_Primary | Primary | 6.5x6.5mm | 4,505 | 27.39 | 30,703 | 7,376 | 2,901 |
| 6 | 6_LN | LN | 6.5x6.5mm | 4,534 | 12.37 | 10,917 | 4,056 | 1,172 |
| 7 | 7_Primary | Primary | 6.5x6.5mm | 3,581 | 9.26 | 11,963 | 3,661 | 576 |
| 7 | 7_LN | LN | 11x11mm | 12,230 | 5.03 | 2,775 | 1,824 | 515 |
| 8 | 8_Primary | Primary | 6.5x6.5mm | 4,175 | 4.30 | 8,615 | 3,306 | 510 |
| 8 | 8_LN_section1 | LN | 6.5x6.5mm | 4,340 | 17.60 | 27,383 | 6,335 | 2,140 |
| 8 | 8_LN_section2 | LN | 6.5x6.5mm | 4,264 | 8.69 | 20,939 | 5,222 | 822 |

| # of CNV clones (ST) | tumor area (mm2) (imaging) | # of CNV clones normalized to tumor area | mean nCount per tumor spot (ST) |
|---|---|---|---|
| 4 | 3.25 | 1.23 | 84,000 |
| 5 | 17.57 | 0.28 | 46,255 |
| 3 | 33.37 | 0.09 | 4,386 |
| 7 | 13.23 | 0.53 | 35,248 |
| 4 | 3.51 | 1.14 | 36,891 |
| 1 | 7.94 | 0.13 | 16,899 |
| 3 | 1.59 | 1.88 | 133,022 |
| 1 | 11.03 | 0.09 | 19,462 |
| 3 | 21.57 | 0.14 | 15,232 |
| 3 | 5.34 | 0.56 | 37,550 |
| 2 | 18.41 | 0.11 | 46,645 |
| 4 | 21.55 | 0.19 | 36,490 |
| 2 | 9.80 | 0.20 | 15,001 |
| 1 | 5.88 | 0.17 | 37,233 |
| 2 | 3.14 | 0.64 | 7,550 |
| 1 | 2.47 | 0.40 | 25,589 |
| 2 | 12.69 | 0.16 | 45,230 |
| 3 | 5.74 | 0.52 | 55,235 |

| mean nFeature per tumor spot (ST) | mean tumor fraction by deconvolution (ST) | mean tumor density (imaging) |
|---|---|---|
| 9,894 | 66.10 | 54.19 |
| 8,756 | 73.62 | 61.47 |
| 2,550 | 86.85 | 90.69 |
| 8,479 | 58.84 | 66.99 |
| 8,329 | 75.31 | 81.10 |
| 5,545 | 69.83 | 47.90 |
| 11,075 | 66.45 | 64.88 |
| 6,026 | 70.94 | 71.86 |
| 5,633 | 62.14 | 65.89 |
| 8,726 | 46.84 | 37.96 |
| 9,146 | 67.42 | 73.70 |
| 8,242 | 73.73 | 81.66 |
| 4,695 | 70.83 | 82.21 |
| 7,722 | 72.70 | 63.48 |
| 3,686 | 46.67 | 59.21 |
| 6,954 | 69.83 | 57.52 |
| 8,745 | 61.01 | 54.68 |
| 9,252 | 62.15 | 68.64 |
