## Supplementary Table 4 for "Melanoma evolution in the lymph node shapes systemic outcomes"

| sample_clone | Tissue | 1p | 1q | 2p | 2q | 3p | 3q | 4p | 4q | 5p | 5q | 6p | 6q | 7p | 7q | 8p | 8q | 9p | 9q | 10p | 10q | 11p |
|---|---|---|---|---|---|---|---|---|---|---|---|---|---|---|---|---|---|---|---|---|---|---|
| 026_A1_LN_Clone_F | LN | 1 | 1 | -1 | 1 | -1 | 0 | 1 | 0 | 0 | -1 | 1 | 0 | 0 | 1 | 1 | 1 | -1 | -1 | -1 | -1 | 0 |
| 026_A1_LN_Clone_G | LN | 0 | 1 | -1 | 1 | -1 | 1 | 0 | 0 | 0 | -1 | 1 | 0 | 0 | 1 | 1 | 1 | -1 | -1 | -1 | -1 | 0 |
| 026_A1_LN_Clone_H | LN | 1 | 1 | -1 | -1 | -1 | 0 | 0 | 0 | 0 | 0 | 1 | 0 | 0 | 1 | 0 | 1 | -1 | -1 | -1 | -1 | 0 |
| 026_A1_LN_Clone_I | LN | 1 | 1 | 0 | 1 | -1 | 0 | 0 | 0 | 0 | 0 | 1 | -1 | 0 | 1 | 0 | 1 | -1 | -1 | -1 | -1 | 0 |
| 026_A1_LN_Clone_J | LN | 0 | 0 | 0 | 0 | 0 | 0 | 0 | 0 | 0 | 0 | 1 | 0 | 0 | 1 | 0 | 1 | 0 | 0 | 0 | -1 | 0 |
| 026_A1_LN_Clone_K | LN | 0 | 1 | 1 | 0 | -1 | 0 | 1 | 0 | 0 | 0 | 1 | -1 | 0 | 1 | 0 | 1 | -1 | -1 | -1 | -1 | 0 |
| 026_A1_LN_Clone_L | LN | 0 | 1 | 0 | 1 | -1 | 0 | 0 | 0 | 0 | 0 | 1 | 0 | 0 | 1 | 0 | 1 | -1 | -1 | -1 | -1 | 0 |
| 026_B1_LN_Clone_M | LN | 1 | 1 | 0 | 0 | 0 | 0 | 1 | 0 | 0 | 0 | 1 | 0 | 0 | 1 | 1 | 1 | -1 | -1 | -1 | -1 | 0 |
| 026_B1_LN_Clone_N | LN | 1 | 1 | 0 | 1 | 1 | 0 | 1 | -1 | 0 | 0 | 1 | 0 | 0 | 1 | 1 | 1 | -1 | -1 | -1 | -1 | 0 |
| 026_B1_LN_Clone_O | LN | 1 | 1 | 0 | 0 | -1 | 0 | 0 | 0 | 0 | 0 | 1 | 0 | 0 | 1 | 0 | 1 | -1 | -1 | -1 | -1 | 0 |
| 026_B1_LN_Clone_P | LN | 1 | 1 | 1 | 0 | -1 | 0 | 1 | 0 | 0 | 0 | 1 | -1 | 1 | 1 | 1 | 1 | -1 | -1 | -1 | -1 | 0 |
| 026_primary_Clone_B | Primary | 0 | 0 | 0 | 1 | -1 | 0 | 0 | 0 | 0 | 0 | 0 | 0 | 1 | 1 | 0 | 1 | -1 | -1 | 0 | -1 | 0 |
| 026_primary_Clone_C | Primary | 0 | 0 | 0 | 0 | -1 | 0 | 0 | 0 | 0 | 0 | 1 | -1 | 1 | 1 | 0 | 1 | -1 | -1 | 0 | -1 | 0 |
| 026_primary_Clone_D | Primary | 0 | 0 | 0 | 0 | -1 | 0 | 0 | 0 | 0 | 0 | 1 | -1 | 1 | 1 | 0 | 0 | -1 | -1 | 0 | -1 | 0 |
| 070_LN_Clone_A | LN | 0 | 1 | 1 | 0 | 0 | 1 | 1 | 0 | 0 | 0 | 1 | -1 | 0 | -1 | 1 | 1 | 0 | 0 | 0 | 0 | 0 |
| 070_LN_Clone_B | LN | 0 | 1 | 0 | 0 | 0 | 1 | 0 | 0 | 0 | 0 | 0 | 0 | 0 | -1 | 1 | 1 | 0 | 0 | 0 | 0 | 0 |
| 070_LN_Clone_C | LN | 1 | 1 | 0 | 0 | 0 | 1 | 0 | 0 | 0 | 1 | 1 | -1 | 0 | -1 | 0 | 1 | 0 | 0 | 0 | 0 | 0 |
| 070_primary_Clone_A | Primary | -1 | 1 | 0 | 0 | 1 | 1 | 0 | -1 | 0 | 1 | 1 | -1 | 0 | -1 | 1 | 1 | 0 | 0 | 0 | 0 | 0 |
| 142_LN_Clone_F | LN | 0 | 1 | 0 | 0 | 0 | 0 | 0 | 0 | 0 | 0 | 0 | 0 | 1 | 1 | 1 | 1 | 0 | 0 | 0 | 0 | 0 |
| 142_LN_Clone_G | LN | 0 | 1 | 0 | 0 | 0 | 0 | 0 | 0 | 0 | 0 | -1 | -1 | 1 | 0 | 1 | 1 | 0 | 0 | 0 | 0 | 0 |
| 142_LN_Clone_H | LN | 0 | 1 | 0 | 0 | 0 | 0 | -1 | -1 | 0 | 0 | 0 | 0 | 1 | 1 | 0 | 0 | 0 | -1 | 0 | 0 | 0 |
| 142_LN_Clone_I | LN | 0 | 1 | 0 | 0 | 0 | 0 | 0 | 0 | 0 | 1 | 0 | 0 | 1 | 1 | 1 | 1 | 1 | 1 | 0 | 1 | 0 |
| 142_LN_Clone_J | LN | 1 | 1 | 0 | 1 | 0 | -1 | 0 | 1 | 0 | 0 | -1 | -1 | 1 | 0 | 1 | 1 | 0 | 1 | 0 | -1 | 0 |
| 142_primary_Clone_A | Primary | 0 | 1 | 0 | 1 | 0 | 0 | 0 | 1 | 0 | 0 | 0 | 0 | 1 | 1 | 1 | 1 | 0 | 0 | 0 | -1 | 0 |
| 142_primary_Clone_B | Primary | 0 | 1 | 0 | -1 | 1 | 0 | 0 | -1 | 0 | 1 | 0 | -1 | 1 | 1 | 1 | 1 | 0 | 0 | 0 | 1 | 0 |
| 142_primary_Clone_C | Primary | 1 | 1 | -1 | -1 | 1 | 0 | -1 | -1 | 0 | 1 | -1 | -1 | 1 | 1 | 1 | 1 | 0 | 1 | 0 | 1 | 0 |
| 142_primary_Clone_D | Primary | 0 | 1 | 0 | 0 | 1 | 0 | 0 | -1 | 0 | 1 | -1 | -1 | 1 | 1 | 1 | 1 | 0 | 1 | 0 | 1 | 0 |
| 183_LN_Clone_E | LN | -1 | 1 | 0 | 0 | 0 | 1 | 0 | 0 | 0 | 0 | 1 | -1 | 1 | 1 | 1 | 1 | -1 | 0 | -1 | -1 | 0 |
| 183_LN_Clone_F | LN | -1 | 1 | 0 | 0 | 0 | 1 | 0 | 0 | 0 | 0 | 1 | 0 | 1 | 1 | 1 | 1 | 0 | 1 | -1 | -1 | -1 |

| | | | | | | | | | | | | | | | | | | | | | | |
|---|---|---|---|---|---|---|---|---|---|---|---|---|---|---|---|---|---|---|---|---|---|---|
| 183_primary_Clone_A | Primary | 0 | 0 | 0 | 0 | 0 | 0 | 0 | 0 | 0 | 0 | 0 | 0 | 1 | 1 | 0 | 0 | -1 | 0 | 0 | 1 | -1 |
| 183_primary_Clone_B | Primary | -1 | 1 | 0 | 0 | 0 | 1 | 0 | 0 | 0 | 0 | 1 | 0 | 1 | 1 | 0 | 1 | 0 | 0 | 0 | -1 | 0 |
| 183_primary_Clone_C | Primary | -1 | 1 | 0 | 1 | 0 | 1 | 0 | 0 | 0 | 0 | 1 | 0 | 1 | 1 | 0 | 0 | 0 | 0 | 0 | -1 | 0 |
| 239_LN_Clone_B | LN | 1 | 1 | 1 | 1 | -1 | -1 | 1 | 0 | 1 | 0 | 1 | -1 | 1 | 1 | 1 | 1 | -1 | -1 | -1 | -1 | -1 |
| 239_LN_Clone_C | LN | 0 | 1 | 0 | 0 | -1 | 0 | 0 | 0 | 0 | 0 | 1 | -1 | 0 | -1 | 1 | 1 | -1 | -1 | -1 | -1 | 0 |
| 239_LN_Clone_E | LN | 0 | 0 | 0 | 0 | 0 | 0 | 0 | 1 | 0 | 0 | 1 | 0 | 1 | 1 | 1 | 1 | 0 | 0 | 0 | 0 | 0 |
| 239_primary_Clone_A | Primary | 1 | 1 | 0 | 1 | -1 | -1 | 0 | 1 | 1 | 0 | -1 | -1 | 1 | 1 | 1 | 1 | -1 | -1 | -1 | -1 | -1 |
| 333_LN_Clone_B | LN | 0 | 1 | 0 | 1 | 0 | 1 | 0 | 0 | 0 | 0 | 0 | 0 | 1 | 1 | 0 | 1 | 0 | -1 | 0 | 0 | 0 |
| 333_LN_Clone_C | LN | 0 | 1 | 0 | 0 | 0 | 0 | 0 | 1 | -1 | -1 | 0 | 1 | 1 | 1 | 0 | 1 | 0 | -1 | 0 | 1 | 0 |
| 333_primary_Clone_A | Primary | -1 | 1 | 0 | 1 | -1 | 0 | 0 | 0 | -1 | -1 | 0 | 0 | 1 | 1 | 0 | 1 | -1 | -1 | 0 | 1 | -1 |
| 515_LN_A2_Clone_B | LN | 1 | 1 | 1 | 0 | 0 | 0 | 0 | 0 | 0 | 0 | -1 | 0 | 0 | 0 | 0 | 0 | -1 | 0 | 0 | 0 | 0 |
| 515_LN_A2_Clone_C | LN | 1 | 1 | 1 | 0 | 0 | 0 | 0 | 0 | -1 | -1 | -1 | 0 | 0 | -1 | 0 | 0 | -1 | -1 | 0 | 0 | 0 |
| 515_LN_A3_Clone_D | LN | 1 | 1 | 1 | 0 | 0 | 0 | 0 | 0 | -1 | -1 | -1 | 0 | 0 | -1 | 0 | 0 | -1 | -1 | 0 | 0 | 0 |
| 515_LN_A3_Clone_E | LN | 1 | 1 | 0 | 0 | 0 | 0 | 0 | 0 | -1 | -1 | 0 | 0 | 0 | 0 | 0 | 0 | -1 | -1 | 0 | 0 | 0 |
| 515_LN_A3_Clone_F | LN | 1 | 1 | 0 | 0 | 0 | 0 | 0 | 0 | -1 | -1 | -1 | 0 | 0 | -1 | 0 | 0 | -1 | -1 | 0 | 0 | 0 |
| 515_primary_Clone_A | Primary | 1 | 1 | 0 | 1 | 0 | 0 | 1 | 0 | -1 | 1 | 0 | 0 | 1 | 0 | 0 | 0 | -1 | -1 | 0 | 0 | 0 |
| 686_LN_Clone_E | LN | 0 | 1 | -1 | -1 | 0 | 1 | 0 | 0 | 0 | 1 | 1 | 1 | 1 | 1 | 1 | 1 | -1 | -1 | -1 | -1 | -1 |
| 686_LN_Clone_F | LN | 0 | 0 | 0 | -1 | 0 | 0 | 1 | 1 | 0 | 1 | 1 | 1 | 1 | 1 | 1 | 1 | 0 | 0 | -1 | -1 | -1 |
| 686_primary_Clone_A | Primary | 0 | -1 | 0 | -1 | 1 | 0 | 1 | 1 | 0 | -1 | 1 | -1 | 1 | 1 | 1 | 1 | -1 | -1 | -1 | -1 | 1 |
| 686_primary_Clone_B | Primary | -1 | -1 | 0 | 0 | 0 | 0 | 1 | 1 | 0 | 0 | 1 | -1 | 1 | 1 | 1 | 1 | -1 | -1 | -1 | -1 | 0 |
| 686_primary_Clone_C | Primary | 0 | -1 | 0 | -1 | 0 | 0 | 0 | 0 | 0 | 0 | 1 | -1 | 1 | 1 | 1 | 1 | -1 | -1 | -1 | -1 | 0 |
| 686_primary_Clone_D | Primary | 0 | 0 | 0 | 0 | 0 | 0 | 0 | 0 | 0 | 1 | 1 | 0 | 1 | 1 | 1 | 1 | 0 | -1 | -1 | -1 | -1 |

| sample_clone | Tissue | 11q | 12p | 12q | 13q | 14q | 15q | 16p | 16q | 17p | 17q | 18p | 18q | 19p | 19q | 20p | 20q | 21p | 21q | 22q |
|---|---|---|---|---|---|---|---|---|---|---|---|---|---|---|---|---|---|---|---|---|
| 026_A1_LN_Clone_F | LN | 0 | 0 | 1 | 0 | -1 | 0 | 1 | -1 | 1 | 1 | 0 | 1 | 0 | 0 | 0 | 0 | 0 | 0 | 0 |
| 026_A1_LN_Clone_G | LN | 0 | -1 | 1 | 0 | 0 | 1 | 1 | -1 | 1 | 1 | 1 | 1 | 0 | 0 | 0 | 0 | 0 | 0 | 0 |
| 026_A1_LN_Clone_H | LN | 0 | 0 | 1 | 0 | 0 | 0 | 1 | -1 | 1 | 1 | 0 | 0 | 0 | 0 | 0 | 0 | 0 | 0 | 0 |
| 026_A1_LN_Clone_I | LN | 0 | 0 | 1 | 0 | 0 | 0 | 1 | -1 | 1 | 1 | 0 | 0 | 0 | 0 | 0 | 0 | 0 | 0 | 0 |
| 026_A1_LN_Clone_J | LN | 0 | 0 | 0 | 0 | 0 | 0 | 1 | -1 | 1 | 1 | 0 | 0 | 0 | 0 | 0 | 0 | 0 | 0 | 0 |
| 026_A1_LN_Clone_K | LN | 0 | 0 | 1 | 0 | -1 | 1 | 1 | -1 | 1 | 1 | 0 | 1 | 0 | 0 | 0 | 0 | 0 | 0 | 0 |
| 026_A1_LN_Clone_L | LN | 0 | 0 | 1 | 0 | -1 | 0 | 1 | -1 | 1 | 1 | 0 | 0 | 0 | 0 | 0 | 0 | 0 | 0 | 0 |
| 026_B1_LN_Clone_M | LN | 0 | -1 | 1 | 0 | -1 | 0 | 1 | -1 | 1 | 1 | 0 | 0 | 0 | 0 | 0 | 0 | 0 | 0 | 0 |
| 026_B1_LN_Clone_N | LN | -1 | -1 | 1 | 0 | -1 | 0 | 1 | -1 | 1 | 1 | 0 | 0 | 0 | 0 | 0 | 0 | 0 | 0 | 0 |
| 026_B1_LN_Clone_O | LN | 0 | 0 | 0 | 0 | 0 | 0 | 1 | -1 | 1 | 1 | 0 | 0 | 0 | 0 | 0 | 0 | 0 | 0 | 0 |
| 026_B1_LN_Clone_P | LN | 0 | -1 | 1 | 0 | -1 | 1 | 1 | -1 | 1 | 1 | 1 | 0 | 0 | 1 | 0 | 0 | 0 | 0 | 0 |
| 026_primary_Clone_B | Primary | 0 | 0 | 0 | 0 | 0 | 0 | 1 | -1 | 1 | 0 | 0 | 0 | 0 | 0 | 0 | 0 | 0 | 0 | 0 |
| 026_primary_Clone_C | Primary | 0 | 0 | 0 | 0 | 0 | 0 | 1 | -1 | 1 | 0 | 0 | 0 | 0 | 0 | 0 | 0 | 0 | 0 | 0 |
| 026_primary_Clone_D | Primary | 0 | 0 | 0 | 0 | 0 | 0 | 1 | 0 | 1 | 0 | 0 | 0 | 0 | 0 | -1 | 0 | 0 | 0 | 0 |
| 070_LN_Clone_A | LN | -1 | -1 | -1 | 0 | -1 | 1 | 1 | -1 | 0 | 1 | 1 | 0 | 1 | -1 | 0 | 0 | -1 | -1 | 0 |
| 070_LN_Clone_B | LN | -1 | 0 | 0 | 0 | -1 | 1 | 1 | 0 | 0 | 1 | 0 | 0 | 1 | -1 | 0 | 0 | 0 | 0 | 1 |
| 070_LN_Clone_C | LN | 0 | 0 | 0 | 0 | -1 | 1 | 1 | 0 | 0 | 1 | 0 | 0 | 0 | 0 | 0 | 0 | 0 | 0 | 0 |
| 070_primary_Clone_A | Primary | -1 | -1 | 1 | 0 | -1 | 1 | 0 | 0 | 0 | 1 | 1 | 1 | 1 | -1 | 0 | 0 | 0 | -1 | 1 |
| 142_LN_Clone_F | LN | -1 | -1 | -1 | 1 | -1 | 0 | 1 | 1 | 1 | 1 | 0 | 0 | 0 | 0 | 0 | 0 | 0 | 0 | 0 |
| 142_LN_Clone_G | LN | -1 | -1 | 0 | 0 | -1 | 0 | 1 | 1 | 0 | 1 | 1 | 1 | 0 | 0 | 0 | 0 | 0 | 0 | 0 |
| 142_LN_Clone_H | LN | -1 | -1 | -1 | 0 | -1 | 0 | 1 | 0 | 1 | 1 | 0 | 0 | 0 | 0 | 0 | 0 | 0 | 0 | 1 |
| 142_LN_Clone_I | LN | -1 | -1 | -1 | 1 | -1 | 0 | 1 | 1 | 1 | 1 | 1 | 1 | -1 | 0 | 0 | 0 | 0 | 0 | 0 |
| 142_LN_Clone_J | LN | -1 | -1 | -1 | 1 | -1 | 0 | 1 | 1 | -1 | 1 | 1 | 1 | 0 | 0 | 1 | 0 | 0 | 0 | 0 |
| 142_primary_Clone_A | Primary | -1 | -1 | 1 | 0 | 0 | 0 | 0 | 0 | 1 | 1 | 1 | 0 | 0 | 0 | 0 | 1 | 0 | 1 | -1 |
| 142_primary_Clone_B | Primary | -1 | -1 | 1 | 0 | -1 | -1 | 0 | 0 | 1 | 1 | 0 | -1 | 0 | 1 | 1 | 1 | 0 | 1 | -1 |
| 142_primary_Clone_C | Primary | -1 | -1 | 1 | 1 | 0 | 0 | 1 | 1 | 1 | 1 | 1 | 1 | 0 | 0 | 1 | 1 | 0 | 1 | -1 |
| 142_primary_Clone_D | Primary | -1 | -1 | 1 | 1 | 0 | 0 | 1 | 1 | 1 | 1 | 1 | 0 | 0 | -1 | 1 | 1 | 0 | 1 | -1 |
| 183_LN_Clone_E | LN | 1 | -1 | 0 | 1 | -1 | 1 | -1 | 0 | 1 | 1 | 1 | 1 | 0 | 0 | 1 | 0 | 0 | 1 | 1 |
| 183_LN_Clone_F | LN | 1 | -1 | 0 | 1 | -1 | 0 | -1 | 0 | 1 | 1 | 1 | 1 | 0 | 0 | 1 | 1 | 0 | 1 | 1 |

| | | | | | | | | | | | | | | | | | | | | |
|---|---|---|---|---|---|---|---|---|---|---|---|---|---|---|---|---|---|---|---|---|
| 183_primary_Clone_A | Primary | 1 | 0 | 0 | 1 | 1 | 0 | 0 | 0 | 1 | 0 | 0 | 0 | 0 | 0 | 0 | 1 | 0 | 1 | 1 |
| 183_primary_Clone_B | Primary | 1 | 0 | 0 | 1 | -1 | 0 | 0 | 0 | 1 | 1 | 1 | 1 | 0 | 0 | 1 | 0 | 0 | 1 | 1 |
| 183_primary_Clone_C | Primary | 1 | 0 | 0 | 1 | -1 | 0 | 0 | 0 | 1 | 1 | 0 | 0 | 1 | 1 | 1 | 0 | 0 | 0 | 0 |
| 239_LN_Clone_B | LN | -1 | 1 | 1 | 1 | -1 | 1 | 1 | 1 | 1 | 1 | 0 | 1 | 0 | -1 | 1 | 1 | 0 | 0 | -1 |
| 239_LN_Clone_C | LN | 0 | 0 | 1 | 0 | 0 | 1 | 0 | 0 | -1 | 1 | 1 | 1 | 0 | -1 | 1 | 0 | 0 | 1 | 0 |
| 239_LN_Clone_E | LN | 0 | 0 | 0 | 0 | 0 | 1 | 0 | 0 | 1 | 1 | 0 | 0 | 0 | 0 | 0 | 0 | 0 | 0 | 0 |
| 239_primary_Clone_A | Primary | -1 | 1 | 1 | 1 | -1 | 1 | 1 | 0 | 1 | 1 | 0 | 1 | 1 | 0 | 0 | 0 | 0 | 0 | 1 |
| 333_LN_Clone_B | LN | 0 | 0 | 0 | 0 | 0 | 0 | 0 | 0 | 1 | 0 | 0 | 0 | 0 | 0 | 0 | 0 | 0 | 1 | 0 |
| 333_LN_Clone_C | LN | 0 | 0 | 0 | 0 | 1 | 0 | -1 | 0 | 1 | 1 | 0 | 0 | 0 | 0 | 0 | 0 | 0 | 1 | 0 |
| 333_primary_Clone_A | Primary | 0 | 0 | -1 | 0 | 0 | 1 | -1 | 1 | 1 | 0 | 0 | 0 | 0 | -1 | 0 | 0 | 0 | 0 | -1 |
| 515_LN_A2_Clone_B | LN | 0 | 0 | 0 | 0 | -1 | 0 | 0 | 0 | 0 | 0 | 1 | 1 | 0 | -1 | 0 | 0 | 0 | 1 | -1 |
| 515_LN_A2_Clone_C | LN | 0 | -1 | 0 | 0 | -1 | 1 | 0 | 0 | 0 | 1 | 1 | 1 | -1 | -1 | 0 | 0 | 0 | 0 | -1 |
| 515_LN_A3_Clone_D | LN | 0 | 0 | 0 | 0 | -1 | 1 | 0 | 0 | 0 | 0 | 1 | 1 | 0 | -1 | 0 | 0 | 0 | 0 | 0 |
| 515_LN_A3_Clone_E | LN | 0 | -1 | 0 | 0 | -1 | 1 | 0 | 0 | 0 | 1 | 1 | 0 | -1 | -1 | 0 | 0 | 0 | 0 | -1 |
| 515_LN_A3_Clone_F | LN | 0 | -1 | 0 | 0 | -1 | 1 | 0 | 0 | 0 | 1 | 1 | 1 | 0 | -1 | 0 | 0 | 0 | 0 | -1 |
| 515_primary_Clone_A | Primary | 0 | -1 | 1 | 0 | 0 | 1 | 0 | 1 | -1 | 1 | 0 | 0 | -1 | 0 | 0 | 0 | 0 | -1 | -1 |
| 686_LN_Clone_E | LN | -1 | 0 | 0 | 1 | -1 | 1 | 0 | 0 | 1 | -1 | 0 | 0 | -1 | 0 | 1 | 1 | 0 | 0 | -1 |
| 686_LN_Clone_F | LN | -1 | 0 | 0 | 1 | -1 | 1 | 0 | 0 | 1 | 1 | 0 | 0 | 0 | 0 | 1 | 1 | 0 | 1 | 0 |
| 686_primary_Clone_A | Primary | 0 | -1 | 1 | 1 | -1 | 1 | 0 | 0 | 1 | 0 | 1 | 1 | 0 | 0 | 0 | 0 | 0 | 0 | 0 |
| 686_primary_Clone_B | Primary | 0 | -1 | 1 | 1 | -1 | 1 | 1 | 0 | 1 | 1 | 0 | 0 | 0 | 0 | 0 | 0 | 0 | 0 | 0 |
| 686_primary_Clone_C | Primary | 0 | -1 | 1 | 0 | -1 | 1 | 1 | 0 | 1 | 1 | 0 | 0 | 0 | 0 | 0 | 0 | 0 | 0 | 0 |
| 686_primary_Clone_D | Primary | -1 | 0 | 1 | 1 | -1 | 1 | 0 | 0 | 1 | 0 | 0 | 0 | 0 | -1 | 0 | 0 | 0 | 0 | 0 |
