## Supplementary Table 5 for "Melanoma evolution in the lymph node shapes systemic outcomes"

**Consensus signatures identity cell states** | **Consensus signatures adaptive cell states**

| Melanocytic | Mesenchymal-like | Neural crest-like | Ferroptosis | Hypoxic stress | IFN response | Mitochondrial stress | Mitotic | p53 |
|---|---|---|---|---|---|---|---|---|
| MITF | PDGFRB | NGFR | SLC7A11 | AK4 | DPP6 | MGP | E2F2 | SON |
| MLANA | COL5A1 | AXL | FTH1 | ATF4 | PKHD1 | APOE | ESCO2 | N4BP2L2 |
| TYR | COL1A1 | SOX2 | CYBB | ENO2 | PCSK2 | MT-ATP8 | PKMYT1 | RBM39 |
| PMEL | THY1 | ANGPTL4 | ZEB1 | FLNB | IFIT1 | MT-ND6 | ASF1B | PLEKHA5 |
| GPR143 | LOXL2 | SERPINE1 | CISD1 | HK2 | OAS2 | DST | RRM2 | FUS |
| DCT | TGFBI | NTM | ACSL3 | HSPA9 | PALMD | APP | KIFC1 | HPS4 |
| SLC24A5 | COL6A3 | TFAP2B | TFRC | NDUFA4L2 | TFAP2B | PCDH7 | CLSPN | ATM |
| SLC45A2 | | GFRA2 | ACSL4 | NXPH4 | CTXND1 | NFIA | FAM111B | MDM4 |
| SOX10 | | TFAP2C | GPX4 | P4HA1 | APCDD1 | KCNJ10 | SPC25 | PPP1R15A |
| EDNRB | | WNT5A | GSS | PGK1 | IDO1 | RNASE1 | EXO1 | FUS |
| GPNMB | | SLITRK6 | PROM2 | SLC16A3 | CXCL10 | | | HSPA1A |
| IRF4 | | | | SLC2A1 | CXCL9 | | | TRPM1 |
| MYO1D | | | | VEGFA | STAT1 | | | PAX3 |
| RAB27A | | | | | IFNG | | | ARID1B |
| RUNX3 | | | | | | | | JUN |
| SLC7A8 | | | | | | | | HNRNPU |
| TFAP2A | | | | | | | | CCNL1 |
| TYRP1 | | | | | | | | |
| MLPH | | | | | | | | |
| TRPM1 | | | | | | | | |

**Other gene signatures**

| *See methods: CIN signat* | *Aybey et al. 2025 Journal of Translational Medicine* | | *GSEA* | | *GSEA* | | *GSEA* |
|---|---|---|---|---|---|---|---|
| **CIN** | **IFN type I response** | **IFN type II response** | **TGFb response** | | **TNFR1/2 response** | | **cGAS-STING** |
| CCL2 | CMPK2 | CD74 | ACVR1 | PPM1A | ARHGDIB | PRKDC | MIR4691 |
| C3 | DDX58 | CXCL9 | APC | PPP1CA | BAG4 | RB1 | TAB1 |
| CXCL1 | GMPR | GBP2 | ARID4B | PPP1R15A | CASP2 | RELA | SMPDL3A |
| CXCL5 | HERC5 | ICAM1 | BCAR3 | RAB31 | CASP3 | RIPK1 | TREX1 |
| IL1RN | HERC6 | IDO1 | BMP2 | RHOA | CASP8 | SPTAN1 | CGAS |
| TFF1 | HRASLS2 | IRF1 | BMPR1A | SERPINE1 | CHUK | TANK | LYPLAL1 |
| TFF2 | HSH2D | | BMPR2 | SKI | CRADD | TNF | PARP1 |
| TFF3 | IFI27 | | CDH1 | SKIL | DFFA | TNFAIP3 | AKT1 |
| CCL17 | IFI6 | | CDK9 | SLC20A1 | DFFB | TNFRSF1A | TBK1 |
| CD36 | IFIT1 | | CDKN1C | SMAD1 | DUSP1 | TNFRSF1B | STING1 |
| CDCP1 | IFIT3 | | CTNNB1 | SMAD3 | ELP1 | TRADD | IRGM |
| | ISG15 | | ENG | SMAD6 | FADD | TRAF1 | IRF3 |
| | LAMP3 | | FKBP1A | SMAD7 | IKBKB | TRAF2 | PCBP2 |
| | MX1 | | FNTA | SMURF1 | IKBKG | TRAF3 | ZDHHC9 |
| | MX2 | | FURIN | SMURF2 | JUN | | DDX41 |
| | OAS1 | | HDAC1 | SPTBN1 | LMNA | | ENPP3 |
| | OAS2 | | HIPK2 | TGFB1 | LMNB1 | | MARCHF5 |
| | OASL | | ID1 | TGFBR1 | LMNB2 | | PPP6C |
| | RSAD2 | | ID2 | TGIF1 | MADD | | PRKDC |
| | USP18 | | ID3 | THBS1 | MAP2K4 | | AARS2 |
| | | | IFNGR2 | TJP1 | MAP3K1 | | RELA |
| | | | JUNB | TRIM33 | MAP3K14 | | SLC19A1 |
| | | | KLF10 | UBE2D3 | MAP3K7 | | MAP3K7 |
| | | | LEFTY2 | WWTR1 | MAPK8 | | BTK |
| | | | LTBP2 | XIAP | NFKB1 | | RNF39 |
| | | | MAP3K7 | | NFKBIA | | ZDHHC18 |
| | | | NCOR2 | | PAK1 | | BANF1 |
| | | | NOG | | PAK2 | | SPSB3 |
| | | | PMEPA1 | | PARP1 | | AURKB |

**25 NMF programs with their 150 ranked genes**

| | NMF1 | NMF2 | NMF3 | NMF4 | NMF5 | NMF6 | NMF7 | NMF8 | NMF9 | NMF10 |
|---|---|---|---|---|---|---|---|---|---|---|
| **V1** | PMEL | DCT | NRAS | CCND1 | FXYD3 | TYRP1 | DCT | NRAS | IGKC | STX3 |
| **V2** | PAEP | PMEL | MDM2 | BMPR2 | THBS2 | PMEL | CCL21 | APCDD1 | LYZ | VPS41 |
| **V3** | QPCT | ADGRG1 | CDK4 | UQCRC1 | SLC20A1 | MCOLN3 | IGKC | CDK4 | IGHG1 | NDUFA3 |
| **V4** | EPHX1 | CAPG | CPM | B4GALT5 | S100A16 | APCDD1 | RNASE1 | MDM2 | CTSS | SDC3 |
| **V5** | GLMP | IFI6 | LMBRD2 | METTL9 | FLOT1 | QPCT | CLU | CTDSP2 | LCP1 | COL1A2 |
| **V6** | CTSK | IFI27 | ABHD5 | FLOT1 | SERPINA3 | TIMP2 | BCAN | IGFBP3 | LSP1 | SPG7 |
| **V7** | MICAL1 | EMP1 | NADK2 | EEF1G | CTSK | PRKD3 | LYZ | NADK2 | CD4 | WWC3 |
| **V8** | ITPKB | S100A4 | CTDSP2 | HNRNPF | TSC22D1 | STXBP1 | IGHM | SLC1A5 | LAPTM5 | CHST11 |
| **V9** | TTYH3 | ECM1 | GDF15 | TUBB2B | SLC5A3 | UGCG | IGHG1 | LMBRD2 | SPI1 | CTSK |
| **V10** | ENO2 | COL1A2 | MITF | GSN | CCND1 | PLPPR4 | PTPRZ1 | MARCH9 | FCER1G | KIFC3 |
| **V11** | S100A16 | TIMP3 | CAPG | PFN2 | PLA1A | PLD3 | MS4A1 | TSFM | ITGB2 | SNX10 |
| **V12** | IFI6 | L1CAM | TSPAN31 | MYC | LOXL4 | IFI16 | PTGDS | VGF | ARHGDIB | RARA |
| **V13** | SLC39A6 | VAMP8 | METTL1 | TGIF1 | CDKN1A | EMILIN2 | APOC1 | S100A4 | LIPA | NBR1 |
| **V14** | SGK1 | CTSK | ANO10 | GLI3 | SERPINF1 | CSTB | VCAM1 | CDH3 | FUCA1 | CAVIN1 |
| **V15** | ADGRG1 | COL3A1 | TSFM | ENO1 | TNC | RNASE1 | CORO1A | IGFBP7 | TRAC | TDRD3 |
| **V16** | RAB32 | MAL | IGKC | GLMP | FRMD4A | SPTAN1 | LCP1 | CEBPD | MPEG1 | NUDCD3 |
| **V17** | ECM1 | SPP1 | CTSK | PLXNA1 | ABCB5 | LYZ | ARRDC3 | METTL1 | CAPG | IFITM2 |
| **V18** | VGF | ITGA3 | EMP1 | VTI1A | ITGA10 | NBL1 | PLP1 | SLC1A3 | CORO1A | SLC2A4RG |
| **V19** | OAF | COL1A1 | SLC1A3 | COL4A1 | IGFBP7 | IFI30 | ADGRG1 | TSPAN31 | IGHG3 | PAX3 |
| **V20** | NUP93 | ITPR3 | VGF | MITF | BHLHE40 | SLC7A11 | FOS | ABHD2 | ITGAX | TMEM165 |
| **V21** | WIPI1 | MFSD12 | SLC20A1 | HK2 | SEMA3B | SGK1 | ARHGDIB | SLC24A4 | FGL2 | PINK1 |
| **V22** | SLC1A4 | FN1 | MARCH9 | SLC25A4 | PERP | GDF15 | CCL19 | EEF1AKMT3 | C3 | NSMF |
| **V23** | L1CAM | CSTB | ABCB5 | VEZF1 | BIRC7 | MYO1D | ELOVL2 | UGCG | CXCR4 | AMFR |
| **V24** | TYRP1 | QDPR | SKP2 | DCXR | CNIH3 | CDKN1A | LTB | COL6A1 | IFI30 | PPARD |
| **V25** | NBL1 | METRN | ZFYVE16 | SDC3 | CD109 | VAMP8 | SELL | SKP2 | ADA2 | JUNB |
| **V26** | ANKRD9 | FXYD3 | EEF1AKMT3 | BAP1 | ECM1 | CD55 | C3 | L1CAM | CTSH | ZNF697 |
| **V27** | ASAH1 | BCAN | ATP23 | EIF4G2 | SNX10 | NQO1 | TSPAN7 | ITGA1 | TRBC2 | TRAPPC6A |
| **V28** | NQO1 | ENO1 | MMP2 | SLC31A1 | SGK1 | CHIT1 | S100A4 | ATP23 | IGHA1 | SLC1A5 |

| | | | | | | | | | | |
|---|---|---|---|---|---|---|---|---|---|---|
| **V29** | TSPAN10 | ITM2C | SERPINF1 | PDZRN3 | BRSK1 | CHL1 | HIST1H1B | IER3 | IKZF1 | SAMD4B |
| **V30** | TRPM1 | CD109 | BHLHE41 | SHOC2 | PCDH7 | PTMS | TRBC2 | CLEC11A | SOD2 | NAMPT |
| **V31** | KCNAB2 | BAMBI | PRLR | COL3A1 | PLAT | ATP6V0B | ENPP2 | ENO2 | CIITA | SYNE3 |
| **V32** | MYO1D | SGK1 | TDRD3 | CREM | SESN3 | MICAL2 | RIPOR2 | MFSD12 | OAS2 | NAPG |
| **V33** | SS18L1 | ATP6V1F | ATP6AP1 | VEGFA | DUSP5 | IFI27 | LGI4 | CXCL8 | CD37 | IFI6 |
| **V34** | CTSH | MYO1D | TSPAN10 | COL6A2 | PCSK6 | ST5 | JCHAIN | BHLHE40 | HMOX1 | INTS1 |
| **V35** | SLC7A5 | MMP9 | TRIM63 | SNX9 | PTPRZ1 | SHC1 | POSTN | SMIM3 | PECAM1 | NPL |
| **V36** | BAMBI | TFRC | SCIN | SNX25 | TFPI2 | TFRC | CXCR4 | TYRP1 | EPSTI1 | SLC39A13 |
| **V37** | IGSF8 | CLEC11A | MME | TSFM | ABHD2 | ITPKB | C2 | STXBP1 | CD52 | PRMT2 |
| **V38** | PIK3CD | ITGB3 | MYO1D | PRKD3 | GJB1 | IGFBP7 | SESN3 | LSAMP | NCF2 | SMAP2 |
| **V39** | COX7A2 | ST3GAL5 | CDKN2A | SLC25A37 | RCAN1 | SLC1A5 | ST6GAL1 | PRKD3 | IL32 | RCAN1 |
| **V40** | EMILIN2 | COL18A1 | MDK | ITPR3 | INPP5F | PDK4 | IKZF1 | COL6A2 | IGHM | ZNRF1 |
| **V41** | FRZB | PTMS | L1CAM | BHLHE40 | STRIP2 | QDPR | HIST1H1D | CCND1 | IGLC1 | ADGRG1 |
| **V42** | SYTL2 | TIMP2 | BHLHE40 | RAB32 | ITGA3 | SRGN | METRN | ITGA3 | IFI6 | COL6A1 |
| **V43** | PIR | CTSH | IGFBP7 | WDR34 | MGP | CHST11 | IRF8 | THBS1 | CCL5 | HIPK1 |
| **V44** | GPR161 | TFAP2A | WIPI1 | AZIN1 | ENO2 | TSNAX | DKK3 | PLPPR4 | PLEK | PDP1 |
| **V45** | KIRREL1 | ENO2 | HEY1 | HILPDA | PRRX1 | CTSH | MET | SPP1 | ISG15 | PDLIM2 |
| **V46** | NAV2 | ATOX1 | SEPTIN3 | H1FX | SCN1B | TMEM59 | AIF1L | PRRX1 | IRF8 | HGS |
| **V47** | CA14 | PFKP | IGHG1 | FGFR1 | MMP9 | ANXA3 | TRAC | B4GALT5 | CPVL | BCAR1 |
| **V48** | IFI27 | HIST1H1B | IER3 | UCK1 | CDH1 | EPHA2 | PLA1A | RACK1 | PTGDS | PSEN2 |
| **V49** | PLXNC1 | PHLDA2 | FN1 | IL16 | LIF | TMEM163 | ALDH1A1 | PLAUR | CXCL12 | CARD19 |
| **V50** | ANXA2 | ENTHD1 | ADAMTS2 | HK1 | DUSP10 | IFI6 | LSP1 | HEY1 | JCHAIN | PLP2 |
| **V51** | MAPK4 | PLP1 | CD55 | LCP1 | SPP1 | IQSEC1 | IFI6 | SNTB1 | CCL19 | SRPRA |
| **V52** | CADM1 | ANXA2 | MCF2L | CTXN1 | SDC2 | EIF4H | CD37 | SOCS3 | NFKBIA | HK1 |
| **V53** | NCSTN | CNIH3 | LY96 | TTYH3 | CSTB | TMEM14C | AKAP12 | CAPG | TRBC1 | IKBKB |
| **V54** | SYT11 | AIF1L | RACK1 | PDLIM7 | CHST11 | ASAH1 | TBX2 | TNFRSF12A | OAS1 | ITGB2 |
| **V55** | HSPB8 | HPCAL1 | DUSP10 | IER5L | TM4SF1 | ADGRL4 | IL7R | ADGRG1 | FERMT3 | HIST2H2BE |
| **V56** | HTRA3 | PLXNC1 | FAM20C | ATP5MC2 | FCRLA | TM9SF2 | CORO2B | SLC16A6 | FBP1 | ETV4 |
| **V57** | CAPG | TNC | CNIH3 | COL6A3 | RXRG | ADGRG1 | FADS2 | KLF6 | SERPINA1 | NKIRAS2 |
| **V58** | ITGB5 | GSN | SFRP1 | SRPRA | APOC1 | CAPG | PLEKHB1 | SLC7A11 | S100A9 | VPS4B |

| | | | | | | | | | | |
|---|---|---|---|---|---|---|---|---|---|---|
| **V59** | MCF2L | TDRD3 | UGCG | ZCCHC14 | FADS2 | RAB6B | STK17B | WIPI1 | RAC2 | DUSP1 |
| **V60** | ZNF697 | WIPI1 | CLEC11A | ATP5PD | LSAMP | JAK1 | VAMP8 | RUNX3 | MX1 | PTPN14 |
| **V61** | EGFL8 | SNX10 | LYZ | IER3 | MICALL2 | WDR63 | IL32 | TGFBI | SELL | S100A4 |
| **V62** | CREM | ARL6IP4 | CCND1 | CIZ1 | SHROOM2 | IVNS1ABP | GDF11 | ZNRF1 | STAB1 | PYCARD |
| **V63** | HES1 | KCNN4 | TGFBI | GAB2 | MXI1 | ARL2BP | SMAP2 | HPCAL1 | PLA2G7 | STARD13 |
| **V64** | TDRD3 | BHLHE41 | NBL1 | PDP1 | OAF | KIT | IFI27 | NPTX1 | CD300A | SLC12A7 |
| **V65** | COX7A1 | PLBD2 | COL3A1 | PKN3 | CADPS | KLF6 | TAGLN | FGFR1 | NPL | ABHD2 |
| **V66** | TSNAX | NRCAM | COL1A1 | MC1R | ATP6V1F | IGSF8 | SELENOP | LINGO1 | ALDH2 | TTYH3 |
| **V67** | LAMA1 | ITGB8 | SH3BP4 | WIPI1 | IGFBP5 | STAT2 | ERBB3 | MICAL2 | MZB1 | AHR |
| **V68** | SELENON | PDLIM4 | PRKCB | ABHD2 | ADCY1 | SLC31A1 | RACK1 | ATP6V0E2 | SERPINB9 | PLK3 |
| **V69** | HSBP1 | NAMPT | PMEL | CDC25B | PLAUR | PIK3CD | ARPC1A | KIT | LTB | SETBP1 |
| **V70** | IFI16 | CTXND1 | IFI6 | CDCA4 | LYPD1 | PLEKHB1 | PMEL | CAVIN1 | CXCL10 | KAZN |
| **V71** | ACADM | PRAME | MFSD12 | PRMT2 | ENPP2 | FUCA1 | KCNN4 | PFKP | SLC40A1 | KLF16 |
| **V72** | MME | OAF | PDLIM4 | PTPN14 | KCNN4 | SYT11 | ATP6V0E2 | AXIN2 | IL2RG | TTYH2 |
| **V73** | SCRN1 | IGFBP4 | EPHX1 | FZD7 | STC1 | CCND1 | CD52 | NKD1 | IL4I1 | RHOB |
| **V74** | PTGFRN | IFI16 | TTYH2 | HNRNPC | TPD52L1 | NRP2 | ADGRG6 | CCN3 | CHST11 | IGF2R |
| **V75** | TIMP2 | BIRC7 | EGR1 | ARL8A | TUSC3 | HPGD | CIITA | PAG1 | PIM2 | TSC22D1 |
| **V76** | FUCA1 | ACADM | VAC14 | TFRC | NBL1 | CCL18 | CSF2RA | IER5L | CSF2RA | METRN |
| **V77** | SLC25A16 | MITF | S100A16 | SLC19A1 | ITGB8 | HYOU1 | DNASE1L3 | TMEM204 | CCL2 | ITGAX |
| **V78** | ATOX1 | ERBB3 | COL6A2 | FAM89A | PHLDA2 | ATP6AP2 | MPEG1 | IGFBP2 | TNFAIP3 | COX7A2 |
| **V79** | MITF | PTPRZ1 | CRYL1 | STXBP1 | ANXA2 | SPAG7 | PCDH7 | TWIST1 | TGFBI | KANSL1 |
| **V80** | HTRA1 | SLC20A1 | ADCY1 | VPS51 | PFKFB4 | ENO2 | SERPINB9 | DCXR | TRAF3IP3 | SLC19A1 |
| **V81** | CENPF | ATP5MC2 | IGHA1 | IKBKB | ADAMTS1 | HNRNPU | COL1A2 | NQO1 | TFRC | SBF1 |
| **V82** | TRIM63 | MMP1 | COL4A2 | FXYD6 | DCT | KCNAB2 | HIST1H1E | MCOLN3 | IL18BP | HIST1H1B |
| **V83** | SH3PXD2B | C2 | PAEP | MYL12A | CADM1 | GSN | SLC40A1 | NBL1 | CXCL9 | ZCCHC14 |
| **V84** | SLC25A4 | SELENON | BIRC7 | SLC39A8 | ERBB3 | IGKC | IL2RG | MC1R | ANXA6 | NRIP1 |
| **V85** | TMEM106C | LSAMP | FUCA1 | GRM3 | TRPM8 | ADGRG6 | IGFBP2 | MME | TREM2 | PHACTR4 |
| **V86** | FH | OLFML3 | RTP4 | KIF1B | SORBS2 | PLA2G7 | LRATD2 | ETV1 | SMAP2 | TGDS |
| **V87** | ANXA6 | TM7SF3 | TNFRSF12A | TLK1 | LRATD2 | CDC42EP3 | CD79A | MMP11 | SELENOP | STARD3NL |
| **V88** | UQCRC1 | IER3 | IER5L | OR8K5 | RND3 | SS18L1 | PRAME | SEPTIN4 | CHI3L1 | PTGFRN |

| | | | | | | | | | | |
|---|---|---|---|---|---|---|---|---|---|---|
| **V89** | RGS5 | MCOLN3 | MX1 | CEBPB | EMP1 | SDC3 | STAT1 | TMEM158 | TBC1D10C | HPS4 |
| **V90** | NUF2 | PTTG1 | CDKN2B | CTSH | ITM2C | IFNAR1 | JAML | SERPINE1 | VCAM1 | KCNN4 |
| **V91** | GSTO1 | LINGO1 | THBS2 | SFXN3 | LDHB | CDH3 | HNRNPH3 | SLC2A1 | LY96 | IER3 |
| **V92** | PMVK | KLF6 | CDH3 | QPCT | NPDC1 | SLC39A6 | PLAAT4 | QPCT | STK17B | BCL2L11 |
| **V93** | HIST1H1B | DUSP10 | SULT1C2 | NR4A1 | BHLHE41 | HSPH1 | TBC1D10C | FAM20C | SGK1 | ICMT |
| **V94** | LAMB2 | ARPC1A | KCNAB2 | HGS | GPR158 | PSMB4 | PLXNC1 | NDRG1 | VAMP8 | KLF6 |
| **V95** | PRKD3 | PLAUR | ITGA3 | EGR1 | ASAH1 | SEMA5A | TRBC1 | SH3BP4 | EMILIN2 | MICALL2 |
| **V96** | BHLHE40 | SEMA3B | COL1A2 | UPK1B | CD55 | HTT | SOX2 | ITPKB | BCL3 | NOP53 |
| **V97** | STK17B | METTL9 | NQO1 | PSEN2 | TNFRSF12A | ARL6IP4 | CTSH | H3F3B | DENND2D | OSBPL3 |
| **V98** | CD55 | SRPK2 | EIF1AY | PALM2-AKAP2 | DHCR7 | RGMB | C7 | SOD2 | ARHGAP15 | CBLN1 |
| **V99** | NOP2 | PLP2 | CTSH | IQSEC1 | COL1A1 | NCL | PKNOX2 | TFRC | CMPK2 | FOS |
| **V100** | KDM1A | COL4A2 | C21orf91 | NRAS | UCN2 | SNX17 | MPZ | FOS | JUNB | LSP1 |
| **V101** | CSTB | IRF4 | MSANTD3 | LSP1 | NT5E | IFRD1 | F5 | NPDC1 | TNC | ZNF804B |
| **V102** | FHOD3 | ATP6V0E2 | RAB32 | LIX1 | HSPA5 | NT5DC3 | JUNB | PFN2 | SPOCK2 | PKN3 |
| **V103** | SLC7A1 | EHD2 | CAVIN1 | USP33 | DCSTAMP | SERPINF1 | ITPR2 | GOLGA7B | ADAM28 | FAM20C |
| **V104** | BIRC7 | ITGB5 | SCRN1 | NFKBIA | RHOB | LAMB2 | CRYAB | PDE4D | STAT1 | CDKN1C |
| **V105** | STX3 | PYCARD | CDK1 | PLP2 | SHOC2 | IAH1 | CCDC50 | TFAP2A | CD48 | PLEKHM2 |
| **V106** | NPL | EIF4H | FXYD6 | STAU2 | CDH19 | LSR | AZGP1 | THBS2 | ADAMDEC1 | ITPR3 |
| **V107** | TIMMDC1 | PLEC | CTNNB1 | VCL | ST3GAL5 | ARL6IP1 | TRAF3IP3 | METRNL | TRAF1 | AHCY |
| **V108** | OCA2 | ST5 | KDELR2 | USP53 | PLP1 | GLMP | CLEC11A | SGK1 | FGR | MC1R |
| **V109** | CDK1 | SLC7A8 | FAM89A | HNRNPH3 | GDF11 | PSAT1 | PPIA | CTNNA2 | KLHL6 | PGD |
| **V110** | VPS72 | SRPRA | BGN | CDKN1C | NME1 | KIZ | ITM2C | CTNNB1 | WDFY4 | ARRDC4 |
| **V111** | TLK1 | GDF15 | ADGRG1 | ZFYVE16 | COL3A1 | KHDRBS1 | MCOLN3 | PALM2-AKAP2 | CSTB | COL4A2 |
| **V112** | SEPTIN4 | SLC39A6 | ENTHD1 | FABP5 | NT5DC3 | ATP6V1H | KCNJ10 | IQSEC1 | UGCG | KIRREL1 |
| **V113** | ATP11A | GPC1 | COL6A1 | IMMP2L | MYO1D | C21orf91 | ARHGAP15 | CD109 | IFI27 | TSPAN31 |
| **V114** | VAMP8 | WFDC1 | MMD | HNRNPA0 | NPL | ZNF697 | LOXL4 | C1QTNF1 | ITGAL | FOXD3 |
| **V115** | TNFAIP3 | PKNOX2 | GRK3 | PMVK | MDM2 | EIF4G1 | SNX10 | TLE1 | GZMB | ENC1 |
| **V116** | IGSF3 | CXCL1 | ZNF365 | NCOA3 | FGFBP2 | ITGB5 | SGK1 | PTPRG | IFIT2 | CIZ1 |
| **V117** | ATP6AP2 | SCIN | SGK1 | PDE4D | MCOLN3 | SLC25A16 | MPZL2 | NNMT | POU2AF1 | USP53 |
| **V118** | GSDME | RACK1 | S100A4 | HABP4 | UGCG | TNFRSF21 | MSC | RHOB | CEBPD | IPO5 |

| | | | | | | | | | | |
|---|---|---|---|---|---|---|---|---|---|---|
| **V119** | RAB1A | RETSAT | NCS1 | SLC35A4 | TRAF1 | SLC1A4 | EGFL8 | CDC42EP3 | CSF2RB | CCL26 |
| **V120** | SPG7 | ARRDC4 | PIGK | AHR | B4GALT5 | SLC39A8 | BHLHE41 | PLP2 | PLAUR | INO80E |
| **V121** | ARL6IP1 | MMP17 | APOC1 | NCF2 | SYTL2 | TLK1 | ADAM19 | AHR | IFIT1 | MEGF6 |
| **V122** | EPHA4 | USP53 | BMP7 | NHLRC2 | AFAP1L2 | CLEC10A | CD4 | GRK6 | DERL3 | HAS1 |
| **V123** | FRMD4A | TNFRSF12A | PRAME | NAMPT | PALM2-AKAP2 | SYTL2 | LAPTM5 | MMP1 | ZAP70 | CSF2 |
| **V124** | PFN2 | LNPK | TYRP1 | RASEF | SLC7A8 | CPVL | DENND2D | EMP1 | RGCC | PPP6R2 |
| **V125** | MOXD1 | SEMA5A | CTXND1 | POLE3 | AKAP12 | CCN3 | TMC8 | PDP1 | CCL18 | SEPTIN4 |
| **V126** | ARL6IP4 | CDK4 | TMEM255B | ZMAT2 | GASK1B | CDCP1 | CD79B | CST7 | TAGLN | HIBADH |
| **V127** | SLC45A2 | PLEKHM2 | ANGPTL2 | ANXA6 | HPCAL1 | TIMM8B | FABP7 | ANXA6 | COL3A1 | NOP2 |
| **V128** | CHL1 | IGSF3 | PALM2-AKAP2 | IGF2R | RNF175 | ADCY2 | PRKCB | MYADM | COL6A2 | LALBA |
| **V129** | HSD17B1 | SPHK1 | ECM1 | MRRF | C1QTNF1 | GSTO1 | ATP1B1 | PPARD | CSTA | NAV2 |
| **V130** | NRAS | LPAR4 | SLC1A5 | INO80E | SLC35G2 | KIRREL1 | NBL1 | ANXA2 | PLAU | ECM1 |
| **V131** | SRPRA | FKBP10 | PHLDA2 | HPS4 | SNTB1 | SRPRA | IRF4 | ZNRF3 | SPN | IMMP2L |
| **V132** | ARL2BP | SLC6A17 | TSPAN7 | AFDN | FAP | SLC12A7 | MDK | SLC2A4RG | PLAAT4 | CTSS |
| **V133** | LGI3 | CITED1 | IGHG3 | SETBP1 | ATP6AP1 | ITGA7 | ADA2 | ENO1 | APOC2 | AFDN |
| **V134** | SDHB | STXBP6 | CD109 | MEGF6 | GNG11 | ATOX1 | ST3GAL5 | ITPR3 | HIST1H1D | SPPL2B |
| **V135** | QDPR | FABP5 | CAMK2N1 | KLF6 | LIPA | TMEM106C | FGL2 | NCS1 | TENT5C | URGCP |
| **V136** | ITM2C | OAS2 | LRP8 | BAG1 | ENC1 | FRMD4A | NAMPT | ZCCHC14 | COL6A3 | TMPRSS11E |
| **V137** | PIGK | SCUBE2 | CKB | SLC7A1 | COL18A1 | JTB | GJB6 | PPIA | FCGR2A | CD4 |
| **V138** | NR4A3 | PCDH7 | SELENOP | KYAT3 | SNORC | BAX | NIBAN3 | FN1 | ATP2A3 | CADM1 |
| **V139** | ADGRG6 | GAPDHS | ASAH1 | ITGAV | SLC25A37 | TMBIM1 | UBD | MAP1B | C15orf48 | ENDOV |
| **V140** | KLF6 | PRMT2 | POLD2 | AHCY | COL11A2 | INPP5F | PAG1 | NAV2 | SLC16A3 | SOCS3 |
| **V141** | ITGA7 | COX7A1 | CHI3L1 | PPARD | MOSPD2 | COX7A1 | RAC2 | ISM1 | ALDH1A1 | AK5 |
| **V142** | ANGPTL2 | HIST1H1D | COMP | FBXW5 | COL7A1 | PXK | CXCL12 | COL4A2 | PLA2G2D | ARPC1A |
| **V143** | SLC6A17 | STMN3 | NT5E | COL4A2 | RNF128 | LIPA | CHL1 | MCF2L | CHIT1 | SLC20A1 |
| **V144** | JMJD1C | NME1 | TMEM38B | ITPKB | C21orf91 | TMEM106B | RASGRP2 | IFITM2 | IQSEC1 | BCL2 |
| **V145** | PTMS | ITGA6 | METRN | E2F1 | ATP1B1 | USP53 | IFI16 | SLC16A3 | NNMT | HES1 |
| **V146** | INPP5F | FCRLA | EEF1G | ARPC5 | QDPR | TIMMDC1 | FDCSP | UQCRC1 | SLC20A1 | HSPG2 |
| **V147** | MMP17 | SLC1A4 | SDC3 | ZNRF3 | TNFRSF21 | ASPH | ARL6IP4 | RAC3 | PGD | ANO10 |
| **V148** | FABP5 | MPZ | TAGLN | VPS54 | CREM | TMEM138 | CTDSP2 | SNX9 | CPM | SLC5A3 |

| | | | | | | | | | | |
|---|---|---|---|---|---|---|---|---|---|---|
| **V149** | ATP6V1H | NCAPG2 | SULF2 | IMPA2 | COL9A3 | LAMTOR5 | EGR1 | ANTXR1 | AMPD3 | NCOA3 |
| **V150** | BHLHE41 | PPIA | TFAP2A | SLC20A1 | MDK | SRPK2 | CXCL13 | SLC25A4 | CR1 | SPI1 |

| | NMF11 | NMF12 | NMF13 | NMF14 | NMF15 | NMF16 | NMF17 | NMF18 | NMF19 | NMF20 |
|---|---|---|---|---|---|---|---|---|---|---|
| **V1** | COL3A1 | TIMP2 | SFRP1 | SERPINF1 | ITM2C | MDM2 | SFRP1 | TOMM7 | ANKRD28 | MGP |
| **V2** | COL1A1 | IFI16 | NDRG1 | QPCT | ASAH1 | METTL9 | IGKC | MICOS10 | EDNRB | EDNRB |
| **V3** | COL1A2 | ASPH | RNASE1 | PMEL | HNRNPH3 | SDC3 | SPP1 | ANXA2 | ENO1 | EMP1 |
| **V4** | COL6A3 | EIF4H | MGP | EMP1 | SLC39A6 | ASAH1 | TYRP1 | ATP6AP1 | KIF1B | MMP2 |
| **V5** | COL6A2 | PMEL | SDC2 | GAPDHS | ANXA2 | RELL1 | SDC2 | HNRNPA0 | PRAME | SPP1 |
| **V6** | LUM | NCL | AKAP12 | TRPM1 | TMBIM1 | PSAT1 | IFI16 | IARS2 | MITF | TYRP1 |
| **V7** | COL6A1 | PSMB4 | SPP1 | ITPKB | NCSTN | IGF2R | TIMP2 | MXI1 | LAMB2 | ABCB5 |
| **V8** | DCN | PLEC | RGS5 | DCT | GSN | ZNF703 | RNASE1 | BAG1 | SLC7A5 | RNASE1 |
| **V9** | COL5A1 | PLD3 | CRYAB | SLC1A4 | SOD2 | ST3GAL5 | PLP1 | ATP5MC2 | SLC16A1 | NBL1 |
| **V10** | VCAN | TMEM147 | CHL1 | IGFBP7 | S100A16 | ETV1 | S100A4 | SELENON | ZFYVE16 | GDF15 |
| **V11** | COL4A1 | ST5 | LRATD2 | RNASE1 | NFKBIA | CRYL1 | HSPA6 | ZMAT2 | SERBP1 | POSTN |
| **V12** | IGFBP4 | IVNS1ABP | IGFBP2 | MFSD12 | TSC22D1 | LNPK | ASPH | ARPC5 | H1FX | CRYL1 |
| **V13** | FN1 | SRGN | SLC26A2 | MMP2 | IFI6 | KDM1A | HMCN1 | UQCRQ | KPNA6 | ARRDC3 |
| **V14** | BGN | TM9SF2 | SOD3 | TSPAN10 | ARL8A | ETV4 | IGHG1 | SFXN3 | HNRNPF | DCT |
| **V15** | MMP2 | HYOU1 | SYNM | RACK1 | SLC20A1 | SRP72 | SELENOP | SLC39A7 | PMVK | MET |
| **V16** | COL5A2 | ATP6V0B | BAALC | TFRC | CDH19 | ARRDC3 | PLD3 | CTSS | PGD | IGFBP7 |
| **V17** | CXCL12 | STAT2 | SPARCL1 | EPHX1 | LIPA | PIR | SPARCL1 | NME1 | SDHB | COL16A1 |
| **V18** | COL4A2 | TM7SF3 | RACK1 | SLC25A4 | CTSK | STAU2 | TMEM14C | KDELR2 | FBXW5 | PRAME |
| **V19** | COL15A1 | SHC1 | SMTN | LDHB | AHR | SLC45A2 | PCDH7 | HERC4 | USP33 | AVPR1A |
| **V20** | THY1 | SOD1 | EDNRB | PIR | VAC14 | SNTB1 | TIMP3 | HELZ | TFAP2A | TBX2 |
| **V21** | FBN1 | TIMP3 | LOXL4 | SLC7A5 | H2AFJ | UBA2 | STAT1 | IGSF3 | STXBP1 | MCC |
| **V22** | TAGLN | SNX17 | TSPAN7 | ANXA6 | BZW2 | INPP5F | TNFRSF21 | ATP5PD | TRIM63 | S100A4 |
| **V23** | PDGFRB | ATP5F1A | TNFRSF21 | ASAH1 | ECE1 | IDH3B | SRGN | CEBPB | EEF1G | HEBP1 |
| **V24** | POSTN | TMEM59 | FCGR2A | SLC45A2 | TM9SF2 | PHLPP1 | CRYAB | ADAM10 | NAV2 | ERBB3 |
| **V25** | S100A4 | SPTAN1 | IGKC | EDNRB | FRMD4A | VEZF1 | AHR | EPHX1 | CDC25B | SGK1 |
| **V26** | CCN2 | ATOX1 | PYROXD2 | STK32A | STARD3NL | HNRNPC | DCT | ATP6V1F | GRK6 | STK32A |
| **V27** | MRC2 | TMEM106C | SELENOP | FAM126A | GGCT | TBX2 | NFATC2 | FAM89A | NOP53 | FN1 |
| **V28** | SFRP2 | PTMS | CD36 | WIPI1 | VPS4B | LDHB | PTMS | VCL | AURKAIP1 | MME |

| | | | | | | | | | | |
|---|---|---|---|---|---|---|---|---|---|---|
| **V29** | MYL9 | STAT1 | SLC1A4 | EEF1G | SPI1 | NBL1 | STAT2 | ATP5MC3 | HSBP1 | GASK1B |
| **V30** | FBLN1 | TLNRD1 | CPN1 | UGCG | TOMM7 | SERBP1 | TM9SF2 | IKBKB | ANXA6 | PLAT |
| **V31** | NOTCH3 | KLC1 | ABHD2 | TNC | TSNAX | NT5DC3 | AIF1L | MICAL1 | PLEKHM2 | IGFBP2 |
| **V32** | LTBP2 | JTB | LPL | PFN2 | VPS13C | PTPRG | CCDC50 | PPP6R2 | HGS | ALDH1A1 |
| **V33** | PLVAP | TMED7 | LRP2 | KCNAB2 | BAX | ISYNA1 | PRMT2 | HSPA5 | SH3PXD2B | PTPRZ1 |
| **V34** | NNMT | SRSF4 | CSGALNACT1 | BIRC7 | SNX10 | NUP93 | IFI30 | POLD2 | FAM126A | COL18A1 |
| **V35** | COL12A1 | KPNB1 | S100A4 | SYTL2 | LAPTM5 | PTGFRN | IGF1R | ITGAV | ARL6IP4 | SEMA6A |
| **V36** | AQP1 | SRP54 | UGCG | TRIM63 | ATP6AP1 | ARL8A | TSPAN7 | PLXNA1 | SAMD4B | RACK1 |
| **V37** | HTRA1 | NRP2 | FMOD | PRKD3 | SLC7A1 | AZIN1 | PLAAT4 | KRTCAP2 | CIZ1 | COL1A1 |
| **V38** | IGFBP7 | POMP | FGFBP2 | PLP1 | PSME4 | PTPN14 | NCL | VWA1 | SLC2A1 | METRN |
| **V39** | PECAM1 | IGSF8 | ETV1 | ANXA2 | NCS1 | AHR | KLHDC3 | PIGK | ACADM | PLP1 |
| **V40** | PCOLCE | TMEM14C | BHLHE40 | CRYL1 | POLE3 | PCDH7 | LYZ | HNRNPR | HABP4 | CLEC11A |
| **V41** | MXRA8 | SPAG7 | F13A1 | GLMP | HNRNPC | NCSTN | TFAP2A | PCBP1 | PDE4D | ARHGEF5 |
| **V42** | COL18A1 | CCDC50 | PLP1 | PRUNE2 | NRIP1 | CPNE2 | AKAP12 | JUNB | ZNF689 | SUGCT |
| **V43** | ISLR | HSD17B11 | ALDH1A3 | LYZ | OAS1 | LIPA | EIF4H | NCOA3 | NCKIPSD | BHLHE41 |
| **V44** | ADAMTS2 | TIMM8B | ATP1B1 | NME1 | MYL12A | C2orf88 | EDNRB | RICTOR | NHLRC2 | TGDS |
| **V45** | ANGPTL2 | CSDE1 | LY96 | OCA2 | PIGT | SLC16A1 | ADGRG1 | SDHB | VEGFA | PCDH7 |
| **V46** | PRRX1 | EIF4G1 | POSTN | CTNNB1 | HK1 | USP33 | CHL1 | NXT2 | PTPN14 | GRK3 |
| **V47** | COL16A1 | IFNAR1 | CORO2B | TOMM7 | STAT3 | VLDLR | LOXL4 | IDH3B | GPR153 | COL3A1 |
| **V48** | TNC | SRPK2 | OLIG1 | PAG1 | COL6A2 | SNX4 | PSMB4 | ACVR2A | SS18L1 | CRYAB |
| **V49** | SULF2 | ARNT2 | SLC25A4 | PDE4D | SLC25A16 | SPPL2B | EIF3L | HES1 | SBF1 | MCOLN3 |
| **V50** | SERPINF1 | KLF6 | SHISA2 | CCN5 | HGS | AKAP12 | BAALC | NAMPT | IDH3B | PI15 |
| **V51** | FOS | HMGN4 | AGMO | MITF | C1QTNF1 | ZNF697 | AKAP6 | IPO5 | BCAR1 | SESN3 |
| **V52** | CRISPLD2 | TIMM50 | ELOVL2 | SLC25A16 | IQSEC1 | MX1 | RACK1 | IER5L | ATP5PD | CTSK |
| **V53** | TGFBI | ATP6AP2 | STK32A | SEMA6A | PALM2-AKAP2 | ADM | GSN | NFIB | ATP5MC2 | SLC45A2 |
| **V54** | COL14A1 | HNRNPU | LGI3 | PDZRN3 | SLC5A3 | CDH19 | TMED7 | HSPG2 | CROCC | CITED1 |
| **V55** | PDGFRA | ITCH | PLEKHB1 | UQCRC1 | ANO10 | SLC7A1 | NOP53 | HNRNPC | MRRF | CDH1 |
| **V56** | LOXL2 | TMEM14A | SLC5A3 | SH3D21 | SPATA13 | AIF1L | POSTN | SLC7A5 | SLC35A4 | PIR |
| **V57** | THBS2 | STAG2 | CFI | MDM2 | TK2 | NRIP1 | C2 | CD4 | ZNF595 | MXI1 |
| **V58** | EHD2 | ARL2BP | DNASE2B | METTL9 | NKIRAS2 | PHACTR4 | ATRX | TMEM165 | DBN1 | KCNJ13 |

| | | | | | | | | | | |
|---|---|---|---|---|---|---|---|---|---|---|
| **V59** | IGFBP5 | JPT2 | QPCT | TTC39A | COX7A2 | IMPACT | ARL6IP4 | ITM2C | SLC9A3R2 | LRRTM4 |
| **V60** | TNXB | KIRREL1 | EYA1 | RAB32 | KLF10 | MPPED2 | ITPRID2 | PMVK | NFATC2IP | FKBP10 |
| **V61** | LSP1 | PLEKHA2 | CDKN2B | UQCRQ | HSPA5 | NPL | CORO2B | PLEKHM2 | KDM1A | SCARF2 |
| **V62** | LRRC15 | ITGB3 | PTPRZ1 | EGR1 | PPHLN1 | BAX | VPS13C | UTP4 | DIPK1B | NOP53 |
| **V63** | EGFL7 | PIGT | GPR37 | MXI1 | PLBD2 | BHLHE41 | HSD17B11 | ACADM | HNRNPC | TRIM63 |
| **V64** | CXCL14 | TMEM167A | ALDH1A1 | MYC | ARHGDIB | ST6GAL1 | HSPH1 | HSPA2 | SRRM2 | ENPP2 |
| **V65** | PALLD | ITPRID2 | BFSP1 | MYBL2 | CDK4 | ITPKB | CRYL1 | S100P | ITGAX | ENC1 |
| **V66** | MFAP4 | HSPH1 | CADPS | PFKP | URGCP | CERS4 | ERBB3 | TUBG1 | BCL2 | APOC1 |
| **V67** | RGS5 | SRPRB | FGL2 | SH3BP4 | ITCH | VTA1 | STAG2 | AK5 | GSTM3 | PAG1 |
| **V68** | PXDN | IFI30 | THBS2 | RAB6B | VPS54 | CCDC50 | HSPB8 | PLP2 | KYAT3 | PAX3 |
| **V69** | SLC9A3R2 | SPCS3 | PDK4 | CDK1 | COL4A2 | NAV2 | BCAN | ECE1 | TSPAN31 | RBMS3 |
| **V70** | CRABP2 | TMCO3 | AKAP6 | BHLHE41 | HELZ2 | GAS1 | SORBS2 | IGHE | HNRNPA0 | KDELR2 |
| **V71** | ANTXR1 | COPE | PYCARD | MET | CREM | CSGALNACT1 | IRF4 | STARD3NL | ZCCHC14 | NME1 |
| **V72** | VWF | IAH1 | AZGP1 | EIF1AY | CDKN1A | SCN1B | ATP5MC2 | EIF4G2 | ICMT | GPRC5D |
| **V73** | CD93 | TSNAX | CD55 | STXBP1 | KLF6 | SLC35A4 | AZGP1 | COL6A2 | TMEM38B | LDHB |
| **V74** | SULF1 | IFI44 | PRELP | DHFR | FGFR1 | SESN3 | PLEC | SLC35A4 | SLC16A3 | SOX9 |
| **V75** | COL5A3 | USP16 | ITIH6 | ATP5MC3 | CTSS | TLK1 | ST6GAL1 | RACK1 | AK1 | ZNF703 |
| **V76** | DIO2 | TJP1 | ANGPTL7 | SLC16A1 | FAM126A | IFI6 | ALDH1A1 | NDUFA3 | ENDOV | PTGFRN |
| **V77** | CALCRL | TIMMDC1 | WNK4 | LAMB2 | HSPH1 | IRF2BP2 | SNX10 | CDH19 | ISYNA1 | SH3BP4 |
| **V78** | PODXL | EIF3L | CRYL1 | ATP6AP1 | BCL2L11 | PIGK | ATP6AP2 | SLC2A4RG | PAG1 | TGFBI |
| **V79** | ID1 | ATP6V1H | ABCA6 | OAS2 | NOP2 | HGS | TMEM140 | KDM1A | LARP1 | GJB2 |
| **V80** | ID3 | VPS13C | PLAT | VAV3 | CD55 | HIBADH | INPP5F | KIFC3 | NRAS | SLITRK2 |
| **V81** | FAP | KLHDC3 | EEF1G | ZFYVE16 | EEF1G | TCF7L2 | NDRG1 | AMFR | PDLIM3 | AKAP6 |
| **V82** | RHOB | FBL | SORBS2 | C2 | MOSPD2 | EEF1G | TMEM59 | OTOR | PDLIM7 | MOSPD2 |
| **V83** | THBS1 | PHB2 | CEBPD | SDHB | ATP6AP2 | COX7A2 | SPTAN1 | ILVBL | LRP8 | ABCA8 |
| **V84** | PLAU | FH | ASAH1 | GAB2 | ANXA6 | SCARF2 | SRPK2 | SEPTIN4 | UCK1 | FAM126A |
| **V85** | C3 | TMEM106B | SNX10 | PLXNC1 | HNRNPF | HSPA2 | SYNM | CT55 | UQCRH | TDRD3 |
| **V86** | GJA1 | TMEM173 | GPX3 | WIPF3 | HSBP1 | NDRG2 | TJP1 | CIZ1 | ATAD3B | LY96 |
| **V87** | CPE | PRMT2 | CLEC11A | IFI27 | FKBP14 | AFF3 | APOC1 | VPS51 | MT1E | TOMM7 |
| **V88** | CAVIN1 | SEPTIN2 | SLC40A1 | ANKRD28 | IPO5 | HEBP1 | TMEM106B | IMPA2 | TGDS | PINX1 |

| | | | | | | | | | | |
|---|---|---|---|---|---|---|---|---|---|---|
| **V89** | PI16 | CSTB | CPM | BHLHE40 | BAG1 | HSPA12A | IFNAR1 | LAPTM5 | SDC3 | SLC7A5 |
| **V90** | STAB1 | ITGAV | RBMS3 | MSC | ATOX1 | VPS54 | SOD1 | ADGRG1 | SLC2A4RG | ARHGDIB |
| **V91** | ACTA2 | ARHGAP18 | ID4 | ABCC2 | IFIT2 | SMPDL3A | TOMM7 | SLC19A1 | KAZN | PRRX1 |
| **V92** | DUSP1 | ATRX | ERBB3 | CENPF | UGCG | PPHLN1 | ALDH1A3 | AURKAIP1 | NCF2 | COL1A2 |
| **V93** | SELENOP | RASA1 | ABCA8 | UQCRH | IFIT1 | FOS | FN1 | MYC | IKBKB | LNPK |
| **V94** | SPON1 | HTT | SERPINA3 | NR4A1 | SPP1 | SATB2 | RBMS3 | SBF1 | FOSL2 | ECM1 |
| **V95** | NPDC1 | KHDRBS1 | TF | FSTL5 | SEPTIN2 | MET | MFSD12 | LDHB | LSP1 | UGCG |
| **V96** | EMP1 | INO80E | RXRG | NQO1 | KCNN4 | GSTM3 | CIITA | ECM1 | TUBB2B | BGN |
| **V97** | DPT | HNRNPLL | PTPRG | UBE2T | IFI16 | KYAT3 | UBA1 | RCAN1 | CARD16 | TTYH3 |
| **V98** | MMP11 | TMEM109 | STAB1 | CDK4 | UBA1 | GAB2 | LRP2 | ARHGDIB | PPARD | SNX10 |
| **V99** | PAM | SRPRA | METRN | MSANTD3 | SDE2 | PIGT | IFI27 | LARP1 | NFIB | MDM2 |
| **V100** | CCN1 | JAK1 | MFSD12 | MICOS10 | PGM2L1 | ST3GAL1 | PTPRZ1 | SRRM2 | KIF26B | PDGFD |
| **V101** | BICC1 | ATP5MC1 | LPAR1 | TFAP2A | MMD | HSPH1 | SNX17 | CRYL1 | CDC42EP3 | PCOLCE |
| **V102** | SPARCL1 | IRAK1 | CYGB | XYLT1 | FLOT1 | SLC25A16 | EEF1G | UQCRH | VPS51 | BZW2 |
| **V103** | JUNB | IFRD1 | TOMM7 | WDR34 | ABHD2 | SRRM2 | ST3GAL1 | NR4A1 | RACK1 | CORO2B |
| **V104** | OLFML2A | TMEM127 | ATP1B2 | LIPA | IQCE | GGCT | HIBADH | KCNAB2 | MC1R | TFAP2A |
| **V105** | HSPG2 | BAHCC1 | HSPA6 | TMEM97 | SFXN3 | IGFBP2 | ATOX1 | STARD13 | PTGES | JAG1 |
| **V106** | FOSL2 | TMEM11 | STXBP1 | LAMA1 | AHCY | ITGAV | SLC26A2 | PAG1 | KRT19 | EEF1G |
| **V107** | SETBP1 | HIBADH | C2 | SNX25 | IARS2 | RNASE1 | ITCH | TTYH3 | IRF2BP2 | RELL1 |
| **V108** | METRNL | PRPF8 | PPIA | MICAL1 | ATRX | FADS2 | ST5 | PIGT | PRKD3 | ALDH2 |
| **V109** | FAM20C | ITGA7 | LYZ | PERP | CMPK2 | IGF1R | TMEM167A | AZIN1 | IFNL1 | AIF1L |
| **V110** | SERPINE1 | PLBD2 | NOP53 | SUSD5 | CHST11 | HIBCH | IFRD1 | SLC20A1 | SATB2 | SHISA2 |
| **V111** | ANPEP | PRRC2A | GASK1B | MTURN | MX1 | OAS1 | SRPRB | NFKBIA | PSME4 | OAF |
| **V112** | ITGBL1 | KANSL1 | FLOT1 | PRAME | IFNAR1 | AFDN | ST3GAL5 | INO80E | IMPA2 | MYADM |
| **V113** | COMP | QDPR | TUBB2B | POLD2 | CD300E | KLF16 | STAU2 | ZFYVE16 | BAG1 | PLXNC1 |
| **V114** | LAPTM5 | SSH1 | GBP2 | NMRK2 | NCL | EDNRB | SPAG7 | IFITM2 | CPNE2 | ADAMTS3 |
| **V115** | CEBPD | AURKAIP1 | PCSK1N | CAPG | RCAN1 | NBR1 | HPGD | IER3 | PTCH1 | CA8 |
| **V116** | CTSK | HNRNPH2 | MRGPRX3 | CPVL | PMAIP1 | LRP4 | TMEM173 | FABP5 | PIGK | GLRA2 |
| **V117** | GAS6 | ICAM1 | MITF | SEPTIN3 | SDHB | VPS4B | FBL | UBA2 | KCNAB2 | CD79B |
| **V118** | OAF | HPS4 | POLD2 | ANKRD37 | INO80E | FRMD5 | FKBP11 | SPPL2B | UBA2 | COL5A2 |

| | | | | | | | | | | |
|---|---|---|---|---|---|---|---|---|---|---|
| **V119** | FBLN5 | SEC61G | APOC2 | VAMP8 | IFT52 | SRPK2 | COPE | GAS1 | IER3 | CA5B |
| **V120** | ELN | FKBP10 | CDKN1A | CDC25B | MX2 | FLOT1 | PLEKHA2 | MOSPD2 | CEBPB | PDLIM4 |
| **V121** | RGCC | STAT3 | ENPP2 | NR4A3 | PRSS33 | ZFYVE16 | LGI4 | TM9SF2 | ILVBL | ATP5MC3 |
| **V122** | ADIRF | LAMTOR5 | COL22A1 | DIPK1B | IMPACT | METRN | IGHA1 | ARPC1A | CCER2 | MYO1D |
| **V123** | SFRP4 | TIMM44 | LDHB | KDM1A | CSTB | CMBL | TMEM147 | KLF16 | TMCC2 | NT5DC3 |
| **V124** | CPXM1 | SEPTIN7 | MMP16 | ZWINT | TWSG1 | HK1 | HCN2 | SLC9A3R2 | NAPG | TUSC3 |
| **V125** | MPEG1 | SAP30BP | IER3 | MLIP | HPCAL1 | NKIRAS2 | HSPA2 | METRN | ADGRG1 | PHLPP1 |
| **V126** | OLFML3 | TIMM17A | SERPINF1 | PPIA | USP53 | CSF2RA | IDH2 | MRRF | UQCRQ | SLC16A1 |
| **V127** | ANKH | CDH1 | CADM1 | ATP5PD | SPCS3 | HNRNPF | TIMM8B | S100A4 | INO80E | SH3D21 |
| **V128** | CD36 | PIK3R1 | FAM89A | FAM89A | ADAM10 | STK32A | STON1 | AHCY | SLC19A1 | AFF3 |
| **V129** | NFIB | ASH2L | SMOX | CTSS | SHOC2 | GDF11 | IBTK | INTS1 | IPO5 | CDC42EP4 |
| **V130** | TMEM255B | AZIN1 | NELL1 | FCER1G | ATP5PD | SLC19A1 | POMP | LNPK | SPPL2B | TNFRSF21 |
| **V131** | MEGF6 | KPNA2 | HMOX1 | MAL | ATP5MC2 | PINX1 | ARL2BP | PPIA | HILPDA | NXT2 |
| **V132** | NREP | EIF3M | CDH19 | MCC | ITGAV | PMVK | SLC1A4 | UPK1B | MYL12A | ZNF697 |
| **V133** | PDLIM7 | ADAM10 | VEGFA | NREP | SH3RF2 | JUNB | VAMP8 | EGR1 | HELZ | MITF |
| **V134** | GAS1 | SAP30L | SNED1 | SERBP1 | CEACAM5 | CRISPLD1 | GPX3 | FOSL2 | SLC39A13 | ABCA6 |
| **V135** | MMP19 | GSTO1 | CA5B | AK4 | CXCL6 | PLXNC1 | IMMP2L | SATB2 | CD4 | ABCA10 |
| **V136** | VWA1 | VPS54 | FAM126A | TTYH3 | SLC35A4 | IMPA2 | STAT3 | PDP1 | COL4A1 | PYCARD |
| **V137** | RAB31 | ITPR2 | MAOB | FUCA1 | PMVK | CDC25B | CRTAC1 | KIRREL1 | IGSF3 | EPHX1 |
| **V138** | ITGA5 | ATP5PD | CCN2 | PTTG1 | MICOS10 | ZMAT2 | KPNB1 | HIBADH | NXT2 | PPIA |
| **V139** | HMOX1 | IFIT3 | PCSK6 | CA14 | SLC9A3R2 | ADAM10 | HSPA1B | GSTM3 | SMAP2 | RND2 |
| **V140** | MXRA5 | PLEKHM2 | CYP39A1 | MMP17 | CADM1 | UQCRQ | FCGR2A | CTDSP2 | ZNF697 | TSC22D1 |
| **V141** | CPA3 | ARL6IP4 | CRISPLD1 | SNX9 | UQCRC1 | RACK1 | RSL1D1 | ENC1 | VEZF1 | ALDH1A2 |
| **V142** | EFEMP1 | NBR1 | ARHGAP6 | CD55 | AMFR | PIEZO2 | TIMMDC1 | IFT52 | NRIP1 | HTR2B |
| **V143** | ECE1 | SPTLC2 | NDRG2 | GJB1 | CCND1 | RARA | HKDC1 | HMCN1 | HSPG2 | TRPM1 |
| **V144** | SOCS3 | TFAP2A | ABCC11 | AFF3 | SOCS3 | TMBIM1 | EYA1 | PSMB3 | IQSEC1 | MSC |
| **V145** | CD4 | ARL6IP1 | PINX1 | RETSAT | SETBP1 | ZNF595 | ASB11 | NFATC2IP | TRAPPC6A | ZFYVE16 |
| **V146** | SDC1 | IFIH1 | SESN3 | MYL12A | SRP72 | NFIB | SMTN | SPATA13 | POLD2 | LGI4 |
| **V147** | C12orf75 | BAMBI | NRCAM | TUBB2B | ICMT | IFT52 | PIK3R1 | PLXNC1 | SLC7A1 | H2AFJ |
| **V148** | CAMK2N1 | TMEM120A | ARRDC3 | SGK1 | FBXW5 | URGCP | SRPRA | TGDS | HNRNPR | CERS4 |

| | | | | | | | | | | |
|---|---|---|---|---|---|---|---|---|---|---|
| **V149** | MAP1B | RAB1A | TUSC3 | HSPA5 | NUDCD3 | ACVR2A | TIMM50 | SPI1 | AZIN1 | NRIP1 |
| **V150** | CDH13 | IFI27 | AHR | GRK3 | COL6A1 | DCXR | COX7A2 | ACAT2 | AK5 | PMEL |

| | NMF21 | NMF22 | NMF23 | NMF24 | NMF25 |
|---|---|---|---|---|---|
| **V1** | IGFBP5 | HIST1H1B | DCT | CRYAB | MDM2 |
| **V2** | COL1A1 | PLXNC1 | MGP | GDF15 | NRAS |
| **V3** | COL3A1 | TOP2A | RACK1 | FN1 | S100A9 |
| **V4** | IGHG1 | NCAPG2 | EEF1G | PLA1A | PRKD3 |
| **V5** | COL1A2 | TTYH3 | EDNRB | ABCB5 | CDK4 |
| **V6** | IGKC | CENPF | IGKC | GBP1 | S100A8 |
| **V7** | MGP | RETSAT | PRAME | CCN3 | UGCG |
| **V8** | THBS2 | SNX10 | ERBB3 | IFI6 | QPCT |
| **V9** | CCN2 | AHCY | IGHG1 | IGKC | TYRP1 |
| **V10** | ADAMTS1 | TK1 | PLP1 | CCND1 | KRT5 |
| **V11** | COL6A1 | NME1 | NOP53 | RACK1 | STXBP1 |
| **V12** | COL4A2 | ARPC1A | MYC | QPCT | CPM |
| **V13** | AFAP1L2 | LDHB | DCLK1 | GBP2 | KRT16 |
| **V14** | COL4A1 | ST3GAL5 | SPP1 | EGR1 | TSFM |
| **V15** | COL6A2 | H2AFJ | MMP17 | SPP1 | TTYH3 |
| **V16** | S100A4 | GJB1 | TFAP2A | SLC26A2 | ABHD2 |
| **V17** | RHOB | EDNRB | IGFBP7 | ANXA2 | BMPR2 |
| **V18** | IGFBP2 | HPS4 | COL3A1 | TNFRSF12A | PERP |
| **V19** | COL18A1 | TUBG1 | FAM126A | IGFBP7 | SLC25A4 |
| **V20** | COL6A3 | MICOS10 | RHOB | GBP5 | CDC42EP3 |
| **V21** | FN1 | PAX3 | MSC | IDO1 | TSPAN31 |
| **V22** | SORCS1 | POLD2 | ID4 | RHOB | KRT6A |
| **V23** | EGR1 | TOMM7 | MXI1 | TNC | C21orf91 |
| **V24** | COL5A1 | UQCRH | STK32A | ZFYVE16 | APCDD1 |
| **V25** | AQP1 | ANLN | SEMA6A | EMP1 | KRT6B |
| **V26** | COL5A2 | ADGRG1 | MMP2 | SPOCK1 | CDK1 |
| **V27** | PLAT | MITF | COL6A1 | IGHG1 | SGK1 |
| **V28** | BGN | CDK4 | COL11A2 | RXRG | LMBRD2 |

| | | | | | |
|---|---|---|---|---|---|
| **V29** | PTN | GLMP | ENPP2 | NDRG1 | CPVL |
| **V30** | NR4A1 | HIST1H4C | COL6A2 | CITED1 | MARCH9 |
| **V31** | TAGLN | IDH3B | BGN | CPN1 | MYO1D |
| **V32** | POSTN | HIST1H1E | COL18A1 | NR4A1 | RGMB |
| **V33** | TNC | MMP17 | GRK3 | WIPI1 | WIPI1 |
| **V34** | TGFBI | MCOLN3 | BCL2 | IER3 | SLC1A5 |
| **V35** | PMEPA1 | E2F1 | PRELP | LYPD1 | LYZ |
| **V36** | VCAN | SDHB | ADAMTS1 | CDKN1A | ANKRD28 |
| **V37** | IGFBP7 | BIRC5 | COL2A1 | CXCL10 | CSTA |
| **V38** | MXI1 | RAB32 | TMOD1 | UGCG | METTL1 |
| **V39** | GPC1 | NRAS | ALDH1A1 | TSPAN10 | CDC25B |
| **V40** | PTPRZ1 | TTYH2 | PTPRZ1 | CIITA | S100A2 |
| **V41** | IGFBP3 | TFRC | ETS2 | CCL5 | CISD1 |
| **V42** | CADM1 | PPIA | PLAT | SERPINF1 | ANXA2 |
| **V43** | IGFBP4 | SPC25 | PDGFD | CEBPB | CLU |
| **V44** | COL7A1 | HIST1H1D | ARRDC3 | RLBP1 | CHST11 |
| **V45** | LOXL2 | MYBL2 | CYGB | MXI1 | MTURN |
| **V46** | COL12A1 | ITM2C | PLEKHB1 | COL11A1 | DSP |
| **V47** | CCN1 | ICMT | ASPA | FOS | KRT17 |
| **V48** | MMP2 | UBE2T | NDRG2 | BHLHE40 | FOS |
| **V49** | ID4 | CDC20 | SFRP1 | ERBB3 | CTDSP2 |
| **V50** | FBN1 | ARL6IP4 | ABCB5 | EEF1G | NPDC1 |
| **V51** | SHOC2 | ZNF703 | RBMS3 | HNRNPF | ASAH1 |
| **V52** | LGI4 | HMCN1 | CRISPLD1 | FLOT1 | SLC1A4 |
| **V53** | ITGA1 | TMEM97 | GAB2 | STXBP1 | CTSH |
| **V54** | ENC1 | GGCT | PHLPP1 | RTP4 | ANXA6 |
| **V55** | ERBB3 | PTTG1 | AQP1 | CDC25B | CDH3 |
| **V56** | NOTCH3 | PBK | CDH19 | C3 | TOP2A |
| **V57** | NRXN2 | COX7A2 | COL1A1 | CXCL9 | ZFYVE16 |
| **V58** | NREP | DHFR | IGHA1 | ISG15 | CAVIN1 |

| | | | | | |
|---|---|---|---|---|---|
| **V59** | CASP7 | ATP5MC2 | NREP | NT5DC3 | PXK |
| **V60** | SULF1 | KPNA2 | PTN | ENO1 | SPRR1B |
| **V61** | PDLIM7 | ETV4 | ETV1 | OAS2 | IGFBP7 |
| **V62** | DCD | METTL9 | GDF11 | MET | IGKC |
| **V63** | TCF7L2 | SLC45A2 | COL15A1 | CCN1 | BHLHE40 |
| **V64** | PDGFA | BZW2 | POSTN | TRAC | PDK4 |
| **V65** | FOXD3 | ATP5MC3 | GASK1B | IFITM2 | FABP5 |
| **V66** | VTI1A | WDR34 | MTURN | PTTG1 | PDE4D |
| **V67** | NHLRC2 | FABP5 | SMOX | LYZ | B4GALT5 |
| **V68** | S100A16 | SLC19A1 | MCC | RNASE1 | CREM |
| **V69** | ATRNL1 | ASAH1 | COL16A1 | SGK1 | SFN |
| **V70** | COL5A3 | ARRDC4 | HMCN1 | TRIM63 | CHL1 |
| **V71** | PGM2L1 | ZWINT | PAG1 | NME1 | CCND1 |
| **V72** | MDK | PSEN2 | AIF1L | FGL2 | USP53 |
| **V73** | PALLD | PRMT2 | WIPF3 | SERPINA1 | RAB6B |
| **V74** | CDH19 | ACAT2 | NFIB | PRDM7 | RAB32 |
| **V75** | MYL9 | BAX | PCBP2 | VGF | NBL1 |
| **V76** | GLI3 | IARS2 | RXRG | FAM126A | NR4A1 |
| **V77** | EFS | UQCRQ | COL1A2 | RGMB | SH3BP4 |
| **V78** | MXRA5 | UTP4 | TMEM97 | ECM1 | MCOLN3 |
| **V79** | ABHD2 | HSPA2 | LPL | ITGB8 | MICAL2 |
| **V80** | STARD13 | PMEL | NDRG1 | SLC16A3 | SNX9 |
| **V81** | PDLIM4 | MPPED2 | FAM118A | SLC7A5 | IQSEC1 |
| **V82** | NGFR | TLK1 | PPIA | TUSC3 | GOLGA7B |
| **V83** | RXRG | RRM2 | SOD3 | MYC | MYL12A |
| **V84** | FOS | MXI1 | CTDSP2 | FCRLA | EPHX1 |
| **V85** | ANKH | ENC1 | PKNOX2 | SOD2 | SNX25 |
| **V86** | COL8A1 | ILVBL | TTC28 | MDM2 | SDC3 |
| **V87** | DUSP1 | ALDH1A3 | SHISA2 | KCNAB2 | DCXR |
| **V88** | STRA6 | CENPH | TAFA5 | SDC3 | GJB1 |

| | | | | | |
|---|---|---|---|---|---|
| **V89** | PRRX1 | ATP5PD | MITF | MRGPRX3 | SLC24A4 |
| **V90** | THY1 | PKMYT1 | SLC19A1 | PRKD3 | FUCA1 |
| **V91** | AKAP12 | HIBADH | C7 | ID4 | CD55 |
| **V92** | PALM2-AKAP2 | STARD3NL | NRIP1 | MAP1B | SEMA5A |
| **V93** | COL15A1 | DIPK1B | IGFBP5 | CD5L | NME1 |
| **V94** | CDH2 | NCAPG | EGR1 | TGFBI | KRTDAP |
| **V95** | MRC2 | TSFM | SETBP1 | CD55 | SBSN |
| **V96** | TRAF1 | PIR | AKAP12 | ENTHD1 | NADK2 |
| **V97** | HSPA12A | POLE3 | CERS4 | C2 | SEMA6D |
| **V98** | MMP16 | ACACA | CYSLTR2 | LAPTM5 | ENO2 |
| **V99** | COMP | ACADM | MYADM | PDP1 | SLC31A1 |
| **V100** | LUM | IMPA2 | IL17D | BIRC7 | ZNF697 |
| **V101** | PXDN | ERBB3 | PCDH7 | VAMP8 | FAM126A |
| **V102** | CDH6 | UCK1 | SDC2 | TRAF1 | TNFRSF21 |
| **V103** | ALDH1A3 | CDK1 | DIPK1B | MCC | PI3 |
| **V104** | ACTA2 | AFF3 | VPS51 | PERP | ITPKB |
| **V105** | SDC2 | PSMB3 | GRIK3 | HMGA2 | FRMD4A |
| **V106** | DCN | ISOC2 | LGI4 | MFSD12 | TRIM29 |
| **V107** | PDGFRB | SLC19A2 | MYO15B | OAS1 | TFRC |
| **V108** | CCDC186 | HMMR | IRF2BP2 | PDE4D | WWC3 |
| **V109** | FRMD4A | KIF18B | CHL1 | TTC39A | MDK |
| **V110** | TCIM | IMMP2L | SCARF2 | PLP2 | AKAP12 |
| **V111** | CTDSP2 | AURKAIP1 | GOLGA8B | JUNB | COL6A3 |
| **V112** | CPXM1 | ANKRD9 | PAX3 | IER5L | EEF1G |
| **V113** | FAM126A | SELENON | EMP1 | DSG2 | MYBL2 |
| **V114** | TNFAIP6 | LRP8 | TUBB2B | SLC20A1 | ZWINT |
| **V115** | ECE1 | NUF2 | LRP5 | LRATD2 | LYPD3 |
| **V116** | RACK1 | IGSF3 | CRYAB | HSPB2 | SLC2A1 |
| **V117** | MAP1B | PTGFRN | CA5B | LCP1 | EDNRB |
| **V118** | FAM20C | PLXNA1 | RND3 | TRBC2 | CNFN |

| | | | | | |
|---|---|---|---|---|---|
| **V119** | EHD2 | PIGT | SOX2 | BHLHE41 | IGHG1 |
| **V120** | SCARF2 | SLC39A6 | ARHGDIB | CCDC50 | DSC2 |
| **V121** | NRXN1 | STAU2 | FGFBP2 | ABCC2 | CAPG |
| **V122** | NRCAM | AK1 | CCDC50 | APOC1 | CTNNB1 |
| **V123** | IGHA1 | PTPN14 | PRKG1 | CREM | KRT6C |
| **V124** | GUCY1A1 | IPO5 | ABCA9 | DCT | ZNF703 |
| **V125** | DENND2A | CDCA4 | STMN3 | ENO2 | TMEM97 |
| **V126** | GFRA1 | CRISPLD1 | PDZRN3 | TNFAIP3 | IGFBP3 |
| **V127** | LAMA5 | VPS41 | CYP7B1 | CXCL13 | SLC39A6 |
| **V128** | COL9A3 | HES1 | CORO2B | EIF1AY | LSAMP |
| **V129** | PRKG1 | SLC7A1 | MX2 | COL3A1 | CADM1 |
| **V130** | ANO1 | SESN3 | ENC1 | GRK3 | TRPM1 |
| **V131** | FAM118A | CCL17 | SORBS2 | GAB2 | NQO1 |
| **V132** | PCDH20 | CDKN1C | CDC42EP1 | COL16A1 | FXYD6 |
| **V133** | ERRFI1 | MYO1D | RGS5 | UBD | MMD |
| **V134** | LPAR1 | AIF1L | FGL2 | SLC1A4 | CALML5 |
| **V135** | SULF2 | ADGRG6 | KCNJ13 | MITF | NCS1 |
| **V136** | TUBB2B | FRMD5 | S100A4 | LDHB | COL3A1 |
| **V137** | SERPINE1 | NFATC2IP | CRMP1 | SERPINE1 | HIST1H1B |
| **V138** | ISLR | HELZ | POLD2 | EPSTI1 | TK1 |
| **V139** | GEM | DBN1 | CTSK | B4GALT5 | PSAT1 |
| **V140** | DBN1 | SATB2 | LRP3 | FCER1G | COL6A2 |
| **V141** | FREM2 | HIBCH | CSGALNACT1 | CARD19 | TTYH2 |
| **V142** | RGMB | SRPK2 | NR4A1 | PLAAT4 | ATP23 |
| **V143** | ANGPTL2 | SLC22A18AS | CCDC8 | LSP1 | BHLHE41 |
| **V144** | ELN | HSDL1 | C2orf88 | SYNM | EEF1AKMT3 |
| **V145** | ANTXR1 | C2orf88 | CRYL1 | PPIA | DHFR |
| **V146** | SLC9A3R2 | MELK | PLXNB1 | ANXA6 | NOP53 |
| **V147** | RND3 | PCDH7 | DLL3 | PALM2-AKAP2 | FAM171B |
| **V148** | SNX22 | TMCC2 | IGFBP3 | AKAP12 | CTSK |

| | | | | | |
|---|---|---|---|---|---|
| **V149** | DAGLA | SNX4 | TUSC3 | CREB5 | GPR161 |
| **V150** | RCAN1 | SLC35A4 | SOX9 | PHLDA2 | COL6A1 |

**Pathway analysis NMF and MP's**

| Meta program (MP) | Defined by pathways | NMF | Pathways | Average NES |
|---|---|---|---|---|
| **MP1** | **Lipid metabolism** | **NMF14** | Response to corticosteroid | 1.57 |
| | **Stress response** | | Retinoic acid | 1.76 |
| | **Melanosomal biogenesis (NMF14)** | | Amino acid transmembrane transport | 1.6 |
| | **Growth regulation (NMF24)** | | Secreted factors (PEDF axis) | 1.59 |
| | **IFN response (NMF24)** | | Melanosomal biogenesis | 1.66 |
| | | | Starvation response | 1.5 |
| | | | Pigmentation / melanin synthesis | 1.54 |
| | | | Lipid metabolism | 1.66 |
| | | | Metal ion transport | 1.6 |
| | | | Oxidative phosporylation/mitochondrial ETC | FALSE |
| | | | Cell cycle | FALSE |
| | | **NMF24** | MAPK cascade signaling; | 1.56 |
| | | | SMAD negative regulation | 1.56 |
| | | | Cellular response to unfolded protein | 1.51 |
| | | | Lipid catabolism | 1.57 |
| | | | Regulation of cell growth / growth arrest | 1.65 |
| | | | Cell adhesion / migration | 1.57 |
| | | | Regulation of inflammatory response | 1.6 |
| | | | Response to IL-1 | 1.59 |
| | | | Response to type II IFN | 1.51 |
| | | | Cell cycle arrest (G1/S checkpoint) | 1.67 |
| | | | Stress survival factors | FALSE |
| | | | T cell chemokines | FALSE |
| | | | Type I IFN response (ISGs) | FALSE |
| | | | Hypoxia | FALSE |
| **MP2** | **NRAS/CDK4/MDM2 axis** | **NMF3** | NRAS/CDK4/MDM2 axis | FALSE |
| | **p53 pathway (NMF25)** | | Lipid metabolism | 1.52 |
| | **Cell cycle (NMF8)** | | Nucleolar biogenesis | 1.61 |

| | | | | |
|---|---|---|---|---|
| | | | Secreted factors (GDF15-VGF axis) | 1.54 |
| | | | Actin remodeling | FALSE |
| | | | Pigmentation transcription factors | 1.51 |
| | | | Intracellular / vesicle transport | 1.51 |
| | | | Cell cycle | 1.56 |
| | | **NMF25** | Innate immune response | 1.65 |
| | | | Pigmentation / melanin synthesis | 1.63 |
| | | | NRAS/CDK4/MDM2 axis | FALSE |
| | | | Mitotic kinases | 1.68 |
| | | | p53 pathway | 1.64 |
| | | | Membrane / vesicle organization | 1.58 |
| | | | Cytokeratin / epidermal differentiation | 1.63 |
| | | | Autophagy | 1.61 |
| | | **NMF8** | NRAS/CDK4/MDM2 axis | FALSE |
| | | | Cell cycle / G1-S phase transition | 1.62 |
| | | | Skin/epidermis development | 1.64 |
| | | | Amino acid transport (SLC1A5 / SLC7A11) | 1.59 |
| **MP3** | **ECM** | **NMF2** | Small GTPase / RHO signaling | 1.75 |
| | **Pigmentation (NMF2)** | | Fibrillar collagens (real ECM) | 1.6 |
| | **DMA damage response (NMF5)** | | Collagen-containing ECM / ECM organization | 1.67 |
| | | | Melanosomal biogenesis / pigment granule | 1.62 |
| | | | Pigmentation / melanin synthesis | 1.66 |
| | | | Type I IFN response (ISGs) | FALSE |
| | | | Integrin-mediated adhesion | 1.65 |
| | | **NMF5** | Cell junction assembly | 1.58 |
| | | | Inorganic ion transport (FXYD3-driven) | 1.57 |
| | | | Reactive oxygen species response | 1.58 |
| | | | Response to retinoic acid | 1.55 |
| | | | Response to corticosteroid | 1.57 |
| | | | DNA damage response / kinase activity | 1.58 |

| | | | | |
|---|---|---|---|---|
| | | | Cell cycle arrest / senescence | 1.62 |
| | | | Centrosome / microtubule organization | 1.65 |
| | | | ECM / matricellular proteins | 1.67 |
| | | | Serine proteases / fibrinolysis / coagulation | 1.64 |
| | | | Regeneration / wound healing | 1.67 |
| **MP4** | **Pigmentation / melanin synthesis** | **NMF1** | Collagen binding / cathepsin-mediated catabolism | 1.53 |
| | **Metal ion transmembrane transport (NMF1)** | | Metal ion transmembrane transport (zinc/iron) | 1.61 |
| | **p53 signaling (NMF6)** | | Melanosomal biogenesis / pigment granule | 1.52 |
| | **IFN response (NMF6)** | | Pigmentation / melanin synthesis | 1.54 |
| | **ROS species metabolism (NMF6)** | | Cell adhesion molecules | FALSE |
| | | **NMF6** | Pigmentation / melanin synthesis | 1.53 |
| | | | Synaptic / neural | 1.53 |
| | | | Negative regulation of molecular function / inhibitors | 1.52 |
| | | | Kinase activity (PKD/SGK) | 1.53 |
| | | | Anion / vesicular transport | 1.55 |
| | | | Lysosomal / granule biology (myeloid-like) | 1.63 |
| | | | Cellular response to chemical stress / oxidative | 1.52 |
| | | | p53 signaling / cell cycle arrest | 1.64 |
| | | | Immune infiltration | FALSE |
| | | | Viral / IFN response | 1.64 |
| | | | Reactive oxygen species metabolism | 1.51 |
| | | | Type I IFN signaling / JAK-STAT | FALSE |
| **MP5** | **Immune** | **NMF7** | Humoral immune response / B cell biology | 1.92 |
| | **Myeloid / macrophages** | | B cell receptor signaling / immunoglobulins | 1.85 |
| | **B cells (NMF7)** | | Inflammatory response | 1.74 |
| | **T cells (NMF9)** | | Leukocyte migration (T/B/NK) | 1.76 |
| | **Complement (NMF9)** | **NMF9** | Myeloid leukocyte / innate immunity | 1.7 |
| | | | Macrophage markers | FALSE |
| | | | Lysosomal / vacuolar / phagocytic granules | 1.73 |
| | | | Actin-based motility / leukocyte migration | 1.72 |

| | | | | |
|---|---|---|---|---|
| | | | Antigen processing & presentation | 1.68 |
| | | | Complement activation | 1.61 |
| | | | Antigen receptor signaling (BCR/TCR) | 1.67 |
| | | | Immunoglobulins / B/plasma cells | 1.64 |
| | | | T cell markers / TCR | FALSE |
| | | | Pro-inflammatory cytokines | FALSE |
| | | | Type I IFN response (ISGs) | FALSE |
| | | | Type II IFN / CXCR3 chemokines | FALSE |
| **MP6** | **Secreted factors (NMF20)** | **NMF20** | Endocrine / hormone secretion | 1.7 |
| | **Cytoskeleton** | | Response to ketone / corticosteroid hormone | 1.74 |
| | **Cell morphogenesis** | | ECM proteins / matricellular | 1.68 |
| | **MYC (NMF23)** | | Secreted growth factor regulators (IGFBPs) | 1.88 |
| | **Programmed cell death (NMF23)** | | Cell morphogenesis / migration | 1.64 |
| | **Mitochondrial (NMF23)** | **NMF23** | Proteasomal protein catabolism | 1.68 |
| | **Response to hypoxia (NMF13)** | | MYC oncogene transcription program | 1.63 |
| | **Stress response (NMF13)** | | Protein kinases / signaling | 1.71 |
| | | | Apoptosis / programmed cell death | 1.65 |
| | | | Mitochondrial | 1.61 |
| | | | Pigmentation / melanocyte identity | 1.64 |
| | | | Hypoxia / NDRG axis | FALSE |
| | | | Cytoskeleton / cell shape | 1.69 |
| | | **NMF13** | Cellular response to hypoxia / oxygen | 1.55 |
| | | | Microtubule / cytoskeleton | 1.5 |
| | | | Negative regulation of cell growth | 1.75 |
| | | | Wnt pathway regulation (SFRP1-driven) | 1.5 |
| | | | Stress response / proteostasis | 1.55 |
| | | | Connective tissue development | 1.54 |
| | | | Response to hormone / steroid | 1.67 |
| **MP7** | **Collagen** | **NMF21** | Collagen-receptor signaling | 1.66 |
| | **ECM** | | Basement membrane collagens | 1.64 |

| | | | | |
|---|---|---|---|---|
| | **Fibroblast/CAF** | | Fibrillar collagens (massive collagen production) | 1.73 |
| | **Neural crest-like** | | Collagen-containing ECM (full matrisome) | 1.89 |
| | | | Fibroblast/CAF markers | FALSE |
| | | | Neural crest-like markers | FALSE |
| | | **NMF11** | Small leucine-rich proteoglycans (SLRPs) | 1.85 |
| | | | Fibrillar/network collagens (extensive) | 2.03 |
| | | | Collagen-containing ECM (matrisome) | 1.85 |
| | | | Fibroblast / CAF markers | FALSE |
| | | | Endothelial / vascular markers | FALSE |
| | | | Neural crest-like markers | FALSE |
| **MP8** | **Cytoskeleton remodeling** | **NMF4** | Cell projection / neural morphogenesis | 1.75 |
| | **Developmental TFs** | | Cytoskeleton remodeling | 1.79 |
| | **Cell cycle** | | Developmental transcription factors | 1.77 |
| | | | Mitochondrial / OXPHOS | FALSE |
| | | | VEGFA / angiogenesis | FALSE |
| | | | Glycolysis / Warburg | FALSE |
| | | | MYC / cell cycle / proliferation | FALSE |
| | | | Cell cycle inhibitors / regulators | FALSE |
| | | | Stress / AP-1 TFs | FALSE |

| | | | | |
|---|---|---|---|---|
| **MP9** | **Lysosome biology** | **NMF18** | Vesicle organization | 1.52 |
| | **Mitochondrial** | | Mitochondrial outer membrane / contact sites | 1.67 |
| | **Cytoskeleton** | | ECM / collagen fibril organization | 1.62 |
| | **OXPHOS** | | Mitochondrial cation transport | 1.54 |
| | **Cell adhesion (NMF18)** | | Endosome / lysosome biology | 1.56 |
| | **Cell cycle (NMF19)** | | Proton-transporting ATPase complex (V-ATPase / F1F0) | 1.63 |
| | **Hypoxia (NMF19)** | | Cytoskeleton / branched actin | FALSE |
| | | | Cell-substrate junction / adhesion | 1.5 |
| | | | OXPHOS / ETC components | FALSE |
| | | **NMF19** | Negative regulation of cell death | 1.58 |
| | | | Pigmentation / melanocyte identity | FALSE |
| | | | Hormone regulation | 1.7 |
| | | | Glycolysis / Warburg metabolism | 1.6 |
| | | | Protein catabolism / ubiquitin | 1.61 |
| | | | Intracellular transport / autophagy | 1.57 |
| | | | Amino acid transport | 1.57 |
| | | | Cytoskeleton / contraction | 1.56 |

| | | | | |
|---|---|---|---|---|
| | | | Mitotic kinases | FALSE |
| | | | Endosomal / lysosomal trafficking | 1.6 |
| | | | OXPHOS / ETC | FALSE |
| | | | Hypoxia / HIF response | FALSE |
| | | | BCL2 / survival | FALSE |
| | | | Mitochondrial protein import / chaperones | FALSE |
| | | | Cell cycle | FALSE |
| | | | ROS / antioxidant | FALSE |
| **MP10** | **Lysosomal lipid metablism (NMF16)** | **NMF16** | Vesicle / endocytosis | 1.51 |
| | **p53 pathway (NMF16)** | | MDM2 / p53 / anti-apoptotic | 1.63 |
| | **protein modification (NMF16)** | | Protein ubiquitination / SUMO conjugation | 1.68 |
| | **Cell cycle (NMF22)** | | Melanocytic marker | FALSE |
| | **Survival (NMF22)** | | Vacuolar / lysosomal | 1.56 |
| | | **NMF22** | Histones (proliferating cells) | FALSE |
| | | | Organelle assembly | 1.64 |
| | | | Positive regulation of proliferation | 1.55 |
| | | | Cell cycle (G1/S / 12q amplicon) | FALSE |
| | | | Anti-apoptotic / survival | FALSE |
| | | | Mitochondrial / OXPHOS | FALSE |
| | | | Pigmentation / melanocyte identity | 1.52 |
| | | | Nucleotide / nucleic acid metabolism | FALSE |
| | | | DNA replication / S phase | FALSE |

| | | | | |
|---|---|---|---|---|
| | | | Mitosis / G2-M progression (massive proliferation signal) | FALSE |
| **MP11** | **Anti-apoptotic (NMF15)** | **NMF15** | Negative regulation of apoptotic signaling | 1.66 |
| | **Mitochondrial** | | Mitochondrial membrane potential / OXPHOS | 1.67 |
| | **Lysosomal** | | Sterol / cholesterol transport | 1.66 |
| | **Tissue remodeling (NMF15)** | | Antigen receptor signaling | 1.62 |
| | **Biosynthesis (NMF10)** | | Tissue remodeling | 1.69 |
| | | | Endo-lysosomal / vacuolar membrane biology | 1.73 |
| | | **NMF10** | Microtubule / kinesin | 2 |
| | | | Lysosomal / phagocytic enzymes | 3 |
| | | | Connective tissue / collagen development | 7 |
| | | | Glycosphingolipid biosynthesis | 4 |
| | | | Mitochondrial / OXPHOS | 3 |
| | | | Retinoic acid / nuclear receptor signaling | 3 |
| | | | Vesicle trafficking / membrane fusion | 10 |
| **MP12** | **Type I IFN signaling** | **NMF17** | Tissue remodeling / matricellular signaling | 1.64 |
| | **Proteostasis** | | Matricellular / secreted factors | 1.76 |
| | **Cell adhesion** | | Protease inhibitors | 1.62 |
| | | | Plasma cell / B cell markers | FALSE |
| | | | DNA damage | 1.72 |
| | | | Cell adhesion molecules | 1.65 |
| | | | Immune cell migration / inflammatory | NA |
| | | | Antigen receptor signaling / B cell | 1.72 |
| | | | Type I IFN / STAT1-STAT2 signaling | FALSE |
| | | **NMF12** | Inflammatory response (intrinsic) | 1.63 |
| | | | Protease inhibitors | 1.58 |

| | |
|---|---|
| Apoptosis / intrinsic mitochondrial pathway | 1.69 |
| Insulin / IGF1 signaling response | 1.52 |
| Cellular component disassembly | 1.61 |
| Cytoskeleton / intermediate filaments | 1.52 |
| Transmembrane protein / vesicle trafficking | FALSE |
| Heat shock / proteostasis | FALSE |
| NRP2 / semaphorin signaling | FALSE |
| Antigen presentation | FALSE |
| Translation / RNA-binding | FALSE |
| Endoplasmic reticulum / proteostasis | 1.51 |
| OXPHOS / mitochondrial | FALSE |
| STING / cGAS-STING innate sensing | FALSE |
| Cell adhesion / integrins | FALSE |
| Wnt / BMP regulators | FALSE |
| Type I IFN / antiviral response (intrinsic) | 1.6 |

| Genes | Median gene rank |
|---|---|
| SERPINF1; IGFBP7; TRIM63 | 10 |
| SERPINF1; IGFBP7; MMP2; TFRC; TNC | 13 |
| SLC1A4; SLC7A5; MFSD12; SLC25A4 | 15 |
| SERPINF1/PEDF; IGFBP7; CCN5; RNASE1; BIRC7; RACK1; ANXA2 | 15 |
| SERPINF1; PMEL; DCT; SLC1A4; MFSD12; TFRC; ANXA6; SLC45A2; SYTL2; ANXA2; OCA2; WIPI1; GLMP; MITF | 23 |
| SLC7A5; WIPI1 | 25 |
| PMEL; DCT; MFSD12; SLC45A2; OCA2; MITF; RAB32; EDNRB; RACK1; MYC | 25 |
| ITPKB; IGFBP7; EPHX1; ASAH1; EDNRB; UGCG; PRKD3; PLP1; CRYL1 | 25 |
| TRPM1; TFRC; ANXA6; EDNRB; KCNAB2; CTNNB1; PDE4D | 25 |
| UQCRC1; UQCRQ; UQCRH (Complex III); SDHB (Complex II); ATP5MC3; ATP5PD; ATP6AP1; TOMM7; SLC25A4 (ANT1); SLC25A16; MI | 75 |
| CDK1; CDK4; CDC25B; MYBL2; PTTG1; CENPF; UBE2T; ZWINT; DHFR; POLD2; NME1 | 97 |
| CRYAB; GDF15; FN1; GBP1 | 3 |
| GDF15; CCN3 | 5 |
| CRYAB; CCND1; RACK1 | 10 |
| PLA1A; SPP1; ANXA2 | 15 |
| CRYAB; GDF15; FN1; CCN3; RACK1; SPP1; TNFRSF12A; IGFBP7; TNC; SPOCK1; CDKN1A | 15 |
| FN1; GBP1; ANXA2; TNFRSF12A; SPOCK1; CCN3; AKAP12; SPP1 | 16 |
| GBP1; GBP2; GBP5; CCN3; IDO1; TNC; IER3; CCL5; CEBPB | 21 |
| GBP1; GBP2; EGR1; CCL5; CEBPB; CITED1 | 22 |
| GBP1; GBP2; GBP5; CIITA; CCL5; STXBP1 | 30 |
| CCND1; RHOB; NR4A1; IER3; CDKN1A | 32 |
| BIRC7; IER3; CRYAB | 34 |
| CCL5; CXCL9; CXCL10; CXCL13 | 49 |
| ISG15; IFI6; IFITM2; OAS1; OAS2; RTP4; EPSTI1; UBD | 63 |
| NDRG1; BHLHE40; BHLHE41; SOD2; SLC16A3 | 80 |
| NRAS; MDM2; CDK4; TSPAN31; METTL1; TSFM; LMBRD2; CTDSP2 | 6.5 |
| ABHD5; GDF15 | 7.5 |
| MDM2; CDK4; METTL1; SKP2 | 8 |

| | |
|---|---|
| GDF15; VGF | 14.5 |
| CAPG; SCIN; PFN2; CAVIN1 | 36 |
| MITF; BHLHE40; BHLHE41; TFAP2A | 36 |
| MDM2; ZFYVE16; ATP6AP1; MYO1D; CDKN2A; WIPI1; IER3 | 38 |
| CDK4; CDK1; SKP2; CCND1; POLD2; CDKN2A | 50.5 |
| S100A9; S100A8 | 4.5 |
| TYRP1; UGCG; QPCT; MFSD12; RAB32; TRPM1 | 9 |
| MDM2; NRAS; CDK4; TSPAN31; TSFM; LMBRD2; METTL1 | 14 |
| PRKD3; CDK4; CDK1; SGK1 | 15.5 |
| MDM2; PERP; SGK1 | 18 |
| UGCG; STXBP1; SLC25A4; CDK1; ANXA2; CLU | 22.5 |
| KRT5; KRT16; KRT6A; KRT6B; KRT17; KRT6C; DSP | 25 |
| S100A9; S100A8; SLC25A4; WIPI1; CISD1; CLU | 26 |
| NRAS; CDK4; MDM2; CTDSP2; LMBRD2; TSPAN31; METTL1 | 5 |
| CDK4; MDM2; CTDSP2; SKP2; IER3; CCND1 | 15 |
| APCDD1; CDH3; UGCG; COL6A1 | 18.5 |
| SLC1A5; SLC1A3; SLC7A11; MFSD12 | 25 |
| ADGRG1; COL1A2; COL3A1; ITGA3 | 12.5 |
| COL1A2; COL3A1; COL1A1; COL18A1; COL4A2; COL6A2 | 19 |
| S100A4; ECM1; COL1A2; TIMP3; L1CAM; COL3A1; COL1A1; FN1; CSTB; BCAN; MMP9; COL18A1; TIMP2; CTSH | 20.5 |
| DCT; PMEL; CAPG; MFSD12; TFRC; ITGB3; ANXA2 | 21 |
| DCT; PMEL; MFSD12; MITF; TFAP2A | 21 |
| IFI6; IFI27; IFI16; OAS2; MX1-like | 40 |
| ITGA3; ITGB3; ITGB5; ITGB8; ITGA6 | 67 |
| THBS2; FLOT1 | 3.5 |
| FXYD3; SLC20A1; SLC5A3; CNIH3; SCN1B | 9 |
| SLC5A3; CDKN1A | 11 |
| SERPINF1; TNC; IGFBP7 | 15 |
| CCND1; SERPINF1; IGFBP7; SGK1 | 16.5 |
| CCND1; CDKN1A; SGK1; BRSK1 | 20.5 |

| | |
|---|---|
| CCND1; CDKN1A; BIRC7; PERP; SESN3 | 22 |
| FLOT1; CCND1; BIRC7; SNX10; BRSK1 | 23 |
| THBS2; SERPINA3; LOXL4; SERPINF1; TNC; IGFBP7; SEMA3B; ECM1; PLAT; PCSK6; PTPRZ1; TFPI2; MGP; MMP9; SDC2 | 26 |
| SERPINA3; CTSK; SERPINF1; CD109; PLAT; PCSK6; TFPI2; MMP9 | 28 |
| CCND1; CDKN1A; TNC; INPP5F; PRRX1; DUSP10; SPP1 | 40 |
| CTSK; ADGRG1; MMP17 | 15 |
| SLC39A6; SGK1; TRPM1; KCNAB2 | 22 |
| PMEL; QPCT; GLMP; RAB32; TYRP1; TRPM1; SYTL2; VAMP8; SLC45A2 | 24 |
| PMEL; TYRP1; RAB32; SLC45A2; MITF; OCA2 | 51.5 |
| L1CAM; CADM1; IGSF8; IGSF3; ADGRG6 | 52 |
| TYRP1; PMEL; UGCG; QPCT; MFSD12-like | 3.5 |
| STXBP1; PLPPR4 | 9 |
| TIMP2; IFI16; CDKN1A | 12 |
| PRKD3; SGK1 | 14 |
| MCOLN3; SGK1; VAMP8 | 21 |
| QPCT; TIMP2; SPTAN1; LYZ; VAMP8; CHIT1; CTSH; ASAH1 | 21 |
| SLC7A11; GDF15; NQO1 | 22 |
| IFI16; SGK1; CDKN1A; CCND1 | 22.5 |
| LYZ; IGKC; CCL18; CD55; IFI30 | 26 |
| IFI16; VAMP8; CD55; IFI27; TFRC; SLC1A5; EPHA2 | 33 |
| CDKN1A; NQO1; SHC1; PDK4; IFI6 | 35 |
| IFI16; IFI27; IFI6; STAT2; JAK1; IFNAR1 | 55 |
| CCL21; CLU; LYZ; IGHM; IGHG1; MS4A1; CCL19; C3; JCHAIN; C2 | 10 |
| IGKC; IGHM; IGHG1; MS4A1; CD79A; CD79B; IGHG3 | 10 |
| CCL21; CLU; LYZ; IGHG1; VCAM1; PLP1; FOS; CCL19; C3; CXCR4 | 16 |
| CCL21; VCAM1; CORO1A; CCL19; SELL; RIPOR2; CXCR4 | 22 |
| SPI1; FCER1G; ITGB2; C3 | 10.5 |
| CD4; LYZ; CTSS; ITGAX; TREM2; CD300A; LAPTM5; CD52 | 14 |
| LYZ; CTSS; LAPTM5; LIPA; FUCA1; MPEG1; C3; CXCR4; IFI30; ADA2; CTSH | 16 |
| LCP1; LSP1; CAPG; CORO1A; CXCR4 | 17 |

| | |
|---|---|
| CTSS; FCER1G; MPEG1; IFI30; FGL2; CTSH | 18.5 |
| IGHG1; IGHG3; C3; IGHA1 | 20.5 |
| IGKC; IGHG1; LAPTM5; LIPA; TRAC; IGHG3; TRBC2; IGHA1; IGHM; IGLC1; NFKBIA; TRBC1 | 23 |
| IGKC; IGHG1; IGHG3; IGHA1; IGHM; IGLC1; JCHAIN | 28 |
| CD4; TRAC; TRBC1; TRBC2; IL2RG; ZAP70 | 40 |
| CCL2; CCL5; CCL18; CXCL12; IL32 | 49 |
| ISG15; IFI6; OAS1; OAS2; MX1; CMPK2; IFIT1; IFIT2; EPSTI1; RTP4-like | 54 |
| CXCL9; CXCL10; CCL5; STAT1; CIITA | 70 |
| EDNRB; SPP1; AVPR1A; MME | 12 |
| SPP1; POSTN; IGFBP7; AVPR1A; TBX2; SGK1 | 17.5 |
| MGP; MMP2; GDF15; POSTN; IGFBP7; COL16A1; S100A4; FN1; PLAT; PTPRZ1; COL18A1; COL1A1 | 19.5 |
| IGFBP7; IGFBP2; GDF15; CLEC11A | 23.5 |
| SPP1; NBL1; POSTN; MET; SGK1; FN1; PTPRZ1; COL18A1; SEMA6A; METRN | 26 |
| RACK1; PRAME; NOP53 | 7 |
| MYC; NOP53; MXI1; RACK1; ID4 | 12 |
| RACK1; ERBB3; DCLK1; STK32A; GRK3 | 13 |
| MYC; TFAP2A; RHOB; MMP2; BCL2; SFRP1; ERBB3 | 20 |
| RACK1; PPIA | 42.5 |
| DCT; PRAME; TFAP2A; MITF; PAX3; SOX2 | 52.5 |
| NDRG1; NDRG2 | 64 |
| TMOD1; RHOB; RND3; CDC42EP1 | 78 |
| SFRP1; NDRG1 | 1.5 |
| NDRG1; CRYAB | 5.5 |
| SFRP1; SPP1; CRYAB; RACK1 | 8 |
| SFRP1; IGFBP2; EDNRB | 12 |
| CRYAB; HSPA6; SOD3; SELENOP | 20.5 |
| MGP; SLC26A2; LPL; CSGALNACT1 | 22.5 |
| SFRP1; SPP1; IGFBP2; EDNRB; CPN1; ABHD2; LRP2; UGCG | 25 |
| COL1A1; COL4A1; COL4A2 | 12 |
| COL4A1; COL4A2; COL15A1; COL18A1 | 16.5 |

| | |
|---|---|
| COL1A1; COL1A2; COL3A1; COL5A1; COL5A2; COL5A3; COL6A1; COL6A2; COL6A3; COL12A1; COL15A1; COL18A1; COL7A1; COL8/ | 24 |
| COL1A1; COL3A1; COL1A2; MGP; THBS2; CCN2; ADAMTS1; COL6A1; COL4A2; COL4A1; COL6A2; S100A4; COL18A1; COL6A3; FN1; | 25 |
| S100A4 (FSP1); THY1; PDGFRB-like (low); FAP-like (low) | 53 |
| TNC; VCAN; GPC1; IGFBP3; CADM1; NOTCH3; FOXD3; NGFR; MRC2; ECE1; MAP1B; SERPINE1; GEM; ANGPTL2 | 72.5 |
| LUM; DCN; BGN; VCAN | 9 |
| COL3A1; COL1A1; COL1A2; COL6A3; COL6A2; COL6A1; COL5A1; COL4A1; COL5A2; COL4A2; COL15A1; COL12A1; COL16A1; COL1 | 11 |
| COL3A1; COL1A1; COL1A2; COL6A3; COL6A2; LUM; COL6A1; DCN; COL5A1; VCAN; COL4A1; FN1; BGN; MMP2; COL5A2; CXCL12; C | 25 |
| FAP; S100A4; PDGFRB; PDGFRA; THY1 | 25 |
| PECAM1; VWF; CD93; PLVAP; EGFL7; CALCRL; PODXL | 72 |
| VCAN; MRC2; NOTCH3; ANGPTL2; TNC; SERPINE1; ECE1; SDC1; MAP1B | 47 |
| BMPR2; B4GALT5; FLOT1; TUBB2B; GSN; PFN2; GLI3; PLXNA1; SHOC2; VEGFA; CDC42-like | 10.5 |
| GSN; PFN2; TUBB2B; ARPC5 | 10.5 |
| BMPR2; GLI3; TGIF1; HES1; SHOC2; SDC3; GAB2 | 19.5 |
| UQCRC1; SLC25A4; SLC25A37; ATP5MC2; ATP5PD; TSFM; SFXN3 | 39 |
| VEGFA; FGFR1 | 40 |
| HK1; HK2; ENO1; PDP1; HILPDA | 45 |
| CCND1; MYC; CDK4; CDC25B; CDCA4; E2F1; TLK1 | 69.5 |
| CDKN1C; CIZ1; CDCA4 | 70 |
| EGR1; NR4A1; FOS; CREM; CEBPB | 91 |

| | |
|---|---|
| ANXA2; ATP6AP1; ATP6V1F | 4 |
| TOMM7; MICOS10; SELENON; SFXN3; IARS2 | 6 |
| ANXA2; SELENON; COL6A2 | 10 |
| ATP6AP1; ATP5MC2; SELENON; SLC39A7; CTSS; ATP5PD; ATP6V1F; ATP5MC3; ITGAV | 16 |
| ANXA2; ATP6AP1; KDELR2; CTSS; LAPTM5 | 16 |
| ATP6AP1; ATP5MC2; ATP5PD; ATP6V1F; ATP5MC3 | 22 |
| ARPC5; ARPC1A; VCL | 28 |
| ARPC5; ADAM10; VCL; HSPA5; ITGAV; HSPG2 | 30.5 |
| UQCRQ; UQCRH; SDHB; ATP5MC2; ATP5MC3; NDUFA3 | 37 |
| EDNRB; ENO1; PRAME; MITF; SLC7A5; TFAP2A; STXBP1 | 6 |
| MITF; TFAP2A; PRAME; EDNRB; MC1R | 6 |
| EDNRB; SLC7A5; SLC16A1 | 8 |
| ENO1; PGD; SLC2A1; SLC16A1; PMVK | 15 |
| PRAME; FBXW5; USP33; NOP53; HGS | 19 |
| KIF1B; ZFYVE16; KPNA6; STXBP1; HSBP1; HGS; PTPN14 | 21 |
| SLC7A5 (LAT1); STXBP1; SLC7A1 | 21 |
| EDNRB; ENO1; TRIM63; HSBP1; ANXA6; PDE4D | 25.5 |

| | |
|---|---|
| CDC25B; GRK6; CDC42EP3 | 26 |
| ZFYVE16; ANXA6; PLEKHM2; HGS | 30.5 |
| SDHB; ATP5PD; ATP5MC2; UQCRH; UQCRQ; ACADM | 52.5 |
| SLC2A1; HILPDA; SLC16A3; SLC16A1 | 56 |
| BCL2 | 66 |
| BAG1; IPO5; KPNA6; AURKAIP1 | 70.5 |
| CDC25B; POLD2; TSPAN31; NRAS | 74.5 |
| MT1E; NCF2 | 89.5 |

| | |
|---|---|
| MDM2; IGF2R; INPP5F; STAU2 | 12.5 |
| MDM2; KDM1A; BAX | 13 |
| MDM2; KDM1A; ARRDC3; UBA2 | 14.5 |
| SLC45A2 | 18 |
| SDC3; ASAH1; ARRDC3; ARL8A; NCSTN; LIPA; VLDLR; SPPL2B | 38.5 |
| HIST1H1B; HIST1H4C; HIST1H1E; HIST1H1D; H2AFJ | 30 |
| CENPF; SNX10; HPS4; ANLN; BIRC5; RAB32; MYBL2; CDC20 | 31 |
| NME1; EDNRB; ADGRG1; CDK4; E2F1; BIRC5; NRAS; TFRC; CDC20; ZNF703 | 36 |
| CDK4; NRAS; TSFM | 39 |
| BIRC5; BAX; ICMT | 47 |
| TOMM7; MICOS10; UQCRH; UQCRQ; SDHB; ATP5MC2; ATP5MC3; ATP5PD; NDUFA-like (low) | 47.5 |
| PAX3; MITF; RAB32; PMEL; SLC45A2; ALDH1A3 | 50.5 |
| AHCY; UBE2T; DHFR; RRM2 | 53 |
| TK1; DHFR; POLD2; POLE3; RRM2; UBE2T | 53 |

| | |
|---|---|
| TOP2A; CENPF; CDK1; CDK4; MYBL2; ZWINT; UBE2T; DHFR; BIRC5; CDC20; PTTG1; CENPH; NCAPG; NCAPG2; KIF18B; PBK; MEL[illegible] | 56 |
| TMBIM1; SOD2; IFI6; BAX | 11 |
| SOD2; IFI6; LIPA; BAX | 15 |
| ANXA2; NFKBIA; LIPA; STARD3NL; VPS4B | 17 |
| SLC39A6; NFKBIA; LIPA; BAX; LAPTM5 | 17 |
| TMBIM1; LIPA; CTSK; BAX; SNX10 | 18 |
| ITM2C; ASAH1; ANXA2; TMBIM1; NCSTN; ARL8A; LIPA; CTSK; VAC14; ECE1; STARD3NL; VPS13C; LAPTM5; ATP6AP1; SNX10; TM9[illegible] | 20 |
| KIFC3; SPG7 | 8 |
| CTSK; CTSS; STX3 | 47.3 |
| COL1A2; CHST11; CTSK; RARA; PPARD; NAMPT; SLC39A13 | 17.7 |
| CHST11; TMEM165; PINK1; ST6GAL-like (low) | 16.3 |
| NDUFA3; PINK1; HK1 | 25.3 |
| RARA; PPARD; HES1 | 60.3 |
| STX3; VPS41; NUDCD3; NBR1; NAPG; TRAPPC6A; PLEKHM2; HGS; VPS4B; AMFR | 32.3 |

| | |
|---|---|
| SFRP1; SPP1 | 2 |
| SFRP1; SPP1; TIMP2; RNASE1; SPARCL1; TIMP3; SELENOP; SRGN | 11.5 |
| TIMP2; TIMP3 | 13.5 |
| IGKC; IGHG1; IGHA1 | 14 |
| SFRP1; IFI16; CRYAB; DCT | 15 |
| HMCN1; SPARCL1; PCDH7 | 17 |
| IGKC; IGHG1; STAT1; LYZ | 17.5 |
| IGKC; IGHG1; TNFRSF21; NFATC2 | 18 |
| IFI16; STAT1; STAT2; IFI30; IFNAR1; IFI27; IFI6 (low) | 31.5 |
| IFI16; PSMB4; PLD3; SOD1 | 8 |
| TIMP2; TIMP3; SRGN | 13 |

| | |
|---|---|
| IFI16; IVNS1ABP; HYOU1; SOD1 | 13.5 |
| NCL; SHC1; STAT1 | 19 |
| ASPH; TIMP3; SPTAN1; KLC1 | 23 |
| PLEC; SPTAN1; KLC1; HTT | 28 |
| TM9SF2; TM7SF3; TMEM147; TMEM59; TMEM106C; TMEM14C; TMED7; TMEM167A; TMEM127; TMEM173 | 30 |
| HSPH1; HYOU1; POMP; KPNB1 | 36.5 |
| NRP2 | 37 |
| IFI30; CIITA-like (none); PSMB4 | 37.5 |
| EIF4H; EIF4G1; EIF3L; EIF3M; SRSF4; CSDE1; HNRNPU; HNRNPLL; HNRNPH2 | 54 |
| HYOU1; ASPH; SRP54; SRP-like; SPCS3; SRPRB; SRPRA; FKBP10 | 67 |
| ATP5F1A; ATP5MC1; ATP5PD; ATP6V0B; ATP6V1H; TIMM50; TIMM8B; TIMM17A; TIMM44; TIMMDC1 | 78 |
| TMEM173 (STING) | 86 |
| ITGB3; ITGAV; ITGA7 | 90 |
| BAMBI; ARNT2 | 98 |
| IFI16; STAT1; STAT2; IFI30; IFNAR1; IFI44; IFIT3; IFIH1; JAK1; IRAK1; IFI27; IFRD1; STAT3 | 100 |

**Genes list used as negative control for spatial restriction**

Negative control

RNA processing signature from Karras et al

| Gene |
|---|
| LUC7L2 |
| SON |
| FUS |
| DDX17 |
| PNISR |
| TRA2A |
| SNRNP70 |
| RBM25 |
| CPSF6 |
