## Supplementary Table 6 for "Melanoma evolution in the lymph node shapes systemic outcomes"

| gene | log2FoldChange | baseMean | lfcSE | stat | pvalue | padj |
|---|---|---|---|---|---|---|
| CD22 | -2.414464002 | 368.1006908 | 0.415323595 | -5.813452528 | 6.12E-09 | 8.86E-05 |
| CXCR3 | -1.69150108 | 87.45245793 | 0.328480906 | -5.149465463 | 2.61E-07 | 0.001260521 |
| GPR174 | -1.669161583 | 114.5066411 | 0.320849124 | -5.202325506 | 1.97E-07 | 0.001260521 |
| ELMO1 | -1.334734204 | 254.8037203 | 0.268981211 | -4.962183781 | 6.97E-07 | 0.001441499 |
| PTK2B | -0.985331603 | 666.23652 | 0.198168623 | -4.972187775 | 6.62E-07 | 0.001441499 |
| ADGRG5 | -1.859980511 | 97.38628802 | 0.366878428 | -5.069746187 | 3.98E-07 | 0.001441499 |
| VPREB3 | -2.645694527 | 119.3998849 | 0.532900349 | -4.964707817 | 6.88E-07 | 0.001441499 |
| SP140 | -1.824738834 | 363.3760043 | 0.369849104 | -4.933738686 | 8.07E-07 | 0.00145973 |
| TBC1D1 | -1.505755773 | 221.387526 | 0.30692586 | -4.905926702 | 9.30E-07 | 0.001495648 |
| FDCSP | -2.863299419 | 663.199666 | 0.596106787 | -4.80333303 | 1.56E-06 | 0.001737632 |
| GAPT | -1.202083761 | 105.4276062 | 0.248396179 | -4.839381051 | 1.30E-06 | 0.001737632 |
| CARD11 | -1.763832919 | 274.6127306 | 0.36543411 | -4.826678388 | 1.39E-06 | 0.001737632 |
| CD37 | -2.016382205 | 1405.417529 | 0.418902014 | -4.813493699 | 1.48E-06 | 0.001737632 |
| TAGAP | -1.440449641 | 337.5261694 | 0.304337412 | -4.733067922 | 2.21E-06 | 0.002286706 |
| CNR2 | -2.141572353 | 59.93108629 | 0.457038426 | -4.685759955 | 2.79E-06 | 0.002612617 |
| SIDT1 | -1.960070901 | 53.10820854 | 0.418939213 | -4.67865227 | 2.89E-06 | 0.002612617 |
| CFI | -1.940590839 | 216.7483194 | 0.418213232 | -4.640194733 | 3.48E-06 | 0.002844532 |
| IGHD | -2.837403812 | 347.9914187 | 0.611920296 | -4.636884627 | 3.54E-06 | 0.002844532 |
| BLNK | -1.674219345 | 206.2596666 | 0.363698763 | -4.603313276 | 4.16E-06 | 0.003168126 |
| CD19 | -2.831018914 | 167.5736034 | 0.618776076 | -4.575191294 | 4.76E-06 | 0.003279745 |
| PIK3CG | -1.446639289 | 219.1953789 | 0.317727149 | -4.553086798 | 5.29E-06 | 0.003327242 |
| IDO1 | -2.854587301 | 457.3911308 | 0.626531514 | -4.556175125 | 5.21E-06 | 0.003327242 |
| FCRL1 | -2.784614562 | 138.2648913 | 0.631285574 | -4.411022013 | 1.03E-05 | 0.003461404 |
| BCL11A | -2.225792846 | 330.2759248 | 0.503205727 | -4.423226376 | 9.72E-06 | 0.003461404 |
| PLCL2 | -1.121304619 | 149.5943917 | 0.252337812 | -4.443664662 | 8.84E-06 | 0.003461404 |
| DAPP1 | -1.579973666 | 290.2117353 | 0.357611407 | -4.418129947 | 9.96E-06 | 0.003461404 |
| CCR6 | -2.181414444 | 135.9623765 | 0.485668425 | -4.491571479 | 7.07E-06 | 0.003461404 |
| MYO1G | -1.310983927 | 355.4431228 | 0.292243583 | -4.485928871 | 7.26E-06 | 0.003461404 |
| DENND3 | -1.184499413 | 304.3005528 | 0.268363071 | -4.41379438 | 1.02E-05 | 0.003461404 |
| CD72 | -1.65395683 | 365.9325479 | 0.373709129 | -4.425786526 | 9.61E-06 | 0.003461404 |
| SEMA4D | -1.365812044 | 333.2942211 | 0.30873866 | -4.423845211 | 9.70E-06 | 0.003461404 |
| SUSD3 | -1.545901687 | 125.6734554 | 0.345740134 | -4.471282138 | 7.78E-06 | 0.003461404 |
| NPFFR1 | -1.916609688 | 41.93592413 | 0.427962073 | -4.478456877 | 7.52E-06 | 0.003461404 |
| PLD4 | -1.430507643 | 231.6672003 | 0.31984377 | -4.472519951 | 7.73E-06 | 0.003461404 |
| P2RX5 | -2.295466369 | 214.5148157 | 0.518493979 | -4.427180374 | 9.55E-06 | 0.003461404 |
| TNFRSF13B | -2.415784085 | 161.2135129 | 0.538156973 | -4.48899523 | 7.16E-06 | 0.003461404 |
| EVI2B | -1.226337644 | 725.9633014 | 0.277332652 | -4.421901399 | 9.78E-06 | 0.003461404 |
| CLEC17A | -2.712472694 | 65.35579426 | 0.615606505 | -4.40617939 | 1.05E-05 | 0.003461404 |
| CD79A | -2.435394644 | 728.0500428 | 0.548878777 | -4.43703555 | 9.12E-06 | 0.003461404 |
| FLT3LG | -1.528032885 | 90.02810398 | 0.345088209 | -4.4279487 | 9.51E-06 | 0.003461404 |
| RAC2 | -1.478691863 | 715.298346 | 0.329407238 | -4.488947702 | 7.16E-06 | 0.003461404 |
| TNFRSF13C | -2.122998714 | 282.0071501 | 0.474393181 | -4.475188092 | 7.63E-06 | 0.003461404 |
| SASH3 | -1.684600227 | 564.7581153 | 0.376135226 | -4.478709009 | 7.51E-06 | 0.003461404 |
| IL22RA2 | -2.59763459 | 55.04781099 | 0.591148795 | -4.39421447 | 1.11E-05 | 0.003576345 |
| RIC3 | -1.545569329 | 123.7743234 | 0.35227033 | -4.387452473 | 1.15E-05 | 0.00360912 |
| CXCL13 | -2.579293299 | 801.2643845 | 0.590416359 | -4.368600663 | 1.25E-05 | 0.003699866 |
| UBD | -2.361352501 | 541.3522218 | 0.539793367 | -4.374548941 | 1.22E-05 | 0.003699866 |
| BLK | -2.534958862 | 329.650796 | 0.580899611 | -4.363850167 | 1.28E-05 | 0.003699866 |
| PCED1B | -1.283328769 | 214.4692722 | 0.293832843 | -4.367547063 | 1.26E-05 | 0.003699866 |
| CARMIL2 | -1.84739741 | 208.8538941 | 0.427257646 | -4.323848679 | 1.53E-05 | 0.004352181 |
| TRANK1 | -1.111706314 | 254.2865102 | 0.258170164 | -4.306099111 | 1.66E-05 | 0.004625601 |
| ADAM28 | -1.800250676 | 407.8453374 | 0.419984403 | -4.286470313 | 1.82E-05 | 0.004692666 |
| MS4A1 | -2.795943026 | 1172.595675 | 0.652138881 | -4.287342936 | 1.81E-05 | 0.004692666 |
| SNX20 | -1.559457402 | 366.9419654 | 0.363216257 | -4.293468069 | 1.76E-05 | 0.004692666 |
| RASAL3 | -1.604825099 | 244.5107848 | 0.373379127 | -4.298111449 | 1.72E-05 | 0.004692666 |

| | | | | | | |
|---|---|---|---|---|---|---|
| FCRL5 | -1.704282252 | 231.910223 | 0.399921937 | -4.261537299 | 2.03E-05 | 0.004906507 |
| FAM49A | -1.246866766 | 244.806658 | 0.292259083 | -4.266306301 | 1.99E-05 | 0.004906507 |
| CCDC69 | -1.350647343 | 275.6835474 | 0.316245504 | -4.270882361 | 1.95E-05 | 0.004906507 |
| NTS | -3.333529224 | 193.0122352 | 0.782304747 | -4.26116451 | 2.03E-05 | 0.004906507 |
| SHISAL2A | -1.490079119 | 42.0897352 | 0.350833038 | -4.247259973 | 2.16E-05 | 0.005007853 |
| WDFY4 | -1.993737045 | 410.8903231 | 0.469593273 | -4.245667812 | 2.18E-05 | 0.005007853 |
| PTPN6 | -1.523431666 | 529.4807824 | 0.358261559 | -4.25228894 | 2.12E-05 | 0.005007853 |
| CD53 | -1.469995529 | 1201.406722 | 0.346789884 | -4.238865074 | 2.25E-05 | 0.005081361 |
| TLR10 | -2.279176948 | 202.2138671 | 0.53853004 | -4.232218782 | 2.31E-05 | 0.005153393 |
| SH2D1A | -1.657711731 | 134.347567 | 0.392716263 | -4.221143575 | 2.43E-05 | 0.005331249 |
| NUAK2 | -1.12161464 | 89.43526897 | 0.266058218 | -4.215673728 | 2.49E-05 | 0.005380612 |
| APBB1IP | -1.414789837 | 645.0685154 | 0.336334174 | -4.206500394 | 2.59E-05 | 0.005521224 |
| HBA2 | -3.179173575 | 125.184504 | 0.756614622 | -4.20183999 | 2.65E-05 | 0.005554467 |
| UNC13D | -1.266616406 | 208.6893608 | 0.301796272 | -4.196925292 | 2.71E-05 | 0.005595246 |
| PLAC8 | -2.110036472 | 210.7797765 | 0.504437078 | -4.182952771 | 2.88E-05 | 0.0058668 |
| SLAMF6 | -1.721357967 | 251.3845484 | 0.413656907 | -4.161318082 | 3.16E-05 | 0.006201069 |
| PNOC | -1.640800941 | 68.7307998 | 0.394410397 | -4.160136125 | 3.18E-05 | 0.006201069 |
| CCND2 | -1.273607034 | 380.3771424 | 0.306314889 | -4.157835874 | 3.21E-05 | 0.006201069 |
| CSF2RB | -2.253697591 | 392.5860876 | 0.541981443 | -4.158256009 | 3.21E-05 | 0.006201069 |
| FCHO1 | -1.622813844 | 185.6607127 | 0.393929549 | -4.119553481 | 3.80E-05 | 0.007136617 |
| BIN2 | -1.292955055 | 222.4924521 | 0.314271373 | -4.114135632 | 3.89E-05 | 0.007212628 |
| HBB | -2.696018946 | 34.00684415 | 0.657183418 | -4.102384315 | 4.09E-05 | 0.007492967 |
| LYN | -1.119076082 | 748.7916204 | 0.273092839 | -4.097786262 | 4.17E-05 | 0.007547789 |
| SIT1 | -1.503045972 | 88.79349606 | 0.367102304 | -4.094351783 | 4.23E-05 | 0.007565964 |
| XCR1 | -1.744849381 | 63.87039995 | 0.427091695 | -4.085421005 | 4.40E-05 | 0.007767074 |
| ATP2A3 | -1.72695821 | 439.2237855 | 0.423705925 | -4.075841543 | 4.58E-05 | 0.007865656 |
| ARHGAP27 | -1.133418382 | 284.0228704 | 0.278027922 | -4.076635084 | 4.57E-05 | 0.007865656 |
| POU2F2 | -1.487171116 | 174.782061 | 0.365027351 | -4.074136126 | 4.62E-05 | 0.007865656 |
| SIGLEC12 | -2.088371706 | 50.63576383 | 0.513958327 | -4.063309407 | 4.84E-05 | 0.0081439 |
| TNFAIP8 | -1.348653522 | 390.6751076 | 0.332720423 | -4.053413706 | 5.05E-05 | 0.00830324 |
| DOCK2 | -1.309187543 | 606.5150199 | 0.322777735 | -4.05600325 | 4.99E-05 | 0.00830324 |
| PCNX2 | -1.182142372 | 217.7869664 | 0.29440583 | -4.015349739 | 5.94E-05 | 0.009442441 |
| PLEK | -1.433106858 | 544.7066053 | 0.356798128 | -4.016576163 | 5.90E-05 | 0.009442441 |
| COCH | -2.012839636 | 59.94020997 | 0.501650088 | -4.012437524 | 6.01E-05 | 0.009455815 |
| ZC3H12D | -1.523276425 | 163.4632892 | 0.380381399 | -4.004602829 | 6.21E-05 | 0.00966962 |
| RAB37 | -1.756351807 | 91.49174175 | 0.439095137 | -3.999934546 | 6.34E-05 | 0.009757441 |
| DENND1C | -1.443830912 | 238.9402537 | 0.361994395 | -3.988544934 | 6.65E-05 | 0.010130124 |
| LAPTM5 | -1.435131314 | 2716.451855 | 0.360125619 | -3.985085306 | 6.75E-05 | 0.010171786 |
| POU2AF1 | -1.617172028 | 327.3112777 | 0.407000117 | -3.973394499 | 7.09E-05 | 0.010574278 |
| GPD1 | -2.040079998 | 25.57739486 | 0.513769773 | -3.970805809 | 7.16E-05 | 0.010580767 |
| COL4A3 | -1.811626207 | 39.89096692 | 0.458752474 | -3.949027655 | 7.85E-05 | 0.011409139 |
| SEPTIN1 | -1.380534064 | 508.5523719 | 0.349681329 | -3.947977628 | 7.88E-05 | 0.011409139 |
| TCL1A | -2.163684303 | 207.466023 | 0.548891167 | -3.941918602 | 8.08E-05 | 0.011585445 |
| DOCK8 | -1.079334953 | 479.5426691 | 0.274084845 | -3.937959263 | 8.22E-05 | 0.011662766 |
| TSHZ2 | -1.16051308 | 179.1167281 | 0.294952208 | -3.934580071 | 8.33E-05 | 0.011713232 |
| TIMD4 | -1.401716409 | 88.06178613 | 0.357494917 | -3.920940804 | 8.82E-05 | 0.011933085 |
| MAPK13 | -1.113001905 | 179.8675715 | 0.283530445 | -3.925511079 | 8.65E-05 | 0.011933085 |
| C12orf42 | -1.223640894 | 30.02841901 | 0.311893919 | -3.923259863 | 8.74E-05 | 0.011933085 |
| SIGLEC6 | -1.473741697 | 33.85224815 | 0.375766722 | -3.921959054 | 8.78E-05 | 0.011933085 |
| PATL2 | -1.395983166 | 95.69406522 | 0.356751882 | -3.913036586 | 9.11E-05 | 0.012104422 |
| WAS | -1.572741468 | 388.0334839 | 0.401903096 | -3.913235515 | 9.11E-05 | 0.012104422 |
| FCRL3 | -2.182809412 | 91.18731488 | 0.55929008 | -3.902821614 | 9.51E-05 | 0.012413259 |
| PSD4 | -1.452202506 | 310.6031693 | 0.372115974 | -3.902553529 | 9.52E-05 | 0.012413259 |
| PARP15 | -2.218565997 | 178.4347515 | 0.569092713 | -3.898426293 | 9.68E-05 | 0.012461537 |
| GPR18 | -1.428950598 | 66.85351028 | 0.366652352 | -3.897290141 | 9.73E-05 | 0.012461537 |
| TBC1D10C | -1.7140705 | 619.9698367 | 0.440821815 | -3.888352261 | 0.000100927 | 0.012815968 |

| | | | | | | |
|---|---|---|---|---|---|---|
| RHOH | -2.007091674 | 189.1655289 | 0.517305262 | -3.879898047 | 0.0001045 | 0.013139622 |
| GIMAP1 | -1.196052789 | 180.9788735 | 0.30841496 | -3.878063469 | 0.000105291 | 0.013139622 |
| VAV1 | -1.510913632 | 359.921374 | 0.389976996 | -3.874366048 | 0.000106903 | 0.013226687 |
| FCRL2 | -2.195111897 | 72.44773868 | 0.567670338 | -3.866877925 | 0.000110238 | 0.013523721 |
| TRBC1 | -1.53393499 | 734.6952963 | 0.397715362 | -3.85686633 | 0.00011485 | 0.013673668 |
| TOX | -1.398246697 | 73.22399875 | 0.362561207 | -3.856581094 | 0.000114984 | 0.013673668 |
| RASGEF1A | -1.348879434 | 43.23135976 | 0.349809408 | -3.856041041 | 0.000115238 | 0.013673668 |
| BTK | -1.444790742 | 277.5904104 | 0.374633946 | -3.856539851 | 0.000115003 | 0.013673668 |
| KIAA0040 | -1.573870588 | 193.7980014 | 0.408624515 | -3.851630359 | 0.000117334 | 0.013809167 |
| LTA | -1.134756091 | 43.75465611 | 0.295089841 | -3.845459695 | 0.000120327 | 0.014047165 |
| PARVG | -1.305068137 | 462.4876788 | 0.340134709 | -3.836915501 | 0.000124589 | 0.01442844 |
| CD24 | -1.742867723 | 177.4722229 | 0.454501087 | -3.834683292 | 0.000125726 | 0.014444542 |
| SYTL1 | -1.638329252 | 205.6833092 | 0.428266056 | -3.825494054 | 0.00013051 | 0.014539682 |
| SELL | -1.847127731 | 1273.371004 | 0.482959034 | -3.824605407 | 0.000130982 | 0.014539682 |
| EAF2 | -1.274586787 | 274.0210756 | 0.333687767 | -3.819698871 | 0.000133615 | 0.014539682 |
| IFIT2 | -1.505836683 | 354.8986316 | 0.393437491 | -3.827384821 | 0.000129512 | 0.014539682 |
| CXCR5 | -1.986546926 | 118.7966156 | 0.519766508 | -3.821998715 | 0.000132374 | 0.014539682 |
| CD27 | -1.49154092 | 257.7522966 | 0.390669948 | -3.817905442 | 0.000134589 | 0.014539682 |
| TMCC3 | -1.338989948 | 72.00331554 | 0.349890485 | -3.826882998 | 0.000129776 | 0.014539682 |
| SLC38A5 | -1.331744963 | 46.23334146 | 0.348740481 | -3.818727784 | 0.000134142 | 0.014539682 |
| DEF6 | -1.342851481 | 397.3028054 | 0.35192429 | -3.815739693 | 0.000135776 | 0.014559165 |
| RCSD1 | -1.497385797 | 481.374075 | 0.393585809 | -3.804471004 | 0.000142108 | 0.014721548 |
| TNFSF10 | -1.503131471 | 691.9786698 | 0.39514435 | -3.804005981 | 0.000142375 | 0.014721548 |
| GIMAP5 | -1.159001865 | 242.2864571 | 0.304117591 | -3.811031987 | 0.000138388 | 0.014721548 |
| CCDC88C | -1.3121377 | 276.8630856 | 0.34488241 | -3.804594444 | 0.000142037 | 0.014721548 |
| CCL5 | -1.660340716 | 827.6484754 | 0.435938512 | -3.808658034 | 0.000139723 | 0.014721548 |
| AIM2 | -1.986673467 | 224.9776337 | 0.523042536 | -3.798301914 | 0.000145691 | 0.014852253 |
| CD96 | -1.478326268 | 292.5198171 | 0.389141654 | -3.798941216 | 0.000145316 | 0.014852253 |
| CSTA | -1.221994125 | 255.2414999 | 0.321944157 | -3.795671072 | 0.000147245 | 0.014905686 |
| KBTBD8 | -1.299784175 | 71.2804922 | 0.342828871 | -3.791349811 | 0.000149831 | 0.015062152 |
| SLC15A2 | -1.173833668 | 86.11542797 | 0.310580537 | -3.779482382 | 0.000157155 | 0.015092448 |
| THEMIS | -1.519808915 | 105.8499927 | 0.402167326 | -3.779046223 | 0.00015743 | 0.015092448 |
| GIMAP8 | -1.28396284 | 269.4976866 | 0.339374833 | -3.783317781 | 0.000154752 | 0.015092448 |
| TMEM255B | -1.756800296 | 220.1328945 | 0.464430081 | -3.782701353 | 0.000155136 | 0.015092448 |
| FAM30A | -1.414596612 | 133.2635147 | 0.374190799 | -3.780415274 | 0.000156567 | 0.015092448 |
| ZBP1 | -1.35647874 | 143.9843833 | 0.358706348 | -3.781585542 | 0.000155833 | 0.015092448 |
| CORO1A | -1.486621555 | 1801.837764 | 0.393680335 | -3.776214919 | 0.00015923 | 0.015164525 |
| C1R | -1.209799421 | 2515.232594 | 0.321343769 | -3.764813695 | 0.000166673 | 0.015566211 |
| SPIB | -2.145855627 | 292.7216776 | 0.570453254 | -3.761667783 | 0.000168784 | 0.015662303 |
| CLSTN3 | -1.61577769 | 385.0779777 | 0.42978036 | -3.75954287 | 0.000170224 | 0.015695313 |
| TLR9 | -1.191521477 | 23.67917646 | 0.317134266 | -3.757151482 | 0.000171858 | 0.015745717 |
| TFEB | -1.06649877 | 296.0311543 | 0.284165166 | -3.753094675 | 0.000174665 | 0.015902191 |
| KLHL4 | -2.091247176 | 30.79863369 | 0.558996347 | -3.741074849 | 0.000183235 | 0.016578181 |
| MAP4K1 | -1.444147461 | 414.0890592 | 0.386877441 | -3.732829339 | 0.000189341 | 0.017024215 |
| CD1D | -1.270440558 | 148.277099 | 0.341205505 | -3.723388221 | 0.000196567 | 0.017564831 |
| KHDRBS2 | -1.369394603 | 26.77207673 | 0.368221217 | -3.718945403 | 0.000200056 | 0.017766962 |
| LY75 | -1.388534484 | 174.4220186 | 0.373883473 | -3.713816157 | 0.000204157 | 0.017803486 |
| AOX1 | -1.213701421 | 53.04175732 | 0.326686123 | -3.715191232 | 0.00020305 | 0.017803486 |
| DCSTAMP | -2.573224179 | 116.8444377 | 0.692854346 | -3.713946796 | 0.000204052 | 0.017803486 |
| CYTIP | -1.378153109 | 639.3960075 | 0.371815535 | -3.706550643 | 0.000210101 | 0.018212129 |
| NIBAN3 | -2.214148092 | 327.4353426 | 0.599957618 | -3.690507505 | 0.000223807 | 0.019284711 |
| ARRDC5 | -1.408332157 | 106.9703321 | 0.381839743 | -3.68828071 | 0.000225774 | 0.019339121 |
| PTPN22 | -1.332843331 | 190.6847447 | 0.361537996 | -3.686592689 | 0.000227277 | 0.019353278 |
| XAF1 | -1.208032589 | 617.3877017 | 0.328051997 | -3.682442418 | 0.00023101 | 0.01955615 |
| ZBTB32 | -1.758064418 | 39.52214581 | 0.478084051 | -3.677312422 | 0.000235704 | 0.01983753 |
| RTN1 | -1.053122946 | 117.996453 | 0.286536447 | -3.675354238 | 0.00023752 | 0.019874763 |

| | | | | | | |
|---|---|---|---|---|---|---|
| EMB | -1.393999261 | 196.4417081 | 0.379917105 | -3.669219532 | 0.000243292 | 0.020045067 |
| PPP1R1A | -1.986126643 | 42.86420753 | 0.541115784 | -3.670428219 | 0.000242144 | 0.020045067 |
| C3 | -1.579682615 | 2488.145385 | 0.430574159 | -3.668781745 | 0.000243709 | 0.020045067 |
| PIK3AP1 | -1.301056936 | 621.9018328 | 0.355189902 | -3.662989655 | 0.000249289 | 0.020388146 |
| GATA3 | -1.443998033 | 107.379307 | 0.394677884 | -3.65867481 | 0.000253523 | 0.02050277 |
| CXorf21 | -1.191369565 | 150.8210252 | 0.325582116 | -3.659198422 | 0.000253005 | 0.02050277 |
| SELE | -1.486946021 | 57.32671877 | 0.40773783 | -3.646818894 | 0.000265507 | 0.021128488 |
| ARHGAP9 | -1.455828667 | 398.9361681 | 0.399219047 | -3.646691402 | 0.000265639 | 0.021128488 |
| JAK3 | -1.350662305 | 554.0140561 | 0.370348964 | -3.646999012 | 0.000265321 | 0.021128488 |
| PTPRC | -1.636231747 | 242.7442353 | 0.448974844 | -3.644372884 | 0.000268045 | 0.021203351 |
| CLDN5 | -2.329595968 | 415.4883903 | 0.639567248 | -3.64245664 | 0.000270048 | 0.02124577 |
| RHOF | -1.544005041 | 334.5960284 | 0.424056239 | -3.641038377 | 0.000271541 | 0.021247689 |
| SCIMP | -1.467436633 | 270.3479136 | 0.403303878 | -3.638538366 | 0.00027419 | 0.021339631 |
| CD69 | -1.661034635 | 172.8615835 | 0.45700931 | -3.63457505 | 0.000278439 | 0.02155447 |
| CD180 | -1.393698109 | 173.2498309 | 0.384157483 | -3.627934295 | 0.000285698 | 0.021767176 |
| TREML2 | -1.495499079 | 36.97047489 | 0.412120032 | -3.628794917 | 0.000284747 | 0.021767176 |
| AKNA | -1.131658966 | 1069.345791 | 0.311856493 | -3.628781161 | 0.000284763 | 0.021767176 |
| NCKAP1L | -1.39127728 | 574.5569525 | 0.383637571 | -3.626540737 | 0.000287244 | 0.021770352 |
| TNFRSF9 | -1.784927547 | 192.4931851 | 0.494058502 | -3.612785815 | 0.000302925 | 0.022839276 |
| STAP1 | -2.431982963 | 152.3122688 | 0.674498968 | -3.605614062 | 0.000311416 | 0.023357789 |
| AOAH | -1.334163316 | 476.7311806 | 0.371117545 | -3.594988524 | 0.000324406 | 0.023959691 |
| ITGB7 | -1.211080891 | 232.3750183 | 0.336737737 | -3.596510751 | 0.000322514 | 0.023959691 |
| SLA2 | -1.301450849 | 78.53738462 | 0.361994757 | -3.595220164 | 0.000324117 | 0.023959691 |
| ADIPOQ | -3.192946316 | 42.85630887 | 0.889513977 | -3.589540353 | 0.000331262 | 0.024105815 |
| BANK1 | -3.237039958 | 405.4889409 | 0.901821322 | -3.589447131 | 0.00033138 | 0.024105815 |
| TNF | -1.414742216 | 101.2678584 | 0.393926718 | -3.591384269 | 0.000328926 | 0.024105815 |
| S1PR4 | -1.369464308 | 118.2746146 | 0.381757154 | -3.587265609 | 0.000334164 | 0.024186782 |
| KIF21B | -1.142132967 | 120.8705476 | 0.318823308 | -3.582338362 | 0.000340532 | 0.024525104 |
| TMEM176B | -1.195373489 | 1545.080614 | 0.333882426 | -3.580222842 | 0.000343301 | 0.024602127 |
| BIRC3 | -1.53877111 | 486.3287285 | 0.430350478 | -3.575623097 | 0.000349395 | 0.024793324 |
| MPP7 | -1.276767545 | 59.8766161 | 0.357821362 | -3.56817027 | 0.000359483 | 0.025384752 |
| LAMC2 | -1.639313014 | 50.37705411 | 0.460463003 | -3.560140559 | 0.000370656 | 0.025429484 |
| TRAF3IP3 | -1.415015886 | 602.0878684 | 0.397091015 | -3.563454805 | 0.000366006 | 0.025429484 |
| ANXA2R | -1.042223194 | 171.4806706 | 0.292272238 | -3.565932914 | 0.000362564 | 0.025429484 |
| CD3D | -1.462462836 | 265.4048875 | 0.410783265 | -3.560181147 | 0.000370599 | 0.025429484 |
| C16orf54 | -1.503756912 | 91.76403935 | 0.421886095 | -3.564367088 | 0.000364735 | 0.025429484 |
| ZNF101 | -0.577167668 | 367.1941269 | 0.162062333 | -3.561393055 | 0.000368892 | 0.025429484 |
| CXCL5 | -2.446552091 | 94.070303 | 0.688309911 | -3.554433912 | 0.000378794 | 0.025865177 |
| ANK3 | -1.470699093 | 106.2159007 | 0.41424244 | -3.550334179 | 0.000384742 | 0.026148035 |
| EXOC3L4 | -1.516306477 | 34.62301958 | 0.427946304 | -3.543216669 | 0.000395278 | 0.026738511 |
| LAX1 | -1.252083497 | 91.25487464 | 0.353773457 | -3.539223965 | 0.000401305 | 0.026999627 |
| 1-Mar | -1.369749387 | 362.6982182 | 0.387131943 | -3.538197791 | 0.000402868 | 0.026999627 |
| ARHGAP15 | -1.778081659 | 555.7778843 | 0.50292079 | -3.535510349 | 0.000406988 | 0.027150058 |
| WNT5A | -1.035922385 | 59.27490455 | 0.29414022 | -3.521865814 | 0.000428521 | 0.028196681 |
| SLFN12L | -1.451016584 | 90.51065669 | 0.411989569 | -3.521974081 | 0.000428346 | 0.028196681 |
| TUBA4A | -1.246755739 | 213.0473177 | 0.354473481 | -3.51720455 | 0.000436118 | 0.028566687 |
| CLECL1 | -1.410431669 | 114.8774841 | 0.401925846 | -3.509183801 | 0.000449484 | 0.029309607 |
| IKZF3 | -2.272010098 | 353.0321696 | 0.648203448 | -3.505087955 | 0.000456456 | 0.029630779 |
| PTGER4 | -0.89505431 | 220.6437884 | 0.255585984 | -3.501969463 | 0.000461833 | 0.029845928 |
| WNT10A | -0.991541459 | 69.78207809 | 0.283624026 | -3.495971319 | 0.000472339 | 0.030389272 |
| RASGRP2 | -1.596930163 | 389.3958126 | 0.458645095 | -3.481842892 | 0.000497976 | 0.031756375 |
| LCK | -1.371083881 | 191.2928572 | 0.394508515 | -3.47542278 | 0.000510049 | 0.032383625 |
| TRGC2 | -1.23724695 | 50.34134484 | 0.356196264 | -3.473497827 | 0.000513722 | 0.032474379 |
| SLC4A8 | -0.980702934 | 65.40625822 | 0.282678541 | -3.469322184 | 0.000521773 | 0.03283996 |
| RASGRP1 | -1.328584953 | 216.1752816 | 0.38342372 | -3.465056759 | 0.00053012 | 0.033220827 |
| CR2 | -2.664762167 | 110.1669061 | 0.770034842 | -3.460573497 | 0.000539026 | 0.033633368 |

| | | | | | | |
|---|---|---|---|---|---|---|
| ABCG1 | -1.11751276 | 349.5674243 | 0.324040033 | -3.448687346 | 0.000563319 | 0.034848715 |
| TRBC2 | -1.543338236 | 1530.42297 | 0.448163953 | -3.443691146 | 0.000573831 | 0.035198198 |
| AC119396.1 | -1.809227 | 54.31908598 | 0.525259413 | -3.444444695 | 0.000572234 | 0.035198198 |
| LTB | -1.838016239 | 1334.003027 | 0.535038668 | -3.43529608 | 0.000591907 | 0.035875437 |
| TRDC | -1.174524456 | 55.90061749 | 0.341917258 | -3.435113113 | 0.000592307 | 0.035875437 |
| INSL3 | -1.079609669 | 30.47154861 | 0.314660147 | -3.431034015 | 0.000601285 | 0.036267519 |
| MILR1 | -1.082444562 | 214.2965495 | 0.316250228 | -3.422747133 | 0.000619917 | 0.037082318 |
| SH2D3C | -1.340156281 | 233.4905334 | 0.391942141 | -3.419270707 | 0.000627892 | 0.037404808 |
| ABCD2 | -1.528961948 | 84.08138357 | 0.447375121 | -3.417628469 | 0.000631693 | 0.037476984 |
| OLFM3 | -2.205000439 | 56.05833716 | 0.64577327 | -3.414511784 | 0.000638964 | 0.037661637 |
| BASP1 | -1.232390977 | 521.37489 | 0.360974461 | -3.414066946 | 0.000640008 | 0.037661637 |
| TIFAB | -1.300481273 | 94.37989958 | 0.38111594 | -3.412298298 | 0.000644176 | 0.037753396 |
| DPYD | -1.004341601 | 474.2485392 | 0.294776245 | -3.407132081 | 0.000656493 | 0.037807723 |
| DYSF | -0.938193262 | 216.0220682 | 0.275242507 | -3.408606009 | 0.000652957 | 0.037807723 |
| TRAF1 | -1.194038435 | 788.2101395 | 0.350138872 | -3.410185304 | 0.000649188 | 0.037807723 |
| IGHM | -2.114499887 | 5351.33786 | 0.620933183 | -3.405358169 | 0.000660773 | 0.037807723 |
| SPN | -1.443346121 | 256.2535039 | 0.423377452 | -3.409123735 | 0.000651719 | 0.037807723 |
| CFB | -1.178090168 | 280.9395629 | 0.346333984 | -3.401601411 | 0.000669923 | 0.038030591 |
| SLC12A3 | -1.149792178 | 36.4879404 | 0.338649601 | -3.395226733 | 0.000685718 | 0.038775203 |
| RNASET2 | -0.915132553 | 437.2927279 | 0.269731128 | -3.392758415 | 0.000691926 | 0.038822967 |
| SIGLEC10 | -1.454958386 | 155.6111719 | 0.428779448 | -3.393255886 | 0.000690671 | 0.038822967 |
| PLA2G4C | -1.23269604 | 570.7012344 | 0.363728263 | -3.389057618 | 0.000701333 | 0.039198813 |
| PTPN7 | -1.388428759 | 356.9420964 | 0.410007356 | -3.38635085 | 0.000708288 | 0.039284185 |
| ABI3 | -1.238166822 | 338.1261341 | 0.365599578 | -3.386674646 | 0.000707452 | 0.039284185 |
| INPP5D | -1.666218979 | 319.2851856 | 0.492558013 | -3.382787275 | 0.000717542 | 0.039645558 |
| DEGS2 | -0.892210181 | 27.89101913 | 0.263929678 | -3.380484484 | 0.000723582 | 0.039827252 |
| LDOC1 | -1.39257491 | 79.62806327 | 0.412100769 | -3.379209685 | 0.000726945 | 0.039860839 |
| TBX1 | -2.42335507 | 93.56403019 | 0.717540572 | -3.377307381 | 0.000731992 | 0.039986099 |
| FOXP3 | -1.245465224 | 91.80315767 | 0.369027905 | -3.374989284 | 0.000738186 | 0.04017284 |
| SEMA4A | -0.805287469 | 160.627006 | 0.238686936 | -3.37382297 | 0.00074132 | 0.04019233 |
| GPR132 | -1.495316743 | 203.7127377 | 0.444660497 | -3.362827937 | 0.000771484 | 0.041671669 |
| GIMAP7 | -1.487049486 | 654.437171 | 0.442446509 | -3.360970095 | 0.000776692 | 0.04179702 |
| ANO9 | -2.040125341 | 292.9960652 | 0.607663147 | -3.357329389 | 0.000786993 | 0.042194482 |
| TMEM132A | -1.630203927 | 80.8935555 | 0.48596093 | -3.354598747 | 0.000794802 | 0.042455907 |
| SNCAIP | -1.705186468 | 48.9193395 | 0.508583031 | -3.352818252 | 0.000799932 | 0.04257286 |
| S1PR1 | -1.441736399 | 261.1712779 | 0.430506362 | -3.348931696 | 0.000811238 | 0.042891973 |
| IKZF1 | -1.588795782 | 1249.841643 | 0.474448494 | -3.348721308 | 0.000811854 | 0.042891973 |
| CD3E | -1.443031787 | 359.0186675 | 0.431128441 | -3.347104137 | 0.000816605 | 0.042986102 |
| SIRPG | -1.130814184 | 80.85886499 | 0.338145142 | -3.344168064 | 0.000825297 | 0.043129975 |
| TESPA1 | -1.520901142 | 102.6924618 | 0.455424798 | -3.339522019 | 0.000839227 | 0.043700179 |
| DAPK2 | -0.994785341 | 79.21810117 | 0.298031736 | -3.337850369 | 0.000844292 | 0.043806349 |
| TGFB1 | -0.94660286 | 357.2987386 | 0.283989373 | -3.333233387 | 0.000858429 | 0.04438078 |
| IGHA2 | -1.505084068 | 35.0680642 | 0.451734754 | -3.331787192 | 0.000862902 | 0.044453279 |
| PRDM8 | -1.011976747 | 56.66911848 | 0.304134225 | -3.327401738 | 0.000876599 | 0.044839742 |
| TBX21 | -1.338699221 | 37.56485223 | 0.402235233 | -3.32815008 | 0.000874248 | 0.044839742 |
| PLCG2 | -1.389798068 | 324.8956181 | 0.417955519 | -3.32522961 | 0.000883457 | 0.045031444 |
| CD6 | -1.414294044 | 231.2553108 | 0.425666937 | -3.322536756 | 0.000892029 | 0.045308823 |
| TTC7A | -0.922958737 | 305.5171087 | 0.277895914 | -3.321238967 | 0.000896188 | 0.045360885 |
| SIGLEC8 | -1.154603622 | 50.18346251 | 0.34806909 | -3.317167925 | 0.000909349 | 0.045707439 |
| TRAC | -1.356670951 | 1709.147503 | 0.409310623 | -3.314526612 | 0.000917984 | 0.045981808 |
| PLAAT4 | -1.528010026 | 786.6704118 | 0.461139683 | -3.313551362 | 0.000921192 | 0.045983356 |
| LAT2 | -1.482177542 | 207.7090291 | 0.448946179 | -3.301459311 | 0.000961833 | 0.047520451 |
| SLC38A1 | -1.59289255 | 603.5815895 | 0.483317316 | -3.295748975 | 0.000981597 | 0.048331957 |
| HSH2D | -2.037756726 | 118.7889232 | 0.619023433 | -3.29188948 | 0.000995167 | 0.048834031 |
| CD2 | -1.403915775 | 515.1564802 | 0.427860926 | -3.281243248 | 0.001033506 | 0.049914925 |
| PM20D1 | -1.30162947 | 20.24411474 | 0.396721549 | -3.280964882 | 0.001034526 | 0.049914925 |

| | | | | | | |
|---|---|---|---|---|---|---|
| LY86 | -1.235530581 | 240.1233384 | 0.376895023 | -3.278182269 | 0.001044779 | 0.049914925 |
| CBLN3 | -0.940988459 | 75.96717973 | 0.286916958 | -3.279654379 | 0.001039343 | 0.049914925 |
| PLIN1 | -1.933292066 | 57.1903182 | 0.588592628 | -3.284601221 | 0.001021268 | 0.049914925 |
| ABAT | -0.967261798 | 117.6609039 | 0.295027239 | -3.278550829 | 0.001043416 | 0.049914925 |
| SDR42E1 | -0.856084798 | 23.86603333 | 0.260730133 | -3.283413339 | 0.001025582 | 0.049914925 |
| LILRB4 | -1.147306666 | 97.53897234 | 0.349974827 | -3.278254828 | 0.001044511 | 0.049914925 |

| gene | log2FoldChange | baseMean | lfcSE | stat | pvalue | padj |
|---|---|---|---|---|---|---|
| CA14 | 2.37092582 | 423.2683058 | 0.516920671 | 4.586633801 | 4.50E-06 | 0.003260355 |
| GABRG3 | 3.701810514 | 126.5725718 | 0.895914318 | 4.131880067 | 3.60E-05 | 0.006853398 |
| ZFYVE9 | 0.76876712 | 680.9922872 | 0.190388775 | 4.037880496 | 5.39E-05 | 0.008772833 |
| ABCB6 | 1.082101879 | 813.9876314 | 0.286204288 | 3.780872347 | 0.00015628 | 0.015092448 |
| QPCTL | 0.941442347 | 417.1367473 | 0.249500028 | 3.773315602 | 0.000161092 | 0.015241647 |
| GSTP1 | 1.076345763 | 7828.564191 | 0.285719589 | 3.767140251 | 0.000165128 | 0.015522055 |
| DDRGK1 | 0.540345104 | 1647.073636 | 0.151054697 | 3.577148648 | 0.000347363 | 0.024770551 |
| WDR12 | 0.834999485 | 628.9534279 | 0.237082826 | 3.521973734 | 0.000428347 | 0.028196681 |
| GAS2L3 | 1.068292505 | 665.3481834 | 0.305740773 | 3.494111999 | 0.000475641 | 0.030466307 |
| KCNIP3 | 1.412245396 | 180.7595207 | 0.409318206 | 3.450238412 | 0.000560092 | 0.034797799 |
| MFSD10 | 0.591801665 | 892.6811835 | 0.172734422 | 3.426078355 | 0.000612364 | 0.036782477 |
| FAM168B | 0.585030829 | 1034.067882 | 0.171795948 | 3.405381997 | 0.000660716 | 0.037807723 |
| ZMYM4 | 0.595900016 | 1258.978768 | 0.175065171 | 3.403875318 | 0.000664371 | 0.0378639 |
| SLC25A15 | 0.907919972 | 372.4047956 | 0.271361336 | 3.345797099 | 0.000820464 | 0.043032747 |
| DHX37 | 0.691732567 | 502.4612231 | 0.208480339 | 3.317975066 | 0.000906726 | 0.045707439 |
| SNX8 | 0.713043433 | 1722.48313 | 0.215375144 | 3.31070438 | 0.000930615 | 0.046294082 |
| TCFL5 | 1.372458277 | 607.7783383 | 0.41496221 | 3.307429549 | 0.000941564 | 0.046678354 |

| ID | Description | setSize | enrichmentScore | NES | pvalue |
|---|---|---|---|---|---|
| GO:0042254 | ribosome biogenesis | 251 | 0.552200422 | 3.003524645 | 1e-10 |
| GO:0006364 | rRNA processing | 187 | 0.562040979 | 2.943297309 | 1e-10 |
| GO:0022613 | ribonucleoprotein complex biogenesis | 388 | 0.513512691 | 2.928293287 | 1e-10 |
| GO:0042773 | ATP synthesis coupled electron transport | 79 | 0.644583299 | 2.924337481 | 1e-10 |
| GO:0042775 | mitochondrial ATP synthesis coupled electron transport | 79 | 0.644583299 | 2.924337481 | 1e-10 |
| GO:0016072 | rRNA metabolic process | 224 | 0.53963313 | 2.894847745 | 1e-10 |
| GO:0019646 | aerobic electron transport chain | 72 | 0.649766645 | 2.89434858 | 1e-10 |
| GO:0006119 | oxidative phosphorylation | 121 | 0.580856999 | 2.842418354 | 1e-10 |
| GO:0034470 | ncRNA processing | 383 | 0.494591681 | 2.81621605 | 1e-10 |
| GO:0009060 | aerobic respiration | 170 | 0.528024266 | 2.727729676 | 1e-10 |
| GO:0007080 | mitotic metaphase chromosome alignment | 58 | 0.629066506 | 2.672406203 | 1e-10 |
| GO:0042273 | ribosomal large subunit biogenesis | 51 | 0.646787698 | 2.669794757 | 2.87261453464023e-10 |
| GO:0033108 | mitochondrial respiratory chain complex assembly | 91 | 0.571750164 | 2.66423209 | 1e-10 |
| GO:0000070 | mitotic sister chromatid segregation | 183 | 0.509681592 | 2.661480059 | 1e-10 |
| GO:0022904 | respiratory electron transport chain | 97 | 0.562233353 | 2.648325197 | 1e-10 |
| GO:0006399 | tRNA metabolic process | 173 | 0.510580996 | 2.644158959 | 1e-10 |
| GO:0140053 | mitochondrial gene expression | 83 | 0.571196894 | 2.61610809 | 1e-10 |
| GO:0042274 | ribosomal small subunit biogenesis | 71 | 0.588020937 | 2.611067134 | 6.91322825273435e-10 |
| GO:1902850 | microtubule cytoskeleton organization involved in mitosis | 158 | 0.506110334 | 2.589992444 | 1e-10 |
| GO:0051310 | metaphase chromosome alignment | 96 | 0.549852838 | 2.586621405 | 1e-10 |
| GO:0000819 | sister chromatid segregation | 224 | 0.479792257 | 2.573832955 | 1e-10 |
| GO:0042776 | proton motive force-driven mitochondrial ATP synthesis | 55 | 0.612242274 | 2.571625984 | 2.99123269346856e-09 |
| GO:0007052 | mitotic spindle organization | 126 | 0.51434396 | 2.534253496 | 1e-10 |
| GO:0051983 | regulation of chromosome segregation | 128 | 0.512579849 | 2.531248253 | 1e-10 |
| GO:0006120 | mitochondrial electron transport, NADH to ubiquinone | 41 | 0.645307262 | 2.529953367 | 8.25358076662769e-08 |
| GO:0008608 | attachment of spindle microtubules to kinetochore | 50 | 0.612606875 | 2.516532034 | 7.41943832976287e-09 |
| GO:0015986 | proton motive force-driven ATP synthesis | 62 | 0.582845677 | 2.514273017 | 1.42878250053289e-08 |
| GO:0045333 | cellular respiration | 210 | 0.470671347 | 2.503621447 | 1e-10 |
| GO:0045839 | negative regulation of mitotic nuclear division | 56 | 0.591654768 | 2.494326109 | 3.4772227981401e-08 |
| GO:0008033 | tRNA processing | 124 | 0.507231068 | 2.492831763 | 1e-10 |
| GO:0007006 | mitochondrial membrane organization | 108 | 0.516332459 | 2.477946289 | 1e-10 |
| GO:0010257 | NADH dehydrogenase complex assembly | 54 | 0.591647748 | 2.474055951 | 2.16000310418915e-08 |
| GO:0032981 | mitochondrial respiratory chain complex I assembly | 54 | 0.591647748 | 2.474055951 | 2.16000310418915e-08 |
| GO:0007059 | chromosome segregation | 405 | 0.432035294 | 2.472700971 | 1e-10 |
| GO:2001251 | negative regulation of chromosome organization | 90 | 0.529837369 | 2.464442937 | 7.0406007850212e-10 |
| GO:0032508 | DNA duplex unwinding | 72 | 0.552066817 | 2.459150249 | 4.64536153057756e-09 |
| GO:0140014 | mitotic nuclear division | 267 | 0.448810792 | 2.458406226 | 1e-10 |
| GO:0007091 | metaphase/anaphase transition of mitotic cell cycle | 94 | 0.519856781 | 2.437909343 | 1.52171105769607e-09 |
| GO:0051303 | establishment of chromosome localization | 106 | 0.508151606 | 2.430263889 | 2.70347171504166e-10 |
| GO:0000075 | cell cycle checkpoint signaling | 186 | 0.464230824 | 2.429325874 | 1e-10 |
| GO:0050000 | chromosome localization | 114 | 0.499590976 | 2.419846945 | 3.75822854583405e-10 |
| GO:0030071 | regulation of mitotic metaphase/anaphase transition | 90 | 0.518957401 | 2.413836728 | 2.27844752705337e-09 |
| GO:0032392 | DNA geometric change | 77 | 0.534704966 | 2.412721294 | 3.49923567932597e-08 |
| GO:0032543 | mitochondrial translation | 50 | 0.586006171 | 2.407258817 | 1.00753658896532e-07 |

| | | | | | |
|---|---|---|---|---|---|
| GO:0051315 | attachment of mitotic spindle microtubules to kinetochore | 24 | 0.70360614 | 2.405344427 | 1.25340883948173e-06 |
| GO:0030490 | maturation of SSU-rRNA | 45 | 0.599018452 | 2.402272999 | 5.79590990248763e-07 |
| GO:0010965 | regulation of mitotic sister chromatid separation | 59 | 0.562036921 | 2.396953489 | 5.69534637692326e-08 |
| GO:0008380 | RNA splicing | 417 | 0.417595988 | 2.396517894 | 1e-10 |
| GO:0051306 | mitotic sister chromatid separation | 62 | 0.553226542 | 2.386502328 | 2.11465982026046e-07 |
| GO:0022618 | protein-RNA complex assembly | 167 | 0.463117092 | 2.384217229 | 1e-10 |
| GO:0006261 | DNA-templated DNA replication | 159 | 0.463425122 | 2.372951909 | 1e-10 |
| GO:0006839 | mitochondrial transport | 167 | 0.459472353 | 2.365453398 | 1e-10 |
| GO:1905818 | regulation of chromosome separation | 74 | 0.528164536 | 2.363894344 | 5.77339085965525e-08 |
| GO:0000460 | maturation of 5.8S rRNA | 34 | 0.628339931 | 2.356472315 | 1.10492972553517e-06 |
| GO:0000959 | mitochondrial RNA metabolic process | 50 | 0.573121866 | 2.354331291 | 2.73972358202894e-07 |
| GO:0007093 | mitotic cell cycle checkpoint signaling | 139 | 0.467258232 | 2.340050507 | 9.16507664374736e-10 |
| GO:0033046 | negative regulation of sister chromatid segregation | 48 | 0.574357979 | 2.338989325 | 8.08789781580666e-07 |
| GO:0033048 | negative regulation of mitotic sister chromatid segregation | 48 | 0.574357979 | 2.338989325 | 8.08789781580666e-07 |
| GO:0045841 | negative regulation of mitotic metaphase/anaphase transition | 48 | 0.574357979 | 2.338989325 | 8.08789781580666e-07 |
| GO:2000816 | negative regulation of mitotic sister chromatid separation | 48 | 0.574357979 | 2.338989325 | 8.08789781580666e-07 |
| GO:0071826 | protein-RNA complex organization | 176 | 0.450436439 | 2.337559264 | 1e-10 |
| GO:0051653 | spindle localization | 61 | 0.542418255 | 2.331313643 | 5.16441097416419e-07 |
| GO:0031577 | spindle checkpoint signaling | 47 | 0.575151431 | 2.329432543 | 1.15589823614764e-06 |
| GO:0007094 | mitotic spindle assembly checkpoint signaling | 46 | 0.577306963 | 2.3271706 | 7.04284620920751e-07 |
| GO:0071173 | spindle assembly checkpoint signaling | 46 | 0.577306963 | 2.3271706 | 7.04284620920751e-07 |
| GO:0071174 | mitotic spindle checkpoint signaling | 46 | 0.577306963 | 2.3271706 | 7.04284620920751e-07 |
| GO:0022616 | DNA strand elongation | 37 | 0.60807255 | 2.326913604 | 3.84038505866067e-06 |
| GO:0033047 | regulation of mitotic sister chromatid segregation | 54 | 0.556338266 | 2.326404522 | 4.50191476153367e-07 |
| GO:0098813 | nuclear chromosome segregation | 306 | 0.417689465 | 2.326194374 | 1e-10 |
| GO:0051304 | chromosome separation | 80 | 0.511055939 | 2.323497304 | 4.24344174237726e-08 |
| GO:0009201 | ribonucleoside triphosphate biosynthetic process | 99 | 0.491300322 | 2.323375727 | 7.09656851206447e-09 |
| GO:0090151 | establishment of protein localization to mitochondrial membrai | 34 | 0.618807272 | 2.320721849 | 2.02226945183388e-06 |
| GO:0009303 | rRNA transcription | 35 | 0.614228114 | 2.320505977 | 7.11260947874432e-06 |
| GO:0044784 | metaphase/anaphase transition of cell cycle | 97 | 0.491626707 | 2.315742012 | 1.09942163383407e-08 |
| GO:0006400 | tRNA modification | 86 | 0.501926956 | 2.313493266 | 4.86122047542221e-08 |
| GO:0043414 | macromolecule methylation | 163 | 0.449723912 | 2.309028006 | 1e-10 |
| GO:0006418 | tRNA aminoacylation for protein translation | 22 | 0.691387804 | 2.304528809 | 1.64702394415132e-05 |
| GO:0007051 | spindle organization | 189 | 0.439215482 | 2.303738914 | 1e-10 |
| GO:0007098 | centrosome cycle | 133 | 0.462517124 | 2.29805514 | 1.02252736753853e-09 |
| GO:0033045 | regulation of sister chromatid segregation | 104 | 0.481401928 | 2.296662558 | 1.43270955065032e-08 |
| GO:0000387 | spliceosomal snRNP assembly | 37 | 0.599762062 | 2.295111826 | 6.38967281203533e-06 |
| GO:0051984 | positive regulation of chromosome segregation | 27 | 0.649922357 | 2.293839407 | 7.78635956359866e-06 |
| GO:1902099 | regulation of metaphase/anaphase transition of cell cycle | 93 | 0.48998236 | 2.293180147 | 4.23373254062375e-08 |
| GO:0071103 | DNA conformation change | 84 | 0.499186841 | 2.291808154 | 9.26179124766922e-08 |
| GO:0007007 | inner mitochondrial membrane organization | 40 | 0.588107309 | 2.291780197 | 4.93247350548898e-06 |
| GO:0051985 | negative regulation of chromosome segregation | 50 | 0.556386561 | 2.285584219 | 1.01050255208496e-06 |
| GO:1902100 | negative regulation of metaphase/anaphase transition of cell cy | 50 | 0.556386561 | 2.285584219 | 1.01050255208496e-06 |
| GO:1905819 | negative regulation of chromosome separation | 50 | 0.556386561 | 2.285584219 | 1.01050255208496e-06 |
| GO:0051293 | establishment of spindle localization | 57 | 0.53964413 | 2.284448048 | 1.0185067942805e-06 |

| | | | | | |
|---|---|---|---|---|---|
| GO:0042255 | ribosome assembly | 36 | 0.600185584 | 2.281849354 | 5.03064982631802e-06 |
| GO:0006360 | transcription by RNA polymerase I | 62 | 0.527119557 | 2.273882315 | 1.51147645977642e-06 |
| GO:0034508 | centromere complex assembly | 28 | 0.637269332 | 2.272196821 | 1.79095628555938e-05 |
| GO:0000470 | maturation of LSU-rRNA | 22 | 0.681255289 | 2.270755184 | 2.57058811218433e-05 |
| GO:0009142 | nucleoside triphosphate biosynthetic process | 106 | 0.474768293 | 2.270606301 | 1.31283379831904e-08 |
| GO:0070199 | establishment of protein localization to chromosome | 22 | 0.680564163 | 2.268451528 | 2.60757435795539e-05 |
| GO:0000375 | RNA splicing, via transesterification reactions | 284 | 0.410723306 | 2.268338004 | 1e-10 |
| GO:0031023 | microtubule organizing center organization | 146 | 0.448561927 | 2.262986456 | 1.2583552192796e-09 |
| GO:0000377 | RNA splicing, via transesterification reactions with bulged ade | 280 | 0.408961983 | 2.255195723 | 1e-10 |
| GO:0000398 | mRNA splicing, via spliceosome | 280 | 0.408961983 | 2.255195723 | 1e-10 |
| GO:0006582 | melanin metabolic process | 24 | 0.658727169 | 2.251921402 | 2.53323339742487e-05 |
| GO:0006397 | mRNA processing | 449 | 0.389090416 | 2.2497205 | 1e-10 |
| GO:0045048 | protein insertion into ER membrane | 28 | 0.630896538 | 2.249474492 | 2.78240307554949e-05 |
| GO:0000462 | maturation of SSU-rRNA from tricistronic rRNA transcript (S | 29 | 0.62385847 | 2.245597387 | 1.50631392896596e-05 |
| GO:0033044 | regulation of chromosome organization | 238 | 0.413984622 | 2.237694398 | 1e-10 |
| GO:0001510 | RNA methylation | 84 | 0.486868969 | 2.235255787 | 2.93245421821848e-07 |
| GO:0006260 | DNA replication | 273 | 0.406949589 | 2.234694364 | 1e-10 |
| GO:0000027 | ribosomal large subunit assembly | 15 | 0.748820147 | 2.227612431 | 1.36690123262063e-05 |
| GO:0051784 | negative regulation of nuclear division | 62 | 0.512784594 | 2.212044317 | 5.39734174442599e-06 |
| GO:0022900 | electron transport chain | 145 | 0.438504179 | 2.210569025 | 3.16300187080951e-09 |
| GO:0009206 | purine ribonucleoside triphosphate biosynthetic process | 93 | 0.47222469 | 2.210071979 | 3.01564797220183e-07 |
| GO:0034502 | protein localization to chromosome | 99 | 0.466879231 | 2.207887567 | 1.00695474599441e-07 |
| GO:0009145 | purine nucleoside triphosphate biosynthetic process | 94 | 0.469102737 | 2.199894254 | 2.1427915130115e-07 |
| GO:0007008 | outer mitochondrial membrane organization | 15 | 0.737748885 | 2.194677314 | 2.57451841062165e-05 |
| GO:0045040 | protein insertion into mitochondrial outer membrane | 15 | 0.737748885 | 2.194677314 | 2.57451841062165e-05 |
| GO:0043039 | tRNA aminoacylation | 25 | 0.63540243 | 2.194289868 | 3.93491372838668e-05 |
| GO:0070198 | protein localization to chromosome, telomeric region | 27 | 0.620976596 | 2.191678085 | 3.56990468240705e-05 |
| GO:0042438 | melanin biosynthetic process | 23 | 0.648945617 | 2.190895382 | 0.00010014 |
| GO:0044550 | secondary metabolite biosynthetic process | 25 | 0.63223439 | 2.183349403 | 4.46974057368328e-05 |
| GO:0051383 | kinetochore organization | 21 | 0.663862429 | 2.182781387 | 0.00010588 |
| GO:0006626 | protein targeting to mitochondrion | 104 | 0.456919744 | 2.179863448 | 2.18242013380798e-07 |
| GO:0043473 | pigmentation | 111 | 0.450497798 | 2.173036 | 1.63782209785616e-07 |
| GO:0043038 | amino acid activation | 26 | 0.620326791 | 2.167679396 | 8.27285400956884e-05 |
| GO:1904814 | regulation of protein localization to chromosome, telomeric reg | 11 | 0.800118786 | 2.165593732 | 2.35499050373172e-05 |
| GO:0006270 | DNA replication initiation | 37 | 0.565833326 | 2.1652766 | 3.86141342670252e-05 |
| GO:0006479 | protein methylation | 43 | 0.545994694 | 2.164198578 | 1.34208867542306e-05 |
| GO:0008213 | protein alkylation | 43 | 0.545994694 | 2.164198578 | 1.34208867542306e-05 |
| GO:0046148 | pigment biosynthetic process | 54 | 0.516182763 | 2.1584888 | 9.16872988206205e-06 |
| GO:0051298 | centrosome duplication | 72 | 0.484352256 | 2.157519588 | 1.48287555658415e-06 |
| GO:0000280 | nuclear division | 424 | 0.371948016 | 2.139239032 | 1e-10 |
| GO:0072655 | establishment of protein localization to mitochondrion | 118 | 0.438896239 | 2.138606094 | 1.08789227417352e-07 |
| GO:0006754 | ATP biosynthetic process | 84 | 0.465659252 | 2.137880217 | 2.05249646296178e-06 |
| GO:1901970 | positive regulation of mitotic sister chromatid separation | 18 | 0.679673092 | 2.136960129 | 0.000131398 |
| GO:0000154 | rRNA modification | 32 | 0.57817728 | 2.134405538 | 7.85302199536832e-05 |
| GO:1990542 | mitochondrial transmembrane transport | 89 | 0.459808425 | 2.133963417 | 1.76809044327173e-06 |

| | | | | | |
|---|---|---|---|---|---|
| GO:0031123 | RNA 3'-end processing | 89 | 0.459211838 | 2.131194666 | 1.90279816782048e-06 |
| GO:0006302 | double-strand break repair | 302 | 0.38232973 | 2.125563273 | 1e-10 |
| GO:1904816 | positive regulation of protein localization to chromosome, telon | 10 | 0.81103867 | 2.124116004 | 0.000128576 |
| GO:0007005 | mitochondrion organization | 477 | 0.365079724 | 2.121768315 | 1e-10 |
| GO:0070585 | protein localization to mitochondrion | 124 | 0.429975553 | 2.113152731 | 1.56304901408407e-07 |
| GO:0006123 | mitochondrial electron transport, cytochrome c to oxygen | 18 | 0.671975593 | 2.112758422 | 0.000174621 |
| GO:0048285 | organelle fission | 466 | 0.363658803 | 2.110668594 | 1e-10 |
| GO:0030318 | melanocyte differentiation | 24 | 0.617285261 | 2.110248304 | 0.000163018 |
| GO:0071459 | protein localization to chromosome, centromeric region | 23 | 0.623567318 | 2.105216095 | 0.000355863 |
| GO:0170036 | import into the mitochondrion | 42 | 0.533841798 | 2.103983495 | 7.63978934492727e-05 |
| GO:0000725 | recombinational repair | 175 | 0.405548729 | 2.103747965 | 5.47968089105948e-09 |
| GO:0007144 | female meiosis I | 11 | 0.777120092 | 2.103345688 | 6.65298826866447e-05 |
| GO:0031570 | DNA integrity checkpoint signaling | 130 | 0.424762176 | 2.103199916 | 1.5799833707296e-07 |
| GO:0017004 | cytochrome complex assembly | 36 | 0.552975424 | 2.10236075 | 9.03750188487867e-05 |
| GO:0015980 | energy derivation by oxidation of organic compounds | 294 | 0.379141453 | 2.102348319 | 1e-10 |
| GO:0044818 | mitotic G2/M transition checkpoint | 52 | 0.505753836 | 2.096898615 | 2.27263884917529e-05 |
| GO:0007088 | regulation of mitotic nuclear division | 116 | 0.431476759 | 2.095525413 | 4.59617594512184e-07 |
| GO:0032509 | endosome transport via multivesicular body sorting pathway | 40 | 0.537698739 | 2.095344341 | 7.19726463198148e-05 |
| GO:0050931 | pigment cell differentiation | 35 | 0.553496346 | 2.091066089 | 0.000153931 |
| GO:0030150 | protein import into mitochondrial matrix | 18 | 0.663122376 | 2.084923025 | 0.000258322 |
| GO:0051988 | regulation of attachment of spindle microtubules to kinetochore | 19 | 0.651553133 | 2.080935776 | 0.000362403 |
| GO:0006122 | mitochondrial electron transport, ubiquinol to cytochrome c | 12 | 0.748420719 | 2.080639748 | 0.000169402 |
| GO:0000724 | double-strand break repair via homologous recombination | 171 | 0.401993633 | 2.077942055 | 4.53968344750473e-08 |
| GO:0000466 | maturation of 5.8S rRNA from tricistronic rRNA transcript (SS | 24 | 0.606586661 | 2.073674125 | 0.000248334 |
| GO:0070200 | establishment of protein localization to telomere | 15 | 0.694538719 | 2.066134428 | 0.000348207 |
| GO:0006282 | regulation of DNA repair | 208 | 0.388457722 | 2.063542745 | 5.61826531994662e-09 |
| GO:0044743 | protein transmembrane import into intracellular organelle | 36 | 0.542553583 | 2.062737885 | 0.00014178 |
| GO:0040001 | establishment of mitotic spindle localization | 40 | 0.528945951 | 2.061235831 | 0.000115554 |
| GO:0098693 | regulation of synaptic vesicle cycle | 13 | 0.722293282 | 2.060692641 | 0.000509029 |
| GO:0018126 | protein hydroxylation | 22 | 0.618174375 | 2.06049434 | 0.000372687 |
| GO:0070202 | regulation of establishment of protein localization to chromosol | 10 | 0.786476944 | 2.05978867 | 0.00038622 |
| GO:0006356 | regulation of transcription by RNA polymerase I | 43 | 0.518815916 | 2.056468093 | 6.11544367718507e-05 |
| GO:0006268 | DNA unwinding involved in DNA replication | 21 | 0.625325875 | 2.056073097 | 0.000501293 |
| GO:0046112 | nucleobase biosynthetic process | 17 | 0.664868905 | 2.054521471 | 0.000481928 |
| GO:0048066 | developmental pigmentation | 48 | 0.503754338 | 2.051466266 | 6.67558091517719e-05 |
| GO:0006354 | DNA-templated transcription elongation | 128 | 0.414730895 | 2.048045497 | 8.72484876039154e-07 |
| GO:0032200 | telomere organization | 155 | 0.401513898 | 2.047204937 | 1.95029710577549e-07 |
| GO:0048023 | positive regulation of melanin biosynthetic process | 10 | 0.780172508 | 2.043277306 | 0.00032894 |
| GO:1900378 | positive regulation of secondary metabolite biosynthetic proces | 10 | 0.780172508 | 2.043277306 | 0.00032894 |
| GO:1990173 | protein localization to nucleoplasm | 12 | 0.73480184 | 2.042778717 | 0.000528592 |
| GO:0032042 | mitochondrial DNA metabolic process | 23 | 0.603990433 | 2.039122875 | 0.000482936 |
| GO:0033059 | cellular pigmentation | 61 | 0.474145182 | 2.03787598 | 7.32084175171808e-05 |
| GO:0051382 | kinetochore assembly | 17 | 0.659213674 | 2.037046157 | 0.00065948 |
| GO:0043162 | ubiquitin-dependent protein catabolic process via the multivesi | 36 | 0.534461109 | 2.031971059 | 0.0002383 |
| GO:0000077 | DNA damage checkpoint signaling | 121 | 0.414730346 | 2.029479115 | 7.48963122162334e-07 |

| | | | | | |
|---|---|---|---|---|---|
| GO:0000723 | telomere maintenance | 150 | 0.399319726 | 2.022952569 | 2.3767220405354e-07 |
| GO:0042407 | cristae formation | 16 | 0.666932073 | 2.022696241 | 0.00065261 |
| GO:0009225 | nucleotide-sugar metabolic process | 35 | 0.535387297 | 2.022651511 | 0.000322619 |
| GO:0008535 | respiratory chain complex IV assembly | 27 | 0.572471632 | 2.020484408 | 0.000416655 |
| GO:0009260 | ribonucleotide biosynthetic process | 201 | 0.382213464 | 2.019851892 | 2.79229464898096e-08 |
| GO:0007062 | sister chromatid cohesion | 56 | 0.478997286 | 2.019379379 | 5.55671985772595e-05 |
| GO:0046390 | ribose phosphate biosynthetic process | 208 | 0.379703876 | 2.017041066 | 2.22713560025509e-08 |
| GO:0051783 | regulation of nuclear division | 143 | 0.400887456 | 2.016197124 | 2.01286452852618e-07 |
| GO:0010824 | regulation of centrosome duplication | 44 | 0.504956917 | 2.014733338 | 0.000177943 |
| GO:0000002 | mitochondrial genome maintenance | 31 | 0.550050727 | 2.014419576 | 0.000398819 |
| GO:0043455 | regulation of secondary metabolic process | 15 | 0.677070623 | 2.014169816 | 0.000895389 |
| GO:0048021 | regulation of melanin biosynthetic process | 15 | 0.677070623 | 2.014169816 | 0.000895389 |
| GO:1900376 | regulation of secondary metabolite biosynthetic process | 15 | 0.677070623 | 2.014169816 | 0.000895389 |
| GO:1904874 | positive regulation of telomerase RNA localization to Cajal boc | 13 | 0.70543743 | 2.01260313 | 0.000848382 |
| GO:1904872 | regulation of telomerase RNA localization to Cajal body | 16 | 0.663205939 | 2.011395485 | 0.000803213 |
| GO:0000245 | spliceosomal complex assembly | 73 | 0.449134907 | 2.006143903 | 6.67756981470048e-05 |
| GO:1903008 | organelle disassembly | 160 | 0.391444901 | 2.005382288 | 2.13421524645102e-07 |
| GO:0036297 | interstrand cross-link repair | 39 | 0.51699414 | 2.003004362 | 0.000275199 |
| GO:0090670 | RNA localization to Cajal body | 17 | 0.647583684 | 2.001108148 | 0.000887762 |
| GO:0090671 | telomerase RNA localization to Cajal body | 17 | 0.647583684 | 2.001108148 | 0.000887762 |
| GO:0090672 | telomerase RNA localization | 17 | 0.647583684 | 2.001108148 | 0.000887762 |
| GO:0090685 | RNA localization to nucleus | 17 | 0.647583684 | 2.001108148 | 0.000887762 |
| GO:0046605 | regulation of centrosome cycle | 51 | 0.484548946 | 2.000109527 | 0.000200178 |
| GO:0090232 | positive regulation of spindle checkpoint | 13 | 0.699708246 | 1.996257847 | 0.001018058 |
| GO:0006301 | postreplication repair | 35 | 0.528287987 | 1.995830873 | 0.000482163 |
| GO:0090307 | mitotic spindle assembly | 69 | 0.451550894 | 1.992110693 | 8.15975913385736e-05 |
| GO:0071816 | tail-anchored membrane protein insertion into ER membrane | 16 | 0.655684759 | 1.988584972 | 0.001029116 |
| GO:0061640 | cytoskeleton-dependent cytokinesis | 114 | 0.410514105 | 1.988389201 | 4.15659596231192e-06 |
| GO:0090184 | positive regulation of kidney development | 10 | 0.759048386 | 1.987953083 | 0.000845845 |
| GO:0001502 | cartilage condensation | 18 | 0.631351991 | 1.985033759 | 0.000997851 |
| GO:1905820 | positive regulation of chromosome separation | 29 | 0.550150388 | 1.980282924 | 0.000449855 |
| GO:0051225 | spindle assembly | 122 | 0.403839927 | 1.979001427 | 2.41240453757381e-06 |
| GO:0009152 | purine ribonucleotide biosynthetic process | 186 | 0.378028516 | 1.978228086 | 2.69960271088376e-07 |
| GO:0062125 | regulation of mitochondrial gene expression | 29 | 0.549210485 | 1.976899712 | 0.000449855 |
| GO:0033617 | mitochondrial cytochrome c oxidase assembly | 23 | 0.585036272 | 1.975132021 | 0.000992702 |
| GO:0061709 | reticulophagy | 19 | 0.617996467 | 1.97376222 | 0.00108721 |
| GO:0006450 | regulation of translational fidelity | 12 | 0.709963223 | 1.973726361 | 0.001081211 |
| GO:0006177 | GMP biosynthetic process | 13 | 0.691520538 | 1.972898432 | 0.001284693 |
| GO:0006298 | mismatch repair | 28 | 0.55321714 | 1.972507012 | 0.000672194 |
| GO:0045022 | early endosome to late endosome transport | 42 | 0.50042411 | 1.972277315 | 0.000431046 |
| GO:0006167 | AMP biosynthetic process | 11 | 0.728173458 | 1.970867205 | 0.001327518 |
| GO:2000232 | regulation of rRNA processing | 16 | 0.649098234 | 1.968609114 | 0.00128012 |
| GO:0006271 | DNA strand elongation involved in DNA replication | 15 | 0.661019622 | 1.96642082 | 0.001417699 |
| GO:0032790 | ribosome disassembly | 31 | 0.536284008 | 1.964002502 | 0.000486228 |
| GO:0007095 | mitotic G2 DNA damage checkpoint signaling | 37 | 0.513226722 | 1.963966704 | 0.000594477 |

| | | | | | |
|---|---|---|---|---|---|
| GO:0006457 | protein folding | 196 | 0.372452553 | 1.962996808 | 1.32873644195224e-07 |
| GO:0042558 | pteridine-containing compound metabolic process | 32 | 0.529369209 | 1.954225134 | 0.000605694 |
| GO:0000963 | mitochondrial RNA processing | 21 | 0.593682874 | 1.952030828 | 0.001503878 |
| GO:1900037 | regulation of cellular response to hypoxia | 17 | 0.630910121 | 1.94958492 | 0.001699429 |
| GO:0051443 | positive regulation of ubiquitin-protein transferase activity | 16 | 0.642775274 | 1.949432607 | 0.001631526 |
| GO:0006903 | vesicle targeting | 62 | 0.451625792 | 1.948218175 | 0.00021581 |
| GO:0030488 | tRNA methylation | 39 | 0.502680641 | 1.947549184 | 0.000666707 |
| GO:0051168 | nuclear export | 154 | 0.382388938 | 1.946548692 | 1.09707556137093e-06 |
| GO:0001833 | inner cell mass cell proliferation | 14 | 0.667632635 | 1.946331583 | 0.001672323 |
| GO:0032511 | late endosome to vacuole transport via multivesicular body so | 26 | 0.556786902 | 1.94564464 | 0.001239772 |
| GO:0009451 | RNA modification | 168 | 0.376939247 | 1.943014289 | 9.32832807545732e-07 |
| GO:0051294 | establishment of spindle orientation | 42 | 0.49200915 | 1.939112178 | 0.00064917 |
| GO:0002098 | tRNA wobble uridine modification | 17 | 0.626993356 | 1.937481666 | 0.00197844 |
| GO:0090267 | positive regulation of mitotic cell cycle spindle assembly checl | 12 | 0.696413885 | 1.936058657 | 0.001681884 |
| GO:0006383 | transcription by RNA polymerase III | 60 | 0.451918691 | 1.935962983 | 0.000115751 |
| GO:0007064 | mitotic sister chromatid cohesion | 30 | 0.53226339 | 1.933430379 | 0.000964101 |
| GO:0006353 | DNA-templated transcription termination | 21 | 0.587689267 | 1.932323832 | 0.00192602 |
| GO:0006998 | nuclear envelope organization | 53 | 0.46383262 | 1.932013697 | 0.000276753 |
| GO:0044774 | mitotic DNA integrity checkpoint signaling | 85 | 0.419919351 | 1.931341556 | 3.9631208340744e-05 |
| GO:1901991 | negative regulation of mitotic cell cycle phase transition | 173 | 0.37273496 | 1.930292138 | 5.64327126290667e-07 |
| GO:0000281 | mitotic cytokinesis | 86 | 0.418760907 | 1.9301624 | 4.63316648800569e-05 |
| GO:0010948 | negative regulation of cell cycle process | 281 | 0.350073154 | 1.930096239 | 7.17672322521571e-09 |
| GO:0002097 | tRNA wobble base modification | 20 | 0.594696667 | 1.928965653 | 0.001749302 |
| GO:0034501 | protein localization to kinetochore | 15 | 0.648103596 | 1.927997842 | 0.002014625 |
| GO:1903083 | protein localization to condensed chromosome | 15 | 0.648103596 | 1.927997842 | 0.002014625 |
| GO:0007099 | centriole replication | 42 | 0.489028595 | 1.927365181 | 0.000741909 |
| GO:0042440 | pigment metabolic process | 76 | 0.427766883 | 1.924528431 | 5.34497200946983e-05 |
| GO:1901988 | negative regulation of cell cycle phase transition | 244 | 0.355225632 | 1.924341073 | 2.85413288041654e-08 |
| GO:0033119 | negative regulation of RNA splicing | 23 | 0.569772553 | 1.923600412 | 0.001663447 |
| GO:0070129 | regulation of mitochondrial translation | 24 | 0.562429912 | 1.922720083 | 0.001809735 |
| GO:0002183 | cytoplasmic translational initiation | 36 | 0.504695596 | 1.918805368 | 0.000886341 |
| GO:0048193 | Golgi vesicle transport | 275 | 0.34896742 | 1.918251985 | 1.23210630035218e-08 |
| GO:1901976 | regulation of cell cycle checkpoint | 44 | 0.480775189 | 1.918250387 | 0.000687178 |
| GO:0045842 | positive regulation of mitotic metaphase/anaphase transition | 14 | 0.6577105 | 1.917405848 | 0.002311741 |
| GO:0051256 | mitotic spindle midzone assembly | 10 | 0.731123128 | 1.914816634 | 0.002067621 |
| GO:0006412 | translation | 436 | 0.331670747 | 1.912531684 | 1e-10 |
| GO:0036257 | multivesicular body organization | 31 | 0.522175808 | 1.912334842 | 0.001086863 |
| GO:0031126 | sno(s)RNA 3'-end processing | 12 | 0.687307775 | 1.910743302 | 0.00225853 |
| GO:0001732 | formation of cytoplasmic translation initiation complex | 11 | 0.704001064 | 1.905442439 | 0.002583918 |
| GO:0042790 | nucleolar large rRNA transcription by RNA polymerase I | 20 | 0.587103319 | 1.904335774 | 0.002323699 |
| GO:0000132 | establishment of mitotic spindle orientation | 35 | 0.504033497 | 1.904199297 | 0.001114631 |
| GO:0043144 | sno(s)RNA processing | 14 | 0.652559607 | 1.902389588 | 0.002656042 |
| GO:0042770 | signal transduction in response to DNA damage | 180 | 0.365220961 | 1.901577929 | 1.18992356080155e-06 |
| GO:0098534 | centriole assembly | 47 | 0.469371857 | 1.901012533 | 0.000638264 |
| GO:0051299 | centrosome separation | 14 | 0.651964095 | 1.900653508 | 0.002705228 |

| | | | | | |
|---|---|---|---|---|---|
| GO:0010972 | negative regulation of G2/M transition of mitotic cell cycle | 65 | 0.436218097 | 1.900590854 | 0.000140548 |
| GO:0072697 | protein localization to cell cortex | 11 | 0.702000224 | 1.900026984 | 0.002749858 |
| GO:0045324 | late endosome to vacuole transport | 36 | 0.499734994 | 1.899945621 | 0.001122699 |
| GO:1904925 | positive regulation of autophagy of mitochondrion in response t | 13 | 0.665884359 | 1.899758771 | 0.002569386 |
| GO:0007030 | Golgi organization | 133 | 0.382303937 | 1.899509188 | 8.01449197804816e-06 |
| GO:1903405 | protein localization to nuclear body | 10 | 0.725276473 | 1.899504204 | 0.002279082 |
| GO:1904867 | protein localization to Cajal body | 10 | 0.725276473 | 1.899504204 | 0.002279082 |
| GO:0045739 | positive regulation of DNA repair | 124 | 0.38637899 | 1.898893582 | 1.34984446037134e-05 |
| GO:2000779 | regulation of double-strand break repair | 129 | 0.38336281 | 1.895584478 | 9.40600800031344e-06 |
| GO:0044380 | protein localization to cytoskeleton | 60 | 0.442219431 | 1.89441257 | 0.000192279 |
| GO:0090235 | regulation of metaphase plate congression | 13 | 0.663500802 | 1.892958517 | 0.002787541 |
| GO:0006888 | endoplasmic reticulum to Golgi vesicle-mediated transport | 122 | 0.386044931 | 1.891797763 | 1.56079901824466e-05 |
| GO:0003197 | endocardial cushion development | 51 | 0.458267399 | 1.891625186 | 0.000520952 |
| GO:0006779 | porphyrin-containing compound biosynthetic process | 33 | 0.508325943 | 1.891592982 | 0.000957021 |
| GO:0033014 | tetrapyrrole biosynthetic process | 33 | 0.508325943 | 1.891592982 | 0.000957021 |
| GO:0006890 | retrograde vesicle-mediated transport, Golgi to endoplasmic re | 49 | 0.462227959 | 1.891120598 | 0.000516429 |
| GO:0070507 | regulation of microtubule cytoskeleton organization | 151 | 0.372665787 | 1.890184355 | 5.41576659267266e-06 |
| GO:0000022 | mitotic spindle elongation | 11 | 0.69800726 | 1.889219666 | 0.003034326 |
| GO:0036503 | ERAD pathway | 110 | 0.39207745 | 1.887881547 | 3.13247958126398e-05 |
| GO:0051028 | mRNA transport | 116 | 0.388375206 | 1.886196876 | 2.25292911759511e-05 |
| GO:0031167 | rRNA methylation | 23 | 0.557453351 | 1.882009742 | 0.002441511 |
| GO:1902749 | regulation of cell cycle G2/M phase transition | 111 | 0.390112373 | 1.881758875 | 3.01972456323521e-05 |
| GO:0006405 | RNA export from nucleus | 81 | 0.412868005 | 1.880832447 | 0.000138143 |
| GO:0000910 | cytokinesis | 179 | 0.361363972 | 1.880620124 | 1.95881018969103e-06 |
| GO:0006368 | transcription elongation by RNA polymerase II | 110 | 0.39044648 | 1.880028307 | 3.5582409290459e-05 |
| GO:0071985 | multivesicular body sorting pathway | 47 | 0.46395628 | 1.879078794 | 0.000765917 |
| GO:0019081 | viral translation | 19 | 0.587901275 | 1.877643948 | 0.003106314 |
| GO:0032259 | methylation | 239 | 0.347308495 | 1.877531665 | 9.59723173199043e-08 |
| GO:0043328 | protein transport to vacuole involved in ubiquitin-dependent prc | 15 | 0.630814764 | 1.87656651 | 0.003830274 |
| GO:0036258 | multivesicular body assembly | 30 | 0.516579005 | 1.876457333 | 0.00215505 |
| GO:0006403 | RNA localization | 178 | 0.360164929 | 1.872899473 | 8.35910100800159e-07 |
| GO:0000463 | maturation of LSU-rRNA from tricistronic rRNA transcript (SS | 12 | 0.673520562 | 1.872414294 | 0.003243633 |
| GO:1902750 | negative regulation of cell cycle G2/M phase transition | 66 | 0.427709094 | 1.868642089 | 0.000268979 |
| GO:1902570 | protein localization to nucleolus | 16 | 0.615629733 | 1.867104607 | 0.003915663 |
| GO:0002091 | negative regulation of receptor internalization | 16 | 0.615366785 | 1.866307131 | 0.003965863 |
| GO:1905508 | protein localization to microtubule organizing center | 37 | 0.487125018 | 1.864083212 | 0.001456469 |
| GO:1903241 | U2-type prespliceosome assembly | 25 | 0.539409416 | 1.862788936 | 0.003337985 |
| GO:0072528 | pyrimidine-containing compound biosynthetic process | 40 | 0.478012951 | 1.862756335 | 0.001433539 |
| GO:0010833 | telomere maintenance via telomere lengthening | 77 | 0.412545828 | 1.861508996 | 0.0003014 |
| GO:0001832 | blastocyst growth | 21 | 0.565951283 | 1.860849285 | 0.003561817 |
| GO:0009226 | nucleotide-sugar biosynthetic process | 18 | 0.591282769 | 1.859052122 | 0.004144919 |
| GO:0016024 | CDP-diacylglycerol biosynthetic process | 13 | 0.65046974 | 1.855781079 | 0.003975276 |
| GO:0046341 | CDP-diacylglycerol metabolic process | 13 | 0.65046974 | 1.855781079 | 0.003975276 |
| GO:0009304 | tRNA transcription | 13 | 0.650452364 | 1.855731504 | 0.003975276 |
| GO:2000234 | positive regulation of rRNA processing | 10 | 0.708431338 | 1.855386676 | 0.003947276 |

| | | | | | |
|---|---|---|---|---|---|
| GO:0046189 | phenol-containing compound biosynthetic process | 43 | 0.467201829 | 1.851881611 | 0.001397819 |
| GO:0006099 | tricarboxylic acid cycle | 30 | 0.509753646 | 1.851664429 | 0.002949016 |
| GO:0032506 | cytokinetic process | 41 | 0.472270472 | 1.851555594 | 0.001382191 |
| GO:0009220 | pyrimidine ribonucleotide biosynthetic process | 23 | 0.5483496 | 1.851274706 | 0.003407384 |
| GO:0010389 | regulation of G2/M transition of mitotic cell cycle | 101 | 0.39015132 | 1.85099459 | 0.000123799 |
| GO:0031109 | microtubule polymerization or depolymerization | 128 | 0.374740394 | 1.850562342 | 3.02508767241991e-05 |
| GO:1901992 | positive regulation of mitotic cell cycle phase transition | 86 | 0.401385966 | 1.850077423 | 0.000141193 |
| GO:0072344 | rescue of stalled ribosome | 24 | 0.540652678 | 1.848272538 | 0.003430393 |
| GO:0044773 | mitotic DNA damage checkpoint signaling | 81 | 0.405655043 | 1.847973582 | 0.000221691 |
| GO:0060236 | regulation of mitotic spindle organization | 39 | 0.476794505 | 1.84725783 | 0.001545548 |
| GO:0048753 | pigment granule organization | 39 | 0.476118525 | 1.84463886 | 0.001575853 |
| GO:0000422 | autophagy of mitochondrion | 93 | 0.394112025 | 1.844494711 | 0.00017043 |
| GO:0006085 | acetyl-CoA biosynthetic process | 18 | 0.586411838 | 1.843737427 | 0.004707809 |
| GO:0007100 | mitotic centrosome separation | 13 | 0.645612268 | 1.841922779 | 0.004532784 |
| GO:0046785 | microtubule polymerization | 88 | 0.397598311 | 1.841108355 | 0.000210736 |
| GO:0006289 | nucleotide-excision repair | 79 | 0.405148335 | 1.838071919 | 0.000151516 |
| GO:0030174 | regulation of DNA-templated DNA replication initiation | 15 | 0.61780465 | 1.837863637 | 0.004924638 |
| GO:0071539 | protein localization to centrosome | 36 | 0.483348799 | 1.837646844 | 0.001890862 |
| GO:0046349 | amino sugar biosynthetic process | 11 | 0.678731272 | 1.837047465 | 0.004930779 |
| GO:0045039 | protein insertion into mitochondrial inner membrane | 13 | 0.643579017 | 1.836121943 | 0.004944855 |
| GO:2000786 | positive regulation of autophagosome assembly | 18 | 0.583932537 | 1.835942256 | 0.004963668 |
| GO:0072698 | protein localization to microtubule cytoskeleton | 56 | 0.435199212 | 1.834733389 | 0.000936392 |
| GO:0044772 | mitotic cell cycle phase transition | 412 | 0.319733885 | 1.832158108 | 1.0013114018382e-09 |
| GO:0033120 | positive regulation of RNA splicing | 43 | 0.462204548 | 1.832073527 | 0.001770571 |
| GO:0042797 | tRNA transcription by RNA polymerase III | 12 | 0.658335849 | 1.830200181 | 0.005526189 |
| GO:0071763 | nuclear membrane organization | 44 | 0.458662879 | 1.830024232 | 0.001530533 |
| GO:0090231 | regulation of spindle checkpoint | 22 | 0.548959697 | 1.829788478 | 0.004498869 |
| GO:0033313 | meiotic cell cycle checkpoint signaling | 10 | 0.698391182 | 1.829091437 | 0.005333521 |
| GO:0019348 | dolichol metabolic process | 20 | 0.563899864 | 1.829072753 | 0.004804052 |
| GO:0006221 | pyrimidine nucleotide biosynthetic process | 31 | 0.498912758 | 1.827139895 | 0.002774361 |
| GO:0044839 | cell cycle G2/M phase transition | 147 | 0.361505869 | 1.825180315 | 2.18651418547608e-05 |
| GO:1901978 | positive regulation of cell cycle checkpoint | 19 | 0.570676625 | 1.822631718 | 0.005203075 |
| GO:2000001 | regulation of DNA damage checkpoint | 21 | 0.554252048 | 1.822382169 | 0.004881009 |
| GO:0043043 | peptide biosynthetic process | 461 | 0.313795075 | 1.819107674 | 2.80756802302322e-10 |
| GO:0033233 | regulation of protein sumoylation | 21 | 0.553235902 | 1.81904108 | 0.004933777 |
| GO:0048668 | collateral sprouting | 27 | 0.515340169 | 1.818844322 | 0.004305436 |
| GO:0051987 | positive regulation of attachment of spindle microtubules to kin | 11 | 0.671886921 | 1.818522611 | 0.005855301 |
| GO:0048026 | positive regulation of mRNA splicing, via spliceosome | 29 | 0.505204332 | 1.818498237 | 0.003598842 |
| GO:0045943 | positive regulation of transcription by RNA polymerase I | 32 | 0.492253365 | 1.817207881 | 0.002884255 |
| GO:0048025 | negative regulation of mRNA splicing, via spliceosome | 20 | 0.559676046 | 1.815372323 | 0.005274014 |
| GO:2000781 | positive regulation of double-strand break repair | 84 | 0.39437152 | 1.810592331 | 0.000232779 |
| GO:0010458 | exit from mitosis | 29 | 0.502998434 | 1.810558041 | 0.003880001 |
| GO:0031468 | nuclear membrane reassembly | 25 | 0.52376718 | 1.808770256 | 0.004924895 |
| GO:0006091 | generation of precursor metabolites and energy | 458 | 0.311898914 | 1.806759864 | 5.18111045219693e-10 |
| GO:0070979 | protein K11-linked ubiquitination | 30 | 0.497156055 | 1.805904068 | 0.003998185 |

| | | | | | |
|---|---|---|---|---|---|
| GO:0009069 | serine family amino acid metabolic process | 31 | 0.492869792 | 1.805009085 | 0.0034322 |
| GO:0009209 | pyrimidine ribonucleoside triphosphate biosynthetic process | 14 | 0.61875969 | 1.803853593 | 0.006615513 |
| GO:0006506 | GPI anchor biosynthetic process | 30 | 0.4965101 | 1.803557654 | 0.004026541 |
| GO:0046320 | regulation of fatty acid oxidation | 34 | 0.480872271 | 1.803422222 | 0.003263974 |
| GO:0051231 | spindle elongation | 13 | 0.631325766 | 1.801163589 | 0.006641619 |
| GO:0043248 | proteasome assembly | 13 | 0.630979039 | 1.80017438 | 0.006690098 |
| GO:0015824 | proline transport | 13 | 0.630513295 | 1.79884562 | 0.006835535 |
| GO:0016074 | sno(s)RNA metabolic process | 16 | 0.592526165 | 1.797035253 | 0.007254016 |
| GO:1902969 | mitotic DNA replication | 17 | 0.581347016 | 1.796429218 | 0.006620165 |
| GO:0007020 | microtubule nucleation | 43 | 0.452721317 | 1.79448416 | 0.002422887 |
| GO:0032438 | melanosome organization | 38 | 0.465638246 | 1.793296979 | 0.002915014 |
| GO:0090224 | regulation of spindle organization | 43 | 0.452075891 | 1.791925839 | 0.002547138 |
| GO:0032784 | regulation of DNA-templated transcription elongation | 100 | 0.378218121 | 1.791908476 | 0.000288974 |
| GO:0031297 | replication fork processing | 49 | 0.436817046 | 1.787156524 | 0.002033439 |
| GO:0045023 | G0 to G1 transition | 36 | 0.470028611 | 1.787004736 | 0.003249919 |
| GO:0070316 | regulation of G0 to G1 transition | 34 | 0.475989266 | 1.785109418 | 0.003875969 |
| GO:0001522 | pseudouridine synthesis | 18 | 0.567756533 | 1.785083284 | 0.007471088 |
| GO:0006760 | folic acid-containing compound metabolic process | 27 | 0.50550239 | 1.78412281 | 0.005722063 |
| GO:0006891 | intra-Golgi vesicle-mediated transport | 32 | 0.482857556 | 1.782522209 | 0.003864902 |
| GO:0090114 | COPII-coated vesicle budding | 41 | 0.454519578 | 1.781962491 | 0.003132967 |
| GO:0007004 | telomere maintenance via telomerase | 67 | 0.406259998 | 1.781680362 | 0.000814332 |
| GO:0090266 | regulation of mitotic cell cycle spindle assembly checkpoint | 21 | 0.541786266 | 1.781394645 | 0.006965332 |
| GO:1903504 | regulation of mitotic spindle checkpoint | 21 | 0.541786266 | 1.781394645 | 0.006965332 |
| GO:0070131 | positive regulation of mitochondrial translation | 15 | 0.598673577 | 1.780951954 | 0.008406705 |
| GO:0098743 | cell aggregation | 22 | 0.534239986 | 1.780724845 | 0.006974577 |
| GO:0006414 | translational elongation | 54 | 0.425376338 | 1.778769316 | 0.001622624 |
| GO:0031055 | chromatin remodeling at centromere | 10 | 0.679132957 | 1.778654011 | 0.007988534 |
| GO:0034080 | CENP-A containing chromatin assembly | 10 | 0.679132957 | 1.778654011 | 0.007988534 |
| GO:0007076 | mitotic chromosome condensation | 19 | 0.556618409 | 1.77773247 | 0.007636354 |
| GO:0036444 | calcium import into the mitochondrion | 12 | 0.639308632 | 1.777303751 | 0.008577607 |
| GO:2000641 | regulation of early endosome to late endosome transport | 19 | 0.556448782 | 1.777190715 | 0.007636354 |
| GO:1901990 | regulation of mitotic cell cycle phase transition | 322 | 0.317301148 | 1.776766431 | 9.23527698844522e-08 |
| GO:0006406 | mRNA export from nucleus | 64 | 0.408835142 | 1.775805235 | 0.000975508 |
| GO:0006783 | heme biosynthetic process | 29 | 0.49328702 | 1.775601513 | 0.005398262 |
| GO:0045931 | positive regulation of mitotic cell cycle | 115 | 0.365899967 | 1.774644097 | 0.000172461 |
| GO:0008053 | mitochondrial fusion | 30 | 0.487707043 | 1.771580822 | 0.005104066 |
| GO:0000086 | G2/M transition of mitotic cell cycle | 133 | 0.35646839 | 1.771143105 | 6.97972151704949e-05 |
| GO:0044786 | cell cycle DNA replication | 45 | 0.441562594 | 1.770820067 | 0.002985544 |
| GO:0051646 | mitochondrion localization | 50 | 0.431049101 | 1.770709593 | 0.002239315 |
| GO:0000291 | nuclear-transcribed mRNA catabolic process, exonucleolytic | 17 | 0.572592207 | 1.769375851 | 0.008547876 |
| GO:0006488 | dolichol-linked oligosaccharide biosynthetic process | 18 | 0.561789736 | 1.766323078 | 0.00877597 |
| GO:0019748 | secondary metabolic process | 53 | 0.423914424 | 1.765741427 | 0.001778246 |
| GO:0019985 | translesion synthesis | 26 | 0.504995476 | 1.76466389 | 0.006942723 |
| GO:0003188 | heart valve formation | 14 | 0.604085384 | 1.76107398 | 0.0094683 |
| GO:0031145 | anaphase-promoting complex-dependent catabolic process | 23 | 0.521363704 | 1.760168035 | 0.008075767 |

| | | | | | |
|---|---|---|---|---|---|
| GO:0000479 | endonucleolytic cleavage of tricistronic rRNA transcript (SSU- | 16 | 0.579074403 | 1.756238253 | 0.010592369 |
| GO:0043161 | proteasome-mediated ubiquitin-dependent protein catabolic pro | 371 | 0.309306876 | 1.755826102 | 7.42822281754469e-08 |
| GO:0032786 | positive regulation of DNA-templated transcription, elongation | 58 | 0.412834787 | 1.753808597 | 0.001861126 |
| GO:0007019 | microtubule depolymerization | 46 | 0.434704726 | 1.752329563 | 0.003356661 |
| GO:0050685 | positive regulation of mRNA processing | 32 | 0.474240471 | 1.750711283 | 0.006172305 |
| GO:1904923 | regulation of autophagy of mitochondrion in response to mitoch | 14 | 0.600142497 | 1.749579388 | 0.010673356 |
| GO:0090148 | membrane fission | 44 | 0.438001712 | 1.747588009 | 0.003560831 |
| GO:0009070 | serine family amino acid biosynthetic process | 16 | 0.576130521 | 1.747309938 | 0.011370482 |
| GO:0065004 | protein-DNA complex assembly | 158 | 0.341434268 | 1.747271522 | 4.5699566629954e-05 |
| GO:1904292 | regulation of ERAD pathway | 24 | 0.510715583 | 1.745929737 | 0.008076279 |
| GO:0018022 | peptidyl-lysine methylation | 21 | 0.530781388 | 1.745210579 | 0.009260725 |
| GO:0050686 | negative regulation of mRNA processing | 22 | 0.52324625 | 1.744080601 | 0.009397045 |
| GO:1901987 | regulation of cell cycle phase transition | 413 | 0.304281333 | 1.743904521 | 3.93381652579642e-08 |
| GO:0006047 | UDP-N-acetylglucosamine metabolic process | 14 | 0.597856645 | 1.742915504 | 0.011288181 |
| GO:0099022 | vesicle tethering | 30 | 0.479467013 | 1.741649167 | 0.006777066 |
| GO:0051438 | regulation of ubiquitin-protein transferase activity | 28 | 0.488444788 | 1.741559869 | 0.007198073 |
| GO:0046321 | positive regulation of fatty acid oxidation | 15 | 0.585262686 | 1.741056836 | 0.011565438 |
| GO:0046599 | regulation of centriole replication | 22 | 0.522061419 | 1.740131333 | 0.009583389 |
| GO:0001843 | neural tube closure | 87 | 0.376699953 | 1.73948449 | 0.000589113 |
| GO:0045292 | mRNA cis splicing, via spliceosome | 28 | 0.487310288 | 1.737514787 | 0.007506162 |
| GO:0001325 | formation of extrachromosomal circular DNA | 13 | 0.608890175 | 1.73715516 | 0.012313659 |
| GO:0090656 | t-circle formation | 13 | 0.608890175 | 1.73715516 | 0.012313659 |
| GO:0090737 | telomere maintenance via telomere trimming | 13 | 0.608890175 | 1.73715516 | 0.012313659 |
| GO:0002181 | cytoplasmic translation | 67 | 0.395933894 | 1.736394542 | 0.00141623 |
| GO:0009199 | ribonucleoside triphosphate metabolic process | 220 | 0.324169666 | 1.734640234 | 5.59259830382628e-06 |
| GO:0048308 | organelle inheritance | 15 | 0.582738551 | 1.733547962 | 0.012361339 |
| GO:0048313 | Golgi inheritance | 15 | 0.582738551 | 1.733547962 | 0.012361339 |
| GO:0031124 | mRNA 3'-end processing | 48 | 0.425575869 | 1.733095825 | 0.003786299 |
| GO:0006743 | ubiquinone metabolic process | 20 | 0.53425247 | 1.732908088 | 0.010548027 |
| GO:0033260 | nuclear DNA replication | 41 | 0.441583083 | 1.731244438 | 0.004638019 |
| GO:0006490 | oligosaccharide-lipid intermediate biosynthetic process | 19 | 0.541549233 | 1.729604412 | 0.011156843 |
| GO:1901028 | regulation of mitochondrial outer membrane permeabilization i | 22 | 0.51882437 | 1.729341625 | 0.010435246 |
| GO:0045930 | negative regulation of mitotic cell cycle | 223 | 0.322587262 | 1.729018622 | 7.54112097134415e-06 |
| GO:0033235 | positive regulation of protein sumoylation | 12 | 0.621651311 | 1.728215689 | 0.013214801 |
| GO:0000101 | sulfur amino acid transport | 12 | 0.621618978 | 1.7281258 | 0.013214801 |
| GO:0018205 | peptidyl-lysine modification | 138 | 0.344990996 | 1.725179836 | 0.00011069 |
| GO:0015937 | coenzyme A biosynthetic process | 10 | 0.658474166 | 1.724548489 | 0.013744978 |
| GO:0042762 | regulation of sulfur metabolic process | 13 | 0.603432563 | 1.721584671 | 0.013768028 |
| GO:1904029 | regulation of cyclin-dependent protein kinase activity | 75 | 0.383838608 | 1.721166747 | 0.001667099 |
| GO:0060606 | tube closure | 88 | 0.371285859 | 1.719266604 | 0.00086495 |
| GO:0033866 | nucleoside bisphosphate biosynthetic process | 56 | 0.40769324 | 1.718772413 | 0.003009832 |
| GO:0034030 | ribonucleoside bisphosphate biosynthetic process | 56 | 0.40769324 | 1.718772413 | 0.003009832 |
| GO:0034033 | purine nucleoside bisphosphate biosynthetic process | 56 | 0.40769324 | 1.718772413 | 0.003009832 |
| GO:0009112 | nucleobase metabolic process | 33 | 0.461743908 | 1.718250953 | 0.007627168 |
| GO:0006515 | protein quality control for misfolded or incompletely synthesize | 20 | 0.529328553 | 1.716936809 | 0.01201013 |

| | | | | | |
|---|---|---|---|---|---|
| GO:0042073 | intraciliary transport | 39 | 0.443155518 | 1.716929392 | 0.005424571 |
| GO:0006997 | nucleus organization | 134 | 0.344768064 | 1.715218047 | 0.00017613 |
| GO:0006241 | CTP biosynthetic process | 13 | 0.600890076 | 1.714330991 | 0.014664889 |
| GO:1902108 | regulation of mitochondrial membrane permeability involved ir | 39 | 0.442293503 | 1.713589663 | 0.00572762 |
| GO:0006303 | double-strand break repair via nonhomologous end joining | 65 | 0.393020759 | 1.712381183 | 0.002171401 |
| GO:0050657 | nucleic acid transport | 142 | 0.340770991 | 1.711938644 | 0.000178918 |
| GO:0050658 | RNA transport | 142 | 0.340770991 | 1.711938644 | 0.000178918 |
| GO:0010498 | proteasomal protein catabolic process | 500 | 0.29309495 | 1.711347777 | 6.26320729096646e-09 |
| GO:0039702 | viral budding via host ESCRT complex | 21 | 0.520144787 | 1.710237408 | 0.01232125 |
| GO:0009141 | nucleoside triphosphate metabolic process | 235 | 0.316945429 | 1.709781563 | 5.9231972759974e-06 |
| GO:0170034 | L-amino acid biosynthetic process | 50 | 0.416191142 | 1.709674481 | 0.003926914 |
| GO:0170038 | proteinogenic amino acid biosynthetic process | 50 | 0.416191142 | 1.709674481 | 0.003926914 |
| GO:0000966 | RNA 5'-end processing | 20 | 0.526653927 | 1.708261358 | 0.0127934 |
| GO:0034471 | ncRNA 5'-end processing | 20 | 0.526653927 | 1.708261358 | 0.0127934 |
| GO:0032986 | protein-DNA complex disassembly | 18 | 0.543283093 | 1.70813634 | 0.013509364 |
| GO:0009208 | pyrimidine ribonucleoside triphosphate metabolic process | 16 | 0.561854484 | 1.704013047 | 0.015235944 |
| GO:0006278 | RNA-templated DNA biosynthetic process | 69 | 0.38551716 | 1.700789139 | 0.002007744 |
| GO:0016226 | iron-sulfur cluster assembly | 27 | 0.481388751 | 1.699016005 | 0.010888586 |
| GO:0031163 | metallo-sulfur cluster assembly | 27 | 0.481388751 | 1.699016005 | 0.010888586 |
| GO:0048199 | vesicle targeting, to, from or within Golgi | 31 | 0.463769437 | 1.698436504 | 0.009038126 |
| GO:0000079 | regulation of cyclin-dependent protein serine/threonine kinase a | 72 | 0.381014142 | 1.697205833 | 0.002031341 |
| GO:0043484 | regulation of RNA splicing | 168 | 0.329032139 | 1.696066813 | 0.000118868 |
| GO:0046755 | viral budding | 24 | 0.496102701 | 1.695974209 | 0.011722759 |
| GO:0035795 | negative regulation of mitochondrial membrane permeability | 17 | 0.548689765 | 1.695514554 | 0.015091947 |
| GO:0006284 | base-excision repair | 41 | 0.431973107 | 1.693568133 | 0.006634518 |
| GO:0051236 | establishment of RNA localization | 145 | 0.335790197 | 1.692771572 | 0.000215718 |
| GO:1903715 | regulation of aerobic respiration | 28 | 0.474325353 | 1.691216736 | 0.010671073 |
| GO:1901989 | positive regulation of cell cycle phase transition | 103 | 0.354944192 | 1.690311232 | 0.000763068 |
| GO:0007346 | regulation of mitotic cell cycle | 480 | 0.290481527 | 1.688653897 | 1.12833397697363e-08 |
| GO:0032402 | melanosome transport | 21 | 0.513308179 | 1.687758623 | 0.014774946 |
| GO:0051904 | pigment granule transport | 21 | 0.513308179 | 1.687758623 | 0.014774946 |
| GO:1902751 | positive regulation of cell cycle G2/M phase transition | 30 | 0.464089661 | 1.685791409 | 0.010463336 |
| GO:0016073 | snRNA metabolic process | 55 | 0.401195352 | 1.685157063 | 0.004359546 |
| GO:2001252 | positive regulation of chromosome organization | 102 | 0.354600072 | 1.685131007 | 0.000930154 |
| GO:0002082 | regulation of oxidative phosphorylation | 21 | 0.51204697 | 1.683611765 | 0.015117936 |
| GO:0016180 | snRNA processing | 26 | 0.481505712 | 1.682580899 | 0.012563022 |
| GO:0061512 | protein localization to cilium | 68 | 0.381839849 | 1.677950678 | 0.003079126 |
| GO:0014020 | primary neural tube formation | 93 | 0.358375464 | 1.677243031 | 0.001729408 |
| GO:1904262 | negative regulation of TORC1 signaling | 44 | 0.419236432 | 1.672716202 | 0.006621896 |
| GO:1902410 | mitotic cytokinetic process | 26 | 0.477689125 | 1.669244159 | 0.013995647 |
| GO:0032465 | regulation of cytokinesis | 86 | 0.361979104 | 1.668442411 | 0.001950078 |
| GO:0021915 | neural tube development | 156 | 0.326570827 | 1.667742216 | 0.000218942 |
| GO:0000209 | protein polyubiquitination | 254 | 0.306240221 | 1.66755989 | 3.2536010731588e-05 |
| GO:0016236 | macroautophagy | 327 | 0.297071066 | 1.665679494 | 4.97945795247255e-06 |
| GO:0051656 | establishment of organelle localization | 449 | 0.288058874 | 1.665556199 | 7.77879284806433e-08 |

| | | | | | |
|---|---|---|---|---|---|
| GO:0003203 | endocardial cushion morphogenesis | 40 | 0.427185144 | 1.664686767 | 0.008967242 |
| GO:1902117 | positive regulation of organelle assembly | 80 | 0.365760659 | 1.66291758 | 0.002316837 |
| GO:0006730 | one-carbon metabolic process | 38 | 0.431337968 | 1.661197467 | 0.009556437 |
| GO:0031047 | regulatory ncRNA-mediated gene silencing | 111 | 0.344215977 | 1.660371511 | 0.000790236 |
| GO:0045005 | DNA-templated DNA replication maintenance of fidelity | 57 | 0.391958617 | 1.659258479 | 0.005178385 |
| GO:0031111 | negative regulation of microtubule polymerization or depolyme | 37 | 0.433433151 | 1.658620333 | 0.01061142 |
| GO:0032007 | negative regulation of TOR signaling | 62 | 0.384426742 | 1.658335682 | 0.00434318 |
| GO:0032968 | positive regulation of transcription elongation by RNA polymer | 48 | 0.406958402 | 1.657278899 | 0.00683459 |
| GO:0014850 | response to muscle activity | 26 | 0.474232524 | 1.657165361 | 0.015180318 |
| GO:0046902 | regulation of mitochondrial membrane permeability | 51 | 0.401360672 | 1.656726964 | 0.006218865 |
| GO:0009205 | purine ribonucleoside triphosphate metabolic process | 213 | 0.310480908 | 1.656114267 | 6.04378245998192e-05 |
| GO:0035825 | homologous recombination | 66 | 0.379033202 | 1.655979273 | 0.003956339 |
| GO:0060260 | regulation of transcription initiation by RNA polymerase II | 73 | 0.370401178 | 1.654465183 | 0.003395152 |
| GO:0006220 | pyrimidine nucleotide metabolic process | 51 | 0.400394533 | 1.652738956 | 0.006446781 |
| GO:0006505 | GPI anchor metabolic process | 31 | 0.451102354 | 1.652046563 | 0.013042359 |
| GO:0051321 | meiotic cell cycle | 273 | 0.300656199 | 1.651002316 | 2.41166205039256e-05 |
| GO:0030705 | cytoskeleton-dependent intracellular transport | 185 | 0.315453241 | 1.649044852 | 9.85310906076902e-05 |
| GO:0019068 | virion assembly | 32 | 0.446671371 | 1.648936893 | 0.01312336 |
| GO:0046034 | ATP metabolic process | 193 | 0.312866398 | 1.646379146 | 7.45048546660253e-05 |
| GO:0006901 | vesicle coating | 36 | 0.432813256 | 1.645515446 | 0.012054244 |
| GO:2000785 | regulation of autophagosome assembly | 45 | 0.410119192 | 1.644721055 | 0.008705217 |
| GO:0032204 | regulation of telomere maintenance | 99 | 0.347116509 | 1.641525634 | 0.002115744 |
| GO:1903046 | meiotic cell cycle process | 206 | 0.309028829 | 1.63934015 | 0.000160238 |
| GO:0045786 | negative regulation of cell cycle | 363 | 0.28880839 | 1.637645158 | 3.65110402642985e-06 |
| GO:0050684 | regulation of mRNA processing | 120 | 0.335014653 | 1.636556902 | 0.001049845 |
| GO:0006605 | protein targeting | 300 | 0.294398342 | 1.635435666 | 9.22898627775333e-06 |
| GO:0009124 | nucleoside monophosphate biosynthetic process | 39 | 0.421995958 | 1.634950339 | 0.012031032 |
| GO:0048704 | embryonic skeletal system morphogenesis | 90 | 0.351213329 | 1.63360544 | 0.002461685 |
| GO:0035794 | positive regulation of mitochondrial membrane permeability | 38 | 0.423523647 | 1.631102432 | 0.012020676 |
| GO:0035967 | cellular response to topologically incorrect protein | 100 | 0.344138471 | 1.630447111 | 0.002290903 |
| GO:0001841 | neural tube formation | 100 | 0.344099484 | 1.630262402 | 0.002290903 |
| GO:0034243 | regulation of transcription elongation by RNA polymerase II | 85 | 0.354189857 | 1.629030879 | 0.002910587 |
| GO:0009144 | purine nucleoside triphosphate metabolic process | 219 | 0.304383094 | 1.627477858 | 0.000113055 |
| GO:0002052 | positive regulation of neuroblast proliferation | 37 | 0.424693686 | 1.625176987 | 0.013732426 |
| GO:0018958 | phenol-containing compound metabolic process | 104 | 0.339844494 | 1.621323222 | 0.001868445 |
| GO:0010970 | transport along microtubule | 159 | 0.316575113 | 1.621011643 | 0.00060375 |
| GO:0080135 | regulation of cellular response to stress | 486 | 0.277958694 | 1.618267217 | 3.18915091397308e-07 |
| GO:0010639 | negative regulation of organelle organization | 333 | 0.28765652 | 1.616907826 | 1.50714239033916e-05 |
| GO:0031647 | regulation of protein stability | 292 | 0.291788589 | 1.616697181 | 1.32579464070547e-05 |
| GO:1902686 | mitochondrial outer membrane permeabilization involved in pr | 36 | 0.424995863 | 1.61579445 | 0.01509735 |
| GO:1902017 | regulation of cilium assembly | 68 | 0.367642247 | 1.615560972 | 0.005155747 |
| GO:0010569 | regulation of double-strand break repair via homologous recon | 71 | 0.363742334 | 1.615173191 | 0.005030035 |
| GO:0006986 | response to unfolded protein | 135 | 0.324300456 | 1.614809012 | 0.001223391 |
| GO:0071364 | cellular response to epidermal growth factor stimulus | 39 | 0.416788585 | 1.614775271 | 0.013970544 |
| GO:0090068 | positive regulation of cell cycle process | 238 | 0.298622716 | 1.614133335 | 4.60516966595045e-05 |

| | | | | | |
|---|---|---|---|---|---|
| GO:0001838 | embryonic epithelial tube formation | 121 | 0.329545337 | 1.612627059 | 0.001329665 |
| GO:0090329 | regulation of DNA-templated DNA replication | 48 | 0.39549475 | 1.610594841 | 0.010749238 |
| GO:0061982 | meiosis I cell cycle process | 130 | 0.325037741 | 1.60941672 | 0.001382928 |
| GO:0006352 | DNA-templated transcription initiation | 172 | 0.310872049 | 1.608486466 | 0.000259309 |
| GO:0031110 | regulation of microtubule polymerization or depolymerization | 89 | 0.345939234 | 1.605498354 | 0.00320386 |
| GO:1901293 | nucleoside phosphate biosynthetic process | 263 | 0.29313101 | 1.602203956 | 2.57600895155759e-05 |
| GO:0071806 | protein transmembrane transport | 64 | 0.368666714 | 1.601330741 | 0.006863394 |
| GO:1902115 | regulation of organelle assembly | 209 | 0.300487937 | 1.597736773 | 0.00021733 |
| GO:0009165 | nucleotide biosynthetic process | 261 | 0.29244977 | 1.597349217 | 7.02576333023921e-05 |
| GO:0045665 | negative regulation of neuron differentiation | 71 | 0.359611807 | 1.596831865 | 0.005789969 |
| GO:0031507 | heterochromatin formation | 93 | 0.339442272 | 1.588633269 | 0.003539254 |
| GO:0140013 | meiotic nuclear division | 185 | 0.303746681 | 1.587848327 | 0.000245427 |
| GO:0006611 | protein export from nucleus | 59 | 0.371352209 | 1.583728649 | 0.009931973 |
| GO:0045787 | positive regulation of cell cycle | 300 | 0.285039412 | 1.583445124 | 3.17751861658401e-05 |
| GO:1901873 | regulation of post-translational protein modification | 227 | 0.294630099 | 1.583017636 | 0.000140364 |
| GO:0045814 | negative regulation of gene expression, epigenetic | 121 | 0.322546778 | 1.578379679 | 0.001925722 |
| GO:0061077 | chaperone-mediated protein folding | 68 | 0.359009754 | 1.577626488 | 0.007411386 |
| GO:1901796 | regulation of signal transduction by p53 class mediator | 93 | 0.336984367 | 1.577129959 | 0.004102317 |
| GO:0006417 | regulation of translation | 330 | 0.280624397 | 1.576615698 | 1.65164384568452e-05 |
| GO:0007032 | endosome organization | 93 | 0.336660882 | 1.575616009 | 0.004102317 |
| GO:0070918 | regulatory ncRNA processing | 67 | 0.358566759 | 1.572518476 | 0.008461974 |
| GO:1903432 | regulation of TORC1 signaling | 88 | 0.338612128 | 1.567968479 | 0.004560645 |
| GO:0098732 | macromolecule deacylation | 52 | 0.377891468 | 1.566770314 | 0.012790317 |
| GO:0015931 | nucleobase-containing compound transport | 210 | 0.294435361 | 1.566177097 | 0.000248646 |
| GO:2000142 | regulation of DNA-templated transcription initiation | 79 | 0.345048433 | 1.565411434 | 0.006040927 |
| GO:2000144 | positive regulation of DNA-templated transcription initiation | 67 | 0.356419954 | 1.563103519 | 0.009134683 |
| GO:0035966 | response to topologically incorrect protein | 149 | 0.308785466 | 1.561804838 | 0.001505086 |
| GO:0007127 | meiosis I | 122 | 0.318305303 | 1.559842421 | 0.00221137 |
| GO:0048024 | regulation of mRNA splicing, via spliceosome | 98 | 0.330374035 | 1.55952632 | 0.004345847 |
| GO:0006164 | purine nucleotide biosynthetic process | 226 | 0.290338904 | 1.559189695 | 0.00046036 |
| GO:0072384 | organelle transport along microtubule | 86 | 0.336955788 | 1.553104367 | 0.006084243 |
| GO:0051650 | establishment of vesicle localization | 187 | 0.296031126 | 1.550256386 | 0.000909863 |
| GO:1904356 | regulation of telomere maintenance via telomere lengthening | 57 | 0.365632707 | 1.547814342 | 0.01395474 |
| GO:0032984 | protein-containing complex disassembly | 233 | 0.286692678 | 1.5454652 | 0.000272162 |
| GO:0051261 | protein depolymerization | 113 | 0.318726779 | 1.541614801 | 0.003411608 |
| GO:0034620 | cellular response to unfolded protein | 88 | 0.332726755 | 1.540715824 | 0.006093965 |
| GO:1903320 | regulation of protein modification by small protein conjugation ( | 222 | 0.287254537 | 1.539009167 | 0.000380753 |
| GO:0010608 | post-transcriptional regulation of gene expression | 475 | 0.264786407 | 1.538010184 | 2.90557054912897e-06 |
| GO:0007033 | vacuole organization | 217 | 0.287521032 | 1.535039836 | 0.00073166 |
| GO:0006900 | vesicle budding from membrane | 81 | 0.336901237 | 1.534763582 | 0.008013432 |
| GO:0022411 | cellular component disassembly | 495 | 0.26297622 | 1.534276045 | 2.49630594874759e-06 |
| GO:0060261 | positive regulation of transcription initiation by RNA polymeras | 62 | 0.35487874 | 1.530871849 | 0.014684085 |
| GO:0032886 | regulation of microtubule-based process | 240 | 0.28254436 | 1.528130541 | 0.000434691 |
| GO:0042147 | retrograde transport, endosome to Golgi | 89 | 0.32888554 | 1.526352439 | 0.007159244 |
| GO:0006413 | translational initiation | 105 | 0.319459764 | 1.52603108 | 0.005343707 |

| | | | | | |
|---|---|---|---|---|---|
| GO:1901607 | alpha-amino acid biosynthetic process | 61 | 0.354758198 | 1.524750725 | 0.015283918 |
| GO:0071479 | cellular response to ionizing radiation | 65 | 0.348430357 | 1.518101966 | 0.014954646 |
| GO:0000045 | autophagosome assembly | 107 | 0.315896253 | 1.512493472 | 0.006273254 |
| GO:0072522 | purine-containing compound biosynthetic process | 234 | 0.279447723 | 1.507068145 | 0.000730746 |
| GO:0072594 | establishment of protein localization to organelle | 417 | 0.26253425 | 1.506642895 | 1.82902144009797e-05 |
| GO:0006367 | transcription initiation at RNA polymerase II promoter | 139 | 0.30052443 | 1.505040036 | 0.003464832 |
| GO:0009259 | ribonucleotide metabolic process | 422 | 0.261400251 | 1.502296539 | 1.90470482051403e-05 |
| GO:0072331 | signal transduction by p53 class mediator | 157 | 0.292954921 | 1.498055114 | 0.003191659 |
| GO:0070828 | heterochromatin organization | 102 | 0.314118081 | 1.492752429 | 0.007864028 |
| GO:0099111 | microtubule-based transport | 194 | 0.282818129 | 1.48878358 | 0.001683292 |
| GO:0007224 | smoothened signaling pathway | 143 | 0.295640996 | 1.486877469 | 0.003883006 |
| GO:0044782 | cilium organization | 369 | 0.261257818 | 1.482930184 | 0.000117634 |
| GO:0009150 | purine ribonucleotide metabolic process | 402 | 0.259146154 | 1.482506767 | 6.47678877096957e-05 |
| GO:0019693 | ribose phosphate metabolic process | 436 | 0.256714043 | 1.480304628 | 5.22534552368454e-05 |
| GO:1905037 | autophagosome organization | 113 | 0.305100355 | 1.47570664 | 0.007709349 |
| GO:0042303 | molting cycle | 114 | 0.302727803 | 1.466309408 | 0.008779749 |
| GO:0042633 | hair cycle | 114 | 0.302727803 | 1.466309408 | 0.008779749 |
| GO:0040029 | epigenetic regulation of gene expression | 190 | 0.279058478 | 1.465196921 | 0.002882728 |
| GO:0060271 | cilium assembly | 346 | 0.259449248 | 1.463489856 | 0.000199137 |
| GO:0050821 | protein stabilization | 190 | 0.277409955 | 1.456541347 | 0.003066732 |
| GO:0030326 | embryonic limb morphogenesis | 121 | 0.297522313 | 1.455922688 | 0.008895002 |
| GO:0035113 | embryonic appendage morphogenesis | 121 | 0.297522313 | 1.455922688 | 0.008895002 |
| GO:0090150 | establishment of protein localization to membrane | 248 | 0.267128423 | 1.450530415 | 0.001310515 |
| GO:0140694 | non-membrane-bounded organelle assembly | 370 | 0.255124092 | 1.448194005 | 0.00018381 |
| GO:0006275 | regulation of DNA replication | 126 | 0.293398343 | 1.44561973 | 0.009317788 |
| GO:0048706 | embryonic skeletal system development | 124 | 0.292160873 | 1.435850344 | 0.010403604 |
| GO:0072175 | epithelial tube formation | 133 | 0.288487894 | 1.43337631 | 0.008735744 |
| GO:0120034 | positive regulation of plasma membrane bounded cell projectic | 108 | 0.297904691 | 1.429683163 | 0.014889738 |
| GO:0009952 | anterior/posterior pattern specification | 211 | 0.267978491 | 1.426629757 | 0.003014469 |
| GO:0043467 | regulation of generation of precursor metabolites and energy | 128 | 0.288354719 | 1.42396815 | 0.011401268 |
| GO:0006310 | DNA recombination | 310 | 0.25503778 | 1.423024545 | 0.000746819 |
| GO:0033865 | nucleoside bisphosphate metabolic process | 117 | 0.291889619 | 1.420183132 | 0.014016586 |
| GO:0033875 | ribonucleoside bisphosphate metabolic process | 117 | 0.291889619 | 1.420183132 | 0.014016586 |
| GO:0034032 | purine nucleoside bisphosphate metabolic process | 117 | 0.291889619 | 1.420183132 | 0.014016586 |
| GO:0016482 | cytosolic transport | 129 | 0.286289937 | 1.415595738 | 0.012555883 |
| GO:0010212 | response to ionizing radiation | 131 | 0.283932095 | 1.407260046 | 0.012676462 |
| GO:0060041 | retina development in camera-type eye | 157 | 0.274683176 | 1.404620663 | 0.009468589 |
| GO:0016331 | morphogenesis of embryonic epithelium | 146 | 0.2780151 | 1.40258093 | 0.010582822 |
| GO:0006790 | sulfur compound metabolic process | 298 | 0.251703284 | 1.397106168 | 0.001490591 |
| GO:0007389 | pattern specification process | 450 | 0.240555058 | 1.39114837 | 0.00021513 |
| GO:0035148 | tube formation | 146 | 0.274094523 | 1.382801693 | 0.013803681 |
| GO:1903311 | regulation of mRNA metabolic process | 281 | 0.248244387 | 1.368672665 | 0.00385424 |
| GO:0006163 | purine nucleotide metabolic process | 488 | 0.233764411 | 1.361105349 | 0.000347546 |
| GO:0051648 | vesicle localization | 205 | 0.256557165 | 1.359483803 | 0.009695382 |
| GO:0031396 | regulation of protein ubiquitination | 179 | 0.260702204 | 1.356753436 | 0.014018967 |

| | | | | | |
|---|---|---|---|---|---|
| GO:0071897 | DNA biosynthetic process | 179 | 0.26048238 | 1.355609427 | 0.014313483 |
| GO:0034248 | regulation of amide metabolic process | 394 | 0.236544469 | 1.351796053 | 0.002487237 |
| GO:0002009 | morphogenesis of an epithelium | 483 | 0.229331157 | 1.334299554 | 0.00099924 |
| GO:0048705 | skeletal system morphogenesis | 219 | 0.249449811 | 1.333760158 | 0.011903114 |
| GO:0060828 | regulation of canonical Wnt signaling pathway | 244 | 0.244400789 | 1.323976746 | 0.011387792 |
| GO:0060562 | epithelial tube morphogenesis | 325 | 0.233083604 | 1.306051908 | 0.008548769 |
| GO:0051052 | regulation of DNA metabolic process | 488 | 0.223891605 | 1.303620428 | 0.003270349 |
| GO:0007018 | microtubule-based movement | 383 | 0.228490613 | 1.30103064 | 0.006082879 |
| GO:0003002 | regionalization | 406 | 0.226574462 | 1.297449892 | 0.005182054 |
| GO:0034504 | protein localization to nucleus | 290 | 0.234206909 | 1.296427996 | 0.011574283 |
| GO:0048562 | embryonic organ morphogenesis | 286 | 0.23422369 | 1.294468872 | 0.011778353 |
| GO:0016197 | endosomal transport | 274 | 0.235028648 | 1.290976084 | 0.013806034 |
| GO:0016050 | vesicle organization | 354 | 0.227508872 | 1.286291293 | 0.010323313 |
| GO:0060070 | canonical Wnt signaling pathway | 299 | 0.231602082 | 1.286093436 | 0.012991868 |
| GO:0007409 | axonogenesis | 438 | 0.221894872 | 1.280590403 | 0.006659603 |
| GO:0043010 | camera-type eye development | 338 | 0.226757922 | 1.275878664 | 0.012136183 |
| GO:0031346 | positive regulation of cell projection organization | 345 | 0.223850301 | 1.262266358 | 0.014836795 |
| GO:0016055 | Wnt signaling pathway | 436 | 0.214384601 | 1.236217985 | 0.014219428 |
| GO:0198738 | cell-cell signaling by wnt | 438 | 0.213479683 | 1.232025014 | 0.014378689 |
| GO:0061564 | axon development | 499 | 0.209585946 | 1.223419049 | 0.014123769 |
| GO:0016032 | viral process | 403 | -0.28194851 | -1.328345431 | 0.013995227 |
| GO:0051051 | negative regulation of transport | 385 | -0.283539626 | -1.331782499 | 0.014023967 |
| GO:0015849 | organic acid transport | 340 | -0.293739968 | -1.366623481 | 0.010703381 |
| GO:0046942 | carboxylic acid transport | 339 | -0.294949434 | -1.37207471 | 0.00991976 |
| GO:0001667 | ameboidal-type cell migration | 414 | -0.291988402 | -1.378258228 | 0.00514637 |
| GO:0051222 | positive regulation of protein transport | 239 | -0.306439004 | -1.382594648 | 0.014755632 |
| GO:0045859 | regulation of protein kinase activity | 434 | -0.293897945 | -1.391143157 | 0.003344948 |
| GO:0010038 | response to metal ion | 344 | -0.299534344 | -1.394968918 | 0.00615063 |
| GO:0006814 | sodium ion transport | 233 | -0.310307574 | -1.396531873 | 0.012840719 |
| GO:0060541 | respiratory system development | 212 | -0.313377573 | -1.396975952 | 0.015170031 |
| GO:0048545 | response to steroid hormone | 323 | -0.302238692 | -1.401043227 | 0.006407698 |
| GO:0043086 | negative regulation of catalytic activity | 389 | -0.298272913 | -1.401900236 | 0.003892358 |
| GO:0009913 | epidermal cell differentiation | 230 | -0.312635249 | -1.405238862 | 0.011763476 |
| GO:0072593 | reactive oxygen species metabolic process | 215 | -0.315045257 | -1.40652352 | 0.012969804 |
| GO:0099177 | regulation of trans-synaptic signaling | 462 | -0.296096656 | -1.407006439 | 0.001819555 |
| GO:0050804 | modulation of chemical synaptic transmission | 461 | -0.296266945 | -1.407563856 | 0.001873496 |
| GO:0071241 | cellular response to inorganic substance | 217 | -0.315449711 | -1.409919732 | 0.012114898 |
| GO:0071695 | anatomical structure maturation | 257 | -0.311298759 | -1.414253477 | 0.008416045 |
| GO:0071466 | cellular response to xenobiotic stimulus | 177 | -0.324871917 | -1.419659979 | 0.015177106 |
| GO:0031589 | cell-substrate adhesion | 340 | -0.305528189 | -1.421468109 | 0.004067505 |
| GO:0042391 | regulation of membrane potential | 424 | -0.300771855 | -1.421740345 | 0.001805054 |
| GO:0003015 | heart process | 239 | -0.317759858 | -1.433672192 | 0.00697371 |
| GO:0007160 | cell-matrix adhesion | 229 | -0.320149163 | -1.438644004 | 0.00724705 |
| GO:0043549 | regulation of kinase activity | 492 | -0.301998041 | -1.440344475 | 0.000666152 |
| GO:0006936 | muscle contraction | 328 | -0.310460188 | -1.44075827 | 0.003010315 |

| | | | | | |
|---|---|---|---|---|---|
| GO:0030003 | intracellular monoatomic cation homeostasis | 498 | -0.302300234 | -1.442557312 | 0.000602486 |
| GO:0030073 | insulin secretion | 190 | -0.329538542 | -1.451199997 | 0.008650147 |
| GO:0007264 | small GTPase-mediated signal transduction | 483 | -0.305839797 | -1.456913446 | 0.000476912 |
| GO:0042886 | amide transport | 296 | -0.316749153 | -1.457555122 | 0.002859546 |
| GO:0045444 | fat cell differentiation | 229 | -0.326840249 | -1.468711522 | 0.00454102 |
| GO:0008016 | regulation of heart contraction | 192 | -0.33313663 | -1.468807118 | 0.006693792 |
| GO:0060048 | cardiac muscle contraction | 133 | -0.348310714 | -1.469300817 | 0.013715883 |
| GO:0050679 | positive regulation of epithelial cell proliferation | 192 | -0.333365559 | -1.469816473 | 0.006610269 |
| GO:0046777 | protein autophosphorylation | 191 | -0.334977907 | -1.476008919 | 0.006267534 |
| GO:0007188 | adenylate cyclase-modulating G protein-coupled receptor sign | 239 | -0.327203923 | -1.476282021 | 0.003533039 |
| GO:0120254 | olefinic compound metabolic process | 152 | -0.346122221 | -1.48495862 | 0.00930066 |
| GO:0010876 | lipid localization | 465 | -0.312927389 | -1.487540347 | 0.000159525 |
| GO:0045598 | regulation of fat cell differentiation | 130 | -0.35407595 | -1.489042474 | 0.011514323 |
| GO:0031334 | positive regulation of protein-containing complex assembly | 186 | -0.339443264 | -1.491301586 | 0.005270913 |
| GO:0045861 | negative regulation of proteolysis | 200 | -0.336655751 | -1.491602668 | 0.004289303 |
| GO:0010469 | regulation of signaling receptor activity | 101 | -0.368522569 | -1.494966653 | 0.014685039 |
| GO:0006887 | exocytosis | 333 | -0.322768018 | -1.499477512 | 0.00066283 |
| GO:0002064 | epithelial cell development | 206 | -0.337496009 | -1.500238543 | 0.003539698 |
| GO:0006898 | receptor-mediated endocytosis | 245 | -0.331808858 | -1.500653272 | 0.002244734 |
| GO:1903034 | regulation of response to wounding | 161 | -0.347773817 | -1.502268205 | 0.006245714 |
| GO:0071248 | cellular response to metal ion | 190 | -0.341462325 | -1.503709165 | 0.004145858 |
| GO:0010952 | positive regulation of peptidase activity | 149 | -0.351455676 | -1.504075249 | 0.007283177 |
| GO:0002027 | regulation of heart rate | 97 | -0.373625542 | -1.506011004 | 0.013972899 |
| GO:0044409 | symbiont entry into host | 158 | -0.349746566 | -1.506969214 | 0.006026764 |
| GO:0043547 | positive regulation of GTPase activity | 206 | -0.339305946 | -1.508284085 | 0.002996944 |
| GO:0050796 | regulation of insulin secretion | 161 | -0.349279016 | -1.508770168 | 0.005755854 |
| GO:0060079 | excitatory postsynaptic potential | 102 | -0.371419498 | -1.50916596 | 0.012848723 |
| GO:0097191 | extrinsic apoptotic signaling pathway | 215 | -0.338073367 | -1.50933281 | 0.002849604 |
| GO:0043087 | regulation of GTPase activity | 269 | -0.330902076 | -1.509963518 | 0.001442526 |
| GO:0050673 | epithelial cell proliferation | 417 | -0.32034071 | -1.512644622 | 0.000204437 |
| GO:0060047 | heart contraction | 229 | -0.336986015 | -1.514303225 | 0.002183406 |
| GO:0009612 | response to mechanical stimulus | 205 | -0.340972454 | -1.515245684 | 0.002820492 |
| GO:0009306 | protein secretion | 339 | -0.325738943 | -1.515304366 | 0.000418834 |
| GO:0120162 | positive regulation of cold-induced thermogenesis | 98 | -0.375886819 | -1.517585071 | 0.012038343 |
| GO:0050678 | regulation of epithelial cell proliferation | 343 | -0.326481626 | -1.520040133 | 0.000352338 |
| GO:0044344 | cellular response to fibroblast growth factor stimulus | 106 | -0.372379689 | -1.522093991 | 0.010359251 |
| GO:0006941 | striated muscle contraction | 176 | -0.348728382 | -1.522840665 | 0.003647123 |
| GO:0051651 | maintenance of location in cell | 213 | -0.341506538 | -1.523107592 | 0.002195506 |
| GO:0035592 | establishment of protein localization to extracellular region | 340 | -0.32744089 | -1.523416826 | 0.000363761 |
| GO:0045165 | cell fate commitment | 274 | -0.333327042 | -1.523700531 | 0.000996964 |
| GO:1901654 | response to ketone | 208 | -0.34260451 | -1.524357813 | 0.002168456 |
| GO:1903305 | regulation of regulated secretory pathway | 116 | -0.368232825 | -1.524653493 | 0.008912495 |
| GO:0046718 | symbiont entry into host cell | 151 | -0.355750922 | -1.525012731 | 0.005514001 |
| GO:0099565 | chemical synaptic transmission, postsynaptic | 110 | -0.37127964 | -1.525564196 | 0.009460983 |
| GO:0071692 | protein localization to extracellular region | 347 | -0.327831821 | -1.527889193 | 0.000263733 |

| | | | | | |
|---|---|---|---|---|---|
| GO:0001890 | placenta development | 146 | -0.357938639 | -1.527976777 | 0.00556923 |
| GO:0010811 | positive regulation of cell-substrate adhesion | 119 | -0.368555359 | -1.531404865 | 0.007780056 |
| GO:0006869 | lipid transport | 419 | -0.325416941 | -1.537191621 | 0.000152363 |
| GO:0106106 | cold-induced thermogenesis | 148 | -0.359942814 | -1.539372605 | 0.004723824 |
| GO:0001938 | positive regulation of endothelial cell proliferation | 90 | -0.386943741 | -1.540717615 | 0.01051914 |
| GO:1905954 | positive regulation of lipid localization | 106 | -0.377410284 | -1.542656438 | 0.008118004 |
| GO:0039531 | regulation of cytoplasmic pattern recognition receptor signaling | 102 | -0.380123106 | -1.544530796 | 0.008447937 |
| GO:0120161 | regulation of cold-induced thermogenesis | 147 | -0.362324299 | -1.548219453 | 0.004195735 |
| GO:0019058 | viral life cycle | 304 | -0.33577482 | -1.548586655 | 0.00024584 |
| GO:0014013 | regulation of gliogenesis | 102 | -0.381134265 | -1.548639377 | 0.0081074 |
| GO:0030856 | regulation of epithelial cell differentiation | 148 | -0.362166051 | -1.548880755 | 0.00401711 |
| GO:0050806 | positive regulation of synaptic transmission | 159 | -0.359214091 | -1.549089931 | 0.003304222 |
| GO:0001892 | embryonic placenta development | 86 | -0.391692078 | -1.549206146 | 0.010529572 |
| GO:0032088 | negative regulation of NF-kappaB transcription factor activity | 75 | -0.400959582 | -1.550582976 | 0.011888945 |
| GO:0097305 | response to alcohol | 244 | -0.342971708 | -1.550604008 | 0.000944374 |
| GO:0032024 | positive regulation of insulin secretion | 83 | -0.394493029 | -1.550827477 | 0.010492584 |
| GO:0023061 | signal release | 463 | -0.326544844 | -1.551905891 | 2.53942720322908e-05 |
| GO:0002244 | hematopoietic progenitor cell differentiation | 129 | -0.369720895 | -1.553487905 | 0.004938209 |
| GO:0008625 | extrinsic apoptotic signaling pathway via death domain recepto | 77 | -0.401182555 | -1.55830623 | 0.010499584 |
| GO:0071900 | regulation of protein serine/threonine kinase activity | 249 | -0.344227563 | -1.559038516 | 0.000678106 |
| GO:1990845 | adaptive thermogenesis | 164 | -0.360298864 | -1.560169604 | 0.002514372 |
| GO:0051965 | positive regulation of synapse assembly | 60 | -0.420101465 | -1.561125408 | 0.013938811 |
| GO:0031960 | response to corticosteroid | 149 | -0.365123407 | -1.562567107 | 0.003257611 |
| GO:0008037 | cell recognition | 146 | -0.366215736 | -1.563310239 | 0.003344024 |
| GO:0071805 | potassium ion transmembrane transport | 198 | -0.353480515 | -1.564201735 | 0.001495993 |
| GO:1901343 | negative regulation of vasculature development | 100 | -0.386346817 | -1.564594021 | 0.006895281 |
| GO:0016042 | lipid catabolic process | 328 | -0.337203315 | -1.564865589 | 0.000289995 |
| GO:1902414 | protein localization to cell junction | 105 | -0.383514097 | -1.56506482 | 0.006396032 |
| GO:0043065 | positive regulation of apoptotic process | 486 | -0.328792827 | -1.566832863 | 2.08496048016033e-05 |
| GO:1903035 | negative regulation of response to wounding | 92 | -0.39222454 | -1.566904595 | 0.007798752 |
| GO:1904019 | epithelial cell apoptotic process | 127 | -0.373912556 | -1.5677287 | 0.004139472 |
| GO:0051056 | regulation of small GTPase mediated signal transduction | 296 | -0.340720557 | -1.567862103 | 0.000179422 |
| GO:0097006 | regulation of plasma lipoprotein particle levels | 75 | -0.405728142 | -1.569023854 | 0.009820707 |
| GO:0051235 | maintenance of location | 314 | -0.339571701 | -1.570238702 | 0.000281425 |
| GO:0052372 | modulation by symbiont of entry into host | 53 | -0.432296067 | -1.57052181 | 0.014877576 |
| GO:0046888 | negative regulation of hormone secretion | 66 | -0.415579129 | -1.572173141 | 0.011437658 |
| GO:0001508 | action potential | 141 | -0.370248836 | -1.573534707 | 0.003237662 |
| GO:0019229 | regulation of vasoconstriction | 66 | -0.416033216 | -1.573890993 | 0.011256329 |
| GO:0030324 | lung development | 185 | -0.358424958 | -1.573903122 | 0.001534367 |
| GO:0055092 | sterol homeostasis | 85 | -0.398775202 | -1.574158271 | 0.008042137 |
| GO:0015833 | peptide transport | 258 | -0.346574874 | -1.575028308 | 0.000388563 |
| GO:0034762 | regulation of transmembrane transport | 436 | -0.332918182 | -1.576262778 | 7.07159016362788e-05 |
| GO:0002262 | myeloid cell homeostasis | 165 | -0.363780622 | -1.576283081 | 0.001817117 |
| GO:0061045 | negative regulation of wound healing | 69 | -0.413669615 | -1.576852867 | 0.010234364 |
| GO:0010721 | negative regulation of cell development | 269 | -0.345608374 | -1.577070903 | 0.000329396 |

| | | | | | |
|---|---|---|---|---|---|
| GO:0086002 | cardiac muscle cell action potential involved in contraction | 56 | -0.430176739 | -1.578760961 | 0.012610556 |
| GO:0045055 | regulated exocytosis | 214 | -0.354654961 | -1.582623247 | 0.000903311 |
| GO:0006509 | membrane protein ectodomain proteolysis | 47 | -0.446279128 | -1.584185516 | 0.014941675 |
| GO:0006813 | potassium ion transport | 220 | -0.354063467 | -1.584768755 | 0.000829294 |
| GO:0030072 | peptide hormone secretion | 230 | -0.352618791 | -1.584957647 | 0.000731588 |
| GO:0032370 | positive regulation of lipid transport | 84 | -0.402756808 | -1.586441468 | 0.007073912 |
| GO:0009308 | amine metabolic process | 109 | -0.386826464 | -1.58761648 | 0.004502108 |
| GO:0030323 | respiratory tube development | 189 | -0.360821731 | -1.588117952 | 0.00102802 |
| GO:0000302 | response to reactive oxygen species | 181 | -0.363099289 | -1.590631287 | 0.001057921 |
| GO:0090276 | regulation of peptide hormone secretion | 189 | -0.361753995 | -1.592221209 | 0.000908482 |
| GO:0002790 | peptide secretion | 235 | -0.353955013 | -1.594359026 | 0.000532297 |
| GO:0062013 | positive regulation of small molecule metabolic process | 131 | -0.378745152 | -1.594575876 | 0.002728099 |
| GO:0090087 | regulation of peptide transport | 194 | -0.361222496 | -1.594781783 | 0.000917082 |
| GO:0051346 | negative regulation of hydrolase activity | 175 | -0.365624839 | -1.595427651 | 0.00126922 |
| GO:0034764 | positive regulation of transmembrane transport | 202 | -0.359733581 | -1.595686638 | 0.000780954 |
| GO:0070509 | calcium ion import | 73 | -0.415532283 | -1.599016443 | 0.007354359 |
| GO:0009914 | hormone transport | 308 | -0.346557154 | -1.599813281 | 0.000183442 |
| GO:0031638 | zymogen activation | 56 | -0.436302299 | -1.601241941 | 0.010527817 |
| GO:0045109 | intermediate filament organization | 70 | -0.419358859 | -1.603072096 | 0.007482814 |
| GO:0032892 | positive regulation of organic acid transport | 46 | -0.453829462 | -1.603562789 | 0.012788098 |
| GO:0090132 | epithelium migration | 300 | -0.348403535 | -1.605149473 | 0.000119323 |
| GO:2000379 | positive regulation of reactive oxygen species metabolic proce | 66 | -0.424541087 | -1.606077032 | 0.008145844 |
| GO:2000181 | negative regulation of blood vessel morphogenesis | 99 | -0.397404514 | -1.606877956 | 0.004347998 |
| GO:1903532 | positive regulation of secretion by cell | 277 | -0.351332306 | -1.607493029 | 0.000118017 |
| GO:0016525 | negative regulation of angiogenesis | 98 | -0.398304878 | -1.608094526 | 0.004377579 |
| GO:0046328 | regulation of JNK cascade | 134 | -0.381021764 | -1.608902417 | 0.002152351 |
| GO:0050909 | sensory perception of taste | 54 | -0.441327772 | -1.609214313 | 0.009899146 |
| GO:0003018 | vascular process in circulatory system | 259 | -0.354022112 | -1.6094561 | 0.000217061 |
| GO:2001056 | positive regulation of cysteine-type endopeptidase activity | 114 | -0.38998488 | -1.610971772 | 0.003016863 |
| GO:1903531 | negative regulation of secretion by cell | 139 | -0.37985913 | -1.611069091 | 0.001804534 |
| GO:0001954 | positive regulation of cell-matrix adhesion | 54 | -0.442057257 | -1.611874237 | 0.009641281 |
| GO:0010631 | epithelial cell migration | 297 | -0.350308861 | -1.612606953 | 0.000166425 |
| GO:0002791 | regulation of peptide secretion | 192 | -0.365868174 | -1.613121254 | 0.000656254 |
| GO:0051047 | positive regulation of secretion | 301 | -0.350073245 | -1.613203983 | 0.000126841 |
| GO:0051347 | positive regulation of transferase activity | 355 | -0.345578241 | -1.613226064 | 5.69209502216621e-05 |
| GO:0030336 | negative regulation of cell migration | 280 | -0.352324417 | -1.61332079 | 0.000155026 |
| GO:0016447 | somatic recombination of immunoglobulin gene segments | 59 | -0.435649809 | -1.614473724 | 0.00866831 |
| GO:0007189 | adenylate cyclase-activating G protein-coupled receptor signal | 154 | -0.375951705 | -1.615445838 | 0.00123338 |
| GO:0060627 | regulation of vesicle-mediated transport | 494 | -0.338964502 | -1.616739737 | 2.12841941216238e-06 |
| GO:0090130 | tissue migration | 305 | -0.350687238 | -1.617732882 | 4.1697542915179e-05 |
| GO:0045214 | sarcomere organization | 47 | -0.455779916 | -1.61791107 | 0.011053344 |
| GO:0043536 | positive regulation of blood vessel endothelial cell migration | 53 | -0.446079987 | -1.620598478 | 0.009276944 |
| GO:1904062 | regulation of monoatomic cation transmembrane transport | 278 | -0.354206264 | -1.62106254 | 0.000206334 |
| GO:0141085 | regulation of inflammasome-mediated signaling pathway | 35 | -0.485657131 | -1.621310141 | 0.014155667 |
| GO:0010950 | positive regulation of endopeptidase activity | 135 | -0.383587514 | -1.621622673 | 0.001797227 |

| | | | | | |
|---|---|---|---|---|---|
| GO:2000116 | regulation of cysteine-type endopeptidase activity | 168 | -0.373493426 | -1.622098668 | 0.000850909 |
| GO:0050708 | regulation of protein secretion | 250 | -0.358235606 | -1.623179165 | 0.000160764 |
| GO:1905517 | macrophage migration | 56 | -0.44235375 | -1.623450939 | 0.008359486 |
| GO:0030522 | intracellular receptor signaling pathway | 351 | -0.348560449 | -1.625840422 | 2.12986677124171e-05 |
| GO:0070988 | demethylation | 50 | -0.452642459 | -1.626304326 | 0.00945237 |
| GO:0062207 | regulation of pattern recognition receptor signaling pathway | 152 | -0.379110399 | -1.62648689 | 0.001025172 |
| GO:0034367 | protein-containing complex remodeling | 36 | -0.484644534 | -1.627387111 | 0.013649762 |
| GO:0006026 | aminoglycan catabolic process | 33 | -0.49358749 | -1.627528179 | 0.014529915 |
| GO:1903169 | regulation of calcium ion transmembrane transport | 152 | -0.37969904 | -1.629012323 | 0.000988118 |
| GO:2000146 | negative regulation of cell motility | 295 | -0.354225034 | -1.629416077 | 6.89901511265213e-05 |
| GO:1903522 | regulation of blood circulation | 243 | -0.360589705 | -1.629572937 | 0.0002074 |
| GO:0042445 | hormone metabolic process | 229 | -0.362652665 | -1.629640622 | 0.000371644 |
| GO:0042554 | superoxide anion generation | 39 | -0.477058279 | -1.629908028 | 0.012029133 |
| GO:0001894 | tissue homeostasis | 253 | -0.359342491 | -1.630164476 | 0.00016044 |
| GO:0060249 | anatomical structure homeostasis | 253 | -0.359342491 | -1.630164476 | 0.00016044 |
| GO:0016445 | somatic diversification of immunoglobulins | 64 | -0.433596838 | -1.63040677 | 0.006690136 |
| GO:0008217 | regulation of blood pressure | 180 | -0.372649266 | -1.631361562 | 0.000577715 |
| GO:0098739 | import across plasma membrane | 188 | -0.371168711 | -1.632806445 | 0.0004426 |
| GO:0042632 | cholesterol homeostasis | 84 | -0.414530908 | -1.632819136 | 0.004198448 |
| GO:1901658 | glycosyl compound catabolic process | 32 | -0.498585597 | -1.633162808 | 0.014112749 |
| GO:0048534 | hematopoietic or lymphoid organ development | 97 | -0.406845855 | -1.639915548 | 0.002931932 |
| GO:0014002 | astrocyte development | 40 | -0.477590034 | -1.64061796 | 0.010637348 |
| GO:0060191 | regulation of lipase activity | 65 | -0.435261571 | -1.641587234 | 0.005906311 |
| GO:0051048 | negative regulation of secretion | 158 | -0.381033771 | -1.641777843 | 0.000676473 |
| GO:0007565 | female pregnancy | 171 | -0.377492668 | -1.642940372 | 0.000533307 |
| GO:1900047 | negative regulation of hemostasis | 47 | -0.462970377 | -1.643435511 | 0.008854326 |
| GO:0051057 | positive regulation of small GTPase mediated signal transduct | 75 | -0.425125023 | -1.644035089 | 0.004095386 |
| GO:0032228 | regulation of synaptic transmission, GABAergic | 34 | -0.495920071 | -1.645398331 | 0.011828645 |
| GO:0061515 | myeloid cell development | 86 | -0.416443558 | -1.647102295 | 0.0035502 |
| GO:0032570 | response to progesterone | 41 | -0.477015173 | -1.647136355 | 0.009607829 |
| GO:0046883 | regulation of hormone secretion | 244 | -0.364491596 | -1.647897239 | 9.67887866317342e-05 |
| GO:0045446 | endothelial cell differentiation | 106 | -0.403440665 | -1.649055064 | 0.002189125 |
| GO:0070588 | calcium ion transmembrane transport | 307 | -0.357483055 | -1.64967108 | 3.85435045663089e-05 |
| GO:0044546 | NLRP3 inflammasome complex assembly | 36 | -0.492583568 | -1.65404558 | 0.010807951 |
| GO:0070293 | renal absorption | 33 | -0.501760817 | -1.654478458 | 0.011111111 |
| GO:0006721 | terpenoid metabolic process | 93 | -0.413378039 | -1.654621531 | 0.002688387 |
| GO:0030212 | hyaluronan metabolic process | 33 | -0.501902367 | -1.654945196 | 0.011065324 |
| GO:0071277 | cellular response to calcium ion | 82 | -0.421982744 | -1.655634141 | 0.003453226 |
| GO:0045777 | positive regulation of blood pressure | 31 | -0.509553371 | -1.657008252 | 0.011777549 |
| GO:0060706 | cell differentiation involved in embryonic placenta developmen | 24 | -0.541293818 | -1.657610859 | 0.014194983 |
| GO:0002562 | somatic diversification of immune receptors via germline recc | 67 | -0.437697876 | -1.659888228 | 0.00459879 |
| GO:0016444 | somatic cell DNA recombination | 67 | -0.437697876 | -1.659888228 | 0.00459879 |
| GO:0051279 | regulation of release of sequestered calcium ion into cytosol | 76 | -0.428394248 | -1.660227251 | 0.003406526 |
| GO:0030195 | negative regulation of blood coagulation | 46 | -0.469903808 | -1.660359943 | 0.007439019 |
| GO:0016101 | diterpenoid metabolic process | 83 | -0.422736341 | -1.661857333 | 0.00305583 |

| | | | | | |
|---|---|---|---|---|---|
| GO:0048015 | phosphatidylinositol-mediated signaling | 22 | -0.554264375 | -1.662481573 | 0.013485593 |
| GO:0042060 | wound healing | 404 | -0.352900235 | -1.662945484 | 6.45465899854198e-06 |
| GO:0098703 | calcium ion import across plasma membrane | 35 | -0.498549726 | -1.664350577 | 0.010104688 |
| GO:1902656 | calcium ion import into cytosol | 35 | -0.498549726 | -1.664350577 | 0.010104688 |
| GO:0045780 | positive regulation of bone resorption | 19 | -0.575835964 | -1.665462376 | 0.014192371 |
| GO:0008228 | opsonization | 16 | -0.602230535 | -1.666148398 | 0.014992853 |
| GO:0002200 | somatic diversification of immune receptors | 73 | -0.433036948 | -1.666376426 | 0.003443052 |
| GO:0006690 | icosanoid metabolic process | 107 | -0.40720721 | -1.667029055 | 0.001573718 |
| GO:0015718 | monocarboxylic acid transport | 170 | -0.383373031 | -1.667105668 | 0.000327678 |
| GO:0034145 | positive regulation of toll-like receptor 4 signaling pathway | 13 | -0.636963778 | -1.667596301 | 0.015217798 |
| GO:0042744 | hydrogen peroxide catabolic process | 25 | -0.53865136 | -1.667621547 | 0.012655036 |
| GO:0043534 | blood vessel endothelial cell migration | 114 | -0.403871268 | -1.668334455 | 0.001405291 |
| GO:0051592 | response to calcium ion | 139 | -0.393363794 | -1.668345447 | 0.000852141 |
| GO:0040013 | negative regulation of locomotion | 330 | -0.359472496 | -1.668901941 | 1.54433066144443e-05 |
| GO:0034110 | regulation of homotypic cell-cell adhesion | 34 | -0.503151975 | -1.66939285 | 0.009423335 |
| GO:0033002 | muscle cell proliferation | 200 | -0.376920382 | -1.670001021 | 0.000201431 |
| GO:0010759 | positive regulation of macrophage chemotaxis | 18 | -0.58577805 | -1.67011528 | 0.014215831 |
| GO:0001659 | temperature homeostasis | 178 | -0.382013029 | -1.670410969 | 0.000277169 |
| GO:0048002 | antigen processing and presentation of peptide antigen | 47 | -0.470880341 | -1.671514013 | 0.006669871 |
| GO:0015669 | gas transport | 19 | -0.57808042 | -1.671953908 | 0.013556892 |
| GO:0046879 | hormone secretion | 297 | -0.363212347 | -1.672006685 | 4.05695033468165e-05 |
| GO:0019079 | viral genome replication | 122 | -0.401094448 | -1.672160652 | 0.001034536 |
| GO:0003334 | keratinocyte development | 12 | -0.652802429 | -1.672758583 | 0.015038882 |
| GO:0010959 | regulation of metal ion transport | 358 | -0.358144656 | -1.672924961 | 4.42933852956908e-06 |
| GO:0060384 | innervation | 26 | -0.53575582 | -1.673185268 | 0.011427674 |
| GO:0032691 | negative regulation of interleukin-1 beta production | 28 | -0.52712905 | -1.674919535 | 0.010622414 |
| GO:0046596 | regulation of viral entry into host cell | 46 | -0.474138692 | -1.675323501 | 0.006445201 |
| GO:0071481 | cellular response to X-ray | 11 | -0.669988543 | -1.675356862 | 0.014856965 |
| GO:0042537 | benzene-containing compound metabolic process | 23 | -0.552418475 | -1.67540692 | 0.012211063 |
| GO:0048485 | sympathetic nervous system development | 21 | -0.565316166 | -1.676734773 | 0.012302738 |
| GO:0007187 | G protein-coupled receptor signaling pathway, coupled to cycli | 53 | -0.461689902 | -1.677308945 | 0.005255978 |
| GO:0009395 | phospholipid catabolic process | 53 | -0.461914612 | -1.67812531 | 0.005241617 |
| GO:1900225 | regulation of NLRP3 inflammasome complex assembly | 33 | -0.509051015 | -1.678516754 | 0.008714896 |
| GO:0048143 | astrocyte activation | 22 | -0.559742234 | -1.67891207 | 0.011948044 |
| GO:0061318 | renal filtration cell differentiation | 22 | -0.559746651 | -1.678925319 | 0.011948044 |
| GO:0072112 | podocyte differentiation | 22 | -0.559746651 | -1.678925319 | 0.011948044 |
| GO:0051933 | amino acid neurotransmitter reuptake | 10 | -0.68892187 | -1.67926918 | 0.014606292 |
| GO:0141084 | inflammasome-mediated signaling pathway | 40 | -0.488965427 | -1.679694727 | 0.007379195 |
| GO:0030224 | monocyte differentiation | 33 | -0.509464365 | -1.679879711 | 0.008623321 |
| GO:0035458 | cellular response to interferon-beta | 23 | -0.553898753 | -1.679896392 | 0.011748765 |
| GO:0043903 | regulation of biological process involved in symbiotic interactio | 59 | -0.453558306 | -1.680840784 | 0.004249171 |
| GO:0050912 | detection of chemical stimulus involved in sensory perception | 34 | -0.506977879 | -1.682086702 | 0.008372914 |
| GO:0001780 | neutrophil homeostasis | 19 | -0.581753827 | -1.682578324 | 0.012318522 |
| GO:1904035 | regulation of epithelial cell apoptotic process | 96 | -0.418255336 | -1.683003482 | 0.00172035 |
| GO:0002357 | defense response to tumor cell | 12 | -0.656847373 | -1.683123457 | 0.013668596 |

| | | | | | |
|---|---|---|---|---|---|
| GO:2001238 | positive regulation of extrinsic apoptotic signaling pathway | 50 | -0.468471719 | -1.683177458 | 0.00539103 |
| GO:0007229 | integrin-mediated signaling pathway | 107 | -0.411549238 | -1.684804498 | 0.001365623 |
| GO:0002433 | immune response-regulating cell surface receptor signaling pa | 22 | -0.56179697 | -1.685075121 | 0.011483575 |
| GO:0038096 | Fc-gamma receptor signaling pathway involved in phagocytos | 22 | -0.56179697 | -1.685075121 | 0.011483575 |
| GO:0002230 | positive regulation of defense response to virus by host | 27 | -0.53490816 | -1.685562371 | 0.010124842 |
| GO:0032604 | granulocyte macrophage colony-stimulating factor production | 17 | -0.600372513 | -1.686807123 | 0.012314905 |
| GO:0032645 | regulation of granulocyte macrophage colony-stimulating facto | 17 | -0.600372513 | -1.686807123 | 0.012314905 |
| GO:0009164 | nucleoside catabolic process | 25 | -0.545062511 | -1.687469958 | 0.010243802 |
| GO:0050688 | regulation of defense response to virus | 59 | -0.45538285 | -1.687602359 | 0.004036713 |
| GO:0030239 | myofibril assembly | 71 | -0.440659683 | -1.688094832 | 0.00294329 |
| GO:0072672 | neutrophil extravasation | 14 | -0.63171685 | -1.688238775 | 0.01289181 |
| GO:1900017 | positive regulation of cytokine production involved in inflamma | 26 | -0.540804495 | -1.688952467 | 0.010046307 |
| GO:0015908 | fatty acid transport | 111 | -0.410576556 | -1.689025917 | 0.001217238 |
| GO:0002689 | negative regulation of leukocyte chemotaxis | 20 | -0.576696861 | -1.689074131 | 0.011134341 |
| GO:0010634 | positive regulation of epithelial cell migration | 146 | -0.395875425 | -1.689922208 | 0.000497253 |
| GO:0003159 | morphogenesis of an endothelium | 16 | -0.610900572 | -1.690135172 | 0.012200392 |
| GO:0061154 | endothelial tube morphogenesis | 16 | -0.610900572 | -1.690135172 | 0.012200392 |
| GO:0060231 | mesenchymal to epithelial transition | 19 | -0.584426716 | -1.69030899 | 0.011617865 |
| GO:0071260 | cellular response to mechanical stimulus | 70 | -0.44225783 | -1.690607392 | 0.002738627 |
| GO:0060397 | growth hormone receptor signaling pathway via JAK-STAT | 10 | -0.693596935 | -1.690664801 | 0.013196149 |
| GO:0072182 | regulation of nephron tubule epithelial cell differentiation | 12 | -0.659807127 | -1.690707611 | 0.012829297 |
| GO:0150105 | protein localization to cell-cell junction | 21 | -0.570350314 | -1.691666119 | 0.010885668 |
| GO:0048799 | animal organ maturation | 31 | -0.520252608 | -1.691800924 | 0.00827196 |
| GO:0007200 | phospholipase C-activating G protein-coupled receptor signalii | 98 | -0.419289847 | -1.692818103 | 0.00151633 |
| GO:0048251 | elastic fiber assembly | 10 | -0.694623229 | -1.693166424 | 0.012691283 |
| GO:0009435 | NAD biosynthetic process | 24 | -0.553023178 | -1.693529824 | 0.010161956 |
| GO:0010745 | negative regulation of macrophage derived foam cell differentia | 13 | -0.64701591 | -1.69391318 | 0.012000613 |
| GO:0061082 | myeloid leukocyte cytokine production | 48 | -0.475623776 | -1.694505952 | 0.005026367 |
| GO:0071888 | macrophage apoptotic process | 13 | -0.647429211 | -1.694995218 | 0.011932524 |
| GO:0010632 | regulation of epithelial cell migration | 222 | -0.378426932 | -1.695534143 | 8.21957022298519e-05 |
| GO:0016553 | base conversion or substitution editing | 19 | -0.586440042 | -1.696132037 | 0.010917208 |
| GO:0072311 | glomerular epithelial cell differentiation | 23 | -0.559315855 | -1.696325696 | 0.010122748 |
| GO:0001523 | retinoid metabolic process | 78 | -0.435923027 | -1.69658984 | 0.002288828 |
| GO:0007218 | neuropeptide signaling pathway | 100 | -0.419147141 | -1.697425946 | 0.001460642 |
| GO:0070857 | regulation of bile acid biosynthetic process | 11 | -0.67899538 | -1.697879136 | 0.011709156 |
| GO:0042573 | retinoic acid metabolic process | 31 | -0.522162123 | -1.698010444 | 0.007810698 |
| GO:0006691 | leukotriene metabolic process | 27 | -0.538915376 | -1.698189608 | 0.009015484 |
| GO:0055002 | striated muscle cell development | 73 | -0.441333496 | -1.698302503 | 0.002327503 |
| GO:0050819 | negative regulation of coagulation | 49 | -0.474651358 | -1.69846535 | 0.004636337 |
| GO:0034115 | negative regulation of heterotypic cell-cell adhesion | 10 | -0.697127565 | -1.699270824 | 0.011960098 |
| GO:0070587 | regulation of cell-cell adhesion involved in gastrulation | 10 | -0.697127565 | -1.699270824 | 0.011960098 |
| GO:0021984 | adenohypophysis development | 15 | -0.624867917 | -1.699939391 | 0.011305104 |
| GO:0046887 | positive regulation of hormone secretion | 137 | -0.401666337 | -1.700362672 | 0.00045208 |
| GO:0090277 | positive regulation of peptide hormone secretion | 106 | -0.4162933 | -1.701589933 | 0.001094562 |
| GO:1903975 | regulation of glial cell migration | 19 | -0.58843157 | -1.701892035 | 0.010363201 |

| | | | | | |
|---|---|---|---|---|---|
| GO:0002793 | positive regulation of peptide secretion | 108 | -0.415272318 | -1.702218239 | 0.001065406 |
| GO:0051345 | positive regulation of hydrolase activity | 438 | -0.359781982 | -1.703981988 | 8.06811333160211e-07 |
| GO:0046639 | negative regulation of alpha-beta T cell differentiation | 25 | -0.550605518 | -1.704630664 | 0.008872709 |
| GO:0048304 | positive regulation of isotype switching to IgG isotypes | 10 | -0.699327034 | -1.704632101 | 0.011437823 |
| GO:0046131 | pyrimidine ribonucleoside metabolic process | 14 | -0.637892747 | -1.704743622 | 0.010684193 |
| GO:0046133 | pyrimidine ribonucleoside catabolic process | 14 | -0.637892747 | -1.704743622 | 0.010684193 |
| GO:0048286 | lung alveolus development | 51 | -0.473104109 | -1.705900764 | 0.00406991 |
| GO:1904064 | positive regulation of cation transmembrane transport | 131 | -0.405189692 | -1.705911492 | 0.000530464 |
| GO:0060046 | regulation of acrosome reaction | 16 | -0.616623566 | -1.705968573 | 0.010721053 |
| GO:0035510 | DNA dealkylation | 35 | -0.511118463 | -1.706309854 | 0.006857836 |
| GO:0051090 | regulation of DNA-binding transcription factor activity | 348 | -0.366053201 | -1.70635902 | 4.67008456241452e-06 |
| GO:0010596 | negative regulation of endothelial cell migration | 51 | -0.473454978 | -1.707165912 | 0.004026613 |
| GO:0043380 | regulation of memory T cell differentiation | 10 | -0.700501754 | -1.707495519 | 0.011159276 |
| GO:0031644 | regulation of nervous system process | 113 | -0.414067366 | -1.707914965 | 0.000839446 |
| GO:0080111 | DNA demethylation | 30 | -0.52941634 | -1.707985116 | 0.007136678 |
| GO:0090715 | immunological memory formation process | 12 | -0.666893775 | -1.708866629 | 0.01087664 |
| GO:0010633 | negative regulation of epithelial cell migration | 65 | -0.453131516 | -1.708983656 | 0.002701227 |
| GO:0043301 | negative regulation of leukocyte degranulation | 12 | -0.666952818 | -1.709017923 | 0.01087664 |
| GO:0010922 | positive regulation of phosphatase activity | 28 | -0.538194763 | -1.710080146 | 0.007496345 |
| GO:0098659 | inorganic cation import across plasma membrane | 113 | -0.415172875 | -1.712474885 | 0.000826532 |
| GO:0099587 | inorganic ion import across plasma membrane | 113 | -0.415172875 | -1.712474885 | 0.000826532 |
| GO:0045646 | regulation of erythrocyte differentiation | 42 | -0.493543237 | -1.712810853 | 0.004877832 |
| GO:0032480 | negative regulation of type I interferon production | 43 | -0.491561524 | -1.713509683 | 0.004792874 |
| GO:0009119 | ribonucleoside metabolic process | 34 | -0.516633168 | -1.714121697 | 0.006241627 |
| GO:0050691 | regulation of defense response to virus by host | 39 | -0.501938484 | -1.714913251 | 0.005134022 |
| GO:0002855 | regulation of natural killer cell mediated immune response to tu | 10 | -0.703697289 | -1.715284737 | 0.010254 |
| GO:0043542 | endothelial cell migration | 212 | -0.384831346 | -1.715502905 | 6.62817665479491e-05 |
| GO:0071320 | cellular response to cAMP | 47 | -0.483343221 | -1.715754293 | 0.004150465 |
| GO:0070527 | platelet aggregation | 70 | -0.448924969 | -1.716093689 | 0.002088549 |
| GO:0019883 | antigen processing and presentation of endogenous antigen | 23 | -0.565900503 | -1.716296 | 0.008432967 |
| GO:0051591 | response to cAMP | 80 | -0.439117384 | -1.716409139 | 0.001628819 |
| GO:0072010 | glomerular epithelium development | 26 | -0.550193758 | -1.718275484 | 0.007330665 |
| GO:1904427 | positive regulation of calcium ion transmembrane transport | 80 | -0.43965312 | -1.718503206 | 0.001601672 |
| GO:0035666 | TRIF-dependent toll-like receptor signaling pathway | 11 | -0.68750918 | -1.719168536 | 0.009650974 |
| GO:0007254 | JNK cascade | 167 | -0.396874787 | -1.722697735 | 0.000188833 |
| GO:0048771 | tissue remodeling | 171 | -0.395969017 | -1.723353959 | 0.000141528 |
| GO:0043312 | neutrophil degranulation | 12 | -0.672721596 | -1.723799995 | 0.00919804 |
| GO:0140632 | canonical inflammasome complex assembly | 38 | -0.507475223 | -1.724205697 | 0.004810718 |
| GO:0060330 | regulation of response to type II interferon | 14 | -0.645443507 | -1.724922734 | 0.008645096 |
| GO:0060334 | regulation of type II interferon-mediated signaling pathway | 14 | -0.645443507 | -1.724922734 | 0.008645096 |
| GO:0010518 | positive regulation of phospholipase activity | 42 | -0.497215185 | -1.725554115 | 0.004227455 |
| GO:0060368 | regulation of Fc receptor mediated stimulatory signaling pathw | 10 | -0.708056131 | -1.725909555 | 0.009104995 |
| GO:0044706 | multi-multicellular organism process | 199 | -0.389830607 | -1.726061396 | 7.96459657212301e-05 |
| GO:0060707 | trophoblast giant cell differentiation | 12 | -0.673636551 | -1.726144502 | 0.008958241 |
| GO:2000320 | negative regulation of T-helper 17 cell differentiation | 10 | -0.70854648 | -1.727104796 | 0.008983131 |

| | | | | | |
|---|---|---|---|---|---|
| GO:0007520 | myoblast fusion | 45 | -0.49100704 | -1.72713494 | 0.003710657 |
| GO:1902041 | regulation of extrinsic apoptotic signaling pathway via death do | 43 | -0.495532815 | -1.727353004 | 0.004070256 |
| GO:0051281 | positive regulation of release of sequestered calcium ion into c | 40 | -0.502852339 | -1.727399067 | 0.004150797 |
| GO:0035767 | endothelial cell chemotaxis | 28 | -0.543746049 | -1.72771901 | 0.006065507 |
| GO:0030431 | sleep | 21 | -0.582932916 | -1.728986274 | 0.007938808 |
| GO:0060338 | regulation of type I interferon-mediated signaling pathway | 42 | -0.498290287 | -1.72928519 | 0.00406486 |
| GO:0071676 | negative regulation of mononuclear cell migration | 22 | -0.576619551 | -1.729534535 | 0.007543604 |
| GO:0010887 | negative regulation of cholesterol storage | 10 | -0.709945045 | -1.730513844 | 0.008721993 |
| GO:0061005 | cell differentiation involved in kidney development | 58 | -0.468547671 | -1.731261937 | 0.00238467 |
| GO:0048755 | branching morphogenesis of a nerve | 12 | -0.676020526 | -1.732253262 | 0.008255969 |
| GO:0048291 | isotype switching to IgG isotypes | 15 | -0.637070459 | -1.733136135 | 0.008144357 |
| GO:0015732 | prostaglandin transport | 28 | -0.54585158 | -1.734409203 | 0.005801114 |
| GO:0002040 | sprouting angiogenesis | 127 | -0.413766124 | -1.734825474 | 0.000279351 |
| GO:0030851 | granulocyte differentiation | 39 | -0.508082194 | -1.735903732 | 0.003940063 |
| GO:0086010 | membrane depolarization during action potential | 26 | -0.556191698 | -1.737007272 | 0.006027784 |
| GO:1904044 | response to aldosterone | 18 | -0.609485985 | -1.737709115 | 0.007666043 |
| GO:0043123 | positive regulation of canonical NF-kappaB signal transductior | 202 | -0.391934843 | -1.73852324 | 5.67827208346126e-05 |
| GO:0035821 | modulation of process of another organism | 14 | -0.650607741 | -1.738723949 | 0.007549714 |
| GO:0002474 | antigen processing and presentation of peptide antigen via MH( | 29 | -0.543331858 | -1.739163003 | 0.005338713 |
| GO:0051924 | regulation of calcium ion transport | 228 | -0.387584637 | -1.740836198 | 2.54805757360682e-05 |
| GO:0055075 | potassium ion homeostasis | 34 | -0.524704063 | -1.740899875 | 0.004688832 |
| GO:0007156 | homophilic cell adhesion via plasma membrane adhesion mol | 158 | -0.404169435 | -1.741463547 | 0.000121404 |
| GO:0034765 | regulation of monoatomic ion transmembrane transport | 315 | -0.376570262 | -1.741636814 | 1.49469661396339e-06 |
| GO:0034368 | protein-lipid complex remodeling | 33 | -0.528543093 | -1.742788859 | 0.004563492 |
| GO:0034369 | plasma lipoprotein particle remodeling | 33 | -0.528543093 | -1.742788859 | 0.004563492 |
| GO:0019369 | arachidonic acid metabolic process | 55 | -0.476538215 | -1.743455271 | 0.002415908 |
| GO:0030214 | hyaluronan catabolic process | 16 | -0.630816636 | -1.745235532 | 0.007280343 |
| GO:0045453 | bone resorption | 66 | -0.461389544 | -1.745478054 | 0.00178539 |
| GO:0042492 | gamma-delta T cell differentiation | 13 | -0.667072244 | -1.746421453 | 0.007387611 |
| GO:0071636 | positive regulation of transforming growth factor beta productic | 22 | -0.58246429 | -1.747065464 | 0.00602207 |
| GO:1905523 | positive regulation of macrophage migration | 25 | -0.564749655 | -1.748419781 | 0.005815328 |
| GO:0014051 | gamma-aminobutyric acid secretion | 11 | -0.699220544 | -1.748453684 | 0.007108513 |
| GO:0019373 | epoxygenase P450 pathway | 17 | -0.622339497 | -1.748525581 | 0.006702214 |
| GO:2000347 | positive regulation of hepatocyte proliferation | 10 | -0.717816967 | -1.749701906 | 0.007033304 |
| GO:0034767 | positive regulation of monoatomic ion transmembrane transpo | 146 | -0.410217747 | -1.751147044 | 0.000211332 |
| GO:0034694 | response to prostaglandin | 31 | -0.538600822 | -1.751467181 | 0.004428112 |
| GO:0071349 | cellular response to interleukin-12 | 11 | -0.700866621 | -1.752569823 | 0.006728009 |
| GO:0070269 | pyroptosis | 34 | -0.528350452 | -1.75299812 | 0.004125563 |
| GO:0006874 | intracellular calcium ion homeostasis | 285 | -0.382433664 | -1.753979554 | 1.99668969344646e-06 |
| GO:0150079 | negative regulation of neuroinflammatory response | 13 | -0.670081432 | -1.754299625 | 0.006706725 |
| GO:0015671 | oxygen transport | 12 | -0.684658741 | -1.754388058 | 0.006628755 |
| GO:0044703 | multi-organism reproductive process | 192 | -0.397927427 | -1.754471245 | 3.44896971330358e-05 |
| GO:0010888 | negative regulation of lipid storage | 21 | -0.591573586 | -1.754614608 | 0.006006441 |
| GO:0050849 | negative regulation of calcium-mediated signaling | 20 | -0.59959808 | -1.756148984 | 0.005785968 |
| GO:0010594 | regulation of endothelial cell migration | 162 | -0.406284542 | -1.756803094 | 0.000116544 |

| | | | | | |
|---|---|---|---|---|---|
| GO:0050878 | regulation of body fluid levels | 357 | -0.376957185 | -1.760542555 | 4.38507540728186e-07 |
| GO:0010743 | regulation of macrophage derived foam cell differentiation | 31 | -0.541703521 | -1.761556797 | 0.004028352 |
| GO:0045672 | positive regulation of osteoclast differentiation | 28 | -0.554487099 | -1.761848022 | 0.004183645 |
| GO:0070671 | response to interleukin-12 | 12 | -0.688247897 | -1.763585008 | 0.005960741 |
| GO:0006816 | calcium ion transport | 397 | -0.375291011 | -1.766292515 | 2.49754873404433e-07 |
| GO:0050714 | positive regulation of protein secretion | 140 | -0.416287793 | -1.767188696 | 5.72734720452923e-05 |
| GO:2001267 | regulation of cysteine-type endopeptidase activity involved in a | 11 | -0.706784982 | -1.767369132 | 0.005655678 |
| GO:0043269 | regulation of monoatomic ion transport | 431 | -0.373847805 | -1.769019991 | 2.88206088870284e-08 |
| GO:0003014 | renal system process | 123 | -0.423951824 | -1.769096101 | 0.000191476 |
| GO:0070243 | regulation of thymocyte apoptotic process | 13 | -0.675973311 | -1.769724797 | 0.005685397 |
| GO:0051043 | regulation of membrane protein ectodomain proteolysis | 27 | -0.562398829 | -1.772188897 | 0.004046812 |
| GO:0006949 | syncytium formation | 61 | -0.47550223 | -1.772338685 | 0.00135553 |
| GO:0048872 | homeostasis of number of cells | 305 | -0.384260456 | -1.772607346 | 4.99384022106487e-07 |
| GO:2001026 | regulation of endothelial cell chemotaxis | 21 | -0.597721882 | -1.772850532 | 0.005088567 |
| GO:0070099 | regulation of chemokine-mediated signaling pathway | 10 | -0.727370665 | -1.772989351 | 0.005309796 |
| GO:0001893 | maternal placenta development | 34 | -0.534437145 | -1.773192978 | 0.003181707 |
| GO:0046135 | pyrimidine nucleoside catabolic process | 16 | -0.641055869 | -1.773563688 | 0.005385459 |
| GO:0043122 | regulation of canonical NF-kappaB signal transduction | 269 | -0.388679711 | -1.773612876 | 1.75909832701996e-06 |
| GO:2001204 | regulation of osteoclast development | 11 | -0.709299216 | -1.773656161 | 0.005344356 |
| GO:0006027 | glycosaminoglycan catabolic process | 27 | -0.563022765 | -1.774154999 | 0.003953063 |
| GO:0060068 | vagina development | 10 | -0.727963729 | -1.774434965 | 0.005187932 |
| GO:0060252 | positive regulation of glial cell proliferation | 21 | -0.598593395 | -1.775435448 | 0.004863124 |
| GO:0032634 | interleukin-5 production | 20 | -0.606204132 | -1.775497295 | 0.004667672 |
| GO:0032674 | regulation of interleukin-5 production | 20 | -0.606204132 | -1.775497295 | 0.004667672 |
| GO:0090205 | positive regulation of cholesterol metabolic process | 13 | -0.678361531 | -1.775977252 | 0.005259843 |
| GO:0045191 | regulation of isotype switching | 36 | -0.528973602 | -1.776239618 | 0.002962739 |
| GO:0019228 | neuronal action potential | 34 | -0.53539386 | -1.776367233 | 0.002999026 |
| GO:0050732 | negative regulation of peptidyl-tyrosine phosphorylation | 44 | -0.507964354 | -1.778291314 | 0.002294556 |
| GO:0033623 | regulation of integrin activation | 16 | -0.643588248 | -1.780569841 | 0.004969915 |
| GO:0014074 | response to purine-containing compound | 135 | -0.4214035 | -1.781490393 | 0.000192344 |
| GO:2000351 | regulation of endothelial cell apoptotic process | 49 | -0.498332838 | -1.783205807 | 0.001811069 |
| GO:0046683 | response to organophosphorus | 121 | -0.428288788 | -1.78353893 | 0.000158032 |
| GO:0045860 | positive regulation of protein kinase activity | 245 | -0.394548519 | -1.784402411 | 7.67922595899337e-06 |
| GO:0061081 | positive regulation of myeloid leukocyte cytokine production inv | 33 | -0.541198673 | -1.784518669 | 0.002823565 |
| GO:0007586 | digestion | 128 | -0.425208449 | -1.78491011 | 0.000103705 |
| GO:0043031 | negative regulation of macrophage activation | 16 | -0.645186066 | -1.78499041 | 0.004720588 |
| GO:0050798 | activated T cell proliferation | 40 | -0.520612628 | -1.788409235 | 0.00200845 |
| GO:1901342 | regulation of vasculature development | 285 | -0.390102229 | -1.789150376 | 5.61460266445653e-07 |
| GO:0043691 | reverse cholesterol transport | 17 | -0.636834203 | -1.78924992 | 0.004341582 |
| GO:0060135 | maternal process involved in female pregnancy | 55 | -0.489216677 | -1.789840494 | 0.001315169 |
| GO:0050918 | positive chemotaxis | 60 | -0.481721689 | -1.790110321 | 0.001061331 |
| GO:0001935 | endothelial cell proliferation | 150 | -0.417992235 | -1.790411372 | 6.07858746194057e-05 |
| GO:0090280 | positive regulation of calcium ion import | 25 | -0.57832329 | -1.79044267 | 0.003482893 |
| GO:0090026 | positive regulation of monocyte chemotaxis | 18 | -0.628929171 | -1.793143697 | 0.004087462 |
| GO:0086012 | membrane depolarization during cardiac muscle cell action pc | 20 | -0.612241179 | -1.793179066 | 0.003889726 |

| | | | | | |
|---|---|---|---|---|---|
| GO:0055074 | calcium ion homeostasis | 308 | -0.388670731 | -1.79422237 | 6.69878453265031e-07 |
| GO:0002887 | negative regulation of myeloid leukocyte mediated immunity | 11 | -0.718288493 | -1.796134528 | 0.003995296 |
| GO:0002829 | negative regulation of type 2 immune response | 15 | -0.660439314 | -1.796710591 | 0.004063817 |
| GO:0060837 | blood vessel endothelial cell differentiation | 12 | -0.701478878 | -1.797488431 | 0.004093728 |
| GO:0032418 | lysosome localization | 82 | -0.458261169 | -1.797971238 | 0.000419804 |
| GO:1990849 | vacuolar localization | 82 | -0.458261169 | -1.797971238 | 0.000419804 |
| GO:0019884 | antigen processing and presentation of exogenous antigen | 31 | -0.553050001 | -1.798454228 | 0.002782946 |
| GO:0097278 | complement-dependent cytotoxicity | 10 | -0.738025729 | -1.798961413 | 0.004125973 |
| GO:0042454 | ribonucleoside catabolic process | 20 | -0.614423166 | -1.799569836 | 0.003533168 |
| GO:0033083 | regulation of immature T cell proliferation | 12 | -0.702650881 | -1.800491603 | 0.003922442 |
| GO:0033084 | regulation of immature T cell proliferation in thymus | 12 | -0.702650881 | -1.800491603 | 0.003922442 |
| GO:0033674 | positive regulation of kinase activity | 289 | -0.392116088 | -1.800513952 | 1.12635805294504e-06 |
| GO:0046456 | icosanoid biosynthetic process | 49 | -0.50321621 | -1.800680188 | 0.001448855 |
| GO:0060415 | muscle tissue morphogenesis | 65 | -0.478651789 | -1.805233263 | 0.0006998 |
| GO:0042501 | serine phosphorylation of STAT protein | 21 | -0.608735232 | -1.805516265 | 0.003365539 |
| GO:0046643 | regulation of gamma-delta T cell activation | 11 | -0.722061678 | -1.805569662 | 0.003441835 |
| GO:0010575 | positive regulation of vascular endothelial growth factor produc | 29 | -0.564157787 | -1.805825182 | 0.002514162 |
| GO:2000317 | negative regulation of T-helper 17 type immune response | 11 | -0.722656263 | -1.807056465 | 0.003355356 |
| GO:0045765 | regulation of angiogenesis | 279 | -0.39476335 | -1.807075727 | 8.13441212370248e-07 |
| GO:0033625 | positive regulation of integrin activation | 10 | -0.741671772 | -1.80784876 | 0.003621107 |
| GO:0001960 | negative regulation of cytokine-mediated signaling pathway | 73 | -0.470365737 | -1.810021936 | 0.000495799 |
| GO:0032754 | positive regulation of interleukin-5 production | 11 | -0.723843181 | -1.810024444 | 0.003234287 |
| GO:0021604 | cranial nerve structural organization | 11 | -0.724084726 | -1.810628444 | 0.003216991 |
| GO:0033139 | regulation of peptidyl-serine phosphorylation of STAT protein | 20 | -0.618452143 | -1.811370215 | 0.003063159 |
| GO:0002478 | antigen processing and presentation of exogenous peptide antig | 23 | -0.597301849 | -1.811531829 | 0.002837558 |
| GO:0052547 | regulation of peptidase activity | 267 | -0.397289861 | -1.811699262 | 1.04098296084137e-06 |
| GO:0141087 | positive regulation of inflammasome-mediated signaling pathw | 21 | -0.611080569 | -1.812472562 | 0.002979066 |
| GO:0055008 | cardiac muscle tissue morphogenesis | 56 | -0.493927374 | -1.812727616 | 0.001012839 |
| GO:0048245 | eosinophil chemotaxis | 24 | -0.592015095 | -1.812935263 | 0.002524611 |
| GO:0001945 | lymph vessel development | 30 | -0.561948999 | -1.812940881 | 0.00239434 |
| GO:0002204 | somatic recombination of immunoglobulin genes involved in ir | 51 | -0.502954064 | -1.813532594 | 0.001053558 |
| GO:0002208 | somatic diversification of immunoglobulins involved in immun | 51 | -0.502954064 | -1.813532594 | 0.001053558 |
| GO:0045190 | isotype switching | 51 | -0.502954064 | -1.813532594 | 0.001053558 |
| GO:0034114 | regulation of heterotypic cell-cell adhesion | 22 | -0.604818878 | -1.814116663 | 0.002786809 |
| GO:0030099 | myeloid cell differentiation | 406 | -0.385135144 | -1.815502534 | 2.31113619350061e-08 |
| GO:0033079 | immature T cell proliferation | 13 | -0.693636107 | -1.815966694 | 0.003370385 |
| GO:0033080 | immature T cell proliferation in thymus | 13 | -0.693636107 | -1.815966694 | 0.003370385 |
| GO:0045628 | regulation of T-helper 2 cell differentiation | 12 | -0.708708765 | -1.816014489 | 0.003425713 |
| GO:0070189 | kynurenine metabolic process | 10 | -0.745637279 | -1.817514811 | 0.003238105 |
| GO:1903670 | regulation of sprouting angiogenesis | 39 | -0.532367238 | -1.818875543 | 0.001522297 |
| GO:0002043 | blood vessel endothelial cell proliferation involved in sprouting | 24 | -0.594019007 | -1.819071866 | 0.002270562 |
| GO:1904646 | cellular response to amyloid-beta | 36 | -0.541833249 | -1.819421008 | 0.001723226 |
| GO:0051928 | positive regulation of calcium ion transport | 114 | -0.440721337 | -1.820556822 | 0.000124349 |
| GO:0070741 | response to interleukin-6 | 31 | -0.559986777 | -1.821011818 | 0.001952675 |
| GO:0010595 | positive regulation of endothelial cell migration | 101 | -0.449252452 | -1.822459438 | 0.000209974 |

| | | | | | |
|---|---|---|---|---|---|
| GO:2000273 | positive regulation of signaling receptor activity | 27 | -0.578447298 | -1.822759627 | 0.002046843 |
| GO:0002756 | MyD88-independent toll-like receptor signaling pathway | 16 | -0.659045926 | -1.8233355 | 0.002908813 |
| GO:0055094 | response to lipoprotein particle | 31 | -0.561531925 | -1.826036458 | 0.001845047 |
| GO:0045638 | negative regulation of myeloid cell differentiation | 78 | -0.469295648 | -1.826474352 | 0.000299728 |
| GO:0071354 | cellular response to interleukin-6 | 27 | -0.579826989 | -1.827107206 | 0.001953094 |
| GO:0010758 | regulation of macrophage chemotaxis | 27 | -0.579877656 | -1.827266862 | 0.001953094 |
| GO:0060761 | negative regulation of response to cytokine stimulus | 77 | -0.470741735 | -1.828493711 | 0.000273071 |
| GO:0032309 | icosanoid secretion | 45 | -0.52035402 | -1.830364001 | 0.001114664 |
| GO:0046006 | regulation of activated T cell proliferation | 37 | -0.542052089 | -1.830570684 | 0.001491885 |
| GO:0008015 | blood circulation | 490 | -0.384599499 | -1.833791835 | 3.35378214151976e-10 |
| GO:0032494 | response to peptidoglycan | 13 | -0.700591674 | -1.834176642 | 0.002706521 |
| GO:0035150 | regulation of tube size | 141 | -0.43158975 | -1.834229806 | 1.90252768461728e-05 |
| GO:0031663 | lipopolysaccharide-mediated signaling pathway | 56 | -0.50015718 | -1.835591184 | 0.000784593 |
| GO:0000768 | syncytium formation by plasma membrane fusion | 58 | -0.496863186 | -1.835886455 | 0.000567779 |
| GO:0140253 | cell-cell fusion | 58 | -0.496863186 | -1.835886455 | 0.000567779 |
| GO:0002420 | natural killer cell mediated cytotoxicity directed against tumor | 10 | -0.753189943 | -1.83592467 | 0.002402465 |
| GO:0060337 | type I interferon-mediated signaling pathway | 77 | -0.472939866 | -1.837031871 | 0.000273071 |
| GO:0098742 | cell-cell adhesion via plasma-membrane adhesion molecules | 265 | -0.403203063 | -1.83712841 | 7.22001570172759e-07 |
| GO:0003158 | endothelium development | 126 | -0.439148815 | -1.83915259 | 8.3912246889175e-05 |
| GO:0045655 | regulation of monocyte differentiation | 18 | -0.645103626 | -1.839258781 | 0.002248925 |
| GO:0010574 | regulation of vascular endothelial growth factor production | 31 | -0.565844308 | -1.840059828 | 0.001537539 |
| GO:0034162 | toll-like receptor 9 signaling pathway | 17 | -0.655179628 | -1.84079324 | 0.002344124 |
| GO:0070371 | ERK1 and ERK2 cascade | 299 | -0.399858025 | -1.841689551 | 1.19191094888558e-07 |
| GO:0060193 | positive regulation of lipase activity | 52 | -0.509037135 | -1.842312522 | 0.000691324 |
| GO:0032616 | interleukin-13 production | 20 | -0.629107877 | -1.842579547 | 0.001977278 |
| GO:0032656 | regulation of interleukin-13 production | 20 | -0.629107877 | -1.842579547 | 0.001977278 |
| GO:0022600 | digestive system process | 100 | -0.455120397 | -1.843107336 | 0.000135825 |
| GO:2000109 | regulation of macrophage apoptotic process | 10 | -0.756467318 | -1.843913378 | 0.002158737 |
| GO:0035296 | regulation of tube diameter | 140 | -0.434393861 | -1.844051001 | 1.01769040845139e-05 |
| GO:0097746 | blood vessel diameter maintenance | 140 | -0.434393861 | -1.844051001 | 1.01769040845139e-05 |
| GO:0051091 | positive regulation of DNA-binding transcription factor activity | 218 | -0.412600394 | -1.845264058 | 2.0110815359124e-06 |
| GO:0010573 | vascular endothelial growth factor production | 35 | -0.552866124 | -1.845679589 | 0.001213776 |
| GO:0060670 | branching involved in labyrinthine layer morphogenesis | 13 | -0.705103266 | -1.845988168 | 0.00229799 |
| GO:1902042 | negative regulation of extrinsic apoptotic signaling pathway via | 26 | -0.591141833 | -1.846157838 | 0.001632525 |
| GO:0007249 | canonical NF-kappaB signal transduction | 293 | -0.401695195 | -1.846701226 | 1.04256929169256e-07 |
| GO:0071404 | cellular response to low-density lipoprotein particle stimulus | 30 | -0.572809933 | -1.847980059 | 0.001544736 |
| GO:0033209 | tumor necrosis factor-mediated signaling pathway | 103 | -0.454498593 | -1.848966155 | 7.30842250084133e-05 |
| GO:0140374 | antiviral innate immune response | 44 | -0.528386283 | -1.849784794 | 0.000882522 |
| GO:0001767 | establishment of lymphocyte polarity | 16 | -0.668614866 | -1.849809207 | 0.002194076 |
| GO:2000696 | regulation of epithelial cell differentiation involved in kidney dev | 16 | -0.668618769 | -1.849820003 | 0.002194076 |
| GO:0051208 | sequestering of calcium ion | 128 | -0.440713471 | -1.849996002 | 3.20456179282497e-05 |
| GO:0002418 | immune response to tumor cell | 22 | -0.616919773 | -1.850412547 | 0.001729744 |
| GO:0042670 | retinal cone cell differentiation | 12 | -0.722155835 | -1.850471624 | 0.002243842 |
| GO:0101023 | vascular endothelial cell proliferation | 23 | -0.61024009 | -1.85077168 | 0.001689782 |
| GO:1905562 | regulation of vascular endothelial cell proliferation | 23 | -0.61024009 | -1.85077168 | 0.001689782 |

| | | | | | |
|---|---|---|---|---|---|
| GO:0010893 | positive regulation of steroid biosynthetic process | 21 | -0.624404774 | -1.851992321 | 0.001835749 |
| GO:0045687 | positive regulation of glial cell differentiation | 43 | -0.531414901 | -1.852432573 | 0.000943828 |
| GO:0097048 | dendritic cell apoptotic process | 12 | -0.72295302 | -1.852514352 | 0.002175328 |
| GO:2000668 | regulation of dendritic cell apoptotic process | 12 | -0.72295302 | -1.852514352 | 0.002175328 |
| GO:0043270 | positive regulation of monoatomic ion transport | 202 | -0.417896679 | -1.853683339 | 4.45802219587165e-06 |
| GO:0046697 | decidualization | 24 | -0.60562343 | -1.854608237 | 0.001444903 |
| GO:0035456 | response to interferon-beta | 30 | -0.57486653 | -1.854614981 | 0.001359367 |
| GO:0034374 | low-density lipoprotein particle remodeling | 16 | -0.670419759 | -1.854802674 | 0.002061102 |
| GO:0035745 | T-helper 2 cell cytokine production | 13 | -0.708879395 | -1.855874223 | 0.002025635 |
| GO:2000551 | regulation of T-helper 2 cell cytokine production | 13 | -0.708879395 | -1.855874223 | 0.002025635 |
| GO:2000563 | positive regulation of CD4-positive, alpha-beta T cell proliferat | 10 | -0.761416148 | -1.855976309 | 0.001740917 |
| GO:0071605 | monocyte chemotactic protein-1 production | 20 | -0.634008238 | -1.856932101 | 0.001717962 |
| GO:0071637 | regulation of monocyte chemotactic protein-1 production | 20 | -0.634008238 | -1.856932101 | 0.001717962 |
| GO:0048659 | smooth muscle cell proliferation | 138 | -0.438563728 | -1.858117662 | 1.87472076814946e-05 |
| GO:0045639 | positive regulation of myeloid cell differentiation | 99 | -0.459590934 | -1.858324488 | 4.99170742656141e-05 |
| GO:0032728 | positive regulation of interferon-beta production | 38 | -0.547359477 | -1.859717057 | 0.000914186 |
| GO:0097396 | response to interleukin-17 | 16 | -0.672392626 | -1.860260866 | 0.001944749 |
| GO:0097398 | cellular response to interleukin-17 | 16 | -0.672392626 | -1.860260866 | 0.001944749 |
| GO:0030316 | osteoclast differentiation | 104 | -0.4565653 | -1.860339625 | 5.24905396478255e-05 |
| GO:1900227 | positive regulation of NLRP3 inflammasome complex assemb | 20 | -0.635666416 | -1.861788701 | 0.001669341 |
| GO:1905521 | regulation of macrophage migration | 42 | -0.536611933 | -1.862278059 | 0.000842535 |
| GO:0036035 | osteoclast development | 18 | -0.653428997 | -1.862995296 | 0.001723628 |
| GO:0098751 | bone cell development | 18 | -0.653428997 | -1.862995296 | 0.001723628 |
| GO:0030100 | regulation of endocytosis | 273 | -0.407923396 | -1.864090342 | 2.66461062629812e-07 |
| GO:0051969 | regulation of transmission of nerve impulse | 14 | -0.697618633 | -1.864358735 | 0.001887428 |
| GO:0042430 | indole-containing compound metabolic process | 23 | -0.614767804 | -1.864503595 | 0.001386896 |
| GO:0090077 | foam cell differentiation | 39 | -0.545913866 | -1.865158688 | 0.000880544 |
| GO:0030277 | maintenance of gastrointestinal epithelium | 23 | -0.615522311 | -1.866791907 | 0.001323131 |
| GO:0050922 | negative regulation of chemotaxis | 61 | -0.500886854 | -1.866954755 | 0.000338883 |
| GO:0002423 | natural killer cell mediated immune response to tumor cell | 11 | -0.746724014 | -1.867239691 | 0.001573904 |
| GO:0046007 | negative regulation of activated T cell proliferation | 14 | -0.699273563 | -1.86878147 | 0.001803168 |
| GO:0001916 | positive regulation of T cell mediated cytotoxicity | 29 | -0.584018255 | -1.869396994 | 0.001101886 |
| GO:0043304 | regulation of mast cell degranulation | 27 | -0.59330414 | -1.869575389 | 0.001171857 |
| GO:0046330 | positive regulation of JNK cascade | 91 | -0.468738596 | -1.86983126 | 0.000134726 |
| GO:0019755 | one-carbon compound transport | 22 | -0.623500582 | -1.870151274 | 0.001297308 |
| GO:0015670 | carbon dioxide transport | 12 | -0.729935985 | -1.870407691 | 0.001849885 |
| GO:0035590 | purinergic nucleotide receptor signaling pathway | 29 | -0.584662048 | -1.871457724 | 0.001055327 |
| GO:0001787 | natural killer cell proliferation | 14 | -0.700787319 | -1.872826926 | 0.001685204 |
| GO:0007599 | hemostasis | 224 | -0.417720602 | -1.872893062 | 9.17880933889278e-07 |
| GO:0048302 | regulation of isotype switching to IgG isotypes | 14 | -0.700910202 | -1.873155325 | 0.001668352 |
| GO:1901739 | regulation of myoblast fusion | 20 | -0.639582135 | -1.87325736 | 0.001426233 |
| GO:0046549 | retinal cone cell development | 10 | -0.76896404 | -1.874374538 | 0.001288278 |
| GO:0010669 | epithelial structure maintenance | 33 | -0.568608149 | -1.874897165 | 0.000854701 |
| GO:0016554 | cytidine to uridine editing | 12 | -0.731946291 | -1.875558953 | 0.001661471 |
| GO:0045671 | negative regulation of osteoclast differentiation | 32 | -0.572930569 | -1.876686575 | 0.00096432 |

| | | | | | |
|---|---|---|---|---|---|
| GO:0048660 | regulation of smooth muscle cell proliferation | 134 | -0.444485902 | -1.876886074 | 1.32036649207651e-05 |
| GO:0032782 | bile acid secretion | 10 | -0.770006478 | -1.876915515 | 0.001288278 |
| GO:0010720 | positive regulation of cell development | 431 | -0.396674182 | -1.877032708 | 3.23123355060581e-10 |
| GO:0001768 | establishment of T cell polarity | 15 | -0.690183106 | -1.877627921 | 0.001605459 |
| GO:0033632 | regulation of cell-cell adhesion mediated by integrin | 11 | -0.750998387 | -1.877928084 | 0.001400948 |
| GO:0072577 | endothelial cell apoptotic process | 55 | -0.51367722 | -1.879331452 | 0.000328792 |
| GO:0002862 | negative regulation of inflammatory response to antigenic stim | 26 | -0.602070825 | -1.880289486 | 0.000988933 |
| GO:0002381 | immunoglobulin production involved in immunoglobulin-media | 56 | -0.512400494 | -1.880524497 | 0.000342368 |
| GO:0034138 | toll-like receptor 3 signaling pathway | 26 | -0.602527799 | -1.881716631 | 0.000973236 |
| GO:1903238 | positive regulation of leukocyte tethering or rolling | 12 | -0.73438047 | -1.88179636 | 0.001541571 |
| GO:0042542 | response to hydrogen peroxide | 92 | -0.471295572 | -1.882786824 | 6.47222741966198e-05 |
| GO:0007596 | blood coagulation | 217 | -0.421476469 | -1.883812127 | 1.33206226132397e-06 |
| GO:0045637 | regulation of myeloid cell differentiation | 193 | -0.427044567 | -1.884081655 | 7.3600313364618e-07 |
| GO:0034109 | homotypic cell-cell adhesion | 93 | -0.470886426 | -1.884809412 | 4.8180402215255e-05 |
| GO:0048644 | muscle organ morphogenesis | 71 | -0.493008873 | -1.888635975 | 0.000167617 |
| GO:0032608 | interferon-beta production | 56 | -0.514678833 | -1.888886065 | 0.000328103 |
| GO:0032648 | regulation of interferon-beta production | 56 | -0.514678833 | -1.888886065 | 0.000328103 |
| GO:0062208 | positive regulation of pattern recognition receptor signaling patl | 62 | -0.505372742 | -1.889330012 | 0.00026764 |
| GO:1901741 | positive regulation of myoblast fusion | 15 | -0.694761016 | -1.890082024 | 0.001454947 |
| GO:0051282 | regulation of sequestering of calcium ion | 124 | -0.452613336 | -1.891195684 | 1.177453831335e-05 |
| GO:0052548 | regulation of endopeptidase activity | 244 | -0.41870769 | -1.893012771 | 2.75654460942772e-07 |
| GO:0071357 | cellular response to type I interferon | 78 | -0.486767633 | -1.894474411 | 0.000132941 |
| GO:0050792 | regulation of viral process | 159 | -0.439329017 | -1.8945809 | 4.45800745791625e-06 |
| GO:0032481 | positive regulation of type I interferon production | 67 | -0.499759013 | -1.895243609 | 9.5109808222236e-05 |
| GO:0045953 | negative regulation of natural killer cell mediated cytotoxicity | 15 | -0.696907594 | -1.895921744 | 0.001270988 |
| GO:0090025 | regulation of monocyte chemotaxis | 25 | -0.612731462 | -1.896967621 | 0.000977101 |
| GO:0002763 | positive regulation of myeloid leukocyte differentiation | 58 | -0.51371909 | -1.898168237 | 0.000269695 |
| GO:0042832 | defense response to protozoan | 24 | -0.619965512 | -1.898528176 | 0.000793903 |
| GO:0050817 | coagulation | 221 | -0.424103078 | -1.899238373 | 3.68169618811561e-07 |
| GO:1903236 | regulation of leukocyte tethering or rolling | 17 | -0.676167779 | -1.899761567 | 0.001287617 |
| GO:0034104 | negative regulation of tissue remodeling | 21 | -0.640592658 | -1.900005784 | 0.000950081 |
| GO:0035850 | epithelial cell differentiation involved in kidney development | 48 | -0.533376857 | -1.900262991 | 0.000305068 |
| GO:0002431 | Fc receptor mediated stimulatory signaling pathway | 31 | -0.584375325 | -1.900320537 | 0.000876397 |
| GO:0010572 | positive regulation of platelet activation | 11 | -0.760266945 | -1.901104811 | 0.001106922 |
| GO:0014009 | glial cell proliferation | 54 | -0.521882984 | -1.90294294 | 0.000243539 |
| GO:0010742 | macrophage derived foam cell differentiation | 38 | -0.560114242 | -1.903052845 | 0.000509547 |
| GO:0050829 | defense response to Gram-negative bacterium | 75 | -0.492269008 | -1.903692985 | 8.11194097307092e-05 |
| GO:0061517 | macrophage proliferation | 11 | -0.761337849 | -1.903782686 | 0.00107233 |
| GO:0045069 | regulation of viral genome replication | 83 | -0.48431122 | -1.903919946 | 7.85543209484673e-05 |
| GO:0060340 | positive regulation of type I interferon-mediated signaling pathv | 19 | -0.658568366 | -1.904745281 | 0.00097766 |
| GO:0002762 | negative regulation of myeloid leukocyte differentiation | 50 | -0.53018729 | -1.904916045 | 0.000260157 |
| GO:0060263 | regulation of respiratory burst | 17 | -0.678062066 | -1.905083758 | 0.001238094 |
| GO:0001936 | regulation of endothelial cell proliferation | 129 | -0.453838807 | -1.90693333 | 5.03310737518003e-06 |
| GO:0002664 | regulation of T cell tolerance induction | 11 | -0.762927787 | -1.907758447 | 0.000968557 |
| GO:0045581 | negative regulation of T cell differentiation | 46 | -0.539981657 | -1.90797329 | 0.000350759 |

| | | | | | |
|---|---|---|---|---|---|
| GO:0043491 | phosphatidylinositol 3-kinase/protein kinase B signal transduct | 267 | -0.418509302 | -1.908462982 | 5.42825380299597e-08 |
| GO:0070372 | regulation of ERK1 and ERK2 cascade | 276 | -0.417337261 | -1.908926581 | 5.22414449765633e-08 |
| GO:0045623 | negative regulation of T-helper cell differentiation | 17 | -0.679679571 | -1.909628302 | 0.00118857 |
| GO:0071353 | cellular response to interleukin-4 | 33 | -0.579393836 | -1.910461293 | 0.0004884 |
| GO:1903900 | regulation of viral life cycle | 137 | -0.451752222 | -1.912389826 | 4.16802504052821e-06 |
| GO:0099509 | regulation of presynaptic cytosolic calcium ion concentration | 13 | -0.730953076 | -1.91366399 | 0.000885152 |
| GO:0045670 | regulation of osteoclast differentiation | 69 | -0.502565483 | -1.915711948 | 0.00010716 |
| GO:0002645 | positive regulation of tolerance induction | 11 | -0.766522957 | -1.916748442 | 0.000830191 |
| GO:0016045 | detection of bacterium | 12 | -0.748070741 | -1.91687668 | 0.000856428 |
| GO:0034340 | response to type I interferon | 83 | -0.48770328 | -1.917254783 | 6.18510440568629e-05 |
| GO:0002283 | neutrophil activation involved in immune response | 19 | -0.663134166 | -1.917950722 | 0.000782128 |
| GO:0006568 | tryptophan metabolic process | 10 | -0.787339162 | -1.919164488 | 0.000609321 |
| GO:0006953 | acute-phase response | 44 | -0.548918365 | -1.921663899 | 0.000294174 |
| GO:0002534 | cytokine production involved in inflammatory response | 59 | -0.518561984 | -1.921737782 | 0.000207134 |
| GO:1900015 | regulation of cytokine production involved in inflammatory resp | 59 | -0.518561984 | -1.921737782 | 0.000207134 |
| GO:0014015 | positive regulation of gliogenesis | 68 | -0.505259779 | -1.922089197 | 4.86725459491763e-05 |
| GO:0048661 | positive regulation of smooth muscle cell proliferation | 81 | -0.490725151 | -1.922277624 | 3.61384229265469e-05 |
| GO:0019882 | antigen processing and presentation | 90 | -0.483084903 | -1.923528772 | 2.07157678275701e-05 |
| GO:0007157 | heterophilic cell-cell adhesion via plasma membrane cell adhe | 51 | -0.533647287 | -1.924205048 | 0.000274214 |
| GO:0034142 | toll-like receptor 4 signaling pathway | 57 | -0.522759578 | -1.925032375 | 0.000199251 |
| GO:0046636 | negative regulation of alpha-beta T cell activation | 42 | -0.55478653 | -1.92535186 | 0.000428658 |
| GO:0043405 | regulation of MAP kinase activity | 127 | -0.459468974 | -1.926446931 | 1.04972802713458e-05 |
| GO:0072677 | eosinophil migration | 28 | -0.606322189 | -1.926550776 | 0.000622103 |
| GO:0070102 | interleukin-6-mediated signaling pathway | 17 | -0.685721438 | -1.926603535 | 0.000957459 |
| GO:0032490 | detection of molecule of bacterial origin | 13 | -0.736079801 | -1.927085959 | 0.000765997 |
| GO:0034135 | regulation of toll-like receptor 2 signaling pathway | 12 | -0.752266371 | -1.927627676 | 0.000753657 |
| GO:0060075 | regulation of resting membrane potential | 10 | -0.791563812 | -1.929462209 | 0.000591912 |
| GO:0086091 | regulation of heart rate by cardiac conduction | 39 | -0.564767561 | -1.929573855 | 0.000313414 |
| GO:0046850 | regulation of bone remodeling | 51 | -0.535154852 | -1.929640968 | 0.000259781 |
| GO:1903672 | positive regulation of sprouting angiogenesis | 21 | -0.650700479 | -1.929985708 | 0.000676329 |
| GO:0001562 | response to protozoan | 27 | -0.612769139 | -1.930912032 | 0.000515617 |
| GO:0033141 | positive regulation of peptidyl-serine phosphorylation of STAT | 17 | -0.68850811 | -1.934432973 | 0.000891427 |
| GO:0071715 | icosanoid transport | 60 | -0.520684844 | -1.934900036 | 8.70016318687904e-05 |
| GO:0010543 | regulation of platelet activation | 50 | -0.538538885 | -1.934922589 | 0.000250687 |
| GO:0051209 | release of sequestered calcium ion into cytosol | 121 | -0.464694658 | -1.93514525 | 5.26038786152977e-06 |
| GO:0034112 | positive regulation of homotypic cell-cell adhesion | 15 | -0.711381292 | -1.935297118 | 0.000752559 |
| GO:0051283 | negative regulation of sequestering of calcium ion | 122 | -0.464347056 | -1.935860443 | 4.23914593373023e-06 |
| GO:0002753 | cytoplasmic pattern recognition receptor signaling pathway | 154 | -0.450612542 | -1.936259757 | 1.29806438291282e-06 |
| GO:1904018 | positive regulation of vasculature development | 163 | -0.447580056 | -1.936671685 | 9.93137503938784e-07 |
| GO:0050830 | defense response to Gram-positive bacterium | 92 | -0.485123646 | -1.938028837 | 2.88516485248686e-05 |
| GO:0045766 | positive regulation of angiogenesis | 160 | -0.44942614 | -1.939813265 | 1.75233211552114e-06 |
| GO:0030168 | platelet activation | 130 | -0.461274515 | -1.939858796 | 2.67648993043515e-06 |
| GO:0050832 | defense response to fungus | 49 | -0.543182044 | -1.943691648 | 0.000128302 |
| GO:0043371 | negative regulation of CD4-positive, alpha-beta T cell differenti | 20 | -0.663662222 | -1.943784972 | 0.000599666 |
| GO:0002719 | negative regulation of cytokine production involved in immune | 33 | -0.589712088 | -1.944484129 | 0.000351038 |

| | | | | | |
|---|---|---|---|---|---|
| GO:0070670 | response to interleukin-4 | 36 | -0.580087259 | -1.947874088 | 0.000287204 |
| GO:0045776 | negative regulation of blood pressure | 46 | -0.551746169 | -1.949542062 | 0.000127773 |
| GO:2000353 | positive regulation of endothelial cell apoptotic process | 18 | -0.683863875 | -1.949768359 | 0.000541712 |
| GO:0070229 | negative regulation of lymphocyte apoptotic process | 36 | -0.580671252 | -1.949835076 | 0.000287204 |
| GO:0045124 | regulation of bone resorption | 43 | -0.559457931 | -1.950186365 | 0.000309693 |
| GO:0002430 | complement receptor mediated signaling pathway | 14 | -0.731587117 | -1.95513819 | 0.000539265 |
| GO:0032736 | positive regulation of interleukin-13 production | 13 | -0.747366215 | -1.956634237 | 0.000510664 |
| GO:0010951 | negative regulation of endopeptidase activity | 104 | -0.481039373 | -1.960062683 | 8.18303236031986e-06 |
| GO:0071402 | cellular response to lipoprotein particle stimulus | 34 | -0.590836153 | -1.960317553 | 0.000319693 |
| GO:0002888 | positive regulation of myeloid leukocyte mediated immunity | 19 | -0.678263352 | -1.961708133 | 0.000439947 |
| GO:0033628 | regulation of cell adhesion mediated by integrin | 47 | -0.552701967 | -1.961961465 | 0.000112533 |
| GO:0002520 | immune system development | 189 | -0.446208717 | -1.963939563 | 6.14346989230229e-08 |
| GO:0001779 | natural killer cell differentiation | 24 | -0.641380732 | -1.964108272 | 0.000381073 |
| GO:0045779 | negative regulation of bone resorption | 16 | -0.710043636 | -1.964427237 | 0.00046541 |
| GO:0070424 | regulation of nucleotide-binding domain, leucine rich repeat co | 15 | -0.722236621 | -1.964828802 | 0.000451535 |
| GO:0002604 | regulation of dendritic cell antigen processing and presentation | 11 | -0.786297351 | -1.966195804 | 0.000380504 |
| GO:0045940 | positive regulation of steroid metabolic process | 25 | -0.635161374 | -1.966408833 | 0.000425512 |
| GO:0045625 | regulation of T-helper 1 cell differentiation | 11 | -0.786433659 | -1.966536652 | 0.000380504 |
| GO:0051896 | regulation of phosphatidylinositol 3-kinase/protein kinase B sig | 223 | -0.439074881 | -1.968187582 | 3.13114612171078e-08 |
| GO:2000508 | regulation of dendritic cell chemotaxis | 10 | -0.807725971 | -1.96885799 | 0.000365593 |
| GO:1904645 | response to amyloid-beta | 46 | -0.557752959 | -1.970766478 | 0.00010634 |
| GO:0045064 | T-helper 2 cell differentiation | 18 | -0.691684309 | -1.972065244 | 0.000426803 |
| GO:0050860 | negative regulation of T cell receptor signaling pathway | 24 | -0.644242846 | -1.972872959 | 0.000365195 |
| GO:0002832 | negative regulation of response to biotic stimulus | 108 | -0.482033637 | -1.975875619 | 8.39794016677867e-06 |
| GO:0006586 | indolalkylamine metabolic process | 12 | -0.771535198 | -1.97700264 | 0.000393957 |
| GO:0070227 | lymphocyte apoptotic process | 77 | -0.509008273 | -1.977131741 | 3.06931223103768e-05 |
| GO:2001057 | reactive nitrogen species metabolic process | 74 | -0.512863983 | -1.978162679 | 3.24142858628183e-05 |
| GO:0032727 | positive regulation of interferon-alpha production | 22 | -0.660003755 | -1.979640274 | 0.000320323 |
| GO:0002716 | negative regulation of natural killer cell mediated immunity | 16 | -0.716748157 | -1.98297616 | 0.000349058 |
| GO:0043406 | positive regulation of MAP kinase activity | 78 | -0.509573972 | -1.983235504 | 2.83439344988416e-05 |
| GO:0010466 | negative regulation of peptidase activity | 111 | -0.482165643 | -1.983528418 | 4.78923141259101e-06 |
| GO:0097553 | calcium ion transmembrane import into cytosol | 172 | -0.4558476 | -1.985243778 | 2.02916337969275e-07 |
| GO:0070234 | positive regulation of T cell apoptotic process | 12 | -0.775350836 | -1.986779933 | 0.000274057 |
| GO:0009074 | aromatic amino acid family catabolic process | 15 | -0.731080954 | -1.988889615 | 0.000351194 |
| GO:0002360 | T cell lineage commitment | 33 | -0.603520571 | -1.990015461 | 0.000228938 |
| GO:0002691 | regulation of cellular extravasation | 40 | -0.579440856 | -1.990496048 | 9.6501922830337e-05 |
| GO:0071356 | cellular response to tumor necrosis factor | 215 | -0.446101419 | -1.991625409 | 2.75506892782329e-08 |
| GO:0150078 | positive regulation of neuroinflammatory response | 15 | -0.733334704 | -1.995020893 | 0.000351194 |
| GO:0043011 | myeloid dendritic cell differentiation | 19 | -0.690749885 | -1.997822326 | 0.000309592 |
| GO:0060143 | positive regulation of syncytium formation by plasma membra | 23 | -0.659098325 | -1.99895178 | 0.000255061 |
| GO:0002577 | regulation of antigen processing and presentation | 18 | -0.701489829 | -2.000021821 | 0.000295479 |
| GO:0010934 | macrophage cytokine production | 36 | -0.596359893 | -2.00251594 | 0.000197068 |
| GO:0010935 | regulation of macrophage cytokine production | 36 | -0.596359893 | -2.00251594 | 0.000197068 |
| GO:0141060 | disruption of anatomical structure in another organism | 81 | -0.51121684 | -2.002548048 | 5.94844434242451e-06 |
| GO:0006809 | nitric oxide biosynthetic process | 66 | -0.529417971 | -2.002835696 | 3.95355429541565e-05 |

| | | | | | |
|---|---|---|---|---|---|
| GO:0046209 | nitric oxide metabolic process | 73 | -0.521228436 | -2.005747505 | 2.63447847016739e-05 |
| GO:0032621 | interleukin-18 production | 11 | -0.802190264 | -2.005937231 | 0.000125845 |
| GO:0032661 | regulation of interleukin-18 production | 11 | -0.802190264 | -2.005937231 | 0.000125845 |
| GO:0070431 | nucleotide-binding oligomerization domain containing 2 signali | 19 | -0.694310036 | -2.008119177 | 0.000277004 |
| GO:0046717 | acid secretion | 40 | -0.5846482 | -2.008384323 | 6.88890080126235e-05 |
| GO:0045730 | respiratory burst | 37 | -0.594714833 | -2.008418676 | 9.43483562548239e-05 |
| GO:0007162 | negative regulation of cell adhesion | 275 | -0.439332619 | -2.008885262 | 1.30594006960103e-09 |
| GO:0033089 | positive regulation of T cell differentiation in thymus | 11 | -0.803863537 | -2.010121376 | 0.000108286 |
| GO:0001937 | negative regulation of endothelial cell proliferation | 41 | -0.582562001 | -2.011590208 | 0.000124608 |
| GO:0002523 | leukocyte migration involved in inflammatory response | 19 | -0.69602946 | -2.013092184 | 0.000277004 |
| GO:0042976 | activation of Janus kinase activity | 12 | -0.785779824 | -2.013503455 | 5.91244235370552e-05 |
| GO:0090594 | inflammatory response to wounding | 26 | -0.644788922 | -2.013699684 | 0.000218882 |
| GO:0002221 | pattern recognition receptor signaling pathway | 238 | -0.446916785 | -2.015448318 | 1.11181891310564e-09 |
| GO:0006957 | complement activation, alternative pathway | 16 | -0.729143886 | -2.01727054 | 0.000232705 |
| GO:0002517 | T cell tolerance induction | 13 | -0.770947073 | -2.018369854 | 0.000154578 |
| GO:0046851 | negative regulation of bone remodeling | 18 | -0.708330727 | -2.019525946 | 0.000329386 |
| GO:0001914 | regulation of T cell mediated cytotoxicity | 39 | -0.59117318 | -2.019790779 | 8.67058968775864e-05 |
| GO:0002752 | cell surface pattern recognition receptor signaling pathway | 71 | -0.52767201 | -2.02142476 | 1.9873032536315e-05 |
| GO:0070555 | response to interleukin-1 | 125 | -0.483566337 | -2.022683724 | 1.36181666097829e-06 |
| GO:0045824 | negative regulation of innate immune response | 79 | -0.518900391 | -2.023811746 | 6.15916247033925e-06 |
| GO:0033630 | positive regulation of cell adhesion mediated by integrin | 21 | -0.68288307 | -2.025439673 | 0.000175296 |
| GO:0060333 | type II interferon-mediated signaling pathway | 26 | -0.648905351 | -2.026555447 | 0.000186852 |
| GO:0032823 | regulation of natural killer cell differentiation | 13 | -0.77415118 | -2.026758332 | 0.000131536 |
| GO:0060100 | positive regulation of phagocytosis, engulfment | 13 | -0.774247509 | -2.027010525 | 0.000131536 |
| GO:1905155 | positive regulation of membrane invagination | 13 | -0.774247509 | -2.027010525 | 0.000131536 |
| GO:0002347 | response to tumor cell | 37 | -0.600506456 | -2.027977637 | 7.99281921642171e-05 |
| GO:0050862 | positive regulation of T cell receptor signaling pathway | 17 | -0.722117518 | -2.028861994 | 0.000231111 |
| GO:2000319 | regulation of T-helper 17 cell differentiation | 21 | -0.684089815 | -2.029018892 | 0.000171663 |
| GO:0070669 | response to interleukin-2 | 11 | -0.811568813 | -2.029389001 | 7.42621383544009e-05 |
| GO:0034154 | toll-like receptor 7 signaling pathway | 10 | -0.832822784 | -2.030032278 | 7.86579581951982e-05 |
| GO:0032817 | regulation of natural killer cell proliferation | 11 | -0.812099321 | -2.030715576 | 7.23206725565593e-05 |
| GO:0032695 | negative regulation of interleukin-12 production | 18 | -0.712399481 | -2.031126393 | 0.000258671 |
| GO:0002524 | hypersensitivity | 15 | -0.746912043 | -2.031957743 | 7.93145926449255e-05 |
| GO:2000108 | positive regulation of leukocyte apoptotic process | 23 | -0.670475281 | -2.033456476 | 0.000144763 |
| GO:0034612 | response to tumor necrosis factor | 235 | -0.452751146 | -2.039377461 | 1.94248238487012e-09 |
| GO:0032479 | regulation of type I interferon production | 115 | -0.493294718 | -2.04024903 | 6.84799389970402e-07 |
| GO:0032606 | type I interferon production | 115 | -0.493294718 | -2.04024903 | 6.84799389970402e-07 |
| GO:0043029 | T cell homeostasis | 45 | -0.58026736 | -2.041111331 | 2.76461506701327e-05 |
| GO:2001185 | regulation of CD8-positive, alpha-beta T cell activation | 17 | -0.726740796 | -2.041851561 | 0.00014246 |
| GO:1903587 | regulation of blood vessel endothelial cell proliferation involvec | 17 | -0.726968467 | -2.042491225 | 0.00014246 |
| GO:0071902 | positive regulation of protein serine/threonine kinase activity | 121 | -0.490521796 | -2.042698158 | 3.27777174004739e-07 |
| GO:0061760 | antifungal innate immune response | 21 | -0.689082233 | -2.043826468 | 0.000128067 |
| GO:0032703 | negative regulation of interleukin-2 production | 27 | -0.648687612 | -2.044095624 | 0.000113687 |
| GO:0045428 | regulation of nitric oxide biosynthetic process | 55 | -0.558724739 | -2.044141602 | 1.73792651565997e-05 |
| GO:0001959 | regulation of cytokine-mediated signaling pathway | 151 | -0.477018806 | -2.044856965 | 1.290027527901e-07 |

| | | | | | |
|---|---|---|---|---|---|
| GO:0045342 | MHC class II biosynthetic process | 19 | -0.707214058 | -2.045440853 | 7.54500262758713e-05 |
| GO:0019226 | transmission of nerve impulse | 70 | -0.535247505 | -2.04607658 | 9.03565324916278e-06 |
| GO:0019731 | antibacterial humoral response | 51 | -0.567670043 | -2.046883004 | 1.45379942529126e-05 |
| GO:0032700 | negative regulation of interleukin-17 production | 14 | -0.765981194 | -2.047055028 | 8.84366181906875e-05 |
| GO:1903707 | negative regulation of hemopoiesis | 109 | -0.499200408 | -2.048822579 | 1.34327875383877e-06 |
| GO:0001913 | T cell mediated cytotoxicity | 47 | -0.577311884 | -2.049320856 | 3.35509789039201e-05 |
| GO:0001771 | immunological synapse formation | 16 | -0.742247802 | -2.053524215 | 6.57713082434127e-05 |
| GO:0035771 | interleukin-4-mediated signaling pathway | 10 | -0.842881226 | -2.054550054 | 4.15664811723342e-05 |
| GO:0072540 | T-helper 17 cell lineage commitment | 17 | -0.731617362 | -2.055552764 | 0.000114527 |
| GO:0002281 | macrophage activation involved in immune response | 18 | -0.721431125 | -2.05687657 | 0.000143514 |
| GO:0002863 | positive regulation of inflammatory response to antigenic stimu | 10 | -0.844462312 | -2.058404 | 3.91357213686096e-05 |
| GO:0043374 | CD8-positive, alpha-beta T cell differentiation | 15 | -0.756775862 | -2.058792045 | 4.69345202229818e-05 |
| GO:0043300 | regulation of leukocyte degranulation | 42 | -0.593349339 | -2.059181668 | 5.10221585877989e-05 |
| GO:0090023 | positive regulation of neutrophil chemotaxis | 25 | -0.665293991 | -2.059697008 | 6.76672232262423e-05 |
| GO:0002761 | regulation of myeloid leukocyte differentiation | 120 | -0.495300905 | -2.060602448 | 7.9044893105794e-07 |
| GO:0001773 | myeloid dendritic cell activation | 27 | -0.654117383 | -2.06120551 | 8.19960457404634e-05 |
| GO:0034105 | positive regulation of tissue remodeling | 11 | -0.824631936 | -2.062054324 | 4.08124663987244e-05 |
| GO:0050850 | positive regulation of calcium-mediated signaling | 53 | -0.567868433 | -2.063053139 | 3.95009170486547e-05 |
| GO:0140888 | interferon-mediated signaling pathway | 100 | -0.50970306 | -2.064151498 | 1.1822407498084e-06 |
| GO:0051897 | positive regulation of phosphatidylinositol 3-kinase/protein kina | 157 | -0.479702387 | -2.065023361 | 2.73337944365868e-08 |
| GO:0060142 | regulation of syncytium formation by plasma membrane fusior | 28 | -0.649909229 | -2.065045863 | 8.24902003767658e-05 |
| GO:0045620 | negative regulation of lymphocyte differentiation | 56 | -0.563123924 | -2.066680936 | 1.27841949096451e-05 |
| GO:0046427 | positive regulation of receptor signaling pathway via JAK-STA | 46 | -0.585001146 | -2.067045326 | 1.79709405040689e-05 |
| GO:0070231 | T cell apoptotic process | 55 | -0.56509983 | -2.067465412 | 1.00304280293851e-05 |
| GO:0048535 | lymph node development | 18 | -0.72521181 | -2.067655703 | 0.000117589 |
| GO:0002643 | regulation of tolerance induction | 16 | -0.747863234 | -2.06906003 | 4.52564950635642e-05 |
| GO:0045916 | negative regulation of complement activation | 12 | -0.80756905 | -2.069336755 | 2.17756748954152e-05 |
| GO:0002864 | regulation of acute inflammatory response to antigenic stimulu | 13 | -0.790633702 | -2.069910226 | 5.58909443816037e-05 |
| GO:0080164 | regulation of nitric oxide metabolic process | 57 | -0.562131909 | -2.070018742 | 7.03462424146811e-06 |
| GO:0071624 | positive regulation of granulocyte chemotaxis | 26 | -0.662846959 | -2.070095604 | 9.65082382488022e-05 |
| GO:0002686 | negative regulation of leukocyte migration | 49 | -0.57857575 | -2.070342468 | 1.41920502400166e-05 |
| GO:0051767 | nitric-oxide synthase biosynthetic process | 19 | -0.716386155 | -2.071968864 | 4.7099558863675e-05 |
| GO:0051769 | regulation of nitric-oxide synthase biosynthetic process | 19 | -0.716386155 | -2.071968864 | 4.7099558863675e-05 |
| GO:0070230 | positive regulation of lymphocyte apoptotic process | 15 | -0.762081789 | -2.073226703 | 3.52593787074096e-05 |
| GO:1904894 | positive regulation of receptor signaling pathway via STAT | 50 | -0.577254777 | -2.074025365 | 1.59646821072002e-05 |
| GO:1902622 | regulation of neutrophil migration | 45 | -0.589722327 | -2.074369516 | 1.57929156594514e-05 |
| GO:0002295 | T-helper cell lineage commitment | 20 | -0.708248234 | -2.074371912 | 6.32211419382736e-05 |
| GO:0043369 | CD4-positive or CD8-positive, alpha-beta T cell lineage comm | 27 | -0.658318762 | -2.074444578 | 6.2703070697258e-05 |
| GO:0140895 | cell surface toll-like receptor signaling pathway | 65 | -0.550094091 | -2.074677613 | 4.50097097421376e-06 |
| GO:0033631 | cell-cell adhesion mediated by integrin | 17 | -0.738571407 | -2.075090853 | 8.0145058884343e-05 |
| GO:0071347 | cellular response to interleukin-1 | 100 | -0.51260707 | -2.075911907 | 9.13126746456665e-07 |
| GO:0045589 | regulation of regulatory T cell differentiation | 31 | -0.638928693 | -2.077721741 | 6.08393955402174e-05 |
| GO:1904037 | positive regulation of epithelial cell apoptotic process | 33 | -0.630310971 | -2.078352647 | 4.49736325896847e-05 |
| GO:0031640 | killing of cells of another organism | 78 | -0.534313681 | -2.079521169 | 3.63159443254e-06 |
| GO:0141061 | disruption of cell in another organism | 78 | -0.534313681 | -2.079521169 | 3.63159443254e-06 |

| | | | | | |
|---|---|---|---|---|---|
| GO:0038094 | Fc-gamma receptor signaling pathway | 30 | -0.644927766 | -2.080644174 | 2.91177991378984e-05 |
| GO:0045649 | regulation of macrophage differentiation | 23 | -0.686178778 | -2.081082956 | 6.39735250708625e-05 |
| GO:0045591 | positive regulation of regulatory T cell differentiation | 20 | -0.710577253 | -2.081193322 | 5.77633002303988e-05 |
| GO:0038095 | Fc-epsilon receptor signaling pathway | 25 | -0.672293947 | -2.081368312 | 4.64386618012725e-05 |
| GO:0043303 | mast cell degranulation | 50 | -0.579395041 | -2.081715145 | 1.41573849537635e-05 |
| GO:0002573 | myeloid leukocyte differentiation | 228 | -0.463743813 | -2.082905095 | 7.49820128981541e-10 |
| GO:0051092 | positive regulation of NF-kappaB transcription factor activity | 135 | -0.492790155 | -2.083278677 | 1.62913960164285e-07 |
| GO:0050848 | regulation of calcium-mediated signaling | 93 | -0.52124121 | -2.086363682 | 4.70771485466404e-07 |
| GO:0002532 | production of molecular mediator involved in inflammatory res | 83 | -0.531015382 | -2.087522934 | 1.26108008981374e-06 |
| GO:0035739 | CD4-positive, alpha-beta T cell proliferation | 21 | -0.704044677 | -2.088205263 | 5.19694003168194e-05 |
| GO:2000561 | regulation of CD4-positive, alpha-beta T cell proliferation | 21 | -0.704044677 | -2.088205263 | 5.19694003168194e-05 |
| GO:0051607 | defense response to virus | 287 | -0.455771972 | -2.091623183 | 1e-10 |
| GO:2000515 | negative regulation of CD4-positive, alpha-beta T cell activatio | 31 | -0.64382667 | -2.09364939 | 4.35803391966718e-05 |
| GO:0009595 | detection of biotic stimulus | 34 | -0.631210044 | -2.094272877 | 1.68155582498337e-05 |
| GO:0002269 | leukocyte activation involved in inflammatory response | 48 | -0.587960035 | -2.094726609 | 1.72573454323207e-05 |
| GO:0043410 | positive regulation of MAPK cascade | 443 | -0.442275985 | -2.096174559 | 1e-10 |
| GO:0002467 | germinal center formation | 14 | -0.784958327 | -2.097770679 | 3.69421114210552e-05 |
| GO:0071639 | positive regulation of monocyte chemotactic protein-1 producti | 14 | -0.785001098 | -2.097884981 | 3.69421114210552e-05 |
| GO:0035589 | G protein-coupled purinergic nucleotide receptor signaling path | 12 | -0.819148535 | -2.099008341 | 1.02423018102142e-05 |
| GO:0002363 | alpha-beta T cell lineage commitment | 25 | -0.679031523 | -2.102227307 | 3.32272675710882e-05 |
| GO:0140894 | endolysosomal toll-like receptor signaling pathway | 45 | -0.598354539 | -2.104733633 | 8.46598210622704e-06 |
| GO:0002638 | negative regulation of immunoglobulin production | 12 | -0.821617619 | -2.105335189 | 8.93982812503917e-06 |
| GO:0045346 | regulation of MHC class II biosynthetic process | 18 | -0.738857554 | -2.106561164 | 5.85057517895505e-05 |
| GO:0030889 | negative regulation of B cell proliferation | 16 | -0.762082357 | -2.108399068 | 1.78982177369888e-05 |
| GO:0045063 | T-helper 1 cell differentiation | 22 | -0.703238473 | -2.109320126 | 4.49492539880747e-05 |
| GO:0034134 | toll-like receptor 2 signaling pathway | 17 | -0.750851176 | -2.109592103 | 4.86434192395818e-05 |
| GO:0009620 | response to fungus | 60 | -0.567733095 | -2.109734514 | 3.1880395090119e-06 |
| GO:1902624 | positive regulation of neutrophil migration | 34 | -0.636168538 | -2.110724514 | 1.25852942773347e-05 |
| GO:0043032 | positive regulation of macrophage activation | 26 | -0.677013803 | -2.114339183 | 4.97790477100082e-05 |
| GO:0032102 | negative regulation of response to external stimulus | 391 | -0.449743024 | -2.114457872 | 1e-10 |
| GO:0060759 | regulation of response to cytokine stimulus | 160 | -0.489937226 | -2.114667232 | 1.4176445934572e-08 |
| GO:0001782 | B cell homeostasis | 33 | -0.641328555 | -2.114681421 | 2.40811315415342e-05 |
| GO:0070374 | positive regulation of ERK1 and ERK2 cascade | 196 | -0.478574784 | -2.115350727 | 6.26029761955972e-10 |
| GO:0035723 | interleukin-15-mediated signaling pathway | 11 | -0.846283647 | -2.116196059 | 1.18965177109847e-05 |
| GO:0071350 | cellular response to interleukin-15 | 11 | -0.846283647 | -2.116196059 | 1.18965177109847e-05 |
| GO:0090022 | regulation of neutrophil chemotaxis | 33 | -0.64197404 | -2.116809809 | 2.19501162788163e-05 |
| GO:0032602 | chemokine production | 93 | -0.529446293 | -2.119206037 | 1.92152562923733e-07 |
| GO:0002468 | dendritic cell antigen processing and presentation | 13 | -0.809966577 | -2.120524453 | 1.83293202289941e-05 |
| GO:0043277 | apoptotic cell clearance | 44 | -0.606136315 | -2.121973591 | 1.29771603855271e-05 |
| GO:0036006 | cellular response to macrophage colony-stimulating factor stin | 13 | -0.811024008 | -2.123292851 | 1.73833042797809e-05 |
| GO:0060907 | positive regulation of macrophage cytokine production | 23 | -0.700834985 | -2.125533153 | 2.78231325485095e-05 |
| GO:2000318 | positive regulation of T-helper 17 type immune response | 17 | -0.756544638 | -2.125588458 | 3.66487268949292e-05 |
| GO:0002448 | mast cell mediated immunity | 53 | -0.585217911 | -2.126083399 | 1.02757565986664e-05 |
| GO:0097529 | myeloid leukocyte migration | 223 | -0.474350287 | -2.126312355 | 1.40415347937054e-10 |
| GO:0002218 | activation of innate immune response | 273 | -0.465331494 | -2.126428518 | 1e-10 |

| | | | | | |
|---|---|---|---|---|---|
| GO:0071887 | leukocyte apoptotic process | 111 | -0.516972264 | -2.126715564 | 1.12033105125472e-07 |
| GO:1904996 | positive regulation of leukocyte adhesion to vascular endothelia | 23 | -0.70253969 | -2.130703282 | 2.47070123670889e-05 |
| GO:0070233 | negative regulation of T cell apoptotic process | 24 | -0.696377422 | -2.132525326 | 1.55531169826285e-05 |
| GO:0070498 | interleukin-1-mediated signaling pathway | 27 | -0.677262298 | -2.134138024 | 2.4216354209895e-05 |
| GO:0002758 | innate immune response-activating signaling pathway | 254 | -0.470426105 | -2.135158226 | 1e-10 |
| GO:0070672 | response to interleukin-15 | 12 | -0.833801049 | -2.136554337 | 4.09447542191828e-06 |
| GO:0045807 | positive regulation of endocytosis | 145 | -0.501025334 | -2.137045873 | 5.31815970664647e-09 |
| GO:0001774 | microglial cell activation | 45 | -0.608596243 | -2.140759196 | 4.41396989170547e-06 |
| GO:0002224 | toll-like receptor signaling pathway | 71 | -0.559182455 | -2.142136097 | 1.61982172093436e-06 |
| GO:0002279 | mast cell activation involved in immune response | 53 | -0.590552276 | -2.145463027 | 6.70097775330255e-06 |
| GO:0002323 | natural killer cell activation involved in immune response | 29 | -0.670485435 | -2.146171709 | 1.21831508177878e-05 |
| GO:0046425 | regulation of receptor signaling pathway via JAK-STAT | 97 | -0.533004243 | -2.148435171 | 1.39190879299842e-07 |
| GO:0098543 | detection of other organism | 15 | -0.790060473 | -2.149342095 | 6.41630110020465e-06 |
| GO:0034103 | regulation of tissue remodeling | 74 | -0.557644197 | -2.150884004 | 7.66083493964642e-07 |
| GO:0045624 | positive regulation of T-helper cell differentiation | 23 | -0.709659511 | -2.152296688 | 1.58365884599743e-05 |
| GO:0032720 | negative regulation of tumor necrosis factor production | 62 | -0.576437467 | -2.155004647 | 2.22976921258194e-06 |
| GO:1904994 | regulation of leukocyte adhesion to vascular endothelial cell | 30 | -0.668748215 | -2.157492901 | 8.70307014100069e-06 |
| GO:0061900 | glial cell activation | 52 | -0.596230713 | -2.157884433 | 2.15660248849927e-06 |
| GO:0032642 | regulation of chemokine production | 92 | -0.540995963 | -2.161234121 | 2.45861799574943e-07 |
| GO:0009615 | response to virus | 375 | -0.461434964 | -2.163174772 | 1e-10 |
| GO:0048525 | negative regulation of viral process | 88 | -0.54564971 | -2.165436897 | 1.56878551704753e-07 |
| GO:0032611 | interleukin-1 beta production | 90 | -0.544115513 | -2.166538094 | 5.96275326098083e-08 |
| GO:0032651 | regulation of interleukin-1 beta production | 90 | -0.544115513 | -2.166538094 | 5.96275326098083e-08 |
| GO:0035743 | CD4-positive, alpha-beta T cell cytokine production | 19 | -0.749117211 | -2.166635305 | 6.58955315094615e-06 |
| GO:0032607 | interferon-alpha production | 28 | -0.682472854 | -2.168514744 | 1.22091095048704e-05 |
| GO:0032647 | regulation of interferon-alpha production | 28 | -0.682472854 | -2.168514744 | 1.22091095048704e-05 |
| GO:0043373 | CD4-positive, alpha-beta T cell lineage commitment | 23 | -0.715240772 | -2.169223864 | 1.10751744969374e-05 |
| GO:0001911 | negative regulation of leukocyte mediated cytotoxicity | 21 | -0.731673117 | -2.170151558 | 8.84986005776891e-06 |
| GO:0032731 | positive regulation of interleukin-1 beta production | 57 | -0.590490008 | -2.174445825 | 8.66428896204956e-07 |
| GO:0002260 | lymphocyte homeostasis | 74 | -0.564176343 | -2.176079083 | 3.46244863086369e-07 |
| GO:0051770 | positive regulation of nitric-oxide synthase biosynthetic proces | 16 | -0.786870959 | -2.176979929 | 3.85899455051296e-06 |
| GO:0002548 | monocyte chemotaxis | 61 | -0.585168304 | -2.181096864 | 9.31735613881973e-07 |
| GO:0045348 | positive regulation of MHC class II biosynthetic process | 14 | -0.81707776 | -2.183608617 | 5.01371531137392e-06 |
| GO:0070228 | regulation of lymphocyte apoptotic process | 56 | -0.595681136 | -2.18616684 | 1.12350721328724e-06 |
| GO:0002924 | negative regulation of humoral immune response mediated by | 11 | -0.874508224 | -2.186773743 | 1.64268826108959e-06 |
| GO:0002444 | myeloid leukocyte mediated immunity | 106 | -0.535020602 | -2.186885234 | 2.5877907905945e-08 |
| GO:0002755 | MyD88-dependent toll-like receptor signaling pathway | 20 | -0.746694271 | -2.186975622 | 8.23486526043102e-06 |
| GO:0001961 | positive regulation of cytokine-mediated signaling pathway | 57 | -0.594082697 | -2.187675696 | 6.72546325370963e-07 |
| GO:0036005 | response to macrophage colony-stimulating factor | 17 | -0.780365781 | -2.192516362 | 1.02135223702138e-05 |
| GO:0060099 | regulation of phagocytosis, engulfment | 15 | -0.806261906 | -2.193417736 | 1.93176801915103e-06 |
| GO:1905153 | regulation of membrane invagination | 15 | -0.806261906 | -2.193417736 | 1.93176801915103e-06 |
| GO:0050901 | leukocyte tethering or rolling | 33 | -0.665416758 | -2.194108535 | 4.33083267474342e-06 |
| GO:0032743 | positive regulation of interleukin-2 production | 33 | -0.665929857 | -2.195800399 | 4.17334981467263e-06 |
| GO:1902106 | negative regulation of leukocyte differentiation | 104 | -0.539089644 | -2.196596687 | 2.6655170800432e-08 |
| GO:0010324 | membrane invagination | 64 | -0.58433632 | -2.197215957 | 5.67417370643898e-07 |

| | | | | | |
|---|---|---|---|---|---|
| GO:0030890 | positive regulation of B cell proliferation | 44 | -0.627756473 | -2.197661854 | 3.06071665437839e-06 |
| GO:0043299 | leukocyte degranulation | 75 | -0.568321199 | -2.197800515 | 1.40046473825392e-07 |
| GO:0032816 | positive regulation of natural killer cell activation | 21 | -0.741135889 | -2.198218259 | 4.9570395102114e-06 |
| GO:0002223 | stimulatory C-type lectin receptor signaling pathway | 21 | -0.742004906 | -2.200795774 | 4.68011289472711e-06 |
| GO:1990840 | response to lectin | 21 | -0.742004906 | -2.200795774 | 4.68011289472711e-06 |
| GO:1990858 | cellular response to lectin | 21 | -0.742004906 | -2.200795774 | 4.68011289472711e-06 |
| GO:0002710 | negative regulation of T cell mediated immunity | 27 | -0.698589462 | -2.201342581 | 6.42677845141069e-06 |
| GO:0051132 | NK T cell activation | 13 | -0.841090661 | -2.202008533 | 2.46146072429191e-06 |
| GO:0033004 | negative regulation of mast cell activation | 10 | -0.903675888 | -2.202738993 | 7.41184914303727e-07 |
| GO:0033627 | cell adhesion mediated by integrin | 85 | -0.558335172 | -2.204018513 | 8.26419107761623e-08 |
| GO:0032495 | response to muramyl dipeptide | 21 | -0.743740569 | -2.205943772 | 3.9601046929832e-06 |
| GO:0150077 | regulation of neuroinflammatory response | 33 | -0.669593189 | -2.207879668 | 3.07097266680053e-06 |
| GO:0002726 | positive regulation of T cell cytokine production | 25 | -0.713291865 | -2.208294588 | 5.06815871805271e-06 |
| GO:1904892 | regulation of receptor signaling pathway via STAT | 104 | -0.54210925 | -2.208900498 | 1.91850525297667e-08 |
| GO:1903556 | negative regulation of tumor necrosis factor superfamily cytoki | 64 | -0.587536053 | -2.209247563 | 4.36704794420591e-07 |
| GO:0071622 | regulation of granulocyte chemotaxis | 49 | -0.617500884 | -2.209629945 | 7.7100950152434e-07 |
| GO:0036230 | granulocyte activation | 47 | -0.622524129 | -2.209813652 | 1.12215474484754e-06 |
| GO:0032757 | positive regulation of interleukin-8 production | 59 | -0.596423372 | -2.210284136 | 1.03110198197749e-06 |
| GO:0051133 | regulation of NK T cell activation | 11 | -0.884104347 | -2.21076957 | 7.43678873046807e-07 |
| GO:0045059 | positive thymic T cell selection | 15 | -0.813732107 | -2.213740255 | 9.75376594902075e-07 |
| GO:0045060 | negative thymic T cell selection | 12 | -0.864131166 | -2.214273049 | 6.56678095517789e-07 |
| GO:0010820 | positive regulation of T cell chemotaxis | 14 | -0.82889212 | -2.215182036 | 2.09080373885293e-06 |
| GO:0050857 | positive regulation of antigen receptor-mediated signaling pathv | 25 | -0.716794115 | -2.219137274 | 4.14668507408624e-06 |
| GO:0098581 | detection of external biotic stimulus | 22 | -0.739878413 | -2.219219351 | 4.70993863318949e-06 |
| GO:0035872 | nucleotide-binding domain, leucine rich repeat containing rece | 29 | -0.693577639 | -2.220088059 | 2.53631321017236e-06 |
| GO:0002827 | positive regulation of T-helper 1 type immune response | 18 | -0.779038439 | -2.221121125 | 6.29812437354866e-06 |
| GO:0140131 | positive regulation of lymphocyte chemotaxis | 19 | -0.768275057 | -2.222044612 | 1.77065672743589e-06 |
| GO:0032612 | interleukin-1 production | 105 | -0.544773463 | -2.223140657 | 7.25600913675951e-09 |
| GO:0032652 | regulation of interleukin-1 production | 105 | -0.544773463 | -2.223140657 | 7.25600913675951e-09 |
| GO:0043030 | regulation of macrophage activation | 55 | -0.607894114 | -2.224031912 | 4.87498224572774e-07 |
| GO:0045123 | cellular extravasation | 73 | -0.578439651 | -2.2259029 | 1.87231541067062e-07 |
| GO:0033081 | regulation of T cell differentiation in thymus | 27 | -0.707069997 | -2.228065804 | 3.57377876218452e-06 |
| GO:0032637 | interleukin-8 production | 79 | -0.572165863 | -2.231557372 | 5.07458400143747e-08 |
| GO:0002713 | negative regulation of B cell mediated immunity | 15 | -0.821680604 | -2.235363964 | 4.56954165782124e-07 |
| GO:0002890 | negative regulation of immunoglobulin mediated immune resp | 15 | -0.821680604 | -2.235363964 | 4.56954165782124e-07 |
| GO:0070232 | regulation of T cell apoptotic process | 38 | -0.658104517 | -2.235986127 | 1.25687775495828e-06 |
| GO:0050920 | regulation of chemotaxis | 217 | -0.500449038 | -2.23678434 | 1e-10 |
| GO:0002438 | acute inflammatory response to antigenic stimulus | 25 | -0.723323309 | -2.239351137 | 2.79158321063479e-06 |
| GO:0032693 | negative regulation of interleukin-10 production | 20 | -0.764992144 | -2.240567841 | 1.98306404062319e-06 |
| GO:0002639 | positive regulation of immunoglobulin production | 51 | -0.621484241 | -2.240924187 | 2.69210919946864e-07 |
| GO:0072539 | T-helper 17 cell differentiation | 32 | -0.684151843 | -2.241002049 | 1.10126027323187e-06 |
| GO:0061844 | antimicrobial humoral immune response mediated by antimic | 60 | -0.604813615 | -2.24752823 | 1.19462121434718e-07 |
| GO:0032677 | regulation of interleukin-8 production | 78 | -0.57767327 | -2.24827444 | 5.72236224283621e-08 |
| GO:0099024 | plasma membrane invagination | 57 | -0.6106526 | -2.248693421 | 1.84486501692095e-07 |
| GO:0042119 | neutrophil activation | 41 | -0.65139135 | -2.249258376 | 1.28135089073999e-06 |

| | | | | | |
|---|---|---|---|---|---|
| GO:0097530 | granulocyte migration | 147 | -0.52672753 | -2.250717965 | 3.69087133794706e-10 |
| GO:0071621 | granulocyte chemotaxis | 121 | -0.540513269 | -2.250879509 | 1.00720846679401e-09 |
| GO:0031348 | negative regulation of defense response | 238 | -0.499212963 | -2.251286951 | 1e-10 |
| GO:0071346 | cellular response to type II interferon | 108 | -0.549861501 | -2.253904811 | 7.13895149171503e-09 |
| GO:0002714 | positive regulation of B cell mediated immunity | 41 | -0.652822872 | -2.254201429 | 1.2182306809887e-06 |
| GO:0002891 | positive regulation of immunoglobulin mediated immune respo | 41 | -0.652822872 | -2.254201429 | 1.2182306809887e-06 |
| GO:0031342 | negative regulation of cell killing | 24 | -0.737010166 | -2.25695549 | 1.45370675462049e-06 |
| GO:0002673 | regulation of acute inflammatory response | 43 | -0.649445766 | -2.263870448 | 3.3423619151416e-07 |
| GO:0001776 | leukocyte homeostasis | 104 | -0.555670384 | -2.264157248 | 3.40848379223227e-09 |
| GO:0061756 | leukocyte adhesion to vascular endothelial cell | 47 | -0.638019436 | -2.264818332 | 3.36229047537429e-07 |
| GO:0150076 | neuroinflammatory response | 75 | -0.585735808 | -2.265145944 | 2.15123238157024e-08 |
| GO:0042130 | negative regulation of T cell proliferation | 67 | -0.597430449 | -2.26564446 | 4.4717993430476e-08 |
| GO:0030449 | regulation of complement activation | 18 | -0.794714907 | -2.265816395 | 1.95237067158613e-06 |
| GO:0038061 | non-canonical NF-kappaB signal transduction | 119 | -0.545616146 | -2.267119986 | 8.57255101523004e-10 |
| GO:0042330 | taxis | 437 | -0.478846358 | -2.267620956 | 1e-10 |
| GO:0002544 | chronic inflammatory response | 19 | -0.784390312 | -2.268653982 | 4.52129691045446e-07 |
| GO:0002922 | positive regulation of humoral immune response | 21 | -0.7658251 | -2.271446765 | 1.00774145404542e-06 |
| GO:0050858 | negative regulation of antigen receptor-mediated signaling path | 31 | -0.699033843 | -2.273176691 | 7.39656871705244e-07 |
| GO:0071677 | positive regulation of mononuclear cell migration | 69 | -0.596585739 | -2.274104502 | 3.11405614600479e-08 |
| GO:0006935 | chemotaxis | 435 | -0.480413834 | -2.2742908 | 1e-10 |
| GO:2000406 | positive regulation of T cell migration | 35 | -0.681412774 | -2.274817705 | 6.39520259110884e-07 |
| GO:0060760 | positive regulation of response to cytokine stimulus | 62 | -0.609518207 | -2.278676601 | 8.34627867684844e-08 |
| GO:0038093 | Fc receptor signaling pathway | 53 | -0.627326494 | -2.279062925 | 2.98141134822769e-07 |
| GO:0045071 | negative regulation of viral genome replication | 54 | -0.625270829 | -2.27992624 | 1.67582989011637e-07 |
| GO:1990266 | neutrophil migration | 123 | -0.546417312 | -2.280128735 | 4.54946387348902e-10 |
| GO:0033003 | regulation of mast cell activation | 19 | -0.78869193 | -2.281095343 | 3.2798466628406e-07 |
| GO:0034113 | heterotypic cell-cell adhesion | 60 | -0.614016368 | -2.281726281 | 5.05774177452492e-08 |
| GO:0043383 | negative T cell selection | 13 | -0.872978087 | -2.285491073 | 1.54409810995286e-07 |
| GO:0032633 | interleukin-4 production | 30 | -0.709047751 | -2.287505906 | 6.24942707450866e-07 |
| GO:0032673 | regulation of interleukin-4 production | 30 | -0.709047751 | -2.287505906 | 6.24942707450866e-07 |
| GO:0002526 | acute inflammatory response | 99 | -0.565754399 | -2.287589195 | 1.81309275549168e-09 |
| GO:0002921 | negative regulation of humoral immune response | 16 | -0.827406279 | -2.289126118 | 1.31604251620679e-07 |
| GO:2000107 | negative regulation of leukocyte apoptotic process | 55 | -0.626148316 | -2.290816449 | 8.91985493936242e-08 |
| GO:0034341 | response to type II interferon | 130 | -0.545000564 | -2.291963033 | 1e-10 |
| GO:0002369 | T cell cytokine production | 36 | -0.683003731 | -2.293457144 | 6.43555327073092e-07 |
| GO:0002724 | regulation of T cell cytokine production | 36 | -0.683003731 | -2.293457144 | 6.43555327073092e-07 |
| GO:0019722 | calcium-mediated signaling | 227 | -0.510921468 | -2.293876418 | 1e-10 |
| GO:0032722 | positive regulation of chemokine production | 70 | -0.600198384 | -2.294362599 | 2.99819716129171e-08 |
| GO:0010819 | regulation of T cell chemotaxis | 15 | -0.844366918 | -2.297081582 | 8.07131780960049e-08 |
| GO:0002220 | innate immune response activating cell surface receptor signa | 89 | -0.578082918 | -2.298428398 | 1.37425463779759e-08 |
| GO:0002833 | positive regulation of response to biotic stimulus | 355 | -0.492398466 | -2.298611273 | 1e-10 |
| GO:0002886 | regulation of myeloid leukocyte mediated immunity | 54 | -0.631081991 | -2.301115493 | 9.69004543483613e-08 |
| GO:0002507 | tolerance induction | 25 | -0.743330176 | -2.301290798 | 8.1181709761526e-07 |
| GO:1901222 | regulation of non-canonical NF-kappaB signal transduction | 90 | -0.578579045 | -2.303763651 | 1.39276273759393e-09 |
| GO:2000404 | regulation of T cell migration | 48 | -0.647030401 | -2.30517674 | 1.69937140854022e-07 |

| | | | | | |
|---|---|---|---|---|---|
| GO:0019730 | antimicrobial humoral response | 103 | -0.566759779 | -2.305660931 | 9.68507997389139e-10 |
| GO:0002711 | positive regulation of T cell mediated immunity | 57 | -0.62723998 | -2.309775506 | 4.12140128718296e-08 |
| GO:0050728 | negative regulation of inflammatory response | 156 | -0.537421526 | -2.312088766 | 1e-10 |
| GO:0007204 | positive regulation of cytosolic calcium ion concentration | 159 | -0.536157863 | -2.312149685 | 1e-10 |
| GO:0002720 | positive regulation of cytokine production involved in immune r | 76 | -0.597672385 | -2.316258879 | 6.98951560163147e-09 |
| GO:0048247 | lymphocyte chemotaxis | 55 | -0.633267465 | -2.316862456 | 3.65540212069295e-08 |
| GO:0022408 | negative regulation of cell-cell adhesion | 182 | -0.52877572 | -2.317522282 | 1e-10 |
| GO:0002831 | regulation of response to biotic stimulus | 482 | -0.486943443 | -2.31929317 | 1e-10 |
| GO:0071222 | cellular response to lipopolysaccharide | 204 | -0.522272387 | -2.319455896 | 1e-10 |
| GO:2000403 | positive regulation of lymphocyte migration | 42 | -0.668945552 | -2.321533584 | 1.85372967270236e-07 |
| GO:0050856 | regulation of T cell receptor signaling pathway | 44 | -0.663166438 | -2.321625738 | 1.28796853108629e-07 |
| GO:0050921 | positive regulation of chemotaxis | 140 | -0.547632453 | -2.324761613 | 1e-10 |
| GO:0045579 | positive regulation of B cell differentiation | 16 | -0.841461116 | -2.328010635 | 3.43045032311276e-08 |
| GO:0032715 | negative regulation of interleukin-6 production | 47 | -0.655950657 | -2.328469934 | 6.42072253933701e-08 |
| GO:0071216 | cellular response to biotic stimulus | 240 | -0.517171865 | -2.334092186 | 1e-10 |
| GO:0010818 | T cell chemotaxis | 24 | -0.76262702 | -2.335402301 | 2.29246733454485e-07 |
| GO:0030225 | macrophage differentiation | 59 | -0.630589817 | -2.336901496 | 3.9538103473735e-08 |
| GO:0036037 | CD8-positive, alpha-beta T cell activation | 25 | -0.75505634 | -2.337594064 | 2.80684524030428e-07 |
| GO:0030593 | neutrophil chemotaxis | 101 | -0.576581845 | -2.338990076 | 8.98504663568241e-10 |
| GO:0002275 | myeloid cell activation involved in immune response | 93 | -0.584384124 | -2.339104797 | 7.00523787637363e-10 |
| GO:0032755 | positive regulation of interleukin-6 production | 95 | -0.58265034 | -2.340375429 | 5.2994209670654e-10 |
| GO:0032732 | positive regulation of interleukin-1 production | 66 | -0.619909214 | -2.345172192 | 1.75656347555014e-08 |
| GO:2000106 | regulation of leukocyte apoptotic process | 84 | -0.596146843 | -2.348196368 | 2.39564592108906e-09 |
| GO:0045066 | regulatory T cell differentiation | 36 | -0.699379069 | -2.348443867 | 1.4980048524133e-07 |
| GO:2000401 | regulation of lymphocyte migration | 67 | -0.61928103 | -2.348508748 | 5.50775142474161e-09 |
| GO:0002313 | mature B cell differentiation involved in immune response | 31 | -0.722214166 | -2.348556402 | 1.16898692347285e-07 |
| GO:0032496 | response to lipopolysaccharide | 321 | -0.507291877 | -2.350111052 | 1e-10 |
| GO:0045576 | mast cell activation | 66 | -0.621278846 | -2.350353632 | 1.5403730758785e-08 |
| GO:1901623 | regulation of lymphocyte chemotaxis | 24 | -0.767591632 | -2.350605496 | 1.59017440351469e-07 |
| GO:0002701 | negative regulation of production of molecular mediator of imm | 44 | -0.671456294 | -2.350647021 | 7.02215014596167e-08 |
| GO:0045429 | positive regulation of nitric oxide biosynthetic process | 36 | -0.700280357 | -2.351470298 | 1.41874609665741e-07 |
| GO:0045089 | positive regulation of innate immune response | 334 | -0.506262963 | -2.352532055 | 1e-10 |
| GO:0002377 | immunoglobulin production | 102 | -0.579774153 | -2.355760595 | 2.01456684388872e-10 |
| GO:0032753 | positive regulation of interleukin-4 production | 25 | -0.765466441 | -2.369822903 | 1.47916013465669e-07 |
| GO:0032814 | regulation of natural killer cell activation | 38 | -0.697858517 | -2.371054933 | 4.33986779728047e-08 |
| GO:0002367 | cytokine production involved in immune response | 116 | -0.572710394 | -2.371284809 | 1e-10 |
| GO:0002690 | positive regulation of leukocyte chemotaxis | 91 | -0.594609194 | -2.371937939 | 5.17627877406471e-10 |
| GO:0071675 | regulation of mononuclear cell migration | 117 | -0.572655862 | -2.373624033 | 1e-10 |
| GO:0002828 | regulation of type 2 immune response | 34 | -0.716201996 | -2.376265126 | 4.24958771990719e-08 |
| GO:0045785 | positive regulation of cell adhesion | 444 | -0.501424431 | -2.377019229 | 1e-10 |
| GO:0045088 | regulation of innate immune response | 404 | -0.504713017 | -2.378321545 | 1e-10 |
| GO:0071219 | cellular response to molecule of bacterial origin | 213 | -0.534377854 | -2.38330713 | 1e-10 |
| GO:0002675 | positive regulation of acute inflammatory response | 24 | -0.778416273 | -2.383753931 | 5.8406722311895e-08 |
| GO:1904407 | positive regulation of nitric oxide metabolic process | 37 | -0.705918227 | -2.383965009 | 3.54041656474828e-08 |
| GO:0042092 | type 2 immune response | 42 | -0.687281732 | -2.385168149 | 3.58781490136361e-08 |

| | | | | | |
|---|---|---|---|---|---|
| GO:0002407 | dendritic cell chemotaxis | 23 | -0.786625855 | -2.385724698 | 3.91076712307104e-08 |
| GO:0098586 | cellular response to virus | 72 | -0.621804196 | -2.386862499 | 8.46874444147984e-10 |
| GO:0002830 | positive regulation of type 2 immune response | 18 | -0.838447604 | -2.390502948 | 4.49669017498518e-08 |
| GO:0032760 | positive regulation of tumor necrosis factor production | 101 | -0.589370531 | -2.390869283 | 1.9413922200683e-10 |
| GO:0046629 | gamma-delta T cell activation | 26 | -0.766738201 | -2.394551799 | 8.31317671204446e-08 |
| GO:0045622 | regulation of T-helper cell differentiation | 40 | -0.697633677 | -2.396512192 | 1.16257702696277e-08 |
| GO:0050855 | regulation of B cell receptor signaling pathway | 21 | -0.80807249 | -2.396753049 | 2.361924027469e-08 |
| GO:0050868 | negative regulation of T cell activation | 118 | -0.577720619 | -2.397435647 | 1e-10 |
| GO:0032740 | positive regulation of interleukin-17 production | 26 | -0.767889999 | -2.398148907 | 7.49415151045437e-08 |
| GO:0002702 | positive regulation of production of molecular mediator of imm | 128 | -0.571512529 | -2.399055083 | 1e-10 |
| GO:0050871 | positive regulation of B cell activation | 83 | -0.610530319 | -2.400111345 | 1.68235272878194e-10 |
| GO:0050906 | detection of stimulus involved in sensory perception | 446 | -0.506314432 | -2.400945267 | 1e-10 |
| GO:1903038 | negative regulation of leukocyte cell-cell adhesion | 129 | -0.571607696 | -2.401772944 | 1e-10 |
| GO:1903706 | regulation of hemopoiesis | 390 | -0.51172402 | -2.405476852 | 1e-10 |
| GO:0002688 | regulation of leukocyte chemotaxis | 118 | -0.580377798 | -2.408462455 | 1e-10 |
| GO:0001818 | negative regulation of cytokine production | 275 | -0.526859091 | -2.4091074 | 1e-10 |
| GO:0030595 | leukocyte chemotaxis | 220 | -0.538693579 | -2.411163062 | 1e-10 |
| GO:0032623 | interleukin-2 production | 61 | -0.64696414 | -2.411428385 | 1.89811838154751e-09 |
| GO:0032663 | regulation of interleukin-2 production | 61 | -0.64696414 | -2.411428385 | 1.89811838154751e-09 |
| GO:2000316 | regulation of T-helper 17 type immune response | 31 | -0.74284329 | -2.415639913 | 1.89655561811855e-08 |
| GO:0002237 | response to molecule of bacterial origin | 338 | -0.519656031 | -2.416854804 | 1e-10 |
| GO:0050731 | positive regulation of peptidyl-tyrosine phosphorylation | 155 | -0.562078529 | -2.416883387 | 1e-10 |
| GO:0042116 | macrophage activation | 103 | -0.59502573 | -2.420650918 | 1e-10 |
| GO:0002440 | production of molecular mediator of immune response | 213 | -0.542820357 | -2.420960408 | 1e-10 |
| GO:0046638 | positive regulation of alpha-beta T cell differentiation | 51 | -0.671700359 | -2.421991551 | 2.07784238762506e-09 |
| GO:1903557 | positive regulation of tumor necrosis factor superfamily cytoki | 105 | -0.593647607 | -2.422588872 | 1e-10 |
| GO:0097028 | dendritic cell differentiation | 41 | -0.70364846 | -2.4297025 | 1.4841884207826e-08 |
| GO:0060326 | cell chemotaxis | 294 | -0.529316192 | -2.433957355 | 1e-10 |
| GO:0002718 | regulation of cytokine production involved in immune response | 114 | -0.590426099 | -2.438965784 | 1e-10 |
| GO:1901224 | positive regulation of non-canonical NF-kappaB signal transdu | 62 | -0.652423836 | -2.439078788 | 7.41445484012004e-10 |
| GO:0002717 | positive regulation of natural killer cell mediated immunity | 29 | -0.762073259 | -2.439337209 | 4.24269868215245e-09 |
| GO:0050730 | regulation of peptidyl-tyrosine phosphorylation | 213 | -0.547040387 | -2.439781599 | 1e-10 |
| GO:0042531 | positive regulation of tyrosine phosphorylation of STAT protein | 59 | -0.658871237 | -2.441709552 | 2.12932982628958e-09 |
| GO:0042742 | defense response to bacterium | 280 | -0.53363021 | -2.443534055 | 1e-10 |
| GO:0006911 | phagocytosis, engulfment | 48 | -0.686154585 | -2.444564564 | 3.78559221592361e-09 |
| GO:0002861 | regulation of inflammatory response to antigenic stimulus | 38 | -0.720188051 | -2.446922105 | 3.95327428928211e-09 |
| GO:0032733 | positive regulation of interleukin-10 production | 40 | -0.713243369 | -2.450134628 | 2.68767581821296e-09 |
| GO:0050766 | positive regulation of phagocytosis | 66 | -0.648797562 | -2.454459402 | 5.79623380534592e-10 |
| GO:0036336 | dendritic cell migration | 29 | -0.766874728 | -2.454706337 | 2.4565702164981e-09 |
| GO:0006968 | cellular defense response | 47 | -0.693614343 | -2.462167123 | 1.21630894574365e-09 |
| GO:0031294 | lymphocyte costimulation | 48 | -0.691128471 | -2.462285039 | 2.11459610009078e-09 |
| GO:0032689 | negative regulation of type II interferon production | 39 | -0.720986082 | -2.463307012 | 5.41898742899214e-09 |
| GO:0032640 | tumor necrosis factor production | 163 | -0.56961463 | -2.464713318 | 1e-10 |
| GO:0032680 | regulation of tumor necrosis factor production | 163 | -0.56961463 | -2.464713318 | 1e-10 |
| GO:0045582 | positive regulation of T cell differentiation | 115 | -0.598301304 | -2.474552453 | 1e-10 |

| | | | | | |
|---|---|---|---|---|---|
| GO:0045954 | positive regulation of natural killer cell mediated cytotoxicity | 25 | -0.799786326 | -2.476074523 | 4.96342389927104e-09 |
| GO:0072538 | T-helper 17 type immune response | 45 | -0.704158665 | -2.476903456 | 1.08542726239325e-09 |
| GO:0002923 | regulation of humoral immune response mediated by circulatir | 18 | -0.871427745 | -2.484532823 | 8.64046034923933e-10 |
| GO:0031349 | positive regulation of defense response | 456 | -0.523288679 | -2.48472532 | 1e-10 |
| GO:0032735 | positive regulation of interleukin-12 production | 40 | -0.725001094 | -2.490524784 | 6.65331725893117e-10 |
| GO:0043372 | positive regulation of CD4-positive, alpha-beta T cell differentia | 33 | -0.755340881 | -2.490619381 | 1.66320830754113e-09 |
| GO:0071706 | tumor necrosis factor superfamily cytokine production | 168 | -0.573697743 | -2.491594977 | 1e-10 |
| GO:1903555 | regulation of tumor necrosis factor superfamily cytokine produ | 168 | -0.573697743 | -2.491594977 | 1e-10 |
| GO:0050900 | leukocyte migration | 371 | -0.532151195 | -2.49289558 | 1e-10 |
| GO:0006909 | phagocytosis | 225 | -0.556619294 | -2.496516437 | 1e-10 |
| GO:0031295 | T cell costimulation | 47 | -0.703539322 | -2.497398455 | 4.04326437619017e-10 |
| GO:0002683 | negative regulation of immune system process | 472 | -0.525135012 | -2.498378302 | 1e-10 |
| GO:0002437 | inflammatory response to antigenic stimulus | 70 | -0.653596577 | -2.498486469 | 1e-10 |
| GO:0050852 | T cell receptor signaling pathway | 134 | -0.59360249 | -2.506545745 | 1e-10 |
| GO:0032635 | interleukin-6 production | 146 | -0.587300859 | -2.507083547 | 1e-10 |
| GO:0032675 | regulation of interleukin-6 production | 146 | -0.587300859 | -2.507083547 | 1e-10 |
| GO:0045061 | thymic T cell selection | 24 | -0.819254205 | -2.508812443 | 9.61432174759448e-10 |
| GO:0033077 | T cell differentiation in thymus | 85 | -0.635927501 | -2.510312902 | 1e-10 |
| GO:0002687 | positive regulation of leukocyte migration | 147 | -0.588027253 | -2.512652987 | 1e-10 |
| GO:0007259 | receptor signaling pathway via JAK-STAT | 159 | -0.58306743 | -2.514444471 | 1e-10 |
| GO:1902107 | positive regulation of leukocyte differentiation | 182 | -0.573721937 | -2.514512906 | 1e-10 |
| GO:1903708 | positive regulation of hemopoiesis | 182 | -0.573721937 | -2.514512906 | 1e-10 |
| GO:0072676 | lymphocyte migration | 115 | -0.608350702 | -2.516116395 | 1e-10 |
| GO:0072678 | T cell migration | 70 | -0.658433705 | -2.516977229 | 1e-10 |
| GO:0002637 | regulation of immunoglobulin production | 71 | -0.657040355 | -2.517013627 | 1e-10 |
| GO:0070664 | negative regulation of leukocyte proliferation | 90 | -0.632446789 | -2.518252151 | 1e-10 |
| GO:0046637 | regulation of alpha-beta T cell differentiation | 70 | -0.660025011 | -2.523060274 | 1e-10 |
| GO:0002335 | mature B cell differentiation | 37 | -0.747883266 | -2.525685651 | 4.12454620950064e-10 |
| GO:0002685 | regulation of leukocyte migration | 220 | -0.565140897 | -2.529539814 | 1e-10 |
| GO:0002312 | B cell activation involved in immune response | 85 | -0.641526363 | -2.532414315 | 1e-10 |
| GO:0007606 | sensory perception of chemical stimulus | 429 | -0.535407879 | -2.532688822 | 1e-10 |
| GO:0032945 | negative regulation of mononuclear cell proliferation | 82 | -0.646332426 | -2.535862063 | 1e-10 |
| GO:0050764 | regulation of phagocytosis | 92 | -0.635037103 | -2.536920699 | 1e-10 |
| GO:0043368 | positive T cell selection | 42 | -0.731166695 | -2.53746816 | 3.69639699703491e-10 |
| GO:0030888 | regulation of B cell proliferation | 63 | -0.677275674 | -2.539566418 | 1e-10 |
| GO:0097696 | receptor signaling pathway via STAT | 167 | -0.586184337 | -2.544425753 | 1e-10 |
| GO:0050672 | negative regulation of lymphocyte proliferation | 81 | -0.649674157 | -2.544915606 | 1e-10 |
| GO:0042509 | regulation of tyrosine phosphorylation of STAT protein | 70 | -0.668228231 | -2.554418505 | 1e-10 |
| GO:0050869 | negative regulation of B cell activation | 32 | -0.779941914 | -2.554771205 | 3.58666014265214e-10 |
| GO:0046641 | positive regulation of alpha-beta T cell proliferation | 22 | -0.852042954 | -2.555649926 | 1.74170644926082e-10 |
| GO:0002821 | positive regulation of adaptive immune response | 117 | -0.616965335 | -2.557284125 | 1e-10 |
| GO:0018212 | peptidyl-tyrosine modification | 258 | -0.562791759 | -2.557637668 | 1e-10 |
| GO:0018108 | peptidyl-tyrosine phosphorylation | 256 | -0.563298212 | -2.558570009 | 1e-10 |
| GO:0002700 | regulation of production of molecular mediator of immune resp | 178 | -0.585272073 | -2.559192533 | 1e-10 |
| GO:0001912 | positive regulation of leukocyte mediated cytotoxicity | 56 | -0.697556241 | -2.560051394 | 1e-10 |

| | | | | | |
|---|---|---|---|---|---|
| GO:0022407 | regulation of cell-cell adhesion | 449 | -0.540384342 | -2.563382444 | 1e-10 |
| GO:2000516 | positive regulation of CD4-positive, alpha-beta T cell activatior | 42 | -0.739767512 | -2.567316755 | 1.11677699790435e-10 |
| GO:0042093 | T-helper cell differentiation | 69 | -0.674724222 | -2.571957877 | 1e-10 |
| GO:0002709 | regulation of T cell mediated immunity | 86 | -0.650715706 | -2.573686904 | 1e-10 |
| GO:0007608 | sensory perception of smell | 368 | -0.551352834 | -2.581083109 | 1e-10 |
| GO:0032615 | interleukin-12 production | 59 | -0.696493219 | -2.581132775 | 1e-10 |
| GO:0032655 | regulation of interleukin-12 production | 59 | -0.696493219 | -2.581132775 | 1e-10 |
| GO:0007260 | tyrosine phosphorylation of STAT protein | 73 | -0.672374371 | -2.5873746 | 1e-10 |
| GO:0046640 | regulation of alpha-beta T cell proliferation | 40 | -0.753356782 | -2.587932282 | 1e-10 |
| GO:0032620 | interleukin-17 production | 39 | -0.757650272 | -2.58857317 | 1.01119898624887e-10 |
| GO:0032660 | regulation of interleukin-17 production | 39 | -0.757650272 | -2.58857317 | 1.01119898624887e-10 |
| GO:0042269 | regulation of natural killer cell mediated cytotoxicity | 42 | -0.746036712 | -2.589073622 | 1e-10 |
| GO:0045621 | positive regulation of lymphocyte differentiation | 129 | -0.616227847 | -2.589257247 | 1e-10 |
| GO:0009593 | detection of chemical stimulus | 414 | -0.548902966 | -2.590959177 | 1e-10 |
| GO:0050727 | regulation of inflammatory response | 359 | -0.555451629 | -2.595248927 | 1e-10 |
| GO:1902105 | regulation of leukocyte differentiation | 311 | -0.563047746 | -2.601248314 | 1e-10 |
| GO:0071674 | mononuclear cell migration | 188 | -0.591845108 | -2.603582895 | 1e-10 |
| GO:0045580 | regulation of T cell differentiation | 175 | -0.597067441 | -2.605342427 | 1e-10 |
| GO:0001819 | positive regulation of cytokine production | 459 | -0.548898592 | -2.606973613 | 1e-10 |
| GO:0002825 | regulation of T-helper 1 type immune response | 31 | -0.801695796 | -2.607021412 | 1e-10 |
| GO:0006956 | complement activation | 58 | -0.705977245 | -2.608553216 | 1e-10 |
| GO:0046633 | alpha-beta T cell proliferation | 43 | -0.749539701 | -2.61278288 | 1e-10 |
| GO:0002294 | CD4-positive, alpha-beta T cell differentiation involved in imm | 71 | -0.682289469 | -2.613738837 | 1e-10 |
| GO:0050866 | negative regulation of cell activation | 199 | -0.591087586 | -2.617171269 | 1e-10 |
| GO:0002824 | positive regulation of adaptive immune response based on son | 112 | -0.636154791 | -2.620539177 | 1e-10 |
| GO:0050907 | detection of chemical stimulus involved in sensory perception | 380 | -0.55838469 | -2.620552067 | 1e-10 |
| GO:0002712 | regulation of B cell mediated immunity | 60 | -0.705268493 | -2.620825341 | 1e-10 |
| GO:0002889 | regulation of immunoglobulin mediated immune response | 60 | -0.705268493 | -2.620825341 | 1e-10 |
| GO:0050777 | negative regulation of immune response | 177 | -0.599797591 | -2.621059536 | 1e-10 |
| GO:0002274 | myeloid leukocyte activation | 230 | -0.583360493 | -2.622099838 | 1e-10 |
| GO:0032613 | interleukin-10 production | 58 | -0.710343773 | -2.624687333 | 1e-10 |
| GO:0032653 | regulation of interleukin-10 production | 58 | -0.710343773 | -2.624687333 | 1e-10 |
| GO:0042267 | natural killer cell mediated cytotoxicity | 68 | -0.690341665 | -2.626170359 | 1e-10 |
| GO:0043370 | regulation of CD4-positive, alpha-beta T cell differentiation | 52 | -0.727201286 | -2.631894501 | 1e-10 |
| GO:0006958 | complement activation, classical pathway | 39 | -0.770739039 | -2.633291995 | 1e-10 |
| GO:0002287 | alpha-beta T cell activation involved in immune response | 72 | -0.686328162 | -2.634544706 | 1e-10 |
| GO:0002293 | alpha-beta T cell differentiation involved in immune response | 72 | -0.686328162 | -2.634544706 | 1e-10 |
| GO:0030101 | natural killer cell activation | 89 | -0.663951594 | -2.639837907 | 1e-10 |
| GO:0050911 | detection of chemical stimulus involved in sensory perception | 343 | -0.567350896 | -2.64148443 | 1e-10 |
| GO:0002715 | regulation of natural killer cell mediated immunity | 46 | -0.749192394 | -2.64719932 | 1e-10 |
| GO:0002292 | T cell differentiation involved in immune response | 80 | -0.677452114 | -2.648004935 | 1e-10 |
| GO:0002286 | T cell activation involved in immune response | 115 | -0.640933816 | -2.650878972 | 1e-10 |
| GO:0042088 | T-helper 1 type immune response | 46 | -0.750326457 | -2.651206422 | 1e-10 |
| GO:0032946 | positive regulation of mononuclear cell proliferation | 133 | -0.629702163 | -2.656311924 | 1e-10 |
| GO:0050671 | positive regulation of lymphocyte proliferation | 130 | -0.631719284 | -2.65665275 | 1e-10 |

| | | | | | |
|---|---|---|---|---|---|
| GO:0042098 | T cell proliferation | 195 | -0.601498224 | -2.656986512 | 1e-10 |
| GO:0070665 | positive regulation of leukocyte proliferation | 150 | -0.620927735 | -2.659657248 | 1e-10 |
| GO:0002707 | negative regulation of lymphocyte mediated immunity | 52 | -0.735622539 | -2.662372787 | 1e-10 |
| GO:0002456 | T cell mediated immunity | 113 | -0.645556709 | -2.662745375 | 1e-10 |
| GO:0002920 | regulation of humoral immune response | 42 | -0.76786389 | -2.664823473 | 1e-10 |
| GO:0031343 | positive regulation of cell killing | 63 | -0.710817215 | -2.665336432 | 1e-10 |
| GO:0001906 | cell killing | 195 | -0.60420138 | -2.668927112 | 1e-10 |
| GO:0042102 | positive regulation of T cell proliferation | 91 | -0.669738377 | -2.671633539 | 1e-10 |
| GO:0051250 | negative regulation of lymphocyte activation | 152 | -0.622985301 | -2.672776657 | 1e-10 |
| GO:2000514 | regulation of CD4-positive, alpha-beta T cell activation | 72 | -0.697272459 | -2.676555572 | 1e-10 |
| GO:0045577 | regulation of B cell differentiation | 34 | -0.807039473 | -2.677652065 | 1e-10 |
| GO:0022409 | positive regulation of cell-cell adhesion | 293 | -0.582994207 | -2.680181712 | 1e-10 |
| GO:0019221 | cytokine-mediated signaling pathway | 476 | -0.564220702 | -2.685683728 | 1e-10 |
| GO:0002228 | natural killer cell mediated immunity | 71 | -0.702550382 | -2.691355064 | 1e-10 |
| GO:0051251 | positive regulation of lymphocyte activation | 299 | -0.58501064 | -2.694476329 | 1e-10 |
| GO:0001910 | regulation of leukocyte mediated cytotoxicity | 83 | -0.685792352 | -2.695980778 | 1e-10 |
| GO:0002704 | negative regulation of leukocyte mediated immunity | 63 | -0.719179245 | -2.696691359 | 1e-10 |
| GO:0001909 | leukocyte mediated cytotoxicity | 126 | -0.64528088 | -2.702432439 | 1e-10 |
| GO:1990868 | response to chemokine | 94 | -0.673975843 | -2.702589939 | 1e-10 |
| GO:1990869 | cellular response to chemokine | 94 | -0.673975843 | -2.702589939 | 1e-10 |
| GO:0050729 | positive regulation of inflammatory response | 135 | -0.639341199 | -2.702825686 | 1e-10 |
| GO:0042129 | regulation of T cell proliferation | 163 | -0.624817315 | -2.703574448 | 1e-10 |
| GO:0002823 | negative regulation of adaptive immune response based on sor | 54 | -0.742011404 | -2.705597625 | 1e-10 |
| GO:1903037 | regulation of leukocyte cell-cell adhesion | 339 | -0.582096795 | -2.707854968 | 1e-10 |
| GO:0046635 | positive regulation of alpha-beta T cell activation | 67 | -0.714791714 | -2.710715349 | 1e-10 |
| GO:0002708 | positive regulation of lymphocyte mediated immunity | 114 | -0.656326305 | -2.711190112 | 1e-10 |
| GO:0002820 | negative regulation of adaptive immune response | 59 | -0.731878644 | -2.712267549 | 1e-10 |
| GO:0007159 | leukocyte cell-cell adhesion | 375 | -0.578703645 | -2.712922127 | 1e-10 |
| GO:0050854 | regulation of antigen receptor-mediated signaling pathway | 63 | -0.725001492 | -2.718522918 | 1e-10 |
| GO:0002695 | negative regulation of leukocyte activation | 177 | -0.622625103 | -2.720813634 | 1e-10 |
| GO:0002764 | immune response-regulating signaling pathway | 482 | -0.57281632 | -2.728302433 | 1e-10 |
| GO:0070098 | chemokine-mediated signaling pathway | 86 | -0.69115374 | -2.733625933 | 1e-10 |
| GO:0002757 | immune response-activating signaling pathway | 457 | -0.575823136 | -2.734543319 | 1e-10 |
| GO:0050863 | regulation of T cell activation | 347 | -0.587285417 | -2.737095622 | 1e-10 |
| GO:0046632 | alpha-beta T cell differentiation | 119 | -0.658882466 | -2.737759172 | 1e-10 |
| GO:0002698 | negative regulation of immune effector process | 113 | -0.665408286 | -2.744627713 | 1e-10 |
| GO:0045619 | regulation of lymphocyte differentiation | 206 | -0.617815862 | -2.746317423 | 1e-10 |
| GO:0050870 | positive regulation of T cell activation | 225 | -0.612458964 | -2.746965268 | 1e-10 |
| GO:0042100 | B cell proliferation | 100 | -0.678596016 | -2.748119624 | 1e-10 |
| GO:0045058 | T cell selection | 55 | -0.751392155 | -2.749031601 | 1e-10 |
| GO:0002696 | positive regulation of leukocyte activation | 329 | -0.592530527 | -2.750476489 | 1e-10 |
| GO:0050867 | positive regulation of cell activation | 344 | -0.591134068 | -2.752985315 | 1e-10 |
| GO:0002366 | leukocyte activation involved in immune response | 284 | -0.60063923 | -2.753947649 | 1e-10 |
| GO:0002263 | cell activation involved in immune response | 288 | -0.600121332 | -2.755215074 | 1e-10 |
| GO:0002705 | positive regulation of leukocyte mediated immunity | 135 | -0.652325161 | -2.757715604 | 1e-10 |

| | | | | | |
|---|---|---|---|---|---|
| GO:1903039 | positive regulation of leukocyte cell-cell adhesion | 247 | -0.609523592 | -2.758872137 | 1e-10 |
| GO:0043367 | CD4-positive, alpha-beta T cell differentiation | 89 | -0.694616485 | -2.761759959 | 1e-10 |
| GO:0002699 | positive regulation of immune effector process | 235 | -0.614978384 | -2.770115695 | 1e-10 |
| GO:0006959 | humoral immune response | 217 | -0.620150974 | -2.77179869 | 1e-10 |
| GO:0030217 | T cell differentiation | 297 | -0.603714767 | -2.779132188 | 1e-10 |
| GO:0035710 | CD4-positive, alpha-beta T cell activation | 113 | -0.674322396 | -2.781395985 | 1e-10 |
| GO:0032729 | positive regulation of type II interferon production | 71 | -0.727816392 | -2.788144996 | 1e-10 |
| GO:0051249 | regulation of lymphocyte activation | 465 | -0.586667616 | -2.78879951 | 1e-10 |
| GO:0002285 | lymphocyte activation involved in immune response | 199 | -0.630186897 | -2.790292133 | 1e-10 |
| GO:0070663 | regulation of leukocyte proliferation | 246 | -0.616876541 | -2.791289423 | 1e-10 |
| GO:0002455 | humoral immune response mediated by circulating immunogl | 53 | -0.770692515 | -2.79990843 | 1e-10 |
| GO:0046634 | regulation of alpha-beta T cell activation | 106 | -0.687158243 | -2.808744579 | 1e-10 |
| GO:0030183 | B cell differentiation | 151 | -0.655946354 | -2.811873359 | 1e-10 |
| GO:0031341 | regulation of cell killing | 96 | -0.699573316 | -2.814989374 | 1e-10 |
| GO:0070661 | leukocyte proliferation | 327 | -0.609051503 | -2.826095485 | 1e-10 |
| GO:0032944 | regulation of mononuclear cell proliferation | 221 | -0.631195365 | -2.826648807 | 1e-10 |
| GO:0050670 | regulation of lymphocyte proliferation | 217 | -0.63387459 | -2.833137143 | 1e-10 |
| GO:0050864 | regulation of B cell activation | 127 | -0.676220107 | -2.835234205 | 1e-10 |
| GO:0032943 | mononuclear cell proliferation | 292 | -0.618862314 | -2.844141783 | 1e-10 |
| GO:0046651 | lymphocyte proliferation | 285 | -0.621320568 | -2.849601578 | 1e-10 |
| GO:0042113 | B cell activation | 268 | -0.626180734 | -2.856573138 | 1e-10 |
| GO:0002819 | regulation of adaptive immune response | 191 | -0.65137025 | -2.870124503 | 1e-10 |
| GO:0002697 | regulation of immune effector process | 356 | -0.617790718 | -2.884713489 | 1e-10 |
| GO:0002822 | regulation of adaptive immune response based on somatic rec | 177 | -0.660792828 | -2.887603033 | 1e-10 |
| GO:1903131 | mononuclear cell differentiation | 464 | -0.608961034 | -2.894564008 | 1e-10 |
| GO:0032609 | type II interferon production | 108 | -0.708263673 | -2.903201803 | 1e-10 |
| GO:0032649 | regulation of type II interferon production | 108 | -0.708263673 | -2.903201803 | 1e-10 |
| GO:0030098 | lymphocyte differentiation | 415 | -0.619550736 | -2.924823372 | 1e-10 |
| GO:0002706 | regulation of lymphocyte mediated immunity | 170 | -0.680116316 | -2.957500065 | 1e-10 |
| GO:0002703 | regulation of leukocyte mediated immunity | 228 | -0.660861406 | -2.968258663 | 1e-10 |
| GO:0002429 | immune response-activating cell surface receptor signaling pa | 301 | -0.649046988 | -2.990931756 | 1e-10 |
| GO:0046631 | alpha-beta T cell activation | 170 | -0.688736879 | -2.994986768 | 1e-10 |
| GO:0002768 | immune response-regulating cell surface receptor signaling pa | 326 | -0.646249376 | -2.998091948 | 1e-10 |
| GO:0016064 | immunoglobulin mediated immune response | 139 | -0.707809256 | -3.001980272 | 1e-10 |
| GO:0050851 | antigen receptor-mediated signaling pathway | 189 | -0.682069608 | -3.00205585 | 1e-10 |
| GO:0050853 | B cell receptor signaling pathway | 71 | -0.783793128 | -3.002582671 | 1e-10 |
| GO:0019724 | B cell mediated immunity | 142 | -0.713716952 | -3.035701538 | 1e-10 |
| GO:0002443 | leukocyte mediated immunity | 379 | -0.648852587 | -3.044658054 | 1e-10 |
| GO:0002460 | adaptive immune response based on somatic recombination o | 300 | -0.68260751 | -3.144879357 | 1e-10 |
| GO:0002250 | adaptive immune response | 447 | -0.665708274 | -3.156988065 | 1e-10 |
| GO:0002449 | lymphocyte mediated immunity | 286 | -0.692498113 | -3.177323788 | 1e-10 |

| p.adjust | qvalue | rank | leading_edge |
|---|---|---|---|
| 2.40078125e-09 | 1.53659539473684e-09 | 6000 | tags=70%, list=33%, signal=47% |
| 2.40078125e-09 | 1.53659539473684e-09 | 5281 | tags=67%, list=29%, signal=48% |
| 2.40078125e-09 | 1.53659539473684e-09 | 6400 | tags=69%, list=36%, signal=45% |
| 2.40078125e-09 | 1.53659539473684e-09 | 4797 | tags=75%, list=27%, signal=55% |
| 2.40078125e-09 | 1.53659539473684e-09 | 4797 | tags=75%, list=27%, signal=55% |
| 2.40078125e-09 | 1.53659539473684e-09 | 6000 | tags=71%, list=33%, signal=48% |
| 2.40078125e-09 | 1.53659539473684e-09 | 4518 | tags=74%, list=25%, signal=55% |
| 2.40078125e-09 | 1.53659539473684e-09 | 4681 | tags=66%, list=26%, signal=49% |
| 2.40078125e-09 | 1.53659539473684e-09 | 5626 | tags=60%, list=31%, signal=42% |
| 2.40078125e-09 | 1.53659539473684e-09 | 3461 | tags=51%, list=19%, signal=42% |
| 2.40078125e-09 | 1.53659539473684e-09 | 4283 | tags=60%, list=24%, signal=46% |
| 6.61239285764003e-09 | 4.23219417147666e-09 | 5871 | tags=90%, list=33%, signal=61% |
| 2.40078125e-09 | 1.53659539473684e-09 | 6523 | tags=79%, list=36%, signal=51% |
| 2.40078125e-09 | 1.53659539473684e-09 | 5935 | tags=62%, list=33%, signal=42% |
| 2.40078125e-09 | 1.53659539473684e-09 | 3804 | tags=61%, list=21%, signal=48% |
| 2.40078125e-09 | 1.53659539473684e-09 | 6554 | tags=69%, list=36%, signal=44% |
| 2.40078125e-09 | 1.53659539473684e-09 | 6999 | tags=83%, list=39%, signal=51% |
| 1.50136752089418e-08 | 9.60934870019277e-09 | 5375 | tags=70%, list=30%, signal=50% |
| 2.40078125e-09 | 1.53659539473684e-09 | 5061 | tags=56%, list=28%, signal=40% |
| 2.40078125e-09 | 1.53659539473684e-09 | 3231 | tags=49%, list=18%, signal=40% |
| 2.40078125e-09 | 1.53659539473684e-09 | 6230 | tags=61%, list=35%, signal=40% |
| 5.79940572052296e-08 | 3.71185009978151e-08 | 4518 | tags=71%, list=25%, signal=53% |
| 2.40078125e-09 | 1.53659539473684e-09 | 4128 | tags=50%, list=23%, signal=39% |
| 2.40078125e-09 | 1.53659539473684e-09 | 5919 | tags=63%, list=33%, signal=43% |
| 1.25722075156013e-06 | 8.04671236504738e-07 | 4797 | tags=78%, list=27%, signal=57% |
| 1.35311180933895e-07 | 8.66045323702135e-08 | 3447 | tags=58%, list=19%, signal=47% |
| 2.54492280297597e-07 | 1.62885171608769e-07 | 6421 | tags=84%, list=36%, signal=54% |
| 2.40078125e-09 | 1.53659539473684e-09 | 3461 | tags=49%, list=19%, signal=40% |
| 5.79160198302684e-07 | 3.70685539770348e-07 | 4902 | tags=68%, list=27%, signal=49% |
| 2.40078125e-09 | 1.53659539473684e-09 | 6544 | tags=65%, list=36%, signal=42% |
| 2.40078125e-09 | 1.53659539473684e-09 | 4673 | tags=54%, list=26%, signal=40% |
| 3.75010708427869e-07 | 2.40021754396517e-07 | 5787 | tags=76%, list=32%, signal=52% |
| 3.75010708427869e-07 | 2.40021754396517e-07 | 5787 | tags=76%, list=32%, signal=52% |
| 2.40078125e-09 | 1.53659539473684e-09 | 5196 | tags=51%, list=29%, signal=37% |
| 1.51829938332422e-08 | 9.71771934759898e-09 | 4872 | tags=53%, list=27%, signal=39% |
| 8.83913063991632e-08 | 5.65739483127533e-08 | 5443 | tags=61%, list=30%, signal=43% |
| 2.40078125e-09 | 1.53659539473684e-09 | 5010 | tags=54%, list=28%, signal=39% |
| 3.07645926335528e-08 | 1.96905617126393e-08 | 4978 | tags=54%, list=28%, signal=39% |
| 6.2700140228853e-09 | 4.01305811285429e-09 | 3231 | tags=46%, list=18%, signal=38% |
| 2.40078125e-09 | 1.53659539473684e-09 | 6290 | tags=64%, list=35%, signal=42% |
| 8.46083246985205e-09 | 5.41526897080465e-09 | 4296 | tags=52%, list=24%, signal=40% |
| 4.47391006430352e-08 | 2.86348021140019e-08 | 4978 | tags=54%, list=28%, signal=40% |
| 5.81251418517227e-07 | 3.72024003802024e-07 | 5726 | tags=61%, list=32%, signal=42% |
| 1.49935105951111e-06 | 9.59644254610275e-07 | 6796 | tags=88%, list=38%, signal=55% |

| | | | |
|---|---|---|---|
| 1.42656494952865e-05 | 9.13058252074702e-06 | 4283 | tags=75%, list=24%, signal=57% |
| 7.37508535417991e-06 | 4.72034767475128e-06 | 5375 | tags=71%, list=30%, signal=50% |
| 8.97528175194111e-07 | 5.74453695291827e-07 | 5327 | tags=66%, list=30%, signal=47% |
| 2.40078125e-09 | 1.53659539473684e-09 | 7057 | tags=61%, list=39%, signal=38% |
| 2.9206065742294e-06 | 1.86930425743663e-06 | 5575 | tags=68%, list=31%, signal=47% |
| 2.40078125e-09 | 1.53659539473684e-09 | 6373 | tags=63%, list=35%, signal=41% |
| 2.40078125e-09 | 1.53659539473684e-09 | 5202 | tags=57%, list=29%, signal=41% |
| 2.40078125e-09 | 1.53659539473684e-09 | 4572 | tags=50%, list=25%, signal=37% |
| 9.05185209781662e-07 | 5.79354501679153e-07 | 5575 | tags=66%, list=31%, signal=46% |
| 1.27888852978138e-05 | 8.18539475532744e-06 | 5263 | tags=76%, list=29%, signal=54% |
| 3.692618669989e-06 | 2.36342267452496e-06 | 6999 | tags=82%, list=39%, signal=50% |
| 1.92906031001614e-08 | 1.23467524937577e-08 | 6290 | tags=63%, list=35%, signal=41% |
| 9.74670979920544e-06 | 6.23828155576254e-06 | 4872 | tags=65%, list=27%, signal=47% |
| 9.74670979920544e-06 | 6.23828155576254e-06 | 4872 | tags=65%, list=27%, signal=47% |
| 9.74670979920544e-06 | 6.23828155576254e-06 | 4872 | tags=65%, list=27%, signal=47% |
| 9.74670979920544e-06 | 6.23828155576254e-06 | 4872 | tags=65%, list=27%, signal=47% |
| 2.40078125e-09 | 1.53659539473684e-09 | 5609 | tags=56%, list=31%, signal=39% |
| 6.62640289085868e-06 | 4.24116114942348e-06 | 5010 | tags=61%, list=28%, signal=44% |
| 1.3278786092268e-05 | 8.49895072992373e-06 | 4872 | tags=64%, list=27%, signal=47% |
| 8.72688161326398e-06 | 5.58555099401707e-06 | 4872 | tags=65%, list=27%, signal=48% |
| 8.72688161326398e-06 | 5.58555099401707e-06 | 4872 | tags=65%, list=27%, signal=48% |
| 8.72688161326398e-06 | 5.58555099401707e-06 | 4872 | tags=65%, list=27%, signal=48% |
| 4.02781682090929e-05 | 2.57796281016974e-05 | 3474 | tags=49%, list=19%, signal=39% |
| 5.87447306250232e-06 | 3.75989618143956e-06 | 5919 | tags=70%, list=33%, signal=47% |
| 2.40078125e-09 | 1.53659539473684e-09 | 4721 | tags=47%, list=26%, signal=35% |
| 6.87314898067095e-07 | 4.39908845132775e-07 | 5919 | tags=68%, list=33%, signal=45% |
| 1.31372018298639e-07 | 8.40833117615248e-08 | 3804 | tags=54%, list=21%, signal=42% |
| 2.19591308321043e-05 | 1.40547162758103e-05 | 5114 | tags=71%, list=28%, signal=51% |
| 6.94977708368245e-05 | 4.44813348206301e-05 | 5950 | tags=83%, list=33%, signal=56% |
| 1.9991258466107e-07 | 1.27951997684145e-07 | 4978 | tags=53%, list=28%, signal=38% |
| 7.76027559531037e-07 | 4.96688473455989e-07 | 7410 | tags=74%, list=41%, signal=44% |
| 2.40078125e-09 | 1.53659539473684e-09 | 6521 | tags=63%, list=36%, signal=40% |
| 0.000149081 | 9.54178510083479e-05 | 3701 | tags=77%, list=21%, signal=61% |
| 2.40078125e-09 | 1.53659539473684e-09 | 5091 | tags=51%, list=28%, signal=37% |
| 2.11597750871779e-08 | 1.35430968367588e-08 | 5099 | tags=48%, list=28%, signal=35% |
| 2.54492280297597e-07 | 1.62885171608769e-07 | 6853 | tags=68%, list=38%, signal=42% |
| 6.31365419658668e-05 | 4.04098955805992e-05 | 5013 | tags=70%, list=28%, signal=51% |
| 7.57198827181604e-05 | 4.84637336595073e-05 | 5892 | tags=81%, list=33%, signal=55% |
| 6.87314898067095e-07 | 4.39908845132775e-07 | 4978 | tags=53%, list=28%, signal=38% |
| 1.39175963345171e-06 | 8.90781466800666e-07 | 5726 | tags=61%, list=32%, signal=42% |
| 4.99423099913266e-05 | 3.19650628457683e-05 | 4673 | tags=62%, list=26%, signal=46% |
| 1.18296165430745e-05 | 7.57142463587264e-06 | 4872 | tags=62%, list=27%, signal=45% |
| 1.18296165430745e-05 | 7.57142463587264e-06 | 4872 | tags=62%, list=27%, signal=45% |
| 1.18296165430745e-05 | 7.57142463587264e-06 | 4872 | tags=62%, list=27%, signal=45% |
| 1.19006516305094e-05 | 7.61688991440109e-06 | 5010 | tags=61%, list=28%, signal=44% |

| | | | |
|---|---|---|---|
| 5.05448985749289e-05 | 3.23507434830543e-05 | 3994 | tags=61%, list=22%, signal=48% |
| 1.68594089324607e-05 | 1.0790691623239e-05 | 6182 | tags=69%, list=34%, signal=46% |
| 0.000160455 | 0.000102698 | 5475 | tags=79%, list=30%, signal=55% |
| 0.00022205 | 0.000142121 | 4570 | tags=86%, list=25%, signal=64% |
| 2.35926214165754e-07 | 1.51002151564119e-07 | 3804 | tags=53%, list=21%, signal=42% |
| 0.000224456 | 0.000143661 | 3080 | tags=55%, list=17%, signal=45% |
| 2.40078125e-09 | 1.53659539473684e-09 | 7057 | tags=62%, list=39%, signal=38% |
| 2.56938577331974e-08 | 1.64450898914071e-08 | 5099 | tags=47%, list=28%, signal=34% |
| 2.40078125e-09 | 1.53659539473684e-09 | 7057 | tags=61%, list=39%, signal=38% |
| 2.40078125e-09 | 1.53659539473684e-09 | 7057 | tags=61%, list=39%, signal=38% |
| 0.000220216 | 0.000140947 | 1421 | tags=46%, list=8%, signal=42% |
| 2.40078125e-09 | 1.53659539473684e-09 | 7057 | tags=60%, list=39%, signal=37% |
| 0.000238171 | 0.000152439 | 5602 | tags=79%, list=31%, signal=54% |
| 0.000137841 | 8.82235449983935e-05 | 5375 | tags=72%, list=30%, signal=51% |
| 2.40078125e-09 | 1.53659539473684e-09 | 5926 | tags=53%, list=33%, signal=36% |
| 3.9265498094054e-06 | 2.51314789897545e-06 | 6521 | tags=65%, list=36%, signal=42% |
| 2.40078125e-09 | 1.53659539473684e-09 | 5217 | tags=51%, list=29%, signal=37% |
| 0.000125952 | 8.0614059911675e-05 | 3808 | tags=87%, list=21%, signal=68% |
| 5.39383127825076e-05 | 3.45226634127855e-05 | 4902 | tags=65%, list=27%, signal=47% |
| 6.11314764087901e-08 | 3.91265739530458e-08 | 4518 | tags=53%, list=25%, signal=40% |
| 4.02042786055368e-06 | 2.57323358194275e-06 | 3804 | tags=52%, list=21%, signal=41% |
| 1.49935105951111e-06 | 9.59644254610275e-07 | 5080 | tags=49%, list=28%, signal=36% |
| 2.94621848746503e-06 | 1.88569689959354e-06 | 3804 | tags=51%, list=21%, signal=40% |
| 0.00022205 | 0.000142121 | 4572 | tags=93%, list=25%, signal=70% |
| 0.00022205 | 0.000142121 | 4572 | tags=93%, list=25%, signal=70% |
| 0.000325491 | 0.000208327 | 4190 | tags=72%, list=23%, signal=55% |
| 0.000298512 | 0.00019106 | 2271 | tags=44%, list=13%, signal=39% |
| 0.000737081 | 0.000471761 | 1421 | tags=43%, list=8%, signal=40% |
| 0.000365306 | 0.00023381 | 1421 | tags=40%, list=8%, signal=37% |
| 0.000777464 | 0.000497608 | 5475 | tags=76%, list=30%, signal=53% |
| 2.99400762106782e-06 | 1.91628384399446e-06 | 4572 | tags=55%, list=25%, signal=41% |
| 2.31403554331586e-06 | 1.4810746956294e-06 | 2707 | tags=31%, list=15%, signal=26% |
| 0.0006178 | 0.000395417 | 4190 | tags=69%, list=23%, signal=53% |
| 0.000206179 | 0.000131963 | 2945 | tags=82%, list=16%, signal=68% |
| 0.000320273 | 0.000204988 | 6814 | tags=76%, list=38%, signal=47% |
| 0.000124037 | 7.93887673930507e-05 | 7380 | tags=91%, list=41%, signal=54% |
| 0.000124037 | 7.93887673930507e-05 | 7380 | tags=91%, list=41%, signal=54% |
| 8.77741648834165e-05 | 5.6178953220636e-05 | 4067 | tags=46%, list=23%, signal=36% |
| 1.6600643298299e-05 | 1.06250713353561e-05 | 8050 | tags=76%, list=45%, signal=42% |
| 2.40078125e-09 | 1.53659539473684e-09 | 5010 | tags=47%, list=28%, signal=35% |
| 1.61112913664348e-06 | 1.03118666546263e-06 | 5114 | tags=55%, list=28%, signal=40% |
| 2.22480480800054e-05 | 1.42396347945571e-05 | 4938 | tags=60%, list=27%, signal=43% |
| 0.000931358 | 0.000596106 | 2500 | tags=56%, list=14%, signal=48% |
| 0.000594575 | 0.000380552 | 4570 | tags=62%, list=25%, signal=47% |
| 1.9537623423377e-05 | 1.25048553159368e-05 | 3819 | tags=47%, list=21%, signal=37% |

| | | | |
|---|---|---|---|
| 2.09580601064958e-05 | 1.34139912340033e-05 | 7306 | tags=70%, list=41%, signal=42% |
| 2.40078125e-09 | 1.53659539473684e-09 | 7054 | tags=61%, list=39%, signal=38% |
| 0.000914617 | 0.000585392 | 2945 | tags=90%, list=16%, signal=75% |
| 2.40078125e-09 | 1.53659539473684e-09 | 5138 | tags=49%, list=29%, signal=36% |
| 2.23406959082807e-06 | 1.42989330860029e-06 | 5114 | tags=55%, list=28%, signal=39% |
| 0.001192468 | 0.000763227 | 2346 | tags=61%, list=13%, signal=53% |
| 2.40078125e-09 | 1.53659539473684e-09 | 5010 | tags=47%, list=28%, signal=35% |
| 0.001121958 | 0.000718098 | 2433 | tags=54%, list=14%, signal=47% |
| 0.00218277 | 0.001397059 | 5327 | tags=74%, list=30%, signal=52% |
| 0.000579681 | 0.000371019 | 5314 | tags=64%, list=30%, signal=45% |
| 1.0299118885765e-07 | 6.59184532106527e-08 | 6769 | tags=63%, list=38%, signal=40% |
| 0.000514978 | 0.000329606 | 2338 | tags=64%, list=13%, signal=55% |
| 2.24781893437595e-06 | 1.43869343479935e-06 | 6749 | tags=65%, list=38%, signal=41% |
| 0.000670827 | 0.000429356 | 6523 | tags=81%, list=36%, signal=51% |
| 2.40078125e-09 | 1.53659539473684e-09 | 3461 | tags=41%, list=19%, signal=33% |
| 0.000199253 | 0.00012753 | 6583 | tags=69%, list=37%, signal=44% |
| 5.94696786499344e-06 | 3.80629573560561e-06 | 1501 | tags=33%, list=8%, signal=30% |
| 0.000550864 | 0.000352575 | 6800 | tags=72%, list=38%, signal=45% |
| 0.001070206 | 0.000684975 | 2433 | tags=46%, list=14%, signal=40% |
| 0.001674734 | 0.001071896 | 4889 | tags=78%, list=27%, signal=57% |
| 0.002220668 | 0.001421316 | 5010 | tags=74%, list=28%, signal=53% |
| 0.00116199 | 0.00074372 | 3461 | tags=75%, list=19%, signal=61% |
| 7.26585793446981e-07 | 4.65043778599923e-07 | 6769 | tags=63%, list=38%, signal=39% |
| 0.0016168 | 0.001034816 | 5263 | tags=71%, list=29%, signal=50% |
| 0.002150833 | 0.001376618 | 2945 | tags=60%, list=16%, signal=50% |
| 1.04953977679003e-07 | 6.71747160474821e-08 | 7302 | tags=60%, list=41%, signal=36% |
| 0.00099359 | 0.000635937 | 6493 | tags=69%, list=36%, signal=44% |
| 0.000839474 | 0.000537297 | 5010 | tags=57%, list=28%, signal=42% |
| 0.002999514 | 0.001919808 | 3176 | tags=54%, list=18%, signal=44% |
| 0.002272358 | 0.001454399 | 4599 | tags=64%, list=26%, signal=47% |
| 0.002343243 | 0.001499769 | 3080 | tags=80%, list=17%, signal=66% |
| 0.000478796 | 0.000306449 | 6634 | tags=72%, list=37%, signal=46% |
| 0.002956762 | 0.001892445 | 4242 | tags=67%, list=24%, signal=51% |
| 0.002860398 | 0.001830768 | 4949 | tags=76%, list=28%, signal=55% |
| 0.000515582 | 0.000329993 | 2433 | tags=40%, list=14%, signal=34% |
| 1.04122175691974e-05 | 6.66423297242375e-06 | 7204 | tags=62%, list=40%, signal=38% |
| 2.71188371314392e-06 | 1.73571333276566e-06 | 5006 | tags=48%, list=28%, signal=35% |
| 0.002042852 | 0.001307506 | 766 | tags=50%, list=4%, signal=48% |
| 0.002042852 | 0.001307506 | 766 | tags=50%, list=4%, signal=48% |
| 0.003096975 | 0.001982187 | 2945 | tags=67%, list=16%, signal=56% |
| 0.002862224 | 0.001831937 | 5200 | tags=70%, list=29%, signal=49% |
| 0.000558237 | 0.000357294 | 5126 | tags=48%, list=29%, signal=34% |
| 0.003784467 | 0.002422209 | 5475 | tags=76%, list=30%, signal=53% |
| 0.001558079 | 0.000997232 | 6333 | tags=67%, list=35%, signal=43% |
| 9.16957639205121e-06 | 5.86889324286149e-06 | 7082 | tags=66%, list=39%, signal=40% |

| | | | |
|---|---|---|---|
| 3.24607414691791e-06 | 2.07761643636978e-06 | 5006 | tags=48%, list=28%, signal=35% |
| 0.003752052 | 0.002401462 | 4673 | tags=81%, list=26%, signal=60% |
| 0.002017104 | 0.001291027 | 5927 | tags=69%, list=33%, signal=46% |
| 0.002517957 | 0.001611592 | 7165 | tags=89%, list=40%, signal=54% |
| 4.7407300863638e-07 | 3.03425562757832e-07 | 3685 | tags=41%, list=21%, signal=33% |
| 0.000441235 | 0.000282408 | 3695 | tags=46%, list=21%, signal=37% |
| 3.85576771807542e-07 | 2.4678445480069e-07 | 3685 | tags=40%, list=21%, signal=32% |
| 2.79256555131421e-06 | 1.78735291507719e-06 | 1501 | tags=30%, list=8%, signal=28% |
| 0.001211114 | 0.000775161 | 8304 | tags=80%, list=46%, signal=43% |
| 0.002412541 | 0.001544122 | 5200 | tags=65%, list=29%, signal=46% |
| 0.00489596 | 0.003133609 | 766 | tags=40%, list=4%, signal=38% |
| 0.00489596 | 0.003133609 | 766 | tags=40%, list=4%, signal=38% |
| 0.00489596 | 0.003133609 | 766 | tags=40%, list=4%, signal=38% |
| 0.004714427 | 0.00301742 | 2945 | tags=69%, list=16%, signal=58% |
| 0.004500042 | 0.002880206 | 2945 | tags=62%, list=16%, signal=52% |
| 0.000515582 | 0.000329993 | 7057 | tags=64%, list=39%, signal=39% |
| 2.94100603244125e-06 | 1.88236072126209e-06 | 7497 | tags=69%, list=42%, signal=40% |
| 0.001758182 | 0.001125306 | 7381 | tags=74%, list=41%, signal=44% |
| 0.004871592 | 0.003118012 | 2945 | tags=59%, list=16%, signal=49% |
| 0.004871592 | 0.003118012 | 2945 | tags=59%, list=16%, signal=49% |
| 0.004871592 | 0.003118012 | 2945 | tags=59%, list=16%, signal=49% |
| 0.004871592 | 0.003118012 | 2945 | tags=59%, list=16%, signal=49% |
| 0.001340187 | 0.000857773 | 7398 | tags=67%, list=41%, signal=39% |
| 0.005431412 | 0.00347632 | 4721 | tags=85%, list=26%, signal=62% |
| 0.002860398 | 0.001830768 | 6396 | tags=71%, list=36%, signal=46% |
| 0.000612331 | 0.000391916 | 4128 | tags=42%, list=23%, signal=32% |
| 0.005476147 | 0.003504952 | 5254 | tags=81%, list=29%, signal=58% |
| 4.32257847451253e-05 | 2.76662198079253e-05 | 6191 | tags=55%, list=34%, signal=36% |
| 0.004704581 | 0.003011119 | 413 | tags=40%, list=2%, signal=39% |
| 0.005337503 | 0.003416214 | 1429 | tags=50%, list=8%, signal=46% |
| 0.00268689 | 0.001719717 | 5476 | tags=69%, list=30%, signal=48% |
| 2.59206963075676e-05 | 1.65902755924059e-05 | 6226 | tags=57%, list=35%, signal=38% |
| 3.64654027716299e-06 | 2.33393067219725e-06 | 3452 | tags=39%, list=19%, signal=32% |
| 0.00268689 | 0.001719717 | 7216 | tags=83%, list=40%, signal=50% |
| 0.005319223 | 0.003404514 | 7165 | tags=91%, list=40%, signal=55% |
| 0.005720883 | 0.003661592 | 4936 | tags=68%, list=27%, signal=50% |
| 0.005699076 | 0.003647635 | 5217 | tags=100%, list=29%, signal=71% |
| 0.006657438 | 0.004261025 | 2065 | tags=54%, list=11%, signal=48% |
| 0.003843071 | 0.002459718 | 4396 | tags=54%, list=24%, signal=41% |
| 0.002587118 | 0.001655858 | 8414 | tags=81%, list=47%, signal=43% |
| 0.006833269 | 0.004373563 | 2065 | tags=55%, list=11%, signal=48% |
| 0.006639342 | 0.004249443 | 5481 | tags=75%, list=31%, signal=52% |
| 0.007224857 | 0.004624195 | 5200 | tags=80%, list=29%, signal=57% |
| 0.002878959 | 0.001842648 | 7711 | tags=84%, list=43%, signal=48% |
| 0.003446847 | 0.002206119 | 7235 | tags=78%, list=40%, signal=47% |

| | | | |
|---|---|---|---|
| 1.93059436695945e-06 | 1.23565710677505e-06 | 4571 | tags=45%, list=25%, signal=34% |
| 0.003498677 | 0.002239292 | 2568 | tags=47%, list=14%, signal=40% |
| 0.0075884 | 0.004856877 | 6136 | tags=76%, list=34%, signal=50% |
| 0.008369145 | 0.005356585 | 2644 | tags=47%, list=15%, signal=40% |
| 0.008117717 | 0.005195661 | 3476 | tags=56%, list=19%, signal=45% |
| 0.001426202 | 0.000912826 | 6947 | tags=63%, list=39%, signal=39% |
| 0.003815253 | 0.002441913 | 6521 | tags=62%, list=36%, signal=39% |
| 1.27459856336214e-05 | 8.15793726563162e-06 | 7036 | tags=68%, list=39%, signal=42% |
| 0.008275441 | 0.005296611 | 5978 | tags=100%, list=33%, signal=67% |
| 0.006446394 | 0.004125948 | 6800 | tags=77%, list=38%, signal=48% |
| 1.1046609701688e-05 | 7.07026914470824e-06 | 4586 | tags=47%, list=26%, signal=35% |
| 0.003735767 | 0.002391039 | 5010 | tags=57%, list=28%, signal=41% |
| 0.0094406 | 0.006042359 | 5618 | tags=76%, list=31%, signal=53% |
| 0.008316056 | 0.005322606 | 4721 | tags=83%, list=26%, signal=61% |
| 0.000839913 | 0.000537578 | 6689 | tags=72%, list=37%, signal=45% |
| 0.005203434 | 0.003330405 | 5327 | tags=60%, list=30%, signal=42% |
| 0.009255135 | 0.005923654 | 5850 | tags=71%, list=33%, signal=48% |
| 0.001763435 | 0.001128668 | 3207 | tags=38%, list=18%, signal=31% |
| 0.000326506 | 0.000208977 | 7082 | tags=67%, list=39%, signal=41% |
| 7.21071625401755e-06 | 4.61514491946214e-06 | 6812 | tags=64%, list=38%, signal=40% |
| 0.000375665 | 0.000240441 | 6947 | tags=63%, list=39%, signal=39% |
| 1.32060302222083e-07 | 8.45238408214028e-08 | 5995 | tags=52%, list=33%, signal=35% |
| 0.00854627 | 0.005469952 | 5618 | tags=70%, list=31%, signal=48% |
| 0.009583501 | 0.006133821 | 5068 | tags=80%, list=28%, signal=57% |
| 0.009583501 | 0.006133821 | 5068 | tags=80%, list=28%, signal=57% |
| 0.004202553 | 0.002689801 | 8050 | tags=76%, list=45%, signal=42% |
| 0.00042497 | 0.000271998 | 3590 | tags=37%, list=20%, signal=30% |
| 4.83236933417081e-07 | 3.09290838744863e-07 | 6241 | tags=55%, list=35%, signal=37% |
| 0.008258115 | 0.005285521 | 5779 | tags=74%, list=32%, signal=50% |
| 0.008785187 | 0.005622869 | 7216 | tags=88%, list=40%, signal=52% |
| 0.004871592 | 0.003118012 | 2540 | tags=44%, list=14%, signal=38% |
| 2.22068191259957e-07 | 1.42132466257636e-07 | 7214 | tags=61%, list=40%, signal=37% |
| 0.003914175 | 0.002505228 | 5367 | tags=64%, list=30%, signal=45% |
| 0.010698764 | 0.006847634 | 2500 | tags=50%, list=14%, signal=43% |
| 0.009782599 | 0.006261252 | 1351 | tags=60%, list=8%, signal=56% |
| 2.40078125e-09 | 1.53659539473684e-09 | 4967 | tags=45%, list=28%, signal=33% |
| 0.005720883 | 0.003661592 | 6928 | tags=71%, list=39%, signal=44% |
| 0.010523823 | 0.006735665 | 4437 | tags=75%, list=25%, signal=57% |
| 0.011798485 | 0.007551499 | 2540 | tags=64%, list=14%, signal=55% |
| 0.010737936 | 0.006872706 | 5726 | tags=75%, list=32%, signal=51% |
| 0.005840344 | 0.003738052 | 5892 | tags=63%, list=33%, signal=42% |
| 0.01211881 | 0.00775652 | 4437 | tags=71%, list=25%, signal=54% |
| 1.36187527089131e-05 | 8.71654287310674e-06 | 6290 | tags=59%, list=35%, signal=39% |
| 0.003676448 | 0.002353073 | 8050 | tags=74%, list=45%, signal=41% |
| 0.012312568 | 0.007880533 | 4902 | tags=71%, list=27%, signal=52% |

| | | | |
|---|---|---|---|
| 0.000988337 | 0.000632575 | 6583 | tags=65%, list=37%, signal=41% |
| 0.012481998 | 0.007988975 | 681 | tags=27%, list=4%, signal=26% |
| 0.005877436 | 0.003761792 | 6800 | tags=69%, list=38%, signal=43% |
| 0.01174085 | 0.007514611 | 2701 | tags=54%, list=15%, signal=46% |
| 7.78152728231975e-05 | 4.98048665867897e-05 | 6633 | tags=56%, list=37%, signal=36% |
| 0.010595489 | 0.006781534 | 2945 | tags=70%, list=16%, signal=59% |
| 0.010595489 | 0.006781534 | 2945 | tags=70%, list=16%, signal=59% |
| 0.000124567 | 7.97276552616992e-05 | 6853 | tags=56%, list=38%, signal=35% |
| 8.97660328725566e-05 | 5.74538278803062e-05 | 6963 | tags=60%, list=39%, signal=37% |
| 0.001294795 | 0.000828721 | 6923 | tags=62%, list=39%, signal=38% |
| 0.012615778 | 0.008074599 | 5892 | tags=85%, list=33%, signal=57% |
| 0.000142114 | 9.09583770944297e-05 | 6836 | tags=63%, list=38%, signal=39% |
| 0.003055125 | 0.001955402 | 964 | tags=25%, list=5%, signal=24% |
| 0.0051755 | 0.003312526 | 7120 | tags=70%, list=40%, signal=42% |
| 0.0051755 | 0.003312526 | 7120 | tags=70%, list=40%, signal=42% |
| 0.003031492 | 0.001940275 | 6629 | tags=63%, list=37%, signal=40% |
| 5.40345803223477e-05 | 3.45842784634616e-05 | 6199 | tags=52%, list=35%, signal=35% |
| 0.01348443 | 0.008630571 | 1351 | tags=55%, list=8%, signal=50% |
| 0.000265548 | 0.000169961 | 7464 | tags=66%, list=42%, signal=39% |
| 0.000197807 | 0.000126604 | 6994 | tags=74%, list=39%, signal=46% |
| 0.011214893 | 0.007177977 | 5159 | tags=61%, list=29%, signal=43% |
| 0.000257053 | 0.000164524 | 6583 | tags=59%, list=37%, signal=38% |
| 0.000973656 | 0.000623178 | 6994 | tags=73%, list=39%, signal=45% |
| 2.14214367007849e-05 | 1.37105706665581e-05 | 5725 | tags=47%, list=32%, signal=32% |
| 0.000297942 | 0.000190695 | 7383 | tags=63%, list=41%, signal=37% |
| 0.004315137 | 0.002761859 | 6928 | tags=66%, list=39%, signal=41% |
| 0.013754613 | 0.008803499 | 2536 | tags=47%, list=14%, signal=41% |
| 1.43864844450764e-06 | 9.20792168997387e-07 | 6634 | tags=58%, list=37%, signal=37% |
| 0.016404784 | 0.010499713 | 6333 | tags=87%, list=35%, signal=56% |
| 0.010164957 | 0.006505976 | 6928 | tags=70%, list=39%, signal=43% |
| 1.00146266657267e-05 | 6.40975899597865e-06 | 6620 | tags=64%, list=37%, signal=41% |
| 0.014270127 | 0.009133448 | 5871 | tags=100%, list=33%, signal=67% |
| 0.001731042 | 0.001107936 | 6583 | tags=64%, list=37%, signal=40% |
| 0.016677521 | 0.010674276 | 4989 | tags=75%, list=28%, signal=54% |
| 0.016786637 | 0.010744115 | 1954 | tags=38%, list=11%, signal=33% |
| 0.007379605 | 0.00472324 | 6226 | tags=62%, list=35%, signal=41% |
| 0.014622419 | 0.009358929 | 6929 | tags=76%, list=39%, signal=47% |
| 0.007293484 | 0.00466812 | 4109 | tags=52%, list=23%, signal=41% |
| 0.001897952 | 0.001214765 | 4619 | tags=49%, list=26%, signal=37% |
| 0.015319055 | 0.009804804 | 5978 | tags=76%, list=33%, signal=51% |
| 0.017237028 | 0.011032383 | 6937 | tags=94%, list=39%, signal=58% |
| 0.016791783 | 0.010747408 | 5273 | tags=69%, list=29%, signal=49% |
| 0.016791783 | 0.010747408 | 5273 | tags=69%, list=29%, signal=49% |
| 0.016791783 | 0.010747408 | 5864 | tags=85%, list=33%, signal=57% |
| 0.016742551 | 0.010715898 | 5159 | tags=90%, list=29%, signal=64% |

| | | | |
|---|---|---|---|
| 0.007147253 | 0.004574526 | 1421 | tags=30%, list=8%, signal=28% |
| 0.013220024 | 0.008461341 | 5509 | tags=60%, list=31%, signal=42% |
| 0.007082896 | 0.004533335 | 3252 | tags=34%, list=18%, signal=28% |
| 0.014820792 | 0.009485896 | 4003 | tags=57%, list=22%, signal=44% |
| 0.000889907 | 0.000569576 | 6583 | tags=60%, list=37%, signal=38% |
| 0.000257153 | 0.000164588 | 6226 | tags=53%, list=35%, signal=35% |
| 0.000991737 | 0.000634751 | 5150 | tags=51%, list=29%, signal=37% |
| 0.014886591 | 0.00952801 | 7711 | tags=83%, list=43%, signal=48% |
| 0.001457235 | 0.000932688 | 7082 | tags=65%, list=39%, signal=40% |
| 0.007741597 | 0.00495493 | 4110 | tags=44%, list=23%, signal=34% |
| 0.007874141 | 0.005039763 | 5126 | tags=46%, list=29%, signal=33% |
| 0.001167738 | 0.000747398 | 3698 | tags=40%, list=21%, signal=32% |
| 0.019174416 | 0.012272388 | 5992 | tags=78%, list=33%, signal=52% |
| 0.018609546 | 0.011910849 | 4902 | tags=69%, list=27%, signal=50% |
| 0.001398689 | 0.000895217 | 6226 | tags=56%, list=35%, signal=37% |
| 0.001055804 | 0.000675756 | 7981 | tags=71%, list=44%, signal=40% |
| 0.019926535 | 0.012753774 | 3474 | tags=53%, list=19%, signal=43% |
| 0.00910042 | 0.005824631 | 6226 | tags=61%, list=35%, signal=40% |
| 0.01993622 | 0.012759973 | 3839 | tags=73%, list=21%, signal=57% |
| 0.019954747 | 0.012771831 | 4572 | tags=62%, list=25%, signal=46% |
| 0.020017522 | 0.012812009 | 6347 | tags=78%, list=35%, signal=50% |
| 0.005088476 | 0.003256827 | 6923 | tags=61%, list=39%, signal=37% |
| 2.08612199176189e-08 | 1.33520096651895e-08 | 5257 | tags=42%, list=29%, signal=31% |
| 0.008643313 | 0.005532064 | 5789 | tags=60%, list=32%, signal=41% |
| 0.021883995 | 0.014006626 | 5864 | tags=92%, list=33%, signal=62% |
| 0.007697752 | 0.004926867 | 3207 | tags=36%, list=18%, signal=30% |
| 0.018495014 | 0.011837544 | 5196 | tags=73%, list=29%, signal=52% |
| 0.021244214 | 0.013597141 | 2856 | tags=60%, list=16%, signal=50% |
| 0.019514676 | 0.012490168 | 7673 | tags=95%, list=43%, signal=54% |
| 0.01258393 | 0.008054215 | 4109 | tags=58%, list=23%, signal=45% |
| 0.000192526 | 0.000123224 | 6583 | tags=54%, list=37%, signal=35% |
| 0.020819076 | 0.013325036 | 4721 | tags=68%, list=26%, signal=50% |
| 0.019775004 | 0.012656788 | 7459 | tags=90%, list=42%, signal=53% |
| 6.48695980056418e-09 | 4.15191203088159e-09 | 4562 | tags=41%, list=25%, signal=32% |
| 0.01993622 | 0.012759973 | 3941 | tags=67%, list=22%, signal=52% |
| 0.01779503 | 0.011389526 | 2530 | tags=44%, list=14%, signal=38% |
| 0.023024106 | 0.014736343 | 5721 | tags=82%, list=32%, signal=56% |
| 0.015467469 | 0.009899795 | 5410 | tags=66%, list=30%, signal=46% |
| 0.012986542 | 0.008311903 | 6634 | tags=69%, list=37%, signal=43% |
| 0.021034451 | 0.013462884 | 5779 | tags=75%, list=32%, signal=51% |
| 0.001523597 | 0.000975163 | 6953 | tags=58%, list=39%, signal=36% |
| 0.016571568 | 0.010606462 | 4973 | tags=55%, list=28%, signal=40% |
| 0.019926535 | 0.012753774 | 8304 | tags=88%, list=46%, signal=47% |
| 1.14543542587059e-08 | 7.33124186287767e-09 | 3461 | tags=34%, list=19%, signal=28% |
| 0.016865372 | 0.010794508 | 3383 | tags=37%, list=19%, signal=30% |

| | | | |
|---|---|---|---|
| 0.014886591 | 0.00952801 | 2568 | tags=45%, list=14%, signal=39% |
| 0.025612444 | 0.016392982 | 4003 | tags=71%, list=22%, signal=56% |
| 0.0169461 | 0.010846178 | 6934 | tags=70%, list=39%, signal=43% |
| 0.014328845 | 0.00917103 | 1903 | tags=35%, list=11%, signal=32% |
| 0.025640321 | 0.016410824 | 1351 | tags=46%, list=8%, signal=43% |
| 0.025760829 | 0.016487954 | 5825 | tags=77%, list=32%, signal=52% |
| 0.026207859 | 0.016774071 | 838 | tags=38%, list=5%, signal=37% |
| 0.027537482 | 0.017625082 | 4437 | tags=62%, list=25%, signal=47% |
| 0.025612444 | 0.016392982 | 4807 | tags=65%, list=27%, signal=47% |
| 0.011137669 | 0.007128551 | 6226 | tags=63%, list=35%, signal=41% |
| 0.013096254 | 0.008382123 | 5126 | tags=45%, list=29%, signal=32% |
| 0.011647848 | 0.007455085 | 4110 | tags=44%, list=23%, signal=34% |
| 0.001829079 | 0.001170683 | 7547 | tags=63%, list=42%, signal=37% |
| 0.009643144 | 0.006171995 | 6098 | tags=63%, list=34%, signal=42% |
| 0.014287554 | 0.009144602 | 7533 | tags=69%, list=42%, signal=40% |
| 0.016565859 | 0.010602808 | 6853 | tags=65%, list=38%, signal=40% |
| 0.028204734 | 0.01805215 | 6142 | tags=78%, list=34%, signal=51% |
| 0.022601414 | 0.014465803 | 2568 | tags=44%, list=14%, signal=38% |
| 0.016530052 | 0.01057989 | 7145 | tags=72%, list=40%, signal=43% |
| 0.013862645 | 0.008872644 | 6825 | tags=71%, list=38%, signal=44% |
| 0.004558184 | 0.002917419 | 4619 | tags=49%, list=26%, signal=37% |
| 0.026605922 | 0.017028847 | 5196 | tags=71%, list=29%, signal=51% |
| 0.026605922 | 0.017028847 | 5196 | tags=71%, list=29%, signal=51% |
| 0.031162613 | 0.01994531 | 7216 | tags=100%, list=40%, signal=60% |
| 0.026608164 | 0.017030282 | 1429 | tags=41%, list=8%, signal=38% |
| 0.008088116 | 0.005176716 | 7246 | tags=63%, list=40%, signal=38% |
| 0.029846523 | 0.019102961 | 3056 | tags=70%, list=17%, signal=58% |
| 0.029846523 | 0.019102961 | 3056 | tags=70%, list=17%, signal=58% |
| 0.028687673 | 0.018361251 | 7238 | tags=84%, list=40%, signal=50% |
| 0.031662446 | 0.020265223 | 2581 | tags=58%, list=14%, signal=50% |
| 0.028687673 | 0.018361251 | 4835 | tags=53%, list=27%, signal=39% |
| 1.39117677379863e-06 | 8.90408413462843e-07 | 5367 | tags=44%, list=30%, signal=31% |
| 0.005249974 | 0.003360192 | 6994 | tags=77%, list=39%, signal=47% |
| 0.0214188 | 0.013708883 | 7120 | tags=69%, list=40%, signal=42% |
| 0.00117903 | 0.000754626 | 5150 | tags=48%, list=29%, signal=34% |
| 0.02051641 | 0.013131318 | 3722 | tags=40%, list=21%, signal=32% |
| 0.000536217 | 0.0003432 | 6583 | tags=55%, list=37%, signal=35% |
| 0.01335455 | 0.008547443 | 5200 | tags=58%, list=29%, signal=41% |
| 0.010466028 | 0.006698674 | 2411 | tags=30%, list=13%, signal=26% |
| 0.031574961 | 0.02020923 | 5865 | tags=71%, list=33%, signal=48% |
| 0.032176706 | 0.02059437 | 7673 | tags=94%, list=43%, signal=54% |
| 0.008673889 | 0.005551634 | 1421 | tags=23%, list=8%, signal=21% |
| 0.026552565 | 0.016994697 | 5949 | tags=65%, list=33%, signal=44% |
| 0.034231733 | 0.021909669 | 964 | tags=29%, list=5%, signal=27% |
| 0.030101159 | 0.019265938 | 2917 | tags=35%, list=16%, signal=29% |

| | | | |
|---|---|---|---|
| 0.037565322 | 0.024043299 | 5281 | tags=62%, list=29%, signal=44% |
| 1.14420695329899e-06 | 7.32337915028743e-07 | 5361 | tags=42%, list=30%, signal=30% |
| 0.008985453 | 0.005751047 | 8347 | tags=74%, list=46%, signal=40% |
| 0.01466243 | 0.009384538 | 4271 | tags=43%, list=24%, signal=33% |
| 0.024070424 | 0.015406028 | 6906 | tags=75%, list=38%, signal=46% |
| 0.037738534 | 0.024154161 | 2701 | tags=50%, list=15%, signal=43% |
| 0.015319055 | 0.009804804 | 6947 | tags=66%, list=39%, signal=41% |
| 0.03970624 | 0.025413571 | 5841 | tags=75%, list=33%, signal=51% |
| 0.000371521 | 0.000237788 | 6680 | tags=56%, list=37%, signal=35% |
| 0.030101159 | 0.019265938 | 7853 | tags=88%, list=44%, signal=49% |
| 0.033658436 | 0.021542737 | 7238 | tags=86%, list=40%, signal=51% |
| 0.034073298 | 0.021808264 | 5779 | tags=68%, list=32%, signal=46% |
| 6.44726303134527e-07 | 4.12650452123543e-07 | 5429 | tags=44%, list=30%, signal=32% |
| 0.039463685 | 0.025258326 | 5858 | tags=79%, list=33%, signal=53% |
| 0.026016144 | 0.016651366 | 6947 | tags=60%, list=39%, signal=37% |
| 0.02735891 | 0.017510789 | 4100 | tags=50%, list=23%, signal=39% |
| 0.040181561 | 0.025717796 | 1903 | tags=53%, list=11%, signal=48% |
| 0.034606056 | 0.022149251 | 8050 | tags=77%, list=45%, signal=43% |
| 0.003422202 | 0.002190345 | 3713 | tags=40%, list=21%, signal=32% |
| 0.028285022 | 0.018103538 | 6974 | tags=71%, list=39%, signal=44% |
| 0.041930454 | 0.026837157 | 5513 | tags=69%, list=31%, signal=48% |
| 0.041930454 | 0.026837157 | 5513 | tags=69%, list=31%, signal=48% |
| 0.041930454 | 0.026837157 | 5513 | tags=69%, list=31%, signal=48% |
| 0.007223361 | 0.004623238 | 2832 | tags=34%, list=16%, signal=29% |
| 5.57084427476764e-05 | 3.5655616926382e-05 | 3708 | tags=38%, list=21%, signal=30% |
| 0.042020349 | 0.026894693 | 3947 | tags=47%, list=22%, signal=36% |
| 0.042020349 | 0.026894693 | 3947 | tags=47%, list=22%, signal=36% |
| 0.016227749 | 0.010386404 | 7306 | tags=67%, list=41%, signal=40% |
| 0.037451286 | 0.023970311 | 7695 | tags=80%, list=43%, signal=46% |
| 0.018940377 | 0.012122594 | 5200 | tags=61%, list=29%, signal=43% |
| 0.039057468 | 0.024998331 | 7673 | tags=89%, list=43%, signal=51% |
| 0.037201287 | 0.023810302 | 2286 | tags=32%, list=13%, signal=28% |
| 7.35678245871129e-05 | 4.70863309438815e-05 | 4258 | tags=38%, list=24%, signal=29% |
| 0.044405776 | 0.028421461 | 5080 | tags=83%, list=28%, signal=60% |
| 0.044405776 | 0.028421461 | 838 | tags=42%, list=5%, signal=40% |
| 0.000808916 | 0.000517738 | 5080 | tags=44%, list=28%, signal=32% |
| 0.045986191 | 0.02943299 | 5885 | tags=90%, list=33%, signal=61% |
| 0.046038249 | 0.029466308 | 1149 | tags=23%, list=6%, signal=22% |
| 0.00826734 | 0.005291426 | 4456 | tags=40%, list=25%, signal=30% |
| 0.004784862 | 0.003062502 | 3713 | tags=40%, list=21%, signal=32% |
| 0.013406808 | 0.00858089 | 6008 | tags=61%, list=33%, signal=41% |
| 0.013406808 | 0.00858089 | 6008 | tags=61%, list=33%, signal=41% |
| 0.013406808 | 0.00858089 | 6008 | tags=61%, list=33%, signal=41% |
| 0.028687673 | 0.018361251 | 3266 | tags=45%, list=18%, signal=37% |
| 0.041283144 | 0.026422852 | 6825 | tags=65%, list=38%, signal=40% |

| | | |
|---|---|---|
| 0.0215093 | 0.013766806 | 5325 tags=51%, list=30%, signal=36% |
| 0.001200106 | 0.000768116 | 5366 tags=41%, list=30%, signal=29% |
| 0.048379178 | 0.030964596 | 4003 tags=69%, list=22%, signal=54% |
| 0.022608832 | 0.014470551 | 4264 tags=41%, list=24%, signal=31% |
| 0.010218551 | 0.006540279 | 8130 tags=72%, list=45%, signal=40% |
| 0.001215063 | 0.000777689 | 6994 tags=68%, list=39%, signal=42% |
| 0.001215063 | 0.000777689 | 6994 tags=68%, list=39%, signal=42% |
| 1.16647490940242e-07 | 7.46590291749336e-08 | 7236 tags=57%, list=40%, signal=35% |
| 0.041930454 | 0.026837157 | 8304 tags=90%, list=46%, signal=49% |
| 5.89061010651781e-05 | 3.77022453081286e-05 | 3708 tags=37%, list=21%, signal=30% |
| 0.016679207 | 0.010675356 | 6491 tags=74%, list=36%, signal=47% |
| 0.016679207 | 0.010675356 | 6491 tags=74%, list=36%, signal=47% |
| 0.043249854 | 0.027681625 | 7203 tags=70%, list=40%, signal=42% |
| 0.043249854 | 0.027681625 | 7203 tags=70%, list=40%, signal=42% |
| 0.045346015 | 0.029023252 | 6497 tags=78%, list=36%, signal=50% |
| 0.049702819 | 0.031811779 | 4003 tags=62%, list=22%, signal=49% |
| 0.009568943 | 0.006124503 | 4619 tags=48%, list=26%, signal=36% |
| 0.03826258 | 0.024489571 | 5901 tags=59%, list=33%, signal=40% |
| 0.03826258 | 0.024489571 | 5901 tags=59%, list=33%, signal=40% |
| 0.032927281 | 0.021074768 | 6825 tags=68%, list=38%, signal=42% |
| 0.009640633 | 0.006170388 | 4456 tags=40%, list=25%, signal=30% |
| 0.000857471 | 0.000548816 | 6837 tags=54%, list=38%, signal=33% |
| 0.040636255 | 0.026008818 | 8304 tags=83%, list=46%, signal=45% |
| 0.049408047 | 0.031623113 | 1651 tags=29%, list=9%, signal=27% |
| 0.025629005 | 0.016403582 | 6396 tags=63%, list=36%, signal=41% |
| 0.001426202 | 0.000912826 | 6994 tags=67%, list=39%, signal=41% |
| 0.037738534 | 0.024154161 | 2307 tags=43%, list=13%, signal=37% |
| 0.004306532 | 0.002756351 | 5467 tags=51%, list=30%, signal=36% |
| 2.04564620132152e-07 | 1.30929485233674e-07 | 4978 tags=41%, list=28%, signal=30% |
| 0.048611787 | 0.031113475 | 7105 tags=81%, list=40%, signal=49% |
| 0.048611787 | 0.031113475 | 7105 tags=81%, list=40%, signal=49% |
| 0.037279804 | 0.023860556 | 5112 tags=53%, list=28%, signal=38% |
| 0.017958291 | 0.01149402 | 7906 tags=71%, list=44%, signal=40% |
| 0.005059049 | 0.003237993 | 5926 tags=52%, list=33%, signal=35% |
| 0.049449086 | 0.03164938 | 4466 tags=67%, list=25%, signal=50% |
| 0.042658747 | 0.027303293 | 7284 tags=73%, list=41%, signal=43% |
| 0.01364406 | 0.00873274 | 5163 tags=43%, list=29%, signal=31% |
| 0.008464175 | 0.005417408 | 3713 tags=39%, list=21%, signal=31% |
| 0.025612444 | 0.016392982 | 7740 tags=70%, list=43%, signal=40% |
| 0.046564721 | 0.029803272 | 8304 tags=85%, list=46%, signal=46% |
| 0.009341415 | 0.005978877 | 4390 tags=40%, list=24%, signal=30% |
| 0.001440706 | 0.000922109 | 3980 tags=37%, list=22%, signal=29% |
| 0.000274302 | 0.000175564 | 7351 tags=58%, list=41%, signal=35% |
| 5.02524607157575e-05 | 3.21635716332036e-05 | 7194 tags=55%, list=40%, signal=34% |
| 1.19223094374572e-06 | 7.63075177141788e-07 | 4335 tags=35%, list=24%, signal=28% |

| | | | |
|---|---|---|---|
| 0.032727239 | 0.020946733 | 964 | tags=25%, list=5%, signal=24% |
| 0.010714281 | 0.006857566 | 6347 | tags=60%, list=35%, signal=39% |
| 0.03452902 | 0.022099945 | 1921 | tags=26%, list=11%, signal=24% |
| 0.004435426 | 0.002838849 | 5626 | tags=46%, list=31%, signal=32% |
| 0.020760832 | 0.013287758 | 6098 | tags=61%, list=34%, signal=41% |
| 0.03761118 | 0.02407265 | 4283 | tags=49%, list=24%, signal=37% |
| 0.017922733 | 0.011471261 | 7740 | tags=63%, list=43%, signal=36% |
| 0.026207859 | 0.016774071 | 7547 | tags=67%, list=42%, signal=39% |
| 0.049573982 | 0.031729318 | 1939 | tags=27%, list=11%, signal=24% |
| 0.024236616 | 0.015512398 | 4443 | tags=39%, list=25%, signal=30% |
| 0.000475001 | 0.00030402 | 3708 | tags=37%, list=21%, signal=29% |
| 0.016757863 | 0.010725698 | 3658 | tags=41%, list=20%, signal=33% |
| 0.014788521 | 0.009465241 | 7547 | tags=66%, list=42%, signal=38% |
| 0.024998056 | 0.015999749 | 4277 | tags=49%, list=24%, signal=37% |
| 0.043922377 | 0.028112067 | 6934 | tags=68%, list=39%, signal=42% |
| 0.000210541 | 0.000134755 | 3471 | tags=30%, list=19%, signal=24% |
| 0.000726106 | 0.000464736 | 5325 | tags=43%, list=30%, signal=31% |
| 0.044170957 | 0.028271168 | 8393 | tags=78%, list=47%, signal=42% |
| 0.000566716 | 0.000362721 | 3452 | tags=35%, list=19%, signal=29% |
| 0.041319233 | 0.026445951 | 6825 | tags=61%, list=38%, signal=38% |
| 0.032018111 | 0.020492863 | 6355 | tags=62%, list=35%, signal=40% |
| 0.009994896 | 0.006397131 | 5006 | tags=41%, list=28%, signal=30% |
| 0.001106694 | 0.000708328 | 4869 | tags=41%, list=27%, signal=30% |
| 3.83584364896373e-05 | 2.45509235209507e-05 | 5578 | tags=45%, list=31%, signal=32% |
| 0.00557679 | 0.003569367 | 7321 | tags=62%, list=41%, signal=37% |
| 8.8213607563098e-05 | 5.6460213996831e-05 | 4499 | tags=39%, list=25%, signal=30% |
| 0.04128572 | 0.026424501 | 5224 | tags=56%, list=29%, signal=40% |
| 0.011299116 | 0.007231883 | 4303 | tags=48%, list=24%, signal=37% |
| 0.04128572 | 0.026424501 | 5902 | tags=50%, list=33%, signal=34% |
| 0.010626332 | 0.006801275 | 7515 | tags=56%, list=42%, signal=33% |
| 0.010626332 | 0.006801275 | 3713 | tags=38%, list=21%, signal=30% |
| 0.01308593 | 0.008375515 | 7383 | tags=61%, list=41%, signal=36% |
| 0.000824242 | 0.000527548 | 3708 | tags=36%, list=21%, signal=29% |
| 0.04596922 | 0.029422128 | 1529 | tags=30%, list=9%, signal=27% |
| 0.009013705 | 0.005769129 | 1421 | tags=20%, list=8%, signal=19% |
| 0.003490734 | 0.002234208 | 6892 | tags=54%, list=38%, signal=34% |
| 4.24253712495206e-06 | 2.71539233664101e-06 | 6228 | tags=47%, list=35%, signal=31% |
| 0.000137841 | 8.82235449983935e-05 | 4283 | tags=36%, list=24%, signal=28% |
| 0.000122901 | 7.86614999177004e-05 | 3584 | tags=31%, list=20%, signal=25% |
| 0.049408047 | 0.031623113 | 5902 | tags=50%, list=33%, signal=34% |
| 0.020683563 | 0.013238302 | 5808 | tags=49%, list=32%, signal=33% |
| 0.020245316 | 0.012957807 | 6658 | tags=59%, list=37%, signal=37% |
| 0.006377407 | 0.004081794 | 6154 | tags=47%, list=34%, signal=31% |
| 0.046564721 | 0.029803272 | 3118 | tags=33%, list=17%, signal=28% |
| 0.000373889 | 0.000239304 | 5517 | tags=48%, list=31%, signal=34% |

| | | | |
|---|---|---|---|
| 0.006838597 | 0.004376974 | 3740 | tags=37%, list=21%, signal=30% |
| 0.037902935 | 0.024259385 | 5100 | tags=52%, list=28%, signal=37% |
| 0.007082896 | 0.004533335 | 2338 | tags=27%, list=13%, signal=24% |
| 0.00167759 | 0.001073724 | 7679 | tags=64%, list=43%, signal=37% |
| 0.01414578 | 0.009053862 | 4314 | tags=39%, list=24%, signal=30% |
| 0.00022205 | 0.000142121 | 3452 | tags=35%, list=19%, signal=29% |
| 0.026281882 | 0.016821449 | 5994 | tags=53%, list=33%, signal=36% |
| 0.001433167 | 0.000917284 | 6355 | tags=53%, list=35%, signal=34% |
| 0.00053908 | 0.000345033 | 3452 | tags=35%, list=19%, signal=29% |
| 0.022810993 | 0.014599942 | 2049 | tags=24%, list=11%, signal=21% |
| 0.015255948 | 0.009764413 | 6634 | tags=52%, list=37%, signal=33% |
| 0.00160127 | 0.001024876 | 4869 | tags=41%, list=27%, signal=30% |
| 0.03569702 | 0.022847511 | 6296 | tags=61%, list=35%, signal=40% |
| 0.000268995 | 0.000172167 | 5179 | tags=43%, list=29%, signal=31% |
| 0.000988175 | 0.000632471 | 5111 | tags=42%, list=28%, signal=30% |
| 0.009255135 | 0.005923654 | 4021 | tags=31%, list=22%, signal=25% |
| 0.028013761 | 0.01792992 | 3131 | tags=40%, list=17%, signal=33% |
| 0.017116658 | 0.010955342 | 4908 | tags=45%, list=27%, signal=33% |
| 0.000149279 | 9.55447840762083e-05 | 4945 | tags=42%, list=28%, signal=31% |
| 0.017116658 | 0.010955342 | 7529 | tags=66%, list=42%, signal=38% |
| 0.031291994 | 0.020028119 | 4605 | tags=42%, list=26%, signal=31% |
| 0.018685691 | 0.011959585 | 7828 | tags=65%, list=44%, signal=37% |
| 0.043249854 | 0.027681625 | 5886 | tags=52%, list=33%, signal=35% |
| 0.001617122 | 0.001035022 | 5879 | tags=51%, list=33%, signal=35% |
| 0.023648114 | 0.015135733 | 7547 | tags=65%, list=42%, signal=38% |
| 0.033239647 | 0.021274695 | 7547 | tags=69%, list=42%, signal=40% |
| 0.0075884 | 0.004856877 | 6154 | tags=48%, list=34%, signal=32% |
| 0.010343288 | 0.006620115 | 2997 | tags=30%, list=17%, signal=25% |
| 0.017922733 | 0.011471261 | 5779 | tags=50%, list=32%, signal=34% |
| 0.00274164 | 0.001754759 | 3452 | tags=35%, list=19%, signal=29% |
| 0.023772257 | 0.015215189 | 7058 | tags=58%, list=39%, signal=35% |
| 0.004961861 | 0.003175788 | 6947 | tags=57%, list=39%, signal=35% |
| 0.046561254 | 0.029801052 | 5006 | tags=47%, list=28%, signal=34% |
| 0.001747866 | 0.001118704 | 6418 | tags=48%, list=36%, signal=31% |
| 0.014828674 | 0.00949094 | 4271 | tags=37%, list=24%, signal=29% |
| 0.023795114 | 0.015229818 | 7515 | tags=56%, list=42%, signal=33% |
| 0.002314305 | 0.001481247 | 5111 | tags=41%, list=28%, signal=30% |
| 3.08955650431603e-05 | 1.977438925896e-05 | 4619 | tags=37%, list=26%, signal=28% |
| 0.004148326 | 0.002655093 | 7030 | tags=56%, list=39%, signal=34% |
| 0.029921357 | 0.019150857 | 7319 | tags=56%, list=41%, signal=33% |
| 2.67287393048827e-05 | 1.7107455218173e-05 | 5062 | tags=38%, list=28%, signal=28% |
| 0.048393701 | 0.030973891 | 7547 | tags=69%, list=42%, signal=40% |
| 0.002606448 | 0.00166823 | 6204 | tags=45%, list=35%, signal=30% |
| 0.027228163 | 0.017427106 | 4772 | tags=40%, list=27%, signal=30% |
| 0.021246064 | 0.013598325 | 6093 | tags=58%, list=34%, signal=39% |

| | | | |
|---|---|---|---|
| 0.049832869 | 0.031895016 | 6491 | tags=69%, list=36%, signal=44% |
| 0.049045492 | 0.031391064 | 5406 | tags=52%, list=30%, signal=37% |
| 0.024386731 | 0.015608477 | 6626 | tags=54%, list=37%, signal=34% |
| 0.004148326 | 0.002655093 | 3452 | tags=34%, list=19%, signal=28% |
| 0.000163389 | 0.000104575 | 5041 | tags=40%, list=28%, signal=29% |
| 0.014985825 | 0.009591523 | 7737 | tags=63%, list=43%, signal=36% |
| 0.000169166 | 0.000108273 | 3122 | tags=28%, list=17%, signal=24% |
| 0.01410204 | 0.009025866 | 5084 | tags=41%, list=28%, signal=30% |
| 0.02943503 | 0.018839589 | 6634 | tags=50%, list=37%, signal=32% |
| 0.008316327 | 0.00532278 | 5325 | tags=38%, list=30%, signal=27% |
| 0.016572883 | 0.010607304 | 4567 | tags=38%, list=25%, signal=28% |
| 0.00085056 | 0.000544392 | 5506 | tags=36%, list=31%, signal=26% |
| 0.000503241 | 0.000322094 | 3122 | tags=28%, list=17%, signal=24% |
| 0.000416537 | 0.0002666 | 3126 | tags=28%, list=17%, signal=23% |
| 0.028926531 | 0.018514129 | 6626 | tags=54%, list=37%, signal=34% |
| 0.032176706 | 0.02059437 | 1104 | tags=18%, list=6%, signal=17% |
| 0.032176706 | 0.02059437 | 1104 | tags=18%, list=6%, signal=17% |
| 0.012986542 | 0.008311903 | 6535 | tags=45%, list=36%, signal=29% |
| 0.001335441 | 0.000854735 | 5591 | tags=37%, list=31%, signal=26% |
| 0.013598943 | 0.008703864 | 4464 | tags=37%, list=25%, signal=28% |
| 0.032521525 | 0.020815068 | 4075 | tags=33%, list=23%, signal=26% |
| 0.032521525 | 0.020815068 | 4075 | tags=33%, list=23%, signal=26% |
| 0.006762741 | 0.004328423 | 5404 | tags=44%, list=30%, signal=31% |
| 0.001244158 | 0.000796311 | 6226 | tags=49%, list=35%, signal=33% |
| 0.033805859 | 0.021637093 | 5323 | tags=40%, list=30%, signal=29% |
| 0.037110012 | 0.023751882 | 3699 | tags=37%, list=21%, signal=30% |
| 0.032072808 | 0.020527872 | 3740 | tags=35%, list=21%, signal=28% |
| 0.048884791 | 0.031288209 | 4335 | tags=40%, list=24%, signal=30% |
| 0.01341559 | 0.00858651 | 3128 | tags=28%, list=17%, signal=23% |
| 0.039768554 | 0.025453455 | 3071 | tags=29%, list=17%, signal=24% |
| 0.004226472 | 0.00270511 | 5132 | tags=43%, list=29%, signal=31% |
| 0.046564721 | 0.029803272 | 6101 | tags=53%, list=34%, signal=35% |
| 0.046564721 | 0.029803272 | 6101 | tags=53%, list=34%, signal=35% |
| 0.046564721 | 0.029803272 | 6101 | tags=53%, list=34%, signal=35% |
| 0.042658075 | 0.027302863 | 5853 | tags=44%, list=33%, signal=30% |
| 0.042972717 | 0.027504246 | 6098 | tags=54%, list=34%, signal=36% |
| 0.034231733 | 0.021909669 | 1497 | tags=16%, list=8%, signal=15% |
| 0.037553132 | 0.024035497 | 3740 | tags=34%, list=21%, signal=27% |
| 0.007540058 | 0.004825937 | 3411 | tags=27%, list=19%, signal=22% |
| 0.00142477 | 0.000911909 | 3215 | tags=26%, list=18%, signal=22% |
| 0.046115154 | 0.029515531 | 3740 | tags=34%, list=21%, signal=27% |
| 0.016495931 | 0.010558051 | 7393 | tags=56%, list=41%, signal=33% |
| 0.002148913 | 0.00137539 | 3156 | tags=27%, list=18%, signal=23% |
| 0.034928379 | 0.022355551 | 6947 | tags=54%, list=39%, signal=33% |
| 0.046564721 | 0.029803272 | 4733 | tags=37%, list=26%, signal=27% |

| | | | |
|---|---|---|---|
| 0.047321499 | 0.03028764 | 5006 | tags=42%, list=28%, signal=31% |
| 0.011407877 | 0.007301495 | 4945 | tags=38%, list=28%, signal=28% |
| 0.005340287 | 0.003417996 | 1851 | tags=18%, list=10%, signal=16% |
| 0.04109918 | 0.026305108 | 3812 | tags=32%, list=21%, signal=25% |
| 0.039744106 | 0.025437807 | 2061 | tags=17%, list=11%, signal=15% |
| 0.031574961 | 0.02020923 | 1851 | tags=18%, list=10%, signal=17% |
| 0.014346584 | 0.009182384 | 5220 | tags=39%, list=29%, signal=28% |
| 0.023772257 | 0.015215189 | 4702 | tags=30%, list=26%, signal=23% |
| 0.020761996 | 0.013288503 | 2990 | tags=24%, list=17%, signal=20% |
| 0.040189574 | 0.025722924 | 4990 | tags=38%, list=28%, signal=28% |
| 0.04073706 | 0.026073337 | 1851 | tags=19%, list=10%, signal=18% |
| 0.046115154 | 0.029515531 | 4839 | tags=36%, list=27%, signal=27% |
| 0.036887837 | 0.023609681 | 5497 | tags=37%, list=31%, signal=26% |
| 0.043776328 | 0.02801859 | 1898 | tags=16%, list=11%, signal=15% |
| 0.025693611 | 0.016444932 | 1463 | tags=15%, list=8%, signal=14% |
| 0.04155375 | 0.026596051 | 1497 | tags=15%, list=8%, signal=14% |
| 0.048789162 | 0.031227002 | 4335 | tags=32%, list=24%, signal=24% |
| 0.047035848 | 0.030104812 | 1931 | tags=15%, list=11%, signal=14% |
| 0.047511518 | 0.03040926 | 1931 | tags=15%, list=11%, signal=14% |
| 0.046845485 | 0.029982972 | 1463 | tags=14%, list=8%, signal=14% |
| 0.046564721 | 0.029803272 | 2688 | tags=20%, list=15%, signal=17% |
| 0.046564721 | 0.029803272 | 2708 | tags=18%, list=15%, signal=16% |
| 0.037784594 | 0.024183642 | 4226 | tags=34%, list=24%, signal=26% |
| 0.035673988 | 0.02283277 | 4226 | tags=34%, list=24%, signal=26% |
| 0.020659432 | 0.013222857 | 2759 | tags=23%, list=15%, signal=20% |
| 0.048600274 | 0.031106106 | 3885 | tags=29%, list=22%, signal=23% |
| 0.014632063 | 0.009365102 | 3161 | tags=22%, list=18%, signal=19% |
| 0.024001125 | 0.015361674 | 3866 | tags=31%, list=22%, signal=24% |
| 0.043362122 | 0.027753481 | 3694 | tags=33%, list=21%, signal=27% |
| 0.049573982 | 0.031729318 | 2881 | tags=24%, list=16%, signal=20% |
| 0.024877898 | 0.015922843 | 3217 | tags=25%, list=18%, signal=21% |
| 0.016589761 | 0.010618106 | 2661 | tags=20%, list=15%, signal=18% |
| 0.040731451 | 0.026069747 | 4342 | tags=35%, list=24%, signal=27% |
| 0.043725953 | 0.027986348 | 1939 | tags=19%, list=11%, signal=17% |
| 0.008812441 | 0.005640312 | 4129 | tags=32%, list=23%, signal=26% |
| 0.009030987 | 0.005780191 | 4129 | tags=33%, list=23%, signal=26% |
| 0.041503994 | 0.026564205 | 3369 | tags=29%, list=19%, signal=23% |
| 0.03117843 | 0.019955434 | 2523 | tags=22%, list=14%, signal=19% |
| 0.049573982 | 0.031729318 | 2755 | tags=38%, list=15%, signal=32% |
| 0.017040732 | 0.010906745 | 2624 | tags=22%, list=15%, signal=19% |
| 0.008769852 | 0.005613054 | 3748 | tags=32%, list=21%, signal=26% |
| 0.026608164 | 0.017030282 | 4119 | tags=33%, list=23%, signal=26% |
| 0.027528041 | 0.01761904 | 2624 | tags=22%, list=15%, signal=19% |
| 0.003815253 | 0.002441913 | 2187 | tags=23%, list=12%, signal=21% |
| 0.013406808 | 0.00858089 | 4198 | tags=35%, list=23%, signal=28% |

| | | | |
|---|---|---|---|
| 0.003486704 | 0.002231629 | 4195 | tags=32%, list=23%, signal=25% |
| 0.031853687 | 0.020387625 | 3214 | tags=26%, list=18%, signal=22% |
| 0.002834722 | 0.001814334 | 2993 | tags=22%, list=17%, signal=19% |
| 0.012894181 | 0.008252788 | 3653 | tags=28%, list=20%, signal=23% |
| 0.018630914 | 0.011924526 | 2657 | tags=24%, list=15%, signal=21% |
| 0.025760829 | 0.016487954 | 4026 | tags=34%, list=22%, signal=27% |
| 0.045938866 | 0.0294027 | 4116 | tags=37%, list=23%, signal=29% |
| 0.025612444 | 0.016392982 | 4176 | tags=38%, list=23%, signal=29% |
| 0.024379915 | 0.015604114 | 1214 | tags=14%, list=7%, signal=13% |
| 0.015249193 | 0.009760089 | 5634 | tags=52%, list=31%, signal=36% |
| 0.033763647 | 0.021610075 | 5222 | tags=47%, list=29%, signal=33% |
| 0.001104102 | 0.000706669 | 2359 | tags=21%, list=13%, signal=19% |
| 0.040026601 | 0.025618615 | 2586 | tags=28%, list=14%, signal=24% |
| 0.021034451 | 0.013462884 | 2007 | tags=19%, list=11%, signal=17% |
| 0.017740279 | 0.011354484 | 2661 | tags=22%, list=15%, signal=19% |
| 0.048393701 | 0.030973891 | 2662 | tags=42%, list=15%, signal=36% |
| 0.003800141 | 0.002432241 | 1901 | tags=20%, list=11%, signal=18% |
| 0.015255948 | 0.009764413 | 3072 | tags=26%, list=17%, signal=22% |
| 0.010475423 | 0.006704687 | 2641 | tags=23%, list=15%, signal=20% |
| 0.024310422 | 0.015559636 | 3968 | tags=39%, list=22%, signal=31% |
| 0.017237028 | 0.011032383 | 3369 | tags=31%, list=19%, signal=25% |
| 0.027614068 | 0.017674101 | 3376 | tags=28%, list=19%, signal=23% |
| 0.046564721 | 0.029803272 | 4116 | tags=38%, list=23%, signal=30% |
| 0.023611703 | 0.015112428 | 3774 | tags=31%, list=21%, signal=25% |
| 0.013395358 | 0.008573561 | 3043 | tags=26%, list=17%, signal=22% |
| 0.022705698 | 0.014532549 | 3214 | tags=28%, list=18%, signal=23% |
| 0.043365322 | 0.027755529 | 3315 | tags=32%, list=18%, signal=27% |
| 0.012858787 | 0.008230134 | 2187 | tags=20%, list=12%, signal=18% |
| 0.007333139 | 0.004693501 | 3043 | tags=26%, list=17%, signal=22% |
| 0.00136573 | 0.000874122 | 4342 | tags=36%, list=24%, signal=28% |
| 0.0102515 | 0.006561368 | 4119 | tags=34%, list=23%, signal=27% |
| 0.012755513 | 0.008164035 | 2746 | tags=26%, list=15%, signal=22% |
| 0.00252864 | 0.00161843 | 3654 | tags=30%, list=20%, signal=24% |
| 0.041287756 | 0.026425804 | 2530 | tags=31%, list=14%, signal=26% |
| 0.002163303 | 0.0013846 | 2766 | tags=26%, list=15%, signal=22% |
| 0.036987359 | 0.02367338 | 2797 | tags=25%, list=16%, signal=21% |
| 0.015653084 | 0.010018596 | 4351 | tags=38%, list=24%, signal=29% |
| 0.010276906 | 0.006577628 | 2236 | tags=19%, list=12%, signal=17% |
| 0.002226766 | 0.001425219 | 3654 | tags=30%, list=20%, signal=24% |
| 0.005337404 | 0.00341615 | 2809 | tags=25%, list=16%, signal=21% |
| 0.010212514 | 0.006536415 | 1988 | tags=23%, list=11%, signal=20% |
| 0.03256611 | 0.020843605 | 1901 | tags=24%, list=11%, signal=22% |
| 0.021849808 | 0.013984745 | 3774 | tags=32%, list=21%, signal=25% |
| 0.034231733 | 0.021909669 | 3722 | tags=35%, list=21%, signal=28% |
| 0.001700845 | 0.001088608 | 3654 | tags=30%, list=20%, signal=25% |

| | | | |
|---|---|---|---|
| 0.022040237 | 0.014106627 | 3318 | tags=31%, list=18%, signal=25% |
| 0.029174025 | 0.018672535 | 2587 | tags=27%, list=14%, signal=23% |
| 0.001060499 | 0.000678761 | 2407 | tags=23%, list=13%, signal=20% |
| 0.019214176 | 0.012297836 | 2578 | tags=22%, list=14%, signal=19% |
| 0.037407368 | 0.023942202 | 4176 | tags=43%, list=23%, signal=33% |
| 0.030220021 | 0.019342014 | 3220 | tags=38%, list=18%, signal=31% |
| 0.031258893 | 0.020006934 | 1188 | tags=17%, list=7%, signal=16% |
| 0.017399635 | 0.011136458 | 2578 | tags=22%, list=14%, signal=19% |
| 0.001602263 | 0.001025512 | 2586 | tags=22%, list=14%, signal=19% |
| 0.030198837 | 0.019328456 | 2371 | tags=27%, list=13%, signal=24% |
| 0.016933579 | 0.010838163 | 1807 | tags=20%, list=10%, signal=18% |
| 0.014484841 | 0.009270874 | 4104 | tags=41%, list=23%, signal=32% |
| 0.037407368 | 0.023942202 | 2766 | tags=28%, list=15%, signal=24% |
| 0.041073332 | 0.026288564 | 2624 | tags=27%, list=15%, signal=23% |
| 0.005122792 | 0.00327879 | 3297 | tags=30%, list=18%, signal=25% |
| 0.037362351 | 0.023913389 | 3214 | tags=31%, list=18%, signal=26% |
| 0.000220442 | 0.000141092 | 3654 | tags=30%, list=20%, signal=25% |
| 0.01994102 | 0.012763045 | 2079 | tags=20%, list=12%, signal=18% |
| 0.037365629 | 0.023915487 | 2187 | tags=23%, list=12%, signal=21% |
| 0.003866087 | 0.002474449 | 1928 | tags=18%, list=11%, signal=16% |
| 0.01151515 | 0.007370154 | 2578 | tags=23%, list=14%, signal=20% |
| 0.046533368 | 0.029783205 | 3063 | tags=35%, list=17%, signal=29% |
| 0.014311136 | 0.009159696 | 3297 | tags=37%, list=18%, signal=30% |
| 0.014632063 | 0.009365102 | 4215 | tags=44%, list=23%, signal=34% |
| 0.007554948 | 0.004835467 | 4293 | tags=39%, list=24%, signal=30% |
| 0.026387546 | 0.016889078 | 2555 | tags=30%, list=14%, signal=26% |
| 0.001833651 | 0.001173609 | 3590 | tags=26%, list=20%, signal=21% |
| 0.024848301 | 0.0159039 | 2193 | tags=24%, list=12%, signal=21% |
| 0.000184377 | 0.000118008 | 2729 | tags=23%, list=15%, signal=20% |
| 0.029226299 | 0.018705993 | 3560 | tags=41%, list=20%, signal=33% |
| 0.017236583 | 0.011032098 | 2265 | tags=24%, list=13%, signal=21% |
| 0.001217142 | 0.000779019 | 2993 | tags=24%, list=17%, signal=20% |
| 0.035359148 | 0.02263126 | 4031 | tags=40%, list=22%, signal=31% |
| 0.001786817 | 0.001143634 | 2756 | tags=23%, list=15%, signal=19% |
| 0.048870968 | 0.031279362 | 4202 | tags=40%, list=23%, signal=30% |
| 0.03982825 | 0.025491663 | 1757 | tags=26%, list=10%, signal=23% |
| 0.014256013 | 0.009124415 | 4116 | tags=40%, list=23%, signal=31% |
| 0.039374729 | 0.025201391 | 4285 | tags=41%, list=24%, signal=31% |
| 0.00771073 | 0.004935173 | 2881 | tags=26%, list=16%, signal=22% |
| 0.030010306 | 0.019207788 | 3299 | tags=35%, list=18%, signal=29% |
| 0.002355134 | 0.001507379 | 3653 | tags=30%, list=20%, signal=24% |
| 0.00054192 | 0.00034685 | 3658 | tags=28%, list=20%, signal=23% |
| 0.008807574 | 0.005637197 | 1953 | tags=20%, list=11%, signal=18% |
| 0.036633898 | 0.02344715 | 3560 | tags=45%, list=20%, signal=36% |
| 0.002042852 | 0.001307506 | 2746 | tags=27%, list=15%, signal=23% |

| | | |
|---|---|---|
| 0.04279651 | 0.027391467 | 4010 tags=48%, list=22%, signal=38% |
| 0.004934886 | 0.003158523 | 2179 tags=24%, list=12%, signal=22% |
| 0.049029116 | 0.031380583 | 1595 tags=32%, list=9%, signal=29% |
| 0.004629283 | 0.002962925 | 4293 tags=39%, list=24%, signal=30% |
| 0.004148326 | 0.002655093 | 3653 tags=31%, list=20%, signal=25% |
| 0.026953664 | 0.017251416 | 3654 tags=38%, list=20%, signal=30% |
| 0.018495961 | 0.01183815 | 4397 tags=43%, list=24%, signal=33% |
| 0.00547505 | 0.00350425 | 2881 tags=26%, list=16%, signal=22% |
| 0.005595511 | 0.003581349 | 2027 tags=19%, list=11%, signal=17% |
| 0.004958732 | 0.003173786 | 3278 tags=29%, list=18%, signal=24% |
| 0.003112744 | 0.00199228 | 3653 tags=31%, list=20%, signal=25% |
| 0.012401553 | 0.007937486 | 3214 tags=31%, list=18%, signal=25% |
| 0.004992368 | 0.003195314 | 3278 tags=28%, list=18%, signal=23% |
| 0.006593934 | 0.00422038 | 2661 tags=26%, list=15%, signal=22% |
| 0.00439537 | 0.002813212 | 3129 tags=27%, list=17%, signal=23% |
| 0.027849593 | 0.017824846 | 3278 tags=29%, list=18%, signal=24% |
| 0.001243038 | 0.000795594 | 3653 tags=30%, list=20%, signal=24% |
| 0.037407368 | 0.023942202 | 4574 tags=46%, list=25%, signal=35% |
| 0.028231662 | 0.018069385 | 4827 tags=61%, list=27%, signal=45% |
| 0.043249854 | 0.027681625 | 4431 tags=52%, list=25%, signal=39% |
| 0.000859739 | 0.000550267 | 2740 tags=26%, list=15%, signal=23% |
| 0.030286968 | 0.019384863 | 3678 tags=35%, list=20%, signal=28% |
| 0.017922733 | 0.011471261 | 2555 tags=30%, list=14%, signal=26% |
| 0.000852327 | 0.000545523 | 3434 tags=30%, list=19%, signal=24% |
| 0.018020498 | 0.011533835 | 2555 tags=31%, list=14%, signal=26% |
| 0.010160024 | 0.006502819 | 3328 tags=30%, list=19%, signal=25% |
| 0.035620698 | 0.022798662 | 5468 tags=50%, list=30%, signal=35% |
| 0.001432929 | 0.000917131 | 3138 tags=31%, list=17%, signal=26% |
| 0.013416529 | 0.008587112 | 2473 tags=25%, list=14%, signal=21% |
| 0.008769852 | 0.005613054 | 1573 tags=20%, list=9%, signal=19% |
| 0.034774246 | 0.022256899 | 2587 tags=31%, list=14%, signal=27% |
| 0.001144129 | 0.000732288 | 2740 tags=27%, list=15%, signal=23% |
| 0.003769474 | 0.002412613 | 3278 tags=29%, list=18%, signal=24% |
| 0.000906467 | 0.000580175 | 3933 tags=33%, list=22%, signal=26% |
| 0.00045024 | 0.000288171 | 2903 tags=25%, list=16%, signal=21% |
| 0.001075385 | 0.000688289 | 4026 tags=33%, list=22%, signal=26% |
| 0.031901456 | 0.020418199 | 1848 tags=24%, list=10%, signal=21% |
| 0.006424029 | 0.004111634 | 4613 tags=44%, list=26%, signal=33% |
| 2.29899221566784e-05 | 1.47144636818996e-05 | 2162 tags=20%, list=12%, signal=18% |
| 0.000341697 | 0.0002187 | 2740 tags=27%, list=15%, signal=23% |
| 0.038797176 | 0.024831734 | 5857 tags=60%, list=33%, signal=40% |
| 0.033697458 | 0.021567712 | 1996 tags=32%, list=11%, signal=29% |
| 0.001376905 | 0.000881274 | 3838 tags=31%, list=21%, signal=25% |
| 0.046925959 | 0.030034478 | 1572 tags=26%, list=9%, signal=24% |
| 0.008752583 | 0.005602001 | 3376 tags=30%, list=19%, signal=24% |

| | | | |
|---|---|---|---|
| 0.004724196 | 0.003023673 | 3454 | tags=29%, list=19%, signal=23% |
| 0.001107688 | 0.000708964 | 3654 | tags=31%, list=20%, signal=25% |
| 0.031025002 | 0.019857234 | 3862 | tags=41%, list=22%, signal=32% |
| 0.000188077 | 0.000120377 | 2903 | tags=25%, list=16%, signal=21% |
| 0.034231733 | 0.021909669 | 1439 | tags=22%, list=8%, signal=20% |
| 0.005464622 | 0.003497575 | 1205 | tags=22%, list=7%, signal=20% |
| 0.045767287 | 0.029292882 | 4031 | tags=53%, list=22%, signal=41% |
| 0.047985414 | 0.030712571 | 3403 | tags=36%, list=19%, signal=30% |
| 0.005303651 | 0.003394548 | 3826 | tags=30%, list=21%, signal=24% |
| 0.00053068 | 0.000339656 | 4026 | tags=33%, list=22%, signal=26% |
| 0.001379521 | 0.000882948 | 4191 | tags=37%, list=23%, signal=29% |
| 0.002268244 | 0.001451766 | 3732 | tags=34%, list=21%, signal=27% |
| 0.04128572 | 0.026424501 | 3233 | tags=41%, list=18%, signal=34% |
| 0.001106694 | 0.000708328 | 2493 | tags=25%, list=14%, signal=21% |
| 0.001106694 | 0.000708328 | 2493 | tags=25%, list=14%, signal=21% |
| 0.025760829 | 0.016487954 | 1848 | tags=23%, list=10%, signal=21% |
| 0.003359162 | 0.002149997 | 5503 | tags=48%, list=31%, signal=34% |
| 0.002648702 | 0.001695275 | 4133 | tags=37%, list=23%, signal=29% |
| 0.017399635 | 0.011136458 | 3299 | tags=36%, list=18%, signal=29% |
| 0.046834209 | 0.029975755 | 4401 | tags=50%, list=25%, signal=38% |
| 0.013162638 | 0.008424612 | 2939 | tags=29%, list=16%, signal=24% |
| 0.037659644 | 0.024103669 | 3711 | tags=48%, list=21%, signal=38% |
| 0.023209838 | 0.014855218 | 2236 | tags=32%, list=12%, signal=28% |
| 0.003860358 | 0.002470783 | 1757 | tags=21%, list=10%, signal=19% |
| 0.003115692 | 0.001994167 | 3037 | tags=29%, list=17%, signal=25% |
| 0.032430684 | 0.020756926 | 3819 | tags=49%, list=21%, signal=39% |
| 0.017110972 | 0.010951702 | 2827 | tags=28%, list=16%, signal=24% |
| 0.040887992 | 0.026169939 | 4236 | tags=44%, list=24%, signal=34% |
| 0.015290492 | 0.009786522 | 3144 | tags=31%, list=18%, signal=26% |
| 0.034673937 | 0.022192697 | 1988 | tags=32%, list=11%, signal=28% |
| 0.000714122 | 0.000457067 | 3650 | tags=30%, list=20%, signal=24% |
| 0.010270504 | 0.006573531 | 3008 | tags=31%, list=17%, signal=26% |
| 0.000320119 | 0.000204889 | 3826 | tags=31%, list=21%, signal=25% |
| 0.038088112 | 0.024377905 | 1572 | tags=33%, list=9%, signal=30% |
| 0.038955441 | 0.02493303 | 2407 | tags=33%, list=13%, signal=29% |
| 0.012257288 | 0.007845152 | 5222 | tags=53%, list=29%, signal=38% |
| 0.038817054 | 0.024844457 | 2195 | tags=30%, list=12%, signal=27% |
| 0.014946145 | 0.009566126 | 3369 | tags=40%, list=19%, signal=33% |
| 0.04073706 | 0.026073337 | 5503 | tags=55%, list=31%, signal=38% |
| 0.047005583 | 0.030085441 | 2705 | tags=46%, list=15%, signal=39% |
| 0.018805167 | 0.012036054 | 1848 | tags=22%, list=10%, signal=20% |
| 0.018805167 | 0.012036054 | 1848 | tags=22%, list=10%, signal=20% |
| 0.014820792 | 0.009485896 | 3520 | tags=43%, list=20%, signal=35% |
| 0.028100929 | 0.017985711 | 3819 | tags=50%, list=21%, signal=39% |
| 0.01357018 | 0.008685455 | 5222 | tags=52%, list=29%, signal=37% |

| | | | |
|---|---|---|---|
| 0.04529096 | 0.028988014 | 2155 | tags=32%, list=12%, signal=28% |
| 6.34725347280624e-05 | 4.06249442990339e-05 | 4202 | tags=36%, list=23%, signal=28% |
| 0.036254182 | 0.023204117 | 3161 | tags=37%, list=18%, signal=31% |
| 0.036254182 | 0.023204117 | 3161 | tags=37%, list=18%, signal=31% |
| 0.047005583 | 0.030085441 | 1907 | tags=37%, list=11%, signal=33% |
| 0.049144572 | 0.031454479 | 5223 | tags=75%, list=29%, signal=53% |
| 0.014912612 | 0.009544665 | 1848 | tags=22%, list=10%, signal=20% |
| 0.007870802 | 0.005037626 | 2621 | tags=31%, list=15%, signal=26% |
| 0.002042852 | 0.001307506 | 3996 | tags=38%, list=22%, signal=30% |
| 0.049669989 | 0.031790766 | 3971 | tags=54%, list=22%, signal=42% |
| 0.042923759 | 0.027472911 | 2524 | tags=36%, list=14%, signal=31% |
| 0.007173521 | 0.004591338 | 2875 | tags=39%, list=16%, signal=33% |
| 0.004726768 | 0.003025319 | 3831 | tags=38%, list=21%, signal=30% |
| 0.000141032 | 9.02661090454028e-05 | 2881 | tags=27%, list=16%, signal=23% |
| 0.034148475 | 0.021856381 | 3185 | tags=47%, list=18%, signal=39% |
| 0.001347113 | 0.000862206 | 3389 | tags=31%, list=19%, signal=25% |
| 0.047035848 | 0.030104812 | 1451 | tags=39%, list=8%, signal=36% |
| 0.001763435 | 0.001128668 | 2666 | tags=25%, list=15%, signal=21% |
| 0.025717079 | 0.016459953 | 3704 | tags=40%, list=21%, signal=32% |
| 0.045480708 | 0.02910946 | 1478 | tags=53%, list=8%, signal=48% |
| 0.000333789 | 0.000213638 | 3653 | tags=31%, list=20%, signal=25% |
| 0.005500221 | 0.00352036 | 2586 | tags=27%, list=14%, signal=23% |
| 0.049269173 | 0.031534228 | 3885 | tags=58%, list=22%, signal=46% |
| 4.55991869392488e-05 | 2.91852915189979e-05 | 3425 | tags=29%, list=19%, signal=24% |
| 0.03982825 | 0.025491663 | 2151 | tags=38%, list=12%, signal=34% |
| 0.037628449 | 0.024083702 | 3883 | tags=68%, list=22%, signal=53% |
| 0.024998056 | 0.015999749 | 4058 | tags=35%, list=23%, signal=27% |
| 0.048829362 | 0.031252732 | 841 | tags=27%, list=5%, signal=26% |
| 0.041740375 | 0.026715499 | 2375 | tags=43%, list=13%, signal=38% |
| 0.041930454 | 0.026837157 | 2420 | tags=43%, list=13%, signal=37% |
| 0.021003406 | 0.013443014 | 4209 | tags=72%, list=23%, signal=55% |
| 0.020959647 | 0.013415007 | 3242 | tags=30%, list=18%, signal=25% |
| 0.032034541 | 0.020503379 | 1572 | tags=27%, list=9%, signal=25% |
| 0.041157202 | 0.026342245 | 2035 | tags=55%, list=11%, signal=48% |
| 0.041157202 | 0.026342245 | 1714 | tags=36%, list=10%, signal=33% |
| 0.041157202 | 0.026342245 | 1714 | tags=36%, list=10%, signal=33% |
| 0.048211745 | 0.030857432 | 4431 | tags=70%, list=25%, signal=53% |
| 0.027926438 | 0.01787403 | 1015 | tags=28%, list=6%, signal=26% |
| 0.031812084 | 0.020360998 | 2527 | tags=39%, list=14%, signal=34% |
| 0.040703442 | 0.026051821 | 2504 | tags=39%, list=14%, signal=34% |
| 0.017586133 | 0.011255824 | 5093 | tags=42%, list=28%, signal=30% |
| 0.031056084 | 0.019877128 | 5468 | tags=56%, list=30%, signal=39% |
| 0.041930454 | 0.026837157 | 2451 | tags=53%, list=14%, signal=45% |
| 0.008447704 | 0.005406866 | 2265 | tags=28%, list=13%, signal=25% |
| 0.045805449 | 0.029317308 | 1425 | tags=50%, list=8%, signal=46% |

| | | | |
|---|---|---|---|
| 0.021403923 | 0.013699361 | 2111 | tags=26%, list=12%, signal=23% |
| 0.007005941 | 0.004484081 | 2439 | tags=29%, list=14%, signal=25% |
| 0.039942306 | 0.025564663 | 1211 | tags=27%, list=7%, signal=25% |
| 0.039942306 | 0.025564663 | 1211 | tags=27%, list=7%, signal=25% |
| 0.036284127 | 0.023223283 | 3539 | tags=52%, list=20%, signal=42% |
| 0.041930454 | 0.026837157 | 1456 | tags=59%, list=8%, signal=54% |
| 0.041930454 | 0.026837157 | 1456 | tags=59%, list=8%, signal=54% |
| 0.036646339 | 0.023455113 | 4401 | tags=60%, list=25%, signal=45% |
| 0.016969656 | 0.010861254 | 1769 | tags=27%, list=10%, signal=25% |
| 0.013203985 | 0.008451075 | 4026 | tags=44%, list=22%, signal=34% |
| 0.043486863 | 0.02783332 | 976 | tags=57%, list=5%, signal=54% |
| 0.036086852 | 0.023097019 | 2898 | tags=54%, list=16%, signal=45% |
| 0.006350718 | 0.004064712 | 2359 | tags=28%, list=13%, signal=24% |
| 0.039014631 | 0.024970914 | 1367 | tags=35%, list=8%, signal=32% |
| 0.002935749 | 0.001878996 | 2740 | tags=29%, list=15%, signal=25% |
| 0.041727107 | 0.026707006 | 862 | tags=38%, list=5%, signal=36% |
| 0.041727107 | 0.026707006 | 862 | tags=38%, list=5%, signal=36% |
| 0.040318125 | 0.025805202 | 1035 | tags=26%, list=6%, signal=25% |
| 0.012440208 | 0.007962228 | 2504 | tags=27%, list=14%, signal=23% |
| 0.044391643 | 0.028412416 | 2666 | tags=40%, list=15%, signal=34% |
| 0.043347365 | 0.027744036 | 1035 | tags=33%, list=6%, signal=31% |
| 0.03826258 | 0.024489571 | 1358 | tags=29%, list=8%, signal=26% |
| 0.03071871 | 0.019661195 | 1528 | tags=26%, list=9%, signal=24% |
| 0.007638824 | 0.004889151 | 5515 | tags=59%, list=31%, signal=41% |
| 0.042999243 | 0.027521224 | 3494 | tags=60%, list=19%, signal=48% |
| 0.036395911 | 0.023294829 | 2996 | tags=46%, list=17%, signal=38% |
| 0.041273512 | 0.026416688 | 980 | tags=46%, list=5%, signal=44% |
| 0.020243807 | 0.012956841 | 1831 | tags=29%, list=10%, signal=26% |
| 0.041157202 | 0.026342245 | 3010 | tags=54%, list=17%, signal=45% |
| 0.000615316 | 0.000393827 | 2740 | tags=29%, list=15%, signal=25% |
| 0.038341236 | 0.024539915 | 3830 | tags=63%, list=21%, signal=50% |
| 0.036284127 | 0.023223283 | 1714 | tags=35%, list=10%, signal=32% |
| 0.010626332 | 0.006801275 | 5222 | tags=55%, list=29%, signal=39% |
| 0.007394649 | 0.004732869 | 5478 | tags=58%, list=31%, signal=41% |
| 0.040612006 | 0.025993298 | 3816 | tags=64%, list=21%, signal=50% |
| 0.02925323 | 0.01872323 | 3466 | tags=61%, list=19%, signal=50% |
| 0.032864274 | 0.021034441 | 2290 | tags=37%, list=13%, signal=32% |
| 0.010747434 | 0.006878785 | 4026 | tags=44%, list=22%, signal=34% |
| 0.018940377 | 0.012122594 | 3819 | tags=49%, list=21%, signal=39% |
| 0.041157202 | 0.026342245 | 1652 | tags=50%, list=9%, signal=45% |
| 0.041157202 | 0.026342245 | 1652 | tags=50%, list=9%, signal=45% |
| 0.039500381 | 0.025281813 | 4217 | tags=47%, list=23%, signal=36% |
| 0.002694938 | 0.001724867 | 3278 | tags=31%, list=18%, signal=26% |
| 0.005754645 | 0.003683202 | 3434 | tags=33%, list=19%, signal=27% |
| 0.036987359 | 0.02367338 | 3747 | tags=58%, list=21%, signal=46% |

| | | |
|---|---|---|
| 0.005625414 | 0.003600489 | 3434 tags=32%, list=19%, signal=26% |
| 9.74670979920544e-06 | 6.23828155576254e-06 | 3376 tags=29%, list=19%, signal=24% |
| 0.032478659 | 0.020787633 | 5559 tags=64%, list=31%, signal=44% |
| 0.03982825 | 0.025491663 | 1515 tags=50%, list=8%, signal=46% |
| 0.037738534 | 0.024154161 | 3699 tags=71%, list=21%, signal=57% |
| 0.037738534 | 0.024154161 | 3699 tags=71%, list=21%, signal=57% |
| 0.017040732 | 0.010906745 | 2881 tags=33%, list=16%, signal=28% |
| 0.003104981 | 0.001987311 | 3747 tags=35%, list=21%, signal=28% |
| 0.037825254 | 0.024209666 | 5751 tags=75%, list=32%, signal=51% |
| 0.026276972 | 0.016818306 | 1439 tags=26%, list=8%, signal=24% |
| 4.76224732632332e-05 | 3.04802751613719e-05 | 2756 tags=24%, list=15%, signal=21% |
| 0.0169461 | 0.010846178 | 4026 tags=45%, list=22%, signal=35% |
| 0.039057468 | 0.024998331 | 4946 tags=80%, list=28%, signal=58% |
| 0.004677458 | 0.002993759 | 3315 tags=39%, list=18%, signal=32% |
| 0.027159148 | 0.017382934 | 1439 tags=30%, list=8%, signal=28% |
| 0.03826258 | 0.024489571 | 4946 tags=75%, list=28%, signal=54% |
| 0.012306704 | 0.00787678 | 2386 tags=34%, list=13%, signal=29% |
| 0.03826258 | 0.024489571 | 1508 tags=58%, list=8%, signal=53% |
| 0.02826536 | 0.018090954 | 3883 tags=46%, list=22%, signal=36% |
| 0.004618058 | 0.002955741 | 4133 tags=38%, list=23%, signal=29% |
| 0.004618058 | 0.002955741 | 4133 tags=38%, list=23%, signal=29% |
| 0.019775004 | 0.012656788 | 2100 tags=33%, list=12%, signal=30% |
| 0.019482146 | 0.012469347 | 2165 tags=33%, list=12%, signal=29% |
| 0.024309911 | 0.015559309 | 4401 tags=53%, list=25%, signal=40% |
| 0.020623332 | 0.013199752 | 1769 tags=33%, list=10%, signal=30% |
| 0.036661479 | 0.023464803 | 1084 tags=60%, list=6%, signal=56% |
| 0.000513705 | 0.000328791 | 2740 tags=30%, list=15%, signal=25% |
| 0.017237028 | 0.011032383 | 2824 tags=55%, list=16%, signal=47% |
| 0.009874017 | 0.006319763 | 3185 tags=37%, list=18%, signal=31% |
| 0.031222297 | 0.019983511 | 5212 tags=57%, list=29%, signal=40% |
| 0.008112417 | 0.005192269 | 5770 tags=55%, list=32%, signal=37% |
| 0.027776983 | 0.017778373 | 1714 tags=35%, list=10%, signal=31% |
| 0.007996651 | 0.005118175 | 3747 tags=32%, list=21%, signal=26% |
| 0.034788789 | 0.022266207 | 1496 tags=45%, list=8%, signal=42% |
| 0.001275352 | 0.000816276 | 3088 tags=29%, list=17%, signal=24% |
| 0.000992961 | 0.000635534 | 3262 tags=31%, list=18%, signal=26% |
| 0.033450389 | 0.021409578 | 1211 tags=58%, list=7%, signal=54% |
| 0.019528848 | 0.012499239 | 1015 tags=29%, list=6%, signal=27% |
| 0.031853687 | 0.020387625 | 2072 tags=50%, list=12%, signal=44% |
| 0.031853687 | 0.020387625 | 2072 tags=50%, list=12%, signal=44% |
| 0.017508044 | 0.011205844 | 2236 tags=36%, list=12%, signal=31% |
| 0.033151243 | 0.021218112 | 1037 tags=30%, list=6%, signal=28% |
| 0.000600619 | 0.00038442 | 3037 tags=30%, list=17%, signal=25% |
| 0.032713813 | 0.02093814 | 2705 tags=33%, list=15%, signal=28% |
| 0.032765769 | 0.020971394 | 3742 tags=60%, list=21%, signal=48% |

| | | | |
|---|---|---|---|
| 0.015914652 | 0.01018601 | 1674 | tags=24%, list=9%, signal=22% |
| 0.017040732 | 0.010906745 | 4541 | tags=47%, list=25%, signal=35% |
| 0.017237028 | 0.011032383 | 2698 | tags=32%, list=15%, signal=28% |
| 0.023729223 | 0.015187645 | 2332 | tags=36%, list=13%, signal=31% |
| 0.029696845 | 0.019007161 | 1757 | tags=62%, list=10%, signal=56% |
| 0.017040732 | 0.010906745 | 1560 | tags=31%, list=9%, signal=28% |
| 0.028408695 | 0.018182693 | 3440 | tags=50%, list=19%, signal=40% |
| 0.032041464 | 0.02050781 | 2180 | tags=60%, list=12%, signal=53% |
| 0.010994885 | 0.007037163 | 1962 | tags=24%, list=11%, signal=22% |
| 0.030677864 | 0.019635052 | 974 | tags=25%, list=5%, signal=24% |
| 0.030286968 | 0.019384863 | 1515 | tags=47%, list=8%, signal=43% |
| 0.022840259 | 0.014618673 | 1903 | tags=46%, list=11%, signal=42% |
| 0.001775481 | 0.001136379 | 2740 | tags=33%, list=15%, signal=28% |
| 0.016723501 | 0.010703705 | 1266 | tags=26%, list=7%, signal=24% |
| 0.023611703 | 0.015112428 | 3306 | tags=46%, list=18%, signal=38% |
| 0.028781612 | 0.018421376 | 1456 | tags=28%, list=8%, signal=26% |
| 0.000449725 | 0.000287842 | 1693 | tags=19%, list=9%, signal=18% |
| 0.028414292 | 0.018186276 | 1646 | tags=64%, list=9%, signal=58% |
| 0.021246064 | 0.013598325 | 5212 | tags=55%, list=29%, signal=39% |
| 0.00022088 | 0.000141372 | 4633 | tags=38%, list=26%, signal=28% |
| 0.019109789 | 0.012231024 | 3052 | tags=44%, list=17%, signal=37% |
| 0.000873712 | 0.000559211 | 4046 | tags=46%, list=23%, signal=36% |
| 1.67025552534891e-05 | 1.06902990361363e-05 | 3838 | tags=33%, list=21%, signal=26% |
| 0.018685691 | 0.011959585 | 4031 | tags=58%, list=22%, signal=45% |
| 0.018685691 | 0.011959585 | 4031 | tags=58%, list=22%, signal=45% |
| 0.011113899 | 0.007113337 | 1898 | tags=29%, list=11%, signal=26% |
| 0.027614068 | 0.017674101 | 1928 | tags=38%, list=11%, signal=34% |
| 0.008701832 | 0.005569518 | 2373 | tags=30%, list=13%, signal=26% |
| 0.027941082 | 0.017883403 | 1211 | tags=46%, list=7%, signal=43% |
| 0.023611703 | 0.015112428 | 6435 | tags=77%, list=36%, signal=50% |
| 0.022881565 | 0.014645111 | 3785 | tags=56%, list=21%, signal=44% |
| 0.027068724 | 0.017325059 | 3694 | tags=73%, list=21%, signal=58% |
| 0.0257771 | 0.016498368 | 5011 | tags=76%, list=28%, signal=55% |
| 0.026815561 | 0.017163025 | 3631 | tags=60%, list=20%, signal=48% |
| 0.001401131 | 0.00089678 | 3747 | tags=36%, list=21%, signal=28% |
| 0.018216318 | 0.011659167 | 2236 | tags=39%, list=12%, signal=34% |
| 0.025843963 | 0.016541163 | 1084 | tags=45%, list=6%, signal=43% |
| 0.017192019 | 0.011003575 | 3159 | tags=44%, list=18%, signal=36% |
| 2.17582532906418e-05 | 1.39261466674308e-05 | 3826 | tags=34%, list=21%, signal=27% |
| 0.025778321 | 0.016499149 | 2734 | tags=69%, list=15%, signal=59% |
| 0.02562285 | 0.016399642 | 1478 | tags=50%, list=8%, signal=46% |
| 0.000289582 | 0.000185344 | 3037 | tags=31%, list=17%, signal=26% |
| 0.023573172 | 0.015087767 | 2180 | tags=43%, list=12%, signal=38% |
| 0.022809851 | 0.014599211 | 1959 | tags=35%, list=11%, signal=31% |
| 0.000844667 | 0.000540621 | 2740 | tags=32%, list=15%, signal=27% |

| | | | |
|---|---|---|---|
| 5.73418584109666e-06 | 3.67010678544509e-06 | 4172 | tags=37%, list=23%, signal=29% |
| 0.0169461 | 0.010846178 | 2072 | tags=35%, list=12%, signal=31% |
| 0.017361703 | 0.01111218 | 2527 | tags=39%, list=14%, signal=34% |
| 0.023408764 | 0.014982539 | 1084 | tags=42%, list=6%, signal=39% |
| 3.39600321226471e-06 | 2.17357699560402e-06 | 3315 | tags=30%, list=18%, signal=25% |
| 0.000452446 | 0.000289583 | 3885 | tags=38%, list=22%, signal=30% |
| 0.022367952 | 0.014316378 | 860 | tags=55%, list=5%, signal=52% |
| 4.86624896207902e-07 | 3.11459269551258e-07 | 3658 | tags=32%, list=20%, signal=26% |
| 0.001291777 | 0.000826788 | 2439 | tags=28%, list=14%, signal=25% |
| 0.022471028 | 0.014382351 | 1347 | tags=38%, list=8%, signal=36% |
| 0.017000482 | 0.010880984 | 3058 | tags=41%, list=17%, signal=34% |
| 0.006965793 | 0.004458384 | 2766 | tags=30%, list=15%, signal=25% |
| 6.42095020892566e-06 | 4.10966326935023e-06 | 2175 | tags=32%, list=12%, signal=29% |
| 0.020467495 | 0.01310001 | 2332 | tags=43%, list=13%, signal=37% |
| 0.021163429 | 0.013545435 | 2890 | tags=70%, list=16%, signal=59% |
| 0.014068183 | 0.009004196 | 3037 | tags=47%, list=17%, signal=39% |
| 0.021395625 | 0.01369405 | 3699 | tags=69%, list=21%, signal=55% |
| 1.94800330051616e-05 | 1.24679951599309e-05 | 1693 | tags=19%, list=9%, signal=18% |
| 0.021246064 | 0.013598325 | 2662 | tags=64%, list=15%, signal=54% |
| 0.016755536 | 0.010724209 | 2206 | tags=30%, list=12%, signal=26% |
| 0.020772006 | 0.013294909 | 1831 | tags=50%, list=10%, signal=45% |
| 0.019728554 | 0.012627059 | 1108 | tags=48%, list=6%, signal=45% |
| 0.019036171 | 0.012183906 | 2842 | tags=50%, list=16%, signal=42% |
| 0.019036171 | 0.012183906 | 2842 | tags=50%, list=16%, signal=42% |
| 0.021005195 | 0.01344416 | 1652 | tags=46%, list=9%, signal=42% |
| 0.013271862 | 0.008494519 | 1820 | tags=28%, list=10%, signal=25% |
| 0.013395358 | 0.008573561 | 4285 | tags=59%, list=24%, signal=45% |
| 0.010635251 | 0.006806983 | 2009 | tags=27%, list=11%, signal=24% |
| 0.02002957 | 0.012819721 | 1244 | tags=38%, list=7%, signal=35% |
| 0.001294795 | 0.000828721 | 4106 | tags=38%, list=23%, signal=29% |
| 0.008785187 | 0.005622869 | 2265 | tags=41%, list=13%, signal=36% |
| 0.001094999 | 0.000700843 | 2895 | tags=31%, list=16%, signal=26% |
| 7.47963910364077e-05 | 4.78726622883614e-05 | 3152 | tags=29%, list=18%, signal=24% |
| 0.012760024 | 0.008166922 | 1456 | tags=33%, list=8%, signal=31% |
| 0.000762402 | 0.000487968 | 4578 | tags=46%, list=25%, signal=35% |
| 0.01921373 | 0.012297551 | 2848 | tags=75%, list=16%, signal=63% |
| 0.009568943 | 0.006124503 | 1825 | tags=35%, list=10%, signal=32% |
| 7.18903082828122e-06 | 4.60126538532326e-06 | 2697 | tags=34%, list=15%, signal=29% |
| 0.017922733 | 0.011471261 | 1652 | tags=41%, list=9%, signal=37% |
| 0.006781063 | 0.00434015 | 3037 | tags=40%, list=17%, signal=33% |
| 0.005608718 | 0.003589802 | 5207 | tags=62%, list=29%, signal=44% |
| 0.000476937 | 0.000305259 | 4176 | tags=43%, list=23%, signal=33% |
| 0.015053347 | 0.00963474 | 3102 | tags=36%, list=17%, signal=30% |
| 0.017101117 | 0.010945394 | 3792 | tags=72%, list=21%, signal=57% |
| 0.016589761 | 0.010618106 | 3056 | tags=50%, list=17%, signal=42% |

| | | | |
|---|---|---|---|
| 8.40218974238139e-06 | 5.37773529506213e-06 | 3826 | tags=35%, list=21%, signal=28% |
| 0.016864757 | 0.010794115 | 1508 | tags=82%, list=8%, signal=75% |
| 0.017040732 | 0.010906745 | 5032 | tags=73%, list=28%, signal=53% |
| 0.017110972 | 0.010951702 | 2438 | tags=50%, list=14%, signal=43% |
| 0.002529524 | 0.001618996 | 1211 | tags=23%, list=7%, signal=22% |
| 0.002529524 | 0.001618996 | 1211 | tags=23%, list=7%, signal=22% |
| 0.012613557 | 0.008073178 | 3298 | tags=48%, list=18%, signal=40% |
| 0.017192019 | 0.011003575 | 1515 | tags=60%, list=8%, signal=55% |
| 0.015249193 | 0.009760089 | 3699 | tags=60%, list=21%, signal=48% |
| 0.016679207 | 0.010675356 | 710 | tags=42%, list=4%, signal=40% |
| 0.016679207 | 0.010675356 | 710 | tags=42%, list=4%, signal=40% |
| 1.29636640325847e-05 | 8.29726009039154e-06 | 3152 | tags=29%, list=18%, signal=25% |
| 0.007353151 | 0.004706309 | 1887 | tags=33%, list=11%, signal=29% |
| 0.003978696 | 0.002546523 | 3742 | tags=46%, list=21%, signal=37% |
| 0.014690771 | 0.009402677 | 2634 | tags=48%, list=15%, signal=41% |
| 0.014912612 | 0.009544665 | 1211 | tags=55%, list=7%, signal=51% |
| 0.01151515 | 0.007370154 | 3220 | tags=55%, list=18%, signal=45% |
| 0.01466243 | 0.009384538 | 2586 | tags=64%, list=14%, signal=55% |
| 9.7644720531788e-06 | 6.24965010408639e-06 | 3315 | tags=34%, list=18%, signal=28% |
| 0.015552287 | 0.009954082 | 1244 | tags=50%, list=7%, signal=47% |
| 0.002929984 | 0.001875306 | 2890 | tags=36%, list=16%, signal=30% |
| 0.014256013 | 0.009124415 | 2842 | tags=64%, list=16%, signal=54% |
| 0.014193559 | 0.009084442 | 2371 | tags=45%, list=13%, signal=39% |
| 0.013592908 | 0.008700001 | 2634 | tags=50%, list=15%, signal=43% |
| 0.012813836 | 0.008201364 | 3112 | tags=52%, list=17%, signal=43% |
| 1.21171993888846e-05 | 7.75548908425879e-06 | 2666 | tags=27%, list=15%, signal=23% |
| 0.01333528 | 0.008535109 | 891 | tags=29%, list=5%, signal=27% |
| 0.00540826 | 0.003461501 | 2883 | tags=43%, list=16%, signal=36% |
| 0.011553432 | 0.007394656 | 4539 | tags=79%, list=25%, signal=59% |
| 0.011031195 | 0.007060403 | 2438 | tags=53%, list=14%, signal=46% |
| 0.005582042 | 0.003572729 | 1848 | tags=27%, list=10%, signal=25% |
| 0.005582042 | 0.003572729 | 1848 | tags=27%, list=10%, signal=25% |
| 0.005582042 | 0.003572729 | 1848 | tags=27%, list=10%, signal=25% |
| 0.012615778 | 0.008074599 | 1652 | tags=45%, list=9%, signal=41% |
| 3.98995591158842e-07 | 2.55373032380597e-07 | 3235 | tags=29%, list=18%, signal=24% |
| 0.014691053 | 0.009402858 | 710 | tags=38%, list=4%, signal=37% |
| 0.014691053 | 0.009402858 | 710 | tags=38%, list=4%, signal=37% |
| 0.014879459 | 0.009523445 | 1373 | tags=75%, list=8%, signal=69% |
| 0.014256013 | 0.009124415 | 1415 | tags=50%, list=8%, signal=46% |
| 0.007662603 | 0.00490437 | 1221 | tags=33%, list=7%, signal=31% |
| 0.010571875 | 0.00676642 | 4023 | tags=75%, list=22%, signal=58% |
| 0.008447704 | 0.005406866 | 3328 | tags=39%, list=19%, signal=32% |
| 0.000892817 | 0.000571438 | 3747 | tags=33%, list=21%, signal=27% |
| 0.009341415 | 0.005978877 | 3247 | tags=32%, list=18%, signal=26% |
| 0.001395134 | 0.000892941 | 2740 | tags=35%, list=15%, signal=30% |

| | | | |
|---|---|---|---|
| 0.009699227 | 0.006207891 | 3129 | tags=56%, list=17%, signal=46% |
| 0.01308593 | 0.008375515 | 1496 | tags=38%, list=8%, signal=34% |
| 0.008921839 | 0.005710332 | 2072 | tags=61%, list=12%, signal=54% |
| 0.001889359 | 0.001209265 | 3144 | tags=37%, list=18%, signal=31% |
| 0.009341415 | 0.005978877 | 1266 | tags=30%, list=7%, signal=28% |
| 0.009341415 | 0.005978877 | 1451 | tags=37%, list=8%, signal=34% |
| 0.001750047 | 0.0011201 | 2890 | tags=35%, list=16%, signal=30% |
| 0.005840344 | 0.003738052 | 5515 | tags=62%, list=31%, signal=43% |
| 0.007540399 | 0.004826155 | 1825 | tags=38%, list=10%, signal=34% |
| 7.66258179991838e-09 | 4.90435682365852e-09 | 4140 | tags=37%, list=23%, signal=29% |
| 0.012312568 | 0.007880533 | 1456 | tags=62%, list=8%, signal=57% |
| 0.000169166 | 0.000108273 | 4140 | tags=41%, list=23%, signal=32% |
| 0.00440778 | 0.002821154 | 1110 | tags=36%, list=6%, signal=34% |
| 0.003304514 | 0.00211502 | 2766 | tags=31%, list=15%, signal=26% |
| 0.003304514 | 0.00211502 | 2766 | tags=31%, list=15%, signal=26% |
| 0.011060338 | 0.007079056 | 1084 | tags=70%, list=6%, signal=66% |
| 0.001750047 | 0.0011201 | 2634 | tags=32%, list=15%, signal=28% |
| 8.92841378326314e-06 | 5.71453959067161e-06 | 4052 | tags=42%, list=23%, signal=33% |
| 0.000625879 | 0.000400588 | 1996 | tags=27%, list=11%, signal=24% |
| 0.01048702 | 0.006712109 | 2527 | tags=50%, list=14%, signal=43% |
| 0.007720355 | 0.004941334 | 3220 | tags=52%, list=18%, signal=42% |
| 0.010816056 | 0.006922705 | 662 | tags=35%, list=4%, signal=34% |
| 1.7523011893503e-06 | 1.12154238864851e-06 | 3328 | tags=34%, list=19%, signal=29% |
| 0.003934145 | 0.002518009 | 2236 | tags=37%, list=12%, signal=32% |
| 0.0094406 | 0.006042359 | 3023 | tags=50%, list=17%, signal=42% |
| 0.0094406 | 0.006042359 | 3023 | tags=50%, list=17%, signal=42% |
| 0.000958415 | 0.000613424 | 4172 | tags=46%, list=23%, signal=36% |
| 0.010174537 | 0.006512108 | 3010 | tags=70%, list=17%, signal=58% |
| 9.66727241165724e-05 | 6.18743847129723e-05 | 4140 | tags=41%, list=23%, signal=32% |
| 9.66727241165724e-05 | 6.18743847129723e-05 | 4140 | tags=41%, list=23%, signal=32% |
| 2.18762957871108e-05 | 1.40016985555746e-05 | 2194 | tags=23%, list=12%, signal=20% |
| 0.006338037 | 0.004056596 | 3220 | tags=49%, list=18%, signal=40% |
| 0.010643138 | 0.006812031 | 2231 | tags=62%, list=12%, signal=54% |
| 0.008117717 | 0.005195661 | 3454 | tags=38%, list=19%, signal=31% |
| 1.54773692433393e-06 | 9.9061313070305e-07 | 1693 | tags=20%, list=9%, signal=19% |
| 0.007741597 | 0.00495493 | 2072 | tags=60%, list=12%, signal=53% |
| 0.000557982 | 0.000357131 | 2239 | tags=28%, list=12%, signal=25% |
| 0.004868921 | 0.003116303 | 1103 | tags=30%, list=6%, signal=28% |
| 0.010276906 | 0.006577628 | 709 | tags=38%, list=4%, signal=36% |
| 0.010276906 | 0.006577628 | 1035 | tags=31%, list=6%, signal=29% |
| 0.000270911 | 0.000173394 | 2727 | tags=29%, list=15%, signal=25% |
| 0.008464175 | 0.005417408 | 2040 | tags=50%, list=11%, signal=44% |
| 0.010475423 | 0.006704687 | 1506 | tags=75%, list=8%, signal=69% |
| 0.008328306 | 0.005330447 | 3529 | tags=48%, list=20%, signal=38% |
| 0.008328306 | 0.005330447 | 3529 | tags=48%, list=20%, signal=38% |

| | | | |
|---|---|---|---|
| 0.008883868 | 0.005686028 | 1595 | tags=38%, list=9%, signal=35% |
| 0.005122792 | 0.00327879 | 2371 | tags=37%, list=13%, signal=32% |
| 0.010221381 | 0.00654209 | 1831 | tags=58%, list=10%, signal=52% |
| 0.010221381 | 0.00654209 | 1831 | tags=58%, list=10%, signal=52% |
| 4.57412427643191e-05 | 2.9276212891613e-05 | 3129 | tags=36%, list=17%, signal=30% |
| 0.007339153 | 0.004697349 | 3037 | tags=54%, list=17%, signal=45% |
| 0.006979675 | 0.004467269 | 2504 | tags=40%, list=14%, signal=34% |
| 0.009759269 | 0.00624632 | 2057 | tags=69%, list=11%, signal=61% |
| 0.009620985 | 0.006157813 | 2023 | tags=54%, list=11%, signal=48% |
| 0.009620985 | 0.006157813 | 2023 | tags=54%, list=11%, signal=48% |
| 0.008512072 | 0.005448064 | 860 | tags=50%, list=5%, signal=48% |
| 0.008446878 | 0.005406337 | 1646 | tags=45%, list=9%, signal=41% |
| 0.008446878 | 0.005406337 | 1646 | tags=45%, list=9%, signal=41% |
| 0.000166986 | 0.000106878 | 3144 | tags=33%, list=18%, signal=28% |
| 0.000399987 | 0.000256008 | 2666 | tags=30%, list=15%, signal=26% |
| 0.004981019 | 0.00318805 | 1988 | tags=37%, list=11%, signal=33% |
| 0.009330545 | 0.00597192 | 2939 | tags=44%, list=16%, signal=37% |
| 0.009330545 | 0.00597192 | 2939 | tags=44%, list=16%, signal=37% |
| 0.000417885 | 0.000267463 | 3104 | tags=36%, list=17%, signal=30% |
| 0.00826734 | 0.005291426 | 891 | tags=30%, list=5%, signal=29% |
| 0.004690415 | 0.003002052 | 1451 | tags=50%, list=8%, signal=46% |
| 0.008447704 | 0.005406866 | 2662 | tags=44%, list=15%, signal=38% |
| 0.008447704 | 0.005406866 | 2662 | tags=44%, list=15%, signal=38% |
| 3.61516488062434e-06 | 2.31384917171514e-06 | 2265 | tags=24%, list=13%, signal=21% |
| 0.009091015 | 0.005818611 | 3138 | tags=71%, list=17%, signal=59% |
| 0.007097306 | 0.004542558 | 4247 | tags=65%, list=24%, signal=50% |
| 0.004862378 | 0.003112115 | 2072 | tags=36%, list=12%, signal=32% |
| 0.006816398 | 0.004362765 | 3276 | tags=57%, list=18%, signal=46% |
| 0.002099569 | 0.001343807 | 2265 | tags=34%, list=13%, signal=30% |
| 0.007870802 | 0.005037626 | 1084 | tags=64%, list=6%, signal=60% |
| 0.008769852 | 0.005613054 | 1496 | tags=43%, list=8%, signal=39% |
| 0.005788196 | 0.003704675 | 4539 | tags=59%, list=25%, signal=44% |
| 0.006129558 | 0.003923161 | 1211 | tags=37%, list=7%, signal=35% |
| 0.000951757 | 0.000609162 | 3328 | tags=37%, list=19%, signal=31% |
| 0.006700213 | 0.004288402 | 2364 | tags=41%, list=13%, signal=36% |
| 0.008938203 | 0.005720805 | 2364 | tags=50%, list=13%, signal=43% |
| 0.005586598 | 0.003575645 | 3722 | tags=59%, list=21%, signal=47% |
| 0.008319087 | 0.005324546 | 1084 | tags=50%, list=6%, signal=47% |
| 1.09115981038366e-05 | 6.9838563574147e-06 | 3006 | tags=33%, list=17%, signal=27% |
| 0.00826734 | 0.005291426 | 1515 | tags=50%, list=8%, signal=46% |
| 0.007262326 | 0.004648177 | 922 | tags=60%, list=5%, signal=57% |
| 0.006659175 | 0.004262137 | 1506 | tags=50%, list=8%, signal=46% |
| 0.004736692 | 0.003031671 | 3276 | tags=48%, list=18%, signal=40% |
| 0.008254972 | 0.00528351 | 3699 | tags=83%, list=21%, signal=66% |
| 0.005203434 | 0.003330405 | 3093 | tags=47%, list=17%, signal=39% |

| | | | |
|---|---|---|---|
| 0.000122583 | 7.84577767672112e-05 | 3144 | tags=34%, list=18%, signal=29% |
| 0.006659175 | 0.004262137 | 3282 | tags=80%, list=18%, signal=65% |
| 7.41013485150123e-09 | 4.74278074572423e-09 | 2176 | tags=23%, list=12%, signal=21% |
| 0.008009049 | 0.00512611 | 709 | tags=40%, list=4%, signal=38% |
| 0.007157294 | 0.004580953 | 2587 | tags=55%, list=14%, signal=47% |
| 0.002042852 | 0.001307506 | 2265 | tags=42%, list=13%, signal=37% |
| 0.005303651 | 0.003394548 | 1508 | tags=42%, list=8%, signal=39% |
| 0.002119027 | 0.001356262 | 1848 | tags=29%, list=10%, signal=26% |
| 0.005242339 | 0.003355305 | 2827 | tags=38%, list=16%, signal=32% |
| 0.007734282 | 0.004950248 | 3228 | tags=83%, list=18%, signal=68% |
| 0.000503241 | 0.000322094 | 1832 | tags=24%, list=10%, signal=22% |
| 1.50217516662332e-05 | 9.61451795377626e-06 | 3006 | tags=33%, list=17%, signal=28% |
| 9.08328365339242e-06 | 5.81366246128889e-06 | 2686 | tags=30%, list=15%, signal=25% |
| 0.000388095 | 0.000248396 | 3185 | tags=38%, list=18%, signal=31% |
| 0.001151033 | 0.000736707 | 3742 | tags=45%, list=21%, signal=36% |
| 0.002042852 | 0.001307506 | 1988 | tags=32%, list=11%, signal=29% |
| 0.002042852 | 0.001307506 | 1988 | tags=32%, list=11%, signal=29% |
| 0.001724227 | 0.001103574 | 2329 | tags=29%, list=13%, signal=25% |
| 0.007377973 | 0.004722196 | 922 | tags=60%, list=5%, signal=57% |
| 0.000110821 | 7.09300389731424e-05 | 2727 | tags=30%, list=15%, signal=25% |
| 3.70716043097216e-06 | 2.3727299793692e-06 | 2666 | tags=28%, list=15%, signal=24% |
| 0.000940222 | 0.00060178 | 2634 | tags=33%, list=15%, signal=29% |
| 4.57412427643191e-05 | 2.9276212891613e-05 | 2688 | tags=28%, list=15%, signal=24% |
| 0.000704271 | 0.000450761 | 2165 | tags=31%, list=12%, signal=28% |
| 0.006597544 | 0.00422269 | 2162 | tags=67%, list=12%, signal=59% |
| 0.005252361 | 0.00336172 | 4804 | tags=76%, list=27%, signal=56% |
| 0.001733833 | 0.001109722 | 1903 | tags=33%, list=11%, signal=29% |
| 0.00445194 | 0.002849419 | 3190 | tags=54%, list=18%, signal=45% |
| 4.83497965217062e-06 | 3.09457909468916e-06 | 3006 | tags=33%, list=17%, signal=27% |
| 0.006659175 | 0.004262137 | 3228 | tags=71%, list=18%, signal=58% |
| 0.005149202 | 0.003295694 | 964 | tags=57%, list=5%, signal=54% |
| 0.00191909 | 0.001228294 | 1714 | tags=27%, list=10%, signal=25% |
| 0.004843829 | 0.003100243 | 1211 | tags=29%, list=7%, signal=27% |
| 0.005809685 | 0.003718429 | 1174 | tags=73%, list=7%, signal=68% |
| 0.001590639 | 0.001018072 | 1451 | tags=30%, list=8%, signal=27% |
| 0.002999687 | 0.001919919 | 2072 | tags=37%, list=12%, signal=33% |
| 0.000609486 | 0.000390096 | 3435 | tags=41%, list=19%, signal=34% |
| 0.005657118 | 0.00362078 | 1451 | tags=55%, list=8%, signal=50% |
| 0.000594575 | 0.000380552 | 2560 | tags=33%, list=14%, signal=28% |
| 0.005252361 | 0.00336172 | 1262 | tags=37%, list=7%, signal=34% |
| 0.001679543 | 0.001074974 | 3093 | tags=42%, list=17%, signal=35% |
| 0.006443119 | 0.004123852 | 3862 | tags=71%, list=22%, signal=55% |
| 5.05448985749289e-05 | 3.23507434830543e-05 | 4176 | tags=46%, list=23%, signal=35% |
| 0.005221709 | 0.003342101 | 597 | tags=55%, list=3%, signal=53% |
| 0.002158439 | 0.001381487 | 2023 | tags=54%, list=11%, signal=48% |

| | | | |
|---|---|---|---|
| 8.57636192113451e-07 | 5.48921241017344e-07 | 2935 | tags=30%, list=16%, signal=25% |
| 8.27515259860716e-07 | 5.29642647524191e-07 | 3328 | tags=36%, list=19%, signal=29% |
| 0.006211693 | 0.00397573 | 4588 | tags=65%, list=26%, signal=48% |
| 0.002889037 | 0.001849098 | 1515 | tags=33%, list=8%, signal=31% |
| 4.32536390572985e-05 | 2.76840476745037e-05 | 2688 | tags=29%, list=15%, signal=25% |
| 0.004871592 | 0.003118012 | 3992 | tags=69%, list=22%, signal=54% |
| 0.000784991 | 0.000502426 | 3093 | tags=41%, list=17%, signal=34% |
| 0.004630087 | 0.00296344 | 2305 | tags=55%, list=13%, signal=48% |
| 0.00474199 | 0.003035062 | 2206 | tags=67%, list=12%, signal=59% |
| 0.000483634 | 0.000309545 | 2634 | tags=34%, list=15%, signal=29% |
| 0.004397951 | 0.002814864 | 1463 | tags=42%, list=8%, signal=39% |
| 0.003516325 | 0.002250588 | 2611 | tags=80%, list=15%, signal=68% |
| 0.001858163 | 0.001189298 | 3220 | tags=57%, list=18%, signal=47% |
| 0.001379249 | 0.000882774 | 2666 | tags=44%, list=15%, signal=38% |
| 0.001379249 | 0.000882774 | 2666 | tags=44%, list=15%, signal=38% |
| 0.000391035 | 0.000250278 | 2371 | tags=35%, list=13%, signal=31% |
| 0.000301775 | 0.000193148 | 3144 | tags=38%, list=18%, signal=32% |
| 0.000183457 | 0.00011742 | 2461 | tags=33%, list=14%, signal=29% |
| 0.001753713 | 0.001122446 | 2110 | tags=31%, list=12%, signal=28% |
| 0.001335441 | 0.000854735 | 1103 | tags=21%, list=6%, signal=20% |
| 0.0025753 | 0.001648294 | 4588 | tags=52%, list=26%, signal=39% |
| 9.9103355680017e-05 | 6.3430085494412e-05 | 2072 | tags=32%, list=12%, signal=29% |
| 0.003586723 | 0.002295645 | 4539 | tags=79%, list=25%, signal=59% |
| 0.0051755 | 0.003312526 | 1266 | tags=35%, list=7%, signal=33% |
| 0.004315137 | 0.002761859 | 2329 | tags=69%, list=13%, signal=60% |
| 0.004257331 | 0.002724861 | 3233 | tags=58%, list=18%, signal=48% |
| 0.003435212 | 0.002198672 | 3221 | tags=70%, list=18%, signal=57% |
| 0.001965554 | 0.001258033 | 4116 | tags=56%, list=23%, signal=44% |
| 0.001678882 | 0.001074551 | 1907 | tags=31%, list=11%, signal=28% |
| 0.003860358 | 0.002470783 | 3010 | tags=57%, list=17%, signal=48% |
| 0.003029619 | 0.001939077 | 3190 | tags=52%, list=18%, signal=43% |
| 0.004887344 | 0.003128095 | 2634 | tags=59%, list=15%, signal=50% |
| 0.000647351 | 0.00041433 | 2217 | tags=45%, list=12%, signal=40% |
| 0.001628672 | 0.001042415 | 3185 | tags=50%, list=18%, signal=41% |
| 5.26552830569413e-05 | 3.37014734074006e-05 | 2727 | tags=31%, list=15%, signal=26% |
| 0.004255038 | 0.002723393 | 1125 | tags=47%, list=6%, signal=44% |
| 4.38616008564074e-05 | 2.80731673832179e-05 | 2727 | tags=30%, list=15%, signal=26% |
| 1.46652641495996e-05 | 9.38635177814476e-06 | 2827 | tags=30%, list=16%, signal=25% |
| 1.17155913612433e-05 | 7.49844398872459e-06 | 4425 | tags=50%, list=25%, signal=38% |
| 0.000246281 | 0.00015763 | 5479 | tags=57%, list=31%, signal=39% |
| 1.94401320974601e-05 | 1.24424569935445e-05 | 4425 | tags=51%, list=25%, signal=38% |
| 2.85585192924556e-05 | 1.82785871162941e-05 | 3185 | tags=38%, list=18%, signal=31% |
| 0.000913723 | 0.000584819 | 3785 | tags=61%, list=21%, signal=48% |
| 0.003473655 | 0.002223277 | 4588 | tags=60%, list=26%, signal=45% |
| 0.002158439 | 0.001381487 | 2023 | tags=42%, list=11%, signal=38% |

| | | | |
|---|---|---|---|
| 0.00181975 | 0.001164712 | 1515 | tags=33%, list=8%, signal=31% |
| 0.000912068 | 0.00058376 | 3129 | tags=46%, list=17%, signal=38% |
| 0.003158786 | 0.002021749 | 3121 | tags=67%, list=17%, signal=55% |
| 0.00181975 | 0.001164712 | 3023 | tags=50%, list=17%, signal=42% |
| 0.001944204 | 0.001244368 | 1907 | tags=35%, list=11%, signal=31% |
| 0.003147506 | 0.002014529 | 1364 | tags=43%, list=8%, signal=40% |
| 0.003003391 | 0.001922289 | 2842 | tags=54%, list=16%, signal=45% |
| 7.93263673289051e-05 | 5.07720271135901e-05 | 2661 | tags=34%, list=15%, signal=29% |
| 0.002002889 | 0.001281928 | 2072 | tags=41%, list=12%, signal=36% |
| 0.002635395 | 0.001686758 | 1468 | tags=42%, list=8%, signal=39% |
| 0.000821414 | 0.000525738 | 2082 | tags=38%, list=12%, signal=34% |
| 9.53478938335603e-07 | 6.10264406898821e-07 | 2175 | tags=25%, list=12%, signal=23% |
| 0.002314305 | 0.001481247 | 2705 | tags=46%, list=15%, signal=39% |
| 0.002769032 | 0.001772291 | 964 | tags=38%, list=5%, signal=36% |
| 0.002694306 | 0.001724463 | 2175 | tags=47%, list=12%, signal=41% |
| 0.002314305 | 0.001481247 | 1508 | tags=64%, list=8%, signal=58% |
| 0.002561406 | 0.001639402 | 1652 | tags=40%, list=9%, signal=36% |
| 0.002314305 | 0.001481247 | 1373 | tags=73%, list=8%, signal=67% |
| 5.24360328720285e-07 | 3.35611445771783e-07 | 2827 | tags=31%, list=16%, signal=27% |
| 0.00223353 | 0.001429548 | 860 | tags=60%, list=5%, signal=57% |
| 0.000779912 | 0.000499175 | 3328 | tags=39%, list=19%, signal=32% |
| 0.002566666 | 0.001642768 | 1848 | tags=72%, list=10%, signal=65% |
| 0.002233324 | 0.001429416 | 700 | tags=33%, list=4%, signal=32% |
| 8.11536796619838e-05 | 5.1941579614783e-05 | 2165 | tags=32%, list=12%, signal=29% |
| 0.002385478 | 0.001526801 | 2611 | tags=75%, list=15%, signal=64% |
| 0.000260552 | 0.000166764 | 1496 | tags=32%, list=8%, signal=30% |
| 0.000273651 | 0.000175148 | 1697 | tags=31%, list=9%, signal=28% |
| 0.002004791 | 0.001283146 | 1368 | tags=41%, list=8%, signal=38% |
| 0.002153923 | 0.001378597 | 2162 | tags=69%, list=12%, signal=61% |
| 0.000242283 | 0.000155071 | 1903 | tags=37%, list=11%, signal=33% |
| 4.85719740293471e-05 | 3.10879933799768e-05 | 2661 | tags=33%, list=15%, signal=29% |
| 2.80883741702514e-06 | 1.7977675556927e-06 | 3826 | tags=37%, list=21%, signal=29% |
| 0.001753713 | 0.001122446 | 1496 | tags=58%, list=8%, signal=54% |
| 0.002158439 | 0.001381487 | 2611 | tags=73%, list=15%, signal=63% |
| 0.00150326 | 0.000962146 | 2111 | tags=42%, list=12%, signal=38% |
| 0.000712908 | 0.00045629 | 3862 | tags=60%, list=22%, signal=47% |
| 4.69048577019445e-07 | 3.00209726877843e-07 | 2275 | tags=29%, list=13%, signal=26% |
| 0.002158439 | 0.001381487 | 1015 | tags=60%, list=6%, signal=57% |
| 0.001944204 | 0.001244368 | 1858 | tags=53%, list=10%, signal=47% |
| 0.00165534 | 0.001059484 | 2681 | tags=43%, list=15%, signal=37% |
| 0.001864492 | 0.001193349 | 2305 | tags=61%, list=13%, signal=53% |
| 0.001323691 | 0.000847215 | 1831 | tags=36%, list=10%, signal=32% |
| 0.001323691 | 0.000847215 | 1831 | tags=36%, list=10%, signal=32% |
| 5.90616137779339e-05 | 3.78017796235701e-05 | 5493 | tags=64%, list=31%, signal=45% |
| 0.000326155 | 0.000208752 | 1697 | tags=30%, list=9%, signal=28% |

| | | | |
|---|---|---|---|
| 0.000226455 | 0.00014494 | 1697 | tags=32%, list=9%, signal=29% |
| 0.0009004 | 0.000576292 | 1988 | tags=64%, list=11%, signal=57% |
| 0.0009004 | 0.000576292 | 1988 | tags=64%, list=11%, signal=57% |
| 0.001763435 | 0.001128668 | 1953 | tags=42%, list=11%, signal=38% |
| 0.000530566 | 0.000339583 | 4062 | tags=55%, list=23%, signal=43% |
| 0.000699475 | 0.000447692 | 3321 | tags=62%, list=18%, signal=51% |
| 2.65771777078408e-08 | 1.70104497737855e-08 | 2629 | tags=30%, list=15%, signal=26% |
| 0.00079229 | 0.000507097 | 1431 | tags=55%, list=8%, signal=50% |
| 0.000893628 | 0.000571957 | 2555 | tags=54%, list=14%, signal=46% |
| 0.001763435 | 0.001128668 | 3747 | tags=68%, list=21%, signal=54% |
| 0.000465274 | 0.000297794 | 1515 | tags=58%, list=8%, signal=53% |
| 0.001440706 | 0.000922109 | 3262 | tags=62%, list=18%, signal=50% |
| 2.28536422740712e-08 | 1.46272391419672e-08 | 2329 | tags=27%, list=13%, signal=24% |
| 0.001523597 | 0.000975163 | 1926 | tags=44%, list=11%, signal=39% |
| 0.001073489 | 0.000687076 | 597 | tags=54%, list=3%, signal=52% |
| 0.002042852 | 0.001307506 | 964 | tags=39%, list=5%, signal=37% |
| 0.000645933 | 0.000413423 | 1508 | tags=64%, list=8%, signal=59% |
| 0.000176248 | 0.000112806 | 3392 | tags=32%, list=19%, signal=26% |
| 1.53011429586336e-05 | 9.7933394824899e-06 | 3170 | tags=36%, list=18%, signal=30% |
| 6.1055181520492e-05 | 3.90777421929673e-05 | 2165 | tags=35%, list=12%, signal=31% |
| 0.001195751 | 0.000765328 | 1714 | tags=52%, list=10%, signal=47% |
| 0.001263356 | 0.000808598 | 2072 | tags=46%, list=12%, signal=41% |
| 0.000931358 | 0.000596106 | 2705 | tags=62%, list=15%, signal=52% |
| 0.000931358 | 0.000596106 | 2827 | tags=62%, list=16%, signal=52% |
| 0.000931358 | 0.000596106 | 2827 | tags=62%, list=16%, signal=52% |
| 0.000602008 | 0.000385309 | 1496 | tags=43%, list=8%, signal=40% |
| 0.001515909 | 0.000970242 | 2111 | tags=59%, list=12%, signal=52% |
| 0.001174881 | 0.00075197 | 4222 | tags=67%, list=24%, signal=51% |
| 0.00056557 | 0.000361987 | 1158 | tags=64%, list=6%, signal=60% |
| 0.000594627 | 0.000380585 | 662 | tags=70%, list=4%, signal=67% |
| 0.000552839 | 0.000353839 | 1084 | tags=55%, list=6%, signal=51% |
| 0.001675228 | 0.001072212 | 2460 | tags=67%, list=14%, signal=58% |
| 0.000598854 | 0.000383291 | 2414 | tags=80%, list=13%, signal=69% |
| 0.001009893 | 0.000646372 | 1496 | tags=39%, list=8%, signal=36% |
| 3.86359117715591e-08 | 2.47285187268256e-08 | 2275 | tags=29%, list=13%, signal=26% |
| 8.53707312526996e-06 | 5.46406601968484e-06 | 2165 | tags=30%, list=12%, signal=27% |
| 8.53707312526996e-06 | 5.46406601968484e-06 | 2165 | tags=30%, list=12%, signal=27% |
| 0.000237309 | 0.000151887 | 1158 | tags=38%, list=6%, signal=35% |
| 0.000996084 | 0.000637533 | 1628 | tags=47%, list=9%, signal=43% |
| 0.000996084 | 0.000637533 | 2740 | tags=76%, list=15%, signal=65% |
| 4.34438310125395e-06 | 2.7805778083111e-06 | 3129 | tags=33%, list=17%, signal=27% |
| 0.000913107 | 0.000584425 | 1646 | tags=48%, list=9%, signal=43% |
| 0.000827871 | 0.00052987 | 3214 | tags=52%, list=18%, signal=43% |
| 0.000156196 | 9.99713882633755e-05 | 1697 | tags=31%, list=9%, signal=28% |
| 1.88325633883124e-06 | 1.20535888780248e-06 | 2239 | tags=29%, list=12%, signal=26% |

| | | | |
|---|---|---|---|
| 0.000573196 | 0.000366868 | 1693 | tags=42%, list=9%, signal=38% |
| 8.6635140201801e-05 | 5.54499321654016e-05 | 4285 | tags=59%, list=24%, signal=45% |
| 0.000133359 | 8.5355042455828e-05 | 4723 | tags=53%, list=26%, signal=39% |
| 0.000657233 | 0.000420655 | 1628 | tags=57%, list=9%, signal=52% |
| 1.51204967419287e-05 | 9.67771872584439e-06 | 2686 | tags=37%, list=15%, signal=31% |
| 0.000282085 | 0.000180546 | 4539 | tags=62%, list=25%, signal=46% |
| 0.000510392 | 0.000326671 | 709 | tags=50%, list=4%, signal=48% |
| 0.000341078 | 0.000218304 | 1515 | tags=60%, list=8%, signal=55% |
| 0.000832997 | 0.000533151 | 1848 | tags=59%, list=10%, signal=53% |
| 0.001002315 | 0.000641521 | 2329 | tags=83%, list=13%, signal=73% |
| 0.000324162 | 0.000207477 | 1468 | tags=70%, list=8%, signal=64% |
| 0.000379552 | 0.000242928 | 917 | tags=53%, list=5%, signal=51% |
| 0.00040831 | 0.000261335 | 1508 | tags=40%, list=8%, signal=37% |
| 0.00052181 | 0.000333979 | 2490 | tags=64%, list=14%, signal=55% |
| 9.61999827778634e-06 | 6.15718114718817e-06 | 2686 | tags=34%, list=15%, signal=29% |
| 0.00061457 | 0.000393349 | 2256 | tags=56%, list=13%, signal=49% |
| 0.000335339 | 0.00021463 | 2040 | tags=64%, list=11%, signal=56% |
| 0.000326155 | 0.000208752 | 3451 | tags=49%, list=19%, signal=40% |
| 1.35560665080642e-05 | 8.67642121373526e-06 | 2158 | tags=34%, list=12%, signal=30% |
| 4.6664861279795e-07 | 2.98673654413815e-07 | 3273 | tags=39%, list=18%, signal=33% |
| 0.00061677 | 0.000394757 | 3437 | tags=50%, list=19%, signal=40% |
| 0.000119048 | 7.61954328187298e-05 | 2206 | tags=34%, list=12%, signal=30% |
| 0.000160771 | 0.0001029 | 1589 | tags=35%, list=9%, signal=32% |
| 9.55767607265128e-05 | 6.11729245953685e-05 | 1496 | tags=35%, list=8%, signal=32% |
| 0.00085056 | 0.000544392 | 1125 | tags=56%, list=6%, signal=52% |
| 0.000368406 | 0.000235794 | 1496 | tags=56%, list=8%, signal=52% |
| 0.000192013 | 0.000122896 | 1933 | tags=67%, list=11%, signal=60% |
| 0.000443233 | 0.000283687 | 1508 | tags=69%, list=8%, signal=63% |
| 6.88452238663423e-05 | 4.40636788306509e-05 | 1697 | tags=32%, list=9%, signal=29% |
| 0.000712908 | 0.00045629 | 3437 | tags=65%, list=19%, signal=53% |
| 0.00013038 | 8.34484961795954e-05 | 1489 | tags=33%, list=8%, signal=30% |
| 0.000379887 | 0.000243143 | 2265 | tags=63%, list=13%, signal=55% |
| 0.000379887 | 0.000243143 | 2265 | tags=63%, list=13%, signal=55% |
| 0.00029564 | 0.000189221 | 1496 | tags=53%, list=8%, signal=49% |
| 0.000144718 | 9.26253951787103e-05 | 1589 | tags=34%, list=9%, signal=31% |
| 0.000143585 | 9.18999156327779e-05 | 1564 | tags=53%, list=9%, signal=49% |
| 0.000493093 | 0.000315599 | 1848 | tags=50%, list=10%, signal=45% |
| 0.000489674 | 0.000313411 | 2111 | tags=48%, list=12%, signal=43% |
| 4.61049460125296e-05 | 2.95089974221699e-05 | 3392 | tags=35%, list=19%, signal=29% |
| 0.000602903 | 0.000385882 | 2587 | tags=53%, list=14%, signal=45% |
| 1.0876118185509e-05 | 6.96114779989506e-06 | 3170 | tags=39%, list=18%, signal=32% |
| 0.000476937 | 0.000305259 | 1628 | tags=42%, list=9%, signal=38% |
| 0.000366589 | 0.000234631 | 3121 | tags=48%, list=17%, signal=40% |
| 3.8218800312313e-05 | 2.44615508190374e-05 | 5493 | tags=67%, list=31%, signal=46% |
| 3.8218800312313e-05 | 2.44615508190374e-05 | 5493 | tags=67%, list=31%, signal=46% |

| | | | |
|---|---|---|---|
| 0.000248208 | 0.000158863 | 1508 | tags=40%, list=8%, signal=37% |
| 0.000498329 | 0.00031895 | 1451 | tags=39%, list=8%, signal=36% |
| 0.000455729 | 0.000291685 | 2305 | tags=55%, list=13%, signal=48% |
| 0.000376037 | 0.000240679 | 1211 | tags=36%, list=7%, signal=34% |
| 0.000130256 | 8.33692839146143e-05 | 1211 | tags=36%, list=7%, signal=34% |
| 1.60571237377023e-08 | 1.02771972198937e-08 | 3120 | tags=35%, list=17%, signal=29% |
| 2.30707188748778e-06 | 1.47661767919945e-06 | 1988 | tags=36%, list=11%, signal=32% |
| 6.07849065058092e-06 | 3.89047554442271e-06 | 3451 | tags=41%, list=19%, signal=33% |
| 1.42999967379986e-05 | 9.15256612086609e-06 | 2666 | tags=42%, list=15%, signal=36% |
| 0.00041481 | 0.000265495 | 1628 | tags=48%, list=9%, signal=43% |
| 0.00041481 | 0.000265495 | 1628 | tags=48%, list=9%, signal=43% |
| 2.40078125e-09 | 1.53659539473684e-09 | 2634 | tags=30%, list=15%, signal=26% |
| 0.000356651 | 0.000228271 | 4588 | tags=58%, list=26%, signal=43% |
| 0.00015176 | 9.7132299527983e-05 | 3667 | tags=59%, list=20%, signal=47% |
| 0.000155519 | 9.95380458104375e-05 | 4817 | tags=73%, list=27%, signal=54% |
| 2.40078125e-09 | 1.53659539473684e-09 | 3328 | tags=37%, list=19%, signal=31% |
| 0.000307234 | 0.000196642 | 1340 | tags=43%, list=7%, signal=40% |
| 0.000307234 | 0.000196642 | 1646 | tags=64%, list=9%, signal=58% |
| 9.69941246927221e-05 | 6.2080093852519e-05 | 3022 | tags=83%, list=17%, signal=69% |
| 0.000279746 | 0.000179049 | 1105 | tags=48%, list=6%, signal=45% |
| 8.16827724095312e-05 | 5.22802199966462e-05 | 1103 | tags=29%, list=6%, signal=27% |
| 8.58502869632667e-05 | 5.49475949066963e-05 | 1508 | tags=58%, list=8%, signal=53% |
| 0.000460995 | 0.000295055 | 1693 | tags=44%, list=9%, signal=40% |
| 0.000160455 | 0.000102698 | 2027 | tags=62%, list=11%, signal=55% |
| 0.000366589 | 0.000234631 | 1858 | tags=55%, list=10%, signal=49% |
| 0.000391035 | 0.000250278 | 3233 | tags=53%, list=18%, signal=43% |
| 3.37240805893066e-05 | 2.15847515991983e-05 | 4466 | tags=57%, list=25%, signal=43% |
| 0.000117374 | 7.51237835866144e-05 | 3437 | tags=62%, list=19%, signal=50% |
| 0.000399402 | 0.000255633 | 1572 | tags=77%, list=9%, signal=70% |
| 2.40078125e-09 | 1.53659539473684e-09 | 2484 | tags=34%, list=14%, signal=30% |
| 2.53280339284534e-07 | 1.62109481204087e-07 | 2239 | tags=30%, list=12%, signal=26% |
| 0.00021053 | 0.000134748 | 1551 | tags=33%, list=9%, signal=31% |
| 1.36924516618555e-08 | 8.76371313140838e-09 | 4209 | tags=50%, list=23%, signal=39% |
| 0.000111627 | 7.14460211907593e-05 | 971 | tags=64%, list=5%, signal=60% |
| 0.000111627 | 7.14460211907593e-05 | 971 | tags=64%, list=5%, signal=60% |
| 0.000192998 | 0.000123526 | 2490 | tags=48%, list=14%, signal=42% |
| 2.67793571820694e-06 | 1.71398526708674e-06 | 2840 | tags=44%, list=16%, signal=37% |
| 0.000163501 | 0.000104647 | 1508 | tags=69%, list=8%, signal=63% |
| 0.000120662 | 7.7228518768556e-05 | 2265 | tags=43%, list=13%, signal=38% |
| 0.000156196 | 9.99713882633755e-05 | 1988 | tags=77%, list=11%, signal=68% |
| 0.000238171 | 0.000152439 | 1456 | tags=43%, list=8%, signal=40% |
| 0.000305622 | 0.00019561 | 1646 | tags=59%, list=9%, signal=53% |
| 9.71612308544668e-05 | 6.21870484359776e-05 | 1211 | tags=36%, list=7%, signal=34% |
| 3.31920280161974e-09 | 2.12442168113672e-09 | 1989 | tags=30%, list=11%, signal=27% |
| 2.40078125e-09 | 1.53659539473684e-09 | 2248 | tags=27%, list=13%, signal=24% |

| | | | |
|---|---|---|---|
| 1.65518140408931e-06 | 1.0593818670392e-06 | 1964 | tags=34%, list=11%, signal=31% |
| 0.000215084 | 0.000137662 | 3262 | tags=65%, list=18%, signal=53% |
| 0.000141824 | 9.07730722537603e-05 | 3023 | tags=58%, list=17%, signal=49% |
| 0.000211112 | 0.00013512 | 980 | tags=30%, list=5%, signal=28% |
| 2.40078125e-09 | 1.53659539473684e-09 | 2248 | tags=28%, list=13%, signal=24% |
| 4.27243564399147e-05 | 2.73452857684007e-05 | 971 | tags=58%, list=5%, signal=55% |
| 1.00570490944767e-07 | 6.43691103602846e-08 | 2265 | tags=32%, list=13%, signal=29% |
| 4.55172130107749e-05 | 2.9132824947551e-05 | 3747 | tags=62%, list=21%, signal=49% |
| 1.80351889435916e-05 | 1.15432375498316e-05 | 2707 | tags=37%, list=15%, signal=31% |
| 6.56845442931379e-05 | 4.20407183146002e-05 | 1211 | tags=36%, list=7%, signal=34% |
| 0.000114038 | 7.29890293068321e-05 | 2684 | tags=55%, list=15%, signal=47% |
| 2.01761118909629e-06 | 1.29135133054496e-06 | 2451 | tags=37%, list=14%, signal=32% |
| 6.32978917525807e-05 | 4.05131658553092e-05 | 2206 | tags=67%, list=12%, signal=59% |
| 9.36053509723e-06 | 5.99111440189571e-06 | 2072 | tags=36%, list=12%, signal=32% |
| 0.000143769 | 9.20179290599767e-05 | 1084 | tags=61%, list=6%, signal=57% |
| 2.40002829781587e-05 | 1.53611347542054e-05 | 3315 | tags=55%, list=18%, signal=45% |
| 8.3838666279922e-05 | 5.36600777378643e-05 | 3262 | tags=60%, list=18%, signal=49% |
| 2.3253471744415e-05 | 1.48831458901603e-05 | 3747 | tags=58%, list=21%, signal=46% |
| 3.35048031083725e-06 | 2.14444052984377e-06 | 2840 | tags=45%, list=16%, signal=38% |
| 2.40078125e-09 | 1.53659539473684e-09 | 2634 | tags=30%, list=15%, signal=26% |
| 2.23706630806825e-06 | 1.43181132670818e-06 | 2688 | tags=35%, list=15%, signal=30% |
| 9.27774216252865e-07 | 5.93812363392015e-07 | 3883 | tags=57%, list=22%, signal=45% |
| 9.27774216252865e-07 | 5.93812363392015e-07 | 3883 | tags=57%, list=22%, signal=45% |
| 6.46955170378834e-05 | 4.14077015723655e-05 | 2023 | tags=63%, list=11%, signal=56% |
| 0.000114038 | 7.29890293068321e-05 | 1368 | tags=50%, list=8%, signal=46% |
| 0.000114038 | 7.29890293068321e-05 | 1368 | tags=50%, list=8%, signal=46% |
| 0.000104399 | 6.68193850420653e-05 | 1105 | tags=52%, list=6%, signal=49% |
| 8.51193112911545e-05 | 5.44797414313194e-05 | 1513 | tags=67%, list=8%, signal=61% |
| 1.03600622491744e-05 | 6.63085149522408e-06 | 3683 | tags=54%, list=21%, signal=43% |
| 4.55678999684973e-06 | 2.91652666145331e-06 | 2175 | tags=34%, list=12%, signal=30% |
| 4.04043960944679e-05 | 2.58604189639862e-05 | 2265 | tags=69%, list=13%, signal=60% |
| 1.1046609701688e-05 | 7.07026914470824e-06 | 4850 | tags=69%, list=27%, signal=50% |
| 5.05152365634494e-05 | 3.23317586170912e-05 | 1693 | tags=57%, list=9%, signal=52% |
| 1.29551131948656e-05 | 8.29178721448488e-06 | 1496 | tags=39%, list=8%, signal=36% |
| 1.82567125726159e-05 | 1.16850214746203e-05 | 1933 | tags=73%, list=11%, signal=65% |
| 4.44261513938375e-07 | 2.84345021595168e-07 | 1508 | tags=32%, list=8%, signal=30% |
| 7.97031210875733e-05 | 5.10131644894003e-05 | 754 | tags=50%, list=4%, signal=48% |
| 8.41847192613022e-06 | 5.38815653963187e-06 | 1953 | tags=39%, list=11%, signal=35% |
| 9.68708464310713e-05 | 6.20011908666164e-05 | 2959 | tags=76%, list=16%, signal=64% |
| 2.12011540101826e-05 | 1.3569580991668e-05 | 1015 | tags=67%, list=6%, signal=63% |
| 2.12011540101826e-05 | 1.3569580991668e-05 | 1015 | tags=67%, list=6%, signal=63% |
| 4.47349539814673e-05 | 2.86321480858313e-05 | 1554 | tags=64%, list=9%, signal=58% |
| 4.32536390572985e-05 | 2.76840476745037e-05 | 1829 | tags=42%, list=10%, signal=38% |
| 4.56330584232465e-07 | 2.92069706146355e-07 | 2686 | tags=38%, list=15%, signal=32% |
| 7.23516008294066e-06 | 4.63078994124535e-06 | 1700 | tags=34%, list=9%, signal=31% |

| | | | |
|---|---|---|---|
| 3.24890579582204e-05 | 2.07942880418363e-05 | 1987 | tags=48%, list=11%, signal=43% |
| 2.02523677207261e-06 | 1.29623200665137e-06 | 1508 | tags=37%, list=8%, signal=34% |
| 5.01084947857883e-05 | 3.20714277175554e-05 | 1831 | tags=52%, list=10%, signal=47% |
| 4.76224732632332e-05 | 3.04802751613719e-05 | 1646 | tags=48%, list=9%, signal=43% |
| 4.76224732632332e-05 | 3.04802751613719e-05 | 1646 | tags=48%, list=9%, signal=43% |
| 4.76224732632332e-05 | 3.04802751613719e-05 | 1646 | tags=48%, list=9%, signal=43% |
| 6.32996480166188e-05 | 4.05142899340785e-05 | 2414 | tags=59%, list=13%, signal=51% |
| 2.64016363202409e-05 | 1.68980963106069e-05 | 1084 | tags=69%, list=6%, signal=65% |
| 9.11064496662141e-06 | 5.83117478895374e-06 | 754 | tags=60%, list=4%, signal=58% |
| 1.25722075156013e-06 | 8.04671236504738e-07 | 2082 | tags=38%, list=12%, signal=33% |
| 4.13925228623721e-05 | 2.6492859358536e-05 | 1456 | tags=38%, list=8%, signal=35% |
| 3.25417207071656e-05 | 2.08279942937088e-05 | 2734 | tags=61%, list=15%, signal=51% |
| 5.08138719105252e-05 | 3.25228971054572e-05 | 1602 | tags=48%, list=9%, signal=44% |
| 3.35929723213521e-07 | 2.1500837098137e-07 | 2451 | tags=38%, list=14%, signal=33% |
| 5.72278820151163e-06 | 3.66281184322691e-06 | 3315 | tags=56%, list=18%, signal=46% |
| 9.40203253247737e-06 | 6.01767440934933e-06 | 1564 | tags=41%, list=9%, signal=37% |
| 1.29551131948656e-05 | 8.29178721448488e-06 | 2827 | tags=53%, list=16%, signal=45% |
| 1.20249578391531e-05 | 7.69645082722438e-06 | 3527 | tags=56%, list=20%, signal=45% |
| 9.12305459829476e-06 | 5.83911744631982e-06 | 1084 | tags=64%, list=6%, signal=60% |
| 1.15282010620541e-05 | 7.37850675131388e-06 | 1079 | tags=53%, list=6%, signal=50% |
| 8.25346334366529e-06 | 5.28254449025934e-06 | 1079 | tags=67%, list=6%, signal=63% |
| 2.26233798925882e-05 | 1.44798620683718e-05 | 1260 | tags=50%, list=7%, signal=47% |
| 4.31958075683627e-05 | 2.76470332236579e-05 | 3769 | tags=64%, list=21%, signal=51% |
| 4.78467484951779e-05 | 3.06238202213643e-05 | 2329 | tags=64%, list=13%, signal=55% |
| 2.71098799821206e-05 | 1.73514003961815e-05 | 2175 | tags=41%, list=12%, signal=36% |
| 6.23321616744445e-05 | 3.98950602321406e-05 | 1602 | tags=67%, list=9%, signal=61% |
| 1.9537623423377e-05 | 1.25048553159368e-05 | 1826 | tags=58%, list=10%, signal=52% |
| 1.32724500459893e-07 | 8.49489540854332e-08 | 3883 | tags=57%, list=22%, signal=45% |
| 1.32724500459893e-07 | 8.49489540854332e-08 | 3883 | tags=57%, list=22%, signal=45% |
| 6.281266432336e-06 | 4.02026010201579e-06 | 1572 | tags=60%, list=9%, signal=55% |
| 2.6152842077231e-06 | 1.6738858109273e-06 | 3862 | tags=55%, list=22%, signal=43% |
| 3.77395949697355e-05 | 2.41548403586246e-05 | 1431 | tags=33%, list=8%, signal=31% |
| 8.05901634956968e-07 | 5.15809068771163e-07 | 2848 | tags=54%, list=16%, signal=46% |
| 5.92497954197665e-06 | 3.79222233517165e-06 | 1933 | tags=60%, list=11%, signal=54% |
| 5.92497954197665e-06 | 3.79222233517165e-06 | 1933 | tags=60%, list=11%, signal=54% |
| 1.42786888761064e-05 | 9.13892824258991e-06 | 1496 | tags=61%, list=8%, signal=56% |
| 2.40078125e-09 | 1.53659539473684e-09 | 3102 | tags=39%, list=17%, signal=33% |
| 2.97349573874548e-05 | 1.90315542427113e-05 | 4158 | tags=76%, list=23%, signal=58% |
| 2.16481555837836e-05 | 1.38556797603232e-05 | 1628 | tags=70%, list=9%, signal=64% |
| 3.6444280043908e-06 | 2.33257873369216e-06 | 1987 | tags=39%, list=11%, signal=35% |
| 1.27704634703454e-05 | 8.17360405375869e-06 | 3742 | tags=62%, list=21%, signal=50% |
| 1.7523011893503e-06 | 1.12154238864851e-06 | 3468 | tags=50%, list=19%, signal=40% |
| 8.99479241546582e-07 | 5.75702455282781e-07 | 2848 | tags=46%, list=16%, signal=39% |
| 2.58870785251054e-06 | 1.65687588758317e-06 | 1700 | tags=37%, list=9%, signal=33% |
| 1.45030986638821e-05 | 9.28255942366067e-06 | 1508 | tags=39%, list=8%, signal=36% |

| | | | |
|---|---|---|---|
| 8.35222644991785e-09 | 5.34575680260041e-09 | 3837 | tags=49%, list=21%, signal=39% |
| 2.09131866112027e-08 | 1.33852704139731e-08 | 2686 | tags=51%, list=15%, signal=44% |
| 2.40078125e-09 | 1.53659539473684e-09 | 3311 | tags=39%, list=18%, signal=33% |
| 1.31759747351593e-07 | 8.43314737617799e-08 | 2072 | tags=37%, list=12%, signal=33% |
| 1.38909939988062e-05 | 8.89078811611125e-06 | 1820 | tags=39%, list=10%, signal=35% |
| 1.38909939988062e-05 | 8.89078811611125e-06 | 1820 | tags=39%, list=10%, signal=35% |
| 1.63037987479882e-05 | 1.04350790280768e-05 | 1515 | tags=62%, list=8%, signal=57% |
| 4.41766802805598e-06 | 2.82748307343162e-06 | 1903 | tags=53%, list=11%, signal=48% |
| 6.56694087368637e-08 | 4.20310309571754e-08 | 2451 | tags=36%, list=14%, signal=31% |
| 4.43447151537562e-06 | 2.83823797300061e-06 | 1554 | tags=60%, list=9%, signal=55% |
| 3.75010708427869e-07 | 2.40021754396517e-07 | 3831 | tags=56%, list=21%, signal=44% |
| 7.19468030428549e-07 | 4.6048812744472e-07 | 2414 | tags=39%, list=13%, signal=34% |
| 2.13890733468241e-05 | 1.36898568340695e-05 | 1933 | tags=67%, list=11%, signal=60% |
| 1.82307607403473e-08 | 1.16684112707757e-08 | 3360 | tags=39%, list=19%, signal=32% |
| 2.40078125e-09 | 1.53659539473684e-09 | 2906 | tags=36%, list=16%, signal=31% |
| 5.88726500246888e-06 | 3.76808353130426e-06 | 1431 | tags=58%, list=8%, signal=53% |
| 1.18296165430745e-05 | 7.57142463587264e-06 | 2602 | tags=62%, list=14%, signal=53% |
| 9.11008243186458e-06 | 5.83081474435713e-06 | 1508 | tags=48%, list=8%, signal=44% |
| 5.22923198725286e-07 | 3.34691625470806e-07 | 1948 | tags=39%, list=11%, signal=35% |
| 2.40078125e-09 | 1.53659539473684e-09 | 2906 | tags=36%, list=16%, signal=31% |
| 8.08743109566974e-06 | 5.17627725427198e-06 | 1554 | tags=46%, list=9%, signal=42% |
| 1.26345391004706e-06 | 8.08660705610127e-07 | 1988 | tags=40%, list=11%, signal=36% |
| 3.98342481439291e-06 | 2.54955016208853e-06 | 1508 | tags=36%, list=8%, signal=33% |
| 2.36230516161817e-06 | 1.51196916933001e-06 | 2560 | tags=44%, list=14%, signal=38% |
| 1.01307989008926e-08 | 6.48411384257379e-09 | 2329 | tags=50%, list=13%, signal=43% |
| 4.34438310125395e-06 | 2.7805778083111e-06 | 1037 | tags=47%, list=6%, signal=45% |
| 8.05307796534461e-07 | 5.15428988584664e-07 | 2590 | tags=48%, list=14%, signal=42% |
| 2.21212750204435e-06 | 1.41584949991261e-06 | 1079 | tags=69%, list=6%, signal=65% |
| 7.91937707215056e-06 | 5.06871600161451e-06 | 1211 | tags=50%, list=7%, signal=47% |
| 7.91937707215056e-06 | 5.06871600161451e-06 | 1211 | tags=50%, list=7%, signal=47% |
| 3.64159087426532e-08 | 2.33076285767885e-08 | 4427 | tags=58%, list=25%, signal=44% |
| 1.91668182573624e-06 | 1.22675252757914e-06 | 1933 | tags=69%, list=11%, signal=61% |
| 1.34696384416023e-06 | 8.62110381699143e-07 | 3023 | tags=55%, list=17%, signal=46% |
| 2.40078125e-09 | 1.53659539473684e-09 | 2072 | tags=37%, list=12%, signal=33% |
| 8.10510459055578e-06 | 5.18758899325311e-06 | 2023 | tags=47%, list=11%, signal=42% |
| 8.10510459055578e-06 | 5.18758899325311e-06 | 2023 | tags=47%, list=11%, signal=42% |
| 2.40078125e-09 | 1.53659539473684e-09 | 3546 | tags=41%, list=20%, signal=33% |
| 5.04847116528735e-07 | 3.23122214614192e-07 | 1646 | tags=40%, list=9%, signal=36% |
| 1.23398804123892e-06 | 7.89801378750381e-07 | 1260 | tags=53%, list=7%, signal=50% |
| 2.46243994282915e-07 | 1.5760594081478e-07 | 3506 | tags=38%, list=20%, signal=31% |
| 2.40078125e-09 | 1.53659539473684e-09 | 2175 | tags=29%, list=12%, signal=26% |
| 1.44902723217768e-06 | 9.27435005506022e-07 | 1508 | tags=41%, list=8%, signal=37% |
| 9.76404673570134e-06 | 6.24937788400088e-06 | 1496 | tags=48%, list=8%, signal=44% |
| 2.82505603473674e-08 | 1.80814811547283e-08 | 3360 | tags=42%, list=19%, signal=34% |
| 2.3900083928806e-06 | 1.52970033812232e-06 | 1554 | tags=42%, list=9%, signal=38% |

| | | | |
|---|---|---|---|
| 2.02464290882777e-08 | 1.29585191057759e-08 | 4756 | tags=55%, list=26%, signal=41% |
| 6.71886798700968e-07 | 4.30034248504893e-07 | 1602 | tags=37%, list=9%, signal=34% |
| 2.40078125e-09 | 1.53659539473684e-09 | 3311 | tags=43%, list=18%, signal=35% |
| 2.40078125e-09 | 1.53659539473684e-09 | 4091 | tags=48%, list=23%, signal=37% |
| 1.29781156760203e-07 | 8.30650971642449e-08 | 1646 | tags=38%, list=9%, signal=35% |
| 6.0230834943107e-07 | 3.85501276281348e-07 | 2265 | tags=49%, list=13%, signal=43% |
| 2.40078125e-09 | 1.53659539473684e-09 | 2451 | tags=35%, list=14%, signal=31% |
| 2.40078125e-09 | 1.53659539473684e-09 | 2256 | tags=30%, list=13%, signal=27% |
| 2.40078125e-09 | 1.53659539473684e-09 | 2329 | tags=34%, list=13%, signal=30% |
| 2.5952215417833e-06 | 1.66104490753836e-06 | 1826 | tags=48%, list=10%, signal=43% |
| 1.88325633883124e-06 | 1.20535888780248e-06 | 1273 | tags=41%, list=7%, signal=38% |
| 2.40078125e-09 | 1.53659539473684e-09 | 3527 | tags=49%, list=20%, signal=39% |
| 5.72922491463342e-07 | 3.66693159538683e-07 | 1366 | tags=62%, list=8%, signal=58% |
| 9.93999010749754e-07 | 6.36198863303789e-07 | 1178 | tags=38%, list=7%, signal=36% |
| 2.40078125e-09 | 1.53659539473684e-09 | 1989 | tags=31%, list=11%, signal=28% |
| 3.13797421784246e-06 | 2.00842818642459e-06 | 1297 | tags=50%, list=7%, signal=46% |
| 6.46279744546742e-07 | 4.13644716353157e-07 | 1595 | tags=41%, list=9%, signal=37% |
| 3.76656568709828e-06 | 2.4107517037502e-06 | 1628 | tags=56%, list=9%, signal=51% |
| 1.89766655061526e-08 | 1.21458199593218e-08 | 2329 | tags=42%, list=13%, signal=36% |
| 1.51599267564057e-08 | 9.70295550185628e-09 | 1595 | tags=39%, list=9%, signal=35% |
| 1.16739215998509e-08 | 7.47177368569077e-09 | 2842 | tags=49%, list=16%, signal=42% |
| 3.09336364490863e-07 | 1.97987564715152e-07 | 2827 | tags=50%, list=16%, signal=42% |
| 4.68905727102337e-08 | 3.00118297254772e-08 | 1964 | tags=40%, list=11%, signal=36% |
| 2.1511069679748e-06 | 1.37679393346496e-06 | 1628 | tags=50%, list=9%, signal=46% |
| 1.03203171513603e-07 | 6.60541305335667e-08 | 1826 | tags=42%, list=10%, signal=38% |
| 1.72292413229355e-06 | 1.10273990483858e-06 | 2939 | tags=68%, list=16%, signal=57% |
| 2.40078125e-09 | 1.53659539473684e-09 | 1989 | tags=31%, list=11%, signal=28% |
| 2.72044049550266e-07 | 1.74119001347791e-07 | 1211 | tags=36%, list=7%, signal=34% |
| 2.25709281385711e-06 | 1.4446290861637e-06 | 2686 | tags=62%, list=15%, signal=53% |
| 1.0843752461578e-06 | 6.94043245053127e-07 | 2023 | tags=48%, list=11%, signal=42% |
| 2.04685763146864e-06 | 1.31007021576692e-06 | 1697 | tags=44%, list=9%, signal=40% |
| 2.40078125e-09 | 1.53659539473684e-09 | 2175 | tags=30%, list=12%, signal=27% |
| 4.68997266005306e-09 | 3.00176885789958e-09 | 1987 | tags=34%, list=11%, signal=31% |
| 2.12902065283372e-06 | 1.36265781417775e-06 | 1211 | tags=56%, list=7%, signal=52% |
| 7.00074212128235e-07 | 4.48075313121622e-07 | 2206 | tags=50%, list=12%, signal=44% |
| 2.40078125e-09 | 1.53659539473684e-09 | 1652 | tags=36%, list=9%, signal=33% |
| 1.14543542587059e-08 | 7.33124186287767e-09 | 3035 | tags=48%, list=17%, signal=40% |
| 2.40078125e-09 | 1.53659539473684e-09 | 1948 | tags=38%, list=11%, signal=34% |
| 6.87314898067095e-07 | 4.39908845132775e-07 | 2023 | tags=53%, list=11%, signal=47% |
| 2.40078125e-09 | 1.53659539473684e-09 | 2144 | tags=32%, list=12%, signal=29% |
| 2.40078125e-09 | 1.53659539473684e-09 | 2175 | tags=30%, list=12%, signal=27% |
| 2.40078125e-09 | 1.53659539473684e-09 | 1989 | tags=33%, list=11%, signal=30% |
| 9.13403855798744e-07 | 5.84614761696937e-07 | 1903 | tags=62%, list=11%, signal=56% |
| 5.86506744122451e-07 | 3.7538762100906e-07 | 1697 | tags=46%, list=9%, signal=42% |
| 5.92761031822064e-07 | 3.79390613650136e-07 | 2023 | tags=50%, list=11%, signal=44% |

| | | |
|---|---|---|
| 6.42662426160284e-07 | 4.11329488852139e-07 | 1725 tags=61%, list=10%, signal=55% |
| 1.80725358810191e-08 | 1.1567141073761e-08 | 2634 tags=43%, list=15%, signal=37% |
| 7.21583754972817e-07 | 4.61842275221097e-07 | 1515 tags=67%, list=8%, signal=61% |
| 4.53680478499611e-09 | 2.90373533175715e-09 | 2686 tags=49%, list=15%, signal=41% |
| 1.26155022400556e-06 | 8.07442270900849e-07 | 1211 tags=54%, list=7%, signal=50% |
| 2.10152894344506e-07 | 1.34506202778944e-07 | 1373 tags=45%, list=8%, signal=42% |
| 4.0662143061133e-07 | 2.60253872641948e-07 | 1964 tags=71%, list=11%, signal=64% |
| 2.40078125e-09 | 1.53659539473684e-09 | 1628 tags=38%, list=9%, signal=35% |
| 1.15147637958131e-06 | 7.36990636699158e-07 | 1987 tags=65%, list=11%, signal=58% |
| 2.40078125e-09 | 1.53659539473684e-09 | 1654 tags=34%, list=9%, signal=31% |
| 3.9615861575072e-09 | 2.53557255392543e-09 | 1515 tags=36%, list=8%, signal=33% |
| 2.40078125e-09 | 1.53659539473684e-09 | 5866 tags=65%, list=33%, signal=45% |
| 2.40078125e-09 | 1.53659539473684e-09 | 1628 tags=33%, list=9%, signal=31% |
| 2.40078125e-09 | 1.53659539473684e-09 | 2305 tags=33%, list=13%, signal=29% |
| 2.40078125e-09 | 1.53659539473684e-09 | 2898 tags=44%, list=16%, signal=37% |
| 2.40078125e-09 | 1.53659539473684e-09 | 2241 tags=36%, list=12%, signal=32% |
| 2.40078125e-09 | 1.53659539473684e-09 | 2399 tags=38%, list=13%, signal=33% |
| 3.78760895226981e-08 | 2.42422022961144e-08 | 1829 tags=41%, list=10%, signal=37% |
| 3.78760895226981e-08 | 2.42422022961144e-08 | 1829 tags=41%, list=10%, signal=37% |
| 3.33035166541617e-07 | 2.13155739696512e-07 | 1848 tags=58%, list=10%, signal=52% |
| 2.40078125e-09 | 1.53659539473684e-09 | 1989 tags=32%, list=11%, signal=29% |
| 2.40078125e-09 | 1.53659539473684e-09 | 1987 tags=37%, list=11%, signal=34% |
| 2.40078125e-09 | 1.53659539473684e-09 | 3546 tags=58%, list=20%, signal=47% |
| 2.40078125e-09 | 1.53659539473684e-09 | 2023 tags=33%, list=11%, signal=30% |
| 4.11949010140116e-08 | 2.63663735231064e-08 | 1829 tags=41%, list=10%, signal=37% |
| 2.40078125e-09 | 1.53659539473684e-09 | 2686 tags=49%, list=15%, signal=42% |
| 2.62876715680976e-07 | 1.68251543408603e-07 | 1858 tags=63%, list=10%, signal=57% |
| 2.40078125e-09 | 1.53659539473684e-09 | 2332 tags=37%, list=13%, signal=32% |
| 2.40078125e-09 | 1.53659539473684e-09 | 1652 tags=37%, list=9%, signal=34% |
| 1.59333005060761e-08 | 1.01979454315527e-08 | 2678 tags=42%, list=15%, signal=36% |
| 8.09802052810838e-08 | 5.18305491180246e-08 | 2127 tags=62%, list=12%, signal=55% |
| 2.40078125e-09 | 1.53659539473684e-09 | 2040 tags=34%, list=11%, signal=31% |
| 4.1945067667871e-08 | 2.68465099893527e-08 | 1987 tags=47%, list=11%, signal=42% |
| 2.40078125e-09 | 1.53659539473684e-09 | 3792 tags=45%, list=21%, signal=36% |
| 7.27070304970829e-08 | 4.65353885227189e-08 | 1700 tags=44%, list=9%, signal=40% |
| 7.56910398190898e-08 | 4.84452730580332e-08 | 1508 tags=50%, list=8%, signal=46% |
| 5.22735936035977e-08 | 3.34571769908789e-08 | 2827 tags=70%, list=16%, signal=59% |
| 1.27227332027343e-08 | 8.14305478593147e-09 | 2265 tags=48%, list=13%, signal=43% |
| 4.79304144463406e-08 | 3.06773697545644e-08 | 1725 tags=59%, list=10%, signal=53% |
| 2.4918115935135e-08 | 1.59485843166457e-08 | 3637 tags=60%, list=20%, signal=48% |
| 4.17887705181926e-08 | 2.67464736031113e-08 | 1496 tags=52%, list=8%, signal=48% |
| 1.02162873431244e-07 | 6.53882984247453e-08 | 2586 tags=59%, list=14%, signal=51% |
| 2.40078125e-09 | 1.53659539473684e-09 | 3328 tags=52%, list=19%, signal=42% |
| 2.40078125e-09 | 1.53659539473684e-09 | 3328 tags=52%, list=19%, signal=42% |
| 2.40078125e-09 | 1.53659539473684e-09 | 2111 tags=37%, list=12%, signal=32% |

| | | | |
|---|---|---|---|
| 9.41518619904932e-08 | 6.02609327861464e-08 | 1236 | tags=60%, list=7%, signal=56% |
| 2.23860266935198e-08 | 1.43279465897688e-08 | 3170 | tags=62%, list=18%, signal=51% |
| 1.83118170022155e-08 | 1.17202904991315e-08 | 1933 | tags=78%, list=11%, signal=69% |
| 2.40078125e-09 | 1.53659539473684e-09 | 2175 | tags=32%, list=12%, signal=28% |
| 1.45004566926918e-08 | 9.28086845712048e-09 | 2707 | tags=60%, list=15%, signal=51% |
| 3.35150106824518e-08 | 2.1450938551445e-08 | 1829 | tags=52%, list=10%, signal=46% |
| 2.40078125e-09 | 1.53659539473684e-09 | 3328 | tags=52%, list=19%, signal=43% |
| 2.40078125e-09 | 1.53659539473684e-09 | 3328 | tags=52%, list=19%, signal=43% |
| 2.40078125e-09 | 1.53659539473684e-09 | 1749 | tags=32%, list=10%, signal=30% |
| 2.40078125e-09 | 1.53659539473684e-09 | 2329 | tags=35%, list=13%, signal=31% |
| 9.06930761170247e-09 | 5.80471723927109e-09 | 1496 | tags=53%, list=8%, signal=49% |
| 2.40078125e-09 | 1.53659539473684e-09 | 2265 | tags=33%, list=13%, signal=30% |
| 2.40078125e-09 | 1.53659539473684e-09 | 1732 | tags=41%, list=10%, signal=38% |
| 2.40078125e-09 | 1.53659539473684e-09 | 1282 | tags=35%, list=7%, signal=33% |
| 2.40078125e-09 | 1.53659539473684e-09 | 2848 | tags=47%, list=16%, signal=40% |
| 2.40078125e-09 | 1.53659539473684e-09 | 2848 | tags=47%, list=16%, signal=40% |
| 2.01671063005855e-08 | 1.29077493697721e-08 | 1079 | tags=54%, list=6%, signal=51% |
| 2.40078125e-09 | 1.53659539473684e-09 | 1477 | tags=33%, list=8%, signal=30% |
| 2.40078125e-09 | 1.53659539473684e-09 | 1564 | tags=38%, list=9%, signal=35% |
| 2.40078125e-09 | 1.53659539473684e-09 | 2734 | tags=41%, list=15%, signal=35% |
| 2.40078125e-09 | 1.53659539473684e-09 | 1903 | tags=34%, list=11%, signal=31% |
| 2.40078125e-09 | 1.53659539473684e-09 | 1903 | tags=34%, list=11%, signal=31% |
| 2.40078125e-09 | 1.53659539473684e-09 | 1826 | tags=43%, list=10%, signal=39% |
| 2.40078125e-09 | 1.53659539473684e-09 | 1554 | tags=44%, list=9%, signal=41% |
| 2.40078125e-09 | 1.53659539473684e-09 | 1987 | tags=41%, list=11%, signal=36% |
| 2.40078125e-09 | 1.53659539473684e-09 | 1628 | tags=39%, list=9%, signal=36% |
| 2.40078125e-09 | 1.53659539473684e-09 | 1848 | tags=43%, list=10%, signal=39% |
| 9.2179858194876e-09 | 5.89987719996321e-09 | 1587 | tags=68%, list=9%, signal=62% |
| 2.40078125e-09 | 1.53659539473684e-09 | 1564 | tags=35%, list=9%, signal=32% |
| 2.40078125e-09 | 1.53659539473684e-09 | 2023 | tags=42%, list=11%, signal=38% |
| 2.40078125e-09 | 1.53659539473684e-09 | 6103 | tags=69%, list=34%, signal=46% |
| 2.40078125e-09 | 1.53659539473684e-09 | 1628 | tags=45%, list=9%, signal=41% |
| 2.40078125e-09 | 1.53659539473684e-09 | 2265 | tags=45%, list=13%, signal=39% |
| 8.35222644991785e-09 | 5.34575680260041e-09 | 1105 | tags=48%, list=6%, signal=45% |
| 2.40078125e-09 | 1.53659539473684e-09 | 2027 | tags=52%, list=11%, signal=47% |
| 2.40078125e-09 | 1.53659539473684e-09 | 2734 | tags=42%, list=15%, signal=36% |
| 2.40078125e-09 | 1.53659539473684e-09 | 1628 | tags=46%, list=9%, signal=42% |
| 2.40078125e-09 | 1.53659539473684e-09 | 2009 | tags=46%, list=11%, signal=41% |
| 8.16430119879261e-09 | 5.22547717469437e-09 | 1508 | tags=53%, list=8%, signal=49% |
| 4.08569764776985e-09 | 2.61500883924777e-09 | 1260 | tags=68%, list=7%, signal=63% |
| 2.40078125e-09 | 1.53659539473684e-09 | 1646 | tags=38%, list=9%, signal=35% |
| 2.40078125e-09 | 1.53659539473684e-09 | 2040 | tags=34%, list=11%, signal=31% |
| 2.40078125e-09 | 1.53659539473684e-09 | 2040 | tags=34%, list=11%, signal=31% |
| 2.40078125e-09 | 1.53659539473684e-09 | 2023 | tags=37%, list=11%, signal=33% |
| 2.40078125e-09 | 1.53659539473684e-09 | 1463 | tags=45%, list=8%, signal=41% |

| | | | |
|---|---|---|---|
| 2.40078125e-09 | 1.53659539473684e-09 | 2324 | tags=36%, list=13%, signal=32% |
| 2.65008163286491e-09 | 1.69615754568931e-09 | 1829 | tags=50%, list=10%, signal=45% |
| 2.40078125e-09 | 1.53659539473684e-09 | 2178 | tags=55%, list=12%, signal=49% |
| 2.40078125e-09 | 1.53659539473684e-09 | 1602 | tags=42%, list=9%, signal=38% |
| 2.40078125e-09 | 1.53659539473684e-09 | 6103 | tags=71%, list=34%, signal=48% |
| 2.40078125e-09 | 1.53659539473684e-09 | 2707 | tags=59%, list=15%, signal=51% |
| 2.40078125e-09 | 1.53659539473684e-09 | 2707 | tags=59%, list=15%, signal=51% |
| 2.40078125e-09 | 1.53659539473684e-09 | 2009 | tags=45%, list=11%, signal=40% |
| 2.40078125e-09 | 1.53659539473684e-09 | 1628 | tags=52%, list=9%, signal=48% |
| 2.40884843778509e-09 | 1.54175871546798e-09 | 1987 | tags=62%, list=11%, signal=55% |
| 2.40884843778509e-09 | 1.54175871546798e-09 | 1987 | tags=62%, list=11%, signal=55% |
| 2.40078125e-09 | 1.53659539473684e-09 | 2162 | tags=64%, list=12%, signal=57% |
| 2.40078125e-09 | 1.53659539473684e-09 | 1831 | tags=39%, list=10%, signal=35% |
| 2.40078125e-09 | 1.53659539473684e-09 | 5484 | tags=68%, list=31%, signal=48% |
| 2.40078125e-09 | 1.53659539473684e-09 | 3328 | tags=45%, list=19%, signal=37% |
| 2.40078125e-09 | 1.53659539473684e-09 | 2230 | tags=35%, list=12%, signal=32% |
| 2.40078125e-09 | 1.53659539473684e-09 | 1948 | tags=40%, list=11%, signal=36% |
| 2.40078125e-09 | 1.53659539473684e-09 | 2175 | tags=38%, list=12%, signal=33% |
| 2.40078125e-09 | 1.53659539473684e-09 | 2707 | tags=40%, list=15%, signal=35% |
| 2.40078125e-09 | 1.53659539473684e-09 | 1602 | tags=68%, list=9%, signal=62% |
| 2.40078125e-09 | 1.53659539473684e-09 | 1957 | tags=50%, list=11%, signal=45% |
| 2.40078125e-09 | 1.53659539473684e-09 | 1628 | tags=53%, list=9%, signal=49% |
| 2.40078125e-09 | 1.53659539473684e-09 | 2178 | tags=56%, list=12%, signal=50% |
| 2.40078125e-09 | 1.53659539473684e-09 | 2451 | tags=41%, list=14%, signal=36% |
| 2.40078125e-09 | 1.53659539473684e-09 | 1646 | tags=40%, list=9%, signal=37% |
| 2.40078125e-09 | 1.53659539473684e-09 | 6103 | tags=71%, list=34%, signal=48% |
| 2.40078125e-09 | 1.53659539473684e-09 | 1933 | tags=45%, list=11%, signal=40% |
| 2.40078125e-09 | 1.53659539473684e-09 | 1933 | tags=45%, list=11%, signal=40% |
| 2.40078125e-09 | 1.53659539473684e-09 | 2206 | tags=40%, list=12%, signal=35% |
| 2.40078125e-09 | 1.53659539473684e-09 | 2035 | tags=39%, list=11%, signal=35% |
| 2.40078125e-09 | 1.53659539473684e-09 | 1988 | tags=62%, list=11%, signal=55% |
| 2.40078125e-09 | 1.53659539473684e-09 | 1988 | tags=62%, list=11%, signal=55% |
| 2.40078125e-09 | 1.53659539473684e-09 | 2162 | tags=51%, list=12%, signal=45% |
| 2.40078125e-09 | 1.53659539473684e-09 | 1848 | tags=48%, list=10%, signal=43% |
| 2.40078125e-09 | 1.53659539473684e-09 | 2243 | tags=64%, list=12%, signal=56% |
| 2.40078125e-09 | 1.53659539473684e-09 | 2178 | tags=50%, list=12%, signal=44% |
| 2.40078125e-09 | 1.53659539473684e-09 | 2178 | tags=50%, list=12%, signal=44% |
| 2.40078125e-09 | 1.53659539473684e-09 | 2705 | tags=51%, list=15%, signal=43% |
| 2.40078125e-09 | 1.53659539473684e-09 | 6103 | tags=73%, list=34%, signal=49% |
| 2.40078125e-09 | 1.53659539473684e-09 | 2162 | tags=65%, list=12%, signal=58% |
| 2.40078125e-09 | 1.53659539473684e-09 | 2178 | tags=49%, list=12%, signal=43% |
| 2.40078125e-09 | 1.53659539473684e-09 | 2666 | tags=51%, list=15%, signal=44% |
| 2.40078125e-09 | 1.53659539473684e-09 | 1858 | tags=63%, list=10%, signal=57% |
| 2.40078125e-09 | 1.53659539473684e-09 | 2144 | tags=48%, list=12%, signal=43% |
| 2.40078125e-09 | 1.53659539473684e-09 | 1515 | tags=44%, list=8%, signal=40% |

| | | | |
|---|---|---|---|
| 2.40078125e-09 | 1.53659539473684e-09 | 2305 | tags=41%, list=13%, signal=36% |
| 2.40078125e-09 | 1.53659539473684e-09 | 1515 | tags=47%, list=8%, signal=43% |
| 2.40078125e-09 | 1.53659539473684e-09 | 2162 | tags=60%, list=12%, signal=53% |
| 2.40078125e-09 | 1.53659539473684e-09 | 2023 | tags=43%, list=11%, signal=39% |
| 2.40078125e-09 | 1.53659539473684e-09 | 1508 | tags=57%, list=8%, signal=52% |
| 2.40078125e-09 | 1.53659539473684e-09 | 1646 | tags=48%, list=9%, signal=43% |
| 2.40078125e-09 | 1.53659539473684e-09 | 1646 | tags=37%, list=9%, signal=34% |
| 2.40078125e-09 | 1.53659539473684e-09 | 1515 | tags=53%, list=8%, signal=49% |
| 2.40078125e-09 | 1.53659539473684e-09 | 1628 | tags=41%, list=9%, signal=38% |
| 2.40078125e-09 | 1.53659539473684e-09 | 1848 | tags=44%, list=10%, signal=40% |
| 2.40078125e-09 | 1.53659539473684e-09 | 1737 | tags=65%, list=10%, signal=59% |
| 2.40078125e-09 | 1.53659539473684e-09 | 2144 | tags=40%, list=12%, signal=36% |
| 2.40078125e-09 | 1.53659539473684e-09 | 2686 | tags=40%, list=15%, signal=35% |
| 2.40078125e-09 | 1.53659539473684e-09 | 2162 | tags=54%, list=12%, signal=47% |
| 2.40078125e-09 | 1.53659539473684e-09 | 1831 | tags=37%, list=10%, signal=34% |
| 2.40078125e-09 | 1.53659539473684e-09 | 1513 | tags=47%, list=8%, signal=43% |
| 2.40078125e-09 | 1.53659539473684e-09 | 2162 | tags=56%, list=12%, signal=49% |
| 2.40078125e-09 | 1.53659539473684e-09 | 1519 | tags=41%, list=8%, signal=38% |
| 2.40078125e-09 | 1.53659539473684e-09 | 2890 | tags=57%, list=16%, signal=48% |
| 2.40078125e-09 | 1.53659539473684e-09 | 2890 | tags=57%, list=16%, signal=48% |
| 2.40078125e-09 | 1.53659539473684e-09 | 2057 | tags=53%, list=11%, signal=48% |
| 2.40078125e-09 | 1.53659539473684e-09 | 2305 | tags=44%, list=13%, signal=38% |
| 2.40078125e-09 | 1.53659539473684e-09 | 2023 | tags=56%, list=11%, signal=49% |
| 2.40078125e-09 | 1.53659539473684e-09 | 1628 | tags=35%, list=9%, signal=33% |
| 2.40078125e-09 | 1.53659539473684e-09 | 1260 | tags=46%, list=7%, signal=43% |
| 2.40078125e-09 | 1.53659539473684e-09 | 1602 | tags=40%, list=9%, signal=37% |
| 2.40078125e-09 | 1.53659539473684e-09 | 1508 | tags=54%, list=8%, signal=50% |
| 2.40078125e-09 | 1.53659539473684e-09 | 1628 | tags=35%, list=9%, signal=32% |
| 2.40078125e-09 | 1.53659539473684e-09 | 2111 | tags=51%, list=12%, signal=45% |
| 2.40078125e-09 | 1.53659539473684e-09 | 1628 | tags=37%, list=9%, signal=34% |
| 2.40078125e-09 | 1.53659539473684e-09 | 2175 | tags=34%, list=12%, signal=30% |
| 2.40078125e-09 | 1.53659539473684e-09 | 2890 | tags=60%, list=16%, signal=51% |
| 2.40078125e-09 | 1.53659539473684e-09 | 2175 | tags=33%, list=12%, signal=30% |
| 2.40078125e-09 | 1.53659539473684e-09 | 2305 | tags=39%, list=13%, signal=35% |
| 2.40078125e-09 | 1.53659539473684e-09 | 1987 | tags=45%, list=11%, signal=41% |
| 2.40078125e-09 | 1.53659539473684e-09 | 2206 | tags=50%, list=12%, signal=44% |
| 2.40078125e-09 | 1.53659539473684e-09 | 2206 | tags=39%, list=12%, signal=35% |
| 2.40078125e-09 | 1.53659539473684e-09 | 1628 | tags=42%, list=9%, signal=38% |
| 2.40078125e-09 | 1.53659539473684e-09 | 2122 | tags=52%, list=12%, signal=46% |
| 2.40078125e-09 | 1.53659539473684e-09 | 1211 | tags=51%, list=7%, signal=48% |
| 2.40078125e-09 | 1.53659539473684e-09 | 1831 | tags=38%, list=10%, signal=35% |
| 2.40078125e-09 | 1.53659539473684e-09 | 1831 | tags=38%, list=10%, signal=35% |
| 2.40078125e-09 | 1.53659539473684e-09 | 2206 | tags=42%, list=12%, signal=37% |
| 2.40078125e-09 | 1.53659539473684e-09 | 2206 | tags=41%, list=12%, signal=37% |
| 2.40078125e-09 | 1.53659539473684e-09 | 1646 | tags=41%, list=9%, signal=37% |

| | | | |
|---|---|---|---|
| 2.40078125e-09 | 1.53659539473684e-09 | 1628 | tags=39%, list=9%, signal=36% |
| 2.40078125e-09 | 1.53659539473684e-09 | 1987 | tags=48%, list=11%, signal=43% |
| 2.40078125e-09 | 1.53659539473684e-09 | 1654 | tags=37%, list=9%, signal=34% |
| 2.40078125e-09 | 1.53659539473684e-09 | 2634 | tags=45%, list=15%, signal=39% |
| 2.40078125e-09 | 1.53659539473684e-09 | 2178 | tags=39%, list=12%, signal=35% |
| 2.40078125e-09 | 1.53659539473684e-09 | 1987 | tags=45%, list=11%, signal=40% |
| 2.40078125e-09 | 1.53659539473684e-09 | 1646 | tags=59%, list=9%, signal=54% |
| 2.40078125e-09 | 1.53659539473684e-09 | 1848 | tags=35%, list=10%, signal=33% |
| 2.40078125e-09 | 1.53659539473684e-09 | 2684 | tags=47%, list=15%, signal=41% |
| 2.40078125e-09 | 1.53659539473684e-09 | 1891 | tags=43%, list=11%, signal=39% |
| 2.40078125e-09 | 1.53659539473684e-09 | 1957 | tags=62%, list=11%, signal=56% |
| 2.40078125e-09 | 1.53659539473684e-09 | 1848 | tags=45%, list=10%, signal=41% |
| 2.40078125e-09 | 1.53659539473684e-09 | 2122 | tags=44%, list=12%, signal=39% |
| 2.40078125e-09 | 1.53659539473684e-09 | 1646 | tags=50%, list=9%, signal=46% |
| 2.40078125e-09 | 1.53659539473684e-09 | 2166 | tags=41%, list=12%, signal=37% |
| 2.40078125e-09 | 1.53659539473684e-09 | 2144 | tags=44%, list=12%, signal=40% |
| 2.40078125e-09 | 1.53659539473684e-09 | 1825 | tags=42%, list=10%, signal=39% |
| 2.40078125e-09 | 1.53659539473684e-09 | 2111 | tags=45%, list=12%, signal=40% |
| 2.40078125e-09 | 1.53659539473684e-09 | 2166 | tags=42%, list=12%, signal=38% |
| 2.40078125e-09 | 1.53659539473684e-09 | 2166 | tags=42%, list=12%, signal=38% |
| 2.40078125e-09 | 1.53659539473684e-09 | 2122 | tags=40%, list=12%, signal=35% |
| 2.40078125e-09 | 1.53659539473684e-09 | 1646 | tags=41%, list=9%, signal=38% |
| 2.40078125e-09 | 1.53659539473684e-09 | 2206 | tags=41%, list=12%, signal=36% |
| 2.40078125e-09 | 1.53659539473684e-09 | 2023 | tags=44%, list=11%, signal=39% |
| 2.40078125e-09 | 1.53659539473684e-09 | 2206 | tags=40%, list=12%, signal=36% |
| 2.40078125e-09 | 1.53659539473684e-09 | 1646 | tags=54%, list=9%, signal=49% |
| 2.40078125e-09 | 1.53659539473684e-09 | 1646 | tags=54%, list=9%, signal=49% |
| 2.40078125e-09 | 1.53659539473684e-09 | 2206 | tags=40%, list=12%, signal=36% |
| 2.40078125e-09 | 1.53659539473684e-09 | 2162 | tags=48%, list=12%, signal=42% |
| 2.40078125e-09 | 1.53659539473684e-09 | 1646 | tags=43%, list=9%, signal=40% |
| 2.40078125e-09 | 1.53659539473684e-09 | 2118 | tags=39%, list=12%, signal=35% |
| 2.40078125e-09 | 1.53659539473684e-09 | 1987 | tags=48%, list=11%, signal=43% |
| 2.40078125e-09 | 1.53659539473684e-09 | 2118 | tags=39%, list=12%, signal=35% |
| 2.40078125e-09 | 1.53659539473684e-09 | 2025 | tags=49%, list=11%, signal=44% |
| 2.40078125e-09 | 1.53659539473684e-09 | 1282 | tags=44%, list=7%, signal=42% |
| 2.40078125e-09 | 1.53659539473684e-09 | 1211 | tags=58%, list=7%, signal=54% |
| 2.40078125e-09 | 1.53659539473684e-09 | 2025 | tags=50%, list=11%, signal=45% |
| 2.40078125e-09 | 1.53659539473684e-09 | 2025 | tags=43%, list=11%, signal=39% |
| 2.40078125e-09 | 1.53659539473684e-09 | 2025 | tags=47%, list=11%, signal=42% |
| 2.40078125e-09 | 1.53659539473684e-09 | 2127 | tags=48%, list=12%, signal=43% |
| 2.40078125e-09 | 1.53659539473684e-09 | 2025 | tags=48%, list=11%, signal=43% |

| core_enrichment |
| --- |
| RPP25/FBLL1/ZNF593/WDR12/NOC2L/GNL2/BYSL/NOP53/WDR3/RBFA/EIF4A3/EBNA1BP2/MRTO4/DHX37/GEMIN4/NPM3/URB1/BOP1/PRKDC/GTF3A/EM |
| RPP25/FBLL1/WDR12/BYSL/NOP53/WDR3/RBFA/EIF4A3/EBNA1BP2/MRTO4/DHX37/GEMIN4/NPM3/URB1/BOP1/PRKDC/EMG1/RIOK1/RRS1/EXOSC10/M |
| RPP25/FBLL1/ZNF593/HSP90AB1/WDR12/NOC2L/EIF3B/GNL2/BYSL/SNRPB2/EIF3E/NOP53/WDR3/RBFA/EIF4A3/EBNA1BP2/MRTO4/DHX37/GEMIN4/NPM |
| UQCRH/CCNB1/CYCS/CHCHD2/SNCA/NDUFA4/NDUFS5/NDUFAB1/NDUFS6/NDUFA5/COX7A1/COX4I1/NDUFB9/NDUFS7/PINK1/SDHB/NDUFA1/NDU |
| UQCRH/CCNB1/CYCS/CHCHD2/SNCA/NDUFA4/NDUFS5/NDUFAB1/NDUFS6/NDUFA5/COX7A1/COX4I1/NDUFB9/NDUFS7/PINK1/SDHB/NDUFA1/NDU |
| RPP25/FBLL1/WDR12/BYSL/NOP53/WDR3/RBFA/EIF4A3/POLR1D/EBNA1BP2/MRTO4/DHX37/GEMIN4/NPM3/URB1/BOP1/PRKDC/GTF3A/EMG1/MTOR/R |
| UQCRH/CYCS/SNCA/NDUFA4/NDUFS5/NDUFAB1/NDUFS6/NDUFA5/COX7A1/COX4I1/NDUFB9/NDUFS7/PINK1/SDHB/NDUFA1/NDUFB10/DNAJC15/C |
| ABCD1/UQCRH/CCNB1/CYCS/CHCHD2/SLC25A23/SNCA/NDUFA4/NIPSNAP2/NDUFS5/SLC25A33/SHMT2/NDUFAB1/NDUFS6/NDUFA13/NDUFA5/COX |
| TDRD12/TERT/TDRD9/RPP25/FBLL1/YRDC/WDR12/BYSL/POP1/PUS7/NOP53/WDR3/RBFA/GPAT2/NSUN2/EIF4A3/EBNA1BP2/MRTO4/DHX37/ZMPSTE2 |
| ABCD1/UQCRH/CCNB1/CYCS/CHCHD2/MDH2/SLC25A23/SNCA/CS/NDUFA4/NIPSNAP2/NDUFS5/NOP53/SLC25A33/SHMT2/MFN2/NDUFAB1/ADSL/ND |
| KIF2C/CCNB1/CDCA8/PINX1/PSRC1/NUF2/AURKB/MAD1L1/KIFC1/NDC80/CDCA5/KIF18A/RRS1/KIF14/BOD1/BIRC5/SKA3/CHAMP1/CENPE/SKA2/SPDL1 |
| WDR12/NOC2L/NOP53/EBNA1BP2/MRTO4/URB1/BOP1/GTF3A/RRS1/PAK1IP1/MAK16/RPF1/EIF6/PES1/NHP2/NLE1/BRIX1/NOP2/PPAN/MALSU1/ZNHIT6/ |
| CHCHD4/COX20/NDUFS5/SLC25A33/NDUFAB1/NDUFA13/NDUFA5/FASTKD3/NDUFAF5/NDUFB9/SAMM50/NDUFS7/NDUFAF2/COA1/NDUFA1/NDUFB |
| KIF25/KIF2C/CENPF/CDC20/NEK2/KIF4A/TRIP13/CCNB1/UBE2C/CDCA8/PINX1/PSRC1/NUF2/AURKB/NUSAP1/NCAPG/TPX2/MAD2L2/MAD1L1/PLK1/PRC |
| UQCRH/CCNB1/CYCS/CHCHD2/PPARGC1A/SNCA/NDUFA4/NDUFS5/NDUFAB1/NDUFS6/NDUFA5/COX7A1/COX4I1/NDUFB9/SLC25A13/NDUFS7/PINK |
| AARS2/RPP25/YRDC/POP1/PUS7/NSUN2/POLR3D/CARS2/TRMT12/TSEN34/TYW1B/NARS2/VARS2/POLR2E/TSEN15/EXOSC10/DALRD3/TRMT6/TARBP |
| AARS2/TBRG4/SLC25A33/SHMT2/TUFM/AURKAIP1/FASTKD3/DAP3/LRPPRC/DARS2/TARS2/EARS2/FASTK/RPUSD3/MALSU1/WARS2/MTERF3/NDUFA |
| BYSL/WDR3/DHX37/PRKDC/EMG1/RIOK1/RRS1/EXOSC10/DCAF13/NOP56/UTP11/DIMT1/DNTTIP2/RPP40/NOL7/UTP15/SNU13/RRP9/HEATR1/RRP36/T |
| GNAI1/INSC/GPSM2/CDC20/NEK2/KIF4A/CCNB1/CDCA8/RCC1/PSRC1/NUF2/GPSM1/AURKB/NUSAP1/CENPA/TPX2/STIL/PLK1/PRC1/KIFC1/WDR62/DL |
| KIF2C/CENPF/FAM83D/NEK2/CCNB1/CDCA8/PINX1/PSRC1/NUF2/AURKB/MAD1L1/KIFC1/KNSTRN/SPC24/SPC25/NDC80/CDCA5/SPAG5/KIF18A/RRS1/ |
| KIF25/KIF2C/CENPF/CDC20/NEK2/KIF4A/TRIP13/CCNB1/UBE2C/CDCA8/PINX1/PSRC1/NUF2/AURKB/NUSAP1/NCAPG/TPX2/MAD2L2/SMARCD3/MAD1L |
| NDUFS5/NDUFAB1/NDUFS6/NDUFA13/NDUFA5/NDUFB9/NDUFS7/ATP5PB/ATP5F1B/SDHB/NDUFA1/NDUFB10/ATP5MF/SDHA/NDUFB7/ATP5F1C/ATP |
| GNAI1/GPSM2/CDC20/NEK2/KIF4A/CCNB1/CDCA8/RCC1/PSRC1/NUF2/AURKB/TPX2/STIL/PLK1/PRC1/KIFC1/WDR62/DLGAP5/AURKA/SPC25/NDC80/KI |
| HORMAD1/KIF2C/CENPF/CDC20/TRIP13/CCNB1/UBE2C/CDCA8/NUF2/AURKB/NCAPG/MAD2L2/SMARCD3/MAD1L1/CDCA2/PLK1/SPC24/DLGAP5/BUB |
| SNCA/NDUFA4/NDUFS5/NDUFAB1/NDUFS6/NDUFA5/NDUFB9/NDUFS7/PINK1/NDUFA1/NDUFB10/DNAJC15/NDUFB7/NDUFC2/NDUFA8/NDUFA7/ND |
| KIF2C/NEK2/CCNB1/CDCA8/NUF2/AURKB/MAD1L1/KNSTRN/SPC24/SPC25/NDC80/SPAG5/BOD1/BIRC5/SKA3/RCC2/CHAMP1/CENPE/PMF1/KNL1/SKA |
| ATP5MC3/NDUFS5/NDUFAB1/NDUFS6/NDUFA13/NDUFA5/NDUFB9/NDUFS7/ATP5PB/ATP5F1B/SDHB/NDUFA1/NDUFB10/ATP5MF/SDHA/NDUFB7/AT |
| ABCD1/UQCRH/CCNB1/OPN3/CHCHD4/CYCS/CHCHD2/MDH2/PPARGC1A/SLC25A23/SNCA/CS/NDUFA4/NIPSNAP2/NDUFS5/NOP53/SLC25A33/SHMT |
| BMP7/CENPF/CDC20/TRIP13/CCNB1/CDCA8/NUF2/AURKB/MAD2L2/MAD1L1/PLK1/SPC24/BUB1/AURKAIP1/SPC25/NDC80/TOM1L2/MAD2L1/BIRC5/BUB |
| RPP25/YRDC/POP1/PUS7/NSUN2/TRMT12/TSEN34/TYW1B/TSEN15/DALRD3/TRMT6/TARBP1/TRMT9B/ANKRD16/AKT1/DTWD2/DDX1/LAGE3/ELP2/WD |
| FZD9/SLC25A4/SLC25A6/SNCA/TOMM20/TOMM40/TIMM13/MFN2/CHCHD3/BCL2/NDUFA13/SAMM50/HSPA9/MTX2/PINK1/TIMM10/AFG3L2/STPG1/DNA |
| NDUFS5/NDUFAB1/NDUFA13/NDUFA5/NDUFAF5/NDUFB9/NDUFS7/NDUFAF2/COA1/NDUFA1/NDUFB10/NDUFB7/NDUFC2/NDUFA8/NDUFS2/NDUF |
| NDUFS5/NDUFAB1/NDUFA13/NDUFA5/NDUFAF5/NDUFB9/NDUFS7/NDUFAF2/COA1/NDUFA1/NDUFB10/NDUFB7/NDUFC2/NDUFA8/NDUFS2/NDUF |
| HORMAD1/KIF25/KIF2C/CENPF/FAM83D/GPSM2/CDC20/MTCL1/NEK2/KIF4A/TRIP13/CCNB1/UBE2C/CDCA8/PINX1/RCC1/PSRC1/DIAPH3/NUF2/AURKB |
| CENPF/CDC20/TRIP13/CCNB1/CDCA8/PINX1/NUF2/AURKB/MAD2L2/MAD1L1/PLK1/SPC24/BUB1/SPC25/NDC80/ESPL1/EXOSC10/TOP2A/MAD2L1/BIRC5 |
| HFM1/RUVBL2/WRNIP1/DSCC1/RAD51/GINS1/TOP2A/CDC45/DDX1/RAD50/RAD54B/PIF1/GINS4/GINS3/MCM4/RFC3/BRIP1/DHX9/RFC2/TWNK/RFC4/W |
| KIF25/BMP7/KIF2C/CCDC8/CENPF/CDC20/NEK2/PKMYT1/EDN3/KIF4A/TRIP13/CCNB1/UBE2C/CDCA8/PINX1/RCC1/PSRC1/LRP5/NUF2/AURKB/ANLN/N |
| CENPF/CDC20/TRIP13/CCNB1/UBE2C/CDCA8/NUF2/AURKB/MAD2L2/SMARCD3/MAD1L1/PLK1/SPC24/DLGAP5/BUB1/CDC6/SPC25/NDC80/ESPL1/CDC |
| KIF2C/CENPF/FAM83D/NEK2/CCNB1/CDCA8/PINX1/PSRC1/NUF2/AURKB/MAD1L1/KIFC1/KNSTRN/SPC24/SPC25/NDC80/CDCA5/SPAG5/KIF18A/RRS1/ |
| HORMAD1/CENPF/CDC20/TRIP13/CCNB1/CDCA8/CLSPN/NUF2/AURKB/MAD2L2/MAD1L1/PLK1/GTSE1/CDK2/NOP53/NSUN2/SPC24/BUB1/CDC6/SPC2 |
| KIF2C/CENPF/FAM83D/NEK2/CCNB1/CDCA8/PINX1/PSRC1/NUF2/AURKB/MAD1L1/KIFC1/KNSTRN/SPC24/DLGAP5/SPC25/NDC80/CDCA5/SPAG5/KIF18 |
| CENPF/CDC20/TRIP13/CCNB1/UBE2C/CDCA8/NUF2/AURKB/MAD2L2/SMARCD3/MAD1L1/PLK1/SPC24/DLGAP5/BUB1/CDC6/SPC25/NDC80/ESPL1/CDC |
| HFM1/RUVBL2/WRNIP1/DSCC1/RAD51/GINS1/TOP2A/HNRNPA2B1/CDC45/DDX1/RAD50/RAD54B/PIF1/GINS4/GINS3/MCM4/RFC3/BRIP1/DHX9/RFC2/T |
| AARS2/SHMT2/TUFM/AURKAIP1/FASTKD3/DAP3/LRPPRC/DARS2/TARS2/EARS2/RPUSD3/MALSU1/WARS2/NDUFA7/PTCD1/C1QBP/UQCC2/MTIF2/R |

KIF2C/CDCA8/NUF2/AURKB/MAD1L1/NDC80/BOD1/BIRC5/SKA3/CHAMP1/CENPE/SKA2/SEH1L/SKA1/RMDN1/CDT1/INCENP/MAPRE1

BYSL/WDR3/DHX37/RIOK1/RRS1/DCAF13/RPP40/NOL7/SNU13/HEATR1/RRP36/TSR3/NOL10/ABT1/TSR2/GTF2H5/RIOK2/WDR43/NOL11/WDR46/UTP23/

CENPF/CDC20/TRIP13/CCNB1/UBE2C/CDCA8/NUF2/AURKB/MAD2L2/MAD1L1/PLK1/SPC24/DLGAP5/BUB1/SPC25/NDC80/ESPL1/CDC16/MAD2L1/BIRC5

AHNAK2/ZC3H13/PPARGC1A/YBX1/SNRPB2/NONO/PUS7/EIF4A3/RBPMS2/MBNL2/RBM11/GEMIN4/TSEN34/RBM28/LSM4/SNRPE/GEMIN8/SNRPD2/TSE

CENPF/CDC20/TRIP13/CCNB1/UBE2C/CDCA8/NUF2/AURKB/MAD2L2/MAD1L1/PLK1/SPC24/DLGAP5/BUB1/SPC25/NDC80/ESPL1/CDC16/MAD2L1/BIRC5

HSP90AB1/EIF3B/SNRPB2/EIF3E/NOP53/GEMIN4/BOP1/PRKDC/NUFIP1/RUVBL2/LSM4/SNRPE/RRS1/GEMIN8/EIF3D/SNRPD2/GEMIN5/EIF3H/SRPK3/TX

POLD2/CDK2/ENDOG/CDC6/ZMPSTE24/WRNIP1/LIG3/TONSL/DSCC1/RAD51/GINS1/CDK2AP1/POLE4/DNAJA3/MGME1/CIZ1/CDC45/CENPS/GMNN/RA

FZD9/SLC25A4/SLC25A15/HSPD1/SLC25A6/CHCHD4/VDAC1/SLC25A23/MRS2/TOMM20/TOMM40/TIMM13/SLC25A33/BCL2/NDUFA13/SAMM50/SLC25A1

CENPF/CDC20/TRIP13/CCNB1/UBE2C/CDCA8/NUF2/AURKB/NCAPG/MAD2L2/MAD1L1/PLK1/SPC24/DLGAP5/BUB1/SPC25/NDC80/ESPL1/CDC16/MAD2L

WDR12/URB1/BOP1/PRKDC/RRS1/EXOSC10/MAK16/RPF1/EIF6/ERI3/PES1/EXOSC8/RPP40/LAS1L/RRP15/ERI2/EXOSC4/NSA2/ABT1/EXOSC2/MTREX/N

AARS2/TBRG4/SLC25A33/FASTKD3/LRPPRC/DARS2/TARS2/EARS2/FASTK/WARS2/MTERF3/TFB2M/FASTKD2/MTERF2/TWNK/FASTKD1/SARS2/POLR

CENPF/CDC20/TRIP13/CCNB1/CDCA8/CLSPN/NUF2/AURKB/MAD2L2/MAD1L1/PLK1/GTSE1/CDK2/NOP53/SPC24/BUB1/CDC6/SPC25/NDC80/PRKDC/M

CENPF/CDC20/TRIP13/CCNB1/CDCA8/NUF2/AURKB/MAD2L2/MAD1L1/PLK1/SPC24/BUB1/SPC25/NDC80/MAD2L1/BIRC5/BUB1B/KNL1/SPDL1/MAD2L1B

CENPF/CDC20/TRIP13/CCNB1/CDCA8/NUF2/AURKB/MAD2L2/MAD1L1/PLK1/SPC24/BUB1/SPC25/NDC80/MAD2L1/BIRC5/BUB1B/KNL1/SPDL1/MAD2L1B

CENPF/CDC20/TRIP13/CCNB1/CDCA8/NUF2/AURKB/MAD2L2/MAD1L1/PLK1/SPC24/BUB1/SPC25/NDC80/MAD2L1/BIRC5/BUB1B/KNL1/SPDL1/MAD2L1B

CENPF/CDC20/TRIP13/CCNB1/CDCA8/NUF2/AURKB/MAD2L2/MAD1L1/PLK1/SPC24/BUB1/SPC25/NDC80/MAD2L1/BIRC5/BUB1B/KNL1/SPDL1/MAD2L1B

HSP90AB1/EIF3B/SNRPB2/EIF3E/NOP53/DYRK3/GEMIN4/BOP1/PRKDC/NUFIP1/RUVBL2/LSM4/SNRPE/RRS1/GEMIN8/EIF3D/SNRPD2/GEMIN5/EIF3H/S

TLE6/KIF25/INSC/GPSM2/SPIRE2/GPSM1/NUSAP1/CENPA/ASPM/PLK1/SPRY2/NDC80/PAX6/ESPL1/SPAG5/MAD2L1/ZBED3/SPDL1/DYNC1H1/NDEL1/NS

CENPF/CDC20/TRIP13/CCNB1/CDCA8/NUF2/AURKB/MAD2L2/MAD1L1/PLK1/SPC24/BUB1/SPC25/NDC80/MAD2L1/BIRC5/BUB1B/KNL1/SPDL1/MAD2L1B

CENPF/CDC20/TRIP13/CCNB1/CDCA8/NUF2/AURKB/MAD2L2/MAD1L1/PLK1/SPC24/BUB1/SPC25/NDC80/MAD2L1/BIRC5/BUB1B/KNL1/SPDL1/MAD2L1B

CENPF/CDC20/TRIP13/CCNB1/CDCA8/NUF2/AURKB/MAD2L2/MAD1L1/PLK1/SPC24/BUB1/SPC25/NDC80/MAD2L1/BIRC5/BUB1B/KNL1/SPDL1/MAD2L1B

CENPF/CDC20/TRIP13/CCNB1/CDCA8/NUF2/AURKB/MAD2L2/MAD1L1/PLK1/SPC24/BUB1/SPC25/NDC80/MAD2L1/BIRC5/BUB1B/KNL1/SPDL1/MAD2L1B

TERT/POLD2/RUVBL2/LIG3/GINS1/RAD50/MCM4/RFC3/DCLRE1B/TFPT/NUCKS1/RFC4/POLE/MCM3/PCNA/UCHL5/MRE11/MCM7

CENPF/CDC20/TRIP13/CCNB1/CDCA8/NUF2/AURKB/MAD2L2/MAD1L1/PLK1/SPC24/BUB1/SPC25/NDC80/MAD2L1/BIRC5/BUB1B/KNL1/SPDL1/MAD2L1B

HORMAD1/KIF25/KIF2C/CENPF/FAM83D/CDC20/NEK2/KIF4A/TRIP13/CCNB1/UBE2C/CDCA8/PINX1/PSRC1/NUF2/AURKB/NUSAP1/NCAPG/TPX2/ASPM/

CENPF/CDC20/TRIP13/CCNB1/UBE2C/CDCA8/NUF2/AURKB/NCAPG/MAD2L2/MAD1L1/PLK1/SPC24/DLGAP5/BUB1/SPC25/NDC80/ESPL1/CDC16/MAD2L

IMPDH2/CTPS1/ENO1/PPARGC1A/NME4/NME1/ATP5MC3/CAD/NDUFS5/UCK2/NDUFAB1/NDUFS6/NDUFA13/NDUFA5/NDUFB9/SLC25A13/NDUFS7/AT

TOMM20/TOMM40/TIMM13/NDUFA13/SAMM50/MTX2/TIMM10/TIMM8A/TOMM22/ROMO1/HSPA4/MTCH2/TOMM6/PDCD5/MTX1/MTCH1/BCS1L/HSP90

NOP53/POLR1D/NPM3/GTF3A/MTOR/POLR2E/GTF3C5/SMARCA4/TP53/GTF3C1/TCOF1/GTF3C3/GTF2H5/PWP1/GTF3C2/NOL11/GTF3C6/NIFK/DEDD/

CENPF/CDC20/TRIP13/CCNB1/UBE2C/CDCA8/NUF2/AURKB/MAD2L2/SMARCD3/MAD1L1/PLK1/SPC24/DLGAP5/BUB1/CDC6/SPC25/NDC80/ESPL1/CDC

YRDC/PUS7/NSUN2/TRMT12/TYW1B/DALRD3/TRMT6/TARBP1/TRMT9B/ANKRD16/AKT1/DTWD2/ELP2/WDR4/PUSL1/URM1/DUS3L/MTFMT/GTPBP3/

FBLL1/ZC3H13/MTRR/NSUN2/ICMT/ZMPSTE24/TRMT12/EMG1/PAX6/SNRPE/DNMT3B/EEF1AKMT3/SNRPD2/BEND3/MRM1/DALRD3/TRMT6/TARBP1/

AARS2/CARS2/NARS2/VARS2/DALRD3/FARSB/DARS2/LRRC47/TARS2/EARS2/WARS2/AARSD1/SARS2/FARSA/RARS2/IARS2/YARS2

GNAI1/GPSM2/TPPP/CDC20/NEK2/KIF4A/CCNB1/CDCA8/RCC1/PSRC1/NUF2/AURKB/TPX2/ASPM/STIL/PLK1/PRC1/KIFC1/AUNIP/KNSTRN/WDR62/DLG

CSAG1/KIF25/SLC16A1/NEK2/CCNF/RTTN/CROCC/STIL/PLK1/CDK2/WDR62/AURKA/KIF3A/NDC80/CEP85/CEP250/CENPJ/ODF2/PLK2/UXT/ARL2/CEP72/

CENPF/CDC20/TRIP13/CCNB1/UBE2C/CDCA8/NUF2/AURKB/MAD2L2/SMARCD3/MAD1L1/PLK1/SPC24/DLGAP5/BUB1/CDC6/SPC25/NDC80/ESPL1/CDC

GEMIN4/LSM4/SNRPE/GEMIN8/SNRPD2/GEMIN5/SNRPD1/PRMT5/PRPF6/AAR2/WDR77/SNRPC/GEMIN6/LSM2/DDX20/SNRPB/SNRPG/PRPF3/CLNS1A

CCNB1/CDCA8/AURKB/NCAPG/CDC6/BIRC5/RCC2/RAD18/MAD2L1BP/CDK1/NCAPG2/NCAPH/NCAPD2/SIRT2/SMC6/INCENP/SMC2/KAT5/NCAPD3/N

CENPF/CDC20/TRIP13/CCNB1/UBE2C/CDCA8/NUF2/AURKB/MAD2L2/SMARCD3/MAD1L1/PLK1/SPC24/DLGAP5/BUB1/CDC6/SPC25/NDC80/ESPL1/CDC

HFM1/RUVBL2/WRNIP1/DSCC1/RAD51/GINS1/TOP2A/TOP1MT/HNRNPA2B1/CDC45/DDX1/RAD50/RAD54B/PIF1/GINS4/GINS3/MCM4/RFC3/BRIP1/TOP

TIMM13/CHCHD3/NDUFA13/SAMM50/HSPA9/MTX2/PINK1/TIMM10/AFG3L2/DNAJC11/MICOS10/TIMM8A/ROMO1/LETM1/APOOL/MTX1/APOO/UQCC3/M

CENPF/CDC20/TRIP13/CCNB1/CDCA8/NUF2/AURKB/MAD2L2/MAD1L1/PLK1/SPC24/BUB1/SPC25/NDC80/MAD2L1/BIRC5/BUB1B/KNL1/SPDL1/MAD2L1B

CENPF/CDC20/TRIP13/CCNB1/CDCA8/NUF2/AURKB/MAD2L2/MAD1L1/PLK1/SPC24/BUB1/SPC25/NDC80/MAD2L1/BIRC5/BUB1B/KNL1/SPDL1/MAD2L1B

CENPF/CDC20/TRIP13/CCNB1/CDCA8/NUF2/AURKB/MAD2L2/MAD1L1/PLK1/SPC24/BUB1/SPC25/NDC80/MAD2L1/BIRC5/BUB1B/KNL1/SPDL1/MAD2L1B

TLE6/KIF25/INSC/GPSM2/SPIRE2/GPSM1/NUSAP1/CENPA/PLK1/SPRY2/NDC80/PAX6/ESPL1/SPAG5/MAD2L1/ZBED3/SPDL1/DYNC1H1/NDEL1/NSFL1C/S

NOP53/DHX37/BOP1/PRKDC/RRS1/EIF6/NLE1/BRIX1/NOP2/PPAN/MTERF3/C1QBP/SBDS/TRAF7/DHX29/FASTKD2/EIF5B/MRM2/ABT1/DHX30/RPF2/XRC

DHX33/NOP53/POLR1D/ZMPSTE24/MTOR/POLR1C/POLR2E/BEND3/POLR1A/MYBBP1A/CARM1/SMARCA4/PRR7/POLR2L/UTP15/DEK/HEATR1/TCOF1/

CENPF/CENPA/ITGB3BP/DLGAP5/OIP5/CENPI/HJURP/CENPE/CENPV/SUGT1/CENPN/CENPX/MIS18A/RNF4/POGZ/CENPO/CENPH/CENPW/KNTC1/TR

WDR12/URB1/BOP1/MAK16/RPF1/EIF6/PES1/NHP2/NOP2/ZNHIT6/LAS1L/RRP15/SNU13/NSA2/DDX18/RPF2/NIFK/NOL9/FTSJ3

IMPDH2/CTPS1/ENO1/PPARGC1A/NME4/NME1/ATP5MC3/CAD/NDUFS5/UCK2/TYMS/NDUFAB1/NDUFS6/NDUFA13/NDUFA5/NDUFB9/SLC25A13/NDUF

TERT/CCT3/CCT6A/RUVBL2/DKC1/NABP2/CCT8/CCT4/CCT2/LRWD1/TCP1/SPIDR

SNRPB2/EIF4A3/RBPMS2/RBM11/GEMIN4/LSM4/SNRPE/GEMIN8/SNRPD2/EXOSC10/GEMIN5/SRPK3/DAZAP1/HNRNPA2B1/TXNL4A/PQBP1/DDX1/SNRF

CSAG1/KIF25/SLC16A1/NEK2/CCNF/RTTN/CROCC/STIL/PLK1/CDK2/WDR62/AURKA/KIF3A/NDC80/CEP85/CEP250/CENPJ/ODF2/PLK2/UXT/ARL2/CEP72/

SNRPB2/EIF4A3/RBPMS2/RBM11/GEMIN4/LSM4/SNRPE/GEMIN8/SNRPD2/EXOSC10/GEMIN5/SRPK3/DAZAP1/HNRNPA2B1/TXNL4A/PQBP1/DDX1/SNRF

SNRPB2/EIF4A3/RBPMS2/RBM11/GEMIN4/LSM4/SNRPE/GEMIN8/SNRPD2/EXOSC10/GEMIN5/SRPK3/DAZAP1/HNRNPA2B1/TXNL4A/PQBP1/DDX1/SNRF

OCA2/TYRP1/CDH3/MFSD12/MC1R/PMEL/OPN3/RAB38/DDT/BCL2/DCT

ZC3H13/PPARGC1A/TBRG4/YBX1/SNRPB2/NONO/PUS7/AHCYL1/EIF4A3/RBPMS2/MBNL2/RBM11/SSU72/GEMIN4/ADARB1/TSEN34/RBM28/LSM4/SNRP

GET1/EMC1/NCLN/UBL4A/MMGT1/NOMO1/EMC2/EMC7/CCDC47/EMC8/TRAM1/GET3/SGTA/BAG6/TMCO1/EMC10/EMC4/WDR83OS/RAB5IF/EMC3/TR

BYSL/DHX37/RRS1/DCAF13/RPP40/HEATR1/RRP36/NOL10/ABT1/TSR2/GTF2H5/WDR43/WDR46/UTP23/UTP25/TSR1/ERCC2/NOP14/UTP20/TBL3/NGDN

CENPF/CDC20/NEK2/TRIP13/CCNB1/MYC/UBE2C/CDCA8/PINX1/NUF2/AURKB/NCAPG/MAD2L2/SMARCD3/MAD1L1/CCT3/PLK1/CDK2/SPC24/DLGAP5/

FBLL1/ZC3H13/NSUN2/TRMT12/EMG1/SNRPE/SNRPD2/MRM1/DALRD3/TRMT6/TARBP1/TRMT9B/AKT1/SNRPD1/WDR4/NOP2/DIMT1/NSUN5/GTPBP3/

POLD2/CDK2/NPM2/CHAF1B/NT5M/ENDOG/CDC6/ZMPSTE24/RUVBL2/WRNIP1/LIG3/RNASEH2A/TONSL/DSCC1/RAD51/GINS1/ESCO2/CDK2AP1/POLE

NOP53/BOP1/RRS1/EIF6/NLE1/BRIX1/NOP2/PPAN/TRAF7/FASTKD2/MRM2/DHX30/RPF2

HORMAD1/BMP7/CENPF/CDC20/TRIP13/CCNB1/CDCA8/NUF2/AURKB/MAD2L2/MAD1L1/PLK1/SPC24/BUB1/AURKAIP1/SPC25/NDC80/TOM1L2/MAD2L1/

MAOB/NOX4/UQCRH/CCNB1/CYCS/CHCHD2/PPARGC1A/SNCA/NDUFA4/NDUFS5/GSR/NDUFAB1/NDUFS6/NDUFA5/COX7A1/COX4I1/NDUFB9/SLC25

IMPDH2/ENO1/PPARGC1A/NME4/NME1/ATP5MC3/NDUFS5/NDUFAB1/NDUFS6/NDUFA13/NDUFA5/NDUFB9/SLC25A13/NDUFS7/ATP5PB/ATP5F1B/PIN

TERT/PINX1/AURKB/CENPA/CCT3/PLK1/GNL3/CCT6A/RUVBL2/TONSL/ESCO2/BOD1/DKC1/BUB1B/RCC2/CHAMP1/TNKS/NABP2/CCT8/CCT4/GNL3L/KN

IMPDH2/ENO1/PPARGC1A/NME4/NME1/ATP5MC3/NDUFS5/NDUFAB1/NDUFS6/NDUFA13/NDUFA5/NDUFB9/SLC25A13/NDUFS7/ATP5PB/ATP5F1B/PIN

TOMM20/TOMM40/SAMM50/MTX2/TOMM22/HSPA4/MTCH2/TOMM6/PDCD5/MTX1/MTCH1/HSP90AA1/TOMM5/TOMM70

TOMM20/TOMM40/SAMM50/MTX2/TOMM22/HSPA4/MTCH2/TOMM6/PDCD5/MTX1/MTCH1/HSP90AA1/TOMM5/TOMM70

AARS2/CARS2/NARS2/VARS2/DALRD3/FARSB/DARS2/LRRC47/TARS2/EARS2/WARS2/AARSD1/SARS2/FARSA/RARS2/IARS2/YARS2/GATC

TERT/PINX1/CCT3/GNL3/CCT6A/DKC1/TNKS/NABP2/CCT8/CCT4/GNL3L/CCT2

OCA2/TYRP1/CDH3/MFSD12/MC1R/PMEL/OPN3/RAB38/DDT/DCT

OCA2/TYRP1/CDH3/MFSD12/MC1R/PMEL/OPN3/RAB38/DDT/DCT

CENPF/NUF2/CENPA/DLGAP5/NDC80/CENPE/SUGT1/CENPX/RNF4/POGZ/SMC2/CENPH/CENPW/KNTC1/TRAPPC12/CENPK

HSPD1/CHCHD4/TOMM20/TOMM40/TIMM13/MIPEP/MFN2/NDUFA13/HPS4/SAMM50/TSPO/MTX2/PINK1/TIMM10/PAM16/DNAJC15/GRPEL1/HAX1/UBE2

OCA2/TYRP1/TPCN2/SNAI2/CDH3/MFSD12/EDNRB/MC1R/KIT/PMEL/ABCB6/EDN3/RAB32/SHROOM3/BACE2/MITF/RACK1/RAB38/BCL2/HPS4/MYO5A/U

AARS2/CARS2/NARS2/VARS2/DALRD3/FARSB/DARS2/LRRC47/TARS2/EARS2/WARS2/AARSD1/SARS2/FARSA/RARS2/IARS2/YARS2/GATC

CCT3/GNL3/CCT6A/DKC1/CCT8/CCT4/GNL3L/CCT2/TCP1

CDK2/CDC6/WRNIP1/CIZ1/CDC45/GMNN/CCNE1/CDC34/GINS3/MCM4/LRWD1/MCM3/MCM10/ORC5/ORC1/CDT1/MCM7/ORC4/CCNE2/ORC6/MCIDA

ICMT/EEF1AKMT3/PCMT1/SMYD2/PRMT5/EHMT2/EEF1AKMT1/GSPT1/SNRPB/PRMT1/METTL21C/SETD4/LCMT1/TRMT112/PRMT6/RAB6A/ETFBKMT/E

ICMT/EEF1AKMT3/PCMT1/SMYD2/PRMT5/EHMT2/EEF1AKMT1/GSPT1/SNRPB/PRMT1/METTL21C/SETD4/LCMT1/TRMT112/PRMT6/RAB6A/ETFBKMT/E

OCA2/TYRP1/CDH3/MFSD12/MC1R/PMEL/OPN3/RAB38/DDT/TSPO/DCT/ATP5IF1/GPR143/TMEM14B/FLVCR1/ALAD/SLC45A2/CPOX/ALAS1/PPOX/ABCB

CCNF/RTTN/STIL/CDK2/WDR62/NDC80/CEP85/CENPJ/PLK2/CEP72/CNTROB/CKAP5/CEP131/CETN2/CHMP1B/CHMP2A/CHMP3/SAC3D1/SASS6/BRCA

TDRD12/HORMAD1/MSX2/KIF25/TDRD9/BMP7/KIF2C/CCDC8/CENPF/CDC20/NEK2/PKMYT1/EDN3/SPIRE2/KIF4A/TRIP13/CCNB1/UBE2C/CDCA8/PDE3

HSPD1/CHCHD4/TOMM20/TOMM40/TIMM13/MIPEP/MFN2/NDUFA13/HPS4/SAMM50/TSPO/MTX2/PINK1/TIMM10/PAM16/AKT1/DNAJC15/GRPEL1/HAX1/

ENO1/PPARGC1A/ATP5MC3/NDUFS5/NDUFAB1/NDUFS6/NDUFA13/NDUFA5/NDUFB9/SLC25A13/NDUFS7/ATP5PB/ATP5F1B/PINK1/LDHC/SDHB/NDUF

CDC20/UBE2C/CDCA8/AURKB/DLGAP5/ESPL1/CDC16/BIRC5/MAD2L1BP/ANAPC11

FBLL1/EMG1/MRM1/DKC1/NHP2/NOP2/DIMT1/RPUSD1/NSUN5/TFB2M/NSUN4/MRM2/TSR3/GAR1/TRMT112/RPUSD2/NAT10/TRMT61B/FBL/FTSJ3

SLC25A4/SLC25A15/HSPD1/SLC25A6/CHCHD4/VDAC1/SLC25A23/MRS2/TOMM20/TOMM40/SLC25A33/SLC25A13/PSEN2/MICU3/AFG3L2/PAM16/SLC25A2

FBLL1/AHCYL1/SSU72/EXOSC10/DKC1/ERI3/ZNF473/EXOSC8/PTCD1/INTS6/EXOSC5/ERI2/EXOSC4/EXOSC2/EXOSC6/ELAC2/POLR2D/INTS14/PARN/LS

EYA1/FOXM1/SPIRE2/TRIP13/MAD2L2/SMARCD3/MORF4L2/PLK1/RAD54L/AUNIP/UBQLN4/HSF2BP/MEIOB/TWIST1/FANCB/PRKDC/RUVBL2/CDCA5/LIG

CCT3/GNL3/CCT6A/DKC1/CCT8/CCT4/GNL3L/CCT2/TCP1

TERT/FZD9/WIPI1/SLC25A4/RAB32/ATAD3A/HSPD1/SLC25A6/SPATA18/CHCHD4/COX20/CHCHD2/PPARGC1A/VDAC1/PYCARD/SNCA/MTFP1/RAB3A/R

HSPD1/CHCHD4/TOMM20/TOMM40/TIMM13/MIPEP/MFN2/NDUFA13/HPS4/SAMM50/TSPO/MTX2/PINK1/TIMM10/PAM16/AKT1/DNAJC15/GRPEL1/HAX1/

CYCS/NDUFA4/COX7A1/COX4I1/COX5B/COX7B/COX7C/COX6B1/COX6C/COX6A1/COX8A

TDRD12/HORMAD1/MSX2/KIF25/TDRD9/BMP7/KIF2C/CCDC8/CENPF/CDC20/NEK2/PKMYT1/EDN3/SPIRE2/KIF4A/TRIP13/CCNB1/UBE2C/CDCA8/PDE3

OCA2/TYRP1/EDNRB/KIT/EDN3/MITF/BCL2/HPS4/USP13/ENPP1/GNA11/SOX10/RAB27A

AURKB/CENPA/BOD1/BUB1B/RCC2/CHAMP1/KNL1/SPDL1/CDK1/MTBP/TTK/ZW10/CENPQ/HASPIN/KNTC1/TRAPPC12/RB1

VDAC1/SLC25A23/TOMM20/TOMM40/SLC25A33/PSEN2/MICU3/AFG3L2/PAM16/DNAJC15/GRPEL1/TIMM50/MCUR1/ROMO1/SLC25A32/MICU2/TIMM17B/

MAD2L2/MORF4L2/RAD54L/AUNIP/UBQLN4/MEIOB/FANCB/RUVBL2/LIG3/AP5S1/TONSL/RAD51/PSMD14/CDC45/NABP2/RAD54B/HUS1/FEN1/SENP3/GI

TRIP13/HSF2BP/AURKA/MEIOB/NDC80/CCNB2/CDC25B

CLSPN/PLK1/GTSE1/CDK2/NOP53/CDC6/PRKDC/RAD51/TIPRL/PLK2/CDC45/CLOCK/FEM1B/SETMAR/CCNG1/CDC5L/HUS1/EME1/DTL/BRIP1/TP53/MS

COX20/SLC25A33/FASTKD3/COA1/HCCS/COA6/LYRM7/UQCC2/PET117/TTC19/TIMM21/TMEM223/COX19/UQCC3/COA5/BCS1L/PET100/COA4/SLC25A46

ABCD1/UQCRH/CCNB1/OPN3/IRS2/CHCHD4/CYCS/CHCHD2/MDH2/PPARGC1A/SLC25A23/SNCA/CS/NDUFA4/NIPSNAP2/NDUFS5/NOP53/GFPT2/SLC

CLSPN/PLK1/NOP53/CDC6/RAD50/NABP2/CCNG1/HUS1/DTL/RINT1/CDK1/RBBP8/INTS3/CDC14B/ORC1/BLM/MRE11/MBTPS1/BRCA1/BABAM1/CHEK1/

BMP7/CCDC8/CENPF/CDC20/NEK2/PKMYT1/EDN3/TRIP13/CCNB1/UBE2C/CDCA8/RCC1/LRP5/NUF2/AURKB/NUSAP1/MAD2L2/MAD1L1/CDCA2/PLK1/

VPS37D/LAPTM4B/SORT1/VPS36/PTPN23/CHMP1B/TSG101/CHMP2A/CHMP3/VPS37B/LEPROT/SNF8/DENND10/UBXN6/VPS37A/STAM/MVB12A/TMEM5

OCA2/TYRP1/EDNRB/KIT/EDN3/RAB32/MITF/RAB38/BCL2/HPS4/USP13/ENPP1/GNA11/SOX10/AP3D1/RAB27A

TOMM20/TOMM40/PAM16/DNAJC15/GRPEL1/TIMM50/ROMO1/TIMM17B/TIMM21/TIMM44/TOMM20L/TIMM17A/TOMM70/TOMM40L

NEK2/CCNB1/CDCA8/AURKB/KNSTRN/SPAG5/BIRC5/RCC2/ECT2/SIRT2/INCENP/ZW10/KNTC1/KAT5

UQCRH/CYCS/UQCRB/CYC1/UQCRC1/UQCR11/UQCRC2/UQCC3/UQCRQ

MAD2L2/MORF4L2/RAD54L/AUNIP/UBQLN4/MEIOB/FANCB/RUVBL2/LIG3/AP5S1/TONSL/RAD51/PSMD14/CDC45/NABP2/RAD54B/HUS1/FEN1/SENP3/GI

WDR12/URB1/BOP1/RRS1/EXOSC10/ERI3/PES1/EXOSC8/RPP40/ERI2/ABT1/EXOSC2/NOL9/FTSJ3/NOP14/ERI1/UTP20

TERT/CCT3/CCT6A/DKC1/NABP2/CCT8/CCT4/CCT2/TCP1

EYA1/FOXM1/SPIRE2/MAD2L2/SMARCD3/MORF4L2/AUNIP/UBQLN4/TWIST1/FANCB/PRKDC/RUVBL2/RAD51/SUPT3H/BCL7C/USP22/POLH/SETMAR/C

HSPD1/CHCHD4/PEX6/TOMM20/TOMM40/PAM16/DNAJC15/GRPEL1/TIMM50/PEX10/ROMO1/TIMM17B/TIMM21/AIFM1/TIMM44/TOMM20L/LONP2/TIMM

INSC/GPSM2/GPSM1/NUSAP1/CENPA/PLK1/SPRY2/NDC80/PAX6/ESPL1/MAD2L1/SPDL1/NDEL1/NSFL1C/CLASP2/CLASP1/ZW10/MAPRE1/FBXW11/NDE

RAPGEF4/RAB3B/LPAR1/CDK5R1/ARHGDIA/PLD1/BSN

P3H3/P3H2/CRTAP/PLOD3/PLOD1/PLOD2/OGFOD1/P3H4/P4HA2/P3H1/P4HA3/P4HB/EGLN1/JMJD4

CCT3/CCT6A/DKC1/CCT8/CCT4/CCT2/TCP1/SPIDR

DHX33/NOP53/ZMPSTE24/MTOR/BEND3/MYBBP1A/CARM1/SMARCA4/PRR7/POLR2L/UTP15/DEK/HEATR1/BAZ1B/WDR43/PWP1/NOL11/POLR2K/LYAR/

RAD51/GINS1/CDC45/GINS4/GINS3/MCM4/TWNK/WRN/MCM3/BLM/MCM7/HMGA1/RECQL4/SSBP1

CTPS1/SHMT1/CAD/GART/PAICS/DHODH/MTOR/CTPS2/PPAT/CMPK1/PRPS1/APRT/CPS1

OCA2/TYRP1/TPCN2/EDNRB/KIT/EDN3/RAB32/MITF/RAB38/BCL2/HPS4/USP13/DCT/ENPP1/GPR143/GNA11/SOX10/AP3D1/RAB27A

TCEA3/POLR1D/GTF2F2/POLR2E/INTS11/ZNF326/ELOA/ELP2/NELFE/CDK13/MED25/SOX10/ADRM1/INTS6/PBRM1/INTS3/NELFCD/POLR2I/INTS13/GTF2

TERT/NEK2/MYC/PINX1/HSP90AB1/AURKB/MAD2L2/CCT3/EXO1/GNL3/CCT6A/PRKDC/RUVBL2/RAD51/EXOSC10/DKC1/HNRNPA2B1/NHP2/RAD50/TNK

TYRP1/CDH3/PMEL/OPN3/RAB38

TYRP1/CDH3/PMEL/OPN3/RAB38

CCT3/NOP53/CCT6A/DKC1/CCT8/CCT4/CCT2/TCP1

CHCHD4/PPARGC1A/ENDOG/LIG3/DNAJA3/MGME1/LONP1/TP53/TWNK/RRM1/TOP3A/MPV17/SSBP1/SESN2/PARP1/DNA2

TYRP1/CDH3/PMEL/ABCB6/RAB32/SHROOM3/BACE2/RAB38/BCL2/HPS4/MYO5A/MLPH/RAB17/GPR143/AP3D1/RAB27A/DCTN2/MKKS/ANKRD27/AP1S1/

CENPF/CENPA/DLGAP5/CENPE/SUGT1/CENPX/RNF4/POGZ/CENPH/CENPW/KNTC1/TRAPPC12/CENPK

UBAP1L/VPS37D/VPS36/PTPN23/HDAC6/CHMP1B/TSG101/CHMP2A/RNF126/CHMP3/VPS37B/SNF8/VPS37A/TOM1L1/STAM/MVB12A/PLAA/VPS25/VPS28/

CLSPN/PLK1/GTSE1/CDK2/NOP53/PRKDC/RAD51/TIPRL/PLK2/CLOCK/FEM1B/CCNG1/CDC5L/HUS1/EME1/DTL/BRIP1/TP53/MSH2/FANCD2/RINT1/CDK

TERT/NEK2/MYC/PINX1/HSP90AB1/AURKB/MAD2L2/CCT3/EXO1/GNL3/CCT6A/PRKDC/RUVBL2/RAD51/EXOSC10/DKC1/HNRNPA2B1/NHP2/RAD50/TNK

CHCHD3/SAMM50/PINK1/AFG3L2/DNAJC11/MICOS10/LETM1/APOOL/APOO/UQCC3/MICOS13/SLC25A46/IMMT

EXTL2/TGDS/GFPT2/AMDHD2/GNPDA1/UAP1L1/GFPT1/PMM1/UAP1/PMM2/NANS/GMDS/UGGT1/GNPNAT1/UGGT2/ENTPD5/DPAGT1/UGDH/UGP2/SL(

COX20/FASTKD3/COA1/COA6/PET117/TIMM21/TMEM223/COX19/COA5/BCS1L/PET100/COA4/SLC25A46/SCO1/STMP1/COX10/COA8/COA3/COX16/TACO

AK5/PDK4/ADCY2/IMPDH2/CTPS1/ENO1/PPARGC1A/NME4/NME1/SNCA/ATP5MC3/CAD/NDUFS5/UCK2/ACSS2/NDUFAB1/GART/PAICS/ADSL/NDUFS6/

HORMAD1/CDC20/PLK1/BUB1/CDCA5/ESPL1/DSCC1/ESCO2/BOD1/BUB1B/NAA10/TNKS/FEN1/SMC1A/GTF2B/SGO2/RAD51C/SGO1/RAD21/ESCO1/KIF2

AK5/PDK4/ADCY2/IMPDH2/CTPS1/ENO1/PPARGC1A/NME4/NME1/SNCA/ATP5MC3/CAD/NDUFS5/UCK2/ACSS2/NDUFAB1/GART/PAICS/ADSL/NDUFS6/

HORMAD1/MSX2/BMP7/CCDC8/CENPF/CDC20/NEK2/PKMYT1/EDN3/TRIP13/CCNB1/UBE2C/CDCA8/PDE3A/RCC1/LRP5/NUF2/AURKB/NUSAP1/MAD2L

CCNF/STIL/CENPJ/PLK2/CEP131/CHMP1B/CHMP2A/CHMP3/SASS6/BRCA1/PPP1R35/NAT10/GEN1/ROCK2/CEP295NL/PLK4/CHMP1A/NUP62/C10orf90/C

SLC25A4/CHCHD4/PPARGC1A/SLC25A33/ENDOG/LIG3/DNAJA3/MGME1/PIF1/LONP1/TP53/TWNK/RRM1/TOP3A/MPV17/SSBP1/AKT3/SESN2/PARP1/DN/

TYRP1/CDH3/MFSD12/PMEL/OPN3/RAB38

TYRP1/CDH3/MFSD12/PMEL/OPN3/RAB38

TYRP1/CDH3/MFSD12/PMEL/OPN3/RAB38

CCT3/CCT6A/RUVBL2/DKC1/NHP2/CCT8/CCT4/CCT2/TCP1

CCT3/CCT6A/RUVBL2/EXOSC10/DKC1/NHP2/CCT8/CCT4/CCT2/TCP1

SNRPB2/SNRPE/SNRPD2/SRPK3/TXNL4A/DDX1/SNRPD1/PRPF6/SNRPC/GEMIN6/PTBP2/NOL3/GCFC2/SNRPB/SNRPG/SF3B4/SF3A3/SF3B6/DDX39B/SI

WIPI1/SLC25A4/SPATA18/VDAC1/RACK1/AURKA/MFN2/DYRK3/KIF9/PLAAT1/TSPO/PINK1/CTTN/RNF14/MAP1LC3B/HAX1/SLC25A5/ATP5IF1/ATG4B/DDR

FANCB/FAAP20/RAD51/CENPS/CENPX/FANCI/FANCD2/DCLRE1B/NUCKS1/MCM8/FANCL/FANCE/FAAP24/FANCA/FANCC/EXO5/ERCC1/FAN1/FAAP10

CCT3/CCT6A/RUVBL2/EXOSC10/DKC1/NHP2/CCT8/CCT4/CCT2/TCP1

CCT3/CCT6A/RUVBL2/EXOSC10/DKC1/NHP2/CCT8/CCT4/CCT2/TCP1

CCT3/CCT6A/RUVBL2/EXOSC10/DKC1/NHP2/CCT8/CCT4/CCT2/TCP1

CCT3/CCT6A/RUVBL2/EXOSC10/DKC1/NHP2/CCT8/CCT4/CCT2/TCP1

CCNF/STIL/WDR62/AURKA/CENPJ/PLK2/CEP131/CHMP1B/CCNL2/CHMP2A/CHMP3/SASS6/MARK4/BRCA1/PPP1R35/NAT10/GEN1/ROCK2/CEP295NL/P

CDCA8/AURKB/MAD1L1/NDC80/MAD2L1/BIRC5/KNL1/TPR/INCENP/DYNC1LI1/GEN1

POLD2/MAD2L2/FAAP20/POLH/RAD18/USP43/DTL/MSH2/UBE2V1/POLDIP2/PCNA/POLD1/BRCA1/UBE2A/REV3L/PCLAF/SPRTN/USP1/POLI/USP10/RCHY

CDC20/NEK2/KIF4A/CDCA8/AURKB/TPX2/PLK1/PRC1/KIFC1/KIF23/BIRC5/MZT1/SMC1A/RANGRF/POLDIP2/CHMP1B/CHMP2A/CHMP3/UHRF1/GOLGA2/

GET1/EMC1/UBL4A/MMGT1/EMC2/EMC7/EMC8/GET3/SGTA/BAG6/EMC10/EMC4/EMC3

MYH10/IQGAP3/SPIRE2/KIF4A/SEPTIN5/CDCA8/CECR2/AURKB/ANLN/NUSAP1/CENPA/SEPTIN3/PLK1/CUL7/CEP55/BIN3/SEPTIN4/SEPTIN10/ESPL1/KIF2

SOX8/FOXD1/MYC/SOX9

MGP/BMPR1B/SOX9/BMP1/COL11A1/SOX6/COL2A1/OTOR/SOX5

CDC20/UBE2C/CDCA8/AURKB/NCAPG/DLGAP5/ESPL1/CDC16/BIRC5/MAD2L1BP/ANAPC11/NCAPG2/NCAPH/NCAPD2/INCENP/SMC2/NSMCE2/NCAPD

GPSM2/CDC20/NEK2/KIF4A/CDCA8/RCC1/AURKB/TPX2/ASPM/STIL/PLK1/PRC1/KIFC1/AURKA/NDC80/TUBB/SPAG5/CCNB2/KIF23/BIRC5/DCAF13/CENF

AK5/PDK4/ADCY2/IMPDH2/ENO1/PPARGC1A/NME4/NME1/SNCA/ATP5MC3/NDUFS5/ACSS2/NDUFAB1/GART/PAICS/ADSL/NDUFS6/NDUFA13/NDUFA5

SHMT2/FASTKD3/LRPPRC/RPUSD3/MALSU1/C1QBP/UQCC2/RMND1/RCC1L/FASTKD2/TRUB2/CDK5RAP1/MTG2/METTL8/CHCHD10/RPUSD4/COA3/F

COX20/FASTKD3/COA1/PET117/TIMM21/TMEM223/COX19/COA5/BCS1L/PET100/COA4/SCO1/STMP1/COA8/COA3/COX16/TACO1/SURF1/COX18/COX14/S

DDRGK1/ATG9A/SNX7/ULK2/ULK1/UFC1/RETREG2/SNX30/UFM1/ULK3/ATG2A/TEX264/RB1CC1

AARS2/YRDC/VARS2/TARS2/AARSD1/DTD1/IARS2/DTD2/GATC/CDK5RAP1/LARS2/DNAJC2

IMPDH2/GART/PAICS/ADSL/ATIC/PFAS/PPAT

EXO1/RNASEH2C/RNASEH2A/MSH5/MSH2/MSH3/MCM8/MSH6/PCNA/RPA3/ABL1/PMS2/RPA1/RPA2/LIG1

KIF16B/MTMR2/SNX12/PTPN23/DNAJC13/MAPK1/CHMP3/MAP2K2/PXK/ANKRD27/SNF8/DENND10/RDX/HOOK2/DAB2/FLNA/RAB5A/WDR81/BECN1/WD

GART/PAICS/ADSL/ATIC/PFAS/PPAT

RIOK1/YTHDF2/DIMT1/USP36/UTP15/HEATR1/RIOK2/WDR43/TRMT112/WDR75/BUD23/METTL18

POLD2/LIG3/GINS1/MCM4/RFC3/RFC4/POLE/MCM3/PCNA/MCM7/LIG1/DNA2

RACK1/RNF14/DENR/MTIF2/ZNF598/PTRH1/EIF2D/GCN1/GFM2/EIF4E2/MCTS1/TRNT1/LTN1/TCF25/RNF25/USP10/MTIF3/RCHY1/PELO/GIGYF2/HBS1L

CLSPN/PLK1/NOP53/CCNG1/DTL/RINT1/CDK1/CDC14B/BLM/MRE11/MBTPS1/BRCA1/BABAM1/CHEK1/IER3/DONSON/TAOK3/MBTPS2/BABAM2/SYF2/AE

FKBP10/HSPD1/HSP90AB1/ST13/CHCHD4/CCT3/TXNDC5/FKBP9/CRTAP/DNAJB6/CANX/CCT6A/PPIA/RUVBL2/HSPA9/DFFA/ANP32E/HSPA6/DNAJA3/PT
SHMT1/MTRR/TYMS/SHMT2/GART/GGH/ATIC/MTHFD1L/PRKG2/SPR/QDPR/AASDHPPT/SLC19A1/SLC25A32/DHFR
TBRG4/FASTKD3/FASTK/FASTKD2/FASTKD1/TRIT1/ELAC2/TRNT1/HSD17B10/TRMT61B/CDK5RAP1/PUS1/FASTKD5/METTL8/RPUSD4/MTPAP
KCNK2/ENO1/CHCHD2/EPHA4/PINK1/DNMT3A/NOL3/USP19
MAGEC2/CDC20/UBE2C/PLK1/PINK1/ARRDC4/UBE2L3/CDC14B/BMI1
WIPI1/SEPTIN5/TMED2/SEC16B/NLGN1/YKT6/TRAPPC2L/TRAPPC3/TMED9/SCRIB/PEF1/EXOC6B/FAM91A1/CLASP2/PDCD6/CLASP1/TRAPPC5/STARD3
NSUN2/TRMT12/DALRD3/TRMT6/TARBP1/TRMT9B/AKT1/WDR4/GTPBP3/TYW3/THUMPD3/FTSJ1/METTL1/TRMT112/METTL2A/TRMT61A/HSD17B10/TI
RANBP17/CHCHD4/NUP93/CHP1/AHCYL1/NSUN2/EIF4A3/PTPN14/XPO5/HSPA9/CCHCR1/HNRNPA2B1/EIF6/GLE1/DDX25/EMD/YWHAE/XPO4/NXT1/UBI
GINS1/GINS4/PALB2/SBDS/ZPR1/NCAPG2/SALL4/NDEL1/INTS1/BRCA2/CHEK1/COPS2/PELO/TAF8
VPS37D/VPS36/PTPN23/TSG101/VPS37B/LEPROT/SNF8/VPS37A/STAM/MVB12A/TMEM50A/VPS25/VTA1/VPS28/CHMP6/HGS/VPS37C/LEPROTL1/CHMP4E
FBLL1/ZC3H13/YRDC/PUS7/NSUN2/ADARB1/TRMT12/EMG1/SNRPE/TYW1B/SNRPD2/MRM1/DALRD3/TRMT6/TARBP1/DKC1/TRMT9B/ANKRD16/AKT1/
KIF25/INSC/GPSM2/GPSM1/CENPA/PLK1/SPRY2/NDC80/PAX6/SPAG5/MAD2L1/SPDL1/NDEL1/NSFL1C/CLASP1/LLGL1/MAP4/ZW10/MAPRE1/FBXW11/ND
TRMT9B/ELP2/URM1/GTPBP3/CTU1/DPH3/ELP5/ELP3/ELP6/CTU2/MOCS3/ELP1/KTI12
CDCA8/AURKB/MAD1L1/NDC80/MAD2L1/BIRC5/TPR/INCENP/DYNC1LI1/GEN1
POLR3G/POLR1D/POLR3D/GTF3A/MTOR/POLR1C/POLR2E/IVNS1ABP/GTF3C5/SNAPC2/POLR3K/MYBBP1A/RO60/POLR3H/POLR2L/DEK/GTF3C1/PRDX
CDC20/BUB1/CDCA5/DSCC1/ESCO2/BOD1/NAA10/TNKS/SMC1A/SGO1/RAD21/ESCO1/PDS5B/POGZ/HASPIN/NSMCE2/MAU2/RB1
SSU72/ZMPSTE24/TTF2/XRN2/POLR3K/PRMT5/MAZ/DHX9/MTERF2/MED18/ZC3H4/TTF1/POLR2A/MTERF1/CAVIN1
RCC1/NUP93/PLK1/LMNB2/ZMPSTE24/PRKCA/EMD/UBXN2A/REEP4/UBE2I/BANF1/TMEM43/CTDNEP1/CHMP1B/CHMP2A/LEMD2/CHMP3/NSFL1C/LMN
CLSPN/PLK1/GTSE1/CDK2/NOP53/CDC6/PRKDC/PLK2/SETMAR/CCNG1/HUS1/EME1/DTL/TP53/MSH2/FANCD2/RINT1/CDK1/CDC14B/ORC1/BLM/MRE1
MYO16/CENPF/CDC20/PKMYT1/TRIP13/CCNB1/CDCA8/CDKN2B/CLSPN/NUF2/AURKB/MAD2L2/MAD1L1/PLK1/GTSE1/CDK2/NOP53/SPC24/BUB1/CDC6
MYH10/IQGAP3/KIF4A/CDCA8/AURKB/ANLN/NUSAP1/CENPA/PLK1/CUL7/CEP55/ESPL1/KIF23/CKAP2/CIT/BIRC5/KIF20A/ECT2/CNTROB/ARF1/EFHC1/PC
MYO16/HORMAD1/KIF25/BMP7/CENPF/CDC20/NEK2/PKMYT1/TRIP13/CCNB1/CCNF/CDCA8/CDKN2B/CLSPN/NUF2/AURKB/MAD2L2/MAD1L1/PLK1/DAB
TRMT9B/ELP2/URM1/GTPBP3/CTU1/DPH3/ADAT2/ELP5/ELP3/ELP6/CTU2/MOCS3/ELP1/KTI12
AURKB/RCC2/CHAMP1/KNL1/SPDL1/CDK1/MTBP/TTK/ZW10/CENPQ/KNTC1/TRAPPC12
AURKB/RCC2/CHAMP1/KNL1/SPDL1/CDK1/MTBP/TTK/ZW10/CENPQ/KNTC1/TRAPPC12
RTTN/STIL/CDK2/WDR62/CEP85/CENPJ/PLK2/CEP72/CNTROB/CETN2/CHMP2A/SASS6/BRCA1/PPP1R35/CEP152/KIAA0753/CEP295NL/PLK4/CCP110/NU
OCA2/TYRP1/CDH3/MFSD12/MC1R/PMEL/ABCB6/OPN3/ABCC2/RAB38/DDT/BCL2/TSPO/DCT/HMOX2/ATP5IF1/GPR143/TMEM14B/FLVCR1/ALAD/SLC45
MYO16/HORMAD1/CENPF/CDC20/PKMYT1/TRIP13/CCNB1/CDCA8/CDKN2B/CLSPN/NUF2/AURKB/MAD2L2/MAD1L1/PLK1/GTSE1/CDK2/NOP53/NSUN2/
HNRNPA2B1/RBM10/RBM20/C1QBP/PTBP2/SAP18/RNPS1/HNRNPK/TMBIM6/SRSF12/RBM42/PTBP1/SRSF9/SRSF10/RBMX/U2AF2/DYRK1A
SHMT2/FASTKD3/LRPPRC/RPUSD3/MALSU1/C1QBP/UQCC2/RMND1/RCC1L/FASTKD2/TRUB2/CDK5RAP1/MTG2/METTL8/RPUSD4/COA3/MPV17L2/TA
EIF3B/EIF3E/MTOR/EIF3D/EIF3H/YTHDF2/EIF4A1/DENR/EIF3G/EIF2B3/EIF3J/DHX29/EIF2B4/EIF2D/EIF2AK4/EIF3I
MYO1B/WIPI1/SPIRE2/KDELR2/CYTH3/RACK1/GOLPH3L/SORT1/SNX8/TMED2/RAB6B/COG6/COPZ1/MYO5A/KDELR3/COPA/KIF16B/AP4M1/YIF1B/SEC16
CCNB1/CDCA8/AURKB/MAD1L1/NDC80/RAD51/MAD2L1/BIRC5/FEM1B/KNL1/MAD2L1BP/ZWINT/CRY1/LCMT1/WDR76/BRCA1/TPR/CUL4A/INCENP/CCA
CDC20/UBE2C/DLGAP5/ESPL1/CDC16/MAD2L1BP/ANAPC11
KIF4A/CDCA8/AURKB/PRC1/KIF23/BIRC5
AARS2/EEF2/WTIP/BZW2/KRT17/SHMT1/SOX4/EIF3B/EEF1G/EIF4EBP1/YBX1/EIF3E/RACK1/RGS2/PUS7/DHX33/EPHA4/EIF4A3/TMED2/TYMS/SHMT2/TU
VPS37D/VPS36/RAB27A/CHMP1B/TSG101/CHMP2A/CHMP3/VPS37B/SNF8/VPS37A/STAM/MVB12A/VPS25/VTA1/VPS28/CHMP6/CHMP1A/HGS/VPS37C/CH
FBLL1/EXOSC10/DKC1/EXOSC5/EXOSC4/EXOSC2/EXOSC6/PARN/FBL
EIF3B/EIF3E/EIF3D/EIF3H/EIF3G/EIF3J/EIF3I
NOP53/POLR1D/MTOR/POLR2E/SMARCA4/TCOF1/GTF2H5/PWP1/NOL11/DEDD/SMARCB1/ERCC2/PIH1D1/TAF1B/DDX11
INSC/GPSM2/GPSM1/CENPA/PLK1/SPRY2/NDC80/PAX6/MAD2L1/SPDL1/NDEL1/NSFL1C/ZW10/MAPRE1/FBXW11/NDE1/HDAC3/DCTN1/KAT5/ENKD1/SA
FBLL1/EXOSC10/DKC1/EXOSC5/EXOSC4/RNF113A/EXOSC2/EXOSC6/PARN/FBL
SNAI2/FOXM1/CLSPN/SPRED1/SOX4/MAD2L2/MIF/PLK1/GTSE1/CDK2/NOP53/PTTG1IP/DYRK3/TWIST1/ZMPSTE24/PRKDC/TFAP4/MYO6/PSMD10/RAD5
RTTN/STIL/CDK2/WDR62/CEP85/CENPJ/PLK2/CEP72/CNTROB/CETN2/CCDC78/CHMP2A/SASS6/BRCA1/PPP1R35/CEP152/MCIDAS/KIAA0753/CEP295N
KIF25/NEK2/AURKA/CEP85/CNTROB/NDEL1/NSFL1C/KIF11/NDE1/CHEK1

PKMYT1/CLSPN/AURKB/PLK1/NOP53/CDC6/RAD50/NABP2/MIIP/CCNG1/HUS1/DTL/RINT1/CDK1/RBBP8/ZFYVE19/INTS3/CDC14B/RAD21/ORC1/BLM/MF

GNAI1/GPSM2/PLK1

VPS37D/VPS36/PTPN23/CHMP1B/TSG101/CHMP2A/CHMP3/VPS37B/LEPROT/SNF8/VPS37A/STAM/MVB12A/TMEM50A/VPS25/VTA1/BECN1/VPS28/CHMF

VDAC1/MFN2/PINK1/ATP5IF1/MUL1/HDAC6/CDC37

FAM174B/PLK1/CUL7/GOLPH3L/TMED2/RAB6B/KIFC3/OBSL1/COG6/SEC16B/TANGO2/PRMT5/RBSN/TBC1D20/UBXN2A/ARHGEF7/GORASP2/SURF4/DY

CCT3/CCT6A/DKC1/CCT8/CCT4/CCT2/TCP1

CCT3/CCT6A/DKC1/CCT8/CCT4/CCT2/TCP1

EYA1/FOXM1/SPIRE2/MAD2L2/SMARCD3/MORF4L2/FANCB/PRKDC/RUVBL2/BCL7C/SETMAR/CBX8/TRIM28/SMARCA4/BCL7A/DHX9/PBRM1/SMARCC

FOXM1/SPIRE2/MAD2L2/SMARCD3/MORF4L2/AUNIP/UBQLN4/TWIST1/FANCB/PRKDC/RUVBL2/RAD51/BCL7C/SETMAR/SENP3/CYREN/SMARCA4/BCL

FAM83D/DVL1/STIL/AURKA/SPAG5/CEP250/KLHL21/CHAMP1/RAB11FIP3/CEP72/FAM83H/CEP131/DCTN2/CRIPT/NSFL1C/MARK4/NUDCD3/ABL1/BICD1/

CDCA8/AURKB/MAD1L1/SPAG5/BIRC5/TTL/DYNC1H1/INCENP/KAT5/BECN1/NUMA1

KDELR2/TMED2/KDELR3/COPA/YIF1B/SEC16B/CIDEB/YKT6/TRAPPC2L/ERGIC3/SEC31A/VAPB/SCAP/SEC13/CNIH4/TBC1D20/TRAPPC3/RINT1/TMED9/S

ROBO2/SNAI2/GATA4/MSX2/BMP7/TBX3/ADAMTS5/SOX9/DKK1/HEYL/HEY1/ERBB3/TWIST1

ABCB6/TSPO/ATP5IF1/RSAD1/TMEM14B/FLVCR1/ALAD/CPOX/ALAS1/PPOX/ABCB7/SLC25A39/SLC11A2/UROD/ABCB10/SUCLA2/COX10/PGRMC1/FXN/N

ABCB6/TSPO/ATP5IF1/RSAD1/TMEM14B/FLVCR1/ALAD/CPOX/ALAS1/PPOX/ABCB7/SLC25A39/SLC11A2/UROD/ABCB10/SUCLA2/COX10/PGRMC1/FXN/N

KDELR2/GOLPH3L/RAB6B/COPZ1/KDELR3/COPA/ERGIC3/RER1/RINT1/SEC22B/KDELR1/LMAN2/COPG2/SCYL1/USE1/RAB6A/ZW10/COPZ2/COG3/ARF4

GNAI1/GPSM2/TPPP/MAP6/PSRC1/DIAPH3/STMN3/TPX2/ATAT1/STIL/SNCA/PLK1/SPAG5/KIF18A/MAP2/WDR47/CKAP2/SKA3/CENPJ/CDK5R1/ARL2/RAC

KIF4A/CDCA8/AURKB/PRC1/KIF23/BIRC5

FBXO17/UBXN8/CANX/FOXRED2/USP13/RCN3/FBXO27/UBE2J2/UBE2G2/UBXN10/UBXN2A/STUB1/ERLIN2/FAF1/RHBDD2/USP19/UBE4B/FAM8A1/MAN1

RANBP17/NUP93/NSUN2/EIF4A3/HNRNPA2B1/LRPPRC/PEG10/GLE1/DDX25/AHCTF1/TNKS/KHSRP/NDC1/SEC13/NXT1/NXF5/DHX9/SMG5/CETN2/POM

FBLL1/EMG1/MRM1/NOP2/DIMT1/NSUN5/TFB2M/NSUN4/MRM2/TRMT112/TRMT61B/FBL/FTSJ3/BUD23

CENPF/PKMYT1/CCNB1/CLSPN/AURKB/SMARCD3/PLK1/CDK2/NOP53/AURKA/DYRK3/CDC6/CDC25C/KIF14/RCC2/RAD50/CDK4/NABP2/MIIP/CCNG1/H

NUP93/NSUN2/EIF4A3/XPO5/HNRNPA2B1/GLE1/DDX25/NXT1/NXF5/DHX9/SMG5/POM121C/MCM3AP/SARNP/RAE1/CASC3/DDX39B/POM121/SRSF3/RBI

MYH10/IQGAP3/SPIRE2/KIF4A/SEPTIN5/CDCA8/CECR2/AURKB/ANLN/NUSAP1/CENPA/SEPTIN3/PLK1/PRC1/CUL7/CEP55/AURKA/PDXP/BIN3/CDC6/SEF

TCEA3/GTF2F2/INTS11/ELOA/ELP2/NELFE/CDK13/MED25/SOX10/ADRM1/INTS6/PBRM1/INTS3/NELFCD/POLR2I/INTS13/MED15/MED18/WDR43/MED8/C

VPS37D/LAPTM4B/SORT1/VPS36/PTPN23/RAB27A/CHMP1B/TSG101/CHMP2A/CHMP3/VPS37B/LEPROT/SNF8/DENND10/UBXN6/VPS37A/STAM/MVB12A

EIF3B/EIF3D/PAIP1/DENR/EIF3G/PCBP2/DHX9/EIF2D/EIF2AK4

FBLL1/GAMT/ZC3H13/DPH5/MTRR/NSUN2/ICMT/TPMT/TYMS/ZMPSTE24/TRMT12/EMG1/PAX6/NDUFAF5/COMT/SNRPE/DNMT3B/EEF1AKMT3/SNRPD

VPS37D/VPS36/PTPN23/TSG101/VPS37B/SNF8/VPS37A/STAM/MVB12A/VPS25/VPS28/HGS/VPS37C

VPS37D/VPS36/CHMP1B/TSG101/CHMP2A/CHMP3/VPS37B/SNF8/VPS37A/STAM/MVB12A/VPS25/VTA1/VPS28/CHMP6/CHMP1A/HGS/VPS37C/CHMP4B/R

RANBP17/NUP93/CCT3/YBX1/TOMM20/NSUN2/EIF4A3/CCT6A/RUVBL2/XPO5/EXOSC10/DKC1/HNRNPA2B1/LRPPRC/PEG10/GLE1/DDX25/AHCTF1/NHP

WDR12/URB1/BOP1/PES1/ZNHIT6/DDX18/RPF2/NIFK/NOL9/FTSJ3/RBM34/GTPBP4

PKMYT1/CLSPN/AURKB/PLK1/NOP53/CDC6/RAD50/NABP2/MIIP/CCNG1/HUS1/DTL/RINT1/CDK1/RBBP8/ZFYVE19/INTS3/CDC14B/RAD21/ORC1/BLM/MF

TERT/PINX1/BYSL/NOP53/RRS1/POLR1A/WRN/RPF2/UTP25/NMD3/NVL/MCRS1

NECAB2/PCSK9/ANKRD13B/WDR54/LRPAP1/MTMR2

STIL/AURKA/SPAG5/CEP250/RAB11FIP3/CEP72/CEP131/DCTN2/NSFL1C/MARK4/NUDCD3/BICD1/PIBF1/MAPRE1/STK3/BBS4/KIAA0753/PCM1/CEP350/NU

SNRPB2/SNRPE/SNRPD2/SNRPD1/SNRPB/SNRPG/SF3B4/SF3A3/SF3B6/SNRPF/SNIP1/RBMX2/SF3B2/DDX42/SF3A2/SNRPD3/HTATSF1/SF3B3/BUD13

AK5/CTPS1/SLC19A2/SHMT1/NME4/NME1/CAD/UCK2/TYMS/DHODH/MTOR/CTPS2/NME2/TK1/DTYMK/CMPK1/SLC4A7/NME6/UCKL1/UCK1/DUT

TERT/NEK2/PINX1/HSP90AB1/AURKB/CCT3/CCT6A/RAD51/EXOSC10/DKC1/HNRNPA2B1/NHP2/RAD50/TNKS/CCT8/PIF1/CCT4/GNL3L/TP53/CCT2/DCLF

GINS1/GINS4/PALB2/SBDS/ZPR1/NCAPG2/SALL4/NDEL1/RTF1/INTS1/ACVR1C/BRCA2/CHEK1/COPS2/PELO/TAF8

GFPT2/AMDHD2/GNPDA1/UAP1L1/GFPT1/PMM1/UAP1/PMM2/NANS/GMDS/GNPNAT1/UGDH/GNPDA2/GMPPB/FCSK/PGM3/UXS1

GPAT4/GPAT2/AGPAT2/AGPAT1/AGPAT5/GPAM/CDS2/TAMM41/LCLAT1

GPAT4/GPAT2/AGPAT2/AGPAT1/AGPAT5/GPAM/CDS2/TAMM41/LCLAT1

POLR3D/POLR2E/GTF3C5/POLR2L/GTF3C1/GTF3C3/GTF3C2/GTF3C6/POLR2F/GTF3C4/POLR3E

RIOK1/DIMT1/UTP15/HEATR1/RIOK2/WDR43/TRMT112/WDR75/BUD23

OCA2/TYRP1/CDH3/MFSD12/MC1R/PMEL/GPR37/OPN3/SNCA/RAB38/DDT/HAND2/DCT
MDH2/CS/SDHB/PDHA1/SDHA/SUCLG1/MDH1/IDH3A/DLAT/ACLY/FH/ACO2/NNT/OGDH/SUCLG2/PDHB/SUCLA2/IDH3B
IQGAP3/SPIRE2/AURKB/ANLN/CEP55/BIN3/KIF20A/ECT2/CNTROB/ARF1/CHMP1B/CHMP2A/CHMP3/SPIRE1
AK5/CTPS1/NME4/NME1/CAD/UCK2/DHODH/CTPS2/NME2/CMPK1/NME6/UCKL1/UCK1
CENPF/PKMYT1/CCNB1/CLSPN/AURKB/SMARCD3/PLK1/CDK2/NOP53/AURKA/CDC6/CDC25C/KIF14/RCC2/RAD50/CDK4/NABP2/MIIP/CCNG1/HUS1/DT
KIF2C/TPPP/NCKAP5/PSRC1/DIAPH3/STMN3/TPX2/SNCA/KIF18B/KIF18A/MAP2/WDR47/CKAP2/SKA3/CENPJ/CDK5R1/TUBG1/MZT1/ARL2/RAC1/ARHGE
TERT/CDC20/CCNB1/UBE2C/SMARCD3/STIL/DLGAP5/CDC6/CDC25C/CDCA5/ESPL1/CDC16/AKT1/CENPJ/RCC2/CDK4/CCNE1/TFDP1/DTL/CDC25B/M
RACK1/RNF14/ZNF598/PTRH1/GCN1/EIF4E2/TRNT1/LTN1/TCF25/RNF25/USP10/RCHY1/PELO/GIGYF2/HBS1L/MTRES1/ABCE1/ELAC1/NEMF/KLHDC10
CLSPN/PLK1/GTSE1/CDK2/NOP53/PRKDC/PLK2/CCNG1/HUS1/EME1/DTL/TP53/MSH2/FANCD2/RINT1/CDK1/CDC14B/BLM/MRE11/MBTPS1/BRCA1/CAS
GNAI1/GPSM2/PSRC1/TPX2/STIL/PLK1/CENPJ/PKD1/CHMP1B/DYNC1H1/CHMP2A/CHMP3/RAE1/TACC3/SASS6/TPR/CCSAP
TYRP1/PMEL/ABCB6/RAB32/BACE2/RAB38/BCL2/HPS4/GPR143/AP3D1/AP1S1/SNAPIN/ZEB2/KIF13A/BLOC1S6/DTNBP1/AP3B1/AP1M1
WIPI1/SLC25A4/SPATA18/VDAC1/MFN2/TSPO/PINK1/CTTN/MAP1LC3B/HAX1/SLC25A5/ATP5IF1/ATG4B/DNM1L/ATG9A/STUB1/SNX7/USP36/TP53/MUL1/H
PDK4/ACSS2/PDK3/ACSS1/PDHA1/DLAT/ACLY/PPCS/DLD/PDHB/DIP2A/MLYCD/PDHX/MPC2
KIF25/NEK2/AURKA/CEP85/NDEL1/NSFL1C/KIF11/NDE1/CHEK1
TPPP/PSRC1/DIAPH3/TPX2/SNCA/MAP2/CENPJ/CDK5R1/TUBG1/MZT1/ARL2/RAC1/ARHGEF7/PRUNE1/CKAP5/AKAP9/EML2/TBCD/RANBP9/NDEL1/CA
SMARCD3/NTHL1/RAD23A/BCL7C/HUS1/BRIP1/SMARCA4/TP53/BCL7A/GTF2H4/PBRM1/SMARCC1/CETN2/POLR2I/POLE/RAD52/RPA3/GTF2H5/SLX4/P
CDK2/WRNIP1/CIZ1/GMNN/MCM4/MCM3/CDT1/MCM7
STIL/AURKA/SPAG5/CEP250/RAB11FIP3/CEP72/CEP131/DCTN2/NSFL1C/MARK4/NUDCD3/BICD1/PIBF1/MAPRE1/STK3/BBS4/KIAA0753/PCM1/CEP350/N
GFPT2/NANP/AMDHD2/GNPDA1/UAP1L1/GFPT1/UAP1/GNPNAT1
TIMM13/NDUFA13/TIMM10/TIMM8A/ROMO1/TIMM9/TIMM29/TOMM70
WIPI1/SNX7/PIP4K2B/ULK1/PIP4K2C/SNX30/SH3GLB1/ATG2A/PIP4K2A/WDR45/SNX18/BECN1/RAB3GAP2/RALB
FAM83D/DVL1/STIL/AURKA/SPAG5/CEP250/CHAMP1/RAB11FIP3/CEP72/CEP131/DCTN2/CRIPT/NSFL1C/MARK4/NUDCD3/ABL1/BICD1/TTK/PIBF1/MAPR
MYO16/TERT/CENPF/CDC20/ID4/FOXM1/PKMYT1/IQGAP3/TRIP13/CCNB1/MYC/UBE2C/CCNF/CDCA8/RCC1/CDKN2B/CLSPN/NUF2/AURKB/ANLN/MA
EXOSC10/DAZAP1/RBM3/PRMT5/WDR77/RBM20/SNRNP70/PRDX6/HABP4/ZPR1/SF3B4/CLNS1A/THRAP3/UPF3A/EIF1/CIRBP/TRA2A/UPF1/NCBP1/UPF
POLR3D/POLR2E/GTF3C5/POLR2L/GTF3C1/GTF3C3/GTF3C2/GTF3C6/POLR2F/GTF3C4/POLR3E
RCC1/PLK1/PRKCA/EMD/UBXN2A/REEP4/UBE2I/BANF1/TMEM43/CTDNEP1/CHMP1B/CHMP2A/LEMD2/CHMP3/NSFL1C/SIRT2
CCNB1/CDCA8/AURKB/MAD1L1/NDC80/MAD2L1/BIRC5/KNL1/MAD2L1BP/ZWINT/LCMT1/TPR/INCENP/DYNC1LI1/GEN1/ANAPC15
HORMAD1/TRIP13/NSUN2/HUS1/KNL1/ZWINT
ALG6/DPM3/ALG14/ALG8/DPM2/ALG1/ALG3/MPDU1/DHDDS/DPAGT1/ALG9/DPM1/RFT1/SRD5A3/ALG2/DOLPP1/ALG12/NUS1/ALG13
AK5/CTPS1/SHMT1/NME4/NME1/CAD/UCK2/TYMS/DHODH/CTPS2/NME2/DTYMK/CMPK1/SLC4A7/NME6/UCKL1/UCK1/DUT
CENPF/FOXM1/PKMYT1/CCNB1/CLSPN/AURKB/SMARCD3/PLK1/CDK2/NOP53/MELK/AURKA/DYRK3/CDC6/NDC80/CDC25C/KIF14/CCNB2/RCC2/RAD5
CDCA8/AURKB/MAD1L1/NDC80/MAD2L1/BIRC5/KNL1/TPR/INCENP/CCAR2/DYNC1LI1/TELO2/GEN1
RAD51/FEM1B/CRY1/WDR76/BRCA1/CUL4A/CCAR2/RPA2/TELO2/FBXO4/BRCA2/BABAM2/TTI2/RFWD3/TTI1/ETAA1/PPP1R10/BRCC3/RNASEH2B
AARS2/EEF2/WTIP/BZW2/KRT17/SHMT1/SOX4/EIF3B/SLC1A1/EEF1G/EIF4EBP1/YBX1/EIF3E/RACK1/RGS2/PUS7/DHX33/EPHA4/EIF4A3/TMED2/TYMS/SH
HMG20B/HDAC4/GNL3/CAPN3/GNL3L/UBA2/UBE2I/SAE1/MUL1/HMG20A/TOLLIP/ARNT/CTNNB1/RWDD3
LPAR3/COBL/RGMA/FSTL4/DVL1/BDNF/SPART/PRKG1/PTPRS/HDAC6/ULK2/ULK1
CCNB1/CDCA8/AURKB/BIRC5/RCC2/SIRT2/INCENP/KAT5/BECN1
EXOSC10/DAZAP1/RBM3/PRMT5/WDR77/SNRNP70/PRDX6/SF3B4/CLNS1A/THRAP3/UPF3A/EIF1/CIRBP/TRA2A/UPF1/NCBP1/UPF3B/RBMX/PRPF19
DHX33/MTOR/MYBBP1A/CARM1/SMARCA4/UTP15/DEK/HEATR1/BAZ1B/WDR43/PWP1/NOL11/LYAR/SMARCB1/WDR75/ATF4/PIH1D1/ERBB2/DDX11/D
HNRNPA2B1/RBM10/RBM20/C1QBP/SAP18/RNPS1/HNRNPK/SRSF12/RBM42/PTBP1/SRSF9/SRSF10/RBMX/U2AF2/DYRK1A
FOXM1/SPIRE2/MAD2L2/SMARCD3/MORF4L2/FANCB/PRKDC/RUVBL2/BCL7C/SETMAR/SMARCA4/BCL7A/PBRM1/SMARCC1/EP400/MEAF6/MRGBP/S
UBE2C/ANLN/NPM2/CDKN1C/CDCA5/EPS8/MAD2L1BP/CHMP2A/CDC14B/SIRT2/PHB2/CLASP2/CLASP1/ZW10/KNTC1/NFIB
RCC1/UBXN2A/REEP4/UBE2I/BANF1/CHMP1B/CHMP2A/CHMP3/NSFL1C/SIRT2/ANKLE2/BROX/CHMP6/CHMP1A/CHMP4B/VPS4A/CHMP2B/CHMP7/SP
TYRP1/MAOB/NOX4/PGM1/ABCD1/SLC25A4/SLC4A4/PKLR/UQCRH/CCNB1/ENO2/OPN3/IRS2/GPI/ENO1/CHCHD4/CYCS/CHCHD2/MDH2/PPARGC1A/S
UBE2C/UBE2T/CDC16/UBE2D4/UBE2L5/UBE2E3/UBE2L3/ANAPC11/ANAPC13/ANAPC1/RNF4

THNSL2/BAAT/SHMT1/CTH/SHMT2/GART/SEPHS2/PSPH/MPST/CSAD/SCLY/PSAT1/RIDA/DHFR

CTPS1/NME4/NME1/CAD/UCK2/CTPS2/NME2/NME6/UCKL1/UCK1

PIGK/DPM3/SLC30A5/PIGF/PIGP/PIGZ/PIGW/PIGX/PGAP1/DPM2/PIGQ/PIGV/GPAA1/PIGU/PIGA/DPM1/PIGO/PIGB/PIGH/MPPE1/PIGS

PDK4/ABCD1/SOX9/IRS2/PPARGC1A/NR4A3/TWIST1/IRS1/AKT1/MTLN/FMO4/AKT2

KIF4A/CDCA8/AURKB/PRC1/KIF23/BIRC5

PSMG1/PSMD10/ADRM1/PSMG3/ECPAS/PSMD9/PSMD11/PSMG4/PSMG2/OTUD6B

SLC1A4/SLC6A17/SLC6A15/SLC7A5/SLC3A2

FBLL1/EXOSC10/DKC1/EXOSC5/EXOSC4/RNF113A/EXOSC2/EXOSC6/PARN/FBL

ZMPSTE24/RAD51/GINS1/CDC45/GINS3/MCM4/TK1/MCM3/RTF2/LIG1/BRCA2

TPPP/TPX2/CENPJ/TUBG1/MZT1/ARHGEF7/CKAP5/AKAP9/EML2/RANBP9/NDEL1/GOLGA2/CLASP2/MECP2/CLASP1/TUBGCP5/HAUS1/HAUS3/NDE1/DC

TYRP1/PMEL/ABCB6/RAB32/BACE2/RAB38/HPS4/GPR143/AP3D1/AP1S1/SNAPIN/ZEB2/KIF13A/BLOC1S6/DTNBP1/AP3B1/AP1M1

GNAI1/GPSM2/PSRC1/TPX2/STIL/PLK1/SPAG5/CENPJ/PKD1/CHMP1B/DYNC1H1/CHMP2A/CHMP3/RAE1/TACC3/SASS6/RNF4/TPR/CCSAP

INTS11/ZNF326/NELFE/CDK13/MED25/INTS6/INTS3/NELFCD/INTS13/MED15/MED18/HMGN1/WDR43/MED8/CTNNB1/SUPT16H/THOC5/MED20/SETD5/I

TONSL/RAD51/CENPS/RAD50/SETMAR/EME1/CARM1/CENPX/RBBP8/NUCKS1/WRN/PCNA/RTEL1/BLM/MRE11/ZNF365/BRCA1/TRAIP/EXD2/GEN1/MU

CDKN2B/SMARCD3/DAB2IP/BCL7C/SMARCA4/BCL7A/PBRM1/SMARCC1/RRM1/SMARCD2/RHNO1/BCL7B/SMARCB1/CHEK1/BRD7/ARID2/RRM2B/SM

CDKN2B/SMARCD3/DAB2IP/BCL7C/SMARCA4/BCL7A/PBRM1/SMARCC1/RRM1/SMARCD2/RHNO1/BCL7B/SMARCB1/CHEK1/BRD7/ARID2/RRM2B/SM

PUS7/DKC1/NHP2/PUSL1/RPUSD1/RPUSD3/TSR3/TRUB2/GAR1/RPUSD2/PUS1/TRUB1/RPUSD4/NOP10

SHMT1/MTRR/TYMS/SHMT2/GART/GGH/ATIC/MTHFD1L/AASDHPPT/SLC19A1/SLC25A32/DHFR

RAB6B/COG6/COPZ1/COPA/TRAPPC3/GOSR1/COPG2/COPG1/COPB1/RAB6A/NAPG/COPZ2/COG3/GOLGA5/COG2/COG4/GABARAPL2/COG1/COPB2/C

TMED2/SEC16B/CIDEB/TRAPPC2L/SEC31A/VAPB/SCAP/SEC13/TBC1D20/TRAPPC3/PEF1/PDCD6/VAPA/TRAPPC5/TRAPPC6A/SEC31B/TRAPPC1/TRAP

TERT/NEK2/PINX1/HSP90AB1/AURKB/CCT3/CCT6A/EXOSC10/DKC1/NHP2/RAD50/TNKS/CCT8/PIF1/CCT4/GNL3L/TP53/CCT2/SMG5/MAPK1/TCP1/RFC1

CCNB1/CDCA8/AURKB/MAD1L1/NDC80/MAD2L1/BIRC5/KNL1/MAD2L1BP/LCMT1/TPR/INCENP/DYNC1LI1/GEN1/ANAPC15

CCNB1/CDCA8/AURKB/MAD1L1/NDC80/MAD2L1/BIRC5/KNL1/MAD2L1BP/LCMT1/TPR/INCENP/DYNC1LI1/GEN1/ANAPC15

FASTKD3/RPUSD3/C1QBP/UQCC2/RMND1/RCC1L/FASTKD2/TRUB2/CDK5RAP1/METTL8/RPUSD4/COA3/MPV17L2/TRMT10C/NGRN

MGP/BMPR1B/SOX9/BMP1/COL11A1/SOX6/COL2A1/OTOR/SOX5

EEF2/EEF1G/RACK1/EIF4A3/TUFM/RNF14/EIF5A2/ZNF598/PTRH1/GCN1/EEF1B2/GFM1/EIF4E2/EEF2K/C6orf52/SRP9/TRNT1/LTN1/SECISBP2L/TCF25/R

CENPA/ITGB3BP/OIP5/CENPI/HJURP/CENPN/MIS18A

CENPA/ITGB3BP/OIP5/CENPI/HJURP/CENPN/MIS18A

NUSAP1/NCAPG/PLK1/CDCA5/NCAPH/NCAPD2/BAZ1B/PHF13/SMC2/NCAPD3/NCAPH2/CHMP1A/TENT4A/SMARCA5/AKAP8/KMT5A

VDAC1/SLC25A23/PSEN2/MICU3/AFG3L2/MCUR1/MICU2

MTMR2/SNX12/PTPN23/DNAJC13/MAPK1/CHMP3/MAP2K2/DENND10/RDX/DAB2

MYO16/TERT/CENPF/CDC20/PKMYT1/TRIP13/CCNB1/UBE2C/CDCA8/CDKN2B/CLSPN/NUF2/AURKB/ANLN/MAD2L2/SMARCD3/MAD1L1/STIL/PLK1/GT

NUP93/NSUN2/EIF4A3/HNRNPA2B1/GLE1/DDX25/NXT1/NXF5/SMG5/MCM3AP/SARNP/RAE1/CASC3/DDX39B/SRSF3/RBM8A/DDX19A/TPR/THOC5/NUP

TSPO/ATP5IF1/TMEM14B/FLVCR1/ALAD/CPOX/ALAS1/PPOX/ABCB7/SLC25A39/SLC11A2/UROD/ABCB10/COX10/PGRMC1/FXN/NFE2L1/TMEM14C/SLC6

TERT/CDC20/CCNB1/UBE2C/SMARCD3/STIL/EIF4EBP1/DLGAP5/AURKA/CDC6/CDC25C/CDCA5/ESPL1/CDC16/AKT1/PRKCA/CENPJ/USP22/RCC2/CD

MFN2/CHCHD3/PLD6/AFG3L2/MIGA1/MFF/MUL1/GDAP1/RCC1L/MTCH2/STOML2/VAT1

CENPF/FOXM1/PKMYT1/CCNB1/CLSPN/AURKB/SMARCD3/PLK1/CDK2/NOP53/MELK/AURKA/CDC6/CDC25C/KIF14/RCC2/RAD50/CDK4/NABP2/MIIP/C

ZMPSTE24/RAD51/GINS1/CDC45/GMNN/FEN1/GINS3/MCM4/TK1/WIZ/ZPR1/WRN/MCM3/PCNA/RTEL1/BLM/CDT1/RTF2/DBF4B/LIG1/UPF1/ATAD5/BR

TLE6/HSBP1/MFN2/CLUH/MSTO1/LRPPRC/UXT/ZBED3/DNM1L/BRAT1/MYO19/TRAK2/MUL1/HDAC6/PKD1

EXOSC8/NT5C3B/DIS3/EXOSC5/EXOSC4/EXOSC2/EXOSC6/POLR2G/DIS3L2/CNOT6/EXOSC7/DCPS

ALG6/ALG14/ALG8/ALG1/ALG3/MPDU1/DHDDS/DPAGT1/ALG9/DPM1/RFT1/SRD5A3/ALG2/DOLPP1/ALG12/NUS1/ALG13

OCA2/TYRP1/CDH3/MFSD12/MC1R/PMEL/OPN3/RAB38/DDT/BCL2/AKR7A2/DCT

POLD2/MAD2L2/FAAP20/POLH/USP43/DTL/POLDIP2/PCNA/POLD1/REV3L/PCLAF/SPRTN/USP1/POLI/USP10/RCHY1/PRIMPOL

GATA4/SOX9/HEY1/TWIST1

CDC20/UBE2C/PLK1/CDK2/CDC16/ANAPC11/ANAPC13/ANAPC1

BOP1/RRS1/RPP40/ABT1/UTP23/TSR1/NOL9/NOP14/UTP20/TBL3

FOXF2/ANKRD9/CDC20/RHOBTB3/FBXO17/PRAME/FBXL2/UBE2C/CCNF/HSP90AB1/PBK/DVL1/KBTBD6/NEDD4L/PLK1/RACK1/RNF180/CDK2/NOP53/UB

CDK13/MED25/MED15/MED18/HMGN1/MED8/CTNNB1/SUPT16H/THOC5/MED20/NCBP1/PARP1/MED29/TCERG1/GTF2F1/MED27/BRD4/MED22/MED14/

KIF2C/NCKAP5/DIAPH3/STMN3/TPX2/KIF18B/KIF18A/WDR47/CKAP2/HDAC6/CKAP5/APC2/NCKAP5L/CAMSAP2/CLASP2/KATNB1/CLASP1/MAP6D1/CCS

EXOSC10/DAZAP1/RBM3/PRMT5/WDR77/SNRNP70/PRDX6/SF3B4/CLNS1A/THRAP3/UPF3A/EIF1/CIRBP/TRA2A/UPF1/NCBP1/UPF3B/RBMX/PRPF19/N

VDAC1/MFN2/PINK1/ATP5IF1/MUL1/HDAC6/CDC37

VPS37D/DNM1L/VPS36/CHMP1B/TSG101/CHMP2A/SH3GLB1/CHMP3/VPS37B/EXOC6B/SNF8/VPS37A/CORO1C/STAM/MVB12A/SLC25A46/VPS25/EXOC8/

SHMT1/CTH/SHMT2/SEPHS2/PSPH/PSAT1/DHFR/GGT1/PHGDH/SERINC5/SERINC3/SEPHS1

CENPF/SOX9/TAF3/TAF13/CENPA/CHAF1B/ITGB3BP/DLGAP5/TAF4B/OIP5/CENPI/RAD51/HJURP/CDC45/GMNN/CENPE/SMYD3/CENPV/MED25/SUGT1/

USP13/UBXN2A/STUB1/USP19/USP14/UBQLN1/SGTA/BAG6/BCAP31/RNF185/RNFT1/UBQLN2/CAV1/WFS1/AQP11/TMX1/RNFT2/UBXN1/NFE2L2/SVIP/T

EEF1AKMT3/SMYD2/EHMT2/EEF1AKMT1/METTL21C/SETD4/ETFBKMT/EEF2KMT/METTL21A/SETD6/CSKMT/METTL18/ANTKMT/EHMT1/ATPSCKMT/M

HNRNPA2B1/RBM10/RBM20/C1QBP/SAP18/RNPS1/HNRNPK/SRSF12/RBM42/PTBP1/SRSF9/SRSF10/RBMX/U2AF2/DYRK1A

MYO16/TERT/HORMAD1/CENPF/FAM83D/CDC20/PKMYT1/TRIP13/CCNB1/UBE2C/CDCA8/CDKN2B/CLSPN/NUF2/AURKB/ANLN/MAD2L2/SMARCD3/M

GFPT2/AMDHD2/GNPDA1/UAP1L1/GFPT1/UAP1/GNPNAT1/DPAGT1/SLC35A3/GNPDA2/MGAT1

TRAPPC2L/TRAPPC3/EXOC6B/FAM91A1/TRAPPC5/STARD3NL/NAGLU/TRAPPC6A/EXOC8/TRAPPC1/TRAPPC8/TRAPPC12/EXOC3/TRAPPC13/EXOC2/

MAGEC2/CDC20/UBE2C/MAD2L2/PLK1/PINK1/MAD2L1/FEM1B/ARRDC4/UBE2L3/CDC14B/BMI1/ABL1/FEM1A

ABCD1/IRS2/PPARGC1A/NR4A3/TWIST1/IRS1/MTLN/AKT2

STIL/CENPJ/PLK2/SASS6/BRCA1/PPP1R35/CEP295NL/PLK4/NUP62/C10orf90/CDK5RAP2/CEP76/ALMS1/CEP295/RBM14/TRIM37/KAT2A

DLC1/PTK7/COBL/LHX2/SEMA4C/CECR2/RGMA/TULP3/DVL1/STIL/FZD1/CDK20/FZD3/TMED2/TWIST1/CC2D2A/DEAF1/DVL2/CLUAP1/IFT172/RARG/MT

EXOSC10/SNRPC/CLNS1A/SNRNP200/UPF3A/PRPF40B/EIF1/PRPF40A/UPF1/SRSF12/UPF3B/CACTIN/C9orf78/NCBP2/SRSF1/DCPS/WBP4/SDE2/CWC1

EXO1/WRN/RTEL1/BLM/SLX4/XRCC5/ERCC1/DNA2/SMARCAL1

EXO1/WRN/RTEL1/BLM/SLX4/XRCC5/ERCC1/DNA2/SMARCAL1

EXO1/WRN/RTEL1/BLM/SLX4/XRCC5/ERCC1/DNA2/SMARCAL1

EIF3B/YBX1/EIF3E/RACK1/YBX3/MTOR/EIF3D/EIF3H/YTHDF2/PAIP1/EIF4A1/DENR/EIF3G/EIF2B3/DHX9/EIF3J/DHX29/EIF2B4/EIF2D/EIF2AK4/EIF3I/RWDD

AK5/PGM1/SLC4A4/PKLR/IMPDH2/ENO2/CTPS1/GPI/ENO1/PPARGC1A/NME4/NME1/ATP5MC3/CAD/NDUFS5/HDAC4/UCK2/NDUFAB1/NDUFS6/NDUFA

PLK1/YWHAZ/CDK1/MAPK1/MAP2K2/GOLGA2/STK25

PLK1/YWHAZ/CDK1/MAPK1/MAP2K2/GOLGA2/STK25

AHCYL1/SSU72/ZNF473/POLR2D/LSM10/CPSF3/LSM11/CSTF1/VIRMA/SLBP/NCBP1/APP/CSTF2/GRSF1/RPRD1A/CPSF7/NCBP2/RPRD1B/CPEB1/CPSF

NQO1/FDXR/COQ9/COQ4/COQ8A/AIFM2/UBIAD1/COQ8B/COQ10A/COQ7/COQ5/NDUFA9/COQ3/COQ6/COQ10B/ADCK2

ZMPSTE24/RAD51/GINS1/CDC45/GMNN/FEN1/GINS3/MCM4/TK1/WIZ/WRN/MCM3/PCNA/RTEL1/BLM/CDT1/RTF2/DBF4B/LIG1/UPF1/ATAD5/BRCA2/DB

ALG6/ALG14/ALG8/ALG1/ALG3/MPDU1/DHDDS/DPAGT1/ALG9/DPM1/RFT1/SRD5A3/ALG2/DOLPP1/ALG12/NUS1/ALG13

FZD9/SLC25A4/SLC25A6/SLC25A5/ATP5IF1/ZNF205/MUL1

MYO16/BMP7/CENPF/CDC20/PKMYT1/TRIP13/CCNB1/CDCA8/CDKN2B/CLSPN/NUF2/AURKB/MAD2L2/MAD1L1/BTG3/PLK1/GTSE1/CDK2/NOP53/FZD3/

HDAC4/GNL3/UBA2/SAE1/MUL1/TOLLIP/ARNT/RWDD3/RASD2/PIAS4

SLC1A4/MFSD12/SLC1A1/SLC7A5/SLC3A2

EYA1/HMG20B/P3H3/SOX4/CTH/ATAT1/HDAC4/GNL3/PLOD3/NDUFAB1/PLOD1/HINT1/EEF1AKMT3/SMYD2/CAPN3/TRIM28/GNL3L/EHMT2/UBA2/SENP3/

ACOT7/PANK2/PANK4/PPCS/PANK3/COASY/PPCDC/DCAKD/PANK1

PDK4/SNCA/PDK3

PKMYT1/CCNB1/CCNF/CDKN2B/PSRC1/HSP90AB1/PLK1/CDKN1C/CDC6/CDC25C/TFAP4/PSMD10/CCNB2/AKT1/CDK5R1/LATS2/CCNE1/CCNG1/CCN

DLC1/PTK7/COBL/LHX2/SEMA4C/CECR2/RGMA/TULP3/DVL1/STIL/FZD1/CDK20/FZD3/TMED2/TWIST1/CC2D2A/DEAF1/DVL2/CLUAP1/IFT172/RARG/MT

PDK4/SNCA/ACSS2/PPT2/PDK3/GCDH/ACSS1/PDHA1/HACD1/DLAT/ACLY/ACOT7/PANK2/PANK4/ELOVL1/PPCS/SLC25A1/MMUT/PANK3/ACACB/DLD/PD

PDK4/SNCA/ACSS2/PPT2/PDK3/GCDH/ACSS1/PDHA1/HACD1/DLAT/ACLY/ACOT7/PANK2/PANK4/ELOVL1/PPCS/SLC25A1/MMUT/PANK3/ACACB/DLD/PD

PDK4/SNCA/ACSS2/PPT2/PDK3/GCDH/ACSS1/PDHA1/HACD1/DLAT/ACLY/ACOT7/PANK2/PANK4/ELOVL1/PPCS/SLC25A1/MMUT/PANK3/ACACB/DLD/PD

GMPR/CTPS1/SHMT1/TTR/CAD/TYMS/GART/PAICS/DHODH/MTOR/CTPS2/PPAT/MAPK1/CMPK1/RRM1

PEX10/STUB1/LONP1/HDAC6/RNF126/CLPP/ZYG11B/LONP2/KLHL15/YME1L1/UBE2W/FBXL17/CUL3

TUB/CLUAP1/IFT22/IFT172/DYNC2LI1/DYNC2H1/CEP131/WDR35/IFT140/DYNLL1/WDR19/IFT81/WDPCP/IFT88/FUZ/LCA5L/TTC21B/SSX2IP/TRAF3IP1/PC

PYGO1/RCC1/NUP93/PLK1/DYRK3/LMNB2/ZMPSTE24/EMG1/BEND3/PRKCA/SERBP1/AHCTF1/EMD/SUN1/NDC1/UBXN2A/REEP4/MFSD14A/UBE2I/USP

CTPS1/NME4/NME1/UCK2/CTPS2/NME2/NME6/UCKL1/UCK1

FZD9/SLC25A4/SLC25A6/STPG1/SLC25A5/ATP5IF1/ZNF205/THEM4/TP53/MUL1/RTL10/BNIP3L/VDAC2/RHOT2/NAIF1/RHOT1

MAD2L2/PLK1/AUNIP/PRKDC/PSMD14/SETMAR/POLB/CYREN/NHEJ1/DCLRE1B/DEK/MRE11/RHNO1/KMT5C/RIF1/XRCC5/USP51/ERCC1/TFIP11/XRCC

RANBP17/NUP93/YBX1/TOMM20/NSUN2/EIF4A3/XPO5/HNRNPA2B1/LRPPRC/PEG10/GLE1/DDX25/AHCTF1/TNKS/KHSRP/NDC1/SEC13/NXT1/NXF5/DH

RANBP17/NUP93/YBX1/TOMM20/NSUN2/EIF4A3/XPO5/HNRNPA2B1/LRPPRC/PEG10/GLE1/DDX25/AHCTF1/TNKS/KHSRP/NDC1/SEC13/NXT1/NXF5/DH

FOXF2/ANKRD9/CDC20/RHOBTB3/FBXO17/PRAME/FBXL2/UBE2C/CCNF/HSP90AB1/PBK/DVL1/KBTBD6/NEDD4L/PLK1/DAB2IP/RACK1/UBXN8/RNF180/

VPS37D/CHMP1B/TSG101/CHMP2A/CHMP3/VPS37B/VPS37A/MVB12A/VPS28/CHMP6/CHMP1A/VPS37C/CHMP4B/PDCD6IP/VPS4A/CHMP2B/CHMP7/CH

AK5/PGM1/SLC4A4/PKLR/IMPDH2/ENO2/CTPS1/GPI/ENO1/PPARGC1A/NME4/NME1/ATP5MC3/CAD/NDUFS5/HDAC4/UCK2/TYMS/NDUFAB1/NDUFS6/N

PYCR1/SHMT1/CTH/MTRR/SHMT2/PLOD3/SEPHS2/ADI1/PSPH/AASDHPPT/PLOD2/ALDH18A1/PSAT1/GOT2/DHFR/MRI1/GOT1/GGT1/GLS/PYCR2/MTR/

PYCR1/SHMT1/CTH/MTRR/SHMT2/PLOD3/SEPHS2/ADI1/PSPH/AASDHPPT/PLOD2/ALDH18A1/PSAT1/GOT2/DHFR/MRI1/GOT1/GGT1/GLS/PYCR2/MTR/

RPP25/POP1/RPP40/ABT1/RPP21/HSD17B10/POP7/NOP14/UTP20/TBL3/POP5/POP4/TRMT10C/THG1L

RPP25/POP1/RPP40/ABT1/RPP21/HSD17B10/POP7/NOP14/UTP20/TBL3/POP5/POP4/TRMT10C/THG1L

MYC/SMARCD3/GRWD1/SMARCA4/SET/SMARCC1/HMGA1/SMARCD2/SUPT16H/SMARCB1/SSRP1/ARID2/SMARCE1/SMARCD1

CTPS1/NME4/NME1/CAD/UCK2/CTPS2/NME2/NME6/UCKL1/UCK1

TERT/NEK2/PINX1/HSP90AB1/AURKB/CCT3/CCT6A/EXOSC10/DKC1/NHP2/RAD50/TNKS/CCT8/PIF1/CCT4/GNL3L/TP53/CCT2/SMG5/MAPK1/TCP1/RFC1

NDUFAB1/HSPA9/NUBP2/BOLA3/LYRM4/CIAPIN1/ABCB7/BOLA1/ISCA1/CIAO1/CIAO3/NFS1/CIAO2A/FDX2/NUBPL/FXN

NDUFAB1/HSPA9/NUBP2/BOLA3/LYRM4/CIAPIN1/ABCB7/BOLA1/ISCA1/CIAO1/CIAO3/NFS1/CIAO2A/FDX2/NUBPL/FXN

WIPI1/TMED2/SEC16B/TRAPPC2L/TRAPPC3/TMED9/PEF1/PDCD6/TRAPPC5/TRAPPC6A/TRAPPC1/TRAPPC8/TRAPPC12/KLHL12/ARFGAP3/AP1AR/CS

PKMYT1/CCNB1/CCNF/CDKN2B/PSRC1/PLK1/CDKN1C/CDC6/CDC25C/TFAP4/PSMD10/CCNB2/AKT1/CDK5R1/LATS2/CCNE1/CCNG1/CCNO/CDC37/IP

AHNAK2/RBPMS2/MBNL2/RBM11/EXOSC10/SUPT3H/DAZAP1/HNRNPA2B1/USP22/PQBP1/ZNF326/RBM3/AHNAK/PRMT5/WDR77/ESRP1/FASTK/RBM10

VPS37D/CHMP1B/TSG101/CHMP2A/CHMP3/VPS37B/VPS37A/MVB12A/VPS28/CHMP6/CHMP1A/VPS37C/CHMP4B/PDCD6IP/NEDD4/VPS4A/CHMP2B/CH

FZD9/SLC25A4/SLC25A6/BCL2L2/SLC25A5

NEIL2/NTHL1/LIG3/UNG/FEN1/POLB/APEX1/POLE/WRN/PCNA/RPA3/POLD1/PRMT6/SMUG1/RPA1/RPA2/LIG1/APEX2/PARP1/NEIL3/DNA2/XRCC1/FAM16

RANBP17/NUP93/YBX1/TOMM20/NSUN2/EIF4A3/XPO5/HNRNPA2B1/LRPPRC/PEG10/GLE1/DDX25/AHCTF1/TNKS/KHSRP/NDC1/SEC13/NXT1/NXF5/DH

ABCD1/SLC25A23/SNCA/NOP53/SLC25A33/SHMT2/COX7A1/PINK1/AK4/DNAJC15/ARL2/UQCC2

TERT/FAM83D/CDC20/CCNB1/UBE2C/SMARCD3/STIL/DLGAP5/DYRK3/CDC6/CDC25C/CDCA5/ESPL1/CDC16/AKT1/CENPJ/RCC2/CDK4/DDRGK1/CCN

MYO16/TERT/BMP7/CCDC8/CENPF/CDC20/NEK2/PKMYT1/EDN3/TRIP13/CCNB1/UBE2C/CDCA8/RCC1/CDKN2B/CLSPN/LRP5/NUF2/AURKB/ANLN/NU

CDH3/MYO5A/MLPH/RAB17/GPR143/RAB27A/DCTN2/MKKS/ARL6/BLOC1S6/DCTN1/BLOC1S3/MREG/BBS7/BLOC1S5/RAB11A/RAB11B

CDH3/MYO5A/MLPH/RAB17/GPR143/RAB27A/DCTN2/MKKS/ARL6/BLOC1S6/DCTN1/BLOC1S3/MREG/BBS7/BLOC1S5/RAB11A/RAB11B

CCNB1/SMARCD3/DYRK3/CDC25C/RCC2/CDK4/DTL/CDC25B/CDK1/RAD51C/RRM1/DBF4B/CDC25A/APP/ATAD5/BRD4

EXOSC10/DKC1/INTS11/SNAPC2/NHP2/EXOSC8/INTS6/EXOSC5/INTS3/EXOSC4/INTS13/EXOSC2/EXOSC6/INTS14/RPAP2/MTREX/SNAPC1/INTS1/TOE1/

NEK2/AURKB/NCAPG/CCT3/CDK2/GNL3/BUB1/CCT6A/RUVBL2/DKC1/SETDB2/HNRNPA2B1/RAD50/TNKS/NABP2/CCT8/TRIM28/CCT4/FEN1/CCT2/MA

ABCD1/SLC25A23/SNCA/SLC25A33/SHMT2/COX7A1/PINK1/AK4/DNAJC15/UQCC2/COX7A2L/NUPR1/PPIF/COX7A2

INTS11/EXOSC8/INTS6/EXOSC5/INTS3/EXOSC4/INTS13/EXOSC2/EXOSC6/INTS14/INTS1/TOE1/INTS7/EXOSC7/USB1/TUT1/INTS9/EXOSC3/EXOSC9

TUB/TULP3/CROCC/CDK20/NPHP4/TCTN2/CC2D2A/ROPN1B/TCTN1/RAB11FIP3/GAS8/DYNC2H1/GGA1/WDR35/BBS9/IFT140/WDR19/CCDC88A/FSIP2/

BMP7/DLC1/PTK7/COBL/LHX2/SEMA4C/CECR2/RGMA/TULP3/DVL1/STIL/FZD1/CDK20/FZD3/TMED2/TWIST1/CC2D2A/DEAF1/DVL2/CLUAP1/IFT172/RA

TBC1D7/NPRL3/SEC13/ITFG2/YWHAG/YWHAZ/RNF152/CASTOR2/SEH1L/PRKACB/SESN2/TSC2/AKT1S1/MIOS/UBE3A/RNF167/UBE2W/PRKACA/DEPT

IQGAP3/CEP55/KIF20A/CNTROB/ARF1/CHMP1B/CHMP2A/CHMP3/IQGAP1/CHMP6/CHMP1A/NUP62/RHOA/KIF20B/CHMP4B/PDCD6IP/KLHDC8B/VPS4A

CDCA8/AURKB/PLK1/PRC1/AURKA/PDXP/CDC6/KIF14/KIF23/CIT/BIRC5/AHCTF1/KLHL21/RAB11FIP3/CENPV/MYO19/PKP4/KIF20A/ECT2/CDC25B/ZFYVE

BMP7/GPR161/DLC1/GBX2/PTK7/COBL/LHX2/SEMA4C/EPHA2/CECR2/RGMA/TULP3/DVL1/STIL/FZD1/CDK20/FZD3/SEMA3C/ZNF358/TMED2/PLOD3/TW

FOXF2/NQO1/RNF144A/PRAME/UBE2C/KBTBD6/NEDD4L/UBE2T/RUSC1/RNF180/BCL2/BFAR/PPIA/NHLRC3/THOP1/CDC16/RNF14/TTC3/TNKS/RNF6/U

ARMC3/HSPB8/WIPI1/SLC25A4/DIAPH3/GPSM1/VDAC1/VPS37D/EFNB1/UBQLN4/MFN2/MTOR/AP4M1/TSPO/PINK1/CTTN/AKT1/CDK5R1/GNAI3/MAP1LC

TLE6/CDH3/KIF25/INSC/STXBP1/GAB2/KIT/KIF2C/CENPF/FAM83D/GPSM2/NEK2/WIPI1/SPIRE2/SEPTIN5/CCNB1/CDCA8/PINX1/PSRC1/NUF2/GPSM1/AP

ROBO2/SNAI2/MSX2/BMP7/TBX3/ADAMTS5/SOX9/HEYL/HEY1/TWIST1

GPSM2/WIPI1/MNS1/CROCC/STIL/FSCN1/SPAG5/UBAP2L/CENPJ/SNX7/PIP4K2B/ULK1/ATMIN/PIP4K2C/ARHGAP35/SNX30/DYNC1H1/SH3GLB1/SASS6/C

CA14/CA5B/SHMT1/AHCYL1/TYMS/SHMT2/AHCY/CA4/MTHFD1L/MAT2A

TDRD12/TERT/TDRD9/WTIP/GPAT2/ZMPSTE24/PLD6/XPO5/TDRD7/HNRNPA2B1/LIMD1/EIF6/ZC3H7A/TARBP2/SND1/PRKRA/ELOC/SMAD2/TP53/DHX9

TONSL/RAD51/CENPS/RAD50/SETMAR/EME1/CARM1/CENPX/RBBP8/NUCKS1/POLE/WRN/MCM8/PCNA/RTEL1/BLM/MRE11/TIMELESS/POLD1/ZNF365

DIAPH3/TPX2/SNCA/MAP2/WDR47/CKAP2/ARHGEF7/HDAC6/EML2/TBCD/APC2/CAMSAP2/CLASP2/KATNB1/CLASP1/MAP6D1/STMN1/MAPRE1

SH3BP4/ENDOG/TBC1D7/NPRL3/SEC13/ITFG2/YWHAG/YWHAZ/EPM2A/RNF152/CASTOR2/SEH1L/MINAR1/PDCD6/PRKACB/SESN2/TSC2/AKT1S1/MIOS

CDK13/MED25/MED15/MED18/MED8/CTNNB1/MED20/NCBP1/MED29/TCERG1/GTF2F1/MED27/BRD4/MED22/MED14/NCBP2/MED4/LEO1/MED21/MED1

RYR2/PPARGC1A/POSTN/SLC7A5/ADSL/CAPN3/PERM1

FZD9/SLC25A4/SLC25A6/BCL2/STPG1/BCL2L2/SLC25A5/ATP5IF1/ZNF205/THEM4/TP53/MUL1/ALKBH7/RTL10/BNIP3L/VDAC2/RHOT2/NAIF1/RHOT1/PPIF

AK5/PGM1/SLC4A4/PKLR/IMPDH2/ENO2/GPI/ENO1/PPARGC1A/NME4/NME1/ATP5MC3/NDUFS5/HDAC4/NDUFAB1/NDUFS6/NDUFA13/NDUFA5/MTOR/

TRIP13/RAD54L/HSF2BP/CCNB1IP1/HFM1/MEIOB/PSMC3IP/RAD51/TOP2A/MSH5/CENPS/RAD50/RAD54B/EME1/BRIP1/CENPX/FANCD2/RBBP8/NUCKS

MYC/TAF3/TAF13/CTNNBIP1/TAF4B/CDK4/MED25/TAF12/TAF4/TP53/ZNHIT1/TAF11/FOXO1/MED15/MED18/MED8/MED20/TAF6/MDC1/ERCC1/GTF2A2/

AK5/CTPS1/SHMT1/NEIL2/NME4/NME1/CAD/NTHL1/UCK2/NT5M/TYMS/DCTPP1/DHODH/CTPS2/UNG/NME2/DTYMK/CMPK1/SLC4A7/NME6/UCKL1/SM

PIGK/DPM3/SLC30A5/PIGF/PIGP/PIGZ/PIGW/PIGX/PGAP1/DPM2/PIGQ/PIGV/GPAA1/PIGU/PIGA/DPM1/PIGO/PIGB/PIGH/MPPE1/PIGS

TDRD12/HORMAD1/MSX2/TDRD9/CDC20/NEK2/PKMYT1/SPIRE2/TRIP13/PDE3A/NUF2/ASPM/MNS1/RSPH1/PLK1/CKS2/RAD54L/CDK2/NPM2/NSUN2/E

TUB/KIF17/MAP6/KIF4A/TUBA1B/AP3M2/TMEM201/FNBP1L/NEFH/HSBP1/CDR2L/KIF3A/TUBB/MYO5A/DYNC1I1/MYO10/F8A1/MAP2/LRPPRC/TUBA1C/C

VPS37D/TBC1D20/CHMP1B/TSG101/CHMP2A/CHMP3/VPS37B/RAB1B/VPS37A/MVB12A/VPS28/PC/CHMP6/CHMP1A/VPS37C/SPCS1/CHMP4B/PDCD6IP/

AK5/PGM1/SLC4A4/PKLR/ENO2/GPI/ENO1/PPARGC1A/ATP5MC3/NDUFS5/HDAC4/NDUFAB1/NDUFS6/NDUFA13/NDUFA5/MTOR/NDUFB9/SLC25A13/N

AP3M2/TMED2/SEC16B/TRAPPC2L/TRAPPC3/AP3D1/TMED9/PEF1/PDCD6/TRAPPC5/TRAPPC6A/TRAPPC1/TRAPPC8/TRAPPC12/KLHL12/ARFGAP3/TR

WIPI1/EFNB1/MTOR/PINK1/EHMT2/SNX7/TBC1D14/PIP4K2B/ULK1/SEC22B/PIP4K2C/SNX30/SH3GLB1/IFT88/RAB1B/NUPR1/ATG2A/PIP4K2A/WDR45/RN

NEK2/MYC/PINX1/AURKB/CCT3/GNL3/CCT6A/RUVBL2/EXOSC10/DKC1/HNRNPA2B1/RAD50/TNKS/NABP2/CCT8/PIF1/CCT4/GNL3L/TP53/CCT2/SMG5/T

TDRD12/HORMAD1/MSX2/TDRD9/CDC20/PKMYT1/SPIRE2/TRIP13/PDE3A/NUF2/ASPM/PLK1/CKS2/RAD54L/NPM2/NSUN2/HSF2BP/CCNB1IP1/AURKA/

MYO16/HORMAD1/KIF25/BMP7/CENPF/CDC20/HPGD/NEK2/PKMYT1/TRIP13/CCNB1/CCNF/CDCA8/CDKN2B/CLSPN/NUF2/AURKB/MAD2L2/MAD1L1/B

AHCYL1/RBPMS2/RBM11/EXOSC10/SRPK3/DAZAP1/HNRNPA2B1/RBM3/PRMT5/WDR77/RBM10/RBM20/SNRNP70/C1QBP/PRDX6/SAP18/RNPS1/DHX9/

ZDHHC11B/MTCL1/HSPD1/AKAP6/CHCHD4/PEX6/SLC1A1/CHP1/VPS37D/ZDHHC9/TOMM20/SORT1/ICMT/TOMM40/TIMM13/MIPEP/MFN2/NDUFA13/H

IMPDH2/SHMT1/CAD/UCK2/TYMS/GART/PAICS/ADSL/DHODH/ATIC/PFAS/PPAT/TK1/CMPK1/UCKL1/UCK1/DUT/DGUOK/PRPS1/APRT/PRPS2/AMPD2

EYA1/MMP16/TFAP2A/BMP7/SATB2/HOXB9/HOXB8/HOXB6/TULP3/IRX5/HOXD3/HOXB7/EIF4A3/COL11A1/TWIST1/HOXC9/COL2A1/HOXB3/HOXB2/SLC39

FZD9/SLC25A4/SLC25A6/STPG1/SLC25A5/ATP5IF1/ZNF205/TP53/MUL1/RTL10/BNIP3L/VDAC2/RHOT2/RHOT1/PPIF/IER3/BCL2L11/CHCHD10/SLC35F6

HSPB8/TMTC4/HSPD1/CTH/DAB2IP/RACK1/TMED2/BFAR/TBL2/TMEM33/NCK2/VAPB/DDRGK1/STUB1/HDAC6/RHBDD2/RNF126/SERP2/PACRG/UFD1/

BMP7/DLC1/PTK7/COBL/LHX2/SEMA4C/CECR2/RGMA/TULP3/DVL1/STIL/FZD1/CDK20/FZD3/TMED2/TWIST1/CC2D2A/DEAF1/DVL2/CLUAP1/TCTN1/IFT

INTS11/NELFE/CDK13/MED25/INTS6/INTS3/NELFCD/INTS13/MED15/MED18/WDR43/MED8/CTNNB1/MED20/INTS14/AXIN1/NCBP1/INTS1/PARP1/MED29

AK5/PGM1/SLC4A4/PKLR/IMPDH2/ENO2/GPI/ENO1/PPARGC1A/NME4/NME1/ATP5MC3/NDUFS5/HDAC4/NDUFAB1/NDUFS6/NDUFA13/NDUFA5/NUDT1

NR2E1/ID4/ACAN/ASPM/VEGFA/SMARCD3/FZD3/WDR62/PAX6/DCT/HAPLN1

OCA2/TYRP1/MAOB/CHRNB2/CDH3/MFSD12/MC1R/PMEL/SLCO1C1/GPR37/OPN3/SLC1A1/SNCA/FAH/RNF180/RAB38/DDT/BCL2/COMT/HAND2/DCT

TUB/KIF17/KIF4A/AP3M2/TMEM201/NEFH/HSBP1/CDR2L/KIF3A/DYNC1I1/MAP2/LRPPRC/CLUAP1/IFT22/UXT/SUN1/IFT172/KIF1B/DYNC2LI1/DST/TRAK2

EYA1/SNAI2/KCNK2/MAPK8IP2/BMP7/ABCD1/GSTP1/FOXM1/SPIRE2/MYC/SEMA4C/SPRED1/SOX4/RGMA/ENO1/PBK/MAD2L2/CHCHD2/SMARCD3/MO

HORMAD1/FZD9/BMP7/CENPF/DLC1/CDC20/SLC25A4/TRIP13/SPTBN2/CCNB1/CCNF/TMOD1/CDCA8/PINX1/SLC25A6/DIAPH3/NUF2/AURKB/TPX2/MA

TERT/INSC/STXBP1/TBX3/PINX1/HSPD1/HSP90AB1/ATP1B2/SOX4/PEX6/DVL1/PPARGC1A/CCT3/NEDD4L/PYCARD/CHP1/SNCA/PLK1/GLMP/OTUD3/CR

FZD9/SLC25A4/SLC25A6/STPG1/SLC25A5/ATP5IF1/ZNF205/MUL1/RTL10/BNIP3L/VDAC2/RHOT2/RHOT1/PPIF/IER3/BCL2L11/CHCHD10/SLC35F6

MNS1/CROCC/LPAR1/TBC1D7/AKT1/CENPJ/ODF2/RAB11FIP3/DYNC2LI1/CNTROB/ATMIN/ARHGAP35/IFT140/MARK4/DYNLL1/CCDC88A/WDPCP/IFT8

MAD2L2/MORF4L2/UBQLN4/FANCB/RUVBL2/RAD51/SENP3/EP400/FIGNL1/MEAF6/MRGBP/SPIDR/RTEL1/TIMELESS/VPS72/ZNF365/ABL1/RIF1/USP51/F

HSPB8/TMTC4/HSPD1/HSP90AB1/KBTBD6/CTH/DAB2IP/RACK1/TMED2/MFN2/BFAR/TBL2/HSPA6/TMEM33/HSPB2/NCK2/HSPE1/UBE2J2/VAPB/DDRGK

SNAI2/SOX9/DAB2IP/CAD/KNSTRN/PTPN12/BAIAP2/AKT1/FOXC1/PDE8A/ZPR1/PTPN11/DUSP3

TERT/MSX2/FAM83D/GPSM2/CDC20/EDN3/CCNB1/UBE2C/CDCA8/LRP5/AURKB/NUSAP1/NCAPG/SMARCD3/MAD1L1/STIL/NPM2/DLGAP5/AURKA/BUB

IRX1/BMP7/SOX8/DLC1/PTK7/COBL/LHX2/SOX9/SEMA4C/CECR2/RGMA/TULP3/DVL1/IRX3/STIL/FZD1/CDK20/FZD3/TMED2/TWIST1/CC2D2A/DEAF1/DV

CDK2/ENDOG/ZMPSTE24/WRNIP1/LIG3/DSCC1/CIZ1/GMNN/MCM4/RFC3/WIZ/RFC2/RFC4/MCM3/PCNA/BLM/TIMELESS/CDT1/MCM7/CHTF18/SSBP1/

HORMAD1/TDRD9/CDC20/PKMYT1/TRIP13/PLK1/CKS2/RAD54L/HSF2BP/CCNB1IP1/AURKA/HFM1/MEIOB/NDC80/PSMC3IP/CDC25C/ESPL1/RAD51/CC

MYC/NAT14/TAF3/TAF13/NOC2L/CTNNBIP1/PPARGC1A/MITF/TAF4B/TWIST1/GTF2F2/GTF3C5/ELOA/CDK4/MED25/TAF12/POLR3H/ELOC/TAF4/MAZ/SM

PSRC1/DIAPH3/STMN3/TPX2/SNCA/MAP2/WDR47/CKAP2/SKA3/CDK5R1/ARL2/RAC1/ARHGEF7/SKA2/PRUNE1/HDAC6/CKAP5/AKAP9/EML2/TBCD/APC

AK5/PDK4/ADCY2/IMPDH2/CTPS1/SHMT1/ENO1/PPARGC1A/NME4/NME1/SNCA/ATP5MC3/CAD/NDUFS5/UCK2/TYMS/ACSS2/NDUFAB1/GART/PAICS/A

EDNRB/HSPD1/CHCHD4/PEX6/TOMM20/TOMM40/AP4M1/PAM16/DNAJC15/GRPEL1/TIMM50/PEX10/TOMM22/ROMO1/RTN2/TIMM17B/TIMM21/AIFM1/S

GPSM2/WIPI1/SPTBN2/MNS1/LRFN4/CROCC/STIL/PLK1/EFNB1/LPAR1/FSCN1/TBC1D7/MTOR/SPAG5/KIF9/UBAP2L/PINK1/PTPRS/AKT1/CENPJ/ODF2/P

AK5/PDK4/ADCY2/IMPDH2/CTPS1/SHMT1/ENO1/PPARGC1A/NME4/NME1/SNCA/ATP5MC3/CAD/NDUFS5/UCK2/TYMS/ACSS2/NDUFAB1/GART/PAICS/A

NR2E1/CNTN4/BMP7/ZNF536/SOX8/ID4/RAP1GAP/SOX9/HEY1/ASPM/IRX3/DLX1/PAX6/SPAG9/NKX2-2/MIB1/MEIS1

TDRD9/CBX3/MYC/CDK2/LMNB2/BEND3/SETDB2/RLIM/DNMT3A/SAMD1/CENPV/TRIM28/CDYL/PCGF3/LRIF1/TET1/SCMH1/HAT1/MBD3/SUV39H1/LMN

TDRD12/HORMAD1/MSX2/TDRD9/CDC20/SPIRE2/TRIP13/PDE3A/NUF2/ASPM/PLK1/CKS2/RAD54L/NPM2/HSF2BP/CCNB1IP1/AURKA/HFM1/MEIOB/ND

RANBP17/CHCHD4/CHP1/AHCYL1/PTPN14/XPO5/HSPA9/CCHCR1/EMD/YWHAE/XPO4/NXT1/PKD1/IFI27/PTPN11/DNAJC27/CSE1L/CDK5/STRADB/DUS

NR2E1/TERT/MSX2/TBX3/FAM83D/GPSM2/CDC20/EDN3/CCNB1/UBE2C/CDCA8/PSRC1/LRP5/AURKB/NUSAP1/NCAPG/SMARCD3/MAD1L1/STIL/EIF4E

MAGEC2/FOXF2/CDC20/HMG20B/UBE2C/PINX1/HSP90AB1/COPS7A/SOX4/MAD2L2/CHP1/PLK1/BIRC7/RNF180/CRTAP/NOP53/PTTG1IP/HDAC4/COPS8/

TDRD9/CBX3/MYC/LHX2/TFAP2C/CDK2/HDAC4/LMNB2/BEND3/SETDB2/RLIM/DNMT3A/SAMD1/CENPV/TRIM28/CDYL/PCGF3/HDAC6/LRIF1/TET1/SCM

ST13/CCT3/CRTAP/DNAJB6/CCT6A/HSPA9/DFFA/HSPA6/TRAP1/HSPE1/CCT8/PEX19/PFDN6/CCT4/CCT2/HSPA8/PFDN2/P3H1/PFDN5/PDCD5/TCP1/PF

SNAI2/MYC/SPRED1/SOX4/AURKB/MIF/NOP53/PTTG1IP/AURKA/DYRK3/TWIST1/ZMPSTE24/BCL2/BOP1/MTOR/PSMD10/RRS1/PAK1IP1/AKT1/SMYD2/P

EEF2/WTIP/BZW2/KRT17/SHMT1/SOX4/EIF3B/EIF4EBP1/YBX1/EIF3E/RACK1/RGS2/PUS7/EPHA4/EIF4A3/TMED2/TYMS/SHMT2/FASTKD3/PRKDC/YBX3/M

VPS37D/LAPTM4B/PHETA2/ATP6V0B/ARFGEF2/VPS36/STX6/RAB27A/CLCN3/DNAJC13/CHMP1B/TSG101/CHMP2A/CHMP3/VPS37B/TMEM9/ATP6V0A1/P

TDRD12/TERT/TDRD9/GPAT2/ZMPSTE24/PLD6/TDRD7/HNRNPA2B1/ZC3H7A/TARBP2/PRKRA/SMAD2/TP53/DROSHA/DGCR8/TDRD6/AGO1/PUM1/TR

DYRK3/TBC1D7/LAMTOR5/AKT1/GPR137B/NPRL3/SEC13/ITFG2/GPR155/YWHAG/YWHAZ/LAMTOR2/RRAGD/LAMTOR4/RNF152/CASTOR2/SEH1L/DGK

NDN/TPPP/SPRED1/HDAC4/CPT1C/PPT2/LYPLAL1/HDAC6/NEK3/SIRT2/WDR5/FNTA/DR1/CCAR2/SIRT5/NOTUM/BRMS1/HDAC3/FLNA/ABHD13/SPRE

RANBP17/ABCD1/SLC25A4/SLC25A6/NUP93/SLC25A23/YBX1/TOMM20/NSUN2/EIF4A3/SLC25A33/SLC35B2/XPO5/SLC35B4/HNRNPA2B1/SLC25A24/LRPP

MYC/TAF3/TAF13/CTNNBIP1/MITF/TAF4B/TWIST1/CDK4/MED25/TAF12/TAF4/TP53/ZNHIT1/TAF11/FOXO1/MED15/MED18/MED8/MED20/TAF6/MDC1/ER

TAF3/TAF13/MITF/TAF4B/TWIST1/MED25/TAF12/TAF4/TP53/ZNHIT1/TAF11/MED15/MED18/MED8/MED20/TAF6/MDC1/ERCC1/GTF2A2/MED29/MED27/M

HSPB8/TMTC4/HSPD1/HSP90AB1/KBTBD6/CTH/DAB2IP/RACK1/TMED2/MFN2/BFAR/TBL2/HSPA6/TMEM33/HSPB2/NCK2/HSPE1/UBE2J2/VAPB/DDRGK

HORMAD1/TDRD9/TRIP13/PLK1/CKS2/RAD54L/HSF2BP/CCNB1IP1/AURKA/HFM1/MEIOB/NDC80/PSMC3IP/ESPL1/RAD51/CCNB2/TOP2A/MSH5/CENPS/

RBPMS2/RBM11/EXOSC10/DAZAP1/HNRNPA2B1/RBM3/PRMT5/WDR77/RBM10/RBM20/SNRNP70/C1QBP/PRDX6/SAP18/RNPS1/SF3B4/CLNS1A/HNRN

AK5/PDK4/ADCY2/IMPDH2/ENO1/PPARGC1A/NME4/NME1/SNCA/ATP5MC3/NDUFS5/ACSS2/NDUFAB1/GART/PAICS/ADSL/NDUFS6/NDUFA13/NDUFA

AP3M2/TMEM201/NEFH/HSBP1/CDR2L/KIF3A/DYNC1I1/MAP2/LRPPRC/UXT/SUN1/TRAK2/AP3D1/HDAC6/NDEL1/KXD1/COPG2/RHOT2/COPG1/MECP2/

CDH3/STXBP1/WIPI1/SEPTIN5/AP3M2/CADPS/FNBP1L/SNCA/CDR2L/RAB3A/TMED2/KIF3A/MYO5A/DYNC1I1/F8A1/PINK1/MAP2/CADPS2/SEC16B/MLPH/

NEK2/PINX1/AURKB/CCT3/CCT6A/EXOSC10/DKC1/HNRNPA2B1/TNKS/CCT8/PIF1/CCT4/GNL3L/TP53/CCT2/SMG5/MAPK1/TCP1/RTEL1/SLX4/CTNNB1/P

KIF2C/NCKAP5/MICAL2/SPTBN2/MYC/TMOD1/DIAPH3/SEMA5A/STMN3/TPX2/SMARCD3/UBQLN4/SCIN/DSTN/PDXP/DYRK3/ZMPSTE24/KIF18B/EPS8/KI

KIF2C/NCKAP5/MICAL2/SPTBN2/TMOD1/DIAPH3/SEMA5A/STMN3/TPX2/SCIN/DSTN/PDXP/KIF18B/EPS8/KIF18A/WDR47/CKAP2/CAPG/DNAJC6/TMOD2/

HSPB8/TMTC4/HSPD1/CTH/DAB2IP/RACK1/TMED2/BFAR/TBL2/TMEM33/NCK2/VAPB/DDRGK1/STUB1/RHBDD2/SERP2/PACRG/MBTPS1/RPAP2/DNAJ

MAGEC2/FOXF2/CDC20/HMG20B/UBE2C/PINX1/HSP90AB1/COPS7A/SOX4/MAD2L2/CHP1/PLK1/BIRC7/RNF180/NOP53/PTTG1IP/HDAC4/COPS8/GNL3/S

EEF2/WTIP/BZW2/GCNT2/KRT17/SHMT1/SOX4/EIF3B/TBRG4/EIF4EBP1/YBX1/EIF3E/RACK1/RGS2/PUS7/NSUN2/EPHA4/EIF4A3/TMED2/TYMS/SHMT2/Z

OCA2/TPCN2/ATP10B/WIPI1/HSD17B1/EFNB1/LAPTM4B/CLN6/MFN2/HPS4/MTOR/AP4M1/PINK1/MAP1LC3B/ATG4B/ATP6V0B/ATG9A/UBXN2A/EHMT2/S

AP3M2/FNBP1L/GOLPH3L/TMED2/SEC16B/CIDEB/TRAPPC2L/SEC31A/VAPB/SCAP/SEC13/TBC1D20/TRAPPC3/AP3D1/TMED9/PEF1/PDCD6/VAPA/TRAP

MMP8/SNAI2/STON1/KIF2C/WIPI1/NCKAP5/SLC25A4/MICAL2/ADAMTS5/SPTBN2/MYC/TMOD1/DIAPH3/DKK1/SEMA5A/DNASE2B/STMN3/CECR2/MMP1

TAF3/TAF13/TAF4B/MED25/TAF12/TAF4/TP53/ZNHIT1/TAF11/MED15/MED18/MED8/MED20/TAF6/MDC1/ERCC1/GTF2A2/MED29/MED27/MED22/MED14/

GNAI1/GPSM2/TPPP/MAP6/CCNF/PSRC1/DIAPH3/STMN3/TPX2/NEFH/ATAT1/STIL/SNCA/PLK1/WDR62/AURKA/SPAG5/KIF18A/KIF9/MAP2/WDR47/CKAP

DENND2A/WIPF3/RHOBTB3/CLTCL1/SNX8/RAB6B/SPAG9/VAMP3/PHETA2/YKT6/RBSN/UBE2O/TBC1D14/SNX12/TMEM87A/SLC66A2/STX6/STX10/GGA1/

BZW2/EIF3B/EIF4EBP1/EIF3E/DHX33/TMED2/MTOR/EIF3D/EIF3H/YTHDF2/PAIP1/EIF4A1/EIF6/DDX1/NCK2/DENR/EIF3G/MTFMT/C8orf88/EIF2B3/EIF4G3/

PYCR1/SHMT1/CTH/CAD/MTRR/SHMT2/PLOD3/SEPHS2/ADI1/PSPH/AASDHPPT/ILVBL/PLOD2/ALDH18A1/PSAT1/GOT2/DHFR/MRI1/GOT1/GGT1/GLS/P

SNAI2/ZMPSTE24/RAD51/CLOCK/HUS1/TP53/ECT2/ELK1/NET1/TSPYL5/NUCKS1/WRN/FIGNL1/SPIDR/HRAS/BLM/GTF2H5/GADD45A/RHNO1/BRCA1/C

WIPI1/EFNB1/MFN2/MTOR/AP4M1/PINK1/MAP1LC3B/ATG4B/ATG9A/UBXN2A/EHMT2/SNX7/TBC1D14/STBD1/PIP4K2B/ULK2/ULK1/SEC22B/PIP4K2C/ATG

AK5/PDK4/ADCY2/IMPDH2/SHMT1/ENO1/PPARGC1A/NME4/NME1/SNCA/ATP5MC3/NDUFS5/ACSS2/NDUFAB1/GART/PAICS/ADSL/NDUFS6/NDUFA13/

TERT/RYR2/RANBP17/CACNG4/HSPD1/CHCHD4/PEX6/NUP93/IPO4/CCT3/CHP1/VPS37D/ZIC1/TOMM20/PTTG1IP/SORT1/TOMM40/TIMM13/MIPEP/MF

MYC/TAF3/TAF13/NOC2L/CTNNBIP1/PPARGC1A/TAF4B/GTF2F2/ELOA/CDK4/MED25/TAF12/ELOC/TAF4/MAZ/SMARCA4/TP53/GTF2B/ZNHIT1/POLR2I/T

AK5/PDK4/ADCY2/PGM1/ABCD1/NUDT11/SLC4A4/PKLR/SULT2B1/IMPDH2/ENO2/CTPS1/BAAT/EPHA2/GPI/ENO1/GPAT4/FAR2/PPARGC1A/NME4/NME1/

SNAI2/FOXM1/MYC/SPRED1/SOX4/AURKB/PYCARD/MIF/GTSE1/NOP53/PTTG1IP/AURKA/DYRK3/TWIST1/ZMPSTE24/BCL2/BOP1/TFAP4/MYO6/MTOR/

TDRD9/CBX3/MYC/CDK2/LMNB2/BEND3/SETDB2/RLIM/DNMT3A/SAMD1/CENPV/TRIM28/CDYL/PCGF3/HP1BP3/LRIF1/TET1/SCMH1/HAT1/MBD3/SUV3

TUB/KIF17/SPEF2/KIF4A/AP3M2/TMEM201/NEFH/HSBP1/CDR2L/KIF3A/DYNC1I1/MAP2/LRPPRC/CLUAP1/IFT22/UXT/SUN1/DPCD/IFT172/KIF1B/DYNC2L

GPR161/TULP3/TEDC2/STIL/RACK1/ZIC1/GPC2/CDK20/TMED2/PAX6/TMEM231/TCTN2/CFAP410/CC2D2A/CENPJ/ENPP1/RPGRIP1L/CLUAP1/TCTN1/N

TUB/CFAP61/SPEF2/DNALI1/DNAAF5/RTTN/FNBP1L/MNS1/RILPL1/CROCC/ATAT1/RSPH1/WDR54/LPAR1/AURKA/WDR90/KIF3A/TBC1D7/TMEM237/GA

AK5/PDK4/ADCY2/PGM1/ABCD1/NUDT11/SLC4A4/PKLR/SULT2B1/IMPDH2/ENO2/BAAT/EPHA2/GPI/ENO1/GPAT4/FAR2/PPARGC1A/NME4/NME1/SNCA/

AK5/PDK4/ADCY2/PGM1/ABCD1/NUDT11/SLC4A4/PKLR/SULT2B1/IMPDH2/ENO2/CTPS1/BAAT/EPHA2/GPI/ENO1/GPAT4/FAR2/PPARGC1A/NME4/NME1/

WIPI1/EFNB1/MFN2/MTOR/AP4M1/PINK1/MAP1LC3B/ATG4B/ATG9A/UBXN2A/EHMT2/SNX7/TBC1D14/STBD1/PIP4K2B/ULK2/ULK1/SEC22B/PIP4K2C/ATG

TERT/APCDD1/CDH3/SOSTDC1/MSX2/TNFRSF19/FST/LHX2/HOXC13/KRT17/SOX9/DKK1/KRT16/KRT27/FARP2/NSUN2/FZD3/ZMPSTE24/BCL2/GAL

TERT/APCDD1/CDH3/SOSTDC1/MSX2/TNFRSF19/FST/LHX2/HOXC13/KRT17/SOX9/DKK1/KRT16/KRT27/FARP2/NSUN2/FZD3/ZMPSTE24/BCL2/GAL

TDRD9/NDN/CBX3/MYC/LHX2/NOC2L/TFAP2C/CDK2/HDAC4/CDKN1C/LMNB2/ZMPSTE24/NIBAN2/BEND3/DCAF13/SETDB2/RLIM/DNMT3A/TASP1/SAM

CFAP61/SPEF2/DNALI1/DNAAF5/FNBP1L/MNS1/RILPL1/CROCC/ATAT1/RSPH1/WDR54/LPAR1/WDR90/KIF3A/TBC1D7/TMEM237/GALNT11/CFAP161/TM

STXBP1/HSPD1/HSP90AB1/ATP1B2/SOX4/PEX6/DVL1/PPARGC1A/CCT3/CHP1/GLMP/OTUD3/CRTAP/NOP53/EPHA4/GET1/CCT6A/PYURF/HPS4/RUVBL

HOXD13/MSX2/TFAP2A/BMP7/TBX3/DLX5/DKK1/LRP5/TULP3/FRAS1/ZNF358/HOXC10/TWIST1/DLX6/TMEM231/HAND2/HOXA13/RPGRIP1L/SALL1/RARG

HOXD13/MSX2/TFAP2A/BMP7/TBX3/DLX5/DKK1/LRP5/TULP3/FRAS1/ZNF358/HOXC10/TWIST1/DLX6/TMEM231/HAND2/HOXA13/RPGRIP1L/SALL1/RARG

ZDHHC11B/MTCL1/RILPL1/SLC1A1/CHP1/VPS37D/RACK1/ZDHHC9/TOMM20/GOLPH3L/ICMT/TOMM40/GET1/TIMM13/NDUFA13/ZDHHC4/SSR2/SAMM

CENPF/GPSM2/CDC20/NEK2/KIF4A/TMOD1/CDCA8/RCC1/RTTN/DIAPH3/AURKB/ANLN/CENPA/TPX2/ASPM/STIL/PLK1/PRC1/KIFC1/CDK2/FARP2/NOP

CDK2/NPM2/ENDOG/CDC6/ZMPSTE24/RUVBL2/WRNIP1/LIG3/DSCC1/ESCO2/CIZ1/GMNN/EHMT2/MCM4/RFC3/TP53/WIZ/FAF1/PTK6/DHX9/CDK1/RFC

EYA1/MMP16/TFAP2A/BMP7/SATB2/HOXB9/HOXB8/HOXB6/TULP3/IRX5/HOXD3/HOXB7/CDK20/DLX1/EIF4A3/COL11A1/TWIST1/HOXC9/COL2A1/HOXB3/

IRX1/BMP7/SOX8/DLC1/PTK7/COBL/LHX2/SOX9/SEMA4C/CECR2/RGMA/TULP3/DVL1/IRX3/STIL/FZD1/CDK20/FZD3/TMED2/TWIST1/CC2D2A/DEAF1/DV

GPM6A/CDC42EP3/KIT/CDC42EP5/COBL/ANLN/FNBP1L/MNS1/CROCC/DPYSL3/FSCN1/EPS8L1/CDC42EP1/EPS8/MTOR/PALM/CENPJ/NLGN1/AGRN/R

GATA4/HOXD13/MSX2/HES2/DLL3/NKD1/TBX3/ACVR2B/HOXB9/GBX2/COBL/HOXB8/HOXC13/HOXB6/HES6/DKK1/LRP5/HEYL/TULP3/HEY1/HOXD3/CTN

ABCD1/SLC4A4/CCNB1/OPN3/IRS2/CHCHD4/CHCHD2/PPARGC1A/SLC25A23/SNCA/NDUFA4/NOP53/HDAC4/SLC25A33/SHMT2/COX7A1/MTOR/IRS1/M

TRIP13/HSPD1/MAD2L2/MORF4L2/RAD54L/NONO/AUNIP/UBQLN4/EXO1/HSF2BP/CCNB1IP1/ENDOG/HFM1/MEIOB/FANCB/PRKDC/RUVBL2/PSMC3IP/

PDK4/ABCD1/SULT2B1/BAAT/GPAT4/FAR2/SNCA/ACSF2/ACSS2/NUDT8/PPT2/PDK3/NUDT19/GCDH/ACSS1/PDHA1/ENPP1/HACD1/SUCLG1/ABHD14B/

PDK4/ABCD1/SULT2B1/BAAT/GPAT4/FAR2/SNCA/ACSF2/ACSS2/NUDT8/PPT2/PDK3/NUDT19/GCDH/ACSS1/PDHA1/ENPP1/HACD1/SUCLG1/ABHD14B/

PDK4/ABCD1/SULT2B1/BAAT/GPAT4/FAR2/SNCA/ACSF2/ACSS2/NUDT8/PPT2/PDK3/NUDT19/GCDH/ACSS1/PDHA1/ENPP1/HACD1/SUCLG1/ABHD14B/

TPCN2/DENND2A/WIPF3/WIPI1/RHOBTB3/CLTCL1/SORT1/SNX8/RAB6B/KIF16B/AP4M1/MAP2/SPAG9/VAMP3/PHETA2/YKT6/RBSN/UBE2O/TBC1D14/SN

EYA1/SNAI2/MYC/RAD54L/ZMPSTE24/BCL2/PRKDC/RAD51/DNMT3A/CLOCK/NABP2/RAD54B/BRAT1/HUS1/POLB/TP53/MSH2/ECT2/FANCD2/NHEJ1/EL

GPM6A/NR2E1/TUB/ATP8A2/TFAP2A/LAMA1/SOX8/ACVR2B/BMPR1B/VAX2/LHX2/SOX9/LRP5/SMARCD3/SLC1A1/GNB1/DLX1/SDK2/PAX6/NPHP4/RDH13

IRX1/TFAP2A/BMP7/SOX8/DLC1/PTK7/COBL/LHX2/SOX9/SEMA4C/CECR2/RGMA/TULP3/DVL1/IRX3/STIL/FZD1/CDK20/FZD3/TMED2/TWIST1/CC2D2A/D

NOX4/PDK4/GSTM1/EDNRB/ABCD1/GSTP1/HS3ST2/MICAL2/SULT2B1/BAAT/ARSG/TKTL1/SLC19A2/EXTL2/GPAT4/FAR2/CTH/SLC1A1/SNCA/ACSF2/ETH

EYA1/IRX1/ROBO2/GATA4/SOSTDC1/HOXD13/MSX2/HES2/BMP7/DLL3/GPR161/NKD1/TBX3/SATB2/ACVR2B/HOXB9/MICAL2/FOXD1/GBX2/FST/COBL/B

IRX1/BMP7/SOX8/DLC1/PTK7/COBL/LHX2/SOX9/SEMA4C/CECR2/RGMA/TULP3/DVL1/VEGFA/IRX3/STIL/FZD1/DAB2IP/CDK20/FZD3/TMED2/TWIST1/CC

PKP3/TBRG4/YBX1/AHCYL1/EIF4A3/RBPMS2/RBM11/FASTKD3/YBX3/EXOSC10/YTHDF2/SRPK3/DAZAP1/PAIP1/HNRNPA2B1/AKT1/PRKCA/SERBP1/RBM

AK5/PDK4/ADCY2/GMPR/PGM1/NQO1/ABCD1/NUDT11/SLC4A4/PKLR/SULT2B1/KCNAB2/IMPDH2/ENO2/BAAT/EPHA2/GPI/ENO1/GPAT4/FAR2/MDH2/PP

CDH3/STXBP1/WIPI1/SEPTIN5/AP3M2/CADPS/FNBP1L/SNCA/CDR2L/RAB3A/TMED2/KIF3A/MYO5A/DYNC1I1/F8A1/PINK1/MAP2/CADPS2/SEC16B/MLPH/

MAGEC2/FOXF2/CDC20/UBE2C/PINX1/HSP90AB1/SOX4/MAD2L2/CHP1/PLK1/BIRC7/RNF180/PTTG1IP/SPRY2/SEPTIN4/AIMP2/PPIA/TBC1D7/PSMD10/N

TERT/NOX4/KCNK2/CENPF/NEK2/POLD2/MYC/PINX1/HSP90AB1/AURKB/MAD2L2/CCT3/TYMS/CCT6A/FAAP20/WRNIP1/LIG3/DSCC1/NIBAN2/EXOSC10/

PDK4/EEF2/WTIP/BZW2/KRT17/SHMT1/SOX4/EIF3B/SNCA/EIF4EBP1/YBX1/EIF3E/RACK1/RGS2/PUS7/EPHA4/EIF4A3/TMED2/TYMS/SHMT2/FASTKD3/PR

EYA1/KIF26B/IRX1/HOXB13/CEACAM1/SNAI2/RYR2/GATA4/SOSTDC1/HOXD13/MSX2/TFAP2A/FOXF2/BMP7/LAMA1/WNK4/NKD1/TBX3/FLRT3/SOX8/CYP

MGP/EYA1/MSX2/MMP16/TFAP2A/BMP7/SATB2/ACVR2B/FREM1/HOXB9/SPEF2/BMPR1B/DLX5/HOXB8/SOX9/HOXB6/LRP5/TULP3/IRX5/HOXD3/HOXB7/B

TLE6/IGFBP2/SNAI2/CDH3/FZD9/SOSTDC1/AMER2/WNK2/NKD1/RBMS3/PTK7/DLX5/SOX9/DKK1/SEMA5A/SOX4/ASPM/MAD2L2/CTNNBIP1/FZD1/DAB2IP

EYA1/KIF26B/IRX1/RYR2/GATA4/SOSTDC1/MSX2/BMP7/LAMA1/WNK4/TBX3/SOX8/DLC1/MICAL2/FOXD1/GBX2/PTK7/COBL/MYC/LHX2/SOX9/SEMA4C/LR

NOX4/EYA1/KCNK2/FOXM1/NEK2/SPIRE2/MYC/PINX1/HSP90AB1/AURKB/MAD2L2/SMARCD3/MORF4L2/PPARGC1A/CCT3/CDK2/NPM2/AUNIP/UBQLN4/

ARMC3/KIF26B/TUB/KIF17/KIF25/CFAP61/KIF2C/SPEF2/SORD/KIF4A/DNALI1/DNAAF5/DNAH14/AP3M2/TMEM201/NEFH/MNS1/HSBP1/PLTP/KIFC1/CDR2

IRX1/ROBO2/GATA4/SOSTDC1/HOXD13/MSX2/HES2/DLL3/GPR161/NKD1/TBX3/ACVR2B/HOXB9/MICAL2/FOXD1/GBX2/COBL/BMPR1B/VAX2/HOXB8/LHX

TERT/DCLK1/RANBP17/PYGO1/PINX1/SOX9/TAF3/DVL1/NUP93/IPO4/CCT3/CHP1/PLK1/BYSL/ZIC1/NOP53/PTTG1IP/MAGED1/DNAJB6/CCT6A/LMNB2/TM

EYA1/ATP8A2/RYR2/GATA4/MMP16/TFAP2A/FOXF2/BMP7/TBX3/SATB2/HOXB9/MICAL2/GBX2/PTK7/DLX5/VAX2/HOXB8/SOX9/HOXB6/EPHA2/TULP3/IRX5

DCLK1/DENND2A/WIPF3/RHOBTB3/DIAPH3/ZFYVE9/CLTCL1/VPS37D/LAPTM4B/SORT1/SNX8/RAB6B/AP5S1/KIF16B/NSG1/SPAG9/VAMP3/PHETA2/RAB1

TYRP1/STXBP1/PMEL/ABCB6/RAB32/F2R/DIAPH3/VAV3/BACE2/AP3M2/FNBP1L/SNCA/VPS37D/RAB3A/LAPTM4B/RAB38/GOLPH3L/SORT1/TMED2/BCL2/H

TLE6/IGFBP2/SNAI2/CDH3/FZD9/SOSTDC1/AMER2/EDNRB/WNK2/NKD1/PYGO1/RBMS3/FOXD1/PTK7/DLX5/SOX9/DKK1/LRP5/SEMA5A/SOX4/ASPM/DVL

NR2E1/CHRNB2/EFNB3/ROBO2/CNTN4/ATP8A2/DCLK1/SEMA6A/BMP7/STXBP1/FLRT3/RAC3/CHL1/NDN/CRPPA/LPAR3/EDN3/MAP6/FOXD1/GBX2/EPH

GPM6A/NR2E1/TUB/ATP8A2/TFAP2A/FOXF2/BMP7/LAMA1/ANGPTL7/SOX8/ACVR2B/BMPR1B/VAX2/LHX2/TMOD1/SOX9/SPRED1/LRP5/COL8A1/EPHA2/S

GPM6A/ROBO2/ATP8A2/BMP7/CDC42EP3/KIT/CDC42EP5/SS18L1/NEGR1/LPAR3/MAP6/PTK7/COBL/LRP8/TUBB2B/SEMA5A/ATP1B2/RGMA/L1CAM/ANLN

TLE6/TERT/IGFBP2/SHISA2/GPC4/SNAI2/APCDD1/CDH3/FZD9/SOSTDC1/AMER2/EDNRB/WNK2/NKD1/PYGO1/RAC3/RBMS3/FOXD1/PTK7/DLX5/VAX2/S

TLE6/TERT/IGFBP2/SHISA2/GPC4/SNAI2/APCDD1/CDH3/FZD9/SOSTDC1/AMER2/EDNRB/WNK2/NKD1/PYGO1/RAC3/RBMS3/FOXD1/PTK7/DLX5/VAX2/S

NR2E1/CHRNB2/EFNB3/ROBO2/CNTN4/ATP8A2/DCLK1/SEMA6A/BMP7/STXBP1/FLRT3/RAC3/CHL1/NDN/CRPPA/LPAR3/EDN3/MAP6/FOXD1/GBX2/EPH

LRRC15/CCL3/ST3GAL1/ZC3H12A/RESF1/TRIM34/CXCL8/PARD6A/CLDN9/PROX1/SERPINB3/CCL2/CH25H/HMGA2/CD209/SIGLEC1/AGTR1/LDLR/IFNB1

TRH/TRDN/NOS3/GPR35/INS/NEUROG1/PPP3R2/ADRA2A/LILRB2/NTSR1/CRHBP/SLC43A2/SERPINE2/TLR2/STC1/PKDCC/MTNR1B/PTGS2/PLA2R1/GA

ABCB1/FOLR1/SSTR4/GRM1/SLC7A11/NPY5R/PLA2G2E/SLC16A11/SLC32A1/SLC47A1/SLC10A2/PLA2G3/SLC16A14/SLC22A7/SLC26A4/FOLR2/SLC26A3/SL

ABCB1/FOLR1/SSTR4/GRM1/SLC7A11/NPY5R/PLA2G2E/SLC16A11/SLC32A1/SLC47A1/SLC10A2/PLA2G3/SLC16A14/SLC22A7/SLC26A4/FOLR2/SLC26A3/SL

MIXL1/FGFBP1/CER1/GPLD1/CD248/NOS3/FBXW7/BMPER/ZC3H12A/ID1/FGF7/ITGA3/PROX1/VASH1/SMOC2/PTN/TACSTD2/RRAS/PLXND1/MMRN2/TE

EXPH5/CD33/SLC30A8/SFN/P2RX7/ISL1/IL13/KCNN4/CHP2/PRR5L/TMEM30B/IL1A/NR1H4/UCN3/PRKCB/GPER1/CASR/F2RL1/ADCY8/IGF1/GPLD1/TRH/F

SLC8A3/CHI3L1/KSR1/ADRB2/PDGFRB/RASSF2/MMD/PAK1/LAT/NRG1/ELANE/NTRK3/TPD52L1/ADCY8/IGF1/FBXW7/CDKN2A/INS/PYDC1/ADRA2A/TLR3

AQP2/SLFN14/ADCY1/SLC30A8/ITPKA/KCNA1/SLC6A1/P2RX7/SLC30A2/TCIRG1/TRPV6/SLC26A5/KCNMB4/CHP2/SYT4/CNGA3/SYT13/MUC2/KCNK3/IL1A

SLC6A1/P2RX7/SLC9A2/SLC6A11/SLC17A6/SCN2A/SLC9A4/BPIFA1/SLC34A3/CHP2/SCN10A/SLC10A1/SCN7A/SLC6A3/SLC8A3/ADRB2/SLC9A3/SLC34A1/PT

NKX2-1/CRISPLD2/IGFBP5/ABCA3/NOS3/STRA6/FBXW7/NOG/PGR/FGF7/ITGA3/PROX1/ASCL1/WNT2/FOXA1/KDR/TCF21/PKDCC/HSD11B1/GNG8/NOD

CST11/CFLAR/UCN3/GPER1/IL22/SSTR2/GJB2/GHSR/HSD3B1/NEFL/ALPL/ATP1A3/PAK1/SLIT3/LMO3/ABCA3/KDM4C/RHOXF1/PGR/NR3C2/SRD5A2/NKX

CARD18/AHSG/CDKN2A/SERPINB5/INS/PYDC1/SERPINB3/FETUB/NEIL1/HP/ANGPTL4/SERPINB8/SERPINA11/DPEP1/SPINK2/PPARG/SERPINE2/HRG/S

GLI1/ATOH1/KRT4/OVOL3/STK4/CLRN2/TCHH/REG3A/SPRR2B/KRT6A/SPRR4/CERS3/SPRR3/DLX3/WNT16/EXPH5/BCR/DSG4/ANXA1/PRKCH/SFN/KRT

MPO/PXDNL/AKR1C3/NCF2/PLA2R1/CLEC7A/NOX3/EDN1/HBZ/CYP1A1/TYROBP/HBD/SOD2/F2/LIPA/THBS1/SYK/CYP1B1/NCF4/GCH1/TGFB1/FBLN5/IN

GRM1/DRD5/DRD1/SLC7A11/NPY5R/FAM107A/NGFR/CALB1/CHRNB3/CAMKV/RAPSN/CA2/BCR/MYOF/ADCY1/ITPKA/LARGE1/CX3CL1/CPLX2/ROR2/SL

GRM1/DRD5/DRD1/SLC7A11/NPY5R/FAM107A/NGFR/CALB1/CHRNB3/CAMKV/RAPSN/CA2/BCR/MYOF/ADCY1/ITPKA/LARGE1/CX3CL1/CPLX2/ROR2/SL

CHP2/SYT4/SYT13/KCNK3/CFLAR/B2M/SLC34A1/SCN5A/LCE1D/ADCY8/CER1/GPLD1/SYT9/RASA4B/AOC1/GUCY1B1/GLRA1/CRHBP/DPEP1/KCNC2/RAS

LGI4/DEFB1/PGR/ASCL1/PTH1R/GLDN/PPARG/WEE2/FOXA1/WNT10B/KDR/IL21/TFCP2L1/DAZL/BAIAP3/KCNQ3/SOX18/AXL/TUBB8/ADGRB3/CLEC7A/BF

CYP26A1/CYP2A13/SLC28A3/MEF2C/ABCA1/GSTA2/UGT2B7/SULT1C3/ABCB1/LPO/CYP2B6/AS3MT/SLC22A7/CYP2A7/SULT1A2/CES3/FMO1/GSTA3/UGT2

CDKN2A/TNN/SKAP1/VWF/PLEKHA2/ITGA3/MUC4/STRC/OTOA/EGFLAM/TACSTD2/RRAS/TEK/FERMT3/SIGLEC1/LIMCH1/HRG/ECM2/KDR/MSLNL/ST6

KCNA1/SMAD7/ADRB1/P2RX2/KCND2/INSYN2A/ABCB5/P2RX7/GNA15/SCN2A/SLC26A5/KCNN4/KCNMB4/BCO2/KCNA4/GABRA2/CHRNA7/SCN10A/GRIN

SGCG/CAMK2D/BMP10/APELA/KCNE3/CORIN/SCN11A/KCNE4/GRK2/SMAD7/ADRB1/KCND2/ISL1/SCN2A/CLIC2/GJD3/SCN10A/SCN7A/CTNNA3/SLC8A3

CDKN2A/TNN/SKAP1/PLEKHA2/ITGA3/MUC4/STRC/OTOA/RRAS/TEK/FERMT3/SIGLEC1/LIMCH1/HRG/ECM2/KDR/MSLNL/ITGAD/ITGA8/NRP1/ITGA4/CC

SLC8A3/CHI3L1/KSR1/ADRB2/PDGFRB/RASSF2/MMD/PAK1/LAT/NRG1/ELANE/NTRK3/TPD52L1/ADCY8/IGF1/FBXW7/CDKN2A/INS/PYDC1/ADRA2A/TLR3

DTNA/TBX2/KBTBD13/CAMK2D/CACNA1S/DRD1/PTGER3/BMP10/KCNE3/TBXA2R/SCN11A/UTS2/MYOF/KCNE4/LARGE1/GRK2/KCNA1/SMAD7/P2RX2/K

CCL15/NCOA4/MT3/SLC40A1/GRM1/DRD5/CAMK2D/CACNA1S/DRD1/CALB1/PRND/SCNN1G/CA2/LACC1/SLC30A8/PML/SLC26A3/RGN/GP1BB/CX3CL1/

NR1H4/MAFA/UCN3/PRKCB/GPER1/CELA2A/GHSR/CASR/F2RL1/ADCY8/GPLD1/TRH/HNF4A/GPR68/ADRA2A/HNF1B/NR0B2/TFAP2B/BAIAP3/RPH3AL/

SRGAP3/KNDC1/ARHGAP44/PREX2/RERG/LAT/SH3BP1/F2RL1/DOK1/IGF1/RAPGEF3/SIPA1/GPR35/CDKN2A/RASA4B/ADRA2A/ARHGEF15/DOK5/RHOU/

VIP/ISL1/TCIRG1/VGF/PER2/CPLX1/RAB8B/SPINK1/NR1H4/MAFA/UCN3/PRKCB/GPER1/CELA2A/GHSR/CASR/F2RL1/ADCY8/GPLD1/TRH/SLC38A3/HNF4

CLIP3/STEAP4/ZC3H12A/KCNJ8/ALOXE3/INS/ZNF385A/PRDM16/EGR2/WNT5B/CCDC3/TLCD3B/RORC/HMGA2/PPARG/NUDT7/DLK2/TFAP2B/WNT10B/

BMP10/APELA/KCNE3/CORIN/KCNE4/GRK2/SMAD7/ADRB1/KCND2/ISL1/CLIC2/GJD3/SCN10A/CTNNA3/SLC8A3/GLP1R/CACNB2/ATP1A3/CACNA1G/ADF

CAMK2D/BMP10/KCNE3/SCN11A/KCNE4/GRK2/SMAD7/SCN2A/CLIC2/SCN10A/SCN7A/CTNNA3/CACNB2/CACNA1G/ADRA1B/SCN5A/PPP1R13L/NOS1AI

CCR3/FGF10/HMX2/APELA/ACVRL1/EPGN/LRG1/GRN/CCL11/VIP/HPN/NRARP/PRKD2/F3/SPINK1/MDK/CFLAR/CAV2/GHSR/SCN5A/VASH2/FGFBP1/IGF

CSF1R/MAPK15/VEGFC/PDGFRA/FGFR2/HCK/FLT3/TXK/CAMK4/PTK2B/DAPK2/PDGFC/JAK2/STK17B/PIM2/LYN/EPHA1/BMX/STK26/LRRK2/LCK/MAP4K

PTH/PF4/GRM8/CRHR1/PSAPL1/RIMS2/CRHR2/LHCGR/GNAT1/MRAP2/GPR4/CALCA/PDE4A/MCHR1/CHRM5/GCG/DRD3/CHRM1/GPR88/GPR176/MC

FSHB/PLB1/PLA2G10/BCO1/CYP11B1/HSD11B2/ALOX12B/PTGS1/CYP1A2/ABCA4/CYP4F2/SCNN1B/MGLL/ADH4/CYP2A13/CYP17A1/CYP4Z1/ALDH8A1/RD

SLC27A6/LBP/SLC51A/C1QTNF1/ESYT3/NKX3-1/GRAMD1C/CIDEA/NTSR1/TTC39B/FABP2/APOC4/APOL5/NMUR2/NR0B2/PPARG/AGTR1/ATP8B4/LDLR/

ZC3H12A/INS/ZNF385A/CCDC3/TLCD3B/RORC/PPARG/DLK2/WNT10B/JDP2/WIF1/ZFP36/FRZB/PTGS2/CCN4/PIM1/RUNX1T1/SOD2/FFAR4/PTPRQ/CEI

WNT10B/HSPA1B/TRABD2A/TENM1/CLEC7A/IFNG/CDH5/NPHS1/P2RY12/SYK/CCL24/EVL/TAL1/ISG15/HCK/TGFB1/GBP5/CDH17/PTK2B/CCL21/LCP1/MF

CARD18/AHSG/SERPINB5/INS/SERPINB3/FETUB/SERPINB8/SERPINA11/DPEP1/SPINK2/SERPINE2/HRG/SERPINA6/SERPINA1/SERPIND1/SERPINF2/CS

GREM1/SLURP2/CBLB/SOCS5/EPGN/PLAU/CGA/CBLC/DLGAP2/GREM2/ADRB2/FBXW7/CACNG2/NOG/NLGN3/ADRA2A/DLGAP3/BEGAIN/CCL2/CRHBP/

BAIAP3/KCNQ3/RPH3AL/ANK1/SYNGR3/P2RX1/IFNG/STX11/ERC2/EXOC3L1/FCGR2B/SYNGR2/LIN7A/SYT8/CCR2/SYTL5/FGA/SYK/RIMS4/TNFAIP2/TRA

DMRT1/MARVELD2/PLEC/KDF1/F2RL1/PRDM1/RAPGEF3/SIPA1/TNMD/PGR/PROX1/PALLD/WNT5B/INSM1/FOXA1/KDR/TFCP2L1/STC1/SOX18/FRZB/P

CACNG2/DNM1/NLGN3/CXCL8/STAB1/FCHSD2/ARRB2/RAB31/SIGLEC1/LDLR/INPP5F/CACNG3/SYNE1/APOA5/PLA2R1/SNAP91/ADM/ITGA4/STAB2/IL4/

TBXA2R/CLDN3/ANXA1/PLAU/HTN3/GRN/ADAMTS18/PROC/ADTRP/F11/F3/MDK/SCARF1/MYLK/KRT1/F2RL1/KIAA0319/NOS3/STAT3/DUOX2/ADRA2A/F

CHP2/SYT4/SYT13/KCNK3/B2M/SLC34A1/SCN5A/LCE1D/ADCY8/CER1/GPLD1/SYT9/RASA4B/AOC1/GLRA1/CRHBP/DPEP1/RASAL1/CPNE6/MT1H/LGMN/

HTRA2/AGER/S100A9/F3/PRSS22/GRIN2B/GPER1/RFPL1/STAT3/BCL2L10/MEFV/SERPINB3/NKX3-1/PPARG/LGMN/MAP3K5/CLEC7A/VSIR/NODAL/GRIN2

CAMK2D/KCNE3/KCNE4/ADRB1/ISL1/GJD3/SCN10A/CTNNA3/CACNB2/CACNA1G/ADRA1B/SCN5A/NOS1AP/KCNJ3/KCND3/SCN2B/KCNJ8/KCNQ1/KCN

TRIM21/GRK2/TRIM10/AVPR1B/TRIM38/GPR15/NCAM1/SLC10A1/NECTIN4/AVP/RFPL1/CAV2/MRC1/LRRC15/TRIM34/CXCL8/CLDN9/SERPINB3/CH25H/

CAV2/RGS16/PREX2/SH3BP1/CCL1/F2RL1/NTRK3/CCL3/RAPGEF3/RGS8/ARHGEF15/CCL2/TBC1D21/RASGRP3/CCL7/RASGEF1B/RGS7/DOCK11/CCL25

NR1H4/UCN3/PRKCB/GPER1/CELA2A/GHSR/CASR/F2RL1/ADCY8/GPLD1/TRH/HNF4A/GPR68/ADRA2A/NR0B2/TFAP2B/BAIAP3/RPH3AL/MTNR1B/ITPR

CHRNA7/GRIN2B/SLC8A3/ADRB2/HTR3E/NRXN1/GRIN1/NLGN4X/CHRNA9/NLGN3/GLRB/CHRNA3/GLRA1/HTR3C/ARRB2/HTR3D/SYNE1/P2RX1/ZACN

TNFRSF10A/CYLD/SKIL/HSPA1B/HYAL2/FAIM2/MAL/NRP1/IFNG/TNFRSF10C/FGFR3/BMP5/IL7/HMOX1/CASP10/FGA/THBS1/IL2/TGFB1/TNFRSF1B/SFR

CAV2/RGS16/ARHGAP44/PREX2/SH3BP1/CCL1/F2RL1/NTRK3/CCL3/RAPGEF3/RGS8/ARHGEF15/PLXNA2/CCL2/PLXND1/NTRK2/TBC1D21/ALS2CL/RAS

GLI1/KRT4/STK4/ID2/REG3A/LIMS2/BCL2L1/OR51E2/TBX2/CCR3/FGF10/SIDT2/HMX2/A4GNT/APELA/NGFR/ACVRL1/EPGN/WNT16/RGCC/SIX1/RGN/LRG

SGCG/CAMK2D/BMP10/APELA/KCNE3/CORIN/SCN11A/KCNE4/GRK2/SMAD7/ADRB1/KCND2/ISL1/SCN2A/CLIC2/GJD3/SCN10A/SCN7A/CTNNA3/SLC8A

KIAA0319/STRBP/STRA6/CHRNA9/CSRP3/NEUROG1/TLR3/STRC/KCNQ1/TNFRSF8/KCNC1/PTN/TLR5/TNFRSF10A/SERPINE2/MPO/TNFRSF11A/KCN

P2RX7/VIP/ISL1/TCIRG1/VGF/PER2/IL13/CPLX1/ADTRP/RAB8B/KCNN4/SYT4/TVP23C/CBLN4/IL1A/NR1H4/MAFA/UCN3/PRKCB/GPER1/CELA2A/GHSR/C

ADRB1/PER2/IL13/PTH2R/DIO2/ADRB2/ALPL/LCN2/PRDM16/YBX2/TSHR/GADD45G/ESRRG/IL4/FFAR4/KSR2/CCR2/FABP4/SYK/GHRL/IL4R/CMKLR1/UC

VASH2/FGFBP1/IGF1/STAT3/FBXW7/VDR/NOG/APLNR/POLD4/NKX2-5/TNMD/SERPINB5/PGR/FGF7/PROX1/VASH1/PTN/NKX3-1/CCL2/TACSTD2/SULF1

KLB/FGFBP1/NOG/FGF20/CXCL8/FGF7/SMOC2/CCL2/SULF1/HYAL2/ZFP36/FGFR3/HHIP/THBS1/FGFR2/SULF2/SFRP1/WNT5A/DLL4/FLRT2/FGF2/GATA

SCN9A/HRC/ACTN3/DTNA/KBTBD13/CAMK2D/CACNA1S/BMP10/KCNE3/SCN11A/KCNE4/LARGE1/GRK2/SMAD7/STRIT1/SCN2A/CLIC2/SCN10A/SCN7A/

P2RY6/NTSR1/PLCB2/RIT2/TMSB4Y/SYNE1/LACRT/TRPM2/ITPR1/RYR3/TMSB4X/JPH3/MYZAP/F2/CASQ1/CD4/PTK2B/CACNA1C/FASLG/THY1/CCL21/L

P2RX7/VIP/ISL1/TCIRG1/VGF/PER2/IL13/CPLX1/ADTRP/RAB8B/KCNN4/SYT4/TVP23C/CBLN4/IL1A/NR1H4/MAFA/UCN3/PRKCB/GPER1/CELA2A/GHSR/C

DMRT3/ISL2/PRDM1/STAT3/NKX2-5/NEUROG1/WNT9A/PROX1/ASCL1/HNF1B/WNT5B/GCM1/WNT2/CYLD/TENM4/PPARG/NKX6-2/FOXA1/WNT10B/OLI

TLR2/FOXP1/CDA/SPP1/AKR1C3/DSG2/TAT/SLC5A5/EDN1/TXNIP/CARD9/AKR1C2/SLIT2/KLF9/LIPA/THBS1/PTGER2/CYP1B1/PTAFR/TRERF1/ADCY5/PT

BAIAP3/RPH3AL/P2RX1/FCGR2B/SYT8/CCR2/SYK/RIMS4/PTAFR/IL4R/SLC4A8/PRAM1/LGALS9/SV2B/CD300A/GATA2/SYT1/WNT7A/LYN/SPI1/ITGB2/LRR

TRIM21/GRK2/TRIM10/AVPR1B/TRIM38/GPR15/NCAM1/SLC10A1/NECTIN4/AVP/RFPL1/CAV2/MRC1/LRRC15/TRIM34/CLDN9/SERPINB3/CH25H/CD209/

P2RX2/INSYN2A/P2RX7/CHRNA7/GRIN2B/SLC8A3/ADRB2/HTR3E/NRXN1/GRIN1/NLGN4X/CHRNA9/NLGN3/GLRB/CHRNA3/GLRA1/NTSR1/HTR3C/ARR

P2RX7/VIP/ISL1/TCIRG1/VGF/PER2/IL13/CPLX1/ADTRP/RAB8B/KCNN4/SYT4/TVP23C/CBLN4/IL1A/NR1H4/MAFA/UCN3/PRKCB/GPER1/CELA2A/GHSR/C

HSD17B2/CDX2/CSF2/GJB2/GHSR/CYP27B1/VASH2/PRDM1/VDR/VASH1/LHX4/PTN/GCM1/WNT2/PPARG/PLAC1/SPP1/STC1/PTGS2/ADM/NODAL/HTRA
SKAP1/PLEKHA2/ITGA3/EGFLAM/RRAS/TEK/HRG/ECM2/KDR/VWC2/FBLN2/APOA1/NRP1/CCL25/CSF1/DMP1/FGA/ABI3BP/PTK2B/SFRP1/FGB/JAK2/ITGA
KCNQ1/SLC27A6/LBP/SLC51A/C1QTNF1/ESYT3/NKX3-1/GRAMD1C/CIDEA/NTSR1/TTC39B/FABP2/APOC4/APOL5/NMUR2/NR0B2/PPARG/AGTR1/ATP8B
ID1/LCN2/PRDM16/YBX2/CIDEA/LNPEP/TSHR/WNT10B/GADD45G/ESRRG/IL4/FFAR4/KSR2/CCR2/FABP4/SYK/GHRL/IL4R/CMKLR1/NR1H3/UCP2/PDGF
CCR3/FGF10/APELA/ACVRL1/LRG1/CCL11/VIP/NRARP/PRKD2/F3/MDK/CAV2/GHSR/VASH2/FGFBP1/STAT3/APLNR/POLD4/FGF7/PROX1/TEK/AGTR1/KD
APOB/PLA2G3/FITM1/P2RX7/AVPR1B/PON1/IL1A/MYB/GALR1/ERFE/ABCA3/ZC3H12A/C1QTNF1/NKX3-1/CIDEA/NTSR1/APOC4/PPARG/ACSL5/SPP1/TNF
DDX60/TREM2/SLC15A3/GBP5/RSAD2/SLC15A2/TLR9/PTPN22/PLCG2/BTK/LILRA4/TLR6/BIRC3/TLR4/OASL/NLRC3/FCRL3
ID1/LCN2/PRDM16/YBX2/CIDEA/LNPEP/TSHR/WNT10B/GADD45G/ESRRG/IL4/FFAR4/KSR2/CCR2/FABP4/SYK/GHRL/IL4R/CMKLR1/NR1H3/UCP2/PDGF
ZC3H12A/RESF1/TRIM34/CXCL8/CLDN9/PROX1/SERPINB3/CCL2/CH25H/HMGA2/CD209/SIGLEC1/AGTR1/LDLR/IFNB1/HSPA1B/HYAL2/AXL/ATG16L2/NR
EGR2/PTN/TENM4/SERPINE2/LDLR/NKX6-2/OLIG2/TLR2/CLCF1/ASCL2/F2/VEGFC/TREM2/IL6/TGFB1/LRP2/TNFRSF1B/CYSLTR1/LYN/LTA/RNF112/IL34/
ZFP36/FRZB/NCOA3/REG3G/IFNG/NODAL/AQP3/ATOH8/MYCL/CDH5/TP63/GDF3/GDF2/DLL1/STAT1/ZBED2/SPRED3/TLR9/BMP2/IL1B/FGF2/TNF/SFRP
DRD1/FAM107A/CALB1/RAPSN/CA2/ADCY1/LARGE1/ROR2/CLSTN2/SHANK2/GFAP/CX3CR1/LAMA2/RASGRF2/AGER/CHRNA7/GRIN2B/SLC8A3/GPER1
VASH2/PRDM1/VASH1/GCM1/WNT2/ADM/NODAL/HAND1/BMP5/ASCL2/CEBPA/IL10/SOCS3/PLG/JUNB/FGFR2/EOMES/GATA2/TFEB/RSPO3/SPINT2/CIT
CDKN2A/ZC3H12A/PYDC1/ADORA3/ARRB2/CYLD/IL10/CYP1B1/NFKBIA/CMKLR1/NLRP3/NLRC5/BCL3/HAVCR2/KLF4/SPI1/IRAK2/FOXP3/NLRC3/AIM2
IGFBP7/SLC10A1/SPINK1/GRIN2B/SLC6A3/UCN3/AVP/PRKCB/HSD3B1/NEFL/SLIT3/GRIN1/ADCY8/GPLD1/CCL3/GLRA1/GRAMD1C/P2RY6/CRHBP/RGS1
NR1H4/UCN3/PRKCB/GPER1/CASR/ADCY8/GPLD1/TRH/GPR68/NR0B2/BAIAP3/RPH3AL/ITPR1/GIPR/ANO1/F2/GHRL/TUNAR/ABAT/FFAR1/JAK2/VSNL1/
P2RX7/VIP/ISL1/TCIRG1/TRPV6/VGF/PER2/CCKAR/POMC/CPLX1/RAB8B/KCNMB4/SYT4/SYT13/UNC13A/SPINK1/UNC13C/IL1A/NR1H4/MAFA/UCN3/MYE
HERC6/KDR/HYAL2/BATF/ZFP36/LIPA/RBM47/PDGFRA/TAL1/TGFB1/FLT3/SFRP1/GATA2/SMPD3/MEOX1/LYN/SPI1/TIFAB/INHBA/PLD4/PLEK/GATA3/PTP
TNFRSF10A/SKIL/FAIM2/MAL/TNFRSF10C/BMP5/HMOX1/CASP10/FGA/THBS1/SFRP1/FGB/FASLG/ICAM1/SPI1/HGF/TNF/TMC8
HYAL2/TNFRSF11A/TCIM/MAP3K5/CCNI2/TENM1/IFNG/IRGM/EDN1/THBS1/HEG1/DEFB114/PIK3R5/FLT3/PKIB/WNT5A/CD300A/PDGFC/PKIA/CD40/THY
ID1/LCN2/PRDM16/YBX2/CIDEA/LNPEP/TSHR/WNT10B/GADD45G/ESRRG/IL4/FFAR4/KSR2/LIPA/CCR2/FABP4/SYK/GHRL/SCT/IL4R/CMKLR1/BMP8A/NR
NRXN1/SLITRK5/IL1RAP/EPHB1/LRRN1/NLGN3/LRRTM3/NTRK2/LINGO2/ADGRB3/EPHB2/SEMA4A/VSTM5/GHRL/SLITRK4/WNT7A/FLRT2/AMIGO2/SEM
SCGB1A1/SSTR4/TBX2/SSTR5/FAM107A/SCNN1G/WNT7B/ANXA1/UCP3/ISL1/IGFBP7/CFLAR/UCN3/GPER1/IL22/SSTR2/GJB2/GHSR/HSD3B1/NEFL/ALPL
FOLR1/ASTL/SPACA4/PTX3/CNTN2/PRSS55/MYPN/AMIGO1/FOLR2/TEX101/PRSS37/NTM/SPACA3/SFTPA1/IZUMO1R/CLGN/ROBO3/DSCAML1/SPPL2C/
SNAP25/KCNMB2/TRPM5/KCNV1/KCNE3/AMIGO1/KCNE4/LARGE1/KCNA1/KCND2/KCNJ1/SLC9A2/HPN/SLC17A6/KCNN4/SLC9A4/KCNMB4/KCNV2/GAL
TNMD/STAB1/VASH1/SULF1/TEK/PPARG/HRG/APOH/ADGRB3/PDE3B/DCN/SEMA4A/CCR2/THBS1/GDF2/GHRL/STAT1/THBS4/KLF2/TIE1/ALOX5/FASLG
AKR1B10/SULT1E1/ENDOU/ADTRP/BCO2/ACOX2/APOC3/CYP26C1/PLA2G4F/CLPS/SCARF1/PNLIPRP3/PLA2G4B/CYP27B1/PLPP6/ABHD16B/GPLD1/CYP
ZDHHC15/LGI1/CACNG3/MARVELD3/NPTX1/GRIN2A/MAP1A/CDH5/NPHS1/HEG1/STX1B/PECAM1/ZDHHC2/C1QL2/WNT5A/ILDR1/PCLO/TJP3/WNT7A/C
GPLD1/CD248/CCL3/FBXW7/VDR/CLIP3/CDKN2A/ZC3H12A/TLR3/ASCL1/BTG1/TNFRSF8/NKX3-1/CCL2/NTSR1/TNFRSF10A/CYLD/PPARG/SKIL/HRG/FO
ADAMTS18/PROC/ADTRP/F11/MDK/KRT1/KIAA0319/NOS3/STAT3/FIGNL2/SH2B3/TNR/C1QTNF1/SERPINE2/HRG/EPPK1/INPP5F/SPP1/APOH/SERPINF2/
CCL2/ANGPTL4/TEK/KDR/AKR1C3/MAP3K5/ZFP36/PLA2R1/ITGA4/IL4/CDH5/HMOX1/FGA/THBS1/IL10/IL6/TGFB1/FGB/FASLG/GATA2/JAK2/CD40/ECSCR/
SRGAP3/ARHGAP44/PREX2/SH3BP1/F2RL1/IGF1/SIPA1/GPR35/RASA4B/ARHGEF15/RHOU/ITGA3/CDC42SE2/ARHGAP19/GPR55/ABRA/TEK/ARHGAP23/
APOB/PLA2G2E/APOC1/PLA2G3/CES3/APOC3/SOAT2/NR1H4/DGAT2/PRKACG/GPLD1/PCSK6/AGTR1/LDLR/MPO/APOA5/APOA1/HMOX1/LIPC/LIPA/NCEH
NFKBIE/CER1/LIME1/TRDN/CCL3/FBXW7/ZC3H12A/APLNR/LTBP1/CIDEA/P2RY6/NTSR1/TTC39B/PLCB2/APOC4/PPARG/RIT2/TMSB4Y/SYNE1/LACRT/F
TRIM8/ITGAV/TRIM6/KRT6A/PTX3/TRIM21/TRIM10/TRIM38/RFPL1/LRRC15/TRIM34/CXCL8/CH25H/TRIM59/CD74/IFITM1/CD4/LGALS9/TRIM22/HS3ST5/
GHSR/F2RL1/ADRA2A/CRHBP/MTNR1B/EDN1/FFAR4/IL11/GHRL/SFRP1/BMP8A/UCP2/VSNL1/IL1B/INHBA/LIF/ADIPOQ
CAMK2D/DRD1/KCNE3/SCN11A/KCNE4/KCNA1/KCND2/GNA15/SCN2A/KCNMB4/SCN10A/SCN7A/CTNNA3/GPER1/PTPN3/CACNB2/CACNA1G/SCN5A/N
KCNMB2/HTR1A/TBXA2R/AVPR1B/PER2/KCNMB4/AVP/ADRA1B/CASR/ADRA2A/AGTR1/MTNR1B/P2RX1/PTGS2/ADRA2B/ADM/EDN1/ITGA4/ADRA1A/FG
NKX2-1/CRISPLD2/IGFBP5/ABCA3/NOS3/STRA6/FBXW7/NOG/PGR/FGF7/ITGA3/PROX1/ASCL1/WNT2/FOXA1/KDR/TCF21/PKDCC/HSD11B1/NODAL/ADA
APOC3/SOAT2/NR1H4/RORA/DGAT2/ABCG5/MINAR2/HNF4A/ACSM1/TTC39B/ACSM3/LDLR/MALL/MALRD1/ABCB11/APOA5/APOA1/ABCG4/LIPC/FABP4/
VIP/ISL1/TCIRG1/VGF/PER2/CPLX1/RAB8B/SPINK1/NR1H4/MAFA/UCN3/PRKCB/GPER1/CELA2A/GHSR/CASR/F2RL1/ADCY8/GPLD1/TRH/HNF4A/CPE/SL
KCNJ1/P2RX7/TCIRG1/STRIT1/PER2/IL13/DHRS7C/SLC26A5/KCNN4/BPIFA1/CLIC2/GALR2/CHP2/SCN10A/GRIN2B/CYBA/PRKCB/ADRB2/GPER1/SLC34A
KLF13/HSPA1B/MAFB/AXL/ZFP36/SLC4A1/CSF1/HMOX1/RHEX/LIPA/CCR2/TAL1/STAT1/ISG15/IL6/KLF2/MERTK/GATA2/GCNT4/JAK2/LYN/SPI1/FOXP3/HC
CLDN3/PLAU/ADAMTS18/PROC/ADTRP/F11/KRT1/NOS3/FIGNL2/SH2B3/C1QTNF1/SERPINE2/HRG/EPPK1/APOH/SERPINF2/MYOZ1/EDN1/F2/FGA/THBS
KIAA0319/IGF1/CCL3/FBXW7/NOG/ZC3H12A/GPR68/TLR3/PROX1/TNR/PTN/GPR55/PGLYRP2/WEE2/LDLR/NKX6-2/IFNB1/MAFB/SPP1/FRZB/SLIT1/NRP1/

KCNE3/SCN11A/KCNE4/SCN2A/SCN10A/SCN7A/CTNNA3/CACNB2/CACNA1G/SCN5A/NOS1AP/KCNJ3/KCND3/SCN2B/KCNJ8/KCNQ1/FGF12/DSG2/SCN
CRHBP/ZP4/CACNB4/RAB31/BAIAP3/RPH3AL/SYNGR3/P2RX1/STX11/ERC2/FCGR2B/SYNGR2/SYT8/CCR2/SYK/RIMS4/PTAFR/STX1B/OTOF/HCK/IL4R/S
P2RX7/ERAP1/PTPN3/GPLD1/ADRA2A/IFNG/IL10/TIMP3/TNFRSF1B/ADAM8/PRKCQ/IL1B/ADAM19/TNF/TIMP4
SNAP25/KCNMB2/TRPM5/DRD1/KCNV1/KCNE3/AMIGO1/KCNE4/LARGE1/KCNA1/KCND2/KCNJ1/VIP/SLC9A2/HPN/SLC17A6/KCNN4/SLC9A4/KCNMB4/K
VIP/ISL1/TCIRG1/VGF/PER2/CPLX1/RAB8B/SPINK1/NR1H4/MAFA/UCN3/PRKCB/GPER1/CELA2A/GHSR/CASR/F2RL1/ADCY8/GPLD1/TRH/HNF4A/CPE/SL
P2RX7/AVPR1B/PON1/IL1A/MYB/GALR1/ERFE/ABCA3/C1QTNF1/NKX3-1/NTSR1/PPARG/ACSL5/SPP1/TNFRSF11A/PLA2R1/APOA1/EDN1/ABCG4/GHRL/T
ITGAM/DAO/CHDH/HTR1A/DRD1/SLC7A11/MOXD1/SULT1A2/AMD1/AOC3/PNMT/PAOX/NR1H4/SLC6A3/ATCAY/TRH/ACMSD/AOC1/INS/HAAO/NAAA/INS
NKX2-1/CRISPLD2/IGFBP5/ABCA3/NOS3/STRA6/FBXW7/NOG/PGR/FGF7/ITGA3/PROX1/ASCL1/WNT2/FOXA1/KDR/TCF21/PKDCC/HECA/HSD11B1/NOD
PAWR/FOXP1/MPO/HYAL2/MAP3K5/AXL/TRPM2/EDN1/TXNIP/SOD2/HMOX1/CYP1B1/PDGFRA/STAT1/GCH1/KCNA5/IL6/KLF2/IL18RAP/FBLN5/PTK2B/AR
SPINK1/NR1H4/UCN3/PRKCB/GPER1/CELA2A/GHSR/CASR/F2RL1/ADCY8/GPLD1/TRH/HNF4A/INS/GPR68/ADRA2A/CRHBP/NR0B2/TFAP2B/BAIAP3/RP
VIP/ISL1/TCIRG1/VGF/PER2/CPLX1/RAB8B/SPINK1/NR1H4/MAFA/UCN3/PRKCB/GPER1/CELA2A/GHSR/CASR/F2RL1/ADCY8/GPLD1/TRH/HNF4A/CPE/SL
NR1H4/GPT/AVP/GPER1/GHSR/IGF1/GPLD1/NOS3/INS/PTH1R/P2RY6/NTSR1/PPARG/DYRK2/PTGS2/APOA5/APOA1/IFNG/ADM/IL4/TMSB4X/ABCG4/STA
SPINK1/NR1H4/UCN3/PRKCB/GPER1/CELA2A/GHSR/CASR/F2RL1/ADCY8/GPLD1/TRH/HNF4A/INS/GPR68/ADRA2A/CRHBP/NR0B2/TFAP2B/BAIAP3/RP
CARD18/AHSG/SERPINB5/SERPINB3/FETUB/NEIL1/ANGPTL4/SERPINB8/SERPINA11/DPEP1/SPINK2/SERPINE2/HRG/SERPINA6/SERPINA1/SERPIND1/S
ADRB2/GPER1/SLC34A1/CACNB2/ERFE/LRRC55/NOS1AP/GRIN1/IGF1/TRDN/CLIP3/NLGN3/APLNR/INS/PPP3R2/KCNQ1/KCNC1/CCL2/P2RY6/NTSR1/SL
SPINK1/SLC8A3/PDGFRB/CACNB2/CACNA1G/CASR/TRPC4/CCL3/PPP3R2/CCL2/TRPV5/STC1/TRPM2/P2RX1/CACNA1E/AKAP5/CACNA1C/CACNA1I/SL
VIP/ISL1/TCIRG1/TRPV6/VGF/PER2/CCKAR/POMC/CPLX1/RAB8B/SPINK1/NR1H4/MAFA/UCN3/MYB/GALR1/PRKCB/GLP1R/GPER1/CELA2A/GHSR/CASR
RUNX1/KLKB1/CASP8/CTSH/PLAU/PRSS3/HPN/F11/CUZD1/MASP1/SERPINE2/C1RL/LGMN/APOH/SERPINF2/VSIR/GRIN2A/NLRC4/FGA/THBS1/THBD/C
SHH/KRT19/BFSP1/NEFM/PKP1/KRT75/KRT15/KRT84/KRT34/KRT4/TCHH/KRT6A/KRT80/KRT71/GFAP/KRT35/KRT37/KRT24/KRT31/PLEC/KRT5/NEFL/KR
SLC17A8/SSTR4/PLA2G3/SLC6A1/P2RX7/AVPR1B/SYT4/IL1A/AVP/ERFE/TRH/SLC38A3/NTSR1/ACSL5/ABCB11/TNFRSF11A/PLA2R1/EDN1/ADORA2A/ABA
FGFBP1/GPLD1/NOS3/FBXW7/BMPER/ZC3H12A/ID1/FGF7/ITGA3/PROX1/VASH1/SMOC2/PTN/TACSTD2/RRAS/PLXND1/MMRN2/TEK/PPARG/HRG/EPP
RIPK3/DUOXA2/CYBA/PDGFRB/F2RL1/GRIN1/ZC3H12A/AGTR1/AKR1C3/CLEC7A/TYROBP/SOD2/F2/THBS1/SYK/CYP1B1/TGFB1/PTK2B/ITGB2/PLCG2/T
TNMD/STAB1/VASH1/SULF1/TEK/PPARG/HRG/APOH/ADGRB3/PDE3B/DCN/SEMA4A/CCR2/THBS1/GDF2/GHRL/STAT1/THBS4/KLF2/TIE1/ALOX5/FASLG
RAB8B/KCNN4/SYT4/SPINK1/IL1A/NR1H4/UCN3/MYB/GALR1/AVP/PRKCB/GPER1/CASR/SLC6A4/F2RL1/ADCY8/IGF1/GPLD1/TRH/INS/GPR68/PARD6A/C
TNMD/STAB1/VASH1/SULF1/TEK/PPARG/HRG/APOH/ADGRB3/PDE3B/DCN/SEMA4A/CCR2/THBS1/GDF2/GHRL/STAT1/THBS4/KLF2/TIE1/ALOX5/FASLG
AGER/IL1A/TNIK/RASSF2/EPHB1/F2RL1/TPD52L1/FZD10/EDAR/MECOM/TLR3/SERPINB3/CYLD/GADD45G/MAP3K5/SERPINF2/NRK/MARVELD3/FCGR2
PIGR/GNAT1/CALHM1/TAS2R7/GNAT3/SCNN1B/TAS1R1/RTP1/LPO/RTP4/SCNN1G/CALHM3/P2RX2/TAS2R38/RGS21/LCN1/TAS1R2/RTP5/TAS1R3/PLC
AVP/ADRB2/GPER1/C2CD4A/KCNMA1/PLEC/MYLK3/ADRA1B/CASR/SLC6A4/F2RL1/NOS3/SLC38A3/SLC16A2/KCNJ8/SCPEP1/INS/ADRA2A/SLC29A2/SLC
BCL2L10/MEFV/NKX3-1/PPARG/LGMN/MAP3K5/CLEC7A/NODAL/GRIN2A/CARD9/NLRC4/CASP10/SYK/NLRP2/LGALS9/NLRP3/FASLG/JAK2/CYFIP2/NLR
MTNR1B/ADRA2B/EDN1/FCGR2B/FFAR4/CCR2/IL11/IL12B/GHRL/ABAT/TNFRSF1B/LGALS9/SFRP1/BMP8A/NR1H3/UCP2/CD300A/RSAD2/SPI1/LILRB1/V
SKAP1/PLEKHA2/RRAS/TEK/HRG/KDR/NRP1/CCL25/CSF1/PTK2B/SFRP1/THY1/CCL21/EPHA1/CD3E/CCR7/MYOC
FGFBP1/GPLD1/NOS3/FBXW7/BMPER/ZC3H12A/ID1/FGF7/ITGA3/PROX1/VASH1/SMOC2/PTN/TACSTD2/RRAS/PLXND1/MMRN2/TEK/PPARG/HRG/EPP
SPINK1/NR1H4/UCN3/PRKCB/GPER1/CELA2A/GHSR/CASR/F2RL1/ADCY8/GPLD1/TRH/HNF4A/INS/GPR68/ADRA2A/CRHBP/NR0B2/TFAP2B/BAIAP3/RP
PLA2G3/EXPH5/CD33/RGCC/SLC30A8/GRK2/SLC6A1/P2RX7/VIP/ISL1/AVPR1B/IL13/RAB8B/KCNN4/SYT4/SPINK1/CYBA/IL1A/NR1H4/UCN3/MYB/GALR1/A
PAK1/LAT/NRG1/ELANE/NTRK3/TPD52L1/ADCY8/IGF1/FBXW7/INS/ADRA2A/TLR3/PROX1/CHRNA3/TNFRSF10A/HMGA2/NTRK2/TEK/AGTR1/KDR/LCP2/
BMP10/ADAMTS9/NGFR/TBXA2R/ACVRL1/CLDN3/BCR/RGCC/CX3CL1/SMAD7/DACH1/MEOX2/CNN2/CX3CR1/ADTRP/RIPOR2/ANGPT4/PTPRT/HAS1/L
BATF/CLCF1/IL4/IL10/IL2/IL27RA/TGFB1/AICDA/CD28/FOXP3/CD40LG/TBX21/PTPRC/CCR6
LPAR2/ADCYAP1/PLN/ABCA1/GCGR/AKAP13/GPR6/GHRH/SSTR4/ADGRE2/APLP1/DRD5/DRD1/PTGER3/GPR62/TBXA2R/GPR101/ADCY1/ADRB1/VIP/FS
ARRB2/CACNB4/RAB31/RIT2/FCGR2A/INPP5F/USP6/TLR2/CFP/BAIAP3/PKDCC/RPH3AL/SYNE1/AXL/PLA2G4E/P2RX1/APOA5/C2/APOA1/CLEC7A/SNAP9
FGFBP1/GPLD1/NOS3/FBXW7/BMPER/ZC3H12A/ID1/FGF7/ITGA3/PROX1/VASH1/SMOC2/ACTC1/PTN/TACSTD2/RRAS/PLXND1/MMRN2/TEK/PPARG/H
LMOD2/TNNT1/KRT19/MYOM1/MYH7/MYH3/SYNPO2L/TCAP/AKAP13/MYOM3/BMP10/MYPN/KRT8/SIX4/PLEC/MYLK3/CSRP3/PROX1/MYBPC3/TNNT3/M
FGFBP1/NOS3/KDR/PTGS2/NRP1/TMSB4X/HMOX1/THBS1/VEGFC/PLG/TGFB1/HDAC9/GATA2/FOXC2/CD40/JCAD/FGF2
KCNE4/PML/CX3CL1/P2RX2/P2RX7/TCIRG1/STRIT1/IL13/DHRS7C/KCNN4/BPIFA1/CLIC2/GALR2/CHP2/GRIN2B/CYBA/GPER1/PTPN3/CACNB2/SCN5A/L
IRGM/TREM2/GBP5/PTPN22/PLCG2/BTK/TLR6/TLR4/NLRC3
HTRA2/AGER/S100A9/F3/GRIN2B/GPER1/RFPL1/STAT3/BCL2L10/MEFV/SERPINB3/NKX3-1/PPARG/LGMN/MAP3K5/CLEC7A/VSIR/NODAL/GRIN2A/CARD

RFFL/HTRA2/S100A9/F3/GRIN2B/AVP/GPER1/CSN2/RFPL1/CARD18/BCL2L10/MEFV/NKX3-1/DPEP1/PPARG/LGMN/PSMB9/MAP3K5/CST4/CLEC7A/NODA

P2RX7/ISL1/TCIRG1/PER2/IL13/ADTRP/KCNN4/SYT4/IL1A/NR1H4/UCN3/PRKCB/GPER1/CELA2A/GHSR/CASR/F2RL1/ADCY8/IGF1/GPLD1/TRH/HNF4A/IN

BCR/RARRES2/CX3CL1/ROR2/CNN2/CX3CR1/MDK/CCL3/CCL2/CSF1/C3AR1/CCR2/P2RY12/THBS1/CSF1R/CD200R1/TREM2/CMKLR1/PTK2B/IL34/SLAN

PAK1/LMO3/CYP27B1/F2RL1/COLEC12/UNC93B1/STAT3/VDR/HNF4A/KDM4C/RHOXF1/KCNJ8/ALOXE3/TLR3/PGR/MEFV/NR3C2/NKX3-1/CYP24A1/RORC

APOBEC3D/CYP2C8/APOBEC3G/A1CF/APOBEC3H/EOMES/AICDA/CYP2C9/TOX/GATA3/APOBEC3A

MEFV/LBP/SLC46A2/DHX58/ARRB2/HSPA1B/TLR1/IRGM/NOD2/DDX60/TREM2/SLC15A3/GBP5/NR1H3/CD300A/IRF7/LYN/RSAD2/SLC15A2/TLR9/CD300LF

APOB/PLA2G2E/APOC1/PLA2G3/APOC3/NR1H4/PCSK6/AGTR1/MPO/APOA5/APOA1/LIPC/PLA2G5/PCSK5/ABCG1/LIPG/PLA2G7/CETP/PLA2G2A

PGLYRP4/CHI3L1/OVGP1/PGLYRP2/HYAL2/STAB2/LYVE1/FUCA1/TGFB1/CHI3L2/FGF2/CEMIP

PML/CX3CL1/P2RX2/P2RX7/STRIT1/IL13/DHRS7C/CLIC2/CYBA/GPER1/CACNB2/NOS1AP/GRIN1/LIME1/TRDN/GPR35/APLNR/PPP3R2/ADRA2A/P2RY6/N

BMP10/ADAMTS9/NGFR/TBXA2R/ACVRL1/CLDN3/BCR/RGCC/RGN/CX3CL1/SMAD7/DACH1/MEOX2/CNN2/CX3CR1/ADTRP/RIPOR2/ANGPT4/PTPRT/H

TBX2/HTR1A/CAMK2D/BMP10/APELA/KCNE3/CORIN/TBXA2R/KCNE4/GRK2/SMAD7/ADRB1/KCND2/ISL1/AVPR1B/PER2/KCNMB4/CLIC2/GJD3/AGER/SC

SDR16C5/CGA/EGR1/HPN/UGT2B11/AKR1B10/SULT1E1/FOXE1/DUOXA2/ATP6AP2/ADH7/DHRS3/NR5A1/UGT2A3/BCO2/HSD17B2/DIO2/CYP26C1/DGAT2

CYBA/NOXO1/F2RL1/DUOX2/NCF2/CLEC7A/NOX3/EDN1/TYROBP/SOD2/SYK/NCF4/TGFB1/CYBB/ITGB2/HVCN1

HTR4/MUC6/MUC4/IL20RB/ASCL3/PTH1R/GPR55/ARMS2/NXNL2/SLC28A2/SPP1/TNFRSF11A/VSTM4/LACRT/PRICKLE1/PTGS2/LDB2/USH1C/CSF1/IL7/C

HTR4/MUC6/MUC4/IL20RB/ASCL3/PTH1R/GPR55/ARMS2/NXNL2/SLC28A2/SPP1/TNFRSF11A/VSTM4/LACRT/PRICKLE1/PTGS2/LDB2/USH1C/CSF1/IL7/C

BATF/CLCF1/IL4/IL10/IL2/IL27RA/TGFB1/AICDA/SAMHD1/CD28/FOXP3/CD40LG/TBX21/PTPRC/CCR6

TACR3/CYP4A11/UCN/CALCA/SLC4A5/OXTR/AR/LEP/DRD3/CYP11B1/HSD11B2/GAS6/MC3R/SMAD3/UTS2B/PTGS1/SPX/NDST2/CARTPT/CYP4F2/SCNN

SLC39A5/GRM1/CACNA1S/SLC7A11/SCNN1G/SLC47A1/SLC30A8/SLC6A1/KCNJ1/SLC9A2/TRPV6/PER2/GFAP/SLC9A4/CNGA3/SLC8A3/SLC9A3/SLC34A1/S

APOC3/SOAT2/NR1H4/RORA/DGAT2/ABCG5/MINAR2/HNF4A/ACSM1/TTC39B/ACSM3/LDLR/MALL/MALRD1/ABCB11/APOA5/APOA1/ABCG4/LIPC/FABP4/

PNP/GDA/APOBEC2/UPP2/APOBEC3F/APOBEC1/ENPP4/CDA/APOBEC3D/APOBEC3G/FUCA1/APOBEC3H/DPYD/AICDA/ADA2/APOBEC3A

NFKBIZ/EPB42/CD248/PBX1/NKX2-5/SLC46A2/CACNB4/TCF21/MAFB/TNFRSF11A/FOXL1/LIPA/NFKB2/PDPN/TGFB1/CDH17/IL15/BCL3/SPNS2/LTA/TOX/G

TTBK1/IFNGR1/DRD1/GRN/ROR2/GFAP/AGER/S100A9/TSPAN2/LDLR/IFNG/ADORA2A/C1QA/DLL1/TREM2/IL6/IL1B/TNF/TLR4

P2RY6/GPR55/ANGPTL4/PLCB2/AGTR1/PNLIP/APOH/APOA5/PLA2R1/FGFR3/ADRA1A/PDGFRA/PLA2G5/FGFR2/TXK/CD86/NR1H3/S1PR4/ITK/SELE/CCL5

MTNR1B/PLA2R1/ADRA2B/EDN1/FCGR2B/FFAR4/CCR2/IL11/IL12B/CD74/GHRL/SCT/ABAT/TNFRSF1B/LGALS9/SFRP1/BMP8A/NR1H3/UCP2/CD300A/RS

GJB2/GHSR/OVGP1/CYP27B1/IGFBP5/PRDM1/SLC38A3/VDR/CLIC5/ADRA2A/EPYC/PGR/ITGA3/TAC3/PTN/CRHBP/LNPEP/SPP1/STC1/DSG2/PZP/PTGS2

PLAU/ADAMTS18/PROC/ADTRP/F11/KRT1/NOS3/SH2B3/C1QTNF1/SERPINE2/HRG/APOH/SERPINF2/EDN1/F2/FGA/THBS1/THBD/PLG/PDGFRA/SERPIN

F2RL1/IGF1/GPR35/GPR55/ABRA/TEK/ARHGEF3/APOA1/CSF1/TGM2/GPR65/RASGRP1/RASGEF1A/SEMA4D/P2RY8/GPR18/PIK3CG/RELN/GPR174/RASC

PLCL1/NPY5R/CA2/SLC6A1/CNTNAP4/CA7/BAIAP3/NALCN/NPS/ADORA2A/ADRA1A/PLCL2/TACR1/SLC38A1/CNR2

MYB/L3MBTL3/GP9/EPB42/FBXW7/GPR68/ZNF385A/TSPAN2/SH2B3/THPO/TLR2/FOXP1/SLC4A1/TYROBP/RHEX/TAL1/FLI1/KLF2/CLDN18/LILRB1/EVI2B/L

TLR2/DSG2/TXNIP/THBS1/PTGER2/CYP1B1/TRERF1/TGFB1/FOSB/NR1H3/TGFB3/TACR1/CD38

ISL1/TCIRG1/PER2/CCKAR/POMC/RAB8B/SPINK1/NR1H4/UCN3/MYB/GALR1/PRKCB/GPER1/CELA2A/GHSR/CASR/F2RL1/ADCY8/GPLD1/TRH/HNF4A/IN

MARVELD2/NRG1/F2RL1/RAPGEF3/TNMD/PROX1/BTG1/KDR/STC1/SOX18/NRP1/ATOH8/CDH5/LIPA/GDF2/HEG1/PDPN/DLL1/PECAM1/TIE1/HPSE/TJP3

PML/CACNA2D4/GP1BB/CX3CL1/P2RX2/P2RX7/TRPV6/STRIT1/IL13/DHRS7C/CLIC2/CHRNA7/GRIN2B/CYBA/SLC8A3/GPER1/CACNB2/GP9/CACNA1G/AN

KCNJ8/MEFV/IRGM/TREM2/GBP5/NLRP3/PTPN22/PLCG2/BTK/TLR6/TLR4/NLRC3

KCNQ1/SLC6A18/HYAL2/AKR1C3/EDN1/AQP3/KLHL3/SLC12A3/SLC15A2/HBB/ADIPOQ

PLB1/BCO1/HMGCS2/CYP1A2/ABCA4/RBP2/ADH4/CYP26A1/ALDH8A1/RDH10/SDR9C7/RBP3/RARRES2/SDR16C5/AKR1B10/ADH7/DHRS3/BCO2/HTRA2/

HAS3/HYAL2/ITIH6/STAB2/LYVE1/TGFB1/SMPD3/IL1B/FGF2/CEMIP

CHP2/SYT4/SYT13/SCN5A/LCE1D/ADCY8/GPLD1/SYT9/RASA4B/CRHBP/DPEP1/RASAL1/CPNE6/LGMN/KCNQ3/AKR1C3/TRPM2/EDN1/RYR3/SYT8/CPNE

TACR3/OXTR/HSD11B2/SPX/CARTPT/ID2/OR51E2/DRD5/TBXA2R/ADRB1/AVP/GLP1R/ADRA1B/TAC3/ADRA2B/ADRA1A/TACR1

CASP8/STK4/KRT8/PRDM1/GCM1/HAND1/PLG/EOMES/SPINT2/ST14/LIF

BATF/CLCF1/IL4/IL10/IL2/IL27RA/TGFB1/AICDA/CD28/FOXP3/CD40LG/TBX21/PTPRC/BCL11B/CCR6

BATF/CLCF1/IL4/IL10/IL2/IL27RA/TGFB1/AICDA/CD28/FOXP3/CD40LG/TBX21/PTPRC/BCL11B/CCR6

PLN/XCL1/JPH2/HRC/CAMK2D/DRD1/CX3CL1/IL13/DHRS7C/CLIC2/CYBA/GPER1/LIME1/TRDN/APLNR/P2RY6/NTSR1/LACRT/JPH3/F2/CASQ1/PTK2B/CA

PLAU/ADAMTS18/PROC/ADTRP/F11/KRT1/NOS3/SH2B3/C1QTNF1/SERPINE2/HRG/APOH/SERPINF2/EDN1/F2/FGA/THBS1/THBD/PLG/PDGFRA/SERPIN

PLB1/BCO1/CYP1A2/ABCA4/RBP2/ADH4/CYP26A1/ALDH8A1/RDH10/SDR9C7/RBP3/RARRES2/SDR16C5/AKR1B10/ADH7/DHRS3/BCO2/CYP26C1/DGAT2/

PLCB2/INPP5F/PIK3C2G/PIK3C2B/PLCG2/PLCH2/PIK3CG

KRT6A/FGF10/NLRP6/DRD5/SLC7A11/GATA1/SPRR3/MCAM/TEC/TBXA2R/ACVRL1/ANXA8/CLDN3/GRHL3/DCBLD2/MYOF/ANXA1/PLAU/CCN1/LRG1/GP

TRPV6/SLC8A3/CACNA1G/PPP3R2/TRPV5/TRPM2/P2RX1/CACNA1E/AKAP5/CACNA1C/CACNA1I/SLC8A1/MS4A1

TRPV6/SLC8A3/CACNA1G/PPP3R2/TRPV5/TRPM2/P2RX1/CACNA1E/AKAP5/CACNA1C/CACNA1I/SLC8A1/MS4A1

SPP1/TNFRSF11A/TF/SYK/ADAM8/TNFSF11/DCSTAMP

MBL2/COLEC11/CRP/COLEC10/PTX3/SFTPA1/LBP/C4B/C4BPA/CFP/C4BPB/PLA2G5

BATF/CLCF1/IL4/IL10/IL2/IL27RA/TGFB1/AICDA/SAMHD1/CD28/FOXP3/CD40LG/TBX21/PTPRC/BCL11B/CCR6

ATP6V1B1/ALOXE3/CPA1/DPEP1/TLR2/CYP4F12/AKR1C3/GSTA1/PTGS2/CYP4A22/EDN1/CYP1A1/AKR1C2/NLRC4/CYP2C8/CYP4F22/SYK/CYP1B1/CD74/

PLA2G2E/SLC16A11/SLC10A2/PLA2G3/SLC16A14/SLC22A7/P2RX7/AVPR1B/SLC6A11/SLC16A13/SLC10A1/PLA2G4F/IL1A/ERFE/CASR/SLCO1A2/SLC16A2/K

IFI35/TICAM2/F2RL1/LBP/NR1H3/PTPN22/LTF

DUOX2/HP/MPO/PXDNL/HBZ/HBD/HBG1/HBB/HBA2

TBXA2R/ACVRL1/RGCC/ANXA1/PRKD2/MEOX2/ADTRP/ANGPT4/SH3BP1/FGFBP1/GPLD1/NOS3/FBXW7/ID1/VASH1/MMRN2/PPARG/HRG/KDR/SOX18/

ADCY1/ITPKA/SLC6A1/P2RX7/TRPV6/KCNMB4/CHP2/SYT4/SYT13/KCNMA1/CASR/SCN5A/LCE1D/ADCY8/GPLD1/SYT9/CACNG2/RASA4B/CRHBP/DPEP

LDLRAD4/GHSR/SRGAP3/NKX2-1/ELANE/IGFBP5/SAP30L/NRG3/STAT3/NOG/TNN/ADORA3/FIGNL2/VASH1/CCL2/TACSTD2/RRAS/SULF1/MMRN2/DPE

ADAMTS18/GP6/CELA2A/SH2B3/C1QTNF1/SERPINE2/SYK/PDPN/IL6/JAK2/LYN/PRKCQ/TNFSF11/ANK3/MMRN1/CCL5

EFEMP2/ZFPM2/TP73/CFLAR/MYB/GPER1/PDGFRB/CAV2/ELANE/EPHB1/IGFBP5/IGF1/NOS3/NOG/FGF20/NKX2-5/PTGIR/P2RY6/WNT2/TENM4/PPARG/

CSF1/C3AR1/THBS1/CSF1R/CMKLR1/IL34/CCL5

STAT3/ID1/LCN2/PRDM16/YBX2/CIDEA/NTSR1/LNPEP/TSHR/WNT10B/GADD45G/TNFRSF11A/TRPM2/ESRRG/PTGS2/NOX3/IL4/FFAR4/KSR2/CCR2/FA

RAET1G/ERAP1/B2M/UNC93B1/ERAP2/LNPEP/CD209/TAPBPL/LGMN/FCGR2B/FCGR1A/CLEC4A/CD74/TREM2/FCER1G/MPEG1/CTSS/RAET1L/IFI30

CA2/HBQ1/MB/RHCG/HBZ/HBD/IPCEF1/HBG1/HBB/HBA2

VIP/ISL1/TCIRG1/TRPV6/VGF/PER2/CCKAR/POMC/CPLX1/RAB8B/SPINK1/NR1H4/MAFA/UCN3/MYB/GALR1/PRKCB/GLP1R/GPER1/CELA2A/GHSR/CAS

ZC3H12A/RESF1/CXCL8/PROX1/CCL2/HMGA2/CD209/IFNB1/ATG16L2/OAS3/APOBEC3D/APOBEC3G/OAS1/APOBEC3H/IFITM1/ISG15/IFIT5/OAS2/CCL8/C

EXPH5/SFN/PLEC/KDF1/PALLD/KRT2/BCL11B

KCNN4/BPIFA1/CLIC2/GALR2/CHP2/SPINK1/CYBA/ADRB2/GPER1/B2M/PDGFRB/PTPN3/CACNB2/CNTN1/MYLK/CASR/SCN5A/LRRC55/FXYD4/NOS1AP/

SULF1/SERPINE2/NRP1/EDN1/NPTX1/ITGA4/VCAM1/LRIG1/SULF2/SEMA3A

CD33/PML/CX3CL1/CX3CR1/GHSR/IGF1/CARD18/ZC3H12A/MEFV/ELF4/ARRB2/APOA1/FFAR4/NLRP7/GHRL/TREM2/NLRP3/FFAR1/LILRB4

PTX3/TRIM21/TRIM10/TRIM38/RFPL1/LRRC15/TRIM34/CH25H/TRIM59/CD74/IFITM1/CD4/LGALS9/TRIM22/HS3ST5/CIITA

SFRP1/CCND2/GATA3

HAAO/SRD5A2/ACSM1/TDO2/CYP1B1/KYNU/CYP2E1/KMO/IDO2/IDO1

ASCL1/INSM1/TFAP2B/NRP1/PLXNA4/TP63/GFRA3/GATA3/SEMA3A

MC5R/OR10J5/AGT/GALR3/HTR2C/HRH2/LHCGR/SSTR3/CHRM5/CHRM1/MC3R/DRD4/SSTR4/HTR1A/SSTR5/DRD1/ANXA1/SSTR2/CHRM2/HTR6/HT

PLA2G4F/PLA2G4B/PLPP6/ABHD16B/LDLR/INPP5F/PNPLA7/PLA2G4E/LIPC/PLA2G5/PLBD1/SMPD3/LIPG/PLA2G4C/PLA2G7/PLCG2

IRGM/TREM2/GBP5/PTPN22/PLCG2/BTK/TLR6/TLR4/NLRC3

TTBK1/IFNGR1/GRN/AGER/LDLR/IFNG/ADORA2A/C1QA/TREM2/IL6/IL1B/TNF

PODXL/EDN1/NPHS1/PTPRO/FOXC2/BASP1/CD24/ADIPOQ

PODXL/EDN1/NPHS1/PTPRO/FOXC2/BASP1/CD24/ADIPOQ

SLC17A8/SLC6A1/PER2/SLC6A11/SLC6A12/SLC6A13/KCNJ10

IRGM/TREM2/GBP5/NLRP3/NLRP1/PTPN22/PLCG2/BTK/TLR6/TLR4/NLRC3

GPR68/PPARG/FOXP1/CSF1/CSF1R/PDE1B/CD74/CD4/IL31RA/IRF7/IL34/INPP5D/DCSTAMP

TLR3/IFNB1/IRGM/OAS1/STAT1/MNDA/IRF1/PYHIN1/AIM2

TRIM8/ARG1/ITGAV/TRIM6/KRT6A/PTX3/TRIM21/TRIM10/TRIM38/BPIFA1/RFPL1/F2RL1/LRRC15/TRIM34/CXCL8/CH25H/TRIM59/CD74/IFITM1/CD4/LGAL

PIGR/GNAT1/TAS2R7/GNAT3/TAS1R1/RTP1/LPO/RTP4/TAS2R38/TAS1R2/RTP5/TAS1R3/PLCB2/RTP3/CST4/FFAR4/PKD1L3/RTP2/PKD2L1

SLC7A11/ANXA1/SH2B3/AXL/CSF1/CCR2/IL6/MERTK/GCNT4/BTK

CCL2/ANGPTL4/TEK/KDR/AKR1C3/ZFP36/PLA2R1/ITGA4/IL4/CDH5/HMOX1/FGA/THBS1/IL6/TGFB1/FGB/FASLG/GATA2/JAK2/CD40/ECSCR/BTC/ICAM1/C

PLK5/KLF4/SPI1/ABI3/PRF1/LAPTM5

CYLD/SKIL/HYAL2/MAL/THBS1/SFRP1/INHBA/TNF/TNFSF10/TLR6/PTPRC/TLR4/LTB

ITGA3/PTN/TSPAN32/FERMT3/ITGAD/FYB2/ITGA8/APOA1/NRP1/ITGA4/PLPP3/FYB1/SYK/ISG15/HCK/CDH17/PRAM1/PTK2B/ITGAX/ITGA5/THY1/ITGBL1/IT

SYK/HCK/LYN/MYO1G/FGR/VAV1

SYK/HCK/LYN/MYO1G/FGR/VAV1

DTX3L/APOBEC3F/CGAS/ZC3H12A/PARP9/APOBEC3G/IL27/IL12B/STAT1/TRIM22/LILRB1/IL12RB1/IL23A/AIM2

CARD9/SYK/IL12B/CD80/TLR9/IL1B/CD84/RASGRP1/IL23A/IL18

CARD9/SYK/IL12B/CD80/TLR9/IL1B/CD84/RASGRP1/IL23A/IL18

PNP/GDA/APOBEC2/UPP2/APOBEC3F/APOBEC1/ENPP4/CDA/APOBEC3D/APOBEC3G/APOBEC3H/DPYD/AICDA/ADA2/APOBEC3A

PARP9/IRGM/APOBEC3G/IL27/IL12B/STAT1/TRIM22/IL15/MMP12/LILRB1/FOXP3/IL1B/IL12RB1/SPN/IL23A/AIM2

BMP10/MYPN/KRT8/SIX4/CFLAR/PDGFRB/PLEC/MYLK3/NRAP/NKX2-5/CSRP3/MYH11/PROX1/MYBPC3/MYL9/ACTC1/TNNT3/MYOZ1/ANKRD1/FLNC/ED

RIPOR2/MDK/PECAM1/FUT7/IL1R1/ADAM8/JAML/PIK3CG

TICAM2/GPSM3/STAT3/IL17RA/TLR3/CLEC7A/CARD9/NOD2/IL6/GBP5/CD6/TNF/TLR6/TLR4

SLC27A6/NTSR1/FABP2/NMUR2/PPARG/FABP9/ACSL5/FABP12/TNFRSF11A/FABP6/OC90/RBP7/PTGS2/PLA2R1/EDN1/SLC27A2/SLC22A8/FABP4/THBS1/S

CCL2/SLIT2/PADI2/SLAMF8/KLRK1/GPR18/STAP1

FGFBP1/GPLD1/NOS3/ZC3H12A/FGF7/ITGA3/PROX1/SMOC2/RRAS/TEK/KDR/FOXP1/LGMN/PTGS2/NRP1/IFNG/EDN1/TMSB4X/PLPP3/ATOH8/HMOX1/T

FOXP1/ITGAX/DLL4/CXCR4/CXCL10/FGF1

FOXP1/ITGAX/DLL4/CXCR4/CXCL10/FGF1

STAT1/CITED1/BASP1/GATA3/LIF

TLR3/TNFRSF8/TLR5/TNFRSF10A/PTGS2/MAP3K14/ANKRD1/TLR8/PTGER4/TMEM150C/TNFSF14/CD40/IL1B/IRF1/PDE2A/FGF2/TLR4/IL33/TLR7

STAT3/JAK2/LYN/JAK3

STAT1/GATA3/CD24/LIF

CDH5/PECAM1/ILDR1/TJP3/CGNL1/MPP7

FGFR3/RFLNA/ALDH1A2/ZBTB16/BMP2/LTF/SEMA4D/GATA3

LHCGR/LTB4R/PLCE1/GPR4/GPR32/ESR1/GPR139/DRD3/CHRM1/MC3R/HCRTR2/GRPR/FPR2/OPRK1/NMBR/GRM1/DRD5/DRD1/PTGER3/FPR1/CCKE

THSD4/EFEMP2/MYH11/FBLN5/MFAP4/TNXB

NMNAT2/ASPDH/ACMSD/HAAO/NMNAT3/PARP9/NNMT/KYNU/KMO/IDO2/IDO1

PPARG/NFKBIA/NR1H3/ABCG1/CETP/ADIPOQ

AXL/CARD9/SYK/NOD2/CD74/TGFB1/WNT5A/TGFB3/LILRB1/SEMA7A/PLCG2/LAPTM5/TLR4/TLR7

GHSR/CDKN2A/ST6GAL1/NOD2/IRF7/CCR5/CCL5

FGFBP1/GPLD1/NOS3/FBXW7/BMPER/ZC3H12A/FGF7/ITGA3/PROX1/VASH1/SMOC2/PTN/TACSTD2/RRAS/MMRN2/TEK/PPARG/HRG/EPPK1/KDR/FOXF

ADAD1/APOBEC2/APOBEC3F/APOBEC1/APOBEC3D/APOBEC3G/RBM47/A1CF/APOBEC3H/AICDA/ADARB2/APOBEC3A

PODXL/EDN1/NPHS1/PTPRO/FOXC2/BASP1/CD24/ADIPOQ

PLB1/BCO1/CYP1A2/ABCA4/RBP2/ADH4/CYP26A1/ALDH8A1/RDH10/SDR9C7/RBP3/RARRES2/SDR16C5/AKR1B10/ADH7/DHRS3/BCO2/CYP26C1/DGAT2/

LTB4R/SSTR3/UCN/CALCA/PYY/MCHR1/GPR139/SORL1/ECEL1/PRLHR/NPB/CARTPT/HCRTR2/PTH2/GRPR/OPRK1/ADCYAP1/NMBR/KISS1R/SSTR4/S

FGF19/NR1H4/PROX1/MALRD1/STAR/STARD4/CES1

CYP26A1/ALDH8A1/RDH10/ADH7/BCO2/CYP26C1/ADH1A/SCPEP1/ADH1B/UGT1A3/AKR1C3/UGT1A8/DHRS9/ADH1C/CYP1A1/CYP2C8/ALDH1A2/CYP2S1

CPA1/DPEP1/TLR2/CYP4F12/SYK/PLA2G5/ALOX5/DPEP2/GGT5/ALOX5AP

BMP10/MYPN/KRT8/LARGE1/SIX4/CFLAR/PDGFRB/PLEC/MYLK3/NRAP/NKX2-5/CSRP3/MYH11/PROX1/MYBPC3/MYL9/ACTC1/TNNT3/MYOZ1/ANKRD1/I

PLAU/ADAMTS18/PROC/ADTRP/F11/KRT1/NOS3/SH2B3/C1QTNF1/SERPINE2/HRG/APOH/SERPINF2/EDN1/F2/FGA/THBS1/THBD/PLG/PDGFRA/HS3ST5

APOA1/IL10/KLF4/IL1RN/ADIPOQ

APOA1/IL10/KLF4/IL1RN/ADIPOQ

GHRH/GSX1/SLC6A3/POU1F1/GATA2/BMP2/FGF2

SPINK1/NR1H4/UCN3/MYB/GALR1/PRKCB/GPER1/CASR/ADCY8/GPLD1/TRH/INS/GPR68/C1QTNF1/NKX3-1/NR0B2/PPARG/SPP1/BAIAP3/RPH3AL/ITPR1

RAB8B/SPINK1/NR1H4/UCN3/PRKCB/GPER1/CASR/ADCY8/GPLD1/TRH/INS/GPR68/NR0B2/BAIAP3/RPH3AL/ITPR1/GIPR/ANO1/FFAR4/F2/FGA/GHRL/TU

CX3CL1/CX3CR1/CCL3/RRAS/BMERB1/CSF1/CCR2/P2RY12/TREM2/GPR183/STAP1

RAB8B/SPINK1/NR1H4/UCN3/PRKCB/GPER1/CASR/ADCY8/GPLD1/TRH/INS/GPR68/NR0B2/BAIAP3/RPH3AL/ITPR1/GIPR/ANO1/FFAR4/F2/FGA/GHRL/TU

HTRA2/CHP2/RASGRF2/AGER/S100A9/F3/PRSS22/GRIN2B/GPER1/PDGFRB/RFPL1/CAV2/RGS16/PREX2/SH3BP1/CCL1/PPP1R14D/F2RL1/NTRK3/GPLD

SHH/IHH/LOXL3/BCL6/RUNX1/SOCS5/ANXA1/SMAD7/ZC3H12A/ASCL2/IL2/IL4R/FOXP3/SOCS1/TBX21/JAK3

IL4/IL2/CD28/TBX21/PTPRC

APOBEC2/UPP2/APOBEC3F/APOBEC1/CDA/APOBEC3D/APOBEC3G/APOBEC3H/AICDA/APOBEC3A

APOBEC2/UPP2/APOBEC3F/APOBEC1/CDA/APOBEC3D/APOBEC3G/APOBEC3H/AICDA/APOBEC3A

NKX2-1/IGFBP5/STRA6/PGR/FOXA1/KDR/TCF21/PKDCC/FGFR2/CLDN18/HOPX/TGFB3/SMPD3/PTGES3/MMP12/FLT4/LIF

CX3CL1/P2RX2/P2RX7/IL13/KCNN4/GALR2/CHP2/GPER1/CACNB2/LRRC55/NOS1AP/GRIN1/TRDN/APLNR/PPP3R2/KCNQ1/KCNC1/P2RY6/NTSR1/KCNC

B4GALT1/CRISP1/IQCF1/PLB1/PLA2G10/PRSS37/SPINK1/GLRA1/ZP4/FAM170B/SPINK13/HVCN1

APOBEC3D/APOBEC3G/A1CF/APOBEC3H/EOMES/AICDA/TOX/GATA3/APOBEC3A

NFKBIE/ADCY8/STAT3/CDKN2A/ZC3H12A/ID1/INS/NEUROG1/FOXS1/PYDC1/TRIM34/ADORA3/PROX1/ABRA/ARRB2/WNT2/CYLD/NR0B2/PPARG/FOXA

BMP10/ADAMTS9/TBXA2R/ACVRL1/RGCC/MEOX2/ANGPT4/VASH1/MMRN2/PPARG/HRG/STC1/APOH/DCN/SLIT2/THBS1/GDF2/TGFB1/DLL4/KLF4/FGF

FGL2/PCK1/BCL6/IL23R/IFNL1/IL12B/IL12RB1/IL23A

CHRNA7/GRIN2B/SLC8A3/CTSC/AVP/ADRB2/NRXN1/GHSR/GRIN1/NLGN4X/NOS3/CCL3/GPR35/NLGN3/EGR2/FGF12/GLRA1/TNR/DLGAP3/BEGAIN/TYM

APOBEC3D/APOBEC3G/A1CF/APOBEC3H/EOMES/AICDA/TOX/GATA3/APOBEC3A

FGL2/PCK1/BCL6/IL23R/ST3GAL1/IFNL1/IL12B/IL12RB1/IL23A

VASH1/TACSTD2/MMRN2/PPARG/HRG/EPPK1/STC1/APOH/MARVELD3/IL4/DCN/SLIT2/THBS1/GDF2/EVL/TGFB1/DLL4/PTPRR/KLF4/FGF2/TNF/CXCL13

FCGR2B/CCR2/LGALS9/CD300A/SPI1/CD84/NCKAP1L

CD33/RIPK3/ITGA1/CHP2/PDGFRB/PPP1R14D/GPLD1/IFNG/CD300A/BMP2/PLEK/PTPRC/FCRL3

SLC39A5/CACNA1S/SCNN1G/SLC30A8/SLC6A1/KCNJ1/SLC9A2/TRPV6/SLC9A4/CNGA3/SLC8A3/SLC9A3/SLC34A1/SLC12A1/SLC12A6/ATP1A3/CACNA1G/

SLC39A5/CACNA1S/SCNN1G/SLC30A8/SLC6A1/KCNJ1/SLC9A2/TRPV6/SLC9A4/CNGA3/SLC8A3/SLC9A3/SLC34A1/SLC12A1/SLC12A6/ATP1A3/CACNA1G/

KLF13/HSPA1B/MAFB/ZFP36/RHEX/TAL1/STAT1/ISG15/GATA2/LYN/SPI1/INHBA/NCKAP1L/INPP5D

DHX58/CYLD/SIGLEC1/NMI/RELB/REL/IRGM/TYROBP/IL10/HAVCR2/LILRB1/XAF1/LILRA4/NLRC3

PNP/GNMT/APOBEC2/UPP2/APOBEC3F/PRTFDC1/NT5C1B/APOBEC1/NT5E/ENPP4/CDA/APOBEC3D/APOBEC3G/APOBEC3H/AICDA/PTGDR/ADA2/APO

PARP9/APOBEC3G/IL27/IL12B/STAT1/TRIM22/IL15/MMP12/LILRB1/IL1B/IL12RB1/IL23A/AIM2

IL12B/CD226/TGFB1/HAVCR2/CRTAM/IL12A

FGFBP1/GPLD1/NOS3/FBXW7/BMPER/ZC3H12A/ID1/PROX1/VASH1/SMOC2/PTN/RRAS/PLXND1/MMRN2/TEK/PPARG/HRG/KDR/FOXP1/LGMN/STC1/V

AHR/CFTR/AANAT/ZFP36L1/HCN4/FBP1/ITPR2/PCK1/WT1/GATA1/PENK/SLC26A3/IGFBP5/RAPGEF3/KCNQ1/CRHBP/WNT10B/STC1/SLC5A5/CYP1B1/P

GP6/CELA2A/PDIA2/SLC6A4/SH2B3/MYL9/C1QTNF1/TSPAN32/FERMT3/SERPINE2/P2RY12/FGA/SYK/PDPN/PDGFRA/IL6/FGB/JAK2/PEAR1/LYN/PRKCQ/

TAP1/RAET1E/MICB/CD1B/RAET1G/ERAP1/CD1E/B2M/ERAP2/CD1C/CD74/RAET1L/CD1D

AQP9/GATA6/PFKFB1/AHR/CFTR/AANAT/ZFP36L1/HCN4/HMGCS2/FBP1/ITPR2/PCK1/SREBF1/FOS/WT1/GATA1/PENK/SLC26A3/VGF/CNGA3/SLC6A3/IG

PODXL/EDN1/NPHS1/PECAM1/PTPRO/FOXC2/BASP1/CD24/ADIPOQ

CX3CL1/P2RX2/P2RX7/IL13/GPER1/CACNB2/GRIN1/TRDN/APLNR/PPP3R2/P2RY6/NTSR1/LACRT/P2RX1/STAC3/F2/CASQ1/AKAP5/THY1/CXCL9/CXCL10

TICAM2/CD274/CD300LF/TLR6/TLR4

TNIK/RASSF2/ARHGEF6/CASR/EPHB1/F2RL1/MAPK10/TPD52L1/FZD10/EDAR/MECOM/TLR3/SERPINB3/CYLD/GADD45G/MAP3K5/SERPINF2/NRK/MAR

MDK/IL1A/CELA1/ADRB2/IAPP/RASSF2/CRB1/IGFBP5/NOS3/VDR/PTH1R/PTN/GPR55/PPARG/HRG/IL21/SPP1/TNFRSF11A/AXL/CYP1A1/IL7/TF/LIPA/CCR

ITGAM/BCR/SYK/PTAFR/PRAM1/SPI1/ITGB2

IRGM/TREM2/GBP5/NLRP3/NLRP1/PTPN22/PLCG2/BTK/TLR6/TLR4/NLRC3

PPARG/PARP9/IRGM/PARP14/TXK/NR1H3/NLRC5

PPARG/PARP9/IRGM/PARP14/TXK/NR1H3/NLRC5

P2RY6/GPR55/PLCB2/AGTR1/FGFR3/ADRA1A/PDGFRA/PLA2G5/FGFR2/TXK/CD86/S1PR4/ITK/SELE/CCL5

CD226/LYN/PTPRC

GJB2/GHSR/PLA2G4B/SLC6A4/OVGP1/CYP27B1/IGFBP5/PRDM1/SLC38A3/VDR/CLIC5/PPP3R2/ADRA2A/EPYC/PGR/ITGA3/TAC3/PTN/CRHBP/LNPEP/SE

PRDM1/HAND1/PLG/LIF

SMAD7/ZC3H12A/ASCL2/IL2/FOXP3/TBX21

ADGRB3/NPHS1/WNT1/JAM2/IL4R/TNFSF14/DOCK2/NFATC2/CD53/CXCL9/CXCL10

SP100/STK4/BCL2L1/ATF3/RFFL/CFLAR/ZSWIM2/NOS3/SKIL/FAIM2/MAL/BMP5/HMOX1/FGA/THBS1/SFRP1/FGB/ICAM1/HGF/TMC8

TRDN/APLNR/P2RY6/NTSR1/LACRT/F2/CASQ1/THY1/CXCL9/CXCL10/CEMIP/CXCL11/CD19

SMOC2/HRG/KDR/LGMN/NRP1/TMSB4X/THBS1/FGF2/FGF1/CXCL13

GHRH/MRGPRX2/ADRB1/CSF2/MTNR1B/KCNA2/ADORA2A/GRIN2A/GHRL/PTGDR/CACNA1I/PTGDS/IL18

USP18/OAS3/RBM47/IKBKE/OAS1/ISG15/WNT5A/SAMHD1/NLRC5/IRF7/MMP12/ZBP1/PTPN6

ADTRP/RIPOR2/CCL2/BMP5/SLIT2/CD200R1/IL27RA/GCSAM/PADI2/SLAMF8/KLRK1

TTC39B/PPARG/TREM2/NR1H3/ABCG1/CES1

TCF21/PODXL/MTSS1/EDN1/NPHS1/STAT1/PTPRO/FOXC2/BASP1/GATA3/CD24/MMP9/LIF/ADIPOQ

FGFR2/LRRK2/BCL11A

IL4/IL2/IL27RA/CD28/FOXP3/TBX21/PTPRC

PLA2G3/SLC22A7/P2RX7/IL1A/TNFRSF11A/PTGS2/EDN1/SLC22A8/SLCO2A1/SLCO2B1/IL1B/TNFSF11/NOS2

FGFBP1/GPLD1/FBXW7/TNN/BMPER/APLNR/MMRN2/TEK/AGTR1/KDR/VSTM4/PTGS2/NRP1/TMEM215/SLIT2/HMOX1/THBS1/IL10/VEGFC/IL12B/DLL1/G[illegible]

CEBPA/TAL1/C1QC/GATA2/CBFA2T3/LYN/EVI2B/HCLS1/INPP5D/ADIPOQ

SCN10A/PTPN3/CACNB2/CACNA1G/SCN5A/SCN2B/SCN3B/SCN4B/CACNA1C/SCN1B/SLC8A1/ANK3

CARD9/CYBB/SLC12A3/INHBA/PTPRC

TMEM106A/REL/CLEC7A/TIFA/EDN1/CARD9/SECTM1/HMOX1/CASP10/IKBKE/NOD2/CD74/IFIT5/CD4/TGFB1/INAVA/LGALS9/TRIM22/FASLG/CD40/CCL2[illegible]

SLPI/S100A9/F2RL1/CLEC7A/IFNG/SYK/PRF1/NOS2/FCER2

TAP1/RAET1E/MFSD6/MICB/RAET1G/ERAP1/B2M/ERAP2/LNPEP/TAPBPL/FCGR1A/CLEC4A/FCER1G/MPEG1/RAET1L/IFI30

CXCL12/CACNG1/PLN/XCL1/CRACR2B/GRIN3B/SLC8B1/JPH2/HRC/PLPP4/CAMK2D/DRD1/GJC2/EPB41/CD33/PML/CX3CL1/P2RX2/P2RX7/STRIT1/IL13/D[illegible]

KCNMA1/SLC12A1/SLC12A6/ATP1A3/ATP6V1B1/KCNQ1/UPK3A/SLC12A8/FXYD2/ATP1A4/KCNA5/KLHL3/ATP4A/KCNJ10/SLC12A3

CNTN2/FAT4/PCDHGA9/MYPN/PCDH1/AMIGO1/CDH4/DSG4/PCDHGA10/ESAM/CDHR2/CLSTN2/PCDHB9/HMCN2/CDH20/CDH12/CDH9/PCDHA1/PCD[illegible]

KCNE4/PML/CX3CL1/P2RX2/KCNJ1/P2RX7/TCIRG1/STRIT1/IL13/DHRS7C/SLC26A5/KCNN4/BPIFA1/CLIC2/GALR2/CHP2/SCN10A/GRIN2B/CYBA/ADRB2/[illegible]

APOB/PLA2G2E/APOC1/PLA2G3/APOC3/NR1H4/PCSK6/AGTR1/MPO/APOA5/APOA1/LIPC/PLA2G5/PCSK5/ABCG1/LIPG/PLA2G7/CETP/PLA2G2A

APOB/PLA2G2E/APOC1/PLA2G3/APOC3/NR1H4/PCSK6/AGTR1/MPO/APOA5/APOA1/LIPC/PLA2G5/PCSK5/ABCG1/LIPG/PLA2G7/CETP/PLA2G2A

CYP4F12/AKR1C3/PTGS2/CYP4A22/CYP1A1/CYP2C8/CYP1B1/CYP2E1/ALOX5/PTGES3/CYP2S1/CYP2C9/FAAH2/PLA2G4C/PTGDS/ALOX15

HYAL2/STAB2/LYVE1/TGFB1/FGF2/CEMIP

PTH1R/GPR55/SPP1/TNFRSF11A/IL7/TF/CSF1R/SYK/IL6/PTK2B/CLDN18/ADAM8/CD38/S1PR1/TNFSF11/RAC2/INPP5D/HAMP/TMEM119/DCSTAMP

SYK/KLRC1/TCF7/NCKAP1L/GPR18/PTPRC

CD200/TSC22D1/AGT/GATA6/SMAD3/FERMT1/XCL1/LUM/CX3CL1/ATP6AP2/MYB/SERPINF2/PTGS2/THBS1/LGALS9/ADAM8/FOXP3

RARRES2/CX3CL1/CX3CR1/MDK/CCL3/CSF1/C3AR1/P2RY12/THBS1/CSF1R/TREM2/CMKLR1/IL34/CCL5

SLC6A1/P2RX7/TRPC4/TRH/NTSR1/CACNB4/ABAT/SV2A

CYP4A11/CYP1A2/CYP4F2/CYP2A13/CYP2B6/CYP2A7/CYP4F12/CYP1A1/CYP2C8/CYP1B1/CYP2E1/CYP2S1/CYP2C9

HPN/MDK/CFLAR/PTN/TNF/FGF1

CX3CL1/P2RX2/P2RX7/IL13/KCNN4/GALR2/CHP2/ADRB2/GPER1/CACNB2/LRRC55/NOS1AP/GRIN1/TRDN/NLGN3/APLNR/PPP3R2/KCNQ1/KCNC1/CCL2[illegible]

P2RY6/AKR1C3/EDN1/AKR1C2/PTGER2/PTGER4/SFRP1/PTGDR/CCL21/A2M/CCL19/CCR7

IL12B/STAT4/JAK2/IL12RB1/IL12A

GBP1/ELANE/GSDMC/MEFV/GSDMA/NLRC4/GSDMB/TREM2/GBP5/NLRP3/GZMA/NLRP1/ZBP1/GZMB/AIM2

PML/RGN/GP1BB/CX3CL1/CCL11/P2RX7/TCIRG1/CCL23/IL13/DHRS7C/CLIC2/CHRNA7/SYPL2/CYBA/SLC8A3/PRKCB/GPER1/CACNB2/CAV2/GP9/CASR/[illegible]

GRN/IGF1/LDLR/IL4/CD200R1/TREM2/TNFRSF1B/CST7/PTPRC

HBZ/HBD/IPCEF1/HBG1/HBB/HBA2

GJB2/GHSR/PLA2G4B/SLC6A4/OVGP1/CYP27B1/IGFBP5/PRDM1/SLC38A3/VDR/CLIC5/ADRA2A/EPYC/PGR/ITGA3/TAC3/PTN/CRHBP/LNPEP/SERPINE2/[illegible]

TTC39B/PPARG/TREM2/NFKBIA/IL6/NR1H3/ABCG1/CES1/TNF

DYRK2/ITPR1/MYOZ1/SLA2/PLEK/TBC1D10C/CD22

FGFBP1/GPLD1/NOS3/FBXW7/BMPER/ZC3H12A/PROX1/VASH1/SMOC2/PTN/RRAS/MMRN2/TEK/PPARG/HRG/KDR/FOXP1/LGMN/STC1/APOH/PTGS2/[illegible]

TPH1/FGF10/NLRP6/STATH/NPPB/SLC7A11/GATA1/TEC/TBXA2R/SCNN1G/ANXA8/ERBB4/AQP2/PLAU/GP1BB/TFPI2/VIP/ADAMTS18/PROC/ADTRP/F11/K

PPARG/AGTR1/CSF1/PLA2G5/NFKBIA/NR1H3/ABCG1/CETP/PLA2G2A/IL18/ADIPOQ

GPR68/CCR1/TNFRSF11A/IFNG/TYROBP/CSF1/IL12B/TREM2/TNF/TNFSF11/IL23A

IL12B/STAT4/JAK2/IL12RB1/IL12A

CHRNA7/SPINK1/GRIN2B/CYBA/SLC8A3/PRKCB/GPER1/PDGFRB/CSN2/CACNB2/GP9/MYLK/CACNA1G/ANXA9/CASR/TRPC4/CYP27B1/CATSPER3/TRF

EXPH5/CD33/SLC30A8/P2RX7/ISL1/IL13/KCNN4/IL1A/NR1H4/UCN3/PRKCB/GPER1/CASR/F2RL1/ADCY8/IGF1/GPLD1/TRH/INS/GPR68/PARD6A/NR0B2/PF

FAS/HTRA2/LGALS9/JAK2/MMP9/ST18

KCNJ1/P2RX7/VIP/TCIRG1/STRIT1/IL13/DHRS7C/SLC26A5/KCNN4/BPIFA1/CLIC2/GALR2/CHP2/SCN10A/SPINK1/GRIN2B/CYBA/BEST3/ADRB2/GPER1/B2

ITGA3/KCNQ1/CHRNA3/SULF1/TRPV5/AGTR1/SLC6A18/LGMN/HYAL2/STC1/CYP4F12/AKR1C3/SERPINF2/SLC22A12/SLC4A1/ADM/EDN1/ADORA2A/CHR

PTCRA/ADAM8/WNT5A/JAK3/BCL11B

PTPN3/GPLD1/ADRA2A/IFNG/IL10/TIMP3/TNFRSF1B/ADAM8/IL1B/TNF/TIMP4

VASH2/RAPGEF3/PLEKHO1/GCM1/ADGRB3/TYROBP/NPHS1/WNT1/JAM2/TREM2/IL4R/TNFSF14/DOCK2/NFATC2/CD53/CXCL9/CXCL10/DCSTAMP

LRRC19/BCR/FAS/ANXA1/P4HTM/RIPK3/KLF1/TRIM10/P2RX7/TCIRG1/AHSP/CX3CR1/MB/MUC2/CFLAR/MYB/B2M/SPRR2A/EMX1/SH2B2/RASSF2/L3MB

SMOC2/HRG/KDR/LGMN/TMSB4X/THBS1/FGF2/FGF1/CXCL13

SLIT3/SH2B3/EDN1/SLIT2/TREM2/PADI2/CCL5

GJB2/GHSR/CYP27B1/PRDM1/VDR/PTN/SPP1/STC1/PTGS2/NODAL/TCF23/GHRL/JUNB/STC2/LDOC1/LIF

APOBEC2/UPP2/APOBEC3F/APOBEC1/CDA/APOBEC3D/APOBEC3G/APOBEC3H/DPYD/AICDA/APOBEC3A

TMEM106A/REL/CLEC7A/TIFA/IRGM/EDN1/TMSB4X/CARD9/TRIM59/SECTM1/HMOX1/CASP10/IKBKE/NOD2/CD74/STAT1/TREM2/IFIT5/NFKBIA/CD4/TGI

FBXW7/GPR68/TYROBP/CLDN18/LILRB1/LTF/TNFSF11

PGLYRP2/HYAL2/STAB2/LYVE1/FUCA1/TGFB1/FGF2/CEMIP

AXL/MERTK/LRP2/WNT5A/RBP4

PRKCH/GFAP/MYB/VEGFC/IL6/CYSLTR1/LYN/LTA/IL1B/TNF

IL1RAP/IL17RA/CRLF2/IL9/IFNL1/LILRB4/IL1RL1/FOXP3/GATA3/IL33

IL1RAP/IL17RA/CRLF2/IL9/IFNL1/LILRB4/IL1RL1/FOXP3/GATA3/IL33

APOA1/ABCG4/STARD4/ABCG1/CES1/FGF1

CLCF1/IL4/IL10/IL2/IL27RA/TGFB1/CD28/FOXP3/TBX21/PTPRC

KCNMB2/DRD1/SCN11A/KCNA1/KCND2/SCN2A/KCNMB4/GPER1/GPR35/FGF12/GLRA1/KCNQ3/MTNR1B/P2RX1/KCNA2/CHRNB4/SCN4B/CACNA1I/HCI

INPP5F/HYAL2/PARP14/SOCS3/SAMSN1/SFRP1/CD300A/LILRB4/SOCS1/SEMA4D/PTPN6/PTPRC

P2RY12/PTGER4/CDH17/SELP/PLEK/CXCL13

CACNA1S/GATA1/PENK/XRN1/SLC26A3/P2RX2/SLC6A1/P2RX7/VGF/CNGA3/SLC6A3/SLC6A4/NT5E/IGFBP5/RAPGEF3/P2RY11/KCNJ8/DUOX2/KCNQ1/P2I

CCL2/ANGPTL4/TEK/KDR/AKR1C3/ITGA4/IL4/CDH5/FGA/THBS1/TGFB1/FGB/FASLG/GATA2/CD40/ECSCR/ICAM1/CD40LG/TNF/GATA3

SLC6A4/NT5E/IGFBP5/RAPGEF3/P2RY11/KCNJ8/DUOX2/KCNQ1/P2RY6/CRHBP/TEK/WNT10B/STC1/CDO1/DNTT/SLC5A5/P2RX1/PTGS2/P2RY2/RYR3/P2

CHI3L1/ADRB2/PDGFRB/RASSF2/MMD/PAK1/LAT/NRG1/ELANE/NTRK3/TPD52L1/ADCY8/IGF1/FBXW7/INS/ADRA2A/TLR3/PROX1/CHRNA3/TNFRSF10A/

CARD9/SYK/NOD2/CD74/WNT5A/LILRB1/SEMA7A/PLCG2/LAPTM5/TLR4/TLR7

PBLD/NPC1L1/ENPP7/PRSS1/DAO/FGF10/STATH/PTGER3/EPB41/CCKBR/MOGAT2/PRSS3/CTRB2/KCNN4/SLC9A4/SOAT2/SPINK1/MUC2/MDK/CLPS/MU

LRFN5/TAFA3/GRN/BPI/ZC3H12A/LDLR/FCGR2B/IL31RA/NR1H3/CST7/PTPRC/ADGRF5

HHLA2/CD274/IL2/IL12B/PDCD1LG2/LGALS9/LILRB4/CRTAM/IL2RA/FOXP3/IL12RB1/IL23A/CD24/IL18

CHRNA7/F3/ANGPT4/MDK/IL1A/CELA1/CHI3L1/PRKCB/GHSR/KRT1/VASH2/NOS3/RAPGEF3/STAT3/TNN/BMPER/ZC3H12A/APLNR/TNMD/CMA1/TLR3/C

APOA1/LIPC/ABCG1/LIPG/CES1/CETP/CLU

GJB2/GHSR/CYP27B1/PRDM1/VDR/PGR/ITGA3/PTN/SPP1/STC1/DSG2/PTGS2/NODAL/TCF23/ADCY7/GHRL/JUNB/STC2/LGALS9/HAVCR2/LDOC1/LIF

ALKBH1/NTF3/MET/SMAD3/COLEC10/FPR2/AZU1/CXCL12/CCL16/ARTN/F7/CCL15/FGF10/WNT7B/CX3CL1/AGER/F3/CASR/F2RL1/NTRK3/CCL3/IL16/CX

CCR3/FGF10/APELA/NGFR/ACVRL1/RGCC/LRG1/CCL11/VIP/NRARP/PRKD2/F3/MDK/CAV2/GHSR/VASH2/FGFBP1/COL8A2/STAT3/BMPER/APLNR/POLD

PDGFRB/CASR/CCL3/PPP3R2/CCL2/STC1/P2RX1/AKAP5/MS4A1

FPR2/CXCL12/DEFB124/CX3CR1/S100A14/CCL1/CCR1/LGMN/CCR2/AIF1/PLA2G7/CCL5/CXCL10

CACNB2/CACNA1G/SCN5A/SCN2B/SCN3B/SCN4B/CACNA1C/SCN1B/SLC8A1/ANK3

PML/RGN/GP1BB/CX3CL1/CCL11/P2RX7/TCIRG1/TRPV6/CCL23/IL13/DHRS7C/CLIC2/CHRNA7/SYPL2/CYBA/SLC8A3/PRKCB/GPER1/CACNB2/CAV2/S100

FOXF1/BCR/CX3CR1/FCGR2B/CCR2/LGALS9/CD300A/SPI1/CD84

ARG1/BCL6/SOCS5/ANXA1/ARG2/IFNB1/IFNL1/ASCL2/CCR2/IL27RA/TBX21

PROX1/KDR/NRP1/LIPA/DLL1/TMEM100

SYK/IL4R/LGALS9/CD300A/GATA2/TFEB/MILR1/PTGDR/LYN/UNC13D/CD84/PTGDS/RASGRP1/CLNK/BTK/PIK3CG/FGR/RAC2/LAT2

SYK/IL4R/LGALS9/CD300A/GATA2/TFEB/MILR1/PTGDR/LYN/UNC13D/CD84/PTGDS/RASGRP1/CLNK/BTK/PIK3CG/FGR/RAC2/LAT2

CD1E/B2M/UNC93B1/LNPEP/LGMN/FCGR2B/FCGR1A/CLEC4A/CD1C/CD74/FCER1G/MPEG1/CTSS/CD1D/IFI30

IL4/CFH/IGHG1/CR1L/C3/CR1

APOBEC2/UPP2/APOBEC3F/APOBEC1/ENPP4/CDA/APOBEC3D/APOBEC3G/APOBEC3H/AICDA/ADA2/APOBEC3A

IL1A/TMEM131L/FOXP3/IL1B/TNFRSF9

IL1A/TMEM131L/FOXP3/IL1B/TNFRSF9

CHI3L1/ADRB2/PDGFRB/RASSF2/MMD/PAK1/LAT/NRG1/ELANE/NTRK3/TPD52L1/ADCY8/IGF1/FBXW7/INS/ADRA2A/TLR3/PROX1/CHRNA3/TNFRSF10A/

AKR1C3/PTGS2/CYP4A22/EDN1/NLRC4/CYP2C8/SYK/CD74/PLA2G5/ALOX5/PTGES3/CYP2C9/GGT5/ALOX5AP/IL1B/PTGDS

SMAD7/ISL1/EFEMP2/ZFPM2/MYLK/NRG1/NOG/NKX2-5/PROX1/XIRP2/MYBPC3/ACTC1/WNT2/TNNC1/ANKRD1/HAND1/HEG1/MYL2/TNNI1/PKP2/FGFR2

IFNK/IFNA8/IFNW1/IFNB1/IFNA4/IFNA1/IFNG/IFNA6/IFNA14/LIF

SYK/NOD2/TCF7/LILRB1/NCKAP1L/PTPRC

ISL1/IL1A/RORA/STAT3/SULF1/PTGS2/NODAL/C3AR1/CYP1B1/IL6/TGFB1/SULF2/HPSE/IL1B/C3/FLT4

SMAD7/ZC3H12A/ASCL2/IL2/IL27RA/FOXP3/TBX21

CHRNA7/F3/ANGPT4/MDK/IL1A/CELA1/CHI3L1/PRKCB/GHSR/KRT1/VASH2/NOS3/RAPGEF3/STAT3/TNN/BMPER/ZC3H12A/APLNR/TNMD/CMA1/TLR3/C

SKAP1/P2RY12/CDH17/PLEK/CXCL13

SLIT3/F2RL1/PYDC1/SH2B3/CCDC3/USP18/CYLD/PPARG/APOA1/OAS3/SLIT2/PARP14/OAS1/TREM2/ISG15/IL6/SAMHD1/NR1H3/NLRC5/PADI2/MMP12/IL1

IL1RAP/IL17RA/CRLF2/IL9/IL1RL1/GATA3/IL33

EGR2/NRP1/KCNA2/PLXNA4/SEMA3A

IFNK/IFNA8/IFNW1/IFNB1/IFNA4/IFNA1/IFNG/IFNA6/IFNA14/LIF

B2M/UNC93B1/LNPEP/LGMN/FCGR2B/FCGR1A/CLEC4A/CD74/FCER1G/MPEG1/CTSS/IFI30

STAT3/CARD18/AHSG/SERPINB5/BCL2L10/MEFV/SERPINB3/FETUB/NKX3-1/SERPINB8/SERPINA11/DPEP1/SPINK2/PPARG/SERPINE2/HRG/LGMN/SERP

GBP5/PTPN22/PLCG2/BTK/TLR6/TLR4

NRG1/NOG/NKX2-5/PROX1/XIRP2/MYBPC3/ACTC1/WNT2/TNNC1/ANKRD1/HAND1/HEG1/MYL2/TNNI1/PKP2/FGFR2/TGFB1/LRP2/WNT5A/FOXC2/DLL4/

XCL1/CCL16/CCL15/CX3CL1/CCL11/CCL23/CCL1/CCL3/CCL2/CCL7/CCL25/CCL14/CCL4/CCL22/CCL24/XCL2/DAPK2/CCL8/CCL5

PROX1/VASH1/KDR/SOX18/SYK/HEG1/SVEP1/PDPN/VEGFC/EFNB2/TIE1/FOXC2/CLEC14A/FGF2/FLT4/TBX1

BATF/CLCF1/IL4/IL10/IL2/IL27RA/TGFB1/AICDA/CD28/FOXP3/CD40LG/TBX21/PTPRC/CCR6

BATF/CLCF1/IL4/IL10/IL2/IL27RA/TGFB1/AICDA/CD28/FOXP3/CD40LG/TBX21/PTPRC/CCR6

BATF/CLCF1/IL4/IL10/IL2/IL27RA/TGFB1/AICDA/CD28/FOXP3/CD40LG/TBX21/PTPRC/CCR6

APOA1/FGA/IL10/FGB/KLF4/IL1B/IL1RN/LCK/TNF/ADIPOQ

NBEAL2/CFLAR/BATF2/MYB/CSF2/B2M/OSCAR/IAPP/RASSF2/L3MBTL3/GP9/EPB42/ERFE/F2RL1/CCL3/STAT3/FBXW7/GPR68/TLR3/ZNF385A/TSPAN2/SH

IL1A/TMEM131L/FOXP3/IL1B/TNFRSF9

IL1A/TMEM131L/FOXP3/IL1B/TNFRSF9

PRKCZ/BCL6/SOCS5/ANXA1/ASCL2/IL4R/CD86/NLRP3/IL18

TDO2/KYNU/KMO/IDO2/IDO1

THBS1/IL10/DLL1/GHRL/SMAD1/KLF2/ALOX5/ITGA5/KLF4/JCAD/PPP1R16B/FGF2/FGF1

APELA/NGFR/ACVRL1/NRARP/FGFBP1/BMPER/APLNR/MMRN2/AGTR1/HMOX1/THBS1/IL12B/GATA2/DLL4/IL12A/JCAD/PPP1R16B/FGF2

AGER/ADRB2/ATP1A3/IGF1/LGMN/ITGA4/FCGR2B/EPHB2/VCAM1/TREM2/ICAM1/TNF/TLR6/TLR4

CX3CL1/P2RX2/P2RX7/STRIT1/IL13/GPER1/PDGFRB/CACNB2/MYLK/CASR/GRIN1/TRDN/CCL3/APLNR/PPP3R2/CCR1/CCL2/P2RY6/NTSR1/STC1/LACRT

PTPRT/CHI3L1/STAT3/CEBPA/SELPLG/IL6/JAK2/CITED1/SPI1/ST18

FGFBP1/GPLD1/NOS3/ZC3H12A/PROX1/SMOC2/RRAS/TEK/KDR/FOXP1/LGMN/PTGS2/NRP1/EDN1/TMSB4X/PLPP3/ATOH8/HMOX1/THBS1/VEGFC/PLG/

ADRB2/FBXW7/NLGN3/ADRA2A/CCL2/NCOA3/NRP1/IFNG/ADRA2B/EDN1/EPHB2/CCR2/IL10/JAK2/RELN

CD274/IRF7/TNIP3/CD300LF/TLR6/TLR4

APOE/ABCA1/MYD88/SOCS5/CD68/ADTRP/CASR/PPARG/LDLR/TXNIP/SYK/TREM2/TGFB1/FCER1G/SMPD3/CES1/ITGB2/TLR6/TLR4

MYB/IAPP/ERFE/CCL3/FBXW7/GPR68/TLR3/GPR55/KLF13/MAFB/ZFP36/IL4/DLL1/C1QC/NFKBIA/ZBTB16/PTK2B/CLDN18/SFRP1/GATA2/LYN/GPR171/LILF

PTPRT/STAT3/CEBPA/SELPLG/IL6/JAK2/SPI1/ST18

CSF1/C3AR1/THBS1/CSF1R/CMKLR1/PTK2B/IL34/SLAMF8/CCL5/STAP1

SLIT3/F2RL1/PYDC1/SH2B3/CCDC3/USP18/CYLD/PPARG/APOA1/OAS3/SLIT2/PARP14/OAS1/TREM2/ISG15/IL6/SAMHD1/NR1H3/NLRC5/KLF4/PADI2/MMF

LHCGR/CYP4A11/LEP/DRD3/PLA2G10/CYP4F2/DRD4/SSTR4/PLA2G2E/PLA2G3/P2RX7/AVPR1B/PLA2G4F/IL1A/NTSR1/NMUR2/TNFRSF11A/OC90/PTGS2

HHLA2/CD274/IL2/IL12B/PDCD1LG2/LGALS9/LILRB4/CRTAM/IL2RA/FOXP3/IL12RB1/IL23A/CD24/IL18

HTR1A/DRD5/SGCG/NPPB/CAMK2D/DRD1/BMP10/APELA/KCNE3/CORIN/TBXA2R/ACVRL1/SCN11A/SCNN1G/UTS2/BCR/MYOF/KCNE4/GRK2/SMAD7/A

MYD88/DEFB124/CARD9/IRF5/NOD2/TREM2/IL6/INAVA

HTR1A/DRD5/NPPB/DRD1/TBXA2R/UTS2/ADRB1/AVPR1B/PER2/ITGA1/KCNMB4/AVP/ADRB2/GPER1/KCNMA1/ADRA1B/CASR/SLC6A4/F2RL1/NOS3/KCI

BPI/NOS3/LBP/TLR2/LACRT/DEFB114/PTAFR/NFKBIA/HCK/LYN/IRAK2/LY86/TIFAB/PTPN22/LTF/PLCG2/CD180/CD6/SCIMP/TLR4

VASH2/RAPGEF3/PLEKHO1/GCM1/ADGRB3/TYROBP/NPHS1/WNT1/JAM2/TREM2/IL4R/TNFSF14/DOCK2/NFATC2/CD53/CXCL9/CXCL10/DCSTAMP

VASH2/RAPGEF3/PLEKHO1/GCM1/ADGRB3/TYROBP/NPHS1/WNT1/JAM2/TREM2/IL4R/TNFSF14/DOCK2/NFATC2/CD53/CXCL9/CXCL10/DCSTAMP

IL12B/CD226/TGFB1/HAVCR2/CRTAM/IL12A/NKG7

IFNK/IFNA8/USP18/IFNW1/IFNB1/IFNA4/IFNA1/OAS3/IFNA6/RBM47/IKBKE/IFNA14/OAS1/STAT1/IFITM1/ISG15/OAS2/WNT5A/SAMHD1/NLRC5/IRF7/MMP1

CLDN23/CNTN2/FAT4/PCDHGA9/MYPN/CLDN3/PCDH1/AMIGO1/CDH4/DSG4/PCDHGA10/ESAM/CX3CL1/CDHR2/CLSTN2/PCDHB9/HMCN2/CDH20/CD

KDR/FOXP1/STC1/SOX18/NRP1/EDN1/ATOH8/CDH5/LIPA/GDF2/HEG1/PDPN/DLL1/PECAM1/ITGAX/TIE1/HPSE/FOXC2/DLL4/TJP3/ICAM1/GJA4/FOXP3/IL

GPR68/FOXP1/CSF1/CD74/CD4/IRF7/IL34/INPP5D/DCSTAMP

IL1A/RORA/STAT3/SULF1/PTGS2/NODAL/C3AR1/CCR2/CYP1B1/IL6/TGFB1/SULF2/HPSE/IL1B/C3/FLT4

HAVCR2/RSAD2/TLR9/PIK3AP1/LILRA4/FCRL3

AGER/CHRNA7/GAREM1/OR2AT4/IL1A/CFLAR/GBP1/CHI3L1/AVP/GPER1/PDGFRB/CASR/NRG1/LMO3/CCL1/EPHB1/F2RL1/IGF1/CCL3/FBXW7/BMPER/F

P2RY6/GPR55/PLCB2/AGTR1/PNLIP/APOH/APOA5/FGFR3/ADRA1A/PDGFRA/PLA2G5/FGFR2/TXK/CD86/NR1H3/S1PR4/ITK/SELE/CCL5

ARG2/IL1RAP/IL17RA/IFNL1/LILRA5/IL4/LGALS9/GATA3/IL33/IL18

ARG2/IL1RAP/IL17RA/IFNL1/LILRA5/IL4/LGALS9/GATA3/IL33/IL18

OPRK1/PBLD/NPC1L1/ENPP7/FGF10/STATH/PTGER3/EPB41/CCKBR/MOGAT2/KCNN4/SLC9A4/SOAT2/SPINK1/MUC2/MDK/MUC13/ABCG5/GHSR/VDR/S

GHSR/CDKN2A/ST6GAL1/NOD2/IRF7/CCR5/CCL5

HTR1A/DRD5/NPPB/DRD1/TBXA2R/UTS2/ADRB1/AVPR1B/PER2/ITGA1/KCNMB4/AVP/ADRB2/GPER1/KCNMA1/ADRA1B/CASR/SLC6A4/F2RL1/NOS3/KCI

HTR1A/DRD5/NPPB/DRD1/TBXA2R/UTS2/ADRB1/AVPR1B/PER2/ITGA1/KCNMB4/AVP/ADRB2/GPER1/KCNMA1/ADRA1B/CASR/SLC6A4/F2RL1/NOS3/KCI

ABRA/WNT2/PPARG/FOXA1/TLR2/HSPA1B/TNFRSF11A/NFAM1/CLEC7A/EDN1/NODAL/TRIM14/CARD9/PLPP3/NLRC4/IKBKE/WNT1/LPAR5/NOD2/IL10/IL

IL1A/RORA/STAT3/SULF1/PTGS2/NODAL/C3AR1/CCR2/CYP1B1/IL6/TGFB1/SULF2/HPSE/IL1B/TNF/C3/FLT4

GCM1/ADM/IL10/SOCS3/FGFR2/RSPO3/SPINT2/ST14

RFFL/CFLAR/NOS3/FAIM2/BMP5/HMOX1/FGA/FGB/ICAM1/HGF

TMEM106A/MAP3K14/REL/CLEC7A/TIFA/IRGM/EDN1/TMSB4X/CARD9/TRIM59/SECTM1/HMOX1/CASP10/IKBKE/NOD2/NFKB2/TLR8/CD74/STAT1/TREM2

ABCA1/MYD88/SOCS5/CD68/ADTRP/CASR/PPARG/LDLR/TXNIP/SYK/TREM2/TGFB1/FCER1G/SMPD3/CES1/ITGB2/TLR6/TLR4

CCDC3/CYLD/HSPA1B/TNFRSF11A/TNFRSF25/APOA1/TIFA/TMSB4X/TNFSF13B/TNFRSF18/SYK/STAT1/NFKBIA/PTK2B/TNFRSF1B/CLDN18/TNFRSF4/

DHX58/USP18/OAS1/USP44/MX1/IFIH1/NLRP1/IFIT1/ZBP1/TNF/IFIT2/IFIT3/CXCL10

DOCK8/CCL21/CRTAM/CCL19/DOCK2/CCR7

STAT1/GATA3/CD24/MMP9/LIF

LIME1/TRDN/CCL3/APLNR/P2RY6/NTSR1/PLCB2/LACRT/TRPM2/ITPR1/RYR3/JPH3/F2/CASQ1/PTK2B/CACNA1C/FASLG/THY1/CCL21/LYN/SLC8A1/CCL

HRG/CD274/IL12B/CD226/TGFB1/HAVCR2/CRTAM/PRF1/IL12A/IL4I1/NKG7

GNAT2/DIO3/GNAT1/CABP4/USH1C/CRB2/THRB/THY1/HCN1

NRARP/MDK/GHSR/STAT3/CCL2/PPARG/ITGA4/GHRL/ALDH1A2/FGF2/COL4A3

NRARP/MDK/GHSR/STAT3/CCL2/PPARG/ITGA4/GHRL/ALDH1A2/FGF2/COL4A3

IFNG/ADM/ABCG4/STAR/STARD4/ABCG1/TNF/FGF1

EGR2/PTN/TENM4/SERPINE2/NKX6-2/OLIG2/TLR2/CLCF1/TGFB1/TNFRSF1B/RNF112/IL34/BMP2/CXCR4/IL33/LIF

AXL/LGALS9/CCL21/LYN/LILRB1/CCL19/CCR7

AXL/LGALS9/CCL21/LYN/LILRB1/CCL19/CCR7

CX3CL1/P2RX2/P2RX7/STRIT1/IL13/KCNN4/GALR2/CHP2/ADRB2/GPER1/PDGFRB/CACNB2/CNTN1/MYLK/CASR/SLC6A4/SCN5A/LRRC55/NOS1AP/GRIN

GJB2/GHSR/CYP27B1/VDR/PTN/SPP1/STC1/PTGS2/TCF23/GHRL/JUNB/STC2/LIF

TLR3/IFNB1/IRGM/IKBKE/OAS1/STAT1/IFITM1/MNDA/XAF1/IRF1/PYHIN1/AIM2

APOB/PLA2G2E/PLA2G3/AGTR1/MPO/LIPC/PLA2G5/ABCG1/PLA2G7/CETP/PLA2G2A

IFNB1/IL4/IL6/NLRP3/RSAD2/TBX21/GATA3

IFNB1/IL4/IL6/NLRP3/RSAD2/TBX21/GATA3

LGALS9/CD28/PRKCQ/CD3E/CARD11

CLEC7A/OAS3/SYK/NOD2/OAS1/LGALS9/IL1B/CD84/ADIPOQ

CLEC7A/OAS3/SYK/NOD2/OAS1/LGALS9/IL1B/CD84/ADIPOQ

MYB/GPER1/PDGFRB/ELANE/IGFBP5/IGF1/NOS3/PTGIR/P2RY6/PPARG/FOXP1/MAP3K5/SERPINF2/PTGS2/IFNG/EDN1/CCN4/CALCRL/SOD2/HMOX1/TH

STAT3/GPR68/CCR1/THPO/HSPA1B/TNFRSF11A/IFNG/TYROBP/CSF1/RHEX/IL12B/CD74/TAL1/STAT1/TREM2/ISG15/CD4/TGFB1/GATA2/IL34/EVI2B/HCL

TLR3/TLR2/OAS3/IRF5/OAS1/TLR8/ISG15/OAS2/IRF7/TLR9/IFIH1/IRF1/TLR4/TLR7

NFKBIZ/STAT3/IL17RA/IKBKE/SOCS3/IL1B/CXCL10

NFKBIZ/STAT3/IL17RA/IKBKE/SOCS3/IL1B/CXCL10

OSCAR/IAPP/RASSF2/ERFE/CCL3/FBXW7/GPR68/TLR3/CCR1/GPR55/FOXP1/MAFB/TNFRSF11A/IFNG/CCN4/IL4/TYROBP/CSF1/TF/CSF1R/IL12B/TREM

GBP5/PTPN22/PLCG2/BTK/TLR6/TLR4

BCR/RARRES2/CX3CL1/CNN2/CX3CR1/MDK/CCL3/CSF1/C3AR1/CCR2/P2RY12/THBS1/CSF1R/CD200R1/TREM2/CMKLR1/PTK2B/IL34/SLAMF8/CCL5/ST

FBXW7/GPR68/FOXP1/TYROBP/CLDN18/LILRB1/LTF/TNFSF11

FBXW7/GPR68/FOXP1/TYROBP/CLDN18/LILRB1/LTF/TNFSF11

CCL2/C4B/ARRB2/RAB31/RIT2/FCGR2A/TLR2/CFP/SYNE1/AXL/APOA5/C2/APOA1/CLEC7A/SNAP91/IFNG/IL4/FCGR2B/FCGR1A/TF/SYK/NOD2/DLL1/STX1

AVP/GHSR/GPR35/FGF12/GLRA1/TNR/TYMP/MTNR1B/CHRNB4/GHRL

TPH2/TPH1/HTR1A/SLC34A1/ACMSD/HAAO/ASMT/GRIN2A/TDO2/PDE1B/KYNU/KMO/IDO2/IL4I1/IDO1

PPARG/AGTR1/CSF1/STAT1/PLA2G5/NFKBIA/TGFB1/WNT5A/NR1H3/ABCG1/CETP/PLA2G2A/IL18/ADIPOQ

MUC2/MUC13/HTR4/MUC6/MUC4/NOD2/INAVA/IL10RA/TLR9/VSIG1/SERPINA3/RBP4/TLR4

CCL2/HRG/ST6GAL1/SLIT1/NRP1/SEMA3G/SLIT2/SEMA4A/THBS1/AIF1/PTPRO/WNT5A/PADI2/SLAMF8/SEMA7A/SEMA4D/KLRK1/GPR18/SEMA3A/STAP1/

IL12B/CD226/TGFB1/HAVCR2/CRTAM/IL12A/NKG7

CD274/PDCD1LG2/LGALS9/LILRB4/CRTAM/FOXP3

XCL1/MICB/SLC22A13/IL23R/CD1B/RAET1G/P2RX7/CD1E/B2M/CD1C/IL12B/RAET1L/IL12RB1/CD1D/IL12A/PTPRC/IL23A

SYK/IL4R/LGALS9/CD300A/GATA2/LYN/UNC13D/CD84/FGR/RAC2

AGER/IL1A/TNIK/RASSF2/F2RL1/TPD52L1/FZD10/EDAR/TLR3/GADD45G/MAP3K5/SERPINF2/NRK/FCGR2B/CARD9/SH3RF2/NOD2/GADD45B/PTK2B/WN

RHCG/UPK3A/SLC14A1/HBZ/HBD/AQP3/HBG1/HBB/HBA2

RHCG/HBZ/HBD/HBG1/HBB/HBA2

P2RX2/P2RX7/GPR34/P2RY11/ADORA3/P2RY6/P2RX1/ADORA2A/P2RY2/P2RY12/P2RY14/PTAFR/ADCY5/GPR171/P2RY13/P2RY8/P2RX5

ELF4/IL12B/JAK2/IL15/PTPN22/IL23A/IL18

PDIA2/GP9/SLC6A4/KRT1/F2RL1/NOS3/HNF4A/JMJD1C/VWF/ADRA2A/ZNF385A/SH2B3/ENPP4/MYL9/C1QTNF1/TSPAN32/FERMT3/SERPINE2/HRG/SER

IL4/IL2/IL27RA/CD28/FOXP3/TBX21/PTPRC

EHD1/SCGB3A1/MYOD1/RIPOR2/CFLAR/PLEKHO1/IL4R/TNFSF14/NFATC2/CD53/CXCL9/CXCL10

USH1C/CRB2/THRB/THY1/HCN1

MUC2/MUC13/HTR4/MUC6/MUC4/LACRT/LDB2/NOD2/INAVA/ILDR1/IL10RA/TLR9/VSIG1/SERPINA3/RBP4/TLR4

APOBEC2/APOBEC3F/APOBEC1/APOBEC3D/APOBEC3G/RBM47/A1CF/APOBEC3H/AICDA/APOBEC3A

IAPP/ERFE/CCL3/FBXW7/TLR3/GPR55/MAFB/IL4/CLDN18/SFRP1/LILRB4/LILRB1/LTF/TLR4/INPP5D

MYB/GPER1/PDGFRB/ELANE/IGFBP5/IGF1/NOS3/PTGIR/P2RY6/PPARG/FOXP1/MAP3K5/SERPINF2/PTGS2/IFNG/EDN1/CCN4/CALCRL/SOD2/HMOX1/TH

SLC10A1/CASR/SLC51A/UGT1A3/ABCB11/SLC51B/CES1/TNF

PLXND1/NTRK2/WNT2/CYLD/TENM4/SERPINE2/SKIL/NKX6-2/WNT10B/OLIG2/TLR2/TNFRSF11A/AXL/CD83/CLCF1/CLEC7A/VSIR/NRP1/IFNG/FOXG1/IL4

DOCK8/CCL21/CRTAM/CCL19/DOCK2/CCR7

SKAP1/FERMT3/PODXL/CD3E/CCL5/CXCL13

CCL2/ANGPTL4/TEK/KDR/AKR1C3/MAP3K5/ITGA4/IL4/CDH5/FGA/THBS1/IL10/TGFB1/FGB/FASLG/GATA2/CD40/ECSCR/ICAM1/CD40LG/TNF/GATA3/COL

IL20RB/FCGR2B/SYK/NOD2/IL10/IL12B/TREM2/HCK/LYN/SPN/FGR

BATF/CLCF1/IL4/IL10/IL2/IL27RA/TGFB1/AICDA/CD28/GAPT/FOXP3/CD40LG/TBX21/BTK/PTPRC/CCR6

F2RL1/COLEC12/UNC93B1/TLR3/OAS1/RFTN1/HAVCR2/PTPN22/TNF/SCIMP

GCNT1/FUT4/ELANE/ITGA4/CHST4/CCR2/PTAFR/FUT7/SELP/SELE

MAP3K5/AXL/TRPM2/EDN1/TXNIP/SOD2/HMOX1/CYP1B1/STAT1/KCNA5/IL6/KLF2/IL18RAP/PTK2B/AREG/SMPD3/KLF4/MAPK13/IL18BP/STK26/HBB/HBA2

PDIA2/GP9/SLC6A4/KRT1/F2RL1/NOS3/HNF4A/JMJD1C/VWF/ADRA2A/SH2B3/ENPP4/MYL9/C1QTNF1/TSPAN32/FERMT3/SERPINE2/HRG/SERPINA1/SE

CCL3/STAT3/FBXW7/GPR68/TLR3/CCR1/GPR55/THPO/KLF13/FOXP1/HSPA1B/MAFB/TNFRSF11A/ZFP36/CSF3R/IFNG/IL4/TYROBP/CSF1/RHEX/IL12B/C

GP6/CTNNA3/CELA2A/PDIA2/SLC6A4/SH2B3/MYL9/C1QTNF1/CDHR5/TSPAN32/FERMT3/SERPINE2/DSG2/DSC2/PLPP3/P2RY12/FGA/SYK/PDPN/PKP2/P

SMAD7/ISL1/GSC/EFEMP2/ZFPM2/MYLK/NRG1/NOG/NKX2-5/PROX1/XIRP2/MYBPC3/ACTC1/WNT2/TNNC1/ANKRD1/HAND1/HEG1/MYL2/TNNI1/PKP2/F

TLR2/NMI/RELB/REL/OAS3/IRF5/OAS1/TLR8/ISG15/OAS2/IRF7/LILRB1/TLR9/IFIH1/IRF1/TLR4/NLRC3/TLR7

TLR2/NMI/RELB/REL/OAS3/IRF5/OAS1/TLR8/ISG15/OAS2/IRF7/LILRB1/TLR9/IFIH1/IRF1/TLR4/NLRC3/TLR7

LBP/SLC46A2/DHX58/HSPA1B/TLR1/DDX60/SLC15A3/GBP5/NR1H3/RSAD2/TLR9/PTPN22/LTF/PLCG2/BTK/TLR6/TLR4/OASL

EHD1/SCGB3A1/MYOD1/RIPOR2/IL4R/TNFSF14/NFATC2/CD53/CXCL9

LIME1/TRDN/CCL3/APLNR/P2RY6/NTSR1/PLCB2/LACRT/TRPM2/ITPR1/RYR3/JPH3/F2/CASQ1/PTK2B/CACNA1C/FASLG/THY1/CCL21/LYN/SLC8A1/CCL

STAT3/CARD18/AHSG/SERPINB5/BCL2L10/MEFV/SERPINB3/FETUB/NKX3-1/SERPINB8/SERPINA11/DPEP1/SPINK2/PPARG/SERPINE2/HRG/LGMN/SER

IFNK/IFNA8/USP18/IFNW1/IFNB1/IFNA4/IFNA1/OAS3/IFNA6/RBM47/IKBKE/IFNA14/OAS1/STAT1/IFITM1/ISG15/OAS2/WNT5A/SAMHD1/NLRC5/IRF7/MMP1

LRRC15/RESF1/TRIM34/CXCL8/PROX1/CH25H/HMGA2/CD209/IFNB1/AXL/ZFP36/OAS3/TRIM14/TRIM59/APOBEC3D/APOBEC3G/OAS1/CD74/APOBEC3H

DHX58/TLR2/OAS3/IRF5/IKBKE/SYK/OAS1/TLR8/STAT1/ISG15/OAS2/IRF7/MMP12/TLR9/IFIH1/IRF1/PTPN22/PLCG2/TRAF3IP3/TLR4/TLR7

ARRB2/CLEC12B/SERPINB4/TGFB1/LGALS9/KLRC1/HAVCR2/LILRB1/SERPINB9/KLRD1

CXCL17/DUSP1/FPR2/CXCL12/GREM1/DEFB124/CX3CR1/S100A14/CCL1/CCR1/LGMN/SLIT2/CCR2/AIF1/LYN/SLAMF8/PLA2G7/CCL5/CXCL10

GPR68/CCR1/TNFRSF11A/IFNG/TYROBP/CSF1/IL12B/CD74/TREM2/CD4/TGFB1/IL34/EVI2B/HCLS1/TNF/TNFSF11/IL23A/LIF/DCSTAMP

BATF2/GBP1/TSPAN32/BATF/CLEC7A/GBP4/IL10/IL12B/IL4R/BCL3/CCDC88B/NKG7/IRF8

PDIA2/GP9/SLC6A4/KRT1/F2RL1/NOS3/HNF4A/JMJD1C/VWF/ADRA2A/SH2B3/ENPP4/MYL9/C1QTNF1/TSPAN32/FERMT3/SERPINE2/HRG/SERPINA1/SE

GCNT1/FUT4/ELANE/ITGA4/CCL25/CHST4/CCR2/PTAFR/FUT7/CCL21/SELP/SELE

GREM1/BCR/P2RX7/IAPP/PPARG/IL6/CLDN18/SFRP1/CD38/INPP5D/HAMP/TMEM119

PODXL/MTSS1/EDN1/NPHS1/STAT1/PTPRO/FOXC2/BASP1/GATA3/CD24/MMP9/LIF/ADIPOQ

SYK/CD226/HCK/FCER1G/LYN/MYO1G/FGR/VAV1/PTPRC

GP1BB/GP9/SVEP1/JAK2/PEAR1/SELP/PLEK/TLR4

CSF1/ASCL2/CSF1R/VEGFC/TREM2/IL6/PTK2B/CYSLTR1/AREG/LYN/LTA/IL34/IL1B/TNF/IL33/CLU

PPARG/AGTR1/CSF1/STAT1/PLA2G5/NFKBIA/TGFB1/WNT5A/NR1H3/ABCG1/CETP/PLA2G2A/IL18/ADIPOQ

DEFA5/DEFB128/CST11/NFKBIZ/BPI/LYPD8/ELANE/F2RL1/IL22RA1/DEFB1/PRB3/LBP/RNASE3/IRGM/DEFB127/F2/DEFB114/IL12B/TREM2/CD4/IL6/LYZ/RN

CSF1/CSF1R/TREM2/IL34/IL33/CLU

RESF1/CXCL8/PROX1/HMGA2/IFNB1/OAS3/APOBEC3D/APOBEC3G/OAS1/APOBEC3H/IFITM1/ISG15/IFIT5/OAS2/AICDA/MX1/NR5A2/RSAD2/CD28/IFIH1/I

RBM47/IKBKE/WNT5A/NLRC5/IRF7/MMP12/ZBP1

IAPP/ERFE/CCL3/FBXW7/GPR68/TLR3/GPR55/MAFB/IL4/C1QC/CLDN18/SFRP1/GATA2/LYN/LILRB4/LILRB1/LTF/INHBA/TLR4/INPP5D/ADIPOQ

BCR/GRN/S100A9/NOXO1/INS/LBP/CLEC7A/CAMK1D/SLAMF8/RAC2/IGHA2/JCHAIN

CCR3/FGF10/APELA/NGFR/ACVRL1/RGCC/LRG1/CCL11/VIP/NRARP/PRKD2/F3/MDK/CAV2/GHSR/VASH2/FGFBP1/STAT3/APLNR/POLD4/TNMD/FGF7/F

LILRB2/LILRB4/IL2RA/FOXP3/CD3E/IDO1

FGL2/RAG2/BCL6/RUNX1/SOCS5/ANXA1/SMAD7/NRARP/MDK/ZC3H12A/IFNB1/IFNL1/ASCL2/LAG3/IL2/CD74/IL4R/DTX1/TMEM131L/LILRB4/FOXP3/SOCS

PREX2/ERFE/F2RL1/IGFBP5/NTRK3/IGF1/LIME1/CCL3/RAPGEF3/FAM110C/STAT3/INS/SH2B3/C1QTNF1/NKX3-1/RRAS/THPO/NTRK2/ARRB2/TEK/SERPI

AGER/CHRNA7/GAREM1/OR2AT4/IL1A/CFLAR/GBP1/CHI3L1/GPER1/PDGFRB/CASR/NRG1/LMO3/CCL1/EPHB1/F2RL1/IGF1/CCL3/FBXW7/BMPER/FGF2

BCL6/SOCS5/ANXA1/SMAD7/ZC3H12A/ASCL2/IL2/IL4R/FOXP3/TBX21/JAK3

IL4/PARP14/RUFY4/IL4R/TCF7/CD300LF/JAK3/GATA3/CORO1A/IL2RG/DCSTAMP

LRRC15/RESF1/TRIM34/CXCL8/PROX1/CH25H/HMGA2/CD209/IFNB1/AXL/OAS3/TRIM59/APOBEC3D/APOBEC3G/OAS1/CD74/APOBEC3H/IFITM1/ISG15/II

CALB1/CACNB2/CACNB4/TSPOAP1/P2RX1/ERC2/ATP2B3/SV2B/CALB2

IAPP/RASSF2/ERFE/CCL3/FBXW7/GPR68/TLR3/CCR1/GPR55/MAFB/TNFRSF11A/IFNG/IL4/TYROBP/CSF1/IL12B/TREM2/CAMK4/CLDN18/SFRP1/LILRB4/

LILRB2/CD274/LILRB4/FOXP3/CD3E/IDO1

PGLYRP4/PGLYRP2/TLR2/TLR1/NLRC4/NOD2/CD1D/TLR6

IFNK/IFNA8/CH25H/USP18/IFNW1/IFNB1/IFNA4/IFNA1/OAS3/IFNA6/RBM47/IKBKE/IFNA14/OAS1/STAT1/IFITM1/ISG15/OAS2/WNT5A/SAMHD1/MX1/NLRC!

TYROBP/SYK/PTAFR/FCER1G/PRAM1/SPI1/ITGB2/DNASE1L3

ACMSD/HAAO/TDO2/KYNU/KMO/IDO2/IL4I1/IDO1

ORM1/ORM2/REG3A/PTGER3/CD163/IL1A/IL22/AHSG/INS/LBP/HP/TNFRSF11A/SERPINA1/SERPINF2/PTGS2/REG3G/SAA4/F2/MRGPRX1/IL6/A2M/IL1B/TN

GPSM3/STAT3/IL17RA/ZC3H12A/TLR3/MEFV/CLEC7A/EPHB2/CARD9/F2/NOD2/NLRP7/ADCY7/TREM2/IL6/GBP5/ALOX5/LILRB4/CD6/TNF/PLD4/TLR6/ABC

GPSM3/STAT3/IL17RA/ZC3H12A/TLR3/MEFV/CLEC7A/EPHB2/CARD9/F2/NOD2/NLRP7/ADCY7/TREM2/IL6/GBP5/ALOX5/LILRB4/CD6/TNF/PLD4/TLR6/ABC

EGR2/PTN/TENM4/SERPINE2/NKX6-2/OLIG2/TLR2/CLCF1/VEGFC/IL6/TGFB1/LRP2/TNFRSF1B/CYSLTR1/LYN/LTA/RNF112/IL34/BMP2/IL1B/CXCR4/TNF/I

MYB/PDGFRB/ELANE/IGFBP5/IGF1/P2RY6/FOXP1/MAP3K5/SERPINF2/PTGS2/EDN1/CCN4/CALCRL/HMOX1/THBS1/IL10/PTAFR/STAT1/FGFR2/IL6/TGFB

ERAP2/LILRB2/LNPEP/CD209/TAPBPL/LGMN/RELB/FCGR2B/FCGR1A/CLEC4A/THBS1/NOD2/CD1C/CD74/TREM2/FCER1G/RFTN1/MPEG1/CTSS/RAET1

TENM4/CD209/TENM1/CRB2/VCAM1/TENM3/ITGA5/ICAM1/JAML/AMIGO2/CRTAM/SELP/CD6/SELE/ITGAL/SELL

OAS1/TREM2/NFKBIA/NR1H3/LYN/TNIP3/IRAK2/PIK3AP1/PTPN22/LTF/SCIMP/TLR4

BCL6/RUNX1/XCL1/CBLB/SOCS5/ANXA1/SMAD7/ARG2/ZC3H12A/VSIR/ADORA2A/CD274/ASCL2/IL2/IL4R/LGALS9/CD300A/LILRB1/FOXP3/SOCS1/TBX21/J

KSR1/PDGFRB/PAK1/ELANE/NTRK3/TPD52L1/ADRA2A/SERPINB3/PPARG/HYAL2/TNFRSF11A/MAP3K5/TENM1/EDN1/THBS1/DEFB114/PIK3R5/FLT3/WN

XCL1/CCL16/CCL15/CX3CL1/CCL11/CCL23/CCL1/CCL3/CCL2/CCL7/CCL25/CCL14/CCL4/CCL22/CCL24/PTGER4/XCL2/DAPK2/CCL8/ADAM8/CD300A/CCL

STAT3/CEBPA/IL6/JAK2/SPI1/ST18

LBP/C4B/TLR2/TLR1/NOD2/TREM2/TLR9/TLR6/TLR4

CYBA/F2RL1/TLR1/NOD2/TREM2/LYN/TLR6

KCNK3/ATP1A3/KCNK1/KCNK6/NALCN/TREM2/KCNJ10

CAMK2D/KCNE3/KCNE4/ISL1/CTNNA3/CACNB2/CACNA1G/SCN5A/KCNJ3/KCND3/SCN2B/KCNQ1/KCNH6/DSG2/SCN3B/DSC2/SCN4B/KCNJ5/PKP2/KCN

SPP1/TNFRSF11A/TF/CSF1R/SYK/IL6/CLDN18/SFRP1/ADAM8/CD38/S1PR1/TNFSF11/INPP5D/HAMP/TMEM119/DCSTAMP

GHSR/TNN/BMPER/IL10/DLL1/GHRL/SMAD1/ITGA5/KLF4/JCAD/FGF2/FGF1

BATF2/GBP1/TSPAN32/BATF/CLEC7A/GBP4/IL10/IL12B/IL4R/BCL3/CCDC88B/SPN/NKG7/IRF8

IFNK/IFNA8/IFNW1/IFNB1/IFNA4/IFNA1/IFNG/IFNA6/IFNA14/LIF

SSTR4/PLA2G2E/PLA2G3/SLC22A7/P2RX7/AVPR1B/PLA2G4F/IL1A/NTSR1/NMUR2/TNFRSF11A/OC90/PTGS2/PLA2R1/EDN1/SLC22A8/SYK/PLA2G5/SLCO2

GP6/CELA2A/GP9/NOS3/SH2B3/C1QTNF1/SERPINE2/HRG/F2/SYK/THBD/SVEP1/PDPN/PDGFRA/IL6/FCER1G/TXK/JAK2/PEAR1/LYN/SELP/PRKCQ/PLEK/I

LIME1/TRDN/CCL3/APLNR/P2RY6/NTSR1/PLCB2/LACRT/TRPM2/ITPR1/RYR3/JPH3/F2/CASQ1/PTK2B/CACNA1C/FASLG/THY1/CCL21/LYN/SLC8A1/CCL1

PDPN/IL6/JAK2/TNFSF11/ANK3/MMRN1/CCL5

LIME1/TRDN/CCL3/APLNR/P2RY6/NTSR1/PLCB2/LACRT/TRPM2/ITPR1/RYR3/JPH3/F2/CASQ1/PTK2B/CACNA1C/FASLG/THY1/CCL21/LYN/SLC8A1/CCL1

F2RL1/COLEC12/UNC93B1/KCNJ8/TLR3/MEFV/SLC46A2/DHX58/CYLD/HSPA1B/TIFA/IRGM/OAS3/NOD2/DDX60/OAS1/TLR8/TREM2/NFKBIA/SLC15A3/INA

ANGPTL3/TJP1/ADAM12/CTSH/GREM1/CCBE1/CCR3/APELA/TBXA2R/ACVRL1/LRG1/GRN/CCL11/ISL1/ERAP1/PRKD2/CX3CR1/CHRNA7/F3/ANGPT4/MD

LCE3B/DEFB118/RNASE7/DEFA6/RNASE13/RNASE11/CRP/GBP7/LCE3C/SSC5D/GSDMD/ACP5/DEFA4/MYD88/DMBT1/KRT6A/NLRP6/RARRES2/P2RX7/I

ANGPTL3/TJP1/ADAM12/CTSH/GREM1/CCBE1/CCR3/APELA/TBXA2R/ACVRL1/LRG1/GRN/CCL11/ISL1/ERAP1/PRKD2/CX3CR1/CHRNA7/F3/ANGPT4/MD

GP6/CELA2A/PDIA2/GP9/SLC6A4/NOS3/VWF/ADRA2A/SH2B3/MYL9/C1QTNF1/TSPAN32/FERMT3/SERPINE2/HRG/AXL/DGKE/P2RX1/ADRA2B/F2/P2RY12

ARG1/S100A12/USP15/DEFA4/CLEC4D/IL36RN/RARRES2/HTN3/CX3CR1/DEFA5/S100A9/ELANE/IL17RA/HRG/MPO/CLEC7A/CARD9/CLEC4A/DEFB114/PL

BCL6/RUNX1/SOCS5/ANXA1/SMAD7/ZC3H12A/ASCL2/IL2/IL4R/FOXP3/TBX21/JAK3

IFNB1/AXL/APOA1/NOD2/IL10/TGFB1/TGFB3/LILRB4/LILRB1/FOXP3/TBX21/JAK3/TNF/CD96

IL4/PARP14/RUFY4/IL4R/TCF7/CITED1/CD300LF/JAK3/GATA3/CORO1A/IL2RG/DCSTAMP

GPR37L1/PRCP/DRD5/NPPB/ADRB1/ADRB2/GLP1R/NOS3/SCPEP1/NTSR1/KCNK6/ADRA1A/SOD2/VEGFC/PTAFR/ADM2/GCH1/ABAT/TNF/NOS2/ADIPO

RGCC/GPER1/CD248/CCL2/AKR1C3/ITGA4/THBS1/TGFB1/FASLG/CD40/ECSCR/CD40LG

ARG2/TSC22D3/ST3GAL1/SLC46A2/FOXP1/PTCRA/IL2/CD74/PDCD1/DOCK8/BCL3/PRKCQ/JAK3/CD27/BCL11B/CCL5/IL7R/IDO1

SPP1/TNFRSF11A/TF/CSF1R/SYK/IL6/CLDN18/ADAM8/CD38/S1PR1/TNFSF11/INPP5D/HAMP/TMEM119/DCSTAMP

C3AR1/FPR3/CMKLR1/C3/CR1/CR2

IL1RAP/IL17RA/IL4/LGALS9/GATA3/IL33/IL18

CARD18/AHSG/SERPINB5/SERPINB3/FETUB/SERPINB8/SERPINA11/DPEP1/SPINK2/SERPINE2/HRG/SERPINA6/SERPINA1/SERPIND1/SERPINF2/CST4/

PPARG/LDLR/TXNIP/ABCG4/SYK/TREM2/TGFB1/FCER1G/SMPD3/ABCG1/CES1/ITGB2/TLR6/TLR4

FCGR1A/TYROBP/PTAFR/SPI1/ITGB2/BTK/C3/STAP1

FERMT3/HRG/PODXL/PDE3B/P2RY12/SYK/LPXN/CYP1B1/FOXC2/CCL21/LYN/NCKAP1L/CD3E/PIK3CG/PTPN6/CCL5/LIF/CXCL13

SLC46A2/CACNB4/TCF21/MAFB/TNFRSF11A/BATF/CLCF1/FOXL1/IL4/CD274/LIPA/NFKB2/IL10/IL2/IL27RA/PDPN/TGFB1/CDH17/LGALS9/CACNA1C/AICDA

PRDM1/PBX1/PGLYRP2/AXL/ZNF683/MERTK/IL15/RASGRP1/TOX/IL15RA/PTPRC

IL6/CLDN18/CD38/INPP5D/HAMP/TMEM119

SLC46A2/HSPA1B/SLC15A3/SLC15A2/PTPN22/BIRC3/TLR4

FCGR2B/THBS1/NOD2/CD74/CCL21/CCL19/CCR7

APOA1/IFNG/ADM/ABCG4/STAR/STARD4/ABCG1/CES1/TNF/FGF1

SOCS5/ANXA1/ASCL2/IL27/IL4R/CD80/CCL19/JAK3

F2RL1/IGFBP5/NTRK3/IGF1/LIME1/CCL3/RAPGEF3/FAM110C/INS/SH2B3/C1QTNF1/NKX3-1/RRAS/THPO/NTRK2/ARRB2/TEK/SERPINE2/INPP5F/KDR/HY

LGALS9/CCL21/SLAMF8/IL12A/CCR7/CCR6

AGER/CHRNA7/ADRB2/ATP1A3/IGF1/LGMN/ITGA4/FCGR2B/EPHB2/RAMP3/VCAM1/TREM2/ICAM1/MMP12/TNF/TLR6/TLR4/MMP9

PRKCZ/BCL6/SOCS5/ANXA1/MYB/BATF/ASCL2/IL4R/CD86/NLRP3/BCL3/GATA3/IL18

THY1/LILRB4/SLA2/UBASH3A/PTPN22/LAPTM5/PTPN6/SH2D1A

DHX58/ARRB2/USP18/PPARG/NMI/LACRT/IRGM/OAS3/CLEC12B/SERPINB4/PARP14/DEFB114/OAS1/ISG15/TGFB1/LGALS9/KLRC1/SERPING1/SAMHD1/

ACMSD/HAAO/ASMT/TDO2/KYNU/KMO/IDO2/IL4I1/IDO1

CD274/PTCRA/LIPA/IL10/IL2/CD74/LGALS9/ADAM8/WNT5A/FASLG/PDCD1/DOCK8/BCL3/LYN/PRKCQ/IL2RA/GIMAP8/JAK3/BTK/CD27/BCL11B/CCL5/CD3

TLR2/PTGS2/TMEM106A/CLEC7A/IFNG/EDN1/CYP1A1/SOD2/IL10/CYP1B1/GCH1/KLF2/PTK2B/JAK2/KLF4/IL1B/KLRK1/TNF/TLR6/TLR4/CLU/NOS2/HBB

IRF5/TLR8/STAT1/IRF7/MMP12/TLR9/IFIH1/TLR4/TLR7

ARRB2/CLEC12B/SERPINB4/TGFB1/LGALS9/KLRC1/HAVCR2/LILRB1/SERPINB9/KLRD1/CD96

PAK1/ELANE/NTRK3/TPD52L1/ADRA2A/TNFRSF11A/MAP3K5/TENM1/EDN1/THBS1/PIK3R5/FLT3/WNT5A/PDGFC/CD40/PIK3R6/IL34/CCL19/LRRK2/IL1B/

CARD18/AHSG/SERPINB5/SERPINB3/FETUB/SERPINB8/SERPINA11/DPEP1/SPINK2/SERPINE2/HRG/SERPINA6/SERPINA1/SERPIND1/SERPINF2/CST4/

PML/GP1BB/CX3CL1/P2RX2/P2RX7/TRPV6/IL13/DHRS7C/CLIC2/GRIN2B/CYBA/SLC8A3/GPER1/GP9/CACNA1G/GRIN1/LIME1/TRDN/CCL3/APLNR/PPP3R

CD274/LGALS9/ADAM8/WNT5A/PDCD1/CCL5/IDO1

ACMSD/HAAO/HGD/TAT/PAH/TDO2/KYNU/KMO/IDO2/IL4I1/IDO1

CYLD/BATF/IL7/SOCS3/IL12B/IL6/FOXP3/IL12RB1/TBX21/TOX/SPN/LY9/IL23A/SLAMF6

BCR/RIPK3/RIPOR2/AGER/MDK/GCNT1/FUT4/ELANE/ITGA4/CCL25/CHST4/LYVE1/CCR2/IL27RA/PTAFR/PTGER4/FUT7/IL1R1/ADAM8/THY1/ICAM1/CCL

NKX3-1/CCL2/CCDC3/CRHBP/CYLD/FOXP1/HSPA1B/HYAL2/TNFRSF11A/MAP3K5/CCL7/ZFP36/TNFRSF25/ANKRD1/APOA1/TIFA/EDN1/HMHB1/CCL25/

TREM2/IL6/LRRK2/IL1B/PLCG2/TNF/IL33/IL18/STAP1

BATF2/CSF2/RELB/BATF/IL4/BATF3/CAMK4/ITGB8/UBD/DCSTAMP

RAPGEF3/GCM1/TYROBP/TREM2/IL4R/TNFSF14/NFATC2/CD53/CXCL9/DCSTAMP

LILRB2/TAPBPL/FCGR2B/THBS1/NOD2/CD74/TREM2/CCL21/CCL19/WAS/CCR7

AXL/CARD9/NOD2/CD74/TGFB1/WNT5A/TGFB3/LILRB1/SEMA7A/PLCG2/LAPTM5/TLR4/TLR7

AXL/CARD9/NOD2/CD74/TGFB1/WNT5A/TGFB3/LILRB1/SEMA7A/PLCG2/LAPTM5/TLR4/TLR7

LYZL6/LCE3B/DEFB118/RNASE7/DEFA6/DEFB132/GBP7/ARG1/LCE3C/S100A12/SCNN1B/AZU1/C8A/DEFA4/MICB/MYD88/KRT6A/NLRP6/CFHR2/C6/C8B/

PTGS2/CLEC7A/IFNG/EDN1/SOD2/IL10/CYP1B1/GCH1/KLF2/PTK2B/JAK2/KLF4/IL1B/KLRK1/TNF/TLR6/TLR4/CLU/NOS2/HBB

TLR2/PTGS2/TMEM106A/CLEC7A/IFNG/EDN1/CYP1A1/SOD2/IL10/CYP1B1/GCH1/KLF2/PTK2B/JAK2/KLF4/IL1B/KLRK1/TNF/TLR6/TLR4/CLU/NOS2/HBB

TLR2/NOD2/IL10/GBP5/TLR9/CD84/TNF

TLR2/NOD2/IL10/GBP5/TLR9/CD84/TNF

HSPA1B/IRGM/NOD2/NFKBIA/SLC15A3/INAVA/PTPN22/TLR4

PTGER3/CCKBR/SLC6A1/P2RX7/SLC9A4/SLC10A1/CASR/TRPC4/TRH/KCNQ1/SLC51A/NTSR1/CACNB4/UGT1A3/ABCB11/SLC51B/GHRL/SCT/ABAT/SV2

BCR/GRN/S100A9/CYBA/NOXO1/INS/LBP/MPO/NCF2/CLEC7A/LIPA/NCF4/TREM2/HCK/CYBB/CAMK1D/SLAMF8/PIK3CG/RAC2/IGHA2/JCHAIN/CD24/CD

SIPA1/CDKN2A/ZC3H12A/PLXNA2/SH2B3/IL20RB/TNR/C1QTNF1/LILRB2/TACSTD2/TSPAN32/PLXND1/SERPINE2/HRG/PAWR/IFNB1/PODXL/APOA1/VSIR

VNN1/ADAM8/RASGRP1/TESPA1/IL2RG/IL7R

NGFR/ACVRL1/RGCC/CAV2/TNMD/VASH1/CCL2/SULF1/MMRN2/PPARG/APOH/ATOH8/THBS1/GDF2/IL12B/GHRL/STAT1/ALOX5/DLL4/IL12A/TNF/COL4

CX3CL1/S100A9/MDK/ELANE/LBP/PTN/FUT7/ALOX5/ADAM8/SLAMF8/TNF/SELE/CCR6

IL4/IL12B/PTK2B/CD300A/JAK2/CCL5/IL23A

MDK/IL1A/NFKBIZ/STAT3/TLR3/PPARG/HMOX1/CCR2/IL6/TGFB1/ALOX5/TNF/SIGLEC10/PTPN6/TLR4/IL33

KCNJ8/TLR3/MEFV/LBP/TLR5/SLC46A2/DHX58/ARRB2/CYLD/TLR2/HSPA1B/NMI/REG3G/TLR1/TIFA/IRGM/OAS3/CD274/NLRC4/NOD2/DDX60/OAS1/TLR

CFP/CFH/C7/CFB/C3/CR1/CR2

LILRB2/LILRB4/IL2RA/FOXP3/ICOS/CD3E/IDO1

IL6/CLDN18/SFRP1/CD38/INPP5D/HAMP/TMEM119

XCL1/MICB/SLC22A13/IL23R/CD1B/RAET1G/RIPK3/P2RX7/AGER/CD1E/B2M/FCGR2B/CD1C/IL12B/KLRC1/RAET1L/LILRB1/IL12RB1/CD1D/IL12A/KLRD1/N

TICAM2/CYBA/S100A14/F2RL1/LBP/TLR5/TLR2/NMI/TLR1/NOD2/OAS1/TREM2/NFKBIA/NR1H3/LYN/TNIP3/IRAK2/PIK3AP1/PTPN22/LTF/SCIMP/TLR6/TLR

RORA/GBP1/CHI3L1/MYLK3/CCL1/IL1RAP/CCL3/ZC3H12A/CXCL8/NKX3-1/CCL2/HYAL2/TNFRSF11A/CCL7/ANKRD1/EDN1/CCL25/CCL14/CCL4/CCL22/NL

DHX58/ARRB2/USP18/PPARG/NMI/OAS3/CLEC12B/SERPINB4/PARP14/OAS1/ISG15/TGFB1/LGALS9/KLRC1/SERPING1/SAMHD1/NR1H3/NLRC5/HAVCR2

PODXL/P2RY12/SYK/FOXC2/CCL21/NCKAP1L/CD3E/PTPN6/CCL5/LIF/CXCL13

PPARG/PARP9/IFNG/IRGM/PARP14/STAT1/HCK/TXK/NR1H3/NLRC5/JAK2/IRF1

PRDM1/PGLYRP2/AXL/ZNF683/IL15/RASGRP1/TOX/IL15RA

F2RL1/RAB31/TREM2/GATA2/PLCG2/NCKAP1L/C3/STAP1

F2RL1/RAB31/TREM2/GATA2/PLCG2/NCKAP1L/C3/STAP1

CD274/TXNIP/PLK5/IL12B/CD226/TGFB1/HAVCR2/KLF4/SPI1/CRTAM/ABI3/PRF1/IL12A/LAPTM5/IL4I1/NKG7

PRKD2/KCNN4/CYLD/LIPA/CD226/LCK/TRAT1/TESPA1/CCR7/CARD11

IL23R/SMAD7/NFKBID/NFKBIZ/ZC3H12A/BATF/ASCL2/IL2/IL12B/JUNB/FOXP3/IL12RB1/TBX21/IL23A

STAT3/IL2/CITED1/IL2RA/JAK3/IL2RB/IL2RG

UNC93B1/HAVCR2/RSAD2/PIK3AP1/SCIMP/LILRA4/TLR7

IL12B/JAK2/IL15/PTPN22/IL23A/IL18

MEFV/ARRB2/LILRA5/THBS1/NOD2/IL10/TLR8/CMKLR1/LILRB1/TIGIT/JAK3/CCR7

FCER1A/FCGR3A/IL20RB/FCGR2A/FCGR2B/FCGR1A/IGHG1/FUT7/SPN/BTK/C3/CCR7

CD274/IL10/LGALS9/ADAM8/WNT5A/PDCD1/LYN/CCL5/IDO1

NKX3-1/CCL2/CCDC3/CRHBP/CYLD/FOXP1/HSPA1B/HYAL2/TNFRSF11A/NMNAT3/MAP3K5/CCL7/ZFP36/PTGS2/TNFRSF25/ANKRD1/APOA1/TIFA/EDN1

DHX58/CYLD/SIGLEC1/TLR2/NMI/RELB/REL/IRGM/OAS3/TYROBP/IRF5/IKBKE/SYK/IL10/OAS1/TLR8/STAT1/ISG15/OAS2/IRF7/HAVCR2/MMP12/LILRB1/TL

DHX58/CYLD/SIGLEC1/TLR2/NMI/RELB/REL/IRGM/OAS3/TYROBP/IRF5/IKBKE/SYK/IL10/OAS1/TLR8/STAT1/ISG15/OAS2/IRF7/HAVCR2/MMP12/LILRB1/TL

IL20RB/SLC46A2/LMO1/IL2/LGALS9/TNFSF14/SPNS2/LGALS2/IL2RA/FOXP3/JAK3/NCKAP1L/CORO1A/SIT1/GPR174/IL7R/AIM2

VSIR/CD274/LILRB4/LILRB1/CRTAM/SOCS1/IRF1/NCKAP1L

FGFBP1/APLNR/MMRN2/AGTR1/HMOX1/THBS1/IL12B/GATA2/DLL4/IL12A/JCAD/PPP1R16B/FGF2

ADRB2/PDGFRB/PAK1/ELANE/NTRK3/TPD52L1/ADRA2A/PROX1/HMGA2/TNFRSF11A/TCIM/MAP3K5/TENM1/IFNG/IRGM/EDN1/THBS1/PIK3R5/FLT3/WN

CLEC7A/CARD9/CLEC4A/DEFB114/PLA2G5/CLEC4E/CLEC6A/SPI1/PLCG2/CLEC4C

NR1H4/GBP1/IL20RB/ZFP36/LAG3/NOD2/HAVCR2/LILRB4/FOXP3/TBX21/LAPTM5/GATA3/PTPRC/CR1

PTGS2/CLEC7A/IFNG/EDN1/SOD2/IL10/KLF2/PTK2B/JAK2/KLF4/IL1B/KLRK1/TNF/TLR6/TLR4/CLU/HBB

CCDC3/USP18/CYLD/PPARG/HSPA1B/AXL/PARP9/APOA1/IRGM/EDN1/OAS3/CSF1/SLIT2/IL7/RBM47/IKBKE/SYK/PARP14/OAS1/CD74/TREM2/ISG15/IL6/IL

TMEM106A/IFNG/IL4/IL10/JAK2/CIITA/TLR4/IL33

KCNMB2/DRD5/DRD1/PENK/SCN11A/KCNA1/KCND2/SCN2A/KCNMB4/AVP/GPER1/GHSR/PLEC/DMRT3/RGS21/CACNG2/GPR35/CLCN1/KCNJ8/OTOA/[illegible]

WFDC3/IGHA1/DEFA4/DMBT1/SLPI/WFDC11/DEFA5/BPIFA1/PGC/SPRR2A/ELANE/KLK7/DEFB1/RNASE3/TF/FGA/NOD2/IGHG1/IGHG3/RNASE6/FGB/IGH[illegible]

VSIR/IFNG/IL27RA/IL12B/FOXP3/IL12A/NCKAP1L/TLR4

CCL3/FBXW7/ZC3H12A/GPR68/TLR3/GPR55/PGLYRP2/IFNB1/MAFB/IFNL1/IL4/FCGR2B/ASCL2/CEBPA/LAG3/IL2/CD74/C1QC/IL4R/CLDN18/SFRP1/GATA[illegible]

XCL1/MICB/CTSH/SLC22A13/IL23R/CD1B/RAET1G/RIPK3/P2RX7/AGER/CD1E/CTSC/B2M/FCGR2B/CD1C/IL12B/GZMM/KLRC1/RAET1L/LILRB1/PRF1/IL12[illegible]

DOCK8/HAVCR2/CCL21/CCL19/PRF1/DOCK2/CD6/CCR7

IL4/PARP14/IL4R/CD300LF/JAK3/IL2RG

STAT3/BATF/SOCS3/IL12B/IL6/IL12RB1/TBX21/LY9/IL23A/SLAMF6

IFI35/GRN/CX3CR1/LBP/NMI/IFNG/TYROBP/SYK/TREM2/DYSF/HAVCR2/SUCNR1/PLCG2/TNF/IL33

FCGR1A/LTA/CD28/TNF/BTK/C3/CCR7

EOMES/TNFSF8/LILRB4/SOCS1/IRF1/NCKAP1L/TOX/GPR18

FCGR2B/CCR2/SYK/PTAFR/IL4R/PRAM1/LGALS9/CD300A/GATA2/LYN/SPI1/ITGB2/UNC13D/CD84/NCKAP1L/FGR/RAC2

RIPOR2/MDK/CXCL8/LBP/EDN1/C3AR1/CD74/THBS4/DAPK2/CCL21/CAMK1D/CCL19/NCKAP1L/RAC2/CCR7/IL23A

CCL3/FBXW7/GPR68/TLR3/CCR1/GPR55/FOXP1/MAFB/TNFRSF11A/IFNG/IL4/TYROBP/CSF1/IL12B/CD74/TAL1/TREM2/C1QC/CD4/TGFB1/CAMK4/CLDN[illegible]

BATF2/CSF2/TSPAN32/RELB/BATF/IL4/IL10/BATF3/CAMK4/HAVCR2/ITGB8/DOCK2/CD2/UBD/DCSTAMP

HRG/IL21/IL2/IL12B/IL15/IL23A/IL18

CA8/CHP2/PDGFRB/IAPP/IGF1/NMUR1/CCL3/PPP3R2/NMUR2/LACRT/CLEC7A/EDN1/C10orf71/RAMP3/CCL4/SYK/PDGFRA/AKAP5/TREM2/CD4/ZAP70/T[illegible]

USP18/IFNW1/PPARG/IFNB1/IFNA4/PARP9/IFNA1/IFNG/IFNL1/IRGM/OAS3/IFNA6/RBM47/IKBKE/IFNA14/PARP14/OAS1/STAT1/IFITM1/ISG15/HCK/OAS2/T[illegible]

PRR5L/CASS4/CHI3L1/GPER1/PDGFRB/ERFE/F2RL1/IGFBP5/IGF1/CCL3/FAM110C/INS/C1QTNF1/NKX3-1/THPO/NTRK2/ARRB2/TEK/KDR/AKR1C3/AXL/[illegible]

RIPOR2/CFLAR/RAPGEF3/PLEKHO1/GCM1/TYROBP/TREM2/IL4R/TNFSF14/NFATC2/CD53/CXCL9/CXCL10/DCSTAMP

PGLYRP2/IFNB1/IFNL1/ASCL2/LAG3/IL2/CD74/IL4R/SFRP1/DTX1/TMEM131L/LILRB4/FOXP3/SOCS1/IRF1/TBX21/JAK3/INHBA/CR1

IFNL1/IL4/F2/IL10/CYP1B1/IL12B/OCIAD2/IL6/PTK2B/CD300A/JAK2/IL10RA/TNF/CCL5/IL23A/IL7R

CD274/PTCRA/LIPA/LGALS9/ADAM8/WNT5A/FASLG/PDCD1/DOCK8/BCL3/PRKCQ/IL2RA/GIMAP8/JAK3/CD27/BCL11B/CCL5/IL7R/IDO1

TNFRSF11A/PDPN/TGFB1/IL15/SPNS2/LTA/TOX/IL7R/LTB/CXCR5

LILRB2/CD274/PDCD1/HAVCR2/LILRB4/IL2RA/FOXP3/CD3E/IDO1

C4BPA/C4BPB/SVEP1/SERPING1/A2M/CR1L/CR1/CR2

IL20RB/FCGR2B/FCGR1A/FUT7/SPN/BTK/C3/PLA2G2D/CCR7

PTGS2/TMEM106A/CLEC7A/IFNG/EDN1/SOD2/IL10/KLF2/PTK2B/JAK2/KLF4/IL1B/KLRK1/TNF/TLR6/TLR4/CLU/HBB

RIPOR2/MDK/S100A14/CXCL8/LBP/EDN1/C3AR1/CD74/THBS4/DAPK2/CCL21/CAMK1D/CCL19/NCKAP1L/RAC2/CCR7/IL23A

CCL25/BMP5/SLIT2/HMOX1/CD200R1/IL27RA/PTGER4/GCSAM/CD300A/CCL21/PADI2/SLAMF8/KLRK1/GPR18/IL33/STAP1

CCL2/KDR/TLR2/IFNG/EDN1/NOD2/STAT1/JAK2/LRRK2/TLR4/IL33/FCER2

CCL2/KDR/TLR2/IFNG/EDN1/NOD2/STAT1/JAK2/LRRK2/TLR4/IL33/FCER2

CD274/IL10/LGALS9/ADAM8/WNT5A/PDCD1/CCL5/IDO1

IFNL1/IL4/F2/IL10/CYP1B1/IL12B/OCIAD2/IL6/TGFB1/PTK2B/CD300A/JAK2/IL10RA/TNF/CCL5/IL23A/IL7R

RIPOR2/MDK/IL1A/FUT4/CXCL8/LBP/EDN1/SLIT2/C3AR1/NOD2/CD74/THBS4/FUT7/IL1R1/DAPK2/ADAM8/CCL21/CAMK1D/CCL19/SLAMF8/NCKAP1L/RA[illegible]

BATF/SOCS3/IL12B/IL6/IL12RB1/TBX21/SPN/LY9/IL23A/SLAMF6

CYLD/BATF/SOCS3/IL12B/IL6/FOXP3/IL12RB1/TBX21/TOX/SPN/LY9/IL23A/SLAMF6

TICAM2/CYBA/S100A14/F2RL1/LBP/TLR5/TLR2/NMI/TLR1/NOD2/OAS1/TREM2/NFKBIA/NR1H3/LYN/TNIP3/IRAK2/PIK3AP1/PTPN22/LTF/SCIMP/TLR6/TLR[illegible]

SKAP1/FERMT3/PODXL/ITGA4/PLPP3/ITGA5/CD3E/CCL5/CXCL13

RORA/GBP1/MYLK3/CCL1/IL1RAP/CCL3/ZC3H12A/CXCL8/NKX3-1/CCL2/HYAL2/CCL7/ANKRD1/EDN1/CCL25/CCL14/CCL4/CCL22/NLRP7/CCL24/NFKBIA/[illegible]

VSIR/IFNG/LAG3/IL2/LGALS9/LILRB4/CD28/FOXP3/SOCS1/IRF1/IL4I1/IL2RG/CR1

GPER1/CD248/CCL2/AKR1C3/PLA2R1/ITGA4/HMOX1/THBS1/IL6/TGFB1/FASLG/JAK2/CD40/ECSCR/CD40LG/SFRP4

LYZL6/LCE3B/DEFB118/RNASE7/DEFA6/DEFB132/GBP7/ARG1/LCE3C/S100A12/SCNN1B/AZU1/C8A/DEFA4/MICB/MYD88/KRT6A/NLRP6/CFHR2/C6/C8B/[illegible]

LYZL6/LCE3B/DEFB118/RNASE7/DEFA6/DEFB132/GBP7/ARG1/LCE3C/S100A12/SCNN1B/AZU1/C8A/DEFA4/MICB/MYD88/KRT6A/NLRP6/CFHR2/C6/C8B/[illegible]

FCGR2B/FCGR1A/SYK/HCK/CLEC4E/FCER1G/LYN/MYO1G/LCK/CD247/FGR/VAV1

CSF1/C1QC/TGFB1/GATA2/IL34/HCLS1/INHBA/LIF/ADIPOQ

LILRB2/VSIR/IFNG/IL2/LGALS9/LILRB4/FOXP3/SOCS1/IL4I1/IL2RG/CR1

SYK/FCER1G/LYN/PRKCQ/PLCG2/BTK/LILRA4/VAV1/FCER2

SYK/IL4R/LGALS9/CD300A/GATA2/MILR1/PTGDR/LYN/UNC13D/CD84/PTGDS/RASGRP1/CLNK/BTK/PIK3CG/FGR/RAC2/LAT2

CSF2/OSCAR/IAPP/RASSF2/L3MBTL3/ERFE/F2RL1/CCL3/FBXW7/GPR68/TLR3/TSPAN2/CCR1/GPR55/CEBPE/PPARG/TLR2/FOXP1/MAFB/TNFRSF11A/R

TRIM21/PRKCH/RIPK3/TRIM38/PRKD2/TICAM2/AGER/S100A9/CFLAR/EDA/RFPL1/IL1RAP/STAT3/INS/TRIM34/TLR2/HSPA1B/TNFRSF11A/CLEC7A/TRIM

CA8/CHP2/GBP1/PDGFRB/IAPP/IGF1/NMUR1/CCL3/PPP3R2/NMUR2/RIT2/DYRK2/LACRT/ITPR1/MYOZ1/CLEC7A/EDN1/C10orf71/RAMP3/CCL4/SYK/PDG

STAT3/IL17RA/ZC3H12A/INS/TLR3/MEFV/LBP/P2RX1/CLEC7A/EPHB2/CARD9/F2/LIPA/SYK/NOD2/NLRP7/ADCY7/TREM2/IL6/IL4R/GBP5/ALOX5/LYN/LILRB

VSIR/CD274/IL2/LGALS9/CD28/PRKCQ/IL2RA/FOXP3/CD3E/CARD11

VSIR/CD274/IL2/LGALS9/CD28/PRKCQ/IL2RA/FOXP3/CD3E/CARD11

IFNK/ZC3H12A/KCNJ8/TLR3/TRIM34/CD207/IFNA8/TSPAN32/DHX58/USP18/IFNW1/IRF9/IFNB1/TLR2/IL21/HYAL2/IFNA4/IRF2/PARP9/IFNA1/MAP3K14/IFN

BCL6/RUNX1/XCL1/CBLB/SOCS5/ANXA1/SMAD7/ARG2/ZC3H12A/VSIR/CD274/ASCL2/IL2/IL4R/LGALS9/FOXP3/TBX21/JAK3

CDHR2/DACH1/PGLYRP4/PAK1/LBP/C4B/PGLYRP2/TLR2/TLR1/CLEC7A/NLRC4/NOD2/TREM2/NLRP3/CLEC6A/CRTAM/TLR9/CD1D/TLR6/TLR4

SCNN1B/AZU1/TTBK1/IFNGR1/TAFA3/ITGAM/MYD88/CX3CL1/GRN/IL13/CX3CR1/AGER/CTSC/CCL3/KCNJ8/TLR3/LDLR/TLR2/IFNG/TYROBP/C1QA/TRE

AGER/CHRNA7/GAREM1/TP73/OR2AT4/IL1A/CFLAR/CHI3L1/KSR1/ADRB2/GPER1/PDGFRB/IAPP/TNIK/RASSF2/CAV2/ADRA1B/CASR/PAK1/NRG1/CCL1/

TNFSF13B/NFKB2/BCL3/UNC13D/KLHL6/CXCL13

CLEC7A/OAS3/SYK/NOD2/OAS1/LGALS9/IL1B/CD84/ADIPOQ

GPR34/P2RY11/P2RY6/P2RY2/P2RY12/P2RY14/PTAFR/GPR171/P2RY13/P2RY8

BATF/SOCS3/IL12B/IL6/FOXP3/IL12RB1/TBX21/TOX/SPN/LY9/IL23A/SLAMF6

OAS1/TLR8/RFTN1/HAVCR2/RSAD2/TLR9/PIK3AP1/PTPN22/TNF/SCIMP/LILRA4/TLR7/FCRL3

FCGR2B/ZPBP2/FOXP3/CR1/IL33/FCRL3/CD22

TMEM106A/IFNG/IL4/IL10/JAK2/CIITA/TLR4/IL33

PAWR/FCGR2B/TYROBP/IL10/MNDA/CD300A/LYN/BTK/INPP5D/TNFRSF13B

RELB/ASCL2/SEMA4A/IL27/IL4R/STAT4/CD80/CCL19/TBX21/JAK3/SPN/IL18R1

CYBA/F2RL1/TLR2/TLR1/NOD2/TREM2/LYN/PIK3AP1/TLR6

DEFA4/MYD88/CLEC4D/IL36RN/PTX3/RARRES2/HTN3/IL25/CX3CR1/DEFA5/S100A9/ELANE/IL17RA/HRG/MPO/CLEC7A/CARD9/CLEC4A/SYK/DEFB114/PL

RIPOR2/MDK/IL1A/FUT4/CXCL8/LBP/EDN1/C3AR1/CD74/THBS4/FUT7/IL1R1/DAPK2/ADAM8/CCL21/CAMK1D/CCL19/NCKAP1L/RAC2/CCR7/IL23A

IL13/SPACA3/CTSC/CCL3/LBP/IRGM/CEBPA/THBS1/IL10/TREM2/IL4R/WNT5A/HAVCR2/IL1RL1/LRRK2/TLR6/TLR4/IL33/HAMP/STAP1

NT5E/KRT1/ELANE/KIAA0319/IGF1/NOS3/STAT3/IL22RA1/INS/FIGNL2/MEFV/SH2B3/IL20RB/TNR/C1QTNF1/CCL2/ELF4/DHX58/ARRB2/USP18/CYLD/TEK/F

CCDC3/USP18/CYLD/PPARG/TLR2/HSPA1B/AXL/PARP9/APOA1/IRGM/EDN1/OAS3/CSF1/SLIT2/IL7/RBM47/IKBKE/SYK/PARP14/OAS1/CD74/TREM2/ISG15

DOCK11/TNFSF13B/CD74/SPNS2/LYN/GAPT/FOXP3/NCKAP1L/SASH3/IL7R/TNFRSF13B

SSTR4/CCL15/FGF10/MT3/NPY5R/APELA/GLIPR2/ERBB4/FGF19/CX3CL1/CCL11/FSHR/CCL23/PRKD2/MOS/AGER/CHRNA7/GAREM1/OR2AT4/IL1A/CFLA

STAT3/CD4/IL15/JAK3/IL15RA/IL2RB/IL2RG

STAT3/CD4/IL15/JAK3/IL15RA/IL2RB/IL2RG

CXCL8/LBP/EDN1/SLIT2/C3AR1/NOD2/CD74/THBS4/DAPK2/CCL21/CAMK1D/CCL19/NCKAP1L/RAC2/CCR7/IL23A

ELANE/F2RL1/IL17RA/TLR3/MEFV/LBP/TLR2/FOXP1/CLEC7A/IFNG/OAS3/CARD9/IL7/HMOX1/CSF1R/SYK/NOD2/IL10/OAS1/CD74/TREM2/IL6/TGFB1/IL4R/

FCGR2B/CLEC4A/THBS1/NOD2/CD74/MPEG1/CCL21/CCL19/CCR7

CCL2/C4B/AXL/XKR5/C2/XKR4/TYROBP/CCR2/THBS1/TGM2/TREM2/MERTK/NR1H3/PEAR1/CD300LF/TIMD4/MARCO/C3/ALOX15

DOK1/TLR2/CSF1/CSF1R/PDE1B/TREM2/PDE2A/TLR4/STAP1/DCSTAMP

CARD9/NOD2/CD74/WNT5A/LILRB1/SEMA7A/PLCG2/LAPTM5/TLR4/TLR7

CLEC7A/ARID5A/CARD9/IL12B/NLRP10/JAK2/CLEC6A/PRKCQ/IL12RB1/IL23A

SYK/IL4R/LGALS9/CD300A/GATA2/MILR1/PTGDR/LYN/UNC13D/CD84/SERPINB9/PTGDS/RASGRP1/CLNK/BTK/PIK3CG/FGR/RAC2/LAT2

PPBP/LGMN/TNFRSF11A/CCL7/CSF3R/EDN1/CCL25/CSF1/CCL14/SLIT2/C3AR1/CCL4/CCR2/P2RY12/THBS1/CSF1R/SYK/NOD2/CCL22/CCL24/CD200R1/

TLR5/SLC46A2/DHX58/ARRB2/CYLD/TLR2/HSPA1B/NMI/REG3G/TLR1/CLEC7A/TIFA/IRGM/OAS3/CD274/TYROBP/NLRC4/SYK/NOD2/DDX60/OAS1/TLR8/

FOXP1/ST6GAL1/AXL/CD274/PTCRA/LIPA/NOD2/IL10/IL2/CD74/IL6/MERTK/LGALS9/ADAM8/WNT5A/FASLG/PDCD1/DOCK8/IRF7/BCL3/CCL21/LYN/LILRB1

MDK/GCNT1/FUT4/ELANE/ITGA4/CHST4/CCR2/PTAFR/IL6/FUT7/ALOX5/SELP/ITGB2/TNF/SELE

ARG2/TSC22D3/ST3GAL1/SLC46A2/PTCRA/DOCK8/BCL3/PRKCQ/JAK3/CD27/BCL11B/CCL5/IL7R/IDO1

NFKBIA/IL6/IL1R1/IRAK2/IL1B/ZBP1/IL1RN/ST18

LBP/TLR5/SLC46A2/DHX58/ARRB2/CYLD/TLR2/HSPA1B/NMI/REG3G/TLR1/CLEC7A/TIFA/IRGM/OAS3/CD274/TYROBP/NLRC4/SYK/NOD2/DDX60/OAS1/TI

STAT3/CD4/IL15/JAK3/IL15RA/IL2RB/IL2RG

CCL2/C4B/ARRB2/RAB31/FCGR2A/CFP/SYNE1/AXL/APOA5/C2/APOA1/CLEC7A/IFNG/IL4/FCGR2B/FCGR1A/TF/SYK/NOD2/DLL1/TREM2/PLA2G5/FCER1G

CX3CL1/GRN/IL13/CX3CR1/AGER/CTSC/CCL3/KCNJ8/TLR3/LDLR/TLR2/IFNG/TYROBP/C1QA/TREM2/IL6/AIF1/JAK2/CST7/TLR9/ITGB2/LRRK2/TNF/TLR6

NFKBIZ/UNC93B1/TLR3/TLR5/ARRB2/TLR2/REG3G/TLR1/CD274/TLR8/CD300A/CTSS/IRF7/TNIP3/IRAK2/TLR9/CD300LF/IRF1/PLCG2/BTK/SCIMP/TLR6/BI

SYK/IL4R/LGALS9/CD300A/GATA2/MILR1/PTGDR/LYN/UNC13D/CD84/PTGDS/RASGRP1/CLNK/BTK/PIK3CG/FGR/RAC2/LAT2/PTPN6

KLRF2/IFNK/IFNA8/PGLYRP2/IFNW1/IFNB1/IFNA4/IFNA1/IFNA6/IFNA14/CD244/IL12B/ZNF683/UNC13D/CORO1A/NKG7

IFNK/SH2B3/IFNA8/IFNW1/IFNB1/INPP5F/IFNA4/IL9/IFNA1/IFNG/IFNL1/IL4/EPHB2/IFNA6/F2/IFNA14/IL10/CYP1B1/PARP14/SOCS3/IL12B/OCIAD2/IL6/PTK2B

PGLYRP2/TLR2/TLR1/CLEC7A/NLRC4/NOD2/CLEC6A/CD1D/TLR6/TLR4

PPARG/HRG/IL21/SPP1/TNFRSF11A/TF/CSF1R/SYK/IL2/IL12B/THBS4/IL6/TGFB1/CLDN18/SFRP1/ADAM8/IL15/CD38/S1PR1/TNFSF11/FLT4/INPP5D/IL23A/I

IL23R/SOCS5/ANXA1/NFKBID/NFKBIZ/IL12B/IL4R/CD86/NLRP3/CD80/CCL19/IL12RB1/IL23A/IL18

CHRNA7/NR1H4/ARG2/GHSR/BPI/IGF1/ZC3H12A/LBP/CIDEA/ELF4/ARRB2/AXL/VSIR/IL4/CLEC4A/NOD2/IL10/IL27RA/DEFB114/GHRL/TREM2/LGALS9/BCL

MDK/GCNT1/FUT4/ELANE/ITGA4/CCL25/CHST4/CCR2/PTAFR/IL6/FUT7/ALOX5/CCL21/KLF4/SELP/ITGB2/TNF/SELE

CX3CL1/GRN/IL13/CX3CR1/AGER/CTSC/CCL3/KCNJ8/TLR3/LDLR/TLR2/IFNG/ADORA2A/TYROBP/C1QA/TREM2/IL6/AIF1/JAK2/CST7/TLR9/ITGB2/LRRK2

ELANE/F2RL1/IL17RA/TLR3/MEFV/LBP/TLR2/FOXP1/CLEC7A/IFNG/OAS3/CARD9/IL7/HMOX1/CSF1R/SYK/NOD2/IL10/OAS1/CD74/TREM2/IL6/TGFB1/IL4R/

IFNK/IL17RA/ZC3H12A/KCNJ8/DUOX2/TLR3/TRIM34/CD207/IFNA8/TSPAN32/HMGA2/DHX58/USP18/IFNW1/IRF9/IFNB1/TLR2/IL21/NMI/HYAL2/IFI44/IFNA4

LRRC15/RESF1/PROX1/CH25H/HMGA2/IFNB1/ZFP36/OAS3/TRIM14/TRIM59/APOBEC3D/APOBEC3G/OAS1/APOBEC3H/STAT1/IFITM1/ISG15/IFIT5/OAS2/

CD33/PML/CX3CL1/EGR1/P2RX7/ISL1/CX3CR1/AGER/ARG2/GHSR/F2RL1/IGF1/CCL3/STAT3/CARD18/ZC3H12A/PYDC1/MEFV/ELF4/ARRB2/FOXP1/TMEM

CD33/PML/CX3CL1/EGR1/P2RX7/ISL1/CX3CR1/AGER/ARG2/GHSR/F2RL1/IGF1/CCL3/STAT3/CARD18/ZC3H12A/PYDC1/MEFV/ELF4/ARRB2/FOXP1/TMEM

IFNB1/ARID5A/IL4/IL6/IL1R1/NLRP3/RSAD2/IL1B/TBX21/GATA3/IL18R1/IL18

NMI/IRF5/IL10/TLR8/STAT1/IRF7/HAVCR2/MMP12/TLR9/IFIH1/LILRA4/TLR4/NLRC3/TLR7

NMI/IRF5/IL10/TLR8/STAT1/IRF7/HAVCR2/MMP12/TLR9/IFIH1/LILRA4/TLR4/NLRC3/TLR7

BATF/SOCS3/IL12B/IL6/FOXP3/IL12RB1/TBX21/TOX/SPN/LY9/IL23A/SLAMF6

ARRB2/CLEC12B/FCGR2B/SERPINB4/TGFB1/LGALS9/KLRC1/HAVCR2/LILRB1/SERPINB9/KLRD1/NCKAP1L/PTPRC/IL7R

EGR1/P2RX7/ISL1/AGER/F2RL1/CCL3/STAT3/PYDC1/MEFV/TMEM106A/CLEC7A/IFNG/LILRA5/TYROBP/NLRC4/NOD2/TLR8/IL6/INAVA/NLRP2/GBP5/LGAL

SLC46A2/SKIL/LMO1/DOCK11/TNFSF13B/IL2/CD74/LGALS9/TNFRSF17/TNFSF14/SPNS2/LYN/GAPT/LGALS2/IL2RA/FOXP3/JAK3/NCKAP1L/CORO1A/SIT1/

CCL2/KDR/TLR2/IFNG/NOD2/STAT1/JAK2/LRRK2/TLR4/IL33/FCER2

S100A12/FPR2/CXCL12/XCL1/CCL16/GREM1/CCL15/ANXA1/DEFB124/CX3CL1/CCL11/CCL23/CX3CR1/S100A14/CCL1/CCL3/CCR1/CCL2/LGMN/TNFRSF1

TMEM106A/IFNG/IL4/IL10/JAK2/CIITA/TLR4/IL33

CD274/PTCRA/IL10/IL2/CD74/LGALS9/ADAM8/WNT5A/PDCD1/DOCK8/BCL3/LYN/PRKCQ/GIMAP8/JAK3/BTK/CD27/BCL11B/CCL5/CD3G/IL7R/IDO1

C4BPA/FCGR2B/C4BPB/SVEP1/CR1L/PTPN6/CR1/CR2

FCGR2B/FCGR1A/TYROBP/CARD9/F2/CCR2/SYK/IGHG1/PTAFR/IL6/IL4R/PRAM1/LGALS9/CD300A/GATA2/MILR1/PTGDR/LYN/SPI1/ITGB2/UNC13D/CD8

REG3G/CD300A/IRF7/IRAK2/TLR9/CD300LF/IRF1/BTK/TLR6/TLR4

HSPA1B/AXL/PARP9/IRGM/EDN1/CSF1/IL7/RBM47/IKBKE/PARP14/CD74/TREM2/IL1R1/TXK/WNT5A/NLRC5/IRF7/MMP12/CD300LF/CXCR4/ZBP1/LAPTM5

ALPL/DOK1/TLR2/SPP1/CSF1/CSF1R/PDE1B/TREM2/PDE2A/TNF/TLR4/STAP1/DCSTAMP

F2RL1/RAB31/TREM2/CD300A/GATA2/PLCG2/NCKAP1L/C3/ALOX15/STAP1

F2RL1/RAB31/TREM2/CD300A/GATA2/PLCG2/NCKAP1L/C3/ALOX15/STAP1

CX3CR1/GCNT1/FUT4/ELANE/ADD2/ITGA4/CCL25/CHST4/VCAM1/CCR2/PTAFR/JAM2/SELPLG/FUT7/CCL21/SELP/ITGB7/TNF/SPN/SELE/SELL

CD83/CLEC7A/CCR2/CD4/CD86/CD80/CD28/PRKCQ/IL1B/PLCG2/CD3E/PTPRC/SASH3/CARD11

CCL3/FBXW7/ZC3H12A/GPR68/TLR3/GPR55/PGLYRP2/IFNB1/MAFB/IFNL1/IL4/FCGR2B/ASCL2/LAG3/IL2/CD74/C1QC/IL4R/CLDN18/SFRP1/GATA2/DTX1/

XKR5/XKR4/FCGR2B/FCGR1A/FCHO2/THBS1/TREM2/AIF1/CD300A/GATA2/ITGB2/BIN2/ELMO1/PLCG2/NCKAP1L/TIMD4/MARCO/C3/ARHGAP25/HAVCR

IL21/CLCF1/IL4/EPHB2/IL7/TNFSF13B/IL2/CD74/TNFRSF4/GPR183/TLR9/CD38/NFATC2/NCKAP1L/BTK/PTPRC/TLR4/SASH3/CARD11/TNFRSF13C/FCRL3
FCGR2B/CCR2/SYK/PTAFR/HCK/IL4R/PRAM1/LGALS9/CD300A/GATA2/MILR1/PTGDR/LYN/SPI1/ITGB2/UNC13D/CD84/PTGDS/RASGRP1/CLNK/NCKAP1
AXL/TYROBP/IL12B/JAK2/IL15/IL12A/RASGRP1/TOX/IL15RA/IL23A/IL18
CLEC7A/TYROBP/SYK/KLRC1/CLEC6A/LYN/KLRC4/KLRD1/PLCG2/KLRK1
CLEC7A/TYROBP/SYK/KLRC1/CLEC6A/LYN/KLRC4/KLRD1/PLCG2/KLRK1
CLEC7A/TYROBP/SYK/KLRC1/CLEC6A/LYN/KLRC4/KLRD1/PLCG2/KLRK1
IL20RB/IFNB1/FCGR2B/NOD2/KLRC1/CD80/LILRB4/LILRB1/FOXP3/TBX21/KLRD1/NCKAP1L/SPN/IL4I1/PTPRC/IL7R
ELF4/IL12B/CD300A/JAK2/IL15/IL12A/RASAL3/IL23A/IL18
CD300A/MILR1/CD300LF/CD84/PTPN6/CNR2
FERMT3/HRG/ITGAD/FYB2/PODXL/ITGA8/ITGA4/PDE3B/PLPP3/P2RY12/SYK/LPXN/CYP1B1/ITGAX/FOXC2/ITGA5/ICAM1/CCL21/LYN/ITGBL1/ITGB2/ITGB7
CARD9/IRF5/NOD2/NFKBIA/INAVA/NLRP1/PTPN22/LDOC1
IGF1/CCL3/CD200R1L/LDLR/PTGS2/IL4/CD200R1/TREM2/IL6/TNFRSF1B/CST7/LRRK2/IL1B/PLCG2/TNF/PTPRC/IL33/MMP9/IL18/STAP1
ARID5A/IL4/IL6/IL1R1/NLRP3/RSAD2/IL1B/TBX21/GATA3/IL18R1/SASH3/IL18
SH2B3/IFNA8/CRLF2/IFNW1/PPARG/IFNB1/INPP5F/IFNA4/IL9/IFNA1/IFNG/IFNL1/IL4/CLEC12B/EPHB2/IFNA6/F2/IFNA14/IL10/CYP1B1/PARP14/SOCS3/IL12
CHRNA7/NR1H4/ARG2/GHSR/BPI/IGF1/ZC3H12A/LBP/CIDEA/ELF4/ARRB2/AXL/VSIR/IL4/CD274/CLEC4A/NOD2/IL10/IL27RA/DEFB114/GHRL/TREM2/LGAL
EDN1/CSF1/SLIT2/C3AR1/THBS1/CSF1R/NOD2/CD74/THBS4/CMKLR1/DAPK2/CCL21/IL34/CAMK1D/CCL19/NCKAP1L/RAC2/CCR7/CCL5/IL23A
BCR/ANXA1/FCER1A/GRN/F2RL1/PLPP6/CCL3/CXCL8/FCGR2B/TYROBP/CCR2/SYK/PTAFR/IL18RAP/FCER1G/PRAM1/CD300A/IL15/SPI1/ITGB2/DNASE1
PRKD2/DEFA5/F3/CHI3L1/ELANE/F2RL1/STAT3/TLR3/LBP/TLR5/TLR2/HSPA1B/HYAL2/TLR1/CLEC7A/SYK/NOD2/GDF2/CD244/TLR8/CD74/IL6/NLRP10/LGA
IL12B/CD300A/JAK2/IL12A/RASAL3/IL23A/IL18
CD74/ZAP70/DOCK2/TOX/CD3E/CD3D/PTPRC/CD3G
CD74/CD28/ZAP70/DOCK2/CD3E/SPN/PTPRC/CCR7
CCR2/XCL2/WNT5A/TNFSF14/CCL21/CCL5/CXCL13
PRKCH/PRKD2/KCNN4/PRKCB/CYLD/FOXP1/NFAM1/LIPA/CD226/LCK/TRAT1/TESPA1/PTPRC/CCR7/CARD11/STAP1
LBP/C4B/PGLYRP2/TLR2/TLR1/CLEC7A/NLRC4/NOD2/TREM2/CLEC6A/TLR9/CD1D/TLR6/TLR4
SLC46A2/CYLD/HSPA1B/IRGM/NOD2/NFKBIA/SLC15A3/INAVA/SLC15A2/PTPN22/BIRC3/TLR4
ARID5A/CCR2/IL27RA/IL12B/NLRP10/IL1R1/IL1B/IL12RB1/TBX21/IL18R1/IL23A/IL18
CCL3/CCL7/CCL4/CCR2/XCL2/PTK2B/WNT5A/TNFSF14/CCL21/CCL5/CXCL13
CD33/ANXA1/PML/CX3CL1/EGR1/P2RX7/ISL1/CX3CR1/AGER/NR1H4/ARG2/GHSR/F2RL1/IGF1/CCL3/IL16/STAT3/CARD18/ZC3H12A/PYDC1/MEFV/PANX3
CD33/ANXA1/PML/CX3CL1/EGR1/P2RX7/ISL1/CX3CR1/AGER/NR1H4/ARG2/GHSR/F2RL1/IGF1/CCL3/IL16/STAT3/CARD18/ZC3H12A/PYDC1/MEFV/PANX3
IL13/SPACA3/RORA/CTSC/BPI/CCL3/ZC3H12A/LBP/LDLR/IRGM/FCGR2B/CEBPA/THBS1/IL10/CD74/TREM2/IL6/IL4R/WNT5A/IL31RA/NR1H3/CST7/HAVCR
BCR/CX3CL1/RIPK3/ITGA1/CX3CR1/RIPOR2/AGER/MDK/GCNT1/FUT4/ELANE/ADD2/CCL2/ITGA4/CCL25/CHST4/LYVE1/VCAM1/CCR2/IL27RA/PTAFR/JA
VNN1/CAMK4/ADAM8/TMEM131L/RASGRP1/TOX/TESPA1/IL2RG/IL7R
CD33/LRRC19/ANXA1/PRKD2/DEFA5/F3/CHI3L1/BPI/ELANE/F2RL1/STAT3/TLR3/LBP/TLR5/TLR2/HSPA1B/HYAL2/TLR1/CLEC7A/SYK/NOD2/GDF2/CD244/I
C4BPA/FCGR2B/C4BPB/SVEP1/FOXP3/CR1L/PTPN6/CR1/CR2
C4BPA/FCGR2B/C4BPB/SVEP1/FOXP3/CR1L/PTPN6/CR1/CR2
RIPK3/P2RX7/PRKD2/ARG2/TSC22D3/ST3GAL1/SLC46A2/CD274/PTCRA/LGALS9/ADAM8/WNT5A/PDCD1/DOCK8/BCL3/PRKCQ/GIMAP8/JAK3/CD27/BCL1
PDGFRB/ROBO3/S100A14/CASR/GPSM3/CCL1/ELANE/F2RL1/NTRK3/CCL3/IL16/CXCL8/CCR1/SMOC2/LBP/PTN/CCL2/HRG/KDR/LGMN/ST6GAL1/CCL7/S
NLRP6/NPY5R/FCER1A/FCGR3A/ELANE/IL20RB/FCGR2A/FCGR2B/FCGR1A/IGHG1/FUT7/IL31RA/A2M/CD6/SPN/BTK/C3/PLA2G2D/CCR7
VSIR/FCGR2B/CD274/TYROBP/IL12B/DLL1/PDCD1LG2/LILRB4/LILRB1/FOXP3/JAK3/BTK/IL23A/IDO1
IL21/CLCF1/MZB1/IL4/EPHB2/IL10/IL2/IL6/TGFB1/IL4R/CD86/TNFRSF4/TLR9/CD28/TBX21/BTK/RBP4/PTPRC/SASH3/IL33
SMAD7/NFKBID/RORA/NFKBIZ/STAT3/ZC3H12A/RORC/BATF/ASCL2/IL2/SOCS3/IL12B/JUNB/IL6/FOXP3/IL12RB1/TBX21/LY9/IL23A/SLAMF6
PF4V1/DEFA5/BPIFA1/PGLYRP4/S100A9/PGC/MUC7/ELANE/KLK7/DEFB1/CXCL8/LEAP2/RNASE3/HRG/PPBP/REG3G/DEFB127/F2/NOD2/RNASE6/CXCL2/
BPI/ELANE/F2RL1/STAT3/TLR3/LBP/TLR5/TLR2/HSPA1B/HYAL2/TLR1/CLEC7A/SYK/NOD2/GDF2/CD244/IL10/TLR8/CD74/IL6/NLRP10/LGALS9/WNT5A/BCL
XKR5/XKR4/FCGR2B/FCGR1A/THBS1/TREM2/AIF1/CD300A/GATA2/ITGB2/BIN2/ELMO1/PLCG2/NCKAP1L/TIMD4/MARCO/C3/ARHGAP25/HAVCR1/ALOX1
FCGR2B/TYROBP/SYK/PTAFR/IL18RAP/FCER1G/PRAM1/CD300A/IL15/SPI1/ITGB2/DNASE1L3/TNF/CCL5/PLA2G2A/IL18

ANXA1/RARRES2/CX3CL1/CCL11/CCL23/ITGA1/PF4V1/RIPOR2/S100A9/MDK/IL1A/GP2/S100A14/FUT4/CCL1/CCL3/IL17RA/CXCL8/LBP/CCL2/DPEP1/PPBP/

ANXA1/RARRES2/CX3CL1/CCL11/CCL23/ITGA1/PF4V1/RIPOR2/S100A9/MDK/S100A14/CCL1/CCL3/IL17RA/CXCL8/LBP/CCL2/DPEP1/PPBP/CCL7/CSF3R/E

MMP26/MDK/NR1H4/RORA/GPER1/IL22/ARG2/GHSR/NLRP4/NT5E/KRT1/ELANE/IGF1/STAT3/IL22RA1/INS/MEFV/IL20RB/ELF4/DHX58/ARRB2/USP18/CYL

PPARG/TLR2/CCL7/PARP9/IFNG/IRGM/EDN1/CCL25/CCL14/GBP4/AQP4/CCL4/CCL22/CCL24/PARP14/IL12B/STAT1/HCK/GBP5/TXK/XCL2/LGALS9/AIF1/C

CLCF1/IL4/FCGR1A/NOD2/IL2/CD226/TREM2/TGFB1/LTA/CD28/TBX21/TNF/BTK/C3/PTPRC/FCER2

CLCF1/IL4/FCGR1A/NOD2/IL2/CD226/TREM2/TGFB1/LTA/CD28/TBX21/TNF/BTK/C3/PTPRC/FCER2

IL4/CLEC12B/FCGR2B/SERPINB4/TGFB1/LGALS9/KLRC1/HAVCR2/LILRB1/SERPINB9/KLRD1/NCKAP1L/PTPRC/CR1/IL7R

INS/IL20RB/TNFRSF11A/PTGS2/FCGR2B/FCGR1A/CREB3L3/IL6/FUT7/NLRP3/C2CD4B/ADAM8/ALOX5AP/IL1B/DNASE1L3/TNF/TNFSF11/SPN/BTK/PIK3C

SH2B3/IL20RB/SLC46A2/SKIL/AXL/LMO1/DOCK11/CSF1/TNFSF13B/LIPA/CCR2/IL2/CD74/IL6/FLT3/MERTK/LGALS9/GPR183/TNFRSF17/GCNT4/TNFSF14/S

NFAT5/LRG1/CX3CR1/MDK/GCNT1/FUT4/ELANE/ADD2/ITGA4/CCL25/CHST4/VCAM1/CCR2/PTAFR/JAM2/SELPLG/IL6/FUT7/ALOX5/CCL21/KLF4/SELP/IT

ADCY1/LARGE1/CX3CL1/GRN/IL13/CX3CR1/AGER/CTSC/ADCY8/IGF1/CCL3/KCNJ8/TLR3/CD200R1L/LDLR/TLR2/PTGS2/IFNG/ADORA2A/IL4/TYROBP/C

IL20RB/LILRB2/TSPAN32/PAWR/VSIR/CD274/IL10/PDCD1LG2/PLA2G5/CD86/LGALS9/TMEM131L/HAVCR2/CD80/LILRB4/LILRB1/CRTAM/IL2RA/FOXP3/SP

C4BPA/CFH/C4BPB/SVEP1/TREM2/SERPING1/A2M/IL1B/CR1L/C3/CR1/CR2

TRIM40/AGER/CHI3L1/EDA/RASSF2/LIME1/EDAR/ZC3H12A/TLR3/NR3C2/TNFRSF10A/CYLD/TLR2/NMI/RELB/TCIM/MAP3K14/REL/EDN1/NOD2/NFKB2/IL

CASR/GPSM3/SLIT3/NRG1/CCL1/ELANE/EPHB1/F2RL1/NTRK3/NRG3/CCR9/CCL3/CMTM8/IL16/IL17RA/DEFB1/ANOS1/CXCL8/FGF7/CCR1/SMOC2/LBP/

VNN1/VCAM1/THBS1/IL10/LTA/FOXP3/UNC13D/TNF/CCL5/CXCL13/IDO1

KLK7/ZP4/FCGR2B/NOD2/TREM2/LTA/IL1B/TNF/C3/PTPRC/CCR7/CR1/FCER2

PAWR/FCGR2B/LPXN/CD300A/THY1/PLCL2/LILRB4/SLA2/UBASH3A/PTPN22/LAPTM5/PTPN6/SH2D1A/FCRL3/CD22

LGMN/CCL7/ITGA4/ASCL2/CCL4/CCR2/JAM2/XCL2/PTK2B/LGALS9/AIF1/ADAM8/WNT5A/TNFSF14/DOCK8/CCL21/IL12A/PLA2G7/TNF/SPN/CORO1A/CCR

CASR/GPSM3/SLIT3/NRG1/CCL1/ELANE/EPHB1/F2RL1/NTRK3/NRG3/CCR9/CCL3/CMTM8/IL16/IL17RA/DEFB1/ANOS1/CXCL8/FGF7/CCR1/SMOC2/LBP/

ITGA4/ASCL2/CCR2/XCL2/AIF1/ADAM8/WNT5A/TNFSF14/DOCK8/CCL21/SPN/CORO1A/CCL5/CXCL10/CCL20/CXCL13

TLR2/HSPA1B/AXL/PARP9/IRGM/EDN1/CSF1/IL7/RBM47/IKBKE/PARP14/CD74/TREM2/IL1R1/TXK/WNT5A/NLRC5/IRF7/MMP12/IFIH1/CD300LF/CXCR4/ZB

FCGR2B/FCGR1A/SYK/CD200R1/HCK/CLEC4E/FCER1G/LYN/PRKCQ/MYO1G/LCK/PLCG2/FCMR/CD247/BTK/FGR/LILRA4/VAV1/FCER2

RESF1/PROX1/HMGA2/IFNB1/OAS3/APOBEC3D/APOBEC3G/OAS1/APOBEC3H/IFITM1/ISG15/IFIT5/OAS2/AICDA/MX1/RSAD2/IFIH1/ISG20/LTF/IFIT1/TNF/

CX3CL1/CCL11/CCL23/ITGA1/PF4V1/RIPOR2/S100A9/MDK/IL1A/GP2/FUT4/CCL1/CCL3/CXCL8/LBP/CCL2/DPEP1/PPBP/CCL7/CSF3R/EDN1/CCL25/CCL14/

CD226/CD300A/MILR1/LYN/CD300LF/CD84/PTPN6/HAVCR1/CNR2

IZUMO1/SKAP1/GLDN/LILRB2/DSG2/ITGAD/DSC2/APOA1/ITGA4/VCAM1/FGA/IL10/CD200R1/PKP2/ITGAX/FGB/ITGA5/THY1/KLF4/ITGB2/ITGB7/IL1B/CD1D/

CD74/CD28/ZAP70/DOCK2/CD3E/SPN/THEMIS/PTPRC/CCR7

CD83/SYK/FCER1G/CD86/LGALS9/NLRP3/HAVCR2/CD28/PRKCQ/FOXP3/CD40LG/CD3E/GATA3/SASH3/IL33

CD83/SYK/FCER1G/CD86/LGALS9/NLRP3/HAVCR2/CD28/PRKCQ/FOXP3/CD40LG/CD3E/GATA3/SASH3/IL33

PRCP/KLKB1/ORM2/REG3A/NLRP6/NPY5R/PTGER3/SCN11A/CD163/FCER1A/FCGR3A/F3/IL1A/IL22/C2CD4A/MYLK3/ELANE/AHSG/INS/IL20RB/LBP/HP/F

C4BPA/FCGR2B/C4BPB/SVEP1/SERPING1/A2M/SPINK5/CR1L/PTPN6/CR1/CR2

ARG2/GHSR/TSC22D3/ST3GAL1/SLC46A2/FOXP1/ST6GAL1/AXL/PTCRA/NOD2/IL2/CD74/MERTK/PDCD1/DOCK8/IRF7/BCL3/CCL21/LILRB1/PRKCQ/CCL1

PPARG/TLR2/CCL7/PARP9/IFNG/IRGM/EDN1/CCL25/CCL14/GBP4/AQP4/CCL4/CXCL16/CCL22/CCL24/PARP14/IL12B/CD74/KYNU/STAT1/GCH1/IFITM1/H

IFNB1/ARID5A/IL4/CCR2/IL6/IL1R1/TNFRSF1B/NLRP3/RSAD2/LILRB4/FOXP3/IL1B/TBX21/GATA3/IL18R1/SASH3/IL18

IFNB1/ARID5A/IL4/CCR2/IL6/IL1R1/TNFRSF1B/NLRP3/RSAD2/LILRB4/FOXP3/IL1B/TBX21/GATA3/IL18R1/SASH3/IL18

FCGR3A/POMC/GNA15/CX3CR1/CA8/NR5A1/CLIC2/CHP2/GRIN2B/GBP1/PDGFRB/IAPP/PLA2G4B/LAT/GRIN1/IGF1/CCR9/NMUR1/CCL3/P2RY11/KCNJ8/

CLEC7A/IFNG/OAS3/CARD9/IL7/HMOX1/CSF1R/SYK/NOD2/OAS1/CD74/IL6/TGFB1/IL4R/LGALS9/AIF1/WNT5A/HAVCR2/TLR9/IL1RL1/IL1B/CD84/TNF/TLR

CCR2/XCL2/WNT5A/TNFSF14/CCL21/CCL5/CXCL10/CXCL13

KLRC3/TICAM2/CYBA/S100A14/PAK1/F2RL1/LBP/TLR5/TLR2/NMI/TLR1/CLEC7A/TYROBP/SYK/NOD2/OAS1/TREM2/NFKBIA/KLRC1/NR1H3/CLEC6A/LYN/

SLC46A2/DHX58/ARRB2/MMRN2/SH2D1B/CYLD/HRG/IFNB1/TLR2/IL21/HSPA1B/NMI/PARP9/REG3G/TLR1/CLEC7A/TIFA/IRGM/OAS3/CD274/TYROBP/CA

FCGR2B/FCGR1A/TYROBP/CCR2/SYK/PTAFR/IL4R/PRAM1/LGALS9/CD300A/GATA2/LYN/SPI1/ITGB2/UNC13D/CD84/DNASE1L3/BTK/FGR/RAC2/C3/STA

CD274/LGALS9/PDCD1/HAVCR2/LYN/LILRB4/CCR4/IL2RA/FOXP3/ICOS/CD3E/IDO1

TRIM40/AGER/EDA/RASSF2/LIME1/EDAR/ZC3H12A/TLR3/NR3C2/CYLD/TLR2/NMI/TCIM/EDN1/NOD2/IL12B/TREM2/NLRP2/CD86/LGALS9/NLRP3/TNFSF

ITGA4/ASCL2/CCR2/CD200R1/IL27RA/XCL2/AIF1/ADAM8/WNT5A/TNFSF14/DOCK8/SPNS2/CCL21/SPN/CORO1A/CCL5/CXCL10/CCR6/CCL20/CXCL13

S100A12/AZU1/WFDC3/IGHA1/PRTN3/DEFA4/DMBT1/REG3A/KRT6A/IL36RN/SLPI/RARRES2/PRSS3/WFDC11/PF4V1/DEFA5/BPIFA1/PGLYRP4/S100A9/CS

ARID5A/IL4/CD1C/IL12B/IL6/IL1R1/NLRP3/RAET1L/RSAD2/FOXP3/IL1B/IL12RB1/CD1D/IL12A/TBX21/GATA3/IL18R1/PTPRC/SASH3/IL23A/IL18

MMP26/MDK/NR1H4/RORA/GPER1/IL22/GHSR/NT5E/KRT1/ELANE/IGF1/STAT3/IL22RA1/INS/MEFV/IL20RB/ELF4/CYLD/TEK/PPARG/LDLR/ZFP36/APOA1/I

GNG3/GATA1/PTGER3/FPR1/TBXA2R/CCKBR/PML/P2RX2/P2RX7/AVPR1B/CX3CR1/GALR2/SPINK1/GALR1/AVP/GLP1R/GPER1/ADRA1B/CCL1/F2RL1/GR

CLEC7A/ARID5A/IL4/CARD9/IRF5/SYK/NOD2/CD74/CD226/IL6/INAVA/IL1R1/NLRP3/WNT5A/RSAD2/LILRB1/IL1B/SEMA7A/TBX21/CLNK/PLCG2/LAPTM5/GA

CCL2/CH25H/CCL7/CCL25/CCL14/CCL4/CXCL16/CCR2/CCL22/CCL24/XCL2/PTK2B/CCL8/ADAM8/WNT5A/GPR183/TNFSF14/CCL21/PADI2/CCL19/KLRK1/

SH2B3/IL20RB/TNR/C1QTNF1/LILRB2/TSPAN32/SERPINE2/PAWR/IFNB1/PODXL/APOA1/VSIR/IFNL1/ADORA2A/CD274/CCL25/ASCL2/LAG3/WNT1/IL10/IL2

TSPAN32/TLR5/SLC46A2/DHX58/ARRB2/USP18/MMRN2/SH2D1B/CYLD/PPARG/HRG/IFNB1/TLR2/IL21/NCR1/FOXP1/HSPA1B/NMI/LACRT/PARP9/REG3G

LBP/LILRB2/CCL2/CEBPE/PPBP/TLR2/AXL/ZFP36/LACRT/IL36A/ANKRD1/ARID5A/IRGM/CD274/EPHB2/NOD2/NUGGC/IL10/NLRP7/DEFB114/PTAFR/IL12B/

CCL7/ITGA4/ASCL2/CCL4/CCR2/JAM2/XCL2/PTK2B/AIF1/ADAM8/WNT5A/TNFSF14/DOCK8/CCL21/SPN/CORO1A/CCL5/CXCL10/CCL20/CXCL13

CYLD/PAWR/LIPA/CD226/CD300A/THY1/LILRB4/SLA2/UBASH3A/PTPN22/LCK/TRAT1/LAPTM5/TESPA1/PTPN6/CCR7/SH2D1A/CARD11

PRKD2/CX3CR1/RIPOR2/AGER/F3/MDK/PDGFRB/S100A14/CASR/GPSM3/CCL1/F2RL1/NTRK3/CCL3/IL16/CXCL8/CCR1/SMOC2/LBP/PTN/KDR/LGMN/CC

IL7/SYK/IL10/IL2/SPI1/NCKAP1L/BTK/CD27/INPP5D/IL2RG

IL10/CD200R1/IL27RA/GHRL/KLF2/HAVCR2/LILRB4/FOXP3/CD84/PTPN22/HGF/NCKAP1L/TNF/PTPN6/TLR4/INPP5D/NLRC3/BANK1

PPBP/TLR2/CDA/AXL/ZFP36/LACRT/IL36A/TLR1/ANKRD1/CLEC7A/ARID5A/IRGM/FCGR2B/CD274/EPHB2/TXNIP/SYK/NOD2/NUGGC/IL10/NLRP7/DEFB11

CXCL16/CCR2/XCL2/WNT5A/GPR183/TNFSF14/CCL21/PIK3CG/CCL5/CXCL10/CXCL11/CXCL13

IFNG/CSF1/CASP10/CEBPA/CSF1R/C1QC/CD4/TGFB1/IL31RA/UCP2/GATA2/IL15/GAB3/IL34/IL1RL1/SOCS1/HCLS1/INHBA/GATA3/TLR4/IL33/MMP9/LIF/AD

VSIR/CD274/CLEC4A/EOMES/TNFSF8/LILRB4/LILRB1/CRTAM/SOCS1/IRF1/NCKAP1L/TOX/GPR18/WDFY4

LBP/CCL2/DPEP1/PPBP/CCL7/CSF3R/EDN1/CCL25/CCL14/SLIT2/C3AR1/CCL4/SYK/NOD2/CCL22/CCL24/CD74/THBS4/FCER1G/XCL2/DAPK2/CCL8/CXCL

IFNG/TYROBP/CCR2/SYK/PTAFR/TREM2/DYSF/FCER1G/IL4R/PRAM1/LGALS9/CD300A/GATA2/MILR1/PTGDR/HAVCR2/LYN/SPI1/SUCNR1/ITGB2/UNC1

IL1RAP/F2RL1/UNC93B1/IL16/STAT3/IL17RA/TLR3/LBP/LILRB2/TLR2/HYAL2/IL36A/TMEM106A/TLR1/CLEC7A/ARID5A/IFNG/LILRA5/TYROBP/CARD9/SYK/N

F2RL1/CCL3/IL16/STAT3/PYDC1/MEFV/PANX3/PANX2/TMEM106A/CLEC7A/IFNG/LILRA5/TYROBP/NLRC4/NOD2/TLR8/IL6/NLRP10/INAVA/NLRP2/GBP5/LC

FOXP1/ST6GAL1/AXL/CD274/PTCRA/NOD2/IL10/IL2/CD74/MERTK/LGALS9/ADAM8/WNT5A/PDCD1/DOCK8/IRF7/BCL3/CCL21/LYN/LILRB1/PRKCQ/CCL19/

VSIR/IFNG/LAG3/IL2/TGFB1/FUT7/LGALS9/USP44/LILRB4/CD28/FOXP3/SOCS1/IRF1/TOX/IL4I1/PLA2G2D/IL2RG/CR1

CCL7/GCSAML/ITGA4/ASCL2/CCL4/CCR2/CD200R1/IL27RA/JAM2/XCL2/PTK2B/AIF1/GCSAM/ADAM8/WNT5A/TNFSF14/DOCK8/SPNS2/CCL21/PADI2/KLR

NFKBIZ/ST3GAL1/C17orf99/IL21/LFNG/DOCK11/FCGR2B/IL10/IL2/DLL1/IL6/CDH17/PTK2B/GPR183/BCL3/PLCG2/MFNG/ITM2A/POU2AF1/CR1/IRF8

PPBP/TLR2/FOXP1/MPO/TNFRSF11A/AXL/ZFP36/LACRT/IL36A/PTGS2/C2/ANKRD1/ARID5A/IRGM/ADM/EDN1/CD274/EPHB2/CYP1A1/SOD2/VCAM1/NOD

SYK/CD226/IL4R/LGALS9/CD300A/GATA2/MILR1/PTGDR/LYN/CD300LF/UNC13D/CD84/PTGDS/RASGRP1/CLNK/BTK/PIK3CG/FGR/RAC2/LAT2/PTPN6/HA

CCL3/CCL2/CCL7/CCL4/CCR2/XCL2/PTK2B/WNT5A/TNFSF14/CCL21/PADI2/KLRK1/CCL5/CXCL10/CXCL13

IFNB1/AXL/APOA1/FCGR2B/ZPBP2/NOD2/IL10/TGFB1/TGFB3/LILRB4/LILRB1/FOXP3/SPINK5/TBX21/JAK3/TNF/CD96/CR1/IL33/FCRL3/CD22

PTGS2/CLEC7A/IFNG/EDN1/SOD2/KLF2/PTK2B/JAK2/KLF4/IL1B/KLRK1/TNF/TLR6/TLR4/CLU/HBB

SLC46A2/DHX58/ARRB2/SH2D1B/CYLD/IFNB1/TLR2/IL21/HSPA1B/NMI/PARP9/REG3G/TLR1/CLEC7A/TIFA/IRGM/OAS3/CD274/TYROBP/CARD9/NLRC4/R

IL21/BATF/CLCF1/MZB1/IL4/FCGR2B/EPHB2/ZPBP2/IL10/IL2/IL27RA/IL6/TGFB1/IL4R/CD86/TNFRSF4/AICDA/SAMHD1/TLR9/CD28/GAPT/FOXP3/CD40LG/T

SYK/FCER1G/CD86/LGALS9/NLRP3/HAVCR2/CD28/PRKCQ/FOXP3/CD40LG/CD3E/GATA3/SASH3/IL33

PGLYRP2/AXL/TYROBP/IL12B/ZNF683/KLRC1/JAK2/IL15/HAVCR2/IL12A/RASGRP1/PTPN22/CLNK/KLRD1/TOX/FGR/IL15RA/IL23A/IL18

APOA1/CLEC7A/ARID5A/IL4/CARD9/IRF5/CCR2/SYK/NOD2/IL10/CD74/CD226/IL6/TGFB1/INAVA/IL1R1/TNFRSF1B/NLRP3/WNT5A/TGFB3/RSAD2/LILRB4/L

S100A14/GPSM3/CCL1/F2RL1/CCL3/CXCL8/CCR1/LBP/PTN/LGMN/CCL7/EDN1/CSF1/C3AR1/CCL4/CCR2/THBS1/CSF1R/VEGFC/CD74/THBS4/IL6/CMKLF

LGMN/CCL7/GCSAML/ITGA4/CSF1/BMP5/ASCL2/SLIT2/C3AR1/CCL4/CCR2/THBS1/CSF1R/CD200R1/IL27RA/JAM2/CMKLR1/XCL2/PTK2B/LGALS9/AIF1/GC

IFNB1/IFNL1/IL4/ASCL2/CCR2/NOD2/IL27RA/CD74/IL6/IL4R/CD86/NLRP3/RSAD2/TBX21/GATA3/IL33/IL18/IDO1

ZP4/CYLD/CD209/TEK/HRG/FOXA1/ECM2/WNT10B/KDR/IL21/CD83/HHLA2/VWC2/SERPINF2/PODXL/FBLN2/APOA1/VSIR/NRP1/IFNG/ITGA4/NODAL/IL4/C

SLC46A2/DHX58/ARRB2/USP18/SH2D1B/CYLD/PPARG/IFNB1/TLR2/IL21/NCR1/HSPA1B/NMI/PARP9/REG3G/TLR1/CLEC7A/TIFA/IRGM/OAS3/CLEC12B/FC

PPBP/TLR2/AXL/ZFP36/LACRT/IL36A/TLR1/ANKRD1/ARID5A/IRGM/FCGR2B/CD274/EPHB2/NOD2/NUGGC/IL10/NLRP7/DEFB114/PTAFR/IL12B/PDCD1LG2

TNFRSF11A/PTGS2/FCGR1A/CREB3L3/IL6/C2CD4B/ADAM8/ALOX5AP/IL1B/TNF/TNFSF11/BTK/PIK3CG/C3/CCR7

PTGS2/TMEM106A/CLEC7A/IFNG/EDN1/SOD2/KLF2/PTK2B/JAK2/KLF4/IL1B/KLRK1/TNF/TLR6/TLR4/CLU/HBB

IFNB1/BATF/IFNL1/IL4/ASCL2/CCR2/NOD2/IL10/IL27RA/CD74/IL6/IL4R/CD86/NLRP3/BCL3/RSAD2/TBX21/GATA3/IL33/IL18/IDO1

TRPM2/CCR2/LGALS9/GPR183/CCL21/CCL19/SLAMF8/IL12A/CXCR4/CCR5/PIK3CG/CCR7/CCL5/CCR6

IFNK/ZC3H12A/TLR3/IFNA8/IFNW1/IFNB1/IL21/IFNA4/IFNA1/IFNG/IFNL1/IRGM/IFNA6/IRF5/IKBKE/IFNA14/OAS1/IL6/LGALS9/NLRP3/JAK2/MMP12/IFIH1/CC

IL4/NOD2/CD74/IL6/IL4R/CD86/NLRP3/RSAD2/GATA3/IL33/IL18/IDO1

CCL3/STAT3/TLR3/TNFRSF8/LBP/FCGR2A/TLR2/TMEM106A/TLR1/CLEC7A/ARID5A/IFNG/OAS3/LILRA5/FCGR2B/EPHB2/TYROBP/CARD9/CCR2/THBS1/

SYK/NOD2/KLRC1/TCF7/JAML/LILRB1/TRDC/NCKAP1L/GPR18/CD247/CD3E/ITK/PTPRC/CD3G

BATF/ASCL2/IL2/IL27/IL12B/JUNB/IL4R/CD86/NLRP3/CD80/CCL19/FOXP3/IL12RB1/TBX21/JAK3/GATA3/IL23A/IL18

FOXP1/GCSAML/NFAM1/FCGR2B/LPXN/GCSAM/CD300A/LYN/PLCL2/PTPN22/FCMR/FCRL3/CD22/STAP1/CD19

IL20RB/LILRB2/TSPAN32/PAWR/IFNB1/VSIR/IFNL1/ADORA2A/CD274/ASCL2/LAG3/IL10/IL2/CD74/PDCD1LG2/PLA2G5/TNFAIP8L2/IL4R/CD86/LGALS9/CD30

IL21/ARID5A/CARD9/NOD2/IL2/IL12B/IL6/TGFB1/RFTN1/JAK2/IL15/PRKCQ/BTK/LY9/IL23A/SLAMF6/IL18

MZB1/CLEC7A/ARID5A/IL4/EPHB2/CARD9/IRF5/SYK/NOD2/CD244/IL10/IL2/CD74/CD226/IL6/TGFB1/IL4R/INAVA/IL1R1/CD86/TNFRSF4/NLRP3/WNT5A/RSA

IL4/EPHB2/IL7/TNFSF13B/SYK/NOD2/IL10/IL2/CD74/IL6/TGFB1/TNFRSF4/GPR183/SPI1/TLR9/CD28/TBX21/CD38/NFATC2/NCKAP1L/BTK/CD27/PTPRC/TLR

OR1G1/OR5H6/OR1S2/OR51A7/SCN1A/OR2T6/OR5M10/OR10G6/OR2A2/OR51A4/OR7A17/OR2K2/OR7D4/OR4K13/PPEF1/OR10P1/OR4Q3/OR7E24/OPN4/

VSIR/IFNL1/ADORA2A/CD274/CCL25/ASCL2/LAG3/IL10/IL2/CD74/PDCD1LG2/PLA2G5/TNFAIP8L2/IL4R/CD86/LGALS9/CD300A/DTX1/TMEM131L/HAVCR2/C

LILRB2/PTN/GPR55/PGLYRP2/THPO/SLC46A2/CYLD/KLF13/IFNB1/WNT10B/FOXP1/HSPA1B/MAFB/TNFRSF11A/TCIM/BATF/AXL/CD83/ZFP36/KAT6B/NFA

GPSM3/CCL1/F2RL1/CCL3/CXCL8/CCR1/LBP/PTN/CCL2/LGMN/CCL7/EDN1/CSF1/SLIT2/C3AR1/CCL4/CCR2/THBS1/CSF1R/NOD2/VEGFC/CD74/THBS4/

CIDEA/PGLYRP2/ELF4/DHX58/ARRB2/CYLD/SIGLEC1/IFNB1/NMI/RELB/AXL/CD83/ZFP36/APOA1/REL/VSIR/IFNG/IFNL1/IRGM/OAS3/LILRA5/IL4/FCGR2B/

CCR1/LBP/PTN/CCL2/CH25H/DPEP1/PPBP/LGMN/TNFRSF11A/CCL7/TRPM2/CSF3R/EDN1/CCL25/CSF1/CCL14/SLIT2/C3AR1/CCL4/CXCL16/CCR2/THBS

CD83/ZFP36/CLEC7A/CARD9/CCR2/LAG3/NOD2/CD4/CD86/HAVCR2/CD80/LILRB4/CD28/PRKCQ/FOXP3/IL1B/TBX21/PLCG2/LAPTM5/CD3E/GATA3/PTPR

CD83/ZFP36/CLEC7A/CARD9/CCR2/LAG3/NOD2/CD4/CD86/HAVCR2/CD80/LILRB4/CD28/PRKCQ/FOXP3/IL1B/TBX21/PLCG2/LAPTM5/CD3E/GATA3/PTPR

BATF/CLEC7A/ARID5A/EPHB2/CARD9/ASCL2/IL2/IL27RA/IL12B/JUNB/NLRP10/JAK2/CLEC6A/PRKCQ/FOXP3/IL12RB1/TBX21/IL23A

PPBP/TLR2/FOXP1/MPO/TNFRSF11A/AXL/ZFP36/LACRT/IL36A/PTGS2/TLR1/C2/ANKRD1/ARID5A/IRGM/ADM/EDN1/FCGR2B/CD274/EPHB2/CYP1A1/CAR

IL21/CTF1/CLCF1/LACRT/PARP9/NRP1/IFNG/IFNL1/LILRA5/FGFR3/GFRA1/IL4/FCGR1A/PLPP3/IL3/IL11/TNFRSF18/CSF1R/SYK/NOD2/IL2/PARP14/IL12B/C

CX3CL1/GRN/FCGR3A/IL13/CX3CR1/SPACA3/AGER/RORA/CTSC/BPI/CCL3/ZC3H12A/KCNJ8/TLR3/LBP/LDLR/TLR2/FOXP1/NMI/TMEM106A/TLR1/IFNG/I

IFNB1/IL21/BATF/AXL/CLCF1/MZB1/APOA1/CLEC7A/ARID5A/IL4/FCGR2B/EPHB2/CARD9/IRF5/ZPBP2/CCR2/SYK/NOD2/CD244/IL10/IL2/IL27RA/CD74/CD2

CD83/IFNG/SYK/IL12B/ZBTB16/IL4R/CD86/LGALS9/NLRP3/CD80/LILRB4/ZAP70/CCL19/FOXP3/SOCS1/IL12RB1/NCKAP1L/SASH3/IL23A/IL2RG/IL18

CCL3/STAT3/TLR3/TNFRSF8/LBP/FCGR2A/TLR2/TMEM106A/TLR1/CLEC7A/ARID5A/IFNG/OAS3/LILRA5/FCGR2B/EPHB2/TYROBP/CARD9/CCR2/THBS1/

AGER/BATF2/CSF2/F2RL1/LILRB2/RELB/BATF/AXL/TRPM2/IL4/FCGR2B/TREM2/BATF3/FLT3/CAMK4/LGALS9/TMEM176A/LYN/LILRB1/TMEM176B/CCL19/

SMOC2/LBP/PTN/CCL2/CH25H/ARRB2/DPEP1/AGTR1/HRG/KDR/PPBP/LGMN/TNFRSF11A/CCL7/TRPM2/CSF3R/NRP1/EDN1/CCL25/TMSB4X/CSF1/CC

APOA1/CLEC7A/ARID5A/IL4/CARD9/IRF5/CCR2/SYK/NOD2/IL10/CD74/CD226/IL6/TGFB1/INAVA/IL1R1/TNFRSF1B/NLRP3/WNT5A/TGFB3/RSAD2/LILRB4/L

EDAR/TLR3/NR3C2/TLR2/NMI/TCIM/EDN1/NOD2/IL12B/TREM2/CD86/LGALS9/NLRP3/TNFSF14/HAVCR2/TLR9/CCL19/IL1B/TNF/LAPTM5/CD27/IL18R1/TL

SH2D1B/IL21/LAG3/NCR3/IL12B/CD226/IL18RAP/KLRC1/KLRC4/CRTAM/IL12A/RASGRP1/CLNK/KLRD1/KLRK1/VAV1/SH2D1A/SLAMF6

HRG/INPP5F/IL21/HYAL2/CTF1/CLCF1/LACRT/IL9/PARP9/NRP1/IFNG/IFNL1/LILRA5/FGFR3/GFRA1/IL4/FCGR1A/PLPP3/IL7/IL3/IL11/TNFRSF18/CSF1R/SY

IL21/CTF1/CLCF1/PARP9/IFNG/IFNL1/FGFR3/IL4/IL3/TNFRSF18/CSF1R/IL2/PARP14/IL12B/IL6/FLT3/IL31RA/JAK2/IL15/CD40/LYN/IL12A/HCLS1/TNF/CCL5/IL

DEFB124/RARRES2/HTN3/GRN/P2RX7/DEFB112/VGF/WFDC11/DEFA5/DEFB123/BPIFA1/PGLYRP4/DEFB128/S100A9/ZNFX1/CYBA/CST11/NR1H4/GBP1/

XKR5/XKR4/FCGR2B/FCGR1A/THBS1/TREM2/AIF1/CD300A/GATA2/ITGB2/BIN2/ELMO1/PLCG2/NCKAP1L/TIMD4/MARCO/C3/ARHGAP25/HAVCR1/ALOX1

FCGR2B/FCGR1A/SYK/NOD2/IL10/IL12B/TREM2/HCK/FUT7/LYN/LTA/CD28/TNF/SPN/BTK/FGR/C3/PLA2G2D/CCR7

F2RL1/STAT3/IL20RB/TLR2/CD83/CLEC7A/LILRA5/IL4/CD274/RBM47/SYK/NOD2/IL12B/TREM2/ISG15/IL6/INAVA/LGALS9/BCL3/TLR9/CD28/CD40LG/TIGIT/

CCL2/C4B/RAB31/FCGR2A/CFP/C2/APOA1/CLEC7A/IFNG/FCGR2B/FCGR1A/NOD2/TREM2/PLA2G5/FCER1G/MERTK/GATA2/IL15/SIRPG/CAMK1D/SIRP

TRPM2/ASB2/CCR2/LGALS9/ALOX5/GPR183/DOCK8/CCL21/CCL19/SLAMF8/IL12A/CXCR4/CCR5/PIK3CG/CCR7/CCL5/CCR6

NCR2/TCIRG1/CX3CR1/KLRC3/CD300C/CCR9/LBP/LILRB2/NCR1/NCF2/ADORA2A/TYROBP/CCR2/PTK2B/MNDA/LSP1/GNLY/PRF1/CCR5/KLRG1/TRAT

CD274/TNFSF13B/PDCD1LG2/EFNB2/CD86/TNFSF14/CCL21/CD80/LYN/LILRB4/CD28/CCL19/ICOS/CD40LG/LCK/CD5/KLRK1/CD3E/SPN/VAV1/PTPN6/CC

ZC3H12A/IL20RB/PGLYRP2/AXL/VSIR/IRGM/CD274/NOD2/IL10/PDCD1LG2/LGALS9/HAVCR2/LILRB4/LILRB1/IL1RL1/FOXP3/INHBA/LAPTM5/GATA3/CD96/

AGER/CHRNA7/CYBA/IL1A/NR1H4/ARG2/GHSR/BPI/IGF1/CCL3/STAT3/ZC3H12A/TLR3/TNFRSF8/LBP/CIDEA/ELF4/ARRB2/FCGR2A/TLR2/FOXP1/AXL/ZF

AGER/CHRNA7/CYBA/IL1A/NR1H4/ARG2/GHSR/BPI/IGF1/CCL3/STAT3/ZC3H12A/TLR3/TNFRSF8/LBP/CIDEA/ELF4/ARRB2/FCGR2A/TLR2/FOXP1/AXL/ZF

CYLD/WNT10B/CD83/VSIR/IFNG/IL4/VNN1/IL7/SYK/IL2/IL12B/CD74/IL36B/ZBTB16/IL4R/CD86/LGALS9/NLRP3/ADAM8/PIK3R6/CD80/LILRB4/ZAP70/CCL19/IL

LAG3/NCR3/IL12B/CD226/IL18RAP/KLRC1/KLRC4/CRTAM/IL12A/RASGRP1/KLRD1/KLRK1/VAV1/SH2D1A/SLAMF6

RORA/NFKBIZ/STAT3/IL17RA/ZC3H12A/RORC/BATF/CLEC7A/ARID5A/EPHB2/CARD9/ASCL2/IL2/IL27RA/SOCS3/IL12B/JUNB/IL6/NLRP10/JAK2/CLEC6A/P

C4BPA/FCGR2B/NOD2/C4BPB/SVEP1/TREM2/LTA/CR1L/TNF/PTPN6/PTPRC/CR1/FCER2/CR2

SLC46A2/DHX58/ARRB2/MMRN2/SH2D1B/CYLD/AGTR1/LDLR/IFNB1/TLR2/IL21/HSPA1B/NMI/HYAL2/TNFRSF11A/PARP9/PTGS2/REG3G/TLR1/CLEC7A/T

UNC93B1/IL16/TLR3/TLR2/CLEC7A/IFNG/IRF5/SYK/NOD2/IL12B/LGALS9/TLR9/CCL19/IRF1/CD40LG/PLCG2/LAPTM5/SCIMP/TLR4/CCR7/IL23A/LTB/IRF8/I

CD83/IFNG/IL12B/IL4R/CD86/LGALS9/NLRP3/CD80/CCL19/FOXP3/SOCS1/IL12RB1/NCKAP1L/SASH3/IL23A/IL2RG/IL18

AGER/CHRNA7/CYBA/IL1A/NR1H4/ARG2/GHSR/BPI/IGF1/CCL3/STAT3/ZC3H12A/TLR3/TNFRSF8/LBP/CIDEA/ELF4/ARRB2/FCGR2A/TLR2/FOXP1/AXL/ZF

AGER/CHRNA7/CYBA/IL1A/NR1H4/ARG2/GHSR/BPI/IGF1/CCL3/STAT3/ZC3H12A/TLR3/TNFRSF8/LBP/CIDEA/ELF4/ARRB2/FCGR2A/TLR2/FOXP1/AXL/ZF

GCSAML/TRPM2/CSF3R/ASB2/EDN1/ITGA4/CCL25/CSF1/CCL14/BMP5/ASCL2/SLIT2/C3AR1/CHST4/CCL4/HMOX1/LYVE1/VCAM1/CXCL16/CCR2/P2RY12

LBP/CCL2/RAB39A/C4B/P2RY6/CEBPE/RAB31/LDLR/FCGR2A/ICAM5/TLR2/C4BPA/CFP/AXL/NCF2/XKR5/C2/APOA1/CLEC7A/CEACAM4/IFNG/XKR4/ADO

CD274/TNFSF13B/PDCD1LG2/EFNB2/CD86/TNFSF14/CCL21/CD80/LYN/LILRB4/CD28/CCL19/ICOS/CD40LG/LCK/CD5/KLRK1/CD3E/SPN/VAV1/PTPN6/CC

CCL2/TSPAN32/GPR55/PGLYRP2/DHX58/ARRB2/USP18/PPARG/TAPBPL/LDLR/PAWR/IFNB1/MAFB/NMI/C4BPA/AXL/APOA1/VSIR/IFNL1/IRGM/OAS3/ADC

IL36A/FCGR2B/FCGR1A/SYK/NOD2/IL10/IGHG1/IL12B/TREM2/IL36B/HCK/FUT7/CYSLTR1/IL31RA/LYN/LTA/CD28/A2M/IL2RA/RASGRP1/IL1RN/CD6/TNF/S

CACNB4/CYLD/PAWR/LCP2/HHLA2/FYB2/FYB1/LIPA/CD226/TXK/PRAM1/RFTN1/CD300A/THY1/LILRB4/TRDC/CD28/ZAP70/CD8A/TRGC2/FOXP3/SLA2/U

BPI/IL1RAP/F2RL1/UNC93B1/IL16/STAT3/IL17RA/ZC3H12A/TLR3/LBP/LILRB2/ELF4/ARRB2/TLR2/HYAL2/IL36A/TMEM106A/TLR1/CLEC7A/ARID5A/IFNG/LIL

BPI/IL1RAP/F2RL1/UNC93B1/IL16/STAT3/IL17RA/ZC3H12A/TLR3/LBP/LILRB2/ELF4/ARRB2/TLR2/HYAL2/IL36A/TMEM106A/TLR1/CLEC7A/ARID5A/IFNG/LIL

CD74/CD28/ZAP70/DOCK2/TOX/CD3E/SPN/GATA3/CD3D/PTPRC/CCR7/CARD11/CD3G

FZD8/VNN1/WNT1/CD74/CAMK4/ADAM8/TMEM131L/CD28/ZAP70/FOXP3/IL1B/DOCK2/RASGRP1/TOX/CD3E/SPN/GATA3/CD3D/TESPA1/PTPRC/CCR7/B

EDN1/ITGA4/CSF1/ASCL2/C3AR1/CCL4/LYVE1/CCR2/P2RY12/TNFRSF18/THBS1/CSF1R/CCL24/VEGFC/PTAFR/JAM2/CD74/TREM2/THBS4/IL6/FUT7/IL1

IGF1/STAT3/IFNK/SH2B3/IFNA8/CCL2/IFNW1/IFNB1/INPP5F/IL21/NMI/CTF1/IFNA4/CLCF1/IL9/PARP9/IFNA1/IFNG/IFNL1/FGFR3/IL4/EPHB2/IFNA6/IL3/F2/C

WNT10B/TNFRSF11A/AXL/CD83/VSIR/IFNG/IL4/TYROBP/CSF1/VNN1/IL7/SYK/IL10/IL2/IL12B/CD74/TREM2/IL36B/CD4/TGFB1/ZBTB16/IL4R/CD86/LGALS9/

WNT10B/TNFRSF11A/AXL/CD83/VSIR/IFNG/IL4/TYROBP/CSF1/VNN1/IL7/SYK/IL10/IL2/IL12B/CD74/TREM2/IL36B/CD4/TGFB1/ZBTB16/IL4R/CD86/LGALS9/

CCL7/GCSAML/ITGA4/CCL25/CCL14/ASCL2/CCL4/CXCL16/CCR2/CCL22/CCL24/CD200R1/IL27RA/JAM2/FUT7/XCL2/PTK2B/AIF1/CCL8/GCSAM/ADAM8/W

ITGA4/ASCL2/CXCL16/CCR2/CD200R1/IL27RA/FUT7/XCL2/AIF1/ADAM8/WNT5A/GPR183/TNFSF14/DOCK8/SPNS2/ICAM1/CCL21/ZAP70/ITGB7/MYO1G/S

IL21/CLCF1/MZB1/IL4/FCGR2B/EPHB2/ZPBP2/IL10/IL2/IL27RA/IL6/TGFB1/IL4R/CD86/TNFRSF4/TLR9/CD28/FOXP3/CD40LG/TBX21/TNF/BTK/RBP4/PTPRC

VSIR/FCGR2B/CD274/TYROBP/LST1/IL10/IL2/PDCD1LG2/PLA2G5/CD86/LGALS9/CCL8/MNDA/CD300A/TMEM131L/HAVCR2/CD80/LYN/LILRB4/LILRB1/CR

BATF/CD83/IFNG/ASCL2/SYK/IL2/IL27/IL12B/ZNF683/JUNB/ZBTB16/IL4R/CD86/LGALS9/NLRP3/CD80/LILRB4/ZAP70/CCL19/FOXP3/SOCS1/IL12RB1/TBX21/

NFKBIZ/ST3GAL1/C17orf99/IL21/LFNG/DOCK11/FCGR2B/IL10/IL2/DLL1/IL6/CDH17/PTK2B/GPR183/BCL3/PLCL2/SLAMF8/PLCG2/MFNG/ITM2A/POU2AF1/IL

EDN1/ITGA4/CCL25/CSF1/BMP5/ASCL2/SLIT2/C3AR1/CHST4/CCL4/HMOX1/LYVE1/CCR2/P2RY12/TNFRSF18/THBS1/CSF1R/NOD2/CCL24/CD200R1/IL27

IFNB1/IL21/BATF/CLCF1/LFNG/DOCK11/IL4/FCGR2B/IL10/IL2/IL27RA/DLL1/IL6/TGFB1/CDH17/PTK2B/AICDA/GPR183/BCL3/PLCL2/CD28/GAPT/FOXP3/CD4

OR4F15/OR8D2/OR2G3/OR10J5/OR2T11/OR4E1/OR5AC2/OR6C76/OR10X1/GNAT2/OR4A5/OR1G1/OR5H6/OR1S2/OR51A7/OR2T6/OR5M10/OR10G6/OR2

IL20RB/LILRB2/TSPAN32/PAWR/VSIR/FCGR2B/CD274/TYROBP/LST1/IL10/IL2/PDCD1LG2/PLA2G5/CD86/LGALS9/MNDA/CD300A/TMEM131L/HAVCR2/CD

CCL2/C4B/RAB31/FCGR2A/TLR2/CFP/C2/APOA1/CLEC7A/IFNG/FCGR2B/FCGR1A/SYK/NOD2/TGM2/TREM2/PLA2G5/DYSF/HCK/FCER1G/MERTK/CD30

SOCS3/IL12B/CD74/IL6/ZAP70/FOXP3/IL12RB1/DOCK2/TBX21/TOX/CD3E/SPN/CD3D/LY9/THEMIS/PTPRC/BCL11B/IL23A/SLAMF6/CD3G

PAWR/IL21/CLCF1/MZB1/IL4/FCGR2B/EPHB2/TYROBP/IL7/TNFSF13B/IL10/IL2/CD74/TNFRSF4/MNDA/CD300A/GPR183/LYN/TLR9/CD38/NFATC2/NCKAP

IGF1/STAT3/IFNK/SH2B3/IFNA8/CRLF2/CCL2/THPO/IFNW1/PPARG/IFNB1/INPP5F/IL21/NMI/CTF1/IFNA4/CLCF1/IL9/PARP9/IFNA1/IFNG/IFNL1/FGFR3/IL4

IL20RB/LILRB2/TSPAN32/PAWR/VSIR/FCGR2B/CD274/TYROBP/LST1/IL10/IL2/PDCD1LG2/PLA2G5/CD86/LGALS9/MNDA/CD300A/TMEM131L/HAVCR2/CD

INPP5F/IL21/CTF1/CLCF1/PARP9/IFNG/IFNL1/FGFR3/IL4/IL3/TNFRSF18/CSF1R/IL2/PARP14/SOCS3/IL12B/IL6/FLT3/IL31RA/JAK2/IL15/CD40/LYN/SOCS1/IL

FCGR2B/TYROBP/IL10/SAMSN1/SFRP1/MNDA/CD300A/LYN/FOXP3/INHBA/LAPTM5/BTK/INPP5D/TBC1D10C/CR1/TNFRSF13B/BANK1

CCR2/SYK/IL12B/LGALS9/JAK2/EBI3/CD28/PRKCQ/ZAP70/CD3E/RASAL3/PTPRC/IL23A/CARD11/IL18

CLCF1/CLEC7A/ARID5A/IL4/CD274/FCGR1A/CARD9/TNFSF13B/CCR2/NOD2/CD1C/IL2/IL27RA/IL12B/CD226/TREM2/CD4/IL6/TGFB1/NLRP10/IL1R1/NLRP

HRG/INPP5F/KDR/IL21/HYAL2/CTF1/PKDCC/CLCF1/LACRT/IL9/PARP9/NRP1/IFNG/IFNL1/LILRA5/FGFR3/GFRA1/IL4/EPHB2/FCGR1A/PLPP3/IL7/IL3/IL11/T

HRG/INPP5F/KDR/IL21/HYAL2/CTF1/PKDCC/CLCF1/LACRT/IL9/PARP9/NRP1/IFNG/IFNL1/LILRA5/FGFR3/GFRA1/IL4/EPHB2/FCGR1A/PLPP3/IL7/IL3/IL11/T

IFNB1/IL21/AXL/CLCF1/MZB1/APOA1/CLEC7A/ARID5A/IL4/FCGR2B/EPHB2/CARD9/IRF5/ZPBP2/CCR2/SYK/NOD2/CD244/IL10/IL2/IL27RA/CD74/CD226/IL6/

TYROBP/LAG3/NCR3/CD1C/IL12B/CD226/IL18RAP/KLRC1/RAET1L/SPI1/KLRC4/CRTAM/IL12RB1/CD1D/IL12A/RASGRP1/KLRD1/KLRK1/VAV1/PTPRC/SH2

TNR/C1QTNF1/LILRB2/CCL2/TSPAN32/ZP4/CYLD/CD209/FERMT3/SERPINE2/FOXA1/PAWR/IFNB1/WNT10B/IL21/CD83/HHLA2/SERPINF2/PODXL/APOA1

CD83/IFNG/IL12B/IL4R/CD86/LGALS9/NLRP3/CD80/CD28/PRKCQ/CCL19/FOXP3/SOCS1/IL12RB1/NCKAP1L/CD3E/SASH3/IL23A/IL2RG/CARD11/IL18

RORA/MYB/NFKBIZ/STAT3/ZC3H12A/RORC/IL21/FOXP1/RELB/BATF/ASCL2/SEMA4A/IL2/IL27/SOCS3/IL12B/PTGER4/JUNB/IL6/IL4R/CD86/NLRP3/STAT4/C

ARID5A/IL4/FCGR2B/CCR2/NOD2/CD1C/IL12B/IL6/FUT7/IL1R1/TNFRSF1B/KLRC1/NLRP3/RAET1L/CD80/RSAD2/LILRB4/LILRB1/FOXP3/IL1B/IL12RB1/CD1

OR4F15/OR8D2/OR2G3/OR10J5/OR2T11/OR4E1/OR5AC2/OR6C76/OR10X1/OR4A5/OR1G1/OR5H6/OR1S2/OR51A7/OR2T6/OR5M10/OR10G6/OR2A2/OR5

UNC93B1/IL16/TLR3/MEFV/ARRB2/TLR2/FOXP1/CLEC7A/IFNG/LILRA5/IRF5/THBS1/SYK/NOD2/IL10/TLR8/IL12B/CMKLR1/LGALS9/LILRB1/TLR9/CCL19/IRF

UNC93B1/IL16/TLR3/MEFV/ARRB2/TLR2/FOXP1/CLEC7A/IFNG/LILRA5/IRF5/THBS1/SYK/NOD2/IL10/TLR8/IL12B/CMKLR1/LGALS9/LILRB1/TLR9/CCL19/IRF

INPP5F/IL21/CTF1/CLCF1/PARP9/IFNG/IFNL1/FGFR3/IL4/IL3/TNFRSF18/CSF1R/IL2/PARP14/SOCS3/IL12B/IL6/FLT3/IL31RA/JAK2/IL15/CD40/LYN/SOCS1/IL

VSIR/CD274/CCR2/SYK/IL2/IL12B/LGALS9/JAK2/EBI3/CD28/PRKCQ/ZAP70/IL2RA/FOXP3/IRF1/CD3E/RASAL3/PTPRC/IL23A/CARD11/IL18

IL21/VSIR/ARID5A/IFNG/CARD9/NOD2/IL2/IL27RA/IL12B/IL6/TGFB1/RFTN1/JAK2/IL15/PRKCQ/FOXP3/IL12A/NCKAP1L/BTK/LY9/TLR4/IL23A/SLAMF6/IL18

IL21/VSIR/ARID5A/IFNG/CARD9/NOD2/IL2/IL27RA/IL12B/IL6/TGFB1/RFTN1/JAK2/IL15/PRKCQ/FOXP3/IL12A/NCKAP1L/BTK/LY9/TLR4/IL23A/SLAMF6/IL18

ARRB2/IL21/NCR1/CLEC12B/SERPINB4/LAG3/NCR3/IL12B/CD226/TGFB1/IL18RAP/LGALS9/KLRC1/PIK3R6/HAVCR2/KLRC4/LILRB1/CRTAM/SERPINB9/IL1

CYLD/WNT10B/AXL/CD83/VSIR/IFNG/IL4/VNN1/IL7/SYK/IL10/IL2/IL12B/CD74/IL36B/ZBTB16/IL4R/CD86/LGALS9/NLRP3/ADAM8/IL15/PIK3R6/CD80/SPI1/LILR

OR1G1/OR5H6/OR1S2/OR51A7/OR2T6/OR5M10/OR10G6/OR2A2/OR51A4/OR7A17/OR2K2/OR7D4/OR4K13/OR10P1/OR4Q3/OR7E24/OR8S1/OR5T2/OR2H

AGER/S100A9/MMP26/MDK/NR1H4/RORA/CTSC/GPER1/IL22/C2CD4A/GHSR/NLRP11/NLRP4/NFKBIZ/GPSM3/NT5E/CCL1/KRT1/ELANE/IGF1/CCL3/IL16/S

GPR55/PGLYRP2/SLC46A2/CYLD/IFNB1/WNT10B/FOXP1/MAFB/TNFRSF11A/BATF/AXL/CD83/NFAM1/VSIR/IFNG/IFNL1/IL4/FCGR2B/TYROBP/CSF1/VNN

LGMN/TNFRSF11A/CCL7/GCSAML/TRPM2/ASB2/ITGA4/CCL25/CSF1/CCL14/BMP5/ASCL2/SLIT2/C3AR1/CCL4/CXCL16/CCR2/THBS1/CSF1R/CCL22/CCL

SLC46A2/CYLD/IFNB1/WNT10B/BATF/CD83/VSIR/IFNG/IFNL1/IL4/VNN1/ASCL2/IL7/CCR2/LAG3/SYK/IL2/IL27/IL12B/CD74/ZNF683/JUNB/IL36B/ZBTB16/IL4R

UNC93B1/CCL3/KLRF2/IL16/STAT3/IL17RA/INS/PYDC1/ADRA2A/TLR3/MEFV/PANX3/IL20RB/CRLF2/TNFRSF8/LBP/LILRB2/TLR5/DHX58/SULF1/FCGR2A/T

ARID5A/ASCL2/CCR2/IL27RA/IL27/IL12B/NLRP10/IL4R/IL1R1/HAVCR2/CD80/IL1RL1/CCL19/IL1B/IL12RB1/TBX21/JAK3/IL18R1/IL23A/IL33/IL18

C1RL/C4BPA/CFP/C2/CFH/C1QA/C4BPB/SVEP1/IGHG1/C1QB/IGHG3/TREM2/C1QC/SERPING1/C1S/IGHE/C7/CFB/A2M/C1R/IL1B/CR1L/IGHA2/C3/CR1/CL

VSIR/CD274/CCR2/SYK/IL2/IL12B/LGALS9/JAK2/IL15/EBI3/CD28/PRKCQ/ZAP70/IL2RA/FOXP3/IRF1/DOCK2/CD3E/RASAL3/PTPRC/IL23A/CARD11/IL18

RORA/MYB/NFKBIZ/STAT3/ZC3H12A/RORC/IL21/FOXP1/RELB/BATF/IFNG/ASCL2/SEMA4A/IL2/IL27/SOCS3/IL12B/PTGER4/JUNB/IL6/IL4R/CD86/LGALS9/N

SH2B3/IL20RB/C1QTNF1/LILRB2/TSPAN32/PGLYRP2/SERPINE2/LDLR/PAWR/IFNB1/AXL/VSIR/IFNL1/ADORA2A/FCGR2B/CD274/TYROBP/ASCL2/LST1/F2

CLCF1/CLEC7A/ARID5A/IL4/CD274/FCGR1A/CARD9/TNFSF13B/CCR2/NOD2/CD1C/IL2/IL27RA/IL12B/CD226/TREM2/CD4/IL6/TGFB1/NLRP10/IL1R1/NLRP

OR4F15/OR8D2/OR2G3/OR10J5/OR2T11/OR4E1/OR5AC2/OR6C76/OR10X1/GNAT2/OR4A5/OR1G1/OR5H6/OR1S2/OR51A7/OR2T6/OR5M10/OR10G6/OR2

C4BPA/CLCF1/IL4/FCGR2B/FCGR1A/NOD2/C4BPB/IL10/SVEP1/IL2/IL27RA/CD226/TREM2/TGFB1/LTA/CD28/FOXP3/CR1L/TBX21/TNF/BTK/PTPN6/C3/PTP

C4BPA/CLCF1/IL4/FCGR2B/FCGR1A/NOD2/C4BPB/IL10/SVEP1/IL2/IL27RA/CD226/TREM2/TGFB1/LTA/CD28/FOXP3/CR1L/TBX21/TNF/BTK/PTPN6/C3/PTP

PGLYRP2/DHX58/ARRB2/USP18/PPARG/IFNB1/NMI/C4BPA/IFNL1/OAS3/CLEC12B/FCGR2B/ASCL2/SERPINB4/CCR2/SYK/NOD2/C4BPB/IL10/SVEP1/IL2/IL

LDLR/TLR2/FOXP1/LCP2/NMI/HYAL2/RELB/BATF/TMEM106A/TLR1/IFNG/IRGM/IL4/FCGR2B/TYROBP/CSF1/CEBPA/C1QA/CCR2/THBS1/SYK/IL10/PTAFR

TLR2/CD83/CLEC7A/VSIR/LILRA5/IL4/FCGR2B/CD274/TYROBP/RBM47/SYK/NOD2/IL12B/DLL1/PDCD1LG2/TREM2/ISG15/IL6/INAVA/LGALS9/BCL3/LILRB4/

TLR2/CD83/CLEC7A/VSIR/LILRA5/IL4/FCGR2B/CD274/TYROBP/RBM47/SYK/NOD2/IL12B/DLL1/PDCD1LG2/TREM2/ISG15/IL6/INAVA/LGALS9/BCL3/LILRB4/

ARRB2/IL21/NCR1/SLAMF7/CLEC12B/SERPINB4/LAG3/NCR3/IL12B/CD226/TGFB1/IL18RAP/LGALS9/KLRC1/PIK3R6/HAVCR2/KLRC4/LILRB1/CRTAM/UNC

BATF/CD83/IFNG/ASCL2/IL2/IL27/IL12B/JUNB/IL4R/CD86/LGALS9/NLRP3/CD80/CCL19/FOXP3/SOCS1/IL12RB1/TBX21/JAK3/NCKAP1L/GATA3/SASH3/IL23A

C4B/C1RL/C4BPA/C2/C1QA/C4BPB/SVEP1/IGHG1/C1QB/IGHG3/TREM2/C1QC/SERPING1/C1S/IGHE/C7/C1R/CR1L/IGHA2/C3/CR1/CLU/CFI/IGHM/CR2

RORC/IL21/FOXP1/RELB/BATF/IFNG/ASCL2/SEMA4A/IL2/IL27/SOCS3/IL12B/PTGER4/JUNB/IL6/IL4R/EOMES/CD86/LGALS9/NLRP3/STAT4/GPR183/BCL3/C

RORC/IL21/FOXP1/RELB/BATF/IFNG/ASCL2/SEMA4A/IL2/IL27/SOCS3/IL12B/PTGER4/JUNB/IL6/IL4R/EOMES/CD86/LGALS9/NLRP3/STAT4/GPR183/BCL3/C

PRDM1/KLRF2/PBX1/IFNK/IFNA8/PGLYRP2/ELF4/IFNW1/IFNB1/NCR1/IFNA4/AXL/IFNA1/SLAMF7/TYROBP/IFNA6/IFNA14/NCR3/CD244/IL2/IL12B/ZNF683/

OR4F15/OR8D2/OR2G3/OR10J5/OR2T11/OR4E1/OR5AC2/OR6C76/OR10X1/OR4A5/OR1G1/OR5H6/OR1S2/OR51A7/OR2T6/OR5M10/OR10G6/OR2A2/OR5

ARRB2/SH2D1B/IL21/NCR1/CLEC12B/SERPINB4/LAG3/NCR3/IL12B/CD226/TGFB1/IL18RAP/LGALS9/KLRC1/PIK3R6/HAVCR2/KLRC4/LILRB1/CRTAM/SERP

RORC/IL21/FOXP1/RELB/BATF/IFNG/IFNL1/ASCL2/SEMA4A/IL2/IL27/SOCS3/IL12B/PTGER4/JUNB/IL6/CLEC4E/FCER1G/IL4R/EOMES/CD86/LGALS9/NLRP

STAT3/IFNK/ZC3H12A/IFNA8/RORC/IFNW1/IFNB1/IL21/FOXP1/IFNA4/RELB/BATF/IFNA1/IFNG/IFNL1/FCGR2B/IFNA6/ASCL2/SEMA4A/IFNA14/CD1C/IL2/IL

RELB/ARID5A/ASCL2/SEMA4A/CCR2/IL27RA/IL27/IL12B/NLRP10/IL4R/IL1R1/STAT4/BCL3/HAVCR2/CD80/IL18BP/EBI3/IL1RL1/CCL19/IL1B/IL12RB1/TBX21/JA

ZP4/CD209/IL21/ST6GAL1/HHLA2/CLCF1/IL4/CD274/EPHB2/CSF1/IL7/TNFSF13B/VCAM1/CCR2/SYK/IL2/IL12B/CD74/PDCD1LG2/IL6/CD86/LGALS9/AIF1/TN

IL4/CD274/EPHB2/IL7/TNFSF13B/VCAM1/CCR2/SYK/IL2/IL12B/CD74/PDCD1LG2/IL6/CD86/LGALS9/AIF1/TNFRSF4/CD70/GPR183/JAK2/IL15/HAVCR2/CD80

LILRB2/TSPAN32/ELF4/ZP4/CD209/SH2D2A/PAWR/IL21/HHLA2/LMO1/VSIR/IL4/CD274/TNFSF13B/VCAM1/LIPA/CCR2/SYK/IL10/IL2/IL27/IL12B/PDCD1LG2/P
ZP4/CD209/IL21/ST6GAL1/HHLA2/CLCF1/IL4/CD274/EPHB2/CSF1/IL7/TNFSF13B/VCAM1/CSF2RA/CCR2/CSF1R/SYK/IL2/IL12B/CD74/PDCD1LG2/IL6/CD86/
IL20RB/ARRB2/IFNB1/C4BPA/CLEC12B/FCGR2B/SERPINB4/NOD2/C4BPB/SVEP1/TGFB1/LGALS9/KLRC1/HAVCR2/CD80/LILRB4/LILRB1/FOXP3/SERPINB
IFNB1/C4BPA/SCART1/ARID5A/IL4/FCGR2B/CCR2/NOD2/CD1C/IL12B/IL6/FUT7/GZMM/IL1R1/TNFRSF1B/KLRC1/RFTN1/CD70/NLRP3/RAET1L/ICAM1/CD
ZP4/C4BPA/FCGR2B/CFH/NOD2/C4BPB/SVEP1/TREM2/SERPING1/SPNS2/LTA/A2M/SPINK5/IL1B/CR1L/TNF/PTPN6/C3/PTPRC/CCR7/CR1/FCER2/CXCL
CLEC7A/IFNG/TYROBP/LAG3/SYK/NCR3/CD1C/IL12B/CD226/IL18RAP/KLRC1/RAET1L/SPI1/KLRC4/CRTAM/PRF1/IL12RB1/CD1D/IL12A/RASGRP1/KLRD1/
CLEC7A/IFNG/SLAMF7/IL4/CLEC12B/FCGR2B/FCGR1A/TYROBP/CFH/SERPINB4/F2/LAG3/GSDMB/SYK/NCR3/CD1C/IGHG1/IL12B/CD226/APOL1/TGFB1/I
ZP4/CD209/IL21/HHLA2/IL4/CD274/TNFSF13B/VCAM1/CCR2/SYK/IL2/IL12B/PDCD1LG2/IL6/CD86/LGALS9/AIF1/CD70/JAK2/IL15/HAVCR2/CD80/EBI3/CD28/F
LILRB2/TSPAN32/PGLYRP2/PAWR/IFNB1/AXL/VSIR/IFNL1/ADORA2A/FCGR2B/CD274/TYROBP/ASCL2/LST1/LAG3/IL10/IL2/CD74/PDCD1LG2/PLA2G5/TNF
BATF/CD83/VSIR/IFNG/CD274/ASCL2/IL2/IL27/IL12B/JUNB/IL4R/CD86/LGALS9/NLRP3/CD80/CD28/PRKCQ/CCL19/IL2RA/FOXP3/SOCS1/IL12RB1/TBX21/JA
CYLD/NFAM1/IL7/SYK/IL10/IL2/SFRP1/SPI1/TLR9/SLAMF8/INHBA/NCKAP1L/BTK/IL4I1/CD27/PTPN6/INPP5D/IL2RG/CR1/CARD11/FCRL3/IKZF3
ZP4/CYLD/CD209/FOXA1/WNT10B/IL21/CD83/HHLA2/SERPINF2/PODXL/VSIR/IFNG/ITGA4/NODAL/IL4/CD274/PLPP3/VNN1/IL7/CHST4/TNFSF13B/VCAM1/
CCL3/EDAR/STAT3/IL22RA1/IFNK/GPR35/IL17RB/IL17RA/DUOX2/PYDC1/CXCL8/SH2B3/IL11RA/IFNA8/IL20RB/CRLF2/CCR1/LILRB2/CCL2/CCDC3/THPO/L
ARRB2/SH2D1B/IL21/NCR1/SLAMF7/CLEC12B/SERPINB4/LAG3/NCR3/IL12B/CD226/TGFB1/IL18RAP/LGALS9/KLRC1/PIK3R6/HAVCR2/KLRC4/LILRB1/CRT
AXL/CD83/HHLA2/CLCF1/CLEC7A/VSIR/IFNG/IL4/CD274/EPHB2/TYROBP/VNN1/IL7/TNFSF13B/VCAM1/CCR2/SYK/NOD2/IL10/IL2/IL12B/CD74/PDCD1LG2/I
CLEC12B/FCGR2B/TYROBP/SERPINB4/LAG3/NCR3/CD1C/IL12B/CD226/TGFB1/IL18RAP/LGALS9/KLRC1/PIK3R6/RAET1L/ICAM1/HAVCR2/SPI1/KLRC4/LIL
ARRB2/IFNB1/C4BPA/CLEC12B/FCGR2B/SERPINB4/CCR2/NOD2/C4BPB/SVEP1/TGFB1/LGALS9/KLRC1/CD300A/HAVCR2/CD80/SPI1/LILRB4/LILRB1/FOX
SLAMF7/CLEC12B/FCGR2B/FCGR1A/TYROBP/SERPINB4/F2/LAG3/NCR3/CD1C/IGHG1/IL12B/CD226/TGFB1/IL18RAP/GZMM/LGALS9/KLRC1/PIK3R6/RAE
SLIT3/CCL1/CCR9/CCL3/GPR35/ZC3H12A/CXCL8/SH2B3/CCR1/CCL2/THPO/PPBP/CCL7/EDN1/CCL25/CCL14/SLIT2/CCL4/CCR2/CCL22/CCL24/CCR8/CC
SLIT3/CCL1/CCR9/CCL3/GPR35/ZC3H12A/CXCL8/SH2B3/CCR1/CCL2/THPO/PPBP/CCL7/EDN1/CCL25/CCL14/SLIT2/CCL4/CCR2/CCL22/CCL24/CCR8/CC
NFKBIZ/GPSM3/CCL1/CCL3/IL16/IL17RA/TLR3/MEFV/LBP/AGTR1/LDLR/TLR2/IL21/NMI/HYAL2/TNFRSF11A/PTGS2/IFNG/CCN4/LILRA5/FCGR1A/CREB3L
LILRB2/TSPAN32/ZP4/CD209/PAWR/IL21/HHLA2/LMO1/VSIR/IL4/CD274/TNFSF13B/VCAM1/CCR2/SYK/IL10/IL2/IL27/IL12B/PDCD1LG2/PLA2G5/IL6/CD86/TN
IFNB1/C4BPA/FCGR2B/ASCL2/NOD2/C4BPB/SVEP1/IL2/IL27RA/IL4R/KLRC1/HAVCR2/CD80/LILRB4/LILRB1/IL1RL1/FOXP3/CR1L/TBX21/JAK3/KLRD1/NCK
VSIR/IFNG/IFNL1/ADORA2A/ITGA4/IL4/CD274/CCL25/VNN1/ASCL2/IL7/CHST4/TNFSF13B/VCAM1/CCR2/LAG3/SYK/NOD2/IL10/IL2/PTAFR/IL12B/CD74/PD
CD83/IFNG/CCR2/SYK/IL12B/ZBTB16/IL4R/CD86/LGALS9/NLRP3/JAK2/CD80/LILRB4/EBI3/CD28/PRKCQ/ZAP70/CCL19/FOXP3/SOCS1/IL12RB1/IL12A/NCK
ARID5A/IL4/FCGR1A/LAG3/NCR3/NOD2/CD1C/IL2/IL12B/CD226/TREM2/IL6/TGFB1/IL18RAP/IL1R1/KLRC1/NLRP3/RAET1L/RSAD2/LTA/KLRC4/CRTAM/CD
IFNB1/C4BPA/FCGR2B/ASCL2/NOD2/C4BPB/SVEP1/IL2/IL27RA/IL4R/SAMSN1/KLRC1/HAVCR2/CD80/LILRB4/LILRB1/IL1RL1/FOXP3/CR1L/TBX21/JAK3/KLR
VSIR/IFNG/IFNL1/ADORA2A/ITGA4/IL4/CD274/CCL25/VNN1/ASCL2/IL7/CHST4/TNFSF13B/VCAM1/OLR1/CCR2/LAG3/SYK/NOD2/IL10/IL2/PTAFR/JAM2/IL12
CYLD/PAWR/FOXP1/GCSAML/NFAM1/FCGR2B/LIPA/LPXN/CD226/GCSAM/CD300A/THY1/LYN/PLCL2/LILRB4/SLA2/UBASH3A/PTPN22/LCK/TRAT1/FCMR
VSIR/IFNL1/ADORA2A/FCGR2B/CD274/TYROBP/ASCL2/LST1/LAG3/IL10/IL2/CD74/PDCD1LG2/PLA2G5/TNFAIP8L2/IL4R/MERTK/SAMSN1/CD86/LGALS9/S
SLC46A2/DHX58/ARRB2/CACNB4/CYLD/BTLA/PAWR/TLR2/FOXP1/HSPA1B/LCP2/NMI/HHLA2/GCSAML/FYB2/NFAM1/REG3G/TLR1/CLEC7A/TIFA/IRGM/
SLIT3/CCL1/CCR9/CCL3/GPR35/CXCL8/SH2B3/CCR1/CCL2/THPO/PPBP/CCL7/EDN1/CCL25/CCL14/SLIT2/CCL4/CCR2/CCL22/CCL24/CCR8/CCRL2/TRE
SLC46A2/DHX58/ARRB2/CACNB4/CYLD/PAWR/TLR2/FOXP1/HSPA1B/LCP2/NMI/HHLA2/GCSAML/FYB2/NFAM1/REG3G/TLR1/CLEC7A/TIFA/IRGM/OAS3/
LILRB2/CCL2/TSPAN32/SLC46A2/ZP4/CYLD/CD209/PAWR/IFNB1/WNT10B/IL21/BATF/CD83/HHLA2/LMO1/VSIR/IFNG/IFNL1/ADORA2A/IL4/CD274/VNN1/A
IL21/FOXP1/RELB/BATF/CD83/IFNG/ASCL2/SEMA4A/SYK/IL2/IL27/SOCS3/IL12B/PTGER4/ZNF683/JUNB/IL6/FUT7/ZBTB16/IL4R/EOMES/CD86/LGALS9/NLR
ZC3H12A/INS/IL20RB/PGLYRP2/ARRB2/IFNB1/C4BPA/AXL/APOA1/IFNL1/CLEC12B/FCGR2B/ASCL2/SERPINB4/ZPBP2/CCR2/NOD2/C4BPB/IL10/SVEP1/IL2
PGLYRP2/SLC46A2/CYLD/IFNB1/WNT10B/BATF/AXL/CD83/NFAM1/VSIR/IFNG/IFNL1/IL4/VNN1/ASCL2/IL7/CCR2/LAG3/SYK/IL10/IL2/IL27/IL12B/CD74/ZNF6
ZP4/CYLD/CD209/WNT10B/IL21/CD83/HHLA2/VSIR/IFNG/IL4/CD274/VNN1/IL7/TNFSF13B/VCAM1/CCR2/SYK/NOD2/IL2/IL12B/CD74/PDCD1LG2/EFNB2/IL3
IFNW1/PAWR/IFNB1/IL21/IFNA4/CLCF1/IL9/IFNA1/MZB1/IL4/FCGR2B/EPHB2/TYROBP/IFNA6/IL7/TNFSF13B/IFNA14/IL10/IL2/CD74/TNFRSF4/CD70/MNDA
SYK/SOCS3/IL12B/CD74/CD4/IL6/IL15/CD28/ZAP70/FOXP3/IL12RB1/CD1D/DOCK2/TBX21/TOX/CD3E/SPN/GATA3/CD3D/LY9/THEMIS/PTPRC/CCR7/BCL11
AXL/CD83/HHLA2/CLCF1/CLEC7A/VSIR/IFNG/IRGM/IL4/CD274/EPHB2/TYROBP/VNN1/IL7/TNFSF13B/VCAM1/CEBPA/CCR2/THBS1/SYK/NOD2/IL10/IL2/IL1
AXL/CD83/HHLA2/CLCF1/CLEC7A/VSIR/IFNG/IRGM/LILRA5/IL4/CD274/EPHB2/TYROBP/VNN1/IL7/TNFSF13B/VCAM1/CEBPA/CCR2/THBS1/SYK/NOD2/IL1
PGLYRP2/RORC/SH2D1B/IFNW1/IFNB1/IL21/FOXP1/NMI/IFNA4/RELB/BATF/CLCF1/IFNA1/IFNG/IFNL1/LFNG/DOCK11/IL4/FCGR2B/TYROBP/IFNA6/ASCL
PGLYRP2/RORC/SH2D1B/IFNW1/IFNB1/IL21/FOXP1/NMI/IFNA4/RELB/TCIM/BATF/CLCF1/IFNA1/IFNG/IFNL1/LFNG/DOCK11/IL4/FCGR2B/TYROBP/IFNA6
CLEC7A/ARID5A/IL4/FCGR1A/TYROBP/LAG3/NCR3/NOD2/CD1C/IL2/PTAFR/IL12B/CD226/TREM2/IL6/TGFB1/IL18RAP/IL1R1/KLRC1/NLRP3/RAET1L/RSAD

VSIR/IFNG/ITGA4/IL4/CD274/VNN1/IL7/CHST4/TNFSF13B/VCAM1/CCR2/SYK/NOD2/IL2/PTAFR/IL12B/CD74/PDCD1LG2/EFNB2/IL36B/CD4/IL6/FUT7/ZBTB1

IL21/FOXP1/RELB/BATF/CD83/IFNG/ASCL2/SEMA4A/IL2/IL27/SOCS3/IL12B/PTGER4/JUNB/IL6/FUT7/IL4R/CD86/LGALS9/NLRP3/STAT4/GPR183/BCL3/CD8

MZB1/CLEC7A/ARID5A/IFNG/IL4/EPHB2/FCGR1A/TYROBP/CARD9/IRF5/CCR2/LAG3/SYK/NCR3/NOD2/DDX60/CD244/IL10/CD1C/IL2/PTAFR/IL12B/CD74/

IFNK/MASP1/KLK7/DEFB1/CXCL8/LEAP2/IFNA8/CCL2/C4B/ZP4/IFNW1/RNASE3/HRG/IFNB1/PPBP/C1RL/C4BPA/CFP/ST6GAL1/IFNA4/CD83/IFNA1/REG3G

RORC/SLC46A2/CYLD/IFNB1/WNT10B/IL21/FOXP1/MAFB/RELB/BATF/CD83/VSIR/IFNG/IFNL1/LFNG/IL4/FZD8/VNN1/ASCL2/IL7/SEMA4A/LIPA/CCR2/LAG3

IL21/FOXP1/RELB/BATF/CD83/VSIR/IFNG/CD274/ASCL2/SEMA4A/IL2/IL27/SOCS3/IL12B/PTGER4/JUNB/IL6/FUT7/IL4R/CD86/LGALS9/NLRP3/STAT4/GPR18

CLEC7A/ARID5A/IFNL1/HMHB1/CCR2/CD244/IL2/IL27RA/IL27/TLR8/IL12B/CD226/ISG15/IL1R1/TXK/LGALS9/WNT5A/JAK2/BCL3/HAVCR2/LTA/EBI3/LILRB1/

BATF/AXL/CD83/HHLA2/CLCF1/LMO1/NFAM1/MZB1/CLEC7A/VSIR/IFNG/IFNL1/ADORA2A/IL4/FCGR2B/CD274/EPHB2/TYROBP/VNN1/ASCL2/IL7/TNFSF1

KLRF2/STAT3/ST3GAL1/IFNK/ZC3H12A/C17orf99/IFNA8/PGLYRP2/RORC/IFNW1/IFNB1/IL21/FOXP1/IFNA4/RELB/BATF/CLCF1/IFNA1/IFNG/IFNL1/LFNG/D

ZP4/CD209/PAWR/IL21/ST6GAL1/HHLA2/CLCF1/LMO1/MZB1/VSIR/IL4/FCGR2B/CD274/EPHB2/TYROBP/CSF1/IL7/TNFSF13B/LST1/VCAM1/CSF2RA/CCR2

C1RL/C4BPA/C2/FCGR2B/C1QA/NOD2/C4BPB/SVEP1/IGHG1/C1QB/IGHG3/TREM2/C1QC/SERPING1/C1S/IGHE/BCL3/C7/LTA/C1R/CR1L/FCMR/TNF/IGH

BATF/CD83/VSIR/IFNG/ADORA2A/CD274/ASCL2/CCR2/SYK/IL2/IL27/IL12B/ZNF683/JUNB/ZBTB16/IL4R/CD86/LGALS9/NLRP3/CD300A/JAK2/CD80/LILRB4/

IFNW1/CYLD/IFNB1/IL21/IFNA4/CLCF1/IL9/IFNA1/NFAM1/LFNG/ITGA4/DOCK11/IL4/FCGR2B/IFNA6/IL7/TNFSF13B/VCAM1/IL11/IFNA14/SYK/IL10/IL2/DLL1/

CLEC7A/IFNG/IL4/CLEC12B/FCGR2B/TYROBP/CFH/SERPINB4/LAG3/SYK/NCR3/CD1C/IL12B/CD226/TGFB1/IL18RAP/LGALS9/KLRC1/PIK3R6/RAET1L/ICA

ELF4/ZP4/IFNW1/CD209/SH2D2A/PAWR/IFNB1/IL21/ST6GAL1/IFNA4/HHLA2/CLCF1/IL9/LMO1/IFNA1/MZB1/VSIR/IL4/FCGR2B/CD274/EPHB2/TYROBP/CS

ZP4/CD209/PAWR/IL21/ST6GAL1/HHLA2/CLCF1/LMO1/MZB1/VSIR/IL4/FCGR2B/CD274/EPHB2/TYROBP/CSF1/IL7/TNFSF13B/LST1/VCAM1/CCR2/SYK/IL1

HHLA2/CLCF1/LMO1/MZB1/VSIR/IL4/FCGR2B/CD274/EPHB2/TYROBP/IL7/TNFSF13B/LST1/VCAM1/CCR2/SYK/IL10/IL2/IL27/IL12B/CD74/PDCD1LG2/PLA2G

CYLD/PAWR/IL21/CLCF1/NFAM1/MZB1/IL4/FCGR2B/EPHB2/TYROBP/IL7/TNFSF13B/SYK/NOD2/IL10/IL2/IL27RA/CD74/IL6/TGFB1/SAMSN1/TNFRSF4/SFR

ELF4/ZP4/IFNW1/CD209/SH2D2A/PAWR/IFNB1/IL21/ST6GAL1/IFNA4/HHLA2/CLCF1/IL9/LMO1/IFNA1/MZB1/VSIR/IL4/FCGR2B/CD274/EPHB2/TYROBP/CS

ELF4/ZP4/IFNW1/CD209/SH2D2A/PAWR/IFNB1/IL21/IFNA4/HHLA2/CLCF1/IL9/LMO1/IFNA1/MZB1/VSIR/IL4/FCGR2B/CD274/EPHB2/TYROBP/IFNA6/IL7/TN

IFNW1/CYLD/PAWR/IFNB1/IL21/IFNA4/BATF/CLCF1/IL9/IFNA1/NFAM1/MZB1/LFNG/ITGA4/DOCK11/IL4/FCGR2B/EPHB2/TYROBP/IFNA6/IL7/TNFSF13B/V

CLEC7A/ARID5A/IL4/FCGR2B/CD274/EPHB2/FCGR1A/CARD9/ASCL2/TNFSF13B/CCR2/NOD2/C4BPB/IL10/SVEP1/CD1C/ADCY7/IL2/IL27RA/IL27/IL12B/CD

PGLYRP2/DHX58/ARRB2/SH2D1B/IFNB1/IL21/NCR1/C4BPA/CFP/BATF/AXL/CLCF1/MZB1/APOA1/CLEC7A/ARID5A/IFNG/IFNL1/IL4/CLEC12B/FCGR2B/EP

IFNB1/C4BPA/BATF/CLCF1/CLEC7A/ARID5A/IL4/FCGR2B/CD274/EPHB2/FCGR1A/CARD9/ASCL2/TNFSF13B/CCR2/NOD2/C4BPB/IL10/SVEP1/CD1C/IL2/IL

PGLYRP2/RORC/SLC46A2/IFNW1/CYLD/PPARG/IFNB1/WNT10B/IL21/FOXP1/MAFB/IFNA4/RELB/BATF/AXL/CD83/CLCF1/IL9/IFNA1/NFAM1/TRPM2/VSIR/

CLEC7A/VSIR/ARID5A/IFNL1/IRGM/HMHB1/CD274/CCR2/NOD2/CD244/IL10/IL2/IL27RA/IL27/TLR8/IL12B/PDCD1LG2/CD226/ISG15/IL1R1/TXK/LGALS9/WN

CLEC7A/VSIR/ARID5A/IFNL1/IRGM/HMHB1/CD274/CCR2/NOD2/CD244/IL10/IL2/IL27RA/IL27/TLR8/IL12B/PDCD1LG2/CD226/ISG15/IL1R1/TXK/LGALS9/WN

PGLYRP2/RORC/SLC46A2/IFNW1/CYLD/IFNB1/WNT10B/IL21/FOXP1/MAFB/IFNA4/RELB/BATF/AXL/CD83/CLCF1/IL9/IFNA1/NFAM1/VSIR/IFNG/IFNL1/LFN

ARRB2/SH2D1B/IFNB1/IL21/NCR1/C4BPA/CLCF1/ARID5A/IL4/CLEC12B/FCGR2B/FCGR1A/SERPINB4/CCR2/LAG3/NCR3/NOD2/C4BPB/IL10/SVEP1/CD1C

CLEC7A/ARID5A/IL4/CLEC12B/FCGR2B/FCGR1A/TYROBP/SERPINB4/CCR2/LAG3/SYK/NCR3/NOD2/C4BPB/IL10/SVEP1/CD1C/IL2/IL27RA/PTAFR/IL12B/C

CACNB4/CYLD/PAWR/TLR2/FOXP1/LCP2/NMI/HHLA2/GCSAML/FYB2/NFAM1/TLR1/CLEC7A/FCGR2B/TYROBP/C3AR1/FYB1/LIPA/SYK/NCR3/NOD2/IGH

IL21/FOXP1/RELB/BATF/CD83/VSIR/IFNG/ADORA2A/CD274/ASCL2/CLEC4A/SEMA4A/CCR2/SYK/IL2/IL27/SOCS3/IL12B/PTGER4/ZNF683/JUNB/IL6/FUT7/Z

CACNB4/CYLD/BTLA/PAWR/TLR2/FOXP1/LCP2/NMI/HHLA2/GCSAML/FYB2/NFAM1/TLR1/CLEC7A/CLEC12B/FCGR2B/FCGR1A/TYROBP/C3AR1/FYB1/LI

FCGR2A/C1RL/C4BPA/BATF/CLCF1/IL9/C2/IL4/FCGR2B/FCGR1A/CARD9/C1QA/NOD2/C4BPB/IL10/SVEP1/IGHG1/IL2/IL27RA/C1QB/TLR8/CD74/IGHG3/IG

CACNB4/CYLD/PAWR/FOXP1/LCP2/HHLA2/GCSAML/FYB2/NFAM1/FCGR2B/FYB1/LIPA/SYK/IGHG1/LPXN/IGHG3/IGKC/CD226/NFKBIA/TXK/PRAM1/RFT

SYK/IGHG1/LPXN/IGHG3/IGKC/NFKBIA/RFTN1/CD79B/MNDA/GCSAM/CD300A/IGHE/LYN/PLCL2/BMX/PTPN22/CD38/NFATC2/LCK/KLHL6/PLCG2/NCKAP

FCGR2A/C1RL/C4BPA/BATF/CLCF1/IL9/C2/IL4/FCGR2B/FCGR1A/CARD9/C1QA/NOD2/C4BPB/IL10/SVEP1/IGHG1/IL2/IL27RA/C1QB/TLR8/CD74/IGHG3/IG

FCGR2A/IFNB1/IL21/NCR1/C1RL/C4BPA/SCART1/BATF/CLCF1/IL9/C2/CLEC7A/ARID5A/SLAMF7/IL4/CLEC12B/FCGR2B/FCGR1A/TYROBP/CARD9/SERP

FCGR2A/IFNB1/C1RL/C4BPA/SCART1/RELB/BATF/CLCF1/IL9/C2/CLEC7A/ARID5A/BACH2/IL4/FCGR2B/CD274/EPHB2/FCGR1A/CARD9/ASCL2/TNFSF13

SH2D1B/IFNW1/CD209/BTLA/FCGR2A/IFNB1/C1RL/C4BPA/SCART1/TNFRSF11A/IFNA4/RELB/FCAMR/BATF/CLCF1/IL9/IFNA1/C2/CLEC7A/ARID5A/IFNG/

FCGR2A/IFNB1/IL21/NCR1/C1RL/C4BPA/SCART1/BATF/CLCF1/IL9/C2/ARID5A/SLAMF7/IL4/CLEC12B/FCGR2B/FCGR1A/CARD9/SERPINB4/C1QA/CCR2/

;/MTERF3/NOP16/RPP40/C1QBP/NOL7/USP36/TFB2M/NSUN4/DIS3/UTP14A/SBDS/LAS1L/TRAF7/DHX29/DDX49/RRP15/EXOSC
TP15/DROSHA/SNU13/MRM2/ERI2/RRP9/HEATR1/EXOSC4/RRP36/TSR3/DDX27/NSA2/RRP1/SUV39H1/NOL10/ABT1/DDX10/TS
LE1/DDX56/MYBBP1A/UTP11/PRMT5/BRIX1/PRPF6/NOP2/AAR2/WDR77/DDX51/DENR/DIMT1/EIF3G/ESF1/PA2G4/SNRPC/DNT
10/UQCRQ/COX7A2L/NDUFS3/NDUFB4/NDUFA2/NDUFB6/COX7A2/NDUFV2/DGUOK/DLD
10/UQCRQ/COX7A2L/NDUFS3/NDUFB4/NDUFA2/NDUFB6/COX7A2/NDUFV2/DGUOK/DLD
SUN4/DIS3/UTP14A/SBDS/LAS1L/DDX49/RRP15/EXOSC5/UTP15/DROSHA/GTF3C1/SNU13/MRM2/ERI2/RRP9/HEATR1/EXOSC4
NDUFB4/NDUFA2/NDUFB6/COX7A2/NDUFV2
A1/UQCC2/COX8A/ATP5ME/UQCR11/NDUFA11/CDK1/NDUFA6/NDUFS8/NDUFB11/NDUFB2/NDUFV1/NDUFB3/COQ9/STOML
7A/PES1/TARBP2/NOP56/DDX1/NHP2/LAGE3/DDX56/ELP2/UTP11/WDR4/BRIX1/NOP2/DDX51/DIMT1/ESF1/PA2G4/PUSL1/RPUS
2/COX6C/MDH1/IDH3A/UQCRC1/NDUFA8/NDUFA7/NDUFS2/DLAT/ACLY/MSH2/NDUFS1/COX6A1/UQCC2/COX8A/ATP5ME/U

PET100/COA4/NDUFB4/NDUFA2/SDHAF4/NDUFB6/CHCHD7/SCO1/NDUFAF8/NDUFAF3/ECSIT/SDHAF1/STMP1/NDUFAF1/UC
N2/NCAPH/NUDC/POLDIP2/CHMP1B/ZWINT/CHMP2A/RAD21/CHMP3/SEH1L/NCAPD2/SKA1/KIF22/RMDN1/UHRF1/BAZ1B/CD
B3/COQ9/NDUFS4/SLC25A22/UQCRC2/UQCC3/NDUFA10/UQCRQ/AIFM2/COX7A2L/NDUFS3
3/FTSJ1/TSEN2/METTL1/SARS2/FARSA/TRUB2/DTD1/GTF3C3/ELP6/DUS1L/EXOSC2/TRIT1/RARS2/IARS2/GTF3C2/OSGEPL1/E
TRF1/MTIF3/FASTKD5/MTERF1/METTL8/CHCHD10/FOXO3/RPUSD4/COA3/MTPAP/PRKAA1/MPV17L2/TACO1/QRSL1/GATB/TS

/RAE1/RMDN1/TACC3/INTS13/UHRF1/SASS6/GOLGA2/CLASP2/TPR/WRAP73/CHEK2/CLASP1/TTK/PIBF1/MYBL2/CCSAP/MAP4

MAD2L1BP/ANAPC11/CDK1/PBRM1/NCAPG2/SMARCC1/RANGRF/DCTN2/NCAPH/NUDC/POLDIP2/CHMP1B/ZWINT/CHMP2A/

SP1/TTK/PIBF1/MYBL2/CCSAP/MAP4/INCENP
/SMARCD2/CSNK2A1/TPR/SMC6/TTK/INCENP/ZW10/SMC2/DYNC1LI1/HASPIN/BCL7B/SMARCB1/GEN1/KNTC1/NSMCE2/KAT

C30/NDUFC1/ANTKMT/ATP5MGL/NDUFA9/ATP5F1E/ATPSCKMT/NDUFB5
FR1/CYC1/SUCLG1/ATP5F1C/LYRM7/ARL2/ATP5PF/COX7C/MTFR1L/COX6B1/NDUFC2/COX6C/MDH1/IDH3A/UQCRC1/NDUFA

MT61B/CDK5RAP1/POP7/PUS1/RTCB/GRSF1/THUMPD2/TRUB1/MOCS3/FAM98A/TSEN54/ELP1/METTL8/KTI12/RTRAF/WDR6/
RHOT1/SLC25A46/TOMM5/PPIF/TOMM70/IMMT

2/CDC16/CCNB2/TOP2A/HJURP/KIF23/BOD1/BCL7C/MAD2L1/BIRC5/MSH5/DCAF13/SETDB2/SKA3/CENPJ/CENPL/BUB1B/CEN

D1/MAD2L1/BIRC5/SKA3/HOXA13/BUB1B/CHAMP1/TUBG1/MZT1/CENPE/PRMT5/KNL1/REEP4/UBE2I/SKA2/SMC1A/BANF1/SPD

E2F1/ORC1/BLM/MRE11/TIMELESS/CDT1/LCMT1/WDR76/RHNO1/MBTPS1/BRCA1/CASP2/TPR/CUL4A/TRIAP1/CRADD/CHEK
RE1/FMN2

IE/LSM5/RBM20/LSM2/DDX20/SNRNP70/SCNM1/C1QBP/PPIL1/PRDX6/PTBP2/SAP18/HSPA8/RNPS1/DHX9/HABP4/ZPR1/NOL3/

1A/AGO1/ABT1/SF3A3/SF3B6/DDX39B/SNRNP200/PUF60/MCTS1/CRNKL1/DHX30/TGS1/POLR2D/SNRPF/RPF2/KHDC4/PRPF

7/HMGA1/POLD1/WDHD1/ZNF365/CHTF18/BRCA1/ORC4/RTF2/RPA1/RECQL4/CCNE2/REV3L/WDR18/SSBP1/DBF4B/LIG1/TR

H2/ALKBH7/RTL10/TOMM6/PDCD5/TIMM21/MTX1/BNIP3L/SLC25A22/AIFM1/PMPCB/SLC25A1/ABCB7/CPT2/SFXN4/SLC25A40/

ENP/ZW10/FBXO31/RPA2/DYNC1LI1/BABAM1/HASPIN/GEN1/MUS81/KNTC1/CHEK1/MDM2/IER3/NAE1/DONSON/ANAPC15/T

F3/CLNS1A/AGO1/ABT1/SF3A3/SF3B6/DDX39B/SNRNP200/PUF60/MCTS1/CRNKL1/DHX30/TGS1/POLR2D/SNRPF/RPF2/KHDC

3/BUB1B/RCC2/CHAMP1/TUBG1/MZT1/SUN1/CENPE/CCNE1/NDC1/PMF1/EHMT2/KNL1/SKA2/BRIP1/SMC1A/SMARCA4/ECT

LHL22

D/NDUFA10/NME6/UCKL1/ATP5MC2/NDUFS3

1/TRMT10A/TPRKB/QTRT2/TRMT10C/TRMT1L/THG1L/TRMU/CDKAL1/OSGEP/TYW5/BCDIN3D

1/SNRPF/TRMT112/METTL2A/PRMT6/MECP2/RAMAC/RBM15B/RAB6A/TRMT61A/VIRMA/ETFBKMT/HSD17B10/TRMT61B/FBL

TN6/RAE1/SAC3D1/RMDN1/TACC3/INTS13/UHRF1/SASS6/DLG1/RNF4/GOLGA2/CLASP2/CHD3/STARD9/TPR/WRAP73/CHEK2

N1/DCTN1/BRCA2/CHEK1/BBS4/KIAA0753/ARHGEF10/HAUS4/ROCK2

NAPC15/RB1/PSMG2/BECN1/BRD7/ARID2/NUMA1/KLHL22/RMI2/ZWILCH/ANAPC5/SMARCE1/SMARCD1/ACTL6A/CDK5RAP2/

S4/UQCC3/ATP5F1D/NDUFA10/NME6/UCKL1/ATP5MC2/NDUFS3

PG/SF3B4/BUD31/PRPF3/PPIH/CLNS1A/HNRNPK/LSM7/CASC3/THRAP3/RNF113A/SF3A3/SF3B6/DDX39B/SNRNP200/UPF3A/PI

V11/HAUS3/MCIDAS/SSX2IP/KIF11/NDE1/GEN1/DCTN1/BRCA2/CHEK1/BBS4/KIAA0753/ARHGEF10/HAUS4/ROCK2

34/BUD31/PRPF3/PPIH/CLNS1A/HNRNPK/LSM7/CASC3/THRAP3/RNF113A/SF3A3/SF3B6/DDX39B/SNRNP200/UPF3A/PRPF40B/

34/BUD31/PRPF3/PPIH/CLNS1A/HNRNPK/LSM7/CASC3/THRAP3/RNF113A/SF3A3/SF3B6/DDX39B/SNRNP200/UPF3A/PRPF40B/

SD3/LSM5/RBM20/LSM2/DDX20/SNRNP70/SCNM1/C1QBP/PPIL1/PRDX6/PTBP2/SAP18/HSPA8/RNPS1/DHX9/HABP4/ZPR1/NOL3

SPDL1/MAD2L1BP/ANAPC11/CDK1/PBRM1/NCAPG2/SMARCC1/SMG5/NCAPH/TFPT/ZWINT/MAPK1/NSMCE1/ANAPC1/TCP1/R

3/SNRPD3/RBM15/TRMT10A

ASEH1/LRWD1/TFPT/NUCKS1/TWNK/RFC4/POLE/WRN/MCM3/MCM8/MCM10/MEAF6/ESCO1/ORC5/PCNA/RTEL1/ORC1/DTI

6/NDUFS8/ETFB/NDUFB2/P4HA2/CIAPIN1/NDUFV1/NDUFB3/COQ9/NDUFS4/SLC25A22/UQCRC2/POR/UQCC3/GLRX2/NDUFA

C2/NDUFS3

C2/NDUFS3

29/TIMM17A/TOMM5/BCAP31/TOMM34/TOMM70

2CD3/CEP192/KAT2A

IN/OBSL1/PSMC3IP/CDC25C/CDCA5/ESPL1/KIF18B/EPS8/SPAG5/TOM1L2/CEP85/KIF18A/RRS1/KIF14/CENPI/RAD51/CDC16/CC

0AA1/TIMM9/TIMM29/TIMM17A/TOMM5/BCAP31/TOMM34/TOMM70/TOMM40L/AP3B1/BAP1/NPEPPS/MAIP1

JFB6/NDUFV2/ATP5MC1/PARP1/ATP6V0C

ZC3H3/INTS9/ELAC1/EXOSC3/EXOSC9/DHX36

NTS3/NUCKS1/WRN/RAD51C/EP400/MCM3/MCM8/NSMCE1/FIGNL1/RAD21/MEAF6/MRGBP/SPIDR/RTEL1/SPIRE1/RAD52/ZS\

NK1/CTTN/TIMM10/AFG3L2/NDUFAF2/DNAJA3/COA1/MSTO1/NDUFA1/AKT1/STPG1/MGME1/NDUFB10/DNAJC11/BCL2L2/MAI

4/MTCH1/BCS1L/HSP90AA1/TIMM9/TIMM29/TIMM17A/TOMM5/BCAP31/TOMM34/TOMM70/TOMM40L/AP3B1/BAP1/NPEPPS/MA

0/TESMIN/OBSL1/PSMC3IP/DDHD2/CDC25C/CDCA5/ESPL1/KIF18B/EPS8/SPAG5/TOM1L2/CEP85/KIF18A/RRS1/KIF14/CENPI/R

/RPA1/RIF1/XRCC5/RECQL4/CHD4/USP51/REV3L/FUS/PPP4R2/FAN1/RPA2/XRCC6/RNF138/PARPBP/SWI5/EXD2/PPP4C/GEN

TRIP/NAE1/DONSON/INTS7/DNA2/TAOK3/MBTPS2/RAD1/BABAM2/PRPF19/GNB1L/SYF2/TTI2/ABRAXAS1/GIGYF2/MAPK14/RFW

/UQCRB/COA6/TRAP1/SDHA/MYBBP1A/NDUFB7/MTFR1/CYC1/SUCLG1/ATP5F1C/LYRM7/ARL2/ATP5PF/COX7C/MTFR1L/CC

IF1/RECQL4/CHD4/USP51/REV3L/FUS/PPP4R2/FAN1/RPA2/RNF138/PARPBP/SWI5/EXD2/PPP4C/GEN1/KDM1A/MUS81/BRCA2

JSP51/FUS/PPP4R2/RPA2/TFIP11/FMN2/BABAM1/PARPBP/TAF6L/BCL7B/USP1/PPP4C/SMARCB1/TMEM161A/KDM1A/PARP1/C

ED4/LEO1/NELFA/MED21/MED10/CCNT1/MED19/MED9/ERCC3/MED16/NELFB/RNF168/CDC73/TSFM/PCID2/PWWP2B/POLR1I

E2/ERCC1/HSP90AA1/NAT10/RPA2/TFIP11/XRCC6/TELO2/TNKS1BP1/UPF1/FBXO4/BRCA2/PARP1/DOT1L/NSMCE2/MCRS1/MA

AM2/PRPF19/GNB1L/SYF2/TTI2/ABRAXAS1/GIGYF2/MAPK14/RFWD3/TAOK1/CCND1/FZR1/RAD17/TOPBP1/TTI1/TICRR/FOXC

SP90AA1/NAT10/RPA2/TFIP11/XRCC6/TELO2/TNKS1BP1/UPF1/FBXO4/BRCA2/PARP1/NSMCE2/MCRS1/MAP2K7

A7/NDUFS2/DLAT/ACLY/ACOT7/NDUFS1/ATP5ME/NDUFA11/NDUFA6/NDUFS8/PANK2/NDUFB11/NDUFB2/PANK4/NDUFV1/ND

A7/NDUFS2/DLAT/ACLY/ACOT7/NDUFS1/ATP5ME/NDUFA11/NDUFA6/G6PD/NDUFS8/PANK2/NDUFB11/NDUFB2/PANK4/NDU

S1/FZD5/ULK3/MAP4/TRNT1/ATG2A/LTN1/TSC2/TCF25/TEX264/FBXL4/FAF2/RB1CC1/WDR45/RNF25/KAT5/SPATA33/CISD2/Z

GAP1/EXOC2/SNX18/USP8/LZTS2/CHMP6/CHMP1A/NUP62/RHOA

HGEF10/HAUS4/AAAS/TUBGCP4/WASHC5/HAUS2/NUMA1/ABRAXAS1/TUBGCP3/PTEN/LZTS2/CHMP6/CHMP1A/RHOA/CSNK

JFS1/ATP5ME/NDUFA11/NDUFA6/NDUFS8/PANK2/NDUFB11/NDUFB2/PANK4/NDUFV1/NDUFB3/ELOVL1/STOML2/NDUFS4/A

/PFDN4/PPIH/PPIB/UNC45A/HSPB1/PDIA4/VBP1/TBCB/PPIL2/NUDCD3/PDRG1/CCDC47/P4HB/FUT10/DNAJC21/CSNK2A1/DN[illegible]

OHB/UPF1/NCBP1/XPOT/NUP155/NMD3/NUP188/NUP42/CHTOP/ALYREF/UHMK1/MDM2/AGFG1/NUP107/KHDRBS1/RANBP3/

L/GAR1/TRIT1/OSGEPL1/TGS1/SNRPF/BAG4/TRMT112/METTL2A/RAMAC/CTU2/PARN/RBM15B/TRMT61A/RPUSD2/NAT10/DU

KN1A

EN/CRADD/CHEK2/TTK/INCENP/ZW10/FBXO31/RPA2/DYNC1LI1/BABAM1/HASPIN/GEN1/KNTC1/CHEK1/MDM2/IER3/NAE1/DO

1/TP53/MSH2/FANCD2/RINT1/SPDL1/MAD2L1BP/CDK1/EIF2AK4/RBBP8/ZFYVE19/INTS3/PTPN11/ZWINT/CHMP2A/CDC14B/RA

FYVE19/INTS3/PTPN11/ZWINT/CDC14B/RAD21/CRY1/E2F1/ORC1/BLM/DLG1/MRE11/TIMELESS/CDT1/LCMT1/WDR76/RHNO1/

CAP29/RAB34/TMED4/EXOC6B/SPIRE1/SPTBN1/SCAMP3/YIF1A/LMAN2/GOLGA2/AP2A1/P4HB/COPG2/RAB1B/ARFRP1/PDCD6/

ARS2/ELP2/TRIM2/EARS2/DENR/EIF3G/PA2G4/AKT2/DUS3L/CAPRIN2/NSUN5/RPUSD3/RRBP1/MALSU1/WARS2/MTFMT/C8orf

DD/CHEK2/MDC1/CCAR2/EEF1E1/FBXO31/RPA2/TELO2/BABAM1/SESN2/FBXO4/MSX1/KDM1A/MUS81/ATAD5/BRCA2/CHEK1/

ЗLU/TMED1/COG2/VMP1/TRAPPC8/TRAPPC12/TJAP1/BLZF1/YIPF7/LMAN1/COG4/DYM/PRKD1/ZFP69B/COG1/YIPF5/GCC2/MA

11/FAM168A/BRD7/ARID2/ZCWPW1/ABRAXAS1/TIGAR/EYA3/UBE2N/SMARCE1/SMARCD1/ACTL6A/MORF4L1/EPC2/RNF168/A
RD8/NUDT16L1/RUVBL1/KLHL15/RAD51AP1/NSD2/DDX11/BRD7/ARID2/ZCWPW1/FBH1/UBE2N/RMI2/POLQ/SMARCE1/PPP4R

ARF4/CREB3L2/GOLT1B/SEC23B/NRBP1/YIPF7/ERGIC2/TRAPPC13/LMAN1/LMAN2L/TEX261/RNF139/SEC23A/YIPF5/SEC24D/C

2L1/HAUS3/DCTN1/BORA/HAUS4/TRAF3IP1/EFNA5/BICD2/PHLDB2/GIT1/HAUS2/DYRK1A/DIXDC1/CLIP1/NAV3/NUMA1/CHMP

N2/CAV1/MAN1B1/RNF139/YOD1/EDEM1/STT3B/TMEM67/DERL2/WFS1/SELENOS/AQP11/TMX1/RNFT2/UBE4A/EDEM2/DERL
TOP/ALYREF/AGFG1/NUP107/UPF3B/BICD2/THOC1/AAAS/NCBP3/THOC7/ENY2/NCBP2/NUTF2/SRSF1/NUP43/NUP62/FYTTD

/TAOK1/CCND1/FZR1/RRM2B/RAD17/TOPBP1/TICRR/CDK6/FOXO4
F4E/DDX19B/PCID2/FMR1/XPO1/NXF1
E1/SPTBN1/PIN1/GIPC1/ORC4/TMEM250/RHOC/INCENP/KIF13A/ENTR1/STMN1/FMN2/CSPP1/PRPF40A/CFL1/EXOC8/BRCA2/
C73/PCID2/PWWP2B/GLYR1/SIRT6/PAF1/ZMYND8/EAF1/INTS9/MED26/MED28/MED1/MED24

RPB/FTSJ1/SNRPG/SMYD5/PRMT1/METTL1/MBD3/PCIF1/MIS18A/SUV39H1/SUV39H2/METTL21C/SETD4/BMI1/LCMT1/TGS1/PF

G3BP2/NUP133/PARN/BICD1/RBM15B/NUP35/THOC2/MAGOH/NUP85/SLBP/NUP153/FBL/POLDIP3/MAGOHB/UPF1/NCBP1/XP

17/TOPBP1/TICRR/CDK6/FOXO4

/TUBGCP4/GIT1/HAUS2/DYRK1A/CLIP1/NAV3/NUMA1/TUBG2/TUBGCP3/TAOK1/HDGFL3/STMN2/CSNK1D

CC4/ARID1A/ACTB

JB1B/RCC2/LATS2/RAD50/CDK4/NABP2/CENPE/CCNE1/MIIP/CTDSPL/CCNG1/TFDP1/HUS1/KNL1/CDC34/DTL/CCNO/SKP2/SI

1/BRD4/NAE1/DONSON/TAOK3/MBTPS2/BABAM2/CCNQ/SYF2/INIP/PAXIP1/PPME1/TAF2/ABRAXAS1/TAOK1/CCND1/MASTL/F

RAP1/RBM3/TARS2/GCLM/ELP2/TRIM2/EARS2/DENR/EIF3G/PA2G4/AKT2/DUS3L/CAPRIN2/NSUN5/RPUSD3/RRBP1/MALSU1/W

XR/ACADM/ACSS1/GPD2/SDHB/AKR7A2/NDUFA1/AK4/AKT1/PDHA1/ENOX2/MTLN/ENPP1/NDUFB10/EIF6/DNAJC15/ATP5MF/C

/EAF1/INTS9/MED26/MED28/MED1/MED24/WDR82/MED17

31/KNL1/DTL/SMARCA4/TP53/TMEM14B/BCL7A/RINT1/CDC25B/SPDL1/PKD1/MAD2L1BP/ANAPC11/CDK1/RBBP8/ZFYVE19/PBF

PS2/BABAM2/SYF2/INIP/PPME1/TAF2/ABRAXAS1/TAOK1/CCND1/MASTL/FZR1/RRM2B/RAD17/TOPBP1/TICRR/CDK6/FOXO4

GID4/ADRM1/SKP2/KBTBD7/FAF1/PSMA5/IFI27/ZNF598/RNF187/ANAPC11/ANAPC13/PSMD7/SMARCC1/TRIM45/MAEA/PSMB5

/SMARCB1/MED29/KNTC1/NAP1L4/TRAPPC12/MED27/SSRP1/MED22/TAF1B/UBN1/RB1/MED14/CENPK/BRF2/TAF2/MED4/TAF

1B/RAD50/CDK4/NABP2/CENPE/DDRGK1/CCNE1/SETMAR/MIIP/CTDSPL/CCNG1/CDC5L/TFDP1/HUS1/KNL1/EME1/DTL/BRIP

E2/NDUFA8/NDUFA7/NDUFS2/NDUFS1/HSPA8/ATP5ME/NDUFA11/PFKM/NDUFA6/NDUFS8/NDUFB11/NDUFB2/PRXL2C/PGK

S3/ZWINT/CDC14B/RAD21/ORC1/BLM/DLG1/MRE11/LCMT1/MTBP/ABL1/MBTPS1/NME6/BRCA1/CTNNB1/CASP2/TPR/TRIAP1/

TL21A/SETD6/MDM2/NSMCE2/RASD2/KAT5/PIAS4

POLR1B

P1/XPOT/SNUPN/NUP155/NUP205/NUP188/IGF2BP2/NUP42/CHTOP/ALYREF/AGFG1/NUP107/KHDRBS1/UPF3B/BICD2/THOC1/

P1/XPOT/SNUPN/NUP155/NUP205/NUP188/IGF2BP2/NUP42/CHTOP/ALYREF/AGFG1/NUP107/KHDRBS1/UPF3B/BICD2/THOC1/

ERC2/DDRGK1/UBXN10/PCBP2/HECTD3/SPSB1/RMND5A/PEX10/UBXN2A/STUB1/CDC34/DTL/GID4/ADRM1/SKP2/ERLIN2/KBT

PF/DNM1L/NDUFC2/NME2/NDUFA8/NDUFA7/NDUFS2/NDUFS1/HSPA8/SMPDL3A/ATP5ME/NDUFA11/PFKM/NDUFA6/NDUF

F12/RBM42/PTBP1/SRSF9/GRSF1/KHDRBS1/SRSF10/UPF3B/SUPT20H/SRRM1/RBPMS/RBMX/U2AF2/PRPF19/POLR2A/TADA3

P1/XPOT/SNUPN/NUP155/NUP205/NUP188/IGF2BP2/NUP42/CHTOP/ALYREF/AGFG1/NUP107/KHDRBS1/UPF3B/BICD2/THOC1/

4/TTLL12/CDC16/BCL7C/MAD2L1/BIRC5/CKS1B/AKT1/PRKCA/CENPJ/HOXA13/USP22/PLK2/BUB1B/RCC2/GMNN/FOXC1/YWHA

H

F144B/ANAPC1/RNF126/IRF2BP1/UBE3C/RNF216/TRAF3IP2/HACE1/GABARAP/PPIL2/MAGEL2/TRIM27/RNF4/TRIM71/PELI3/RF

SG101/SNX30/CHMP2A/SH3GLB1/CHMP3/TBC1D5/NSFL1C/VPS37B/UBQLN1/CDK5/BNIP3L/CAPN1/GABARAP/ATP6V0A1/PHB2

NK1/MAP2/BOD1/CADPS2/MAD2L1/SEC16B/DOCK7/BIRC5/SKA3/LRPPRC/EIF6/MLPH/SYT11/PIK3CD/NLGN1/RCC2/RAB17/CH

FS2/NDUFS1/HSPA8/ATP5ME/NDUFA11/PFKM/NDUFA6/NDUFS8/NDUFB11/NDUFB2/PRXL2C/PGK1/NDUFV1/NDUFB3/FIGNL

D2/P3H4/CDC25B/ANAPC11/ANAPC13/RBBP8/NCAPH/SGO2/PRDM7/ZWINT/RAD51C/SGO1/ANAPC1/FIGNL1/RAD21/NCAPD2

N/MAP6D1/RAB6A/FUZ/KIF13A/LCA5L/RHOT1/WASL/HOOK2/BLOC1S6/BAG3/FBXW11/TTC21B/SSX2IP/NDE1/APP/DTNBP1/TAI

PFKM/NDUFA6/NDUFS8/NDUFB11/NDUFB2/PRXL2C/PGK1/NDUFV1/NDUFB3/FIGNL1/PRKAG1/STOML2/NDUFS4/FOXK2/UQ

O1/FIGNL1/RAD21/NCAPD2/MSH6/SIRT2/SPIRE1/FANCA/MRE11/SLX4/GOLGA2/FBXO43/MND1/ORC4/PTTG1/TTK/CCNE2/INC

50/NLE1/TNKS/NABP2/SETMAR/MIIP/MAPK12/CTDSPL/CCNG1/CDC5L/HUS1/KNL1/EME1/DTL/BRIP1/TP53/MSH2/FANCD2/RIN

ON/RBM23/NCBP2/RBM39/DYRK1A/CPEB1/SRSF2/ZC3H14/RBM15/NCL/SNW1/SF1/CWC22/CDC73/HNRNPL/FMR1/RBMXL1/S

2/SAE1/YWHAG/HSPA8/AP3D1/UBE2L3/GDI1/GDAP1/CSNK2A2/HSPA4/ZFYVE16/CDC37/MICALL1/ZDHHC20/TIMM17B/MTCH2

EIF2S1

FA7/NDUFS2/NDUFS1/HSPA8/ATP5ME/NDUFA11/PFKM/NDUFA6/NDUFS8/NDUFB11/NDUFB2/PRXL2C/PGK1/NDUFV1/NDUF

PP/DTNBP1/AP3B1/TRIM58/TRAK1/KIFAP3/TRAF3IP1/BICD2/MAPK8IP3/PCM1/BLOC1S3/ARL8B/UBB/KIF1C/PEX14/RABGEF1/CL

D24/TMEM33/USP22/POLH/DNMT3A/NCK2/FEM1B/SMYD2/DDRGK1/SETMAR/CBX8/TRIM28/RTN4RL1/TAF12/UBXN2A/STUB

AR/CTNNA2/PIF1/GNL3L/EHMT2/KNL1/ARHGEF7/TMOD2/TP53/MUL1/SPDL1/HDAC6/MYADM/MAD2L1BP/EML2/SEC22B/NOL3/

WIZ/CCT2/MUL1/HSPA8/HDAC6/USP19/CDC37/PFDN2/P3H1/DDI2/MAPK1/MYLIP/MCM8/TCP1/SH3GLB1/URI1/PPIB/DPM2/GO

3/COPS5/CHAC1/ERMP1/CREBZF/YOD1/PARP6/HERPUD2/COMP/DNAJB1/DNAJA1/PTPN1/CCND1

M7/POLDIP2/RAD51C/DYNC1H1/CDC14B/RAD21/CHMP3/NSFL1C/PRKCE/NCAPD2/SIRT2/RRM1/SASS6/PHB2/EGF/GIPC1/MT

RBBP5/MED27/MED22/TAF1B/MED14/BRF2/MED12/PAXIP1/BRD7/TAF2/USP21/CAVIN1/E2F2/MED4/MEN1/TAF8/GTF2E2/CDK7/

DUFC2/NME2/PPAT/NDUFA8/NDUFA7/NDUFS2/DLAT/TK1/ACLY/ACOT7/NDUFS1/SLC52A3/AK6/NADK2/ATP5ME/NDUFA11/N

WDPCP/IFT88/BRCA1/RAB1B/TPR/G3BP2/ODF2L/WRAP73/PPP1R35/RDX/CCSAP/MAP4/NUPR1/ATG2A/FUZ/ENTR1/STAM/PA

DUFC2/NME2/PPAT/NDUFA8/NDUFA7/NDUFS2/DLAT/ACLY/ACOT7/NDUFS1/SLC52A3/AK6/NADK2/ATP5ME/NDUFA11/NDUF

SLX4/GOLGA2/FBXO43/MND1/ORC4/PTTG1/TTK/CCNE2/INCENP/SMC2/BAG6/FMN2/ZSCAN21/C14orf39/MSX1/FOXJ3/PSMD13

RP6/ECT2/CDC25B/PTK6/MAD2L1BP/MEIS2/ANAPC11/CDK1/ZPR1/NCAPG2/USP19/NCAPH/PRDM7/POLDIP2/TTL/PTPN11/RAD

C14B/NDFIP2/CRY1/PRKCE/UBQLN1/MINAR1/CDK5/HMG20A/GABARAP/TOLLIP/ARNT/BMI1/MTBP/ABL1/CTNNB1/PRMT3/FBX

MND1/ZNF598/ELP5/HABP4/GSPT1/DHX29/PUM3/EIF2B4/RIDA/EIF2AK4/DHFR/RCC1L/MSI2/FASTKD2/GUF1/EIF5B/PDIK1L/ILF

3/CHMP4B/EEA1/RAB11A/PDCD6IP/VPS33B/RNF26/HOOK1/ATP6V1A/SQSTM1/SNX10

2/ATM/SESN1/NLK/BMT2/CTNS

9B/UPF3A/POM121/CETN3/SRSF3/RBM8A/SLC35C2/DDX19A/TPR/THOC5/TOMM20L/G3BP2/NUP133/RBM15B/NUP35/SLC29A4/

5/BCL2L11/DNAJC4/ERLEC1/EDEM3/UBE2W/COPS5/CHAC1/CAV1/ERMP1/CREBZF/YOD1/PARP6/HERPUD2/COMP/DNAJB1/D

DLAT/ACLY/ACOT7/NDUFS1/NADK2/ATP5ME/NDUFA11/NDUFA6/NDUFS8/PANK2/NDUFB11/NDUFB2/PANK4/NDUFV1/NDUFB

ARL6/KIF13A/STAM/WASL/BLOC1S6/STARD3NL/FBXW11/NAGLU/NDE1/DTNBP1/TRAPPC6A/EXOC8/DCTN1/TRAPPC1/TANC2/

N1/SPTBN5/CARMIL1/CAMSAP2/EEF2K/SH3GL3/HMGA1/CLASP2/SMARCD2/MICAL1/SUPT16H/LIX1L/KATNB1/CLASP1/SNAPIN

RKCE/UBQLN1/MINAR1/CDK5/HMG20A/GABARAP/TOLLIP/ARNT/BMI1/MTBP/ABL1/CTNNB1/PRMT3/FBXO2/PDCD6/RWDD3/A

RBM10/DUS3L/CAPRIN2/ELOC/NSUN5/RPUSD3/EXOSC8/MALSU1/C8orf88/NT5C3B/C1QBP/TP53/EIF5A2/OGFOD1/UQCC2/EIF4

B/SNAPIN/NUPR1/LAMP2/TP53INP2/ATG2A/HOOK2/BAG3/VPS18/SCARB2/NAGLU/PIP4K2A/TMEM199/VMP1/RB1CC1/ATP6V0C

I/LAMC1/MAP1LC3B/HAX1/MELTF/SLC25A5/CAPG/ATP5IF1/MAP4K4/NAPB/ATG4B/DDRGK1/DNAJC6/DENR/DNM1L/ATG9A/MN

A1/TPR/CAMSAP1/MECP2/KATNB1/CLASP1/BICD1/PPP1R35/GBA2/HAUS1/CCDC40/MAP6D1/CCSAP/KLHL42/NAT10/STMN1/M

AD/GIGYF2/EIF3A/EIF3L

39A/TOM1/ATG4C/ARFIP2/RAB7A/PACS2

DUFS2/DLAT/ACLY/ACOT7/NDUFS1/NADK2/ATP5ME/NDUFA11/NDUFA6/NDUFS8/PANK2/NDUFB11/NDUFB2/PANK4/NDUFV

USP36/TP53/VPS36/TNPO2/CCT2/SAE1/ECT2/HSPA8/AP3D1/PTPN23/UBE2L3/GDAP1/ANGPT1/CDK1/ZPR1/RANGRF/LRWD1/

/GTF2E2/CDK7/SRF/MED21/CAND1/MED10/MED19/MED9/ERCC3/TAF7/PHF2/SMARCD1/MED16/PCID2/GLYR1/MED26/N6AM

CSS1/LDHC/SDHB/NMNAT1/NDUFA1/AK4/PDHA1/ENPP1/NDUFB10/EIF6/ATIC/ATP5MF/GNAI3/HACD1/ATP6V1B2/SDHA/FOX

2/TP53BP2/CHEK1/MDM2/ERCC2/KAT5/RRP8/CDIP1/SPRED2

/RHOT1/CABYR/BLOC1S6/BAG3/FBXW11/TTC21B/SSX2IP/NDE1/APP/DTNBP1/AP3B1/TRIM58/TRAK1/KIFAP3/TRAF3IP1/BICD

DB/DYNC2H1/PTPN23/POC1A/HDAC6/ARMC9/CEP89/ATMIN/CEP131/VDAC3/WDR35/MKKS/CPLANE2/ARHGAP35/BBS9/IFT14

A1/AK4/PDHA1/ENPP1/NDUFB10/EIF6/ATIC/ATP5MF/GNAI3/HACD1/ATP6V1B2/SDHA/FOXK1/ATP5IF1/ITPA/NDUFB7/SUCLG1/

CSS1/LDHC/SDHB/NMNAT1/NDUFA1/AK4/PDHA1/ENPP1/NDUFB10/EIF6/ATIC/ATP5MF/GNAI3/HACD1/ATP6V1B2/SDHA/FOX

AB23/ATG4D/RALB/TMEM39A/TOM1/ATG4C/ARFIP2/RAB7A/PACS2

CB1/DDB1/KDM1A/CHEK1/DOT1L/RRP8/MTA1/RBBP5/RNF8/RB1/CGGBP1/EZH2/HDAC1/BAHD1/DNMT1/MORC2/BRD7/DYR

/HDAC6/ARMC9/CEP89/ATMIN/CEP131/VDAC3/WDR35/MKKS/CPLANE2/ARHGAP35/BBS9/IFT140/TTLL1/CDC14B/RAB34/AHI

1/BAG4/CHEK2/NAPG/HSP90AA1/LAMP2/HIP1/COG3/BAG6/TELO2/PER3/BAG3

S37B/ITGB1BP1/HRAS/MTX1/RAB34/CDK5/EMC2/SPTBN1/EMC7/CCDC47/ATP6AP1/SEC61B/CLSTN1/SSR3/CHM/VPS37A/AR

1A/TMOD2/MOSPD2/CENPX/BSCL2/OGFOD1/CNTROB/GTF2B/SBDS/TRAF7/DHX29/SH3PXD2B/RANGRF/CETN2/FASTKD2/

A3/GLI3

/EP400/MCM3/MCM8/NSMCE1/FIGNL1/RAD21/MEAF6/MRGBP/SPIDR/MSH6/RTEL1/RAD52/UCHL5/ZSWIM7/RPA3/XRCC2/BL

NK1/MPC2/ACOT11/TECR/BPNT1/DLST

NK1/MPC2/ACOT11/TECR/BPNT1/DLST

NK1/MPC2/ACOT11/TECR/BPNT1/DLST

3B/KIF1C/AP1AR/SNX5

8/INTS7/RAD1/BABAM2/RAD51AP1/INIP/PAXIP1/ABRAXAS1/TIGAR/EYA3/MEN1/XRCC4/MAPK14/RFWD3

4B/DLAT/ST3GAL4/CSAD/ACLY/ACOT7/B3GALT6/ANGPT1/GLO1/G6PD/NDST1/MRI1/MMACHC/ELOVL1/HAGH/GGT1/XYLT1/B

NT11/HOXB3/TDRD7/ANKS6/CC2D2A/SETDB2/NCLN/HAND2/DVL2/ALG5/TSHZ1/RPGRIP1L/CLUAP1/TCTN1/NKX2-2/FOXC1/N

SC2/CAPRIN1/PABPC4/UPF3A/PATL1/TRIM71/DDX17/PUF60/CNOT11/SRSF3/RBM8A/ELAVL1/SAFB/BAG4/YTHDF3/PARN/RB

3/ACSS1/GPD2/LDHC/SDHB/NMNAT1/NDUFA1/AK4/PDHA1/ENPP1/NDUFB10/EIF6/ATIC/ATP5MF/GNAI3/DERA/HACD1/NAXD

/TRAPPC5/ARL6/KIF13A/STAM/WASL/BLOC1S6/STARD3NL/FBXW11/NAGLU/NDE1/DTNBP1/TRAPPC6A/EXOC8/DCTN1/TRAP

N1/HSP90AA1/TGFBR1/PLAA/DNAJB2/TCF25/SPRTN/HSPBP1/FBXO4/HDAC3/GPS2/KDM1A

:5/REV3L/PCLAF/HSP90AA1/NAT10/CCNA2/LIG1/TELO2/TNKS1BP1/SPRTN/FBXO4/USP1/NVL/CHEK1/POLI/RFC5/MAP2K7
/EIF5A2/OGFOD1/UQCC2/RTN2/EIF4G3/DHX9/MTIF2/RMND1/ZNF598/ELP5/HABP4/GSPT1/DHX29/PUM3/EIF2B4/RIDA/EIF2AK4
2B/PLOD3/RNF207/SPRY2/TWIST1/BCL2/MTOR/GZF1/CC2D2A/NPHP1/SETDB2/HAND2/DEAF1/DVL2/HOXA13/RPGRIP1L/PIK3C
5/HOXA3/MMP14/GLI3/RYK/PPARGC1B/DLG1/BMI1/WDR19/SKI/NKX3-2/CTNNB1/HOXA2/GLG1/HOXC11

3/MTHFD1L
TRIM28/CCT4/TFDP1/TAF12/GNL3L/EHMT2/SENP3/TAF4/MCM4/RFC3/CYREN/SMARCA4/TP53/MSH2/WIZ/CCT2/BCL7A/UBE2\
IDAC6/CEP131/VDAC3/WDR35/MKKS/TMEM108/IFT140/DYNC1H1/IGBP1/TTLL1/NDEL1/SPA17/HSPB1/KIF22/KIF15/DYNLRB1/D
AND2/DVL2/ALG5/TSHZ1/RPGRIP1L/CLUAP1/TCTN1/NKX2-2/FOXC1/NLE1/DPCD/HOXB2/IFT172/MIB1/RARG/CELSR2/DYNC2L
NA/GLI3/CSE1L/TRAF3IP2/CDK5/NUP58/INTS13/PHB2/PIN1/POM121/ATP13A2/KPNA6/IPO9/UFM1/SMO/ELAVL1/TPR/RPF2/NUP

CHMP3/VPS51/TBC1D5/VPS37B/MAP2K2/LEPROT/PXK/MAGEL2/PHETA1/ANKRD27/TRIM27/ATP6AP1/SNF8/EHD4/DENND10/U
CLCN3/TMED9/SEC22B/DNAJC13/ZFYVE16/CPLANE2/GOSR1/ZDHHC20/RAB13/CHMP1B/STX1A/TSG101/CHMP2A/PEF1/CHM

P2/CTTN/WDR47/AFG3L2/PTPRS/DOCK7/ARHGEF25/CDK5R1/LAMB2

N2/ARHGEF7/FKBP1B/FN1/ROR1/PTK6/HDAC6/ADNP/GDI1/AKIRIN1/ATMIN/ITGA6/PLEK2/ARHGAP35/P3H1/NDEL1/STK24/RAL
LE1/TNKS/CBY1/SALL1/CCNE1/ZBED3/RAC1/RARG/CDH2/CELSR2/MAGI2
LE1/TNKS/CBY1/SALL1/CCNE1/ZBED3/RAC1/RARG/CDH2/CELSR2/MAGI2
:HG4B/NTNG1/C12orf57/NIBAN2/CERS2/TSPO/MAP2/CTTN/WDR47/AFG3L2/PTPRS/DOCK7/ARHGEF25/CDK5R1/TBC1D24/LAM
/PCSK5/AICDA/MX1/NR5A2/ITGA5/IRF7/CIITA/ICAM1/CD80/RSAD2/CD28/ITGB7/IFIH1/ISG20/CXCR4/CCR5/LTF/IFIT1/TNF/LAMI
HBA/NCKAP1L/TNF/SFRP4/PACSIN1/LRRTM1/ANK3/CORO1A/MMP9/HAMP/ANO9/LIF/ADIPOQ
CNB4/SLC43A2/NMUR2/NR0B2/PPARG/SLC6A18/UGT1A3/FABP9/ACSL5/FABP12/ABCB11/TNFRSF11A/FABP6/SLC25A2/OC90/S
CNB4/SLC43A2/NMUR2/NR0B2/PPARG/SLC6A18/UGT1A3/FABP9/ACSL5/FABP12/ABCB11/TNFRSF11A/FABP6/SLC25A2/OC90/S
GFC/PLG/S100A2/EFNB2/TGFB1/PTK2B/WNT5A/HDAC9/GATA2/FOXC2/INSL3/DLL4/PTPRR/CD40/KLF4/WNT7A/PDLIM1/CLEC1
/SNL1/IL1B/CD38/HCLS1/TNF/DOC2B/RBP4/TMEM132A/TLR4/BLK
B/AGAP2/WNT5A/CD300A/PDGFC/PKIA/JAK2/CD40/THY1/PIK3R6/HHEX/LYN/LILRB4/EPHA1/IL34/SLC8A1/BMP2/CCL19/LRRK2/I
I/NPTX1/RYR3/CYP1A1/TXNIP/SYT8/AQP3/SOD2/MT1M/HMOX1/TF/VCAM1/CPNE4/CEBPA/FABP4/FGA/THBS1/ADCY7/JUNB/F
D2/SCN4B/SLC23A1/SLC6A13/SLC9A9/ATP1A4/PKP2/SLC9C1/SPTBN4/PKD2L1/SLC4A8/KLHL3/SLC38A11/SCN1B/ATP4A/FGF11/S

NR1H3/AREG/UCP2/JAK2/TGFB3/PTGES3/CYBB/PADI2/TFPI/TACR1/SLC12A3/PAPPA/A2M/PTGDS/CD38/IL1RN/INHBA/TNF/PT
/KLF4/LYN/GZMA/LILRB4/EPHA1/SLC8A1/BMP2/CSTA/LRRK2/SPINK5/IL1B/LAX1/SERPINB9/PTPN22/LTF/IFIT1/SPOCK2/NCKAP
:/TXNIP/AQP3/KRT14/MYCL/KRT72/TP63/GDF3/DLL1/CDH23/ZBED2/WNT5A/ACER1/KLF4/ST14/CSTA/SPINK5/KRT2/LIPK/FGF2

GN4X/CACNG2/NOG/DNM1/CYP46A1/NLGN3/INS/ADRA2A/ADORA3/CA7/GUCY1B1/EGR2/CHRNA3/TNR/DLGAP3/BEGAIN/LILR
GN4X/CACNG2/NOG/DNM1/CYP46A1/NLGN3/INS/ADRA2A/ADORA3/CA7/GUCY1B1/EGR2/CHRNA3/TNR/DLGAP3/BEGAIN/LILR
NK3/RASGRP2/ALOX15/MMP9/HVCN1

B1/NCEH1/CYP2E1/SLCO2B1/ABCC3/CYP2S1/CYP2C9/GSTO2/AOX1/IL1B/CXCR4/VAV1/CD69/NOS2/AIM2/ADIPOQ
GB7/ITGB8/TNXB/UNC13D/PARVG/MYO1G/SPOCK2/CD3E/CD96/RAC2/CORO1A/ITGAL/CCR7/ALOX15/MYOC
RA1/GLRB/KCNK17/KCNQ1/KCNH5/FGF12/CHRNA3/GLRA1/KCNC1/NTSR1/KCNH6/HTR3C/NTRK2/ARRB2/GRIK4/KCNK6/CA
ADRA1A/RAMP3/DES/SCN4B/THRB/FOXN4/MYL2/KCNJ5/TNNI1/PKP2/ADM2/GCH1/SPTBN4/KCNA5/ATP2B3/CACNA1C/HOPX/

I2B/CD74/PDGFRA/TREM2/FGFR2/CD4/FLT3/MERTK/CD86/PTK2B/TIE1/PTPRO/PKIB/AGAP2/WNT5A/NLRC5/CD300A/PDGFC/
2-5/CSRP3/NEUROG1/MYH11/ADRA2A/KCNQ1/MYBPC3/MYL9/FGF12/CHRNA3/SSPN/GLRA1/TNNT3/MYL4/SULF1/NMUR2/GU

SS3/ADCY8/LIME1/TRDN/CCL3/VDR/STEAP4/ATP6V1B1/APLNR/CSRP3/ATP13A4/CP/CCR1/CA7/PTH1R/RAB39A/P2RY6/NTSR

G1/GMIP/PDPN/ARHGAP22/TGM2/DOCK3/AIF1/IQSEC3/CYTH4/GPR65/DOCK8/RGL4/CGNL1/ARHGAP27/ARHGAP45/ARHGAF

/CDH17/SFRP1/ALOX5/FGB/BMP8A/ILDR1/PCLO/UCP2/FFAR1/JAK2/SMPD3/SLC15A2/VSNL1/IL1B/SLC38A5/CD38/IL1RN/TNF/D

3/CACNA1C/HOPX/SCN1B/FXYD1/SLC8A1/TNNT2/TNF/PIK3CG/ATP2A3

2/TGFA/JCAD/PPP1R16B/FGF2/TNF/BTK/FLT4/CCL5/FGF1/TBX1

IA15/PTH2R/GALR2/UCN3/GALR1/ADRB2/GLP1R/GPER1/SSTR2/GABBR2/ADRA1B/CASR/CHRM2/ADCY8/ADGRF3/ADRA2A/AI

P2C8/BMP5/RDH5/RDH12/RDH8/DKK3/CYP1B1/ALDH1A2/CYP2E1/AKR1B15/ALOX5/PTGES3/CYP2S1/CYP2C9/FAAH2/BMP2/PL/

L3/APOL4/GHRL/CIDEC/TREM2/PLA2G5/APOL1/APOBR/NFKBIA/IL6/ATP8A1/ANO3/SLCO2A1/SLCO2B1/NR1H3/ABCC3/FASLG/S

R/RAC2/LAT2/CORO1A/EXOC3L4/SYTL1/CCL5/NKG7

10/MMRN1/PTPN6/IL33

N1

C/CCL20/CXCL13

2/RASGEF1A/FGD5/SEMA4D/KLRK1/ARHGAP9/TBC1D1/VAV1/RASGRP2/ARHGAP25/CCL5/RASGRF1/TBC1D10C/ARHGAP15/

ASH1/PTN/NKX3-1/HNF1B/CCL2/TACSTD2/SULF1/WNT2/MMRN2/IGFBP4/TEK/TGM1/PPARG/AGTR1/EPPK1/WNT10B/KDR/RE

DRA1A/DES/SCN4B/THRB/FOXN4/MYL2/KCNJ5/TNNI1/PKP2/ADM2/GCH1/SPTBN4/KCNA5/ATP2B3/CACNA1C/HOPX/SCN1B/

CY5/GHRL/TUNAR/TREM2/KCNA5/IL6/TGFB1/ABAT/SLC4A8/SFRP1/PCSK5/ALOX5/ADAM8/FGB/BMP8A/NR1H3/PCLO/TVP23A/

A2/TGFB1/SULF2/TIE1/SFRP1/ALOX5/WNT5A/AREG/GATA2/DLL4/WNT7A/TACR1/JAML/MMP12/SGPP2/TGFA/EAF2/IL12A/JCAI

CNJ5/TNNI1/PKP2/KCNA5/CACNA1C/SCN1B/PGAM2/TNNI2/SLC8A1/TNNT2/CXCR4/TNF/PIK3CG/RCSD1

B/ADCY5/GHRL/TUNAR/TREM2/KCNA5/IL6/TGFB1/ABAT/SLC4A8/SFRP1/PCSK5/ALOX5/ADAM8/FGB/BMP8A/NR1H3/PCLO/TV

2/TOX/SPN/GATA3/LY9/BCL11B/IL23A/SLAMF6/TBX1

A/PPY/VEGFC/IL12B/ADCY5/GHRL/TUNAR/TREM2/KCNA5/IL6/TGFB1/ABAT/SLC4A8/SFRP1/PCSK5/ALOX5/ADAM8/FGB/BMP

8L3/APOL4/GHRL/CIDEC/TREM2/PLA2G5/APOL1/APOBR/NFKBIA/ATP8A1/ANO3/SLCO2A1/SLCO2B1/NR1H3/ABCC3/FASLG/STA

CR5/LTF/IFIT1/TNF/LAMP3/APOBEC3A/HAVCR1/CCL5/OASL/CR1/CR2

C2/SLC4A8/PTK2B/SYT1/KMO/TACR1/CALB2/TNF/LRRTM1/RELN/CLSTN3

GDR/CYBB/CCL21/KLF4/TACR1/SLC12A3/CES1/A2M/CCL19/FOXP3/CCR5/INHBA/TNF/MAP4K1/CD27/PTPRC/CCR7/CLDN5/AD

/TFAP2B/SPP1/TNFRSF11A/BAIAP3/RPH3AL/MTNR1B/P2RX1/PTGS2/ITPR1/SNCG/IFNG/GIPR/KCNA2/FAM3B/STX11/ADRA2B/E

N/CRTAM/CCL19/PRF1/CXCR4/DOCK2/CD5/EMB/CD6/CCR7

NN3/NALCN/RGS7/SLC1A3/KCNN1/KCNA2/KEL/KCNK13/LRRC26/FXYD2/KCNJ5/SLC9A9/ATP1A4/TREM2/SLC9C1/PKD2L1/KCN

1/FUCA1/SCT/CIDEC/PLA2G5/PLBD1/DPEP2/SMPD3/ACER1/AKR1D1/LIPG/FAAH2/CES1/LIPH/PLA2G4C/IL1B/LIPK/PLA2G7/AOA

1/IL10/CYP1B1/IL12B/GADD45B/TGM2/GCH1/IL6/ALDH1A2/TGFB1/ZBTB16/TNFRSF1B/LGALS9/DAPK2/HTRA4/SFRP1/MNDA/AD

RASGEF1A/FGD5/SEMA4D/P2RY8/GPR18/TAGAP/PIK3CG/PSD4/ARHGAP9/VAV1/RHOF/RELN/RASAL3/ARHGAP25/GPR174/RAS

ORO1A/PTPN6/C3/PTPRC/CXCL9/CCR7/XCR1/CXCL10/CEMIP/CXCL11/CD19

1/GNA14/TNF/ANK3/CNR2

LO/UCP2/FFAR1/JAK2/SMPD3/SLC15A2/VSNL1/IL1B/CD38/IL1RN/TNF/DOC2B/TNFSF11/RBP4/CCL5/NOS2/BLK

LC43A2/KCNC2/CACNG3/ACSL5/KCNMB1/LACRT/SCN3B/RGS7/P2RX1/IFNG/EDN1/GRIN2A/PIM1/EPHB2/TMSB4X/STAC3/JPH3

1/SEMA7A/TBX21/LTF/JAK3/SEMA4D/INHBA/TNF/TLR4/INPP5D/CR1/SEMA3A/BCL11A/ADIPOQ

NC2/KCNMB1/KCNQ3/KCNN3/NALCN/RGS7/SLC1A3/KCNN1/KCNA2/KEL/KCNK13/LRRC26/FXYD2/KCNJ5/SLC9A9/ATP1A4/TRE
1B/CD38/IL1RN/TNF/DOC2B/TNFSF11/RBP4/CCL5/NOS2/BLK

NL1/IL1B/CD38/IL1RN/TNF/DOC2B/TNFSF11/RBP4/CCL5/NOS2/BLK

T/KCNA5/IL6/ABAT/SFRP1/ALOX5/FGB/BMP8A/ILDR1/PCLO/UCP2/FFAR1/JAK2/SMPD3/SELENOM/LYN/TACR1/VSNL1/IL1B/CD3

C2/INSL3/DLL4/PTPRR/CD40/KLF4/WNT7A/CLEC14A/TACR1/KRT2/JCAD/FGF2/PLCG2/TNF/GATA3/PIK3CG/CORO1A/FLT4/MMI

1/JAK2/TGFB3/SMPD3/ACHE/SYT1/KMO/ABCG1/TACR1/VSNL1/BMP2/ITGB2/IL1B/UNC13D/CD38/INHBA/DOC2B/TNFSF11/FGF

SLC4A8/LRP2/ATP8A1/CYSLTR1/C2CD4B/SLCO2B1/FGB/ABCC3/FOXC2/TJP3/TACR1/SLC15A2/SLC8A1/PDE2A/SLC38A5/CD38/S

C2/INSL3/DLL4/PTPRR/CD40/KLF4/WNT7A/CLEC14A/TACR1/KRT2/JCAD/FGF2/PLCG2/TNF/GATA3/PIK3CG/CORO1A/FLT4/MMI

FGA/SYK/VEGFC/PTAFR/GHRL/STX1B/TUNAR/SCT/TREM2/TGFB1/IL4R/ABAT/SLC4A8/ADAM8/FGB/ILDR1/GATA2/FFAR1/JAK
R6/PTGES3/KLF4/PARM1/EPHA1/IL34/PRKCQ/CCL19/LRRK2/IL1B/CCND2/RASGRP1/LTF/FGF2/TNF/TNFSF11/PIK3CG/TLR6/PL
GDF2/CDH11/EVL/CYP1B1/CD200R1/IL27RA/MAPK15/CD74/PTGER4/IFITM1/CHRD/TGFB1/AIF1/TIE1/SFRP1/GCSAM/CD300A/D

RE1/ADCY4/ADGRG4/GPR65/ADGRG2/ADGRL3/S1PR4/S1PR1/CXCL9/CXCL10/ADGRG5/ADGRF5/CXCL11/CNR2
T1/NDRG4/CCL21/WNT7A/LYN/SIRPG/SPI1/LILRB1/VSNL1/CAMK1D/GSG1L/ITGB2/CCL19/LRRK2/SIRPB1/CD300LF/UNC13D/CI
C9/GATA2/FOXC2/INSL3/DLL4/PTPRR/CD40/KLF4/WNT7A/CLEC14A/TACR1/KRT2/JCAD/FGF2/PLCG2/TNF/GATA3/PIK3CG/COF

CN4B/CASQ1/AKAP5/TREM2/PTK2B/CACNA1C/SCN1B/CTSS/THY1/ATP4A/LYN/FGF11/TLR9/FXYD1/SLC8A1/TSPAN13/NETO1/I

AR1/JAK2/TGFB3/ACHE/ABCG1/RSAD2/VSNL1/IL1B/IL12A/CD38/TNF/DOC2B/RBP4/TMEM132A/TLR4/CCL5/NOS2/BLK

/NR1H3/NR5A2/JAK2/PTGES3/IRF7/HAVCR2/PADI2/RSAD2/SLC15A2/TLR9/BMP2/IFIH1/NLRP1/PIK3AP1/PTPN22/PLCG2/TNF/TF

HBS1/GDF2/CDH11/EVL/CYP1B1/CD200R1/IL27RA/MAPK15/CD74/PTGER4/IFITM1/CHRD/TGFB1/AIF1/TIE1/SFRP1/GCSAM/CD

2/POPDC2/ADRA2B/ADM/EDN1/ITGA4/ADRA1A/DES/SCN4B/FGA/THRB/FOXN4/SVEP1/HRH1/MYL2/KCNJ5/PKP2/ADM2/GHRL

/BMP5/RDH5/RDH12/RDH8/MEP1A/DKK3/ECE2/CYP1B1/PDGFRA/ALDH1A2/STC2/AKR1B15/PCSK5/GCNT4/AKR1D1/CYP2S1/S

MP/TMEM119/CLDN5/DCSTAMP

MP/TMEM119/CLDN5/DCSTAMP

KCNK6/ACSM3/PPARG/AGTR1/GUCY1A1/CYP4F12/SERPINF2/CPA3/P2RX1/PTGS2/ADRA2B/ADM/EDN1/ADRA1A/SOD2/VEGF

/KCNJ5/SLC9A9/ATP1A4/AKAP5/SLC9C1/SLC15A3/LRP2/CACNA1C/CACNA1I/ATP4A/KCNJ10/SLC12A3/SLC15A2/SLC9A7/SLC8A

1/IL1B/CD38/INHBA/TNF/DOC2B/TNFSF11/RBP4/CCL5/NOS2/LIF/BLK/ADIPOQ

TAC3/ADRA1A/JPH3/CACNG8/F2/PKD1L3/CASQ1/AKAP5/PKD2L1/ATP2B3/CACNA2D3/PTK2B/CACNA1C/FASLG/THY1/CACNA

PAN32/FERMT3/PPARG/SERPINE2/HRG/EPPK1/KDR/SERPINA1/SERPIND1/APOH/AXL/LACRT/SERPINF2/DGKE/P2RX1/REG3

C3/CES1/ABCC11/IL1B/EMB/SLC22A3/TNF/RBP5/TNFSF11/ABCD2/RBP1/PLA2G2D/PLA2G2A/NOS2

1/AIF1/TIE1/PTPRO/SFRP1/GCSAM/WNT5A/CD300A/DLL4/PTPRR/THY1/SPINT2/NDRG4/CCL21/KLF4/PADI2/ARHGAP4/EPHA1/

IPOQ

T/KCNA5/IL6/ABAT/SFRP1/ALOX5/FGB/BMP8A/ILDR1/PCLO/UCP2/FFAR1/JAK2/SMPD3/SELENOM/LYN/TACR1/VSNL1/IL1B/CD

A5/STAC3/JPH3/KEL/SLC31A2/CCL4/TF/LRRC26/FXYD2/F2/SCN4B/CASQ1/MRLN/PKP2/AKAP5/TREM2/SPTBN4/KCNA5/CD4/S

S1PR1

3/PIK3CG/FLT4/MMP9/FGF1/CXCL13

PCSK1N/NTS

CARD9/ADRA1A/NLRC4/CCL14/CCL4/CASP10/TBC1D9/SGSM1/SYK/ODAM/CCL22/CCL24/GMIP/PDGFRA/ARHGAP22/TGM2/PL

CR2/KLF4/SPI1/IRAK2/CAMK1D/TLR9/TRAF1/PRKCQ/BMP2/FOXP3/IL1B/CD40LG/LTF/PLCG2/TNF/SFRP4/TNFSF11/BTK/TLR6/R

3CG/FLT4/FGF1/CXCL13

/LIF/IDO1

2/CASQ1/MRLN/AKAP5/CD4/STC2/PTK2B/CACNA1C/FFAR1/THY1/LYN/LILRB1/TLR9/SLC8A1/TSPAN13/CD84/CXCR4/PIK3CG/[illegible]

CDHB13/PCDHAC1/CDH18/PCDHA12/PCDHB3/CDH5/PCDHB10/CDH11/CDH23/PECAM1/PCDHB5/CDH17/PCDHB7/TENM3/P[illegible]

1/IFNG/EDN1/GRIN2A/EPHB2/TMSB4X/STAC3/JPH3/KEL/SLC31A2/LRRC26/FXYD2/F2/SCN4B/CCR2/CASQ1/KCNJ5/AKAP5/TRE[illegible]

R2/CASQ1/TGM2/TUNAR/CDH23/ATP2B3/STC2/PTK2B/CACNA1C/CCL8/WNT5A/SV2B/FASLG/CD40/THY1/CCL21/SV2A/LYN/C[illegible]

:RPINE2/HRG/HYAL2/CDO1/CYP4F12/SERPINA1/SERPIND1/APOH/AXL/SLC4A9/LACRT/SERPINF2/PRICKLE1/DGKE/P2RX1/SE

/PLCZ1/EDN1/ADORA2A/LILRA5/TMEM37/GRIN2A/RYR3/STAC3/CALCRL/ADRA1A/JPH3/RAMP3/CACNG8/CCL4/F2/PKD1L3/CA

P2RY6/NTSR1/KCNJ13/CACNB4/SERPINE2/GRAMD2A/KCNC2/KCNMB1/STC1/LACRT/SCN3B/RGS7/P2RX1/PTGS2/ATP8B1/G/

:RTK/LGALS9/GPR183/TNFRSF17/GATA2/GCNT4/JAK2/TNFSF14/SPNS2/LYN/SPI1/CCR4/GAPT/LGALS2/IL2RA/FOXP3/JAK3/HC

SF11/PIK3CG/TLR6/RELN/PTPRC/TLR4/CCR7/CCL5/IL23A/CD24/FGF1/CEMIP/ADIPOQ

IAMP

6/TGFB1/KLF2/RLN2/PTK2B/ITGAX/TIE1/SFRP1/CYSLTR1/ALOX5/WNT5A/FASLG/GATA2/FOXC2/ITGA5/CD40/ECSCR/PIK3R6/C

F2/TNF/FLT4/COL4A3

2/P2RX1/ITPR1/EDN1/ERC2/RYR3/JPH3/CCL14/KEL/CDH5/F2/KCTD17/CCR2/FAM20A/CASQ1/TGM2/TUNAR/CDH23/ATP2B3/S

/CD40/PIK3R6/EPHA1/IL34/CCL19/LRRK2/IL1B/CCND2/RASGRP1/LTF/FGF2/TNF/TNFSF11/PIK3CG/TLR6/RELN/FLT4/PTPRC/TL

6/KLF2/RLN2/PTK2B/ITGAX/TIE1/SFRP1/CYSLTR1/ALOX5/WNT5A/FASLG/GATA2/FOXC2/ITGA5/CD40/ECSCR/PIK3R6/CYBB/HH

YFIP2/SPINK5/SERPINB9/NLRP1/LTF/LCK/SPOCK2/TNF/LAPTM5/LAMP3/TNFSF10/BIRC3/SERPINA3/ATP2A3/CR1/COL4A3/MM

/FLI1/NFKBIA/BATF3/CD4/TGFB1/KLF2/ZBTB16/CAMK4/PTK2B/CLDN18/SFRP1/IL31RA/UCP2/GPR183/GATA2/JAK2/IL15/GAB3/IF

2/SCN5A/PPP1R13L/F2RL1/NOS1AP/KCNJ3/KCND3/SCN2B/TRDN/NOS3/ZC3H12A/KCNJ8/SCPEP1/NKX2-5/SLC2A5/CSRP3/INS

ITS

DHB1/TENM4/CD209/PCDH11Y/PCDHA4/CADM2/CELSR3/CLDN16/DSG2/TENM1/PCDHA10/DSC2/APOA1/PCDHA11/PCDHB1

/CD4/TGFB1/XCL2/PTK2B/LGALS9/CCL8/FGB/PDGFC/GPR183/PTPRR/ICAM1/NDRG4/HAVCR2/CCL21/SPRED3/LYN/TLR9/BMI

ITS

ITS

IPOQ

/TREM2/FFAR1/SCN1B/CTSS/THY1/FXYD1/ANK3/RELN/CXCL9/CCL5/CXCL10/CEMIP/CXCL11/P2RX5/MS4A1/CD19

_2RG/CLU/CD22/STAP1/ADIPOQ

P/PRKCQ/DGKA/CD40LG/SCUBE1/LCK/PLCG2/PLEK/PIK3CG/MMRN1/VAV1/PTPN6/WAS/TLR4/HBB

70/CCL19/EVI2B/IL2RA/FOXP3/SOCS1/IL1B/IL12RB1/CXCR4/RASGRP1/SEMA7A/SEMA4D/HCLS1/FGF2/NCKAP1L/TOX/TNF/TNF

A/CD40LG/SCUBE1/LCK/PLCG2/PLEK/PIK3CG/MMRN1/VAV1/PTPN6/WAS/TLR4/HBB

NB9/NLRP1/LTF/LCK/SPOCK2/TNF/LAPTM5/LAMP3/TNFSF10/BIRC3/SERPINA3/ATP2A3/CR1/COL4A3/MMP9/ST18/AIM2/TIMP4

CQ/DGKA/CD40LG/SCUBE1/LCK/PLCG2/PLEK/PIK3CG/MMRN1/VAV1/PTPN6/WAS/TLR4/HBB

PPP1R16B/SEMA4D/HCLS1/FGF2/HGF/TNF/TNFSF11/GATA3/PIK3CG/FGR/PTPN6/RELN/CCR7/CCL5/NLRC3/MYOC/IL18/AIM2/

2B/LGALS9/CCL8/FGB/PDGFC/GPR183/PTPRR/ICAM1/NDRG4/HAVCR2/CCL21/SPRED3/LYN/TLR9/BMP2/CCL19/IL1B/ACKR3/F

SMAD1/TGFB1/RLN2/PTK2B/ITGAX/TIE1/CYSLTR1/WNT5A/GATA2/FOXC2/ITGA5/CD40/PIK3R6/CYBB/KLF4/EPHA1/ITGB8/IL1B/

SMAD1/RLN2/PTK2B/ITGAX/TIE1/CYSLTR1/WNT5A/GATA2/FOXC2/ITGA5/CD40/PIK3R6/CYBB/KLF4/EPHA1/ITGB8/IL1B/CXCR4/

K3CG/FGR/PTPN6/RELN/CCR7/CCL5/NLRC3/MYOC/IL18/AIM2/CD19/BANK1/NTS

10/CEMIP/CXCL11/P2RX5/MS4A1/CD19

/BIRC3/CCL5/ST18/AIM2/GPD1/TNFRSF13C/CCL20/DCSTAMP/ADIPOQ

MP12/LILRB1/CRTAM/BMP2/IL2RA/FOXP3/SOCS1/IRF1/LAX1/TIGIT/PTPN22/TBX21/JAK3/IL1RN/SEMA4D/LAPTM5/SPN/ADAMDE

/AIM2/FCRL3/TLR10

9/TNF/LAPTM5/TNFSF11/GATA3/SELE/BIRC3/CCL5/ST18/AIM2/GPD1/TNFRSF13C/CCL20/UBD/DCSTAMP/ADIPOQ

3/CD19/NTS

TGB8/SOCS1/LTF/HCLS1/INHBA/TNF/TNFSF11/GATA3/TLR4/CCR7/INPP5D/IL23A/IL33/MMP9/LIF/UBD/DCSTAMP/ADIPOQ

/LILRB1/TLR9/IFIH1/FOXP3/IL1B/IRF1/PRF1/IL12RB1/UNC13D/NLRP1/ISG20/IFIT1/ZBP1/TNF/SPN/IFI44L/IFIT2/APOBEC3A/PTPR

/TMEM106A/NRP1/ADRA2B/EDN1/NODAL/LILRA5/FGFR3/FCGR2B/CCL25/CARD9/ADRA1A/RAMP3/CCL14/FFAR4/DOK6/KSR2/

I0/IL2/CD200R1/GPR31/PARP14/DEFB114/SOCS3/OAS1/PLG/IL12B/PTGER4/PDGFRA/GHRL/TREM2/ISG15/HCK/TGFB1/TNFAIP

IP8L3/CD74/PDGFRA/TREM2/PLA2G5/FGFR2/CD4/TGFB1/XCL2/PTK2B/LGALS9/CCL8/FGB/PDGFC/GPR183/ICAM1/NDRG4/HA

L5/IL23A/CXCL10/CXCL11/CCL20/STAP1/CXCL5/CXCL13

TNF/TRAF3IP3/BTK/SCIMP/LILRA4/TLR6/BIRC3/TLR4/OASL/NLRC3/TLR7/AIM2/FCRL3/TLR10

P/LILRA4/TLR6/BIRC3/TLR4/OASL/NLRC3/TLR7/AIM2/FCRL3/TLR10

CL8/RNASE6/AICDA/NLRP3/SAMHD1/MX1/NLRC5/JAK2/IL15/IRF7/BCL3/PIM2/RSAD2/MMP12/LILRB1/TLR9/IFIH1/CCL19/ITGB8

.34/CAMK1D/CCR4/SUCNR1/CCL19/SLAMF8/IL12A/CXCR4/PLA2G7/SEMA7A/SEMA4D/FGF2/NCKAP1L/KLRK1/GPR18/S1PR1/R

/PIK3CG/RAC2/VAV1/CXCL9/CCR7/CCL5/IL23A/CXCL10/CXCL11/CCL20/CXCL5/CXCL13

CXCL5/CXCL13

PING1/ALOX5/NLRP3/SAMHD1/NR1H3/NLRC5/CST7/IL10RA/HAVCR2/KLF4/LYN/MMP12/LILRB1/A2M/IL2RA/FOXP3/SPINK5/SER

SEMA4A/CCR2/P2RY12/THBS1/CSF1R/SYK/NOD2/CCL22/IL10/CCL24/DEFB114/VEGFC/PTAFR/FPR3/CD74/PDGFRA/CCR8/CC

SEMA4A/CCR2/P2RY12/THBS1/CSF1R/SYK/NOD2/CCL22/IL10/CCL24/DEFB114/VEGFC/PTAFR/FPR3/CD74/PDGFRA/CCR8/CC

CR6/NR5A2/ACKR2/SELP/CCR4/SLC8A1/ZAP70/LRRK2/ACKR3/SLA2/CXCR4/CCR5/NFATC2/TRAT1/PLCG2/ACKR4/TNF/PLEK/T

RSAD2/SPI1/MMP12/TNIP3/IRAK2/KLRC4/SLC15A2/CRTAM/TLR9/IFIH1/LY86/CD300LF/IRF1/CD1D/NLRP1/PYHIN1/IL12A/PIK3AP

PTPRC/NLRC3/IL33/HAMP/IL22RA2/ADIPOQ

R2/CCR4/SLC8A1/ACKR3/CXCR4/CCR5/CD38/S1PR4/ACKR4/PIK3CG/RIC3/CCR7/CXCR3/CD24/XCR1/CD52/CXCR5/CCR6/P2

PLA2G2D/PLA2G2A/CR1/IDO1/ADIPOQ

AP/CLEC4E/SLC15A3/INAVA/GBP5/TXK/LGALS9/TRIM22/KLRC1/RFTN1/SERPING1/MNDA/NLRP3/ADAM8/WNT5A/SAMHD1/NR

CXCL10/IRF8/NOS2/CXCL11/STAP1/CXCL5/CXCL13

P1L/S1PR1/RAC2/CCR7/CCL5/IL23A/CXCL10/CCR6/CXCL13

SCIMP/TLR6/CXCL9/TLR4/MMP9/CXCL10/IRF8/NOS2/CXCL11/TLR10/STAP1/CXCL5/CXCL13

P/LTA/TNIP3/IRAK2/LILRB1/SELP/TLR9/LY86/FOXP3/IL1B/IL12A/TIFAB/CCR5/PTPN22/LTF/CMPK2/PLCG2/KLRK1/LDOC1/CD180/C

MP12/TNIP3/IRAK2/KLRC4/SLC15A2/CRTAM/TLR9/IFIH1/CD300LF/IRF1/CD1D/NLRP1/PYHIN1/IL12A/PIK3AP1/RASGRP1/PTPN2

RP3/ADAM8/WNT5A/FGB/JAK2/IL15/FOXC2/TNFSF14/ITGA5/DOCK8/THY1/PIK3R6/HAVCR2/CCL21/CD80/LYN/SIRPG/LILRB4/EP

A/NLRP3/ADAM8/WNT5A/SAMHD1/NR1H3/NLRC5/CD300A/CTSS/PIK3R6/IRF7/HAVCR2/CLEC6A/LYN/RSAD2/SPI1/MMP12/TNI

P9/CXCL10/IRF8/NOS2/CXCL11/TLR10/STAP1/CXCL5/CXCL13

OR2L13/OR8K1/OR2J2/OR4A16/OR2L3/OR2T12/TMC2/OR4X2/OR8H2/TAS1R1/SEMA5B/OR5AP2/OR4F4/OR5D18/OR5AU1/CXC

LRP3/TCF7/ADAM8/GATA2/IL15/DTX1/TMEM131L/PIK3R6/IRF7/CD80/TMEM176A/LYN/SPI1/GPR171/LILRB4/LILRB1/IL34/CRTAM/

RB1/XAF1/IL1RL1/FOXP3/CD84/TIGIT/IL12A/PTPN22/TBX21/LTF/JAK3/INHBA/HGF/NCKAP1L/TNF/SRGN/GPR18/LAPTM5/GATA3

CR5/NCKAP1L/KLRK1/GPR18/S1PR1/TNFSF11/PIK3CG/RAC2/CORO1A/VAV1/CXCL9/CCR7/CCL5/IL23A/CXCL10/CXCR5/CXCL1

RA/CITED1/HAVCR2/KMO/CD80/LYN/IL18BP/LTA/TNIP3/IRAK2/LILRB1/SELP/TLR9/LY86/FOXP3/IL1B/IL12A/TIFAB/CCR5/PTPN22/

SASH3/CR1/IL33/TLR7/IL18/CCR6/FCRL3/CD22/VPREB3

SF14/CXCL2/CCL21/PADI2/LYN/JAML/IL34/ACKR2/CAMK1D/CCR4/PRKCQ/ITGB2/CCL19/IL1B/SLAMF8/BIN2/ACKR3/IL12A/CXCR

ELN/PTPRC/CCL5/IL23A/IL18/LIF/IL22RA2/ADIPOQ/BANK1

ARD9/NLRC4/DEFB127/GBP4/TF/F2/FGA/GSDMB/SYK/NOD2/IL10/IGHG1/IL27RA/DEFB114/OAS1/SLC9A9/IL12B/IGHG3/TREM2/

/IFIH1/CCL19/LRRK2/FOXP3/CD84/RASGRP1/PTPN22/LTF/PLCG2/CD2/GPR18/SPN/BTK/LILRA4/PTPN6/PTPRC/TLR4/SASH3/IL

/IFIH1/CCL19/LRRK2/FOXP3/CD84/RASGRP1/PTPN22/LTF/PLCG2/CD2/GPR18/SPN/BTK/LILRA4/PTPN6/PTPRC/TLR4/SASH3/IL

C4E/SLC15A3/INAVA/GBP5/TXK/KLRC1/RFTN1/MNDA/NLRP3/C2CD4B/ADAM8/WNT5A/NR1H3/NLRC5/CD300A/IL15/CTSS/IRF7

/LILRB4/LILRB1/TLR9/IFIH1/CCL19/LRRK2/FOXP3/CD84/RASGRP1/PTPN22/LTF/PLCG2/CD2/GPR18/SPN/BTK/LILRA4/PTPN6/P

/LILRB4/LILRB1/TLR9/IFIH1/CCL19/LRRK2/FOXP3/CD84/RASGRP1/PTPN22/LTF/PLCG2/CD2/GPR18/SPN/BTK/LILRA4/PTPN6/P

300A/GPR183/TNFSF14/SMPD3/DOCK8/THY1/CXCL2/SPNS2/ICAM1/CCL21/PADI2/LYN/TACR1/JAML/IL34/CRTAM/SELP/CAMK

AP1L/TIMD4/PLD4/BTK/FGR/CORO1A/VAV1/MARCO/ITGAL/C3/IL15RA/ARHGAP25/HAVCR1/PTPRC/TLR4/IL2RB/ALOX15/IL2RG

HCK/TGFB1/TNFAIP8L2/IL4R/MERTK/SAMSN1/CD86/LGALS9/CLDN18/KLRC1/SFRP1/SERPING1/MNDA/GCSAM/SAMHD1/IL31

J2AF1/TLR4/INPP5D/NLRC3/IL33/TLR7/NOS2/BANK1

J2AF1/TLR4/INPP5D/NLRC3/IL33/TLR7/NOS2/BANK1

A2G7/NCKAP1L/KLRK1/TNF/GPR18/SPN/RAC2/CORO1A/SELE/CCR7/CCL5/CXCR3/IL23A/IL33/CXCL10/CCR6/CCL20/STAP1/CX

OR8G5/OR14C36/OBP2A/OR4C12/OR2B6/OR6Q1/OR2M2/GNAT3/OR4M1/OR10W1/OR2L13/OR8K1/OR2J2/OR4A16/OR2L3/OR2

IL18/HAMP/CSF2RB/LIF/IL22RA2

A/IL12A/JAK3/SEMA4D/LCK/HCLS1/HGF/PLCG2/TNF/CD3E/BTK/FGR/PTPN6/RELN/FLT4/PTPRC/CCL5/IL23A/IL18/EFEMP1/LIF/

A/IL12A/JAK3/SEMA4D/LCK/HCLS1/HGF/PLCG2/TNF/CD3E/BTK/FGR/PTPN6/RELN/FLT4/PTPRC/CCL5/IL23A/IL18/EFEMP1/LIF/

RL3/CD22

FAIP8L2/IL4R/CD86/LGALS9/AIF1/CD70/ALOX5/NLRP3/ADAM8/WNT5A/FGB/TENM3/CD300A/JAK2/IL15/TNFSF14/DTX1/TMEM13

M2/OR4M1/OR10W1/OR2L13/OR8K1/OR2J2/OR4A16/OR2L3/OR2T12/OR4X2/OR8H2/OR5AP2/OR4F4/OR5D18/OR5AU1/OR4S1/C

2/OR4X2/OR8H2/TAS1R1/OR5AP2/OR4F4/OR5D18/OR5AU1/OR4S1/SSC5D/OR5AS1/OR2T10/OR5D14/OR14K1/OR52E5/OR6C4/

FCGR1A/CREB3L3/NLRC4/FFAR4/CDH5/CEBPA/CCR2/FABP4/SYK/NOD2/IL10/NLRP7/CCL24/IL2/CD200R1/GPR31/DEFB114/SC

RTAM/TLR9/TMEM176B/CD28/ZAP70/CCL19/EVI2B/IL2RA/FOXP3/SOCS1/SPINK5/IRF1/IL12RB1/SLAMF8/RASGRP1/TBX21/LTF/J

O1G/TBX21/KLRK1/TNF/S1PR1/TNFSF11/SPN/GATA3/PIK3CG/CORO1A/ITGAL/CCR7/CCL5/CXCL10/CXCL11/CCR6/CCL20/CXC

/IL7R/IL18/RHOH

CCR2/THBS1/CSF1R/SYK/NOD2/GDF2/CD244/HEG1/IL10/IL2/CYP1B1/IL27RA/IL27/PTAFR/OAS1/TLR8/IL12B/CD74/PTGER4/CD2

34/TIGIT/PTPN22/TBX21/CLNK/JAK3/INHBA/RASSF5/LAPTM5/SPN/BTK/FGR/IL4I1/PTPN6/PLA2G2D/PTPRC/INPP5D/TBC1D10C/

6Q1/OR2M2/GNAT3/OR4M1/OR10W1/OR2L13/OR8K1/OR2J2/OR4A16/OR2L3/OR2T12/OR4X2/OR8H2/TAS1R1/OR5AP2/OR4F4/C

/FGR/CD96/IL4I1/PTPN6/PTPRC/INPP5D/ALOX15/CR1/NLRC3/IL33/IL7R/CR2

2/DNASE1L3/RASGRP1/CLNK/PLCG2/CD2/TNF/BTK/PIK3CG/FGR/RAC2/LAT2/PTPN6/TLR6/HAVCR1/PTPRC/TLR4/CCL5/PLA2G

OR2L13/OR8K1/OR2J2/OR4A16/OR2L3/OR2T12/OR4X2/OR8H2/OR5AP2/OR4F4/OR5D18/OR5AU1/OR4S1/OR5AS1/OR2T10/OR5

NFRSF13C/FCRL3

CD3E/SPN/PIK3CG/RAC2/CORO1A/IL4I1/PTPN6/RASAL3/PLA2G2D/PTPRC/CCL5/SASH3/IL23A/PLA2G2A/CR1/CD24/CARD11/TN
ASH3/IL23A/CD24/CARD11/IL18/TNFRSF13C/FCRL3/CSF2RB

31/C3/PTPRC/SH2D1A/IL23A/PLA2G2A/SLAMF6/CR1/IL7R/NKG7/IL18/HAMP/NOS2/FCER2/STAP1

10C/PLA2G2A/CR1/TNFRSF13B/IDO1/BANK1

/CD28/PRKCQ/CCDC88B/ITGB2/ZAP70/CCL19/SIRPB1/IL2RA/FOXP3/SOCS1/IL1B/IL12RB1/ICOS/CD1D/CD40LG/IL12A/ADAM19/
CSF2RA/CEBPA/RBM47/IKBKE/CCR2/IL11/TNFRSF18/CSF1R/IFNA14/SYK/CCL22/CCL24/IL2/IL27RA/PARP14/OAS1/IL12B/CD74/

OXP3/SOCS1/IL1B/IL12RB1/ICOS/CD1D/CD40LG/IL12A/RASGRP1/TBX21/CD38/NFATC2/LCK/CD5/NCKAP1L/KLRK1/TOX/CD6/TN

CG/C3/TLR4/CCR7/IL23A/PLA2G2A/IL33/TLR7/NKG7/IL18/AIM2/STAP1/IDO1
CCL5/SASH3/IL23A/PLA2G2A/CR1/CD24/CARD11/TNFRSF9/IL18/TNFRSF13C/IDO1

Q/CCDC88B/ITGB2/ZAP70/CCL19/SIRPB1/IL2RA/FOXP3/SOCS1/IL1B/IRF1/LAX1/IL12RB1/ICOS/CD1D/CD40LG/TIGIT/IL12A/RASG

RB4/EBI3/LILRB1/CRTAM/SELP/CD28/PRKCQ/CCDC88B/ITGB2/ZAP70/ITGB7/CCL19/SIRPB1/IL2RA/FOXP3/SOCS1/IL1B/IRF1/LA

BC1D10C/PLA2G2A/CR1/ADGRF5/CNR2/TNFRSF13B/IDO1/BANK1
NDA/GCSAM/NLRP3/NR1H3/CD300A/IGHE/CTSS/CD40/THY1/IRF7/HAVCR2/CLEC6A/LYN/PLCL2/RSAD2/LILRB4/TNIP3/IRAK2/

0A/IGHE/CTSS/THY1/IRF7/HAVCR2/CLEC6A/LYN/PLCL2/RSAD2/LILRB4/TNIP3/IRAK2/KLRC4/BMX/SLC15A2/TRDC/TLR9/CD28/
M131L/DOCK8/THY1/PIK3R6/HAVCR2/CCL21/CD80/LYN/SIRPG/LILRB4/EBI3/LILRB1/CRTAM/CD28/PRKCQ/CCDC88B/ZAP70/C

BTK/IL4I1/CD27/TESPA1/PTPN6/IL15RA/PTPRC/INPP5D/SASH3/IL23A/IL2RG/CR1/CARD11/IL7R/IL18/RHOH/FCRL3/IKZF3
ASGRP1/LCK/CD5/NCKAP1L/KLRK1/CD6/TNFSF11/CD3E/SPN/GATA3/CORO1A/IL4I1/CD27/VAV1/TESPA1/PTPN6/RASAL3/FCH

CQ/CCDC88B/ZAP70/IL1RL1/CCL19/LRRK2/SIRPB1/IL2RA/FOXP3/SOCS1/IL1B/IL12RB1/ICOS/CD1D/CD40LG/IL12A/RASGRP1/T
LRB1/SELP/TLR9/CD28/PRKCQ/CCDC88B/ZAP70/IL1RL1/CCL19/LRRK2/SIRPB1/IL2RA/FOXP3/SOCS1/IL1B/IL12RB1/ICOS/CD1D
/PTGDR/ICAM1/HAVCR2/CD80/LYN/PLCL2/SPI1/LILRB1/SUCNR1/CD28/GAPT/ITGB2/CCL19/FOXP3/LCP1/IL12RB1/UNC13D/CD
/BCL3/PTGDR/ICAM1/HAVCR2/CD80/LYN/PLCL2/SPI1/LILRB1/SUCNR1/CD28/GAPT/ITGB2/CCL19/FOXP3/LCP1/IL12RB1/UNC1

/RASGRP1/LCK/CD5/NCKAP1L/KLRK1/CD6/TNF/TNFSF11/CD3E/SPN/GATA3/CORO1A/SELE/IL4I1/CD27/VAV1/TESPA1/PTPN6/

KLRD1/PLCG2/KLRK1/TNF/LAPTM5/GATA3/BTK/FGR/SCIMP/VAV1/PTPN6/C3/IL18R1/RBP4/PTPRC/TLR4/SH2D1A/SASH3/IL23A

B/GNLY/CR1L/LTF/CXCL14/FCMR/TNF/GATA3/POU2F2/IGHA2/PTPN6/C3/POU2AF1/PTPRC/CXCL9/CCR7/SH2D1A/BLNK/JCHA

I1/LILRB4/CRTAM/CD28/ZAP70/CD8A/CCL19/IL2RA/FOXP3/SOCS1/SPINK5/IL1B/IRF1/IL12RB1/CD1D/DOCK2/RASGRP1/PTPN22

AM8/CD300A/GPR183/JAK2/TNFSF8/IL15/TNFSF14/DTX1/TMEM131L/DOCK8/THY1/PIK3R6/HAVCR2/CCL21/CD80/LYN/SIRPG/S

LRB1/CD28/GAPT/CCL19/FOXP3/LCP1/IL12RB1/UNC13D/CD40LG/TBX21/JAK3/PLCG2/NCKAP1L/CD180/APBB1IP/SPN/GATA3/M

70/CCL19/IL2RA/FOXP3/IL1B/IRF1/IL12RB1/CD1D/CD40LG/IL12A/PTPN22/CD38/NFATC2/NCKAP1L/CD6/RASSF5/CD3E/SPN/BT

A1/CD19/PAX5

CD40/HAVCR2/HHEX/CD80/LYN/PLCL2/TACR1/LILRB4/EBI3/LILRB1/IL34/CRTAM/TLR9/CD28/PRKCQ/GAPT/CCDC88B/ZAP70/C

IL12RB1/CD1D/CD40LG/IL12A/PTPN22/CD38/NFATC2/NCKAP1L/CD6/RASSF5/CD3E/SPN/BTK/RAC2/CORO1A/IL4I1/PTPN6/RAS

CD38/NFATC2/NCKAP1L/CD6/RASSF5/CD3E/SPN/BTK/RAC2/CORO1A/IL4I1/PTPN6/RASAL3/PLA2G2D/PTPRC/TLR4/CCL5/INPP

R1/LILRB4/EBI3/LILRB1/CRTAM/TLR9/CD28/PRKCQ/GAPT/CCDC88B/ZAP70/CCL19/IL2RA/FOXP3/IL1B/IRF1/IL12RB1/CD1D/CD4

LILRB1/CRTAM/TLR9/CD28/PRKCQ/GAPT/CCDC88B/ZAP70/CCL19/IL2RA/FOXP3/IL1B/IRF1/IL12RB1/CD1D/CD40LG/IL12A/DOC

P1/TBX21/JAK3/CD38/NFATC2/LYL1/INHBA/PLCG2/NCKAP1L/CD180/LAPTM5/BTK/MFNG/LAT2/IL4I1/CD27/PTPN6/ITM2A/POU2

IF/SPN/GATA3/BTK/IL4I1/PTPN6/WAS/C3/IL18R1/PTPRC/SASH3/ALOX15/IL23A/CR1/IL33/IL7R/IL18/FCER2/CR2

RSF1B/LGALS9/KLRC1/TNFRSF4/SERPING1/NLRP3/WNT5A/CD300A/GATA2/TGFB3/PIK3R6/RAET1L/ICAM1/HAVCR2/CD80/LY

IF/SPN/GATA3/BTK/IL4I1/PTPN6/WAS/C3/IL18R1/PTPRC/SASH3/IL23A/CR1/IL33/IL7R/IL18/FCER2/CR2

LT3/CLEC4E/ZBTB16/FCER1G/IL4R/EOMES/MERTK/CD86/CAMK4/CDH17/PTK2B/LGALS9/USP44/KLRC1/SFRP1/CD79B/AICDA

CD86/CAMK4/CDH17/PTK2B/LGALS9/USP44/KLRC1/SFRP1/CD79B/AICDA/NLRP3/TCF7/ADAM8/STAT4/HDAC9/GPR183/TNFSF

KLRK1/TNF/SPN/GATA3/BTK/CD96/IL4I1/VAV1/PTPN6/KLRB1/WAS/C3/IL18R1/PTPRC/SH2D1A/SASH3/IL23A/SLAMF6/CR1/IL7R/I

/CR1L/IL12A/DNASE1L3/RASGRP1/TBX21/CLNK/JAK3/KLRD1/PLCG2/NCKAP1L/KLRK1/TNF/SPN/GATA3/BTK/FGR/SCIMP/CD9

LA2/MYO1G/UBASH3A/PTPN22/LTF/CD38/TRAC/NFATC2/LCK/KLHL6/TRAT1/KLRD1/PLCG2/NCKAP1L/KLRK1/FCMR/CD247/LA

ATA3/CD3D/LY9/ITK/TRBC1/TRBC2/RASAL3/IL18R1/PLA2G2D/PTPRC/BCL11B/SASH3/IL23A/SLAMF6/IL2RG/CARD11/CD3G/NK

TRDC/CD28/PRKCQ/ZAP70/CD8A/TRGC2/FOXP3/LAX1/PIK3AP1/SLA2/MYO1G/UBASH3A/PTPN22/LTF/CD38/TRAC/NFATC2/LC

D19/IGHD

S/TESPA1/PTPN6/TRBC1/TRBC2/FCHO1/PTPRC/CCR7/SH2D1A/INPP5D/BLNK/CARD11/CD3G/IGLC7/IGHM/FCRL3/CD22/STA

RB/FCER2/CR2/CD19/IGHD

DA/NLRP3/C1S/CD300A/IGHE/GATA2/CD40/MILR1/PIK3R6/IRF7/BCL3/PTGDR/RAET1L/ICAM1/C7/HAVCR2/CD80/LYN/RSAD2/L

RP3/STAT4/C1S/IGHE/JAK2/CD40/IRF7/BCL3/RAET1L/ICAM1/C7/HAVCR2/CD80/CLEC6A/IL18BP/RSAD2/LTA/LILRB4/EBI3/LILRB

74/IGHG3/PDCD1LG2/IGKC/ZNF683/CD226/ADGRE1/TREM2/C1QC/JUNB/CD4/IL6/TGFB1/NLRP10/FUT7/FCER1G/IL4R/GZMM

BCL3/RAET1L/ICAM1/C7/HAVCR2/CD80/RSAD2/LTA/LILRB4/KLRC4/LILRB1/CRTAM/CD28/GAPT/CD8A/C1R/FOXP3/IL1B/IL9R/P

5/UTP15/DROSHA/FASTKD2/EIF5B/SNU13/RSL24D1/MRM2/ERI2/RRP9/HEATR1/EXOSC4/RRP36/TSR3/URB2/DDX27/NSA2/RF

SR2/REXO4/DDX18/EXOSC2/GTF2H5/EXOSC6/GAR1/RIOK2/DDX17/WDR43/PWP1/WDR74/NOL11/WDR46/UTP23/RRP12/RPF2

TIP2/RPUSD1/GEMIN6/GNL3L/PPAN/GRWD1/PRKRA/NSUN5/EXOSC8/MALSU1/LSM2/ZNHIT6/MTERF3/DDX20/NOP16/RPP40

I/RRP36/TSR3/DDX27/NSA2/RRP1/SUV39H1/NOL10/ABT1/DDX10/TSR2/REXO4/TCOF1/GTF3C3/DDX18/EXOSC2/GTF2H5/EXOS

2/NDUFS4/UQCRC2/UQCC3/ATP5F1D/NDUFA10/UQCRQ/COX7A2L/NDUFS3/NUPR1/NDUFB4/NDUFA2/NDUFB6/PPIF/COX7A

SD1/PPAN/TP53RK/URM1/DUS3L/PRKRA/NSUN5/EXOSC8/SMAD2/MTFMT/ZNHIT6/GTPBP3/PTCD1/RPP40/NOL7/QTRT1/USP

QCR11/NDUFA11/CDK1/NDUFA6/NDUFS8/PANK2/NDUFB11/NDUFB2/FH/ACO2/NDUFV1/NDUFB3/COQ9/STOML2/NDUFS4/N

QCC1/NDUFC1/NUBPL/OXA1L/NDUFA9/COA8/COA3/COX16/TACO1/NDUFB5/SURF1/COX18

DT1/GOLGA2/LCMT1/CLASP2/TPR/PHF13/WRAP73/KATNB1/CHEK2/CLASP1/TTK/PIBF1/MYBL2/CCSAP/INCENP/ZW10/SMC2/M

LAC2/GTF3C6/YARS2/TRMT112/METTL2A/CTU2/DTD2/RPP21/GATC/TRNT1/TRMT61A/NAT10/POLR2F/DUS4L/HSD17B10/TRM

FM/MTRES1/TRMT10C/TFAM

4/INCENP/ZW10/STMN1/MAPRE1/CENPH/FBXW11/KIF11/NDE1/HDAC3/DCTN1/FLNA/BORA/KAT5/ARHGEF10

/ANAPC1/RAD21/CHMP3/SEH1L/NCAPD2/SKA1/KIF22/RMDN1/TACC3/UHRF1/BAZ1B/CDT1/RIOK2/GOLGA2/LCMT1/CLASP2/SM

5/NCAPD3/ANAPC15/RB1/NCAPH2/C9orf78/PSMG2/PUM2/BECN1/BRD7/ARID2/NUMA1/KLHL22

A8/NDUFA7/NDUFS2/DLAT/ACLY/MSH2/NDUFS1/COX6A1/UQCC2/COX8A/ATP5ME/UQCR11/NDUFA11/CDK1/NDUFA6/CHCH

TRMT2A/RPUSD4/TRMT13/TRMT1/FAM98B/ADAT1/POP5/POP4/TRMT10A/TPRKB/C2orf49

IPU/RCC2/CENPS/CHAMP1/SLC25A5/TUBG1/MZT1/TNKS/SUN1/CENPE/CCNE1/NDC1/RAD18/PMF1/EHMT2/KNL1/CENPN/UE

DL1/MAD2L1BP/ANAPC11/CDK1/NCAPG2/RANGRF/DCTN2/NCAPH/NUDC/POLDIP2/CHMP1B/ZWINT/CHMP2A/NDEL1/CDC14

2/TTK/MDC1/INCENP/CCAR2/ZW10/FBXO31/RPA2/DYNC1LI1/TELO2/BABAM1/HASPIN/FBXO4/GEN1/MUS81/BRCA2/KNTC1/CI

GCFC2/DHX15/SNU13/SNRNP25/SNRPB/RALY/SNRPG/SF3B4/CCNL2/TSEN2/PRMT1/PPP1R8/BUD31/PRPF3/CRIPT/PPIH/CL[illegible]

31/RAMAC/XRCC5/SNIP1/HSP90AA1/RBMX2/CELF2/STRAP/NCBP1/SRSF12/SRSF9/SRSF10/CD2BP2/SF3B2/PIH1D1/U2AF2/RU

AIP/ORC6/UPF1/EXD2/MCIDAS/GEN1/MUS81/ATAD5/BRCA2/PARP1/OOEP/RFC5/DBF4/DONSON/GINS2/DNA2/PRIM2

SLC25A39/TIMM44/VDAC2/TOMM20L/MTCH1/RHOT2/NAIF1/HSP90AA1/TIMM9/TIMM29/RHOT1/TIMM17A/SLC25A46/CCDC51/T

AOK3/MBTPS2/BABAM2/PSMG2/SYF2/INIP/ABRAXAS1/KLHL22/GIGYF2/RFWD3/TAOK1/CCND1/ZWILCH/FZR1/RAD17

4/PRPF31/RAMAC/XRCC5/SNIP1/HSP90AA1/RBMX2/TFIP11/CELF2/STRAP/NCBP1/SRSF12/SRSF9/FAF2/SRSF10/CD2BP2/SF3[illegible]

2/FANCD2/BCL7A/P3H4/SPDL1/MAD2L1BP/ANAPC11/CDK1/PBRM1/NCAPG2/SMARCC1/RANGRF/DCTN2/NCAPH/DSN1/NUD[illegible]

./FTSJ3/EEF2KMT/PARP1/METTL21A/SETD6/RNMT/THUMPD2/RLF/NTMT1/HEMK1/BUD23/FAM98A/EZH2/CSKMT/METTL18/P

/CLASP1/TTK/TUBGCP5/PIBF1/HAUS1/MYBL2/CCSAP/MAP4/INCENP/STMN1/MAPRE1/CENPH/HAUS3/KIF11/HDAC3/DDB1/D[illegible]

PCID2/CUL3/DPF2

RPF40B/DDX17/PUF60/CRNKL1/EFTUD2/SNRNP40/SRSF3/RBM8A/TGS1/SNRPF/DHX16/KHDC4/PRPF38A/PRPF31/RBM15B/EI

DDX17/PUF60/CRNKL1/EFTUD2/SNRNP40/SRSF3/RBM8A/TGS1/SNRPF/DHX16/KHDC4/PRPF38A/PRPF31/RBM15B/EIF1/SNIF

DDX17/PUF60/CRNKL1/EFTUD2/SNRNP40/SRSF3/RBM8A/TGS1/SNRPF/DHX16/KHDC4/PRPF38A/PRPF31/RBM15B/EIF1/SNIF

3/GCFC2/DHX15/SNU13/SNRNP25/SNRPB/RALY/SNRPG/SF3B4/TSEN2/PPP1R8/BUD31/PRPF3/CRIPT/PPIH/CLNS1A/HNRNPK

AD21/NCAPD2/RTEL1/UCHL5/TACC3/RNF4/BAZ1B/MRE11/SLX4/RIOK2/LCMT1/SMARCD2/CTNNB1/TPR/SMC6/SETDB1/PARN

D1/RFC1/UCHL5/RRM1/RPA3/BLM/POLRMT/CHAF1A/MRE11/TIMELESS/CDT1/SLX4/MCM7/HMGA1/POLD1/WDHD1/OBI1/ZNF

A10/UQCRQ/AIFM2/COX7A2L/NDUFS3/PHGDH/PDIA5/NQO2/NDUFB4/NDUFA2/NDUFB6/COX7A2/NDUFV2

CNB2/TOP2A/KIF23/BOD1/MAD2L1/BIRC5/MSH5/DCAF13/SKA3/HOXA13/GPR3/BUB1B/CENPS/CHAMP1/TUBG1/RAD50/MZT1/S

WIM7/RPA3/XRCC2/BLM/UHRF1/TOP3A/MRE11/TIMELESS/SLX4/MCM7/VPS72/POGZ/SMARCD2/ZNF365/ABL1/RHNO1/BRCA1

P1LC3B/HAX1/MICOS10/SLC25A5/GCLM/ATP5IF1/NDUFB7/MTFR1/TIMM8A/GGCT/ATG4B/MIGA1/LYRM7/PIF1/DNM1L/MTFR1L
AIP1

AD51/PINK1/CDC16/CCNB2/TOP2A/KIF23/BOD1/MAD2L1/SEC16B/BIRC5/MSH5/DCAF13/SKA3/HOXA13/GPR3/BUB1B/CENPS/C

1/KDM1A/MUS81/BRCA2/PARP1/OOEP/DMAP1/CHEK1/NSMCE2/KAT5/PIAS4/BRD8/GINS2/RUVBL1/KLHL15/RAD51AP1/MCMD

VD3/TAOK1/CCND1/FZR1/RAD17/TOPBP1/TTI1/TICRR/FOXO4/SDE2/PIDD1

OX6B1/AKT2/NDUFC2/COX6C/MDH1/IDH3A/GFPT1/PPP1R3F/UQCRC1/NDUFA8/NDUFA7/NDUFS2/DLAT/ACLY/TP53/MSH2/ST

2/PARP1/OOEP/DMAP1/CHEK1/NSMCE2/KAT5/PIAS4/BRD8/GINS2/RUVBL1/KLHL15/RAD51AP1/MCMDC2/XRCC1/INIP/ARID2/SI

OOEP/DMAP1/CHEK1/IER3/ATRIP/MCRS1/KAT5/PIAS4/SUPT20H/RNF8/BRD8/NUDT16L1/RUVBL1/KLHL15/BABAM2/RAD51AP1/

E/GLYR1/SIRT6/PAF1/ZMYND8/EAF1/INTS9/MED26/MED28
AP2K7

04/SDE2/PIDD1/ETAA1/UIMC1/PPP1R10/CDKN1A

)UFB3/ELOVL1/CMPK1/STOML2/NDUFS4/ADCY6/PPCS/UQCC3/ATP5F1D/SLC25A1/NDUFA10/NME6/UCKL1/ATP5MC2

FV1/NDUFB3/ELOVL1/CMPK1/STOML2/NDUFS4/ADCY6/PYGL/PPCS/UQCC3/ATP5F1D/SLC25A1/NDUFA10/NME6/UCKL1/ATP5

FAND1/USP10/FIS1/MTIF3/GABARAPL2/RCHY1/BECN1/PELO/USP30/TIGAR/GIGYF2/FBXO7/WDR45B/PRKAA1/ATG4D/ATG4C/I

1D

DCY6/PPCS/UQCC3/ATP5F1D/SLC25A1/NDUFA10

AJC3/UGGT1/BAG4/NUDCD2/HSP90B1/HSPA4L/HSP90AA1/PDIA5/DNAJB4/DNAJC10/TTC4/QSOX2/PPIF/BAG3/DNAJB2/MPDU1

THOC1/CTDSPL2/NCBP3/THOC7/ENY2/MDN1/NCBP2/PRKACA/PRKD1/DESI1/NUTF2/TSC1/LZTS2/RANGAP1/CPSF6/LSG1/EI

JS4L/VIRMA/HSD17B10/TRMT61B/FBL/CDK5RAP1/FTSJ3/ADAD2/PUS1

NSON/ANAPC15/TAOK3/MBTPS2/RB1/TPRA1/BABAM2/EZH2/PSMG2/SYF2/INIP/BRD7/RBL1/PKD2/ABRAXAS1/KLHL22/GIGYF2/

AD21/CRY1/E2F1/ORC1/BLM/DLG1/BAZ1B/MRE11/TIMELESS/CDT1/LCMT1/MTBP/WDR76/FBXO43/RHNO1/MBTPS1/NME6/BRC

/MBTPS1/BRCA1/CASP2/TPR/CUL4A/TRIAP1/AVEN/CRADD/CHEK2/TTK/MDC1/INCENP/CCAR2/ZW10/FBXO31/RPA2/DYNC1L

COPG1/IER3IP1/SCYL1/YIPF6/VAPA/LLGL1/TRAPPC5/USE1/COPB1/RAB6A/NAPG/ZW10/KIF13A/COPZ2/COG3/BBS1/GOLGA5/B

38/NDUFA7/PTCD1/C1QBP/EIF5A2/OGFOD1/UQCC2/EIF2B3/EIF4G3/DHX9/MTIF2/RMND1/EIF3J/ZNF598/ELP5/HABP4/GSPT1/F

MDM2/IER3/DOT1L/ATRIP/KAT5/SPRED2/DONSON/RNF8/INTS7/TAOK3/USP10/MBTPS2/RAD1/BABAM2/PRPF19/GNB1L/SYF2/

PK3/CSNK1D/RAB2B/COG5/HTT/FHDC1/PLEKHM2

CTR8/PARP3/DPF2

3B/SMARCD1/ACTL6A/MORF4L1/EPC2/PARP3/DPF2/SIRT6/HSF1

SNK1D/VAMP7/COPB2/PPP6C/TFG/ARCN1/MIA3/SEC16A/CUL3/MPPE1

6/CHMP1A/TAOK1/NUP62/HDGFL3/RHOA/STMN2

1/OS9/VCP/UBXN1/NFE2L2

1/FXR2/NUP214/DDX39A/MX2/SMG7/SMG1/NUP50/EIF4E/NUP37/DDX19B/ARC/PCID2/FMR1/XPO1/HSF1/NXF1

SEPTIN11/BBS4/ROCK2/DCTN3/EXOC3/GIT1/RTKN/WASHC5/SON/IQGAP1/EXOC2/SNX18/BECN1/CCP110

RMT3/KMT5C/SNRPF/TRMT112/METTL2A/MTR/PRMT6/SETD5/MECP2/RAMAC/HNMT/SETDB1/RBM15B/RAB6A/TRMT61A/SP

T/SNUPN/NUP155/NUP205/NUP188/IGF2BP2/NUP42/CHTOP/ALYREF/AGFG1/NUP107/KHDRBS1/UPF3B/BICD2/THOC1/PIH1D

MARCA4/TP53/TMEM14B/ARPP19/BCL7A/RINT1/CDC25B/SPDL1/PKD1/GSPT1/MAD2L1BP/ANAPC11/CDK1/RBBP8/ZFYVE19/PB

ZR1/RRM2B/RAD17/TOPBP1/TICRR/CDK6/FOXO4

ARS2/MTFMT/C8orf88/NDUFA7/PTCD1/C1QBP/EIF5A2/OGFOD1/UQCC2/EIF2B3/EIF4G3/DHX9/MTIF2/RMND1/EIF3J/ZNF598/

COX5B/COX7B/XYLB/UQCRB/DERA/COA6/TRAP1/SDHA/FOXK1/ATP5IF1/MYBBP1A/NDUFB7/MTFR1/CYC1/SUCLG1/ATP5F1C/

RM1/SMARCC1/INTS3/ZWINT/RAD51C/CDC14B/RAD21/E2F1/CDKN2C/SIRT2/ORC1/RRM1/BLM/SASS6/PHB2/DLG1/MRE11/RI

/UBE4B/PSMC3/ANAPC1/RNF126/APC2/PSMD1/USP14/NSFL1C/RNF216/PPP2CB/SIRT2/UCHL5/GABARAP/FBXL19/RNF4/TBL1

F8/RSF1/MED21/BDP1/CAND1/MED10/DNAJC9/MED19/MED9/TAF7/MED16/DMC1/MCM2/SMARCA5/NAA60/SHPRH

1/SMARCA4/TP53/MSH2/TMEM14B/FANCD2/BCL7A/RINT1/CDC25B/SPDL1/PKD1/MAD2L1BP/ANAPC11/CDK1/EIF2AK4/RBBP8/

K1/NDUFV1/NDUFB3/FIGNL1/PRKAG1/STOML2/NDUFS4/FOXK2/UQCC3/ATP5F1D/NDUFA10/NME6/UCKL1/ENO4/ATP5MC2/O

AVEN/CRADD/CHEK2/TTK/MDC1/TOM1L1/INCENP/ZW10/FBXO31/RPA2

AAAS/NCBP3/THOC7/ENY2/NCBP2/NUTF2/SRSF1/NUP43/RTRAF/CPSF6/NUP62/FYTTD1/FXR2/NUP214/DDX39A/MX2/SMG7/S

AAAS/NCBP3/THOC7/ENY2/NCBP2/NUTF2/SRSF1/NUP43/RTRAF/CPSF6/NUP62/FYTTD1/FXR2/NUP214/DDX39A/MX2/SMG7/S

BD7/FAF1/PSMA5/PKD1/IFI27/ZNF598/RHBDD2/RNF187/ANAPC11/ANAPC13/PSMD7/SMARCC1/TRIM45/MAEA/USP19/PSMB5/

58/NDUFB11/DTYMK/NDUFB2/PRXL2C/PGK1/NDUFV1/NDUFB3/FIGNL1/PRKAG1/STOML2/NDUFS4/FOXK2/UQCC3/ATP5F1D/

/SON/ENY2/RBM23/PRX/RBM39/DYRK1A/SRSF1/SRSF2/RBM12/RBM15/NCL/CLK1/FAM50A/SNW1/SF1/SF3B3/ILDR2/CWC22/H

AAAS/NCBP3/THOC7/ENY2/NCBP2/NUTF2/SRSF1/NUP43/RTRAF/CPSF6/NUP62/FYTTD1/FXR2/NUP214/DDX39A/MX2/SMG7/S

AE/RAD50/NLE1/CDK4/NABP2/CENPE/PRMT5/CCNE1/SETMAR/MIIP/CTDSPL/CCNG1/TFDP1/HUS1/KNL1/EME1/DTL/TTC28/SM

PL4AL1/BRCA1/CTNNB1/RNF20/MGRN1/AXIN1/UBE2A/KLHL42/PLAA/ZFP91/FBXW11/RNF185/TCF25/FBXO4/GPS2/CUL1/FBXC

/WIPI2/ATP13A2/UFM1/CTSK/SNF8/IFT88/UBXN6/FZD5/RAB1B/ULK3/CSNK2A1/VPS37A/LIX1L/ATP6V1E1/SNX32/SNAPIN/NUPR

AMP1/YKT6/TMEM230/MYO5B/UXT/TRAPPC2L/SUN1/CENPE/GPR143/ZBED3/DNM1L/PMF1/MYO19/KNL1/SKA2/TRAPPC3/TRA

L1/PRKAG1/STOML2/NDUFS4/FOXK2/UQCC3/ATP5F1D/NDUFA10/NME6/ENO4/ATP5MC2/OGDH

/MSH6/SIRT2/SPIRE1/XRCC2/PRR19/FANCA/TOP3A/MRE11/SLX4/GOLGA2
NC2/AP3B1/TRIM58/TRAK1/KIFAP3/TRAF3IP1/BICD2/MAPK8IP3/PCM1

CC3/ATP5F1D/NDUFA10

CENP/SMC2/BAG6/FMN2/ZSCAN21/CDC25A/C14orf39/MSX1/FOXJ3/PSMD13/DDB1/MUS81/BRCA2/OOEP
T1/SPDL1/MAD2L1BP/CDK1/EIF2AK4/RBBP8/ZFYVE19/SCRIB/INTS3/PTPN11/IPO5/ZWINT/CHMP2A/CDC14B/RAD21/CRY1/E2F
RPK1/TIA1/SRSF7/DHX36/SRSF4
/TOMM6/ARL6IP1/PDCD5/SH3GLB1/TIMM21/VPS51/GOLGA7/VPS37B/ITGB1BP1/HRAS/LEPROT/MTX1/PXK/CDK5/BNIP3L/GABA

B3/FIGNL1/PRKAG1/STOML2/NDUFS4/FOXK2/UQCC3/ATP5F1D/NDUFA10/NME6/ENO4/ATP5MC2/OGDH

N3/MREG/BLOC1S1/LCA5/STAU1/BORCS8/FLOT2/HTT/ARL8A/BLOC1S5/BBS12/KIF3B/KIF5B/LAMP1/OPA1
I/SENP3/TGFB2/TAF4/CYREN/SMARCA4/TP53/FKBP1B/LETM1/BCL7A/UBE2V1/DHX9/HDAC6/ZNHIT1/RTCA/TXNDC12/DHFR/F
MKKS/TBCD/CAPZA1/ZWINT/CORO2B/RAD21/APC2/TWF1/LMNA/RTEL1/STYXL1/SPTBN1/SPTBN5/ANKRD27/CARMIL1/CAMS
LGA7/LMNA/SERF2/ZSWIM7/PHB2/TESC/PIN1/TBL1X/PDRG1/SMO/TMEM183A/GIPC1/USP27X/CTNND1/CD81

BP/RHNO1/ADAMTS1/TPR/CUL4A/SMC6/PPP1R35/LSM10/CPSF3/CCNE2/RDX/LSM11/INCENP/SMC2/DBF4B/DYNC1LI1/CSPP

/RSF1/SRF/MED21/BDP1/CAND1/MED10/MED19/MED9/ERCC3/TAF7/PHF2/SMARCD1/MED16/SMARCA5/FOSL1/SNAPC5/PCID

DUFA6/NDUFS8/PANK2/NDUFB11/DTYMK/NDUFB2/PANK4/NDUFV1/NDUFB3/ELOVL1/CMPK1/GUK1/STOML2/NDUFS4/ADCY

N3/MCIDAS/PIP4K2A/YAP1/WDR45/BBS4/PTPRD/TRAPPC12/CNOT6/ARF4/KIF24/USP10/RNF5/CEP295NL/PLK4/MPHOSPH9/PA

A6/NDUFS8/PANK2/NDUFB11/DTYMK/NDUFB2/PANK4/NDUFV1/NDUFB3/ELOVL1/CMPK1/GUK1/STOML2/NDUFS4/ADCY6/SL

3/DDB1/MUS81/BRCA2/OOEP

51C/DYNC1H1/CDC14B/RAD21/CHMP3/NSFL1C/PRKCE/NCAPD2/DUSP3/SIRT2/RRM1/SASS6/PHB2/MARK4/EGF/GIPC1/MTE

O2/PDCD6/RWDD3/AXIN1/HSP90AA1/TGFBR1/PLAA/GPS1/DNAJB2/TCF25/SPRTN/HSPBP1/FBXO4/HDAC3/GPS2/KDM1A/SVBI

3/ELP3/PRMT1/EIF4B/EIF2AK1/GCN1/PCIF1/AGO1/HSPB1/PLD1/SARNP/LSM14B/PUM1/TRUB2/CASC3/CNOT9/TCOF1/DAP/ELI

THOC2/MAGOH/SLC29A1/NUP85/SLBP/NUP153/SLC25A17/POLDIP3/MAGOHB/UPF1/NCBP1/XPOT/SNUPN/NUP155/NUP205/NU

NAJA1/PTPN1/CCND1

33/ELOVL1/GUK1/STOML2/NDUFS4/ADCY6/SLC4A7/PPCS/UQCC3/ATP5F1D/SLC25A1/NDUFA10

/AP3B1/TRAPPC8/TRAPPC12/TRAK1/KLHL12/KIFAP3/EXOC3/SNAP29/ARFGAP3/TRAPPC13/UNC13B/BLOC1S3/EXOC2/COPS5/

/MAP6D1/RDX/CCSAP/LAMP2/WDR1/STMN1/TFIP11/MTRF1L/CFL1/UPF1/JMJD4/SMARCB1/VMP1/FAF2/MTRF1/SSRP1/KIF24

XIN1/HSP90AA1/TGFBR1/PLAA/GPS1/DNAJB2/TCF25/SPRTN/HSPBP1/FBXO4/HDAC3/GPS2/KDM1A/SVBP/DCUN1D5/ARRB1/F

4G3/DHX9/MTIF2/RMND1/ZNF598/DIS3/ELP5/DXO/HABP4/GSPT1/DHX29/PUM3/EIF2B4/RIDA/EIF2AK4/DDX49/EXOSC5/DHFR/F

/WDR45/LAMTOR1/AP3B1/CCDC115/RNF5/ATP6V1F/TMEM165/WASHC5/NPR2/CLN5/VPS16/GABARAPL2/UBQLN2/PHF23/ATI

MP1/PEX10/STUB1/GRWD1/SNX7/WDR72/USP36/TMOD2/SMARCA4/TP53/C1QL1/EIF5A2/MUL1/HSPA8/OGFOD1/MTIF2/HDAC6

APRE1/GAS2L1/BBS1/HAUS3/TTC21B/GEN1/DCTN1/BORA/BBS4/TRIM58/HAUS4/ROCK2/TRAF3IP1/EFNA5/BICD2/PHLDB2/CE

I/NDUFB3/ELOVL1/GUK1/STOML2/NDUFS4/ADCY6/SLC4A7/PPCS/UQCC3/ATP5F1D/SLC25A1/NDUFA10

CSNK2A2/HSPA4/ZFYVE16/SCRIB/POM121C/TNPO1/TIMM17B/MAVS/MTCH2/IPO5/TOMM6/TSG101/PDCD5/TCP1/SH3GLB1/RA

T1/MED28/CCNH/RBM14/TAF10/TAF9/DHX36/MED1/MED24/SUB1/MED17/TAF11L4/SGF29/SPHK2/TAF1

K1/ATP5IF1/ITPA/NDUFB7/SUCLG1/ADCY9/ATP5F1C/ARL2/ATP5PF/DNM1L/PFAS/NDUFC2/NME2/PPAT/NDUFA8/NDUFA7/NDU

2/MAPK8IP3/PCM1

0/TTLL1/CDC14B/CEP78/RAB34/AHI1/B9D1/DNAAF4/PTPDC1/MARK4/NUDCD3/DYNLL1/WDR19/FAM161A/CCDC88A/IFT81/WD

ADCY9/ATP5F1C/ARL2/ATP5PF/DNM1L/PFAS/NDUFC2/NME2/PPAT/NDUFA8/NDUFA7/NDUFS2/ABHD14B/DLAT/ACLY/ACOT7

K1/ATP5IF1/ITPA/NDUFB7/SUCLG1/ADCY9/ATP5F1C/ARL2/ATP5PF/DNM1L/PFAS/NDUFC2/NME2/PPAT/NDUFA8/NDUFA7/NDU

K1A/CREBZF/USP21/MPHOSPH8/MEN1/ZDBF2/EHMT1/METTL3/L3MBTL1/RBM15/PHF2/SMARCD1/MTF2/TRIP12

/B9D1/DNAAF4/PTPDC1/MARK4/NUDCD3/DYNLL1/WDR19/FAM161A/CCDC88A/IFT81/WDPCP/EHD4/IFT88/UNC119B/TMEM2

FRP1/NECTIN3/EMC8/MTCH1/BCS1L/TRAM1/ZDHHC7/GET3/ARL6/RDX/SGTA/SRP9/HSP90AA1/KIF13A/TIMM9/TIMM29/BAG6/

EIF5B/MRM2/CFL2/POLDIP2/CHMP1B/CCDC78/DYNC1H1/CHMP2A/CHMP3/NEB/ABT1/SAC3D1/STYXL1/UHRF1/SASS6/CAPRI

M/UHRF1/TOP3A/EXOSC6/MRE11/TIMELESS/SLX4/MCM7/VPS72/POGZ/ZNF365/ABL1/MND1/RHNO1/BRCA1/SLC15A4/KMT5C

TD/HEXB/B3GAT3/SGSH/PPCS/GSS/HGSNAT/GLRX2/SLC25A1

LE1/DPCD/WDR77/HOXB2/IFT172/MIB1/RARG/CELSR2/DYNC2LI1/CAPRIN2/MEIS1/SMAD2/TP53/POFUT1/LRP6/GAS8/DYNC2

I15B/POLR2G/EIF1/SAMD4B/PTCD2/FUS/RBFOX2/MAGOH/CDK11B/CELF2/TNKS1BP1/CIRBP/MAGOHB/CNOT3/TRA2A/DIS3L2

/ATP6V1B2/LDHB/SDHA/FOXK1/ATP5IF1/ITPA/NDUFB7/SUCLG1/ADCY9/QPRT/ATP5F1C/ARL2/ATP5PF/DNM1L/PFAS/NDUFC2

PC1/TANC2/AP3B1/TRAPPC8/TRAPPC12/TRAK1/KLHL12/KIFAP3/EXOC3/SNAP29/ARFGAP3/TRAPPC13/UNC13B/BLOC1S3/EXC

4/DHFR/RCC1L/MSI2/FASTKD2/GUF1/EIF5B/PDIK1L/ILF3/ELP3/PRMT1/EIF4B/EIF2AK1/GCN1/PCIF1/AGO1/HSPB1/PLD1/SARNP/
CD/CLUAP1/TCTN1/TGFB1I1/LGR4/FEM1B/SALL1/HOXB2/IFT172/MIB1/RARG/MTHFD1L

V1/FAF1/PTK6/DHX9/CDK1/RFC2/PBRM1/SMARCC1/DEK/SMG5/TFPT/NUCKS1/PRDM7/MSH3/RFC4/MAPK1/WRN/EP400/MCM
NAL4/DNAAF4/INTS13/DYNC1I2/KXD1/DYNLL1/WDR19/IFT81/WDPCP/COPG2/IFT88/SLC9B2/STARD9/RHOT2/COPG1/MECP2/
LI1/CAPRIN2/SMAD2/TP53/POFUT1/LRP6/GAS8/DYNC2H1/PALB2/RNF2/HOXA5/NOMO1/HOXD4/HOXD8/MKKS/PGAP1/HOXC5/
133/BANP/CD2AP/KPNA2/UTP25/ARL2BP/NUP35/WWTR1/EFCAB7/HSP90AA1/HDGF/E2F3/NUP85/NUP153/SESN2/BAG3/TSC2/

UBXN6/VPS37A/ARFRP1/PLEKHJ1/AP5Z1/CORO1C/BVES/SNX32/EPS15L1/DPY30/RDX/RAB6A/VPS26C/ENTR1/REPS2/STAM/AL
IP3/VPS37B/TMEM9/RAB34/ATP6V0A1/PHETA1/ANKRD27/AP1S1/ATP13A2/ATP6AP1/SNF8/STX7/RAB22A/VPS37A/PLEKHJ1/STX

LA/ABI2/TWF1/MAP2K2/HRAS/VLDLR/STYXL1/DBN1/ANKRD27/CAPRIN1/CARMIL1/MARK4/DYNLL1/EEF2K/CCDC88A/CYFIP1/A

MB2
P3/APOBEC3A/HAVCR1/CCL5/OASL/CR1/CR2

SLC51B/RBP7/SLC6A5/SLC1A3/PTGS2/PLA2R1/ATP8B1/SLC22A4/EDN1/ADORA2A/SLC38A4/FOLR3/SLC27A2/SLC22A8/FABP4/SL
SLC51B/RBP7/SLC6A5/SLC1A3/PTGS2/PLA2R1/ATP8B1/SLC22A4/EDN1/ADORA2A/SLC38A4/FOLR3/SLC27A2/SLC22A8/FABP4/SL
4A/ARHGAP4/TACR1/ITGB7/KRT2/JCAD/SEMA7A/SEMA4D/FGF2/PLCG2/TNF/GATA3/PIK3CG/CORO1A/FLT4/MMP9/SEMA3A/F

L1B/LAX1/CCND2/RASGRP1/PTPN22/LTF/FGF2/TNF/TNFSF11/PIK3CG/FGR/TLR6/RELN/PTPRC/TLR4/CCR7/CCL5/IL23A/CD2
IBIN/FOSB/ABAT/CPNE5/PTK2B/WNT5A/FGB/OTC/UCP2/CRIP1/S100A16/SMPD3/ACER1/CYBB/SYT1/PGAM2/KCNJ10/CPNE9/T
SLC12A3/SLC9A7/FXYD1/SLC8A1/NETO1/HCN1/SLC38A5/FXYD3/SLC6A7/ANK3/SLC38A1

PRC/ADIPOQ
1L/KLRK1/TNF/LAMP3/SERPINA3/PTPRC/CR1/MMP9/TIMP4/ADIPOQ
2/SFRP4/PLAAT4/BCL11B

B2/FRRS1L/PTN/SLC18A3/CCL2/NTRK2/ARRB2/GRIK4/LGI1/CACNB4/SERPINE2/GUCY1A1/CACNG3/LGMN/BAIAP3/KCNQ3/SY
B2/FRRS1L/PTN/SLC18A3/CCL2/NTRK2/ARRB2/GRIK4/LGI1/CACNB4/SERPINE2/GUCY1A1/CACNG3/LGMN/BAIAP3/KCNQ3/SY

CNB4/KCNC2/KDR/KCNQ3/HTR3D/GABRG1/SYNE1/NALCN/DSG2/SCN3B/MTNR1B/GLRX/P2RX1/ALB/ZACN/SCN3A/DSC2/NP
SCN1B/TNNI2/FXYD1/SLC8A1/TNNT2/CXCR4/TNF/PIK3CG/ATP2A3

PKIA/JAK2/CD40/THY1/PIK3R6/HHEX/LYN/LILRB4/EPHA1/IL34/SLC8A1/BMP2/CCL19/LRRK2/IL1B/LAX1/CCND2/SLAMF8/RASGR
CY1A1/STC1/DSG2/TNNC1/SCN3B/MYH1/MYH4/P2RX1/PTGS2/SCN3A/DSC2/ADRA2B/SYNM/EDN1/CHRNB4/ANKRD2/RYR3/S

:1/PLCB2/MCUB/CACNB4/TRPC7/TSPOAP1/MT1H/CLDN16/STC1/CALCB/CCL7/SLC4A9/LACRT/TRPM2/P2RX1/SLC1A3/ITPR1/S

P4/DENND3/ARHGAP30/LRRK2/FGD3/SH2D3A/DOK3/DOCK2/CCDC88C/RASGRP1/FGD2/ELMO1/SH2D3C/DEF6/RASGEF1A/F(

)OC2B/TNFSF11/RBP4/CCL5/NOS2/BLK

DGRE3/HTR1E/NPY1R/NPY2R/S1PR5/PTH1R/PTGIR/TSHR/RIT2/GPR26/GPHB5/CALCB/ADGRB3/ADGRF1/GIPR/ADRA2B/ADM

A2G4C/PTGDS/RBP4/ALOX15

TARD4/SPNS2/SELENOM/ABCG1/LIPG/OSBP2/CES1/SPNS3/ABCC11/APOL3/PLA2G4C/IL1B/CETP/INHBA/TNF/RBP5/TNFSF11/

CCL20/CXCL13

ETNLB/EGFL7/VSTM4/APOH/ZFP36/LACRT/REG3G/APOA1/MTSS1/NRP1/MARVELD3/ITGA4/NODAL/CLEC12B/ATOH8/BMP5/HM

TNNI2/FXYD1/SLC8A1/TNNT2/CXCR4/TNF/PIK3CG/ATP2A3

UCP2/FFAR1/JAK2/TGFB3/ACHE/ABCG1/RSAD2/VSNL1/SUCNR1/IL1B/PRF1/IL12A/CD38/IL1RN/TNF/DOC2B/PLEK/RBP4/TMEM

)/PPP1R16B/FGF2/TNF/GATA3/BTK/FLT4/BCL11B/CCL5/NLRC3/COL4A3/FGF1/TBX1

P23A/UCP2/FFAR1/JAK2/TGFB3/ACHE/ABCG1/RSAD2/VSNL1/SUCNR1/IL1B/PRF1/IL12A/CD38/IL1RN/TNF/DOC2B/PLEK/RBP4/T

3A/NR1H3/PCLO/TVP23A/UCP2/FFAR1/JAK2/TGFB3/ACHE/ABCG1/RSAD2/VSNL1/SUCNR1/IL1B/PRF1/IL12A/CD38/IL1RN/TNF/D

ARD4/SPNS2/SELENOM/ABCG1/LIPG/OSBP2/CES1/SPNS3/ABCC11/APOL3/IL1B/CETP/INHBA/RBP5/TNFSF11/ABCD2/RBP4/RBI

DIPOQ

EDN1/ERC2/ADORA2A/CHRNB4/EXOC3L1/LIN7A/SYT8/ANO1/FFAR4/F2/IL11/FGA/SYK/RIMS4/ADCY5/GHRL/STX1B/TGM2/TUN

A5/KCNT2/KCNK12/KCNK15/ATP4A/KCNJ10/SLC12A3/SLC9A7/NETO1/HCN1/ANK3

AH/PLCG2/PLA1A/PLCH2/TNF/PIK3CG/PLAAT4/ABCD2/PLA2G2D/PLCXD3/PLA2G2A/PLIN1/ADIPOQ

AM8/WNT5A/FASLG/JAK2/TGFB3/CTSS/PDCD1/CD40/ECSCR/STK17B/LYN/LTA/GZMA/LILRB1/CAMK1D/BMP2/LGALS2/CD40L

SGRF1/ARHGAP15/MYOC/P2RY10

3/KEL/CACNG8/FFAR4/SLC31A2/LRRC26/FXYD2/F2/SCN4B/CCR2/THBS1/CASQ1/KCNJ5/AKAP5/TREM2/PTK2B/CACNA1C/SCN

EM2/SLC9C1/PKD2L1/KCNA5/KCNT2/KCNK12/PTK2B/KCNK15/ATP4A/KCNJ10/SLC12A3/SLC9A7/FXYD1/NETO1/HCN1/FXYD3/A

38/IL1RN/INHBA/TNF/DOC2B/TNFSF11/GATA3/BTK/RBP4/CCL5/NOS2/LIF/BLK/ADIPOQ

P9/FGF1/CXCL13

R/RBP4/TMEM132A/TLR4/BLK

SH3GL2/SLC22A3/TNF/SLC38A1/ATP2A3/CLDN5/HBB/NTS

P9/FGF1/CXCL13

2/TGFB3/SMPD3/ACHE/SYT1/KMO/ABCG1/TACR1/VSNL1/BMP2/ITGB2/IL1B/UNC13D/CD38/INHBA/DOC2B/TNFSF11/FGR/RBP

LAAT4/RELN/FLT4/PTPRC/TLR4/CCR7/CCL5/IL23A/CD24/FGF1/CEMIP/ADIPOQ

DLL4/PTPRR/THY1/NDRG4/CCL21/KLF4/PADI2/ARHGAP4/EPHA1/STK26/SLAMF8/FGF2/KLRK1/TNF/GPR18/IL33/CLDN5/STAP1/

D84/DOCK2/SEPTIN1/PLCG2/NCKAP1L/DOC2B/SFRP4/PACSIN1/BTK/FGR/LRRTM1/RAC2/CORO1A/SELE/C3/IL15RA/PTPRC/IL

RO1A/FLT4/MMP9/FGF1/CXCL13

FXYD3/PIK3CG/ANK3/CORO1A/PTPN6/CXCL9/MMP9/CXCL10/HAMP/CEMIP/CXCL11/P2RX5/MS4A1/CD19

RAF3IP3/BTK/SCIMP/LILRA4/TLR6/BIRC3/TLR4/ALOX15/OASL/NLRC3/TLR7/FCRL3/TBX1

300A/DLL4/PTPRR/THY1/SPINT2/NDRG4/CCL21/KLF4/PADI2/ARHGAP4/EPHA1/STK26/SLAMF8/FGF2/KLRK1/TNF/GPR18/GATA
L/GCH1/SPTBN4/KCNA5/ATP2B3/CACNA1C/CYSLTR1/HOPX/FGB/SCN1B/TACR1/FXYD1/SLC8A1/TNNT2/CD38/TNF/PIK3CG/AT
ELENOM/CYP2C9/BMP2/PLA2G7/GATA3/IL4I1/RBP4/RBP1/PCSK1N

C/PTAFR/DLL1/ADM2/GCH1/ABAT/PTPRO/PCSK5/KLK1/TACR1/SUCNR1/TNF/NOS2/HBB/ADIPOQ
A1/SLC38A5/TNF/SLC6A7/SLC38A1/MS4A1

1I/CCL21/LYN/TLR9/SLC8A1/TSPAN13/CCL19/CCR5/LCK/FGF2/PLCG2/PLCH2/PIK3CG/CORO1A/PTPN6/PTPRC/CXCL9/CCR7/A

G/MYOZ1/CLEC7A/SERPINA10/SLC4A1/F10/ADRA2B/EDN1/CCN4/ADORA2A/EPHB2/PLPP3/HMOX1/F2/P2RY12/FGA/THBS1/SY

STK26/SLAMF8/SEMA7A/SEMA4D/FGF2/KLRK1/TNF/GPR18/GATA3/WAS/IL33/SEMA3A/CLDN5/STAP1/CXCL13/ADIPOQ

38/IL1RN/INHBA/TNF/DOC2B/TNFSF11/GATA3/BTK/RBP4/CCL5/NOS2/LIF/BLK/ADIPOQ

TC2/PTK2B/CACNA1C/FFAR1/SCN1B/THY1/LYN/FGF11/LILRB1/TLR9/FXYD1/SLC8A1/TSPAN13/CD84/NETO1/CXCR4/FXYD3/PI

A2G5/FGFR2/NLRP2/TXK/CD86/XCL2/LGALS9/SFRP1/CCL8/AGAP2/NLRP3/NR1H3/CD300A/FASLG/JAK2/GPR65/CTSS/DOCK8/

ELN/IL18R1/TLR4/NLRC3/CARD11/IL18/CLU/AIM2/NTS

CORO1A/PTPN6/CXCL9/CCL5/CXCL10/CEMIP/CXCL11/FCRL3/P2RX5/MS4A1/CD19

CDHB16/NEXN/AMIGO2/CD84/PCDH15/CDHR1/MPZL2/EMB/CLSTN3

EM2/PTK2B/CACNA1C/SCN1B/CTSS/THY1/ATP4A/LYN/KCNJ10/FGF11/TLR9/FXYD1/SLC8A1/TSPAN13/NETO1/FXYD3/PIK3CG/A

ALB2/SLC8A1/CCL19/CCR5/LCK/FGF2/PLCG2/PLCH2/CORO1A/PTPN6/PTPRC/CXCL9/CCR7/CCL5/ATP2A3/XCR1/CXCL10/CEM

ERPINA10/SLC4A1/F10/ADRA2B/ADM/EDN1/ADORA2A/P2RY2/EPHB2/AQP3/ANO1/AQP4/F2/P2RY12/FGA/THBS1/SYK/C4BPB/TH

ASQ1/MRLN/CDH23/AKAP5/PKD2L1/CD4/ATP2B3/STC2/CACNA2D3/PTK2B/CACNA1C/CCL8/CYSLTR1/FASLG/FFAR1/THY1/CA

ABRE/IFNG/EDN1/ADORA2A/LILRA5/GRIN2A/EPHB2/TMSB4X/STAC3/JPH3/KEL/SLC31A2/CCL4/TF/LRRC26/FXYD2/F2/SCN4B/

LS1/INHBA/NCKAP1L/GATA3/BTK/RAC2/CORO1A/SIT1/IKZF1/CCR7/INPP5D/GPR174/SASH3/PLA2G2A/CARD11/IL7R/ADGRF5/

YBB/HHEX/KLF4/EPHA1/ITGB8/SPINK5/IL1B/CXCR4/JCAD/PPP1R16B/FGF2/TNF/PIK3CG/C3/COL4A3/CXCL10/FGF1/CLDN5/CX

STC2/PTK2B/CACNA1C/CCL8/WNT5A/SV2B/FASLG/CD40/THY1/CCL21/SV2A/LYN/CALB2/SLC8A1/CCL19/CCR5/LCK/FGF2/PLC

LR4/CCR7/CCL5/IL23A/CD24/FGF1/CEMIP/ADIPOQ

IEX/KLF4/EPHA1/ITGB8/SPINK5/IL1B/CXCR4/JCAD/PPP1R16B/FGF2/TNF/PIK3CG/C3/COL4A3/CXCL10/FGF1/CLDN5/CXCL13

P9/ST18/AIM2/TIMP4

RF7/CBFA2T3/LYN/SPI1/GPR171/LILRB4/LILRB1/IL34/BMP2/IL1RL1/CCL19/EVI2B/ITGB8/SOCS1/TIFAB/LTF/HCLS1/INHBA/NCKAP

S/CMA1/ADRA2A/ADORA3/NPY1R/ERAP2/KCNQ1/MYBPC3/GUCY1B1/FGF12/ACTC1/TAC3/NTSR1/KCNH6/MYL4/LNPEP/KCNK

3/PCDHAC1/LRRC4C/CDH18/PCDHA12/PCDHB3/CDH5/CRB2/VCAM1/PCDHB10/NTNG2/CDH11/IL10/MDGA1/CDH23/PECAM1

P2/CCL19/IL1B/ACKR3/RASGRP1/SEMA7A/PTPN22/INHBA/FGF2/TNF/TNFSF11/SCIMP/ALKAL2/PTPN6/MARCO/FLT4/PTPRC/T

[illegible]SF11/GATA3/BTK/IL4I1/CD27/TESPA1/RELN/IL15RA/PTPRC/INPP5D/SASH3/IL23A/IL2RG/CR1/IL33/IL7R/IL18/RHOH/LIF/CLDN5/[illegible]

TMEM100/CD19/BANK1/NTS

RASGRP1/SEMA7A/PTPN22/INHBA/FGF2/TNF/SCIMP/ALKAL2/PTPN6/MARCO/FLT4/PTPRC/TLR4/CCR7/CCL5/ALOX15/TBC1D1

CXCR4/JCAD/FGF2/C3/FGF1

/JCAD/FGF2/C3/FGF1

EC1/IL4I1/PTPN6/PLA2G2D/PTPRC/PLA2G2A/CR1/MYOC/IDO1/ADIPOQ

C/CXCL9/IL23A/OASL/IFIT3/IL33/TLR7/CXCL10/AIM2

/CCL4/SH3RF2/DOK2/IL11/FGA/THBS1/CSF1R/SYK/NOD2/CCL22/PDE6G/CCL24/TNFAIP8L3/PIK3R5/CD74/PDGFRA/GADD45B/[illegible]

8L2/TNFRSF1B/LGALS9/AIF1/KLRC1/PTPRO/SERPING1/ALOX5/NLRP3/WNT5A/SAMHD1/FGB/NR1H3/NLRC5/CST7/IL10RA/HA[illegible]

[illegible]VCR2/CCL21/BMP2/CCL19/ACKR3/RASGRP1/SEMA7A/PTPN22/INHBA/FGF2/TNF/SCIMP/ALKAL2/MARCO/FLT4/PTPRC/TLR4/[illegible]

/FOXP3/IL1B/IRF1/PRF1/IL12RB1/UNC13D/NLRP1/IL12A/ISG20/CXCR4/TBX21/IFIT1/ZBP1/TNF/SPN/GATA3/FGR/POU2F2/IFI44L/

AC2/CCR7/CCL5/IL23A/CXCL10/SEMA3A/FGF1/CCR6/STAP1/CXCL13

PINB9/SLAMF8/AOAH/KLRD1/HGF/SPN/GATA3/SIGLEC10/FGR/CD96/PTPN6/PTPRC/CR1/NLRC3/IL33/HAMP/IL22RA2/ADIPOQ

RL2/EFNB2/TREM2/THBS4/IL6/TGFB1/FCER1G/DEFB133/CMKLR1/XCL2/PTK2B/LGALS9/DAPK2/AIF1/PTPRO/CCL8/CXCR6/CY

RL2/EFNB2/TREM2/THBS4/IL6/TGFB1/FCER1G/DEFB133/CMKLR1/XCL2/PTK2B/LGALS9/DAPK2/AIF1/PTPRO/CCL8/CXCR6/CY

NFSF11/CD3E/BTK/LAT2/SELE/MCTP2/PTPRC/CCR7/CXCR3/TBC1D10C/XCR1/CXCR5/TMEM100/CCR6/CCL20/P2RX5/CD22

1/RASGRP1/PTPN22/CLNK/LTF/ZBP1/KLRD1/PLCG2/KLRK1/CD180/TNF/TRAF3IP3/BTK/SCIMP/LILRA4/VAV1/TLR6/BIRC3/TLR4

RX5/CXCL13

1H3/NLRC5/CD300A/IL15/CTSS/PIK3R6/IRF7/HAVCR2/CLEC6A/LYN/RSAD2/SPI1/MMP12/TNIP3/IRAK2/KLRC4/LILRB1/SLC15A2/

CD6/TNF/BTK/SCIMP/CD96/SELE/GJB6/CXCL9/TLR4/CCR7/MMP9/CXCL10/IRF8/NOS2/CXCL11/CNR2/CSF2RB/STAP1/CXCL5/C

2/CLNK/LTF/ZBP1/KLRD1/PLCG2/KLRK1/TNF/TRAF3IP3/BTK/SCIMP/LILRA4/VAV1/TLR6/BIRC3/TLR4/SH2D1A/CCL5/OASL/SLAM

HA1/EBI3/LILRB1/SELP/CD28/PRKCQ/CCDC88B/ITGB2/ZAP70/CCL19/SIRPB1/IL2RA/FOXP3/SOCS1/IL1B/IL12RB1/ICOS/UNC13D

IP3/IRAK2/KLRC4/LILRB1/SLC15A2/CRTAM/TLR9/A2M/IFIH1/CD300LF/IRF1/SERPINB9/CD1D/SLAMF8/NLRP1/PYHIN1/IL12A/PIK

L12/OR4S1/OR5AS1/OR2T10/OR5D14/OR14K1/OR52E5/OR6C4/PPEF2/OR10C1/OR10R2/OR10G3/RTP1/SCN9A/OR1L4/OR51Q1/

TLR9/TMEM176B/CD28/ZAP70/CCL19/EVI2B/IL2RA/FOXP3/SOCS1/SPINK5/IRF1/IL12RB1/SLAMF8/RASGRP1/TBX21/LTF/JAK3/H

/BTK/CD96/LILRA4/PTPN6/TLR6/ABCD2/PTPRC/TLR4/CCR7/INPP5D/IL23A/CR1/CD24/NLRC3/IL33/RNF128/IDO1/ADIPOQ/BANK
1/CNR2/CCR6/CCL20/STAP1/CXCL5/CXCL13

LTF/CMPK2/PLCG2/KLRK1/LDOC1/CD180/CD6/TNF/BTK/SCIMP/CD96/SELE/TLR6/GJB6/CXCL9/TLR4/CCR7/CD24/MMP9/CXCL

4/PLA2G7/CCR5/CXCL14/FGF2/HGF/NCKAP1L/KLRK1/ACKR4/GPR18/S1PR1/TNFSF11/PIK3CG/RAC2/CORO1A/VAV1/CXCL9/C

ISG15/CD4/IL6/NLRP10/CLEC4E/OAS2/FCER1G/DEFB133/LYZ/GBP5/RNASE6/AICDA/FGB/MPEG1/IGHE/TFEB/TNFSF8/BCL3/HA

23A/NLRC3/IL33/CLU/ADIPOQ
23A/NLRC3/IL33/CLU/ADIPOQ

[illegible]/HAVCR2/MAPK13/CLEC6A/LYN/RSAD2/LTA/SPI1/MMP12/TNIP3/IRAK2/KLRC4/SLC15A2/CRTAM/TLR9/SUCNR1/CD28/ALOX5A[illegible]

TPRC/TLR4/SASH3/IL23A/NLRC3/IL33/CLU/ADIPOQ

TPRC/TLR4/SASH3/IL23A/NLRC3/IL33/CLU/ADIPOQ

[illegible]1D/ITGB2/ZAP70/ITGB7/CCL19/IL1B/SLAMF8/IL12A/CXCR4/PLA2G7/CCR5/MYO1G/TBX21/LCK/NCKAP1L/KLRK1/TNF/GPR18/S1[illegible]

[illegible]/IRF8/STAP1/ADIPOQ

RA/NR1H3/NLRC5/CD300A/GATA2/CST7/TGFB3/DTX1/PDCD1/TMEM131L/MILR1/THY1/HAVCR2/CCL21/CD80/TMEM176A/PAD[illegible]

[illegible]CL13

T12/SCNN1B/OR4X2/OR8H2/TAS1R1/OR5AP2/OR4F4/OR5D18/OR5AU1/OR4S1/OR5AS1/OR2T10/OR5D14/OR14K1/OR52E5/OR[illegible]

[illegible]LK/IL22RA2/ADIPOQ/BANK1

[illegible]LK/IL22RA2/ADIPOQ/BANK1

1L/DOCK8/THY1/PIK3R6/SPINT2/HAVCR2/CCL21/CD80/KLF4/LYN/SIRPG/AKNA/SPI1/LILRB4/EBI3/LILRB1/CRTAM/SELP/CD28/PI

)R5AS1/OR2T10/OR5D14/OR14K1/OR52E5/OR6C4/OR10C1/CNGA2/OR10R2/OR10G3/CNGA4/OR1L4/OR51Q1/OR1L1/OR4L1/OF

OR10C1/OR10R2/OR10G3/RTP1/OR1L4/OR51Q1/OR1L1/OR4L1/OR3A1/KCNMB2/OR2V2/REG3A/OR5B3/OR2B3/OR5T3/OR8D1/

)CS3/IL12B/PTGER4/GHRL/TREM2/NFKBIA/IL6/HCK/TGFB1/NLRP10/FUT7/TNFAIP8L2/NLRP2/IL1R1/TNFRSF1B/ALOX5/NLRP3/

AK3/HCLS1/INHBA/NCKAP1L/TOX/CD2/TNF/TNFSF11/GATA3/BTK/IL4I1/CD27/TESPA1/PTPN6/IL15RA/PTPRC/TLR4/INPP5D/SA

;L13

26/STAT1/TREM2/ISG15/CD4/IL6/TGFB1/NLRP10/SULF2/CLEC4E/OAS2/FCER1G/IL4R/INAVA/NLRP2/IL1R1/GBP5/TXK/CD86/LG

PLA2G2A/CR1/ADGRF5/CNR2/TNFRSF13B/IDO1/BANK1

)R5D18/OR5AU1/OR4S1/OR5AS1/OR2T10/OR5D14/OR14K1/OR52E5/OR6C4/OR10C1/OR10R2/OR10G3/RTP1/OR1L4/OR51Q1/O

;2A/IL33/IL18/ADGRF5/HAMP/CLU/RHOH/CNR2/UBD/FCER2/STAP1/DCSTAMP

D14/OR14K1/OR52E5/OR6C4/OR10C1/OR10R2/OR10G3/OR1L4/OR51Q1/OR1L1/OR4L1/OR3A1/OR2V2/OR5B3/OR2B3/OR5T3/OI

IFRSF9/IL18/TNFRSF13C/IDO1

RASGRP1/LCK/CD5/NCKAP1L/KLRK1/CD6/TNF/TNFSF11/CD3E/SPN/GATA3/ANK3/MMRN1/CORO1A/SELE/IL4I1/CD27/VAV1/TE

CCR8/CCRL2/STAT1/TREM2/IFITM1/ISG15/IL36B/NFKBIA/CD4/IL6/HCK/FLT3/IL18RAP/OAS2/FCER1G/IL4R/IL1R1/TXK/CMKLR1/

NFSF11/CD3E/SPN/GATA3/BTK/CORO1A/IL4I1/CD27/VAV1/TESPA1/PTPN6/RASAL3/IL15RA/FCHO1/PTPRC/TLR4/CCR7/CCL5/IN

RP1/PTPN22/TBX21/JAK3/LCK/CD5/NCKAP1L/KLRK1/CD6/TNF/LAPTM5/TNFSF11/CD3E/SPN/GATA3/CORO1A/SELE/IL4I1/CD27

X1/IL12RB1/ICOS/CD1D/CD40LG/TIGIT/IL12A/RASGRP1/PTPN22/TBX21/JAK3/SEMA4D/LCK/CD5/NCKAP1L/KLRK1/CD6/TNF/LAP

KLRC4/LILRB1/BMX/SLC15A2/TRDC/TLR9/CD28/PRKCQ/ZAP70/CD8A/IFIH1/TRGC2/CD300LF/FOXP3/IRF1/LAX1/NLRP1/PIK3AP

ZAP70/CD8A/IFIH1/TRGC2/CD300LF/FOXP3/IRF1/LAX1/NLRP1/PIK3AP1/SLA2/MYO1G/UBASH3A/PTPN22/LTF/CD38/TRAC/NFAT

CL19/SIRPB1/IL2RA/FOXP3/SOCS1/SPINK5/IL1B/IRF1/LAX1/IL12RB1/ICOS/CD1D/CD40LG/TIGIT/IL12A/RASGRP1/PTPN22/TBX21/

O1/PTPRC/CCR7/CCL5/SASH3/IL23A/IL2RG/CR1/CD24/CARD11/IL7R/IL18/RHOH/TNFRSF13C

BX21/CD38/NFATC2/LCK/CD5/NCKAP1L/KLRK1/TOX/CD2/CD6/TNF/TNFSF11/CD3E/SPN/GATA3/BTK/CORO1A/IL4I1/CD27/VAV

/CD40LG/IL12A/RASGRP1/TBX21/CD38/NFATC2/LCK/CD5/NCKAP1L/KLRK1/TOX/CD2/CD6/TNF/PLEK/TNFSF11/CD3E/SPN/GAT

84/CD40LG/PTGDS/DOCK2/DNASE1L3/RASGRP1/TBX21/CLNK/JAK3/PLCG2/NCKAP1L/CD180/TNF/APBB1IP/SPN/GATA3/BTK/P

3D/CD84/CD40LG/PTGDS/DOCK2/DNASE1L3/RASGRP1/TBX21/CLNK/JAK3/PLCG2/NCKAP1L/CD180/TNF/APBB1IP/SPN/GATA3

RASAL3/FCHO1/PTPRC/CCR7/CCL5/SASH3/IL23A/IL2RG/CR1/CD24/CARD11/IL7R/IL18/RHOH/TNFRSF13C

/SLAMF6/CR1/IL33/TLR7/IL18/NOS2/FCER2/STAP1

AIN/CR1/CXCL10/CLU/CFI/CXCL11/IGHM/CCR6/FCER2/CXCL5/CXCL13/CR2/MS4A1

2/TBX21/JAK3/LCK/NCKAP1L/TOX/CD2/GPR18/CD3E/SPN/GATA3/CD3D/LY9/ITK/IL4I1/CD27/VAV1/THEMIS/TESPA1/IL18R1/PLA2

SPI1/TACR1/LILRB4/EBI3/LILRB1/CRTAM/TLR9/CD28/PRKCQ/CCDC88B/ZAP70/CCL19/SIRPB1/IL2RA/FOXP3/SOCS1/SPINK5/IL1

IFNG/LY9/CORO1A/ITGAL/ITM2A/IL18R1/POU2AF1/PTPRC/TLR4/IL23A/SLAMF6/CR1/NKG7/IL18/IRF8/CCR6/CD19

K/RAC2/CORO1A/IL4I1/PTPN6/RASAL3/PLA2G2D/PTPRC/TLR4/CCL5/INPP5D/SASH3/IL23A/PLA2G2A/CR1/CD24/CARD11/TNFR

CL19/IL2RA/FOXP3/IL1B/IRF1/IL12RB1/CD1D/CD40LG/IL12A/DOCK2/RASGRP1/PTPN22/CD38/NFATC2/NCKAP1L/CD180/CD6/R

SAL3/PLA2G2D/PTPRC/TLR4/CCL5/INPP5D/SASH3/IL23A/PLA2G2A/CR1/CD24/CARD11/TNFRSF9/IL18/TNFRSF13C/FCRL3/IKZF

P5D/SASH3/IL23A/PLA2G2A/CR1/CD24/CARD11/TNFRSF9/IL18/TNFRSF13C/FCRL3/IKZF3/CD22/TNFRSF13B/IDO1

40LG/IL12A/DOCK2/RASGRP1/PTPN22/CD38/NFATC2/NCKAP1L/CD180/CD6/RASSF5/CD3E/SPN/BTK/PIK3CG/RAC2/CORO1A/I

K2/RASGRP1/PTPN22/CD38/NFATC2/NCKAP1L/CD180/CD6/RASSF5/CD3E/SPN/BTK/PIK3CG/RAC2/CORO1A/IL4I1/PTPN6/RAS

AF1/PTPRC/TLR4/INPP5D/BLNK/SASH3/TBC1D10C/IL2RG/CR1/CARD11/IL7R/IRF8/CXCR5/TNFRSF13C/CCR6/FCRL3/IKZF3/C

N/RSAD2/LTA/SPI1/LILRB4/KLRC4/LILRB1/CRTAM/TLR9/CD28/A2M/ITGB2/CCL19/FOXP3/SPINK5/IL1B/IL12RB1/UNC13D/CD84/

/NLRP3/TCF7/ADAM8/IL31RA/STAT4/HDAC9/GPR183/TNFSF8/IL15/DTX1/TMEM131L/DLL4/PIK3R6/IRF7/BCL3/HHEX/CD80/TME

F8/IL15/DTX1/TMEM131L/DLL4/PIK3R6/BCL3/HHEX/CD80/PLCL2/RSAD2/SPI1/LILRB4/CRTAM/TLR9/CD28/ZAP70/CD8A/CCL19/IL

IL18/FCER2/CR2

6/RAC2/IL4I1/VAV1/PTPN6/KLRB1/WAS/C3/IL18R1/PTPRC/TLR4/SH2D1A/SASH3/IL23A/SLAMF6/CR1/IL7R/IL18/NOS2/FCER2/ST

PTM5/CD3E/GATA3/BTK/CD3D/FGR/SCIMP/LAT2/ITK/IGHA2/VAV1/THEMIS/TESPA1/PTPN6/TLR6/TRBC1/TRBC2/C3/FCHO1/P

G7/IL18/WDFY4

K/KLHL6/TRAT1/KLRD1/PLCG2/NCKAP1L/KLRK1/FCMR/CD247/LAPTM5/CD3E/GATA3/BTK/CD3D/FGR/SCIMP/LAT2/ITK/IGHA2

P1/CD79A/BLK/MS4A1/CD19/IGHD/BANK1

LTA/SPI1/LILRB4/KLRC4/LILRB1/CRTAM/TLR9/CD28/GAPT/ITGB2/CD8A/C1R/FOXP3/IL1B/IL9R/PRF1/IL12RB1/UNC13D/CD84/SE

I/CD28/PRKCQ/GAPT/CD8A/C1R/IL1RL1/CCL19/FOXP3/IL1B/IL9R/PRF1/IL12RB1/UNC13D/CD1D/CD40LG/CR1L/IL12A/SLA2/MY

/EOMES/SAMSN1/IL1R1/TXK/CD86/CAMK4/PTK2B/TNFRSF1B/KLRC1/RFTN1/CD79B/CD70/SERPING1/AICDA/NLRP3/FGB/STA

RF1/IL12RB1/UNC13D/SERPINB9/CD1D/CD40LG/CR1L/IL12A/SLA2/MYO1G/RASGRP1/TBX21/CLNK/KLRD1/NCKAP1L/KLRK1/FC

RP1/SUV39H1/NOL10/ABT1/DDX10/TSR2/REXO4/DDX18/EXOSC2/GTF2H5/DDX31/EXOSC6/GAR1/RIOK2/DDX17/PNO1/WDR43/

/TRMT112/UTP25/NIFK/RRP1B/DDX54/UTP18/WDR18/RPUSD2/NAT10/MTREX/TRMT61B/TSR1/FBL/NOL9/IMP4/POP7/LYAR/F

/C1QBP/NOL7/USP36/PTBP2/TFB2M/DHX9/NSUN4/EIF3J/DIS3/UTP14A/SBDS/LAS1L/TRAF7/DHX29/EIF2D/DDX49/RRP15/EIF3I/

C6/GAR1/RIOK2/DDX17/WDR43/PWP1/GTF3C2/WDR74/NOL11/GTF3C6/WDR46/UTP23/RRP12/RPF2/TRMT112/UTP25/NIFK/

A2/NDUFV2/C2orf69/DGUOK

36/TP53/TFB2M/CTU1/DPH3/ADAT2/NSUN4/DIS3/ELP5/TYW3/UTP14A/SBDS/LAS1L/INTS6/DDX49/RRP15/EXOSC5/UTP15/DRO

INT/UQCRC2/UQCC3/ATP5F1D/NDUFA10/UQCRQ

APRE1/DYNC1LI1/HASPIN/DIS3L2/KIF11/GEN1/FLNA/KNTC1/NSMCE2/KAT5/ARHGEF10/NCAPD3/ANAPC15/AAAS/RB1/CENP

MT61B/CDK5RAP1/POP7/PUS1/RTCB/ANG/LARS2/GRSF1/THUMPD2/GTF3C4/TRUB1/MOCS3/FAM98A/PARS2/TSEN54/ELP1/

MARCD2/TPR/PHF13/WRAP73/KATNB1/CHEK2/CLASP1/TTK/PIBF1/MYBL2/CCSAP/INCENP/ZW10/SMC2/MAPRE1/DYNC1LI1/

D5/NDUFS8/PANK2/NDUFB11/ETFB/NDUFB2/FH/ACO2/NDUFV1/NDUFB3/IFNLR1/COQ9/STOML2/CISD1/NDUFS4/NNT/SLC2

E2I/SKA2/BRIP1/SMC1A/SMARCA4/CENPX/TOP3B/ECT2/FANCD2/BCL7A/P3H4/SPDL1/GTF2B/MAD2L1BP/ANAPC11/CDK1/PB

B/RAD21/CHMP3/NSFL1C/SEH1L/NCAPD2/SKA1/KIF22/SIRT2/RMDN1/UHRF1/MKI67/PHB2/BAZ1B/PIN1/CDT1/EGF/GOLGA2/L

HEK1/MDM2/IER3/DOT1L/ATRIP/NAE1/DONSON/INTS7/ANAPC15/DNA2/TAOK3/MBTPS2/RAD1/BABAM2/PRPF19/GNB1L/PSM

NS1A/HNRNPK/LSM7/CASC3/THRAP3/RNF113A/PPIL2/SF3A3/SF3B6/DDX39B/SNRNP200/UPF3A/PRPF40B/DDX17/PUF60/CRNI

VBL1/PRPF19/DDX42/AGO4/CPSF7/MDN1/AGO2/SRSF1/LUC7L2/LUC7L/EIF3A/CPSF6/EIF3L/TSSC4/YJU2/GEMIN2/SF3A2/SNRI

TOMM5/PPIF/SFXN1/TOMM70

B2/PIH1D1/ZFAND1/U2AF2/RUVBL1/PRPF19/DDX42/AGO4/CPSF7/MDN1/AGO2

C/POLDIP2/TTL/CHMP1B/ZWINT/SGO1/DYNC1H1/CHMP2A/ANAPC1/RAD21/CHMP3/SEH1L/NCAPD2/SKA1/KIF22/SIRT2/RMDN

RMT8/METTL8/DNMT1/ANTKMT/MPHOSPH8/MEPCE/WDR6/METTL22/TRMT13/EHMT1/SPOCD1/TRMT1/CBLL1/FAM98B/MRI

CTN1/FLNA/BORA/NTMT1/ARHGEF10/HAUS4

IF1/SNIP1/SNRNP27/RBFOX2/MAGOH/RBMX2/MTREX/TFIP11/CELF2/DHX35/PRPF40A/STRAP/ESS2/CIRBP/RBM41/MAGOHB/

P1/SNRNP27/RBFOX2/MAGOH/RBMX2/MTREX/TFIP11/CELF2/DHX35/PRPF40A/STRAP/ESS2/CIRBP/RBM41/MAGOHB/TRA2A/U

P1/SNRNP27/RBFOX2/MAGOH/RBMX2/MTREX/TFIP11/CELF2/DHX35/PRPF40A/STRAP/ESS2/CIRBP/RBM41/MAGOHB/TRA2A/U

/LSM7/TRUB2/CASC3/THRAP3/RNF113A/PPIL2/SF3A3/SF3B6/DDX39B/SNRNP200/UPF3A/PRPF40B/DDX17/PUF60/CRNKL1/ZR

N/XRCC5/TTK/INCENP/ERCC1/ZW10/SMC2/NAT10/DYNC1LI1/HASPIN/BCL7B/UPF1/FBXO4/SMARCB1/GEN1/PARP1/KNTC1/N

365/CHTF18/BRCA1/SUPT16H/ORC4/RTF2/S100A11/RPA1/RECQL4/CCNE2/DUT/REV3L/FAM111B/PCLAF/WDR18/SSBP1/RPA2

UN1/RAD54B/CENPE/PRMT5/CCNE1/NDC1/EHMT2/KNL1/REEP4/EME1/UBE2I/SKA2/BRIP1/SMC1A/CENPX/FANCD2/BANF1/P

/CSNK2A1/KMT5C/AP5Z1/CHEK2/SMC6/RPA1/RIF1/XRCC5/RECQL4/CHD4/USP51/REV3L/FUS/ERCC1/PPP4R2/FAN1/RPA2/T

/MFF/NDUFC2/TIMM50/ATG9A/TOMM22/MYO19/STUB1/MFSD14A/LONP1/SNX7/NDUFA8/NDUFS2/ZNF205/ROMO1/THEM4/U

HAMP1/TUBG1/RAD50/MZT1/SUN1/MTFR1/RAD54B/CENPE/PRMT5/CCNE1/PEX19/DNM1L/NDC1/MTFR1L/MFF/MYO19/EHM

C2/XRCC1/INIP/ARID2/SFPQ/FBH1/RMI1/RFWD3/UBE2N/RMI2/POLQ/TOPBP1/DMC1/MCM2/HTATSF1/ACTL6A/MORF4L1/EPC

BD1/NDUFS1/COX6A1/UQCC2/COX8A/GYS1/ATP5ME/UQCR11/NDUFA11/CDK1/PFKM/NDUFA6/CHCHD5/NDUFS8/PANK2/N

FPQ/FBH1/RMI1/RFWD3/UBE2N/RMI2/POLQ/TOPBP1/DMC1/MCM2/HTATSF1/ACTL6A/MORF4L1/EPC2/MMS22L/RAD51D/MC

TADA3/ENY2/XRCC1/NSD2/DDX11/FAM168A/BRD7/TAF2/ARID2/ZCWPW1/ABRAXAS1/TIGAR/EYA3/FBH1/UBE2N/RMI2/POLQ/

SMC2

HTT/ARFIP2/HBS1L/KIF5B/NEDD9/MTRES1/ATG14/FKBP8/ABCE1/RNF41/VPS13D/GSK3A/PLIN3/CAMKK2/RRP7A/ELAC1/VCP/P

/HSPBP1

F4ENIF1/XPO6/NUP62/FYTTD1/SMURF1/NUP214/DDX39A/SMG7/SMG1/SDAD1/EIF4E/DDX19B/PCID2/FMR1/XPO1/SIRT6/ABCE

PTEN/FBXO7/RFWD3/TAOK1/CCND1/ZWILCH/FZR1/RAD17/TOPBP1/CTDSP1/TICRR/FOXO4/ZNF655/CDK5RAP2/SDE2/PRMT

CA1/CASP2/TPR/CUL4A/TRIAP1/AVEN/CRADD/CHEK2/TTK/MDC1/TOM1L1/INCENP/CCAR2/ZW10/FBXO31/RPA2/DYNC1LI1/TE

I1/TELO2/BABAM1/HASPIN/FBXO4/GEN1/MUS81/BRCA2/KNTC1/CHEK1/MDM2/IER3/DOT1L/ATRIP/NAE1/DONSON/INTS7/ANA

CAP31/VAMP2/OSBPL5/TMED1/ANKFY1/TRAPPC6A/SEC22C/EXOC8/SEC31B/COG2/TRAPPC1/TRAPPC8/TRAPPC12/RABIF/KL

PTRH1/DHX29/PUM3/EIF2B4/EIF2D/RIDA/EIF2AK4/EIF3I/DHFR/RCC1L/MSI2/FASTKD2/GUF1/EIF5B/RSL24D1/PDIK1L/ILF3/RWD

TTI2/PAXIP1/DYRK1A/ABRAXAS1/GIGYF2/FOXO3/MAPK14/COPS3/RFWD3/TAOK1/SNAI1/CCND1/MAPK3/YJU2/FZR1/RAD17

OUT1/METTL9/VIRMA/ETFBKMT/HSD17B10/TRMT61B/FBL/FTSJ3/EEF1AKMT4/EEF2KMT/PARP1/METTL21A/SETD6/PRDM13/

1/AAAS/RUVBL1/NCBP3/THOC7/ENY2/NCBP2/PUM2/NUTF2/SRSF1/NUP43/RTRAF/CPSF6/NOP10/NUP62/FYTTD1/FXR2/NUP

RM1/SMARCC1/INTS3/POLE/ZWINT/RAD51C/CHMP2A/ANAPC1/CDKN3/CDC14B/RAD21/E2F1/CDKN2C/SIRT2/ORC1/RRM1/

ELP5/HABP4/GSPT1/PTRH1/DHX29/PUM3/EIF2B4/EIF2D/RIDA/EIF2AK4/EIF3I/DHFR/RCC1L/MSI2/FASTKD2/GUF1/EIF5B/RSL24

LYRM7/ARL2/ATP5PF/COX7C/MTFR1L/COX6B1/AKT2/NDUFC2/GNPDA1/COX6C/MDH1/PPARD/IDH3A/QDPR/GFPT1/PPP1R3

OK2/LCMT1/SMARCD2/MTBP/MBTPS1/BRCA1/ADAMTS1/CASP2/TPR/CUL4A/TRIAP1/AVEN/CRADD/CHEK2/TTK/LSM10/CPSF

X/TRIM71/GIPC1/RAD23B/PSMD3/PSMC2/FBXL8/DET1/CTNNB1/GID8/FBXO2/CUL4A/ZYG11B/PSMF1/AXIN1/PSMA2/DMAC2/RI

ZFYVE19/PBRM1/SMARCC1/INTS3/PTPN11/ZWINT/RAD51C/CDC14B/RAD21/CRY1/E2F1/CDKN2C/SIRT2/ORC1/RRM1/BLM/S

GDH

SMG1/NUP50/EIF4E/NUP37/DDX19B/ARC/PCID2/FMR1/XPO1/HSF1/NXF1

SMG1/NUP50/EIF4E/NUP37/DDX19B/ARC/PCID2/FMR1/XPO1/HSF1/NXF1

UBE4B/DDI2/PSMC3/FAM8A1/ANAPC1/RNF126/MAN1A2/APC2/PSMD1/USP14/NSFL1C/ALAD/RNF216/UBQLN1/PPP2CB/SIRT2/

NDUFA10/NME6/UCKL1/ENO4/ATP5MC2/OGDH

INRNPL/SUPT7L/FMR1

SMG1/NUP50/EIF4E/NUP37/DDX19B/ARC/PCID2/FMR1/XPO1/HSF1/NXF1

MARCA4/TP53/MSH2/TMEM14B/FANCD2/BCL7A/RINT1/CDC25B/SPDL1/PKD1/MAD2L1BP/MEIS2/ANAPC11/ANAPC13/CDK1/RB

09/UBE2E1/MDM2/RNF25/CUL4B/TMEM129/NPEPPS/TRIM58/RNF8/FBXO28/TPP2/UBE3A/RNF5/UBE3D/PRPF19/BLMH/UBOX5/

1/SPTLC1/AUP1/LAMP2/TP53INP2/ATG2A/NRBP2/STAM/MVB12A/RHEB/TRIM13/PLAA/SESN2/BAG3/TSC2/TEX264/VPS25/PIP4K

K2/ECT2/SPDL1/YWHAZ/AP3D1/HDAC6/PKD1/RAB27A/TMED9/DCTN2/MKKS/SCRIB/DSN1/NUDC/TTL/CHMP1B/ZWINT/SGO1

'1/SUV39H1/CDK5/ORC1/BLM/DLG1/BAZ1B/MRE11/TIMELESS/CDT1/LCMT1/MTBP/WDR76/FBXO43/ABL1/RHNO1/MBTPS1/NM

RAP/AIFM1/PMPCB/PMPCA/GIPC1/SEC61B/FUT10/SSR3/CHM/TIMM44/FZD5/VPS37A/TOMM20L/BAG4/MTCH1/LONP2/KATNB

PBRM1/CD44/NOL3/SMARCC1/DEK/USP19/TFPT/MAPK1/EP400/IGBP1/FIGNL1/STK24/CORO2B/EIF2AK1/MEAF6/CRY1/MRGBP

SAP2/BAZ1B/SLX4/ARHGAP28/LCMT1/CLASP2/MTBP/FBXO43/NME6/BRCA1/VAT1/FHOD3/TPR/ODF2L/TRIAP1/KATNB1/KANK1/

1/CDC25A/MSX1/APP/GEN1/PAGR1/ATAD5/ANKRD17/OOEP/MDM2/CUL4B/NSMCE2/KAT5/ROCK2/DBF4/BRD4/RRM2/STXBP4

͡2/POLR1E/GLYR1/TFAM/MED26/N6AMT1/NCOA6/MED28/CCNH/RBM14/TAF10/TAF9/DHX36/MED1/MED24/SUB1/MED17/TAF

'6/SLC4A7/RRM1/PPCS/UQCC3/ATP5F1D/SLC25A1/NDUFA10

AN2/UBQLN2/PHF23/ARHGEF5/SNX18/LRRTM2/RAB5A/BECN1/CCP110/TCHP/TAPT1/NUMA1/RAB3GAP2/SMAD4/CHMP6/CHM

.C4A7/RRM1/PPCS/UQCC3/ATP5F1D/SLC25A1/NDUFA10

3P/ABL1/RHNO1/ADAMTS1/HCFC1/TPR/CUL4A/SMC6/PPP1R35/LSM10/CPSF3/CCNE2/RDX/LSM11/INCENP/NUPR1/SMC2/DB

P/DCUN1D5/ARRB1/RASD2/COPS7B/MTA1/PIAS4/COPS2

P6/CAPRIN1/UPF3A/EIF4E2/EEF2K/TRIM71/CNOT11/CYFIP1/BZW1/RBM8A/ELAVL1/CSNK2A1/LARP4/DNAJC3/TPR/C6orf52/YTH

UP188/IGF2BP2/SLC35D1/NUP42/CHTOP/ALYREF/SLC35A5/AGFG1/NUP107/KHDRBS1/UPF3B/BICD2/THOC1/LRRC8A/AAAS/N

KIF1C/AP1AR/SYNJ1/CLN3/ACTN4/MREG/CSNK1D/BLOC1S1/HGS/PPP6C/TFG/HTT/RAB7A/BBS7/EXOC7/EXOC4/BLOC1S5/KIF

/ADD1/ZFAND1/SNAP29/VTA1/VPS16/GABARAPL2/PHF23/ATP2A2/SH3GL1/BECN1/CAPZB/MICAL3/ARID2/PEX14/NAV3/CHMP6

RASD2/COPS7B/MTA1/PIAS4/COPS2

RCC1L/MSI2/FASTKD2/GUF1/EIF5B/DCP1B/CDC37/PDIK1L/ILF3/APEX1/ELP3/PRMT1/DGCR8/EPB41L5/FASTKD1/EIF4B/METTL1

P2A2/SNX18/RAB5A/BECN1/ATG16L1/ARL8B/PPT1/RNASEK/RAB3GAP2/DNAJC16/CHMP6/WDR45B/CHMP1A/CLN3/B4GALNT1/

S/ULK2/CKAP5/ZNF598/GSPT1/PTRH1/SH3PXD2B/EIF2D/SET/CDK1/ULK1/EPG5/UFC1/SMARCC1/CSNK2A2/RETREG2/CDC37/

EP295NL/GIT1/PLK4/CFAP69/ERBB2/HAUS2/DYRK1A/DIXDC1/CLIP1/NAV3/NUMA1/CHMP6/CHMP1A/TAOK1/NUP62/HDGFL3/RH

LA/TIMM21/SPIDR/VPS37B/LMNA/GLI3/CSE1L/LEPROT/MTX1/TRAF3IP2/PXK/BNIP3L/NUP58/AIFM1/PMPCB/PHB2/POM121/KPN

UFS2/ABHD14B/DLAT/ACLY/ACOT7/NDUFS1/HSPA8/PDE8A/ATP5ME/NDUFA11/PFKM/NDUFA6/NDUFS8/PANK2/NDUFB11/M

PCP/EHD4/IFT88/UNC119B/TMEM250/DUSP23/ODF2L/WRAP73/FBXW8/PPP1R35/PIBF1/FSIP2/ARL6/CCDC40/WWTR1/MAP4/

/NDUFS1/HSPA8/PDE8A/ATP5ME/NDUFA11/PFKM/NDUFA6/NDUFS8/PANK2/NDUFB11/MCCC2/NDUFB2/PRXL2C/PANK4/PDE

UFS2/ABHD14B/DLAT/ACLY/ACOT7/NDUFS1/HSPA8/PDE8A/ATP5ME/NDUFA11/PFKM/NDUFA6/G6PD/NDUFS8/PANK2/NDUF

50/DUSP23/ODF2L/WRAP73/FBXW8/PPP1R35/PIBF1/FSIP2/ARL6/CCDC40/WWTR1/MAP4/FUZ/ENTR1/MAPRE1/TMEM107/BBS

GORASP1/TMCO1/BBS1/TOMM5/MYO1C/TOMM70/VAMP2/EMC10/SCARB2/SRP19/SEC61G/PDZK1/EMC4/AP3B1/WDR83OS/M

N1/RNF4/PATL1/GOLGA2/POGZ/WNK3/ATXN2L/CLASP2/DHX30/BRCA1/FHOD3/STARD9/TPR/RPF2/SQLE/G3BP2/LDB3/WRAP

/KPNA2/AP5Z1/SMC6/PMS2/RPA1/RIF1/XRCC5/RECQL4/TSN/CHD4/USP51/REV3L/FUS/ERCC1/PPP4R2/FAN1/RPA2/XRCC6/R

H1/PALB2/MEIS2/RNF2/HOXA5/NOMO1/SMAD5/HOXD4/HOXD8/MKKS/HOXD12/PGAP1/HOXC5/IFT140/HOXC4/RNF220/TDRD6

/UPF1/NCBP1/LSM1/PAN3/SRSF12/RBM42/IGF2BP2/PTBP1/SRSF9/KHDRBS1/SRSF10/UPF3B/PIAS4/CNOT6/ROCK2/SRRM1/R

/MDH1/NME2/PPAT/NDUFA8/NDUFA7/NDUFS2/ABHD14B/DLAT/ACLY/TP53/ACOT7/NDUFS1/HSPA8/DCXR/NADK2/PDE8A/AT

C2/COPS5/TPGS1/KIF1C/AP1AR/PTEN/SYNJ1/CLN3/ACTN4/MREG/CSNK1D/BLOC1S1/HGS/PPP6C/TFG/HTT/RAB7A/BBS7/EX

/LSM14B/PUM1/TRUB2/CASC3/CNOT9/TCOF1/DAP/ELP6/CAPRIN1/UPF3A/EIF4E2/PIN1/EEF2K/TRIM71/CNOT11/CYFIP1/BZW1

/I3/NSMCE1/FIGNL1/TCP1/MBD3/MEAF6/MRGBP/MIS18A/SPIDR/ESCO1/ORC5/MSH6/PCNA/RTEL1/SPIRE1/RAD52/UCHL5/AIF

KATNB1/BICD1/SNAPIN/FSIP2/CCDC40/MAP4/RAB6A/FUZ/KIF13A/KIF7/LCA5L/RHOT1/FMN2/DYNC1LI1/CABYR/BBS1/BLOC1S

IFT140/HOXC4/RNF220/TDRD6/EPB41L5/HOXB5/ETS2

SNUPN/NUP155/FBXO4/MSX1/NMD3/NUP188/HDAC3/NVL/FLNA/PARP1/STK3/DMAP1/YAP1/MDM2/NUP107/MCRS1/BBS4

S2/MVB12A/HOOK2/ARL1/ANKRD50/ERC1/SCARB2/TMEM50A/VPS25/ANKFY1/SNX27/DCTN1/DAB2/FLNA

2/ATP6V1E1/PDCD6/CORO1C/SNAPIN/VAPA/TRAPPC5/ZEB2/KIF13A/STAM/WASL/ALS2/MVB12A/HOOK2/TSNARE1/BLOC1S6/I

P2A1/ABL1/IFT88/WRAP73/KATNB1/CORO1C/STK25/FBXW8/DEF8/ARSB/PPP1R35/MEGF8/CORO1B/HSP90AA1/FUZ/NRDC/EN

LC23A1/SLC6A13/THBS1/SYK/SLC16A9/SLC7A7/PLA2G5/ABAT/LRP2/SLCO2A1/SLCO2B1/ABCC3/UCP2/SLC38A11/KMO/SV2A/KC

LC23A1/SLC6A13/THBS1/SYK/SLC16A9/SLC7A7/PLA2G5/ABAT/LRP2/SLCO2A1/SLCO2B1/ABCC3/UCP2/SLC38A11/KMO/SV2A/KC

GF1/TBX1/CXCL13

I/FGF1/CEMIP/ADIPOQ

TLR9/SUCNR1/ALOX5AP/TNNT2/LRRK2/NFATC2/PLCG2/ANK3/RASGRP2/ALOX15/MMP9/HAMP/HVCN1

NE1/NALCN/DGKE/P2RX1/SLC1A3/PTGS2/NPS/SNCG/EDN1/ERC2/ADORA2A/CHRNB4/GRIN2A/EPHB2/TYROBP/ADRA1A/JPH

NE1/NALCN/DGKE/P2RX1/SLC1A3/PTGS2/NPS/SNCG/EDN1/ERC2/ADORA2A/CHRNB4/GRIN2A/EPHB2/TYROBP/ADRA1A/JPH

S/GLRA3/GABRE/KCNA2/POPDC2/EDN1/ADORA2A/CHRNB4/GRIN2A/DCN/SEZ6/ADRA1A/GRIA4/KCNK13/SOD2/LIPA/SCN4B/I

P1/PTPN22/LTF/FGF2/TNF/TNFSF11/PIK3CG/FGR/TLR6/RELN/FLT4/PTPRC/TLR4/CCR7/CCL5/IL23A/CD24/FGF1/CEMIP/ADIP(

STAC3/HTR1D/ADRA1A/DES/ACTA1/SCN4B/HTR7/CASQ1/MYL2/KCNJ5/TNNI1/PTAFR/PKP2/GHRL/KCNA5/SULF2/ABAT/CACN

SLC22A17/IFNG/SLC4A1/TMPRSS6/EDN1/ERC2/RYR3/JPH3/CCL14/KEL/SLC31A2/CDH5/MT1M/HMOX1/TF/FXYD2/F2/KCTD17/[illegible]

GD5/SEMA4D/FGF2/P2RY8/GPR18/TAGAP/PIK3CG/PSD4/ARHGAP9/RAC2/VAV1/RHOF/WAS/RELN/RASGRP2/RASAL3/ARHGA[illegible]

/EDN1/ADORA2A/HTR1D/CALCRL/ADRA1A/RAMP3/P2RY12/ADCY7/PTGER2/HTR1F/ADCY5/PTGER4/ADM2/ADGRE1/AKAP5/

ABCD2/C3/RBP4/RBP1/PLA2G2D/PLA2G2A/CLU/NOS2/ANO9/OSBPL10/ADIPOQ

MOX1/KLF9/LIPA/TP63/THBS1/NOD2/ODAM/GDF2/IL10/CCL24/VEGFC/IL12B/GHRL/STAT1/EFNB2/THBS4/FGFR2/IL6/ALDH1A2/T

132A/TLR4/CCL5/NOS2/BLK

TMEM132A/TLR4/CCL5/NOS2/BLK

OC2B/PLEK/RBP4/TMEM132A/TLR4/CCL5/NOS2/BLK

P1/PLA2G2D/PLA2G2A/CLU/NOS2/ANO9/OSBPL10/ADIPOQ

AR/SCT/OTOF/KCNA5/DYSF/IL6/ABAT/SLC4A8/SFRP1/PCSK5/ALOX5/FGB/BMP8A/SV2B/ILDR1/PCLO/UCP2/FFAR1/JAK2/SMPD

G/IL12A/TNFAIP8/ZBP1/LCK/INHBA/TNF/SFRP4/TNFSF10/IFIT2/TLR6/PTPRC/TLR4/CCL5/INPP5D/MMP9/LTB/IRF8/CLU/UBD/ID

N1B/CTSS/THY1/ATP4A/LYN/KCNJ10/FGF11/TLR9/FXYD1/SLC8A1/TSPAN13/IL1B/NETO1/FXYD3/TNF/SHISA8/PIK3CG/ANK3/CC

NK3

4/TMEM132A/TLR4/BLK

CXCL13/ADIPOQ

.2RB/ALOX15/IL2RG/CLU/CD22/STAP1/ADIPOQ

3/WAS/IL33/CLDN5/STAP1/CXCL13/ADIPOQ

TP2A3

ATP2A3/XCR1/CACNA2D2/CXCL10/CEMIP/CXCL11/P2RX5/MS4A1/CD19

K/C4BPB/ODAM/THBD/SVEP1/PDPN/PLG/PDGFRA/IL6/TGFB1/FCER1G/MERTK/TXK/HPSE/SERPING1/ALOX5/WNT5A/FGB/GA

IK3CG/ANK3/CORO1A/PTPN6/CXCL9/CCL5/CXCL10/HAMP/CEMIP/CXCL11/FCRL3/P2RX5/MS4A1/CD19

CD40/THY1/CCL21/LYN/RGL4/ARHGAP27/ARHGAP45/EPHA1/BMP2/CCL19/CYFIP2/NLRP1/RASGRP1/RASGEF1A/SEMA4D/S1[illegible]

\NK3/CORO1A/PTPN6/RELN/CXCL9/MMP9/CXCL10/HAMP/CEMIP/ANO9/CXCL11/P2RX5/MS4A1/CD19

/IIP/CXCL11/CD19

BD/HEG1/SVEP1/PDPN/CSN3/PLG/PDGFRA/SCT/FLI1/IL6/OAS2/FCER1G/MERTK/TXK/PTPRO/HPSE/SERPING1/FGB/NR1H3/J

CNA1I/CCL21/LYN/GJA4/LILRB1/TLR9/SLC8A1/TSPAN13/CCL19/CD84/CXCR4/CCR5/LCK/FGF2/PLCG2/PLCH2/PIK3CG/CORO1A

CCR2/P2RY12/CASQ1/MRLN/KCNJ5/PKP2/AKAP5/TREM2/SPTBN4/KCNA5/CD4/STC2/PTK2B/CACNA1C/FFAR1/SCN1B/CTSS/T

HAMP/AIM2/TNFRSF13B

CL13

G2/PLCH2/TNFSF11/CORO1A/PTPN6/PTPRC/CXCL9/CCR7/CCL5/ATP2A3/XCR1/CXCL10/CEMIP/CXCL11/CD19

1L/TNF/TNFSF11/GATA3/RAC2/PTPN6/IKZF1/TLR4/CCR7/INPP5D/IL23A/IL33/MMP9/IRF8/ADGRF5/LIF/UBD/DCSTAMP/ADIPO

6/TEK/ACSM3/PPARG/AGTR1/GUCY1A1/KCNMB1/STC1/CYP4F12/VSTM4/DSG2/TNNC1/SCN3B/MTNR1B/SERPINF2/CPA3/P2I

I/PCDHB5/CDH17/CLDN18/PCDHB7/TENM3/ITGA5/ICAM1/KLF4/PCDHB16/JAML/NEXN/AMIGO2/CRTAM/SELP/ADGRL3/BMP2/

LR4/CCR7/CCL5/ALOX15/TBC1D10C/PLA2G2A/FGF1/CCL20/LIF/ADIPOQ

ƆCSTAMP/ADIPOQ

0C/PLA2G2A/FGF1/CCL20/LIF/ADIPOQ

GHRL/TREM2/PLA2G5/FGFR2/CD4/IL6/TGFB1/FLT3/INAVA/XCL2/PTK2B/LGALS9/CCL8/ADAM8/WNT5A/FGB/PDGFC/GPR183/JA

VCR2/KLF4/PADI2/LYN/TFPI/MMP12/LILRB1/A2M/IL2RA/FOXP3/SPINK5/SERPINB9/SLAMF8/SEMA7A/AOAH/LTF/SEMA4D/KLRD

CCR7/CCL5/ALOX15/PLA2G2A/FGF1/CCL20

IFIT2/APOBEC3A/POU2AF1/PTPRC/CXCL9/CCL5/IL23A/OASL/IFIT3/IL33/TLR7/CXCL10/CLU/AIM2/WDFY4

SLTR1/ALOX5/ADAM8/WNT5A/GPR183/TNFSF14/ECSCR/CXCL2/CCL21/PADI2/LYN/FLRT2/JAML/IL34/ACKR2/CAMK1D/CCR4/S

SLTR1/ALOX5/ADAM8/WNT5A/GPR183/TNFSF14/ECSCR/CXCL2/CCL21/PADI2/LYN/FLRT2/JAML/IL34/ACKR2/CAMK1D/CCR4/S

/SH2D1A/CCL5/OASL/SLAMF6/NLRC3/TLR7/AIM2/FCRL3/TLR10

CRTAM/TLR9/A2M/IFIH1/LY86/CD300LF/FOXP3/SPINK5/IL1B/IRF1/IL12RB1/SERPINB9/CD1D/SLAMF8/NLRP1/PYHIN1/IL12A/PIK

CXCL13/IDO1

IF6/NLRC3/TLR7/AIM2/FCRL3/TLR10

/CD1D/CD40LG/IL12A/ADAM19/RASGRP1/LCK/CD5/SPOCK2/NCKAP1L/KLRK1/CD6/TNF/APBB1IP/TNFSF11/CD3E/SPN/GATA3

3AP1/RASGRP1/PTPN22/CLNK/LTF/ZBP1/KLRD1/PLCG2/KLRK1/TNF/TRAF3IP3/BTK/FGR/SCIMP/CD96/LILRA4/VAV1/PTPN6/TL

OR1L1/OR4L1/OR3A1/OR2V2/OR5B3/OR2B3/OR5T3/OR8D1/OR51E2/OR4A15/OR14J1/PRPH2/OR51A2/OR3A3/LPO/OR6C74/OR[illegible]

CLS1/INHBA/NCKAP1L/TOX/CD2/TNF/TNFSF11/GATA3/BTK/IL4I1/CD27/TESPA1/PTPN6/IL15RA/PTPRC/TLR4/INPP5D/SASH3/IL

K1

L10/IRF8/NOS2/CXCL11/CNR2/CSF2RB/TLR10/STAP1/CXCL5/CXCL13/IDO1

CR7/CCL5/CXCR3/IL23A/XCR1/CXCL10/FGF1/CXCR5/CXCL11/CNR2/CCR6/CCL20/STAP1/CXCL5/CXCL13

AVCR2/LTA/SLC15A2/SELP/TLR9/SPINK5/IL1B/GNLY/SLAMF8/NLRP1/IL12A/LTF/KLRK1/TNF/SPN/FGR/IGHA2/TLR6/TLR4/JCHAIN

\P/IL1RL1/IFIH1/LGALS2/LRRK2/CD300LF/IL1B/IRF1/CD1D/NLRP1/PYHIN1/IL12A/PIK3AP1/PLA2G7/PDE2A/RASGRP1/PTPN22/CLI

PR1/TNFSF11/SPN/GATA3/PIK3CG/RAC2/CORO1A/SELE/VAV1/ITGAL/CXCL9/CCR7/CCL5/CXCR3/IL23A/IL33/SELL/CXCL10/CX(

I2/LYN/PLCL2/SPI1/LILRB4/MMP12/LILRB1/CRTAM/TMEM176B/A2M/IL1RL1/CD300LF/IL2RA/FOXP3/SOCS1/SPINK5/IRF1/LAX1/C

6C4/OR10C1/CNGA2/OR10R2/OR10G3/CNGA4/RTP1/OR1L4/OR51Q1/OR1L1/OR4L1/OR3A1/GFY/BEST2/OR2V2/OR5B3/OR2B3/(

RKCQ/BMP2/CCDC88B/ITGB2/ZAP70/CCL19/SIRPB1/IL2RA/FOXP3/SOCS1/IL1B/IRF1/LAX1/IL12RB1/ICOS/CD1D/CD40LG/TIGIT/[illegible]

[illegible]3A1/GFY/BEST2/OR2V2/OR5B3/OR2B3/OR5T3/OR8D1/OR51E2/OR4A15/OR14J1/OR51A2/OR3A3/OR6C74/OR8J1/OR5H2/OR2F[illegible]

OR51E2/OR4A15/OR14J1/OR51A2/OR3A3/LPO/OR6C74/OR8J1/OR5H2/OR2F2/RTP4/OR52A1/OR4C5/OR11L1/OR6B2/OR4A47/O[illegible]
[illegible]2CD4B/ADAM8/WNT5A/NR1H3/JAK2/CST7/IL15/IL10RA/KLF4/MAPK13/LYN/AKNA/LTA/TLR9/SUCNR1/CD28/ALOX5AP/IL1RL1/L[illegible]
SH3/IL23A/IL2RG/CR1/CARD11/IL7R/IL18/RHOH/FCRL3/LIF/IKZF3/DCSTAMP/ADIPOQ

[illegible]LS9/AIF1/HPSE/RFTN1/NLRP3/ADAM8/WNT5A/JAK2/IL15/IRF7/BCL3/CYBB/HAVCR2/CD80/MAPK13/CLEC6A/RSAD2/LTA/MMP

R1L1/OR4L1/OR3A1/OR2V2/OR5B3/OR2B3/OR5T3/OR8D1/OR51E2/OR4A15/OR14J1/OR51A2/OR3A3/LPO/OR6C74/OR8J1/OR5H

[illegible]8D1/OR51E2/OR4A15/OR14J1/OR51A2/OR3A3/OR6C74/OR8J1/OR5H2/OR2F2/OR52A1/OR4C5/OR11L1/OR6B2/OR4A47/OR13C

ESPA1/PTPN6/RASAL3/FCHO1/PTPRC/CCR7/CCL5/SASH3/IL23A/IL2RG/CR1/CD24/CARD11/IL7R/IL18/RHOH/TNFRSF13C/CXC

XCL2/PTK2B/TNFRSF1B/CLDN18/TNFRSF4/CCL8/CD70/CXCR6/WNT5A/SAMHD1/IL31RA/STAT4/NR1H3/MX1/NLRC5/TNFRS

NPP5D/SASH3/IL23A/IL2RG/CR1/CD24/CARD11/IL7R/IL18/RHOH/TNFRSF13C/FCRL3

/VAV1/TESPA1/PTPN6/RASAL3/FCHO1/PLA2G2D/PTPRC/CCR7/CCL5/SASH3/IL23A/PLA2G2A/IL2RG/CR1/CD24/CARD11/IL7R/I

PTM5/TNFSF11/CD3E/SPN/GATA3/RAC2/CORO1A/SELE/IL4I1/CD27/VAV1/TESPA1/PTPN6/ITGAL/RASAL3/FCHO1/PLA2G2D/PT

1/SLA2/MYO1G/UBASH3A/PTPN22/LTF/CD38/TRAC/NFATC2/LCK/KLHL6/TRAT1/KLRD1/PLCG2/NCKAP1L/KLRK1/FCMR/TNF/

TC2/LCK/KLHL6/TRAT1/KLRD1/PLCG2/NCKAP1L/KLRK1/FCMR/TNF/TRAF3IP3/CD247/LAPTM5/CD3E/GATA3/BTK/CD3D/FGR/

/JAK3/LCK/CD5/NCKAP1L/KLRK1/TOX/CD2/CD6/LAPTM5/TNFSF11/CD3E/SPN/GATA3/RAC2/CORO1A/IL4I1/CD27/SIT1/VAV1/TE

1/TESPA1/PTPN6/TLR6/RASAL3/IL15RA/HAVCR1/FCHO1/PTPRC/TLR4/CCR7/CCL5/INPP5D/SASH3/IL23A/IL2RG/CR1/CD24/CA

TA3/BTK/CORO1A/IL4I1/CD27/VAV1/TESPA1/PTPN6/TLR6/RASAL3/IL15RA/HAVCR1/FCHO1/PTPRC/TLR4/CCR7/CCL5/INPP5D/

PIK3CG/FGR/MFNG/LY9/RAC2/LAT2/CORO1A/PTPN6/ITGAL/ITM2A/IL18R1/POU2AF1/PTPRC/TLR4/IL23A/SLAMF6/CR1/IL33/NK

/BTK/PIK3CG/FGR/MFNG/LY9/RAC2/LAT2/CORO1A/PTPN6/ITGAL/ITM2A/IL18R1/POU2AF1/PTPRC/TLR4/IL23A/SLAMF6/CR1/IL

G2D/PTPRC/CCR7/BCL11B/SASH3/IL23A/SLAMF6/IL2RG/CR1/CARD11/CD3G/TNFRSF9/IL7R/IL18/RHOH/CCR6/IKZF3

B/IRF1/LAX1/IL12RB1/ICOS/CD1D/CD40LG/SLAMF8/TIGIT/IL12A/RASGRP1/PTPN22/TBX21/CLNK/JAK3/CD38/NFATC2/LCK/KLR

SF9/IL33/IL18/TNFRSF13C/FCRL3/CSF2RB/IKZF3/CD22/TNFRSF13B/IDO1

ASSF5/TNFSF11/CD3E/SPN/BTK/PIK3CG/RAC2/CORO1A/IL4I1/PTPN6/RASAL3/PLA2G2D/PTPRC/TLR4/CCL5/INPP5D/SASH3/IL

3/CD22/TNFRSF13B/IDO1

L4I1/PTPN6/RASAL3/PLA2G2D/PTPRC/TLR4/CCL5/INPP5D/SASH3/IL23A/PLA2G2A/CR1/CD24/CARD11/TNFRSF9/IL7R/IL18/TNF

AL3/PLA2G2D/PTPRC/TLR4/CCL5/INPP5D/SASH3/IL23A/PLA2G2A/CR1/CD24/CARD11/TNFRSF9/IL7R/IL18/TNFRSF13C/FCRL

D22/TNFRSF13B/CD79A/CR2/FCRL1/MS4A1/CD19/PAX5/BANK1

SERPINB9/CD1D/CD40LG/SLAMF8/CR1L/IL12A/DNASE1L3/RASGRP1/SEMA7A/TBX21/CLNK/JAK3/KLRD1/PLCG2/NCKAP1L/KLR

EM176A/LYN/PLCL2/RSAD2/SPI1/LILRB4/LILRB1/IL34/CRTAM/TLR9/TMEM176B/CD28/ZAP70/CD8A/CCL19/IL2RA/ITGB8/FOXP3/S

2RA/FOXP3/SOCS1/SPINK5/IL1B/IRF1/IL12RB1/CD1D/CD40LG/SLAMF8/DOCK2/RASGRP1/PTPN22/TBX21/JAK3/LYL1/LCK/INHB

AP1/CR2

TPRC/TLR4/CCR7/SH2D1A/INPP5D/BLNK/CR1/CARD11/CD3G/IGLC7/IGHM/FCRL3/CD22/STAP1/CD79A/BLK/CR2/MS4A1/CD1

/LILRA4/VAV1/THEMIS/TESPA1/PTPN6/TLR6/TRBC1/TRBC2/C3/FCHO1/PTPRC/TLR4/CCR7/SH2D1A/INPP5D/BLNK/CR1/CD24

RPINB9/CD1D/CD40LG/CR1L/IL12A/SLA2/PTGDS/MYO1G/DNASE1L3/RASGRP1/TBX21/CLNK/JAK3/KLRD1/PLCG2/NCKAP1L/K

1G/TBX21/JAK3/KLHL6/KLRD1/NCKAP1L/FCMR/TNF/SPN/GATA3/BTK/LY9/IL4I1/CD27/IGHA2/PTPN6/WAS/C3/IL18R1/PTPRC/T

T4/C1S/MPEG1/IGHE/LAIR1/GPR183/TNFRSF17/JAK2/TFEB/CTSS/PDCD1/CD40/IRF7/BCL3/RAET1L/ICAM1/C7/HAVCR2/CD80/

MR/CD2/TNF/SPN/GATA3/BTK/GZMB/CD96/CORO1A/IL4I1/CD27/IGHA2/VAV1/PTPN6/KLRB1/WAS/C3/IL18R1/PTPRC/SH2D1A/

PWP1/DHX30/WDR74/NOL11/WDR46/UTP23/NOM1/RRP12/CUL4A/RPF2/TRMT112/UTP25/RIOX2/NIFK/XRCC5/RRP1B/DDX54/[illegible]
TSJ3/NOC4L/NVL/WDR75/RIOK3/WDR55/ERCC2/DDX52/RRP8/KRR1/NOP14/ERI1/BUD23/PIH1D1/UTP20/TBL3
EXOSC5/UTP15/NOL3/DROSHA/GCFC2/FASTKD2/EIF5B/SNU13/RSL24D1/MRM2/ERI2/RRP9/SNRPB/HEATR1/SNRPG/SF3B4/[illegible]

RRP1B/DDX54/UTP18/WDR18/RPUSD2/NAT10/MTREX/TRMT61B/TSR1/FBL/NOL9/IMP4/POP7/LYAR/FTSJ3/DEDD/NOC4L/SMA

SHA/SNU13/MRM2/ERI2/THUMPD3/RRP9/INTS3/HEATR1/ELP3/FTSJ1/TSEN2/DGCR8/TDRD6/EXOSC4/RRP36/TSR3/DDX27/[illegible]

K/NCAPH2/PSMG2/NSL1/BECN1/PPP2R1A/NUMA1/ABRAXAS1/KLHL22/TUBG2

IETTL8/KTI12/EXOSC7/POLR3E/RTRAF/WDR6/TRMT2A/RPUSD4/TRMT13/TRMT1/FAM98B/ADAT1/POP5/POP4/MARS2/TRMT[illegible]

HASPIN/BCL7B/DIS3L2/KIF11/SMARCB1/GEN1/FLNA/KNTC1/NSMCE2/KAT5/ARHGEF10/NCAPD3/ANAPC15/AAAS/RB1/CENPK/[illegible]

5A22/UQCRC2/UQCC3/ATP5F1D/NDUFA10/UQCRQ

RM1/NCAPG2/SMARCC1/RANGRF/DCTN2/CSNK2A2/NCAPH/SGO2/DSN1/NUDC/POLDIP2/TTL/CHMP1B/ZWINT/SGO1/DYNC

CMT1/CLASP2/MTBP/FBXO43/NME6/TPR/PHF13/WRAP73/KATNB1/CHEK2/ANKLE2/CLASP1/TTK/PIBF1/MYBL2/TOM1L1/CCSA[illegible]

G2/SYF2/TTI2/INIP/ABRAXAS1/KLHL22/GIGYF2/MAPK14/RFWD3/TAOK1/CCND1/ZWILCH/FZR1/RAD17

KL1/RRAGC/ZRANB2/EFTUD2/SNRNP40/SRSF3/ZC3H10/RBM8A/TGS1/SNRPF/THOC5/DHX16/KHDC4/PRPF38A/PRPF31/SRS

PD3

N1/TACC3/UHRF1/BAZ1B/MRE11/CDT1/RIOK2/GOLGA2/LCMT1/CLASP2/SMARCD2/MND1/TPR/PHF13/PTTG1/WRAP73/KATNE

M3/METTL3/SNRPD3/ATPSCKMT/BHMT/RBM15/TRMT10A

TRA2A/UPF1/NCBP1/LSM1/SRSF12/RBM42/PTBP1/SRSF9/KHDRBS1/SRSF10/UPF3B/HNRNPA3/CD2BP2/SF3B2/SRRM1/RBPM

JPF1/NCBP1/SRSF12/RBM42/PTBP1/SRSF9/KHDRBS1/SRSF10/UPF3B/HNRNPA3/CD2BP2/SF3B2/SRRM1/RBPMS/RBMX/U2AI

JPF1/NCBP1/SRSF12/RBM42/PTBP1/SRSF9/KHDRBS1/SRSF10/UPF3B/HNRNPA3/CD2BP2/SF3B2/SRRM1/RBPMS/RBMX/U2AI

ANB2/EFTUD2/SNRNP40/SRSF3/RBM8A/TGS1/POLR2D/SAFB/SNRPF/THOC5/DHX16/KHDC4/PRPF38A/PRPF31/SRSF11/RAM

SMCE2/MCRS1/MAP2K7/KAT5/TRAPPC12/NCAPD3/ANAPC15/RUVBL1/RB1/MAPKAPK5/NCAPH2/PSMG2/XRCC1/MORC2/BEC

2/DBF4B/CCNA2/LIG1/CDC25A/TRAIP/ORC6/UPF1/EXD2/MCIDAS/GEN1/MUS81/ATAD5/ANKRD17/BRCA2/PARP1/OOEP/TTF1/(

3H4/CDC25B/SPDL1/MAD2L1BP/ANAPC11/CDK1/NCAPG2/RANGRF/DCTN2/NCAPH/PRDM7/NUDC/POLDIP2/CHMP1B/ZWINT

FIP11/FMN2/XRCC6/RNF138/BABAM1/PARPBP/TNKS1BP1/BCL7B/C14orf39/SWI5/EXD2/PPP4C/SMARCB1/GEN1/KDM1A/MUS8

SP36/TP53/TRAK2/NDUFS1/LETM1/MUL1/UQCC2/HDAC6/PKD1/ULK2/GDAP1/RALBP1/NDUFA11/BIK/SDHAF3/CEP89/ULK1/ND

T2/KNL1/REEP4/EME1/UBE2I/SKA2/BRIP1/SMC1A/CENPX/MUL1/FANCD2/BANF1/P3H4/CDC25B/SPDL1/GDAP1/MAD2L1BP/RA

2/MMS22L/RAD51D/MCM6

DUFB11/AGL/ETFB/NDUFB2/FH/ACO2/NDUFV1/NDUFB3/IFNLR1/PHLDA2/COQ9/STOML2/CISD1/NDUFS4/NNT/PYGL/SLC25A2

M6

SMARCE1/TAF7/PPP4R3B/SMARCD1/ACTL6A/MORF4L1/EPC2/SF3B3/RNF168/SUPT7L/ACTR8/PARP3/DPF2/SIRT6/HSF1/UIM

ARL/ADCY10/NEMF/SQSTM1

E1/NXF1/PPM1A

T2/PIDD1/CDC73/PCID2

ELO2/BABAM1/HASPIN/FBXO4/GEN1/MUS81/BRCA2/KNTC1/CHEK1/MDM2/IER3/DOT1L/ATRIP/NAE1/DONSON/INTS7/ANAPC1

APC15/DNA2/TAOK3/MBTPS2/RB1/RAD1/TPRA1/BABAM2/PRPF19/EZH2/GNB1L/PSMG2/SYF2/TTI2/INIP/BRD7/RBL1/PKD2/ABRA

HL12/ARF4/CREB3L2/RABEP1/BICD2/GOLT1B/BLZF1/SEC23B/BET1L/ARF5/NRBP1/YIPF7/ERGIC2/ACSL3/ARFGAP3/TRAPPC13

D1/GADD45GIP1/AARSD1/ELP3/FTSJ1/PRMT1/EIF4B/EIF2AK1/SARS2/GCN1/PCIF1/AGO1/EEF1B2/FARSA/HSPB1/ABCF1/PLD1/

RNMT/THUMPD2/RLF/RRP8/NTMT1/HEMK1/BUD23/SETD1A/FAM98A/COQ5/EZH2/CSKMT/METTL18/PRMT8/METTL8/NSD2/D

P214/DDX39A/NOP58/MX2/STAU1/SMG7/SMG1/NUP50/EIF4E/NUP37/DDX19B

TACC3/BLM/SASS6/PHB2/DLG1/MRE11/TRIM71/RIOK2/LCMT1/CLASP2/SMARCD2/MTBP/MBTPS1/CCNI/BRCA1/ADAMTS1/CA

D1/PDIK1L/ILF3/RWDD1/GADD45GIP1/AARSD1/ELP3/FTSJ1/PRMT1/EIF4B/EIF2AK1/SARS2/GCN1/PCIF1/AGO1/EEF1B2/FARSA

F/UQCRC1/NDUFA8/NDUFA7/NDUFS2/DLAT/ACLY/TP53/MSH2/STBD1/NDUFS1/COX6A1/UQCC2/COX8A/GYS1/ATP5ME/UQC

3/CCNE2/RDX/LSM11/INCENP/ZW10/FBXO31/RPA2/DBF4B/DYNC1LI1/CDC25A/BABAM1/HASPIN/BCL7B/APP/SMARCB1/DDB1

NF145/FBXW8/WWTR1/FEM1A/UBE2A/TRIB1/CCAR2/PSMC4/FBXO31/KLHL42/RNF122/BAG6/GNA12/LTN1/TRIM13/PLAA/TRAF

ASS6/PHB2/DLG1/MRE11/TIMELESS/CDT1/RIOK2/LCMT1/SMARCD2/MTBP/WDR76/RHNO1/MBTPS1/BRCA1/ADAMTS1/CASP

/UCHL5/DNAAF4/GABARAP/FBXL19/RNF4/TBL1X/TRIM71/CCDC47/GIPC1/SEC61B/RAD23B/UFD1/PSMD3/PSMC2/FBXL8/UBXN

BP8/ZFYVE19/PBRM1/SMARCC1/MAEA/SCRIB/INTS3/POLDIP2/TTL/PTPN11/ZWINT/RAD51C/ANAPC1/CDC14B/RAD21/E2F1/F

/RNF167/UBE2W/RNF146/RNF115/FBXO7/UBE2D2/DDA1/RBX1/UBE2B/UBE2N/NHLRC1/C10orf90/ZNF738/TRIM65/FZR1/SMUR

2A/EXOC8/VMP1/PGAM5/RB1CC1/ATP6V0C/WDR45/SPATA33/ACBD5/PJVK/RNF5/SNAP29/CAPNS1/ATP6V1C1/LZTS1/SUPT5H

/TSG101/DYNC1H1/CHMP2A/PEF1/NDEL1/CHMP3/NSFL1C/SEH1L/LMNA/SKA1/KIF22/CDK5/SIRT2/EXOC6B/RMDN1/SPIRE1/K

E6/BRCA1/CTNNB1/CASP2/TPR/CUL4A/TAF6/TRIAP1/AVEN/CRADD/CHEK2/TTK/MDC1/TOM1L1/INCENP/NUPR1/CCAR2/ZW

1/TRAM1/ZDHHC7/ARL6/SGTA/SRP9/HSP90AA1/LAMP2/MON1A/TIMM9/TIMM29/BAG6/YWHAB/TIMM17A/GNPTG/TOMM5/MY

/USP14/SPIDR/MAP2K2/HNRNPK/UBQLN1/PCNA/RTEL1/FOXO1/SPIRE1/RAD52/UCHL5/LRIG2/TIMELESS/VPS72/SMARCD2/AI

/ARPIN/CLASP1/TTK/TOM1L1/MAP6D1/RDX/MAP4/INCENP/CORO1B/NUPR1/ERCC1/ZW10/NAT10/STMN1/MAPRE1

/NCAPD3/RB1/RAD51AP1/EZH2/PLK4/ZNF16/NCAPH2/NPR2

11L4/TAF9B/SGF29/SPHK2

MP1A/NBDY/NUP62/RHOA/C10orf90/MAPK9/PRKAA1/RALB/TMEM39A

F4B/DYNC1LI1/CSPP1/CDC25A/MSX1/APP/GEN1/PAGR1/ATAD5/ANKRD17/BRCA2/OOEP/CHEK1/MDM2/CUL4B/NSMCE2/KAT[illegible]

HDF3/POLR2G/EIF1/SAMD4B/SRP9/MAGOH/DAPK3/INPP5E/NAT10/SECISBP2L/CDK5RAP1/SESN2/STK35/AJUBA/CIRBP/POLD[illegible]

CBP3/SLC25A25/ABCC4/THOC7/ENY2/NCBP2/SLC35A3/NUTF2/SLC29A3/SRSF1/NUP43/RTRAF

3B/KIF5B/SEC16A/CUL3/RAB11A/PDCD6IP/EXOC5

/CHMP1A/SYNJ1/TAOK1/CLN3/HDGFL3/STMN2/MTPN/TMEM39A/PEX5/GSPT2/SMARCE1/TOM1

/EIF2AK1/GCN1/PCIF1/AGO1/E2F1/HSPB1/PLD1/SARNP/LSM14B/TRAF3IP2/PUM1/TRUB2/CASC3/CNOT9/TCOF1/DAP/THRAP[illegible]

/SMURF1/EXT1/RAB23/VPS54/ATG4D/RALB/TMEM39A/ATP6V1D/TOM1/ATG4C/ARFIP2/RAB7A/PACS2/RAB14/HOOK3/FNIP1/C[illegible]

ATG10/CTDNEP1/CAPZA1/CFL2/FUNDC2/FUNDC1/CHMP1B/TSG101/SNX30/CHMP2A/GCN1/APC2/CHMP3/TWF1/MMP14/NC[illegible]

HOA/C10orf90/STMN2/CHORDC1

NA6/IPO9/PMPCA/UFM1/SMO/SEC61B/SNF8/SSR3/TIMM44/ELAVL1/FZD5/VPS37A/TPR/TOMM20L/NUP133/BAG4/KPNA2/MTCH

CCC2/NDUFB2/PRXL2C/PANK4/PDE4D/PGK1/NDUFV1/NDUFB3/FIGNL1/ELOVL1/CMPK1/PRKAG1/GUK1/STOML2/NDUFS4/AD

FUZ/LCA5L/ENTR1/MAPRE1/TMEM107/BBS1/SPATA6/LRRC61/TTC21B/GORAB/CEP41/MCIDAS/SPACA9/SSX2IP/DCTN1/ATXN

E4D/PGK1/NDUFV1/NDUFB3/FIGNL1/ELOVL1/PRKAG1/GUK1/STOML2/NDUFS4/ADCY6

B11/MCCC2/NDUFB2/PRXL2C/PANK4/PDE4D/PGK1/NDUFV1/NDUFB3/FIGNL1/ELOVL1/CMPK1/PRKAG1/GUK1/STOML2/NDUF

1/SPATA6/LRRC61/TTC21B/GORAB/CEP41/MCIDAS/SPACA9/SSX2IP/DCTN1/ATXN10/HYLS1/FLNA/YAP1/BBS4/DNAL1/KIAA075

AIP1/RAB5IF/EMC3/BLZF1/SEC61A2/FIS1/HSPA5/ACSL3/TRAM2

73/CHEK2/YTHDF3/CLASP1/BICD1/XRCC5/PPP1R35/TUBGCP5/PIBF1/HAUS1/MYBL2/CCSAP/INCENP/ACTG1/AUP1/CEP152/W

NF138/PARPBP/C14orf39/SWI5/EXD2/PPP4C/GEN1/KDM1A/PAGR1/MUS81/ATAD5/TCF3/BRCA2/APEX2/LEF1/PARP1/OOEP/DM

/EPB41L5/HOXB5/ETS2/APC2/HOXA3/GLI3/CHRDL1/BCOR/AHI1/RFNG

BPMS/RBMX/U2AF2/VIM/PRPF19/FASTKD5/SON/PAN2/RBM23/AGO2/NCBP2/PUM2/RBM39/TOB1/DYRK1A/EXOSC7/DCPS/YT

TP5ME/NDUFA11/PFKM/NDUFA6/SARM1/G6PD/NDUFS8/PANK2/NDUFB11/MCCC2/NDUFB2/PRXL2C/PANK4/PDE4D/PGK1/NE

OC7/EXOC4/BLOC1S5/KIF3B/KIF5B/SEC16A/CUL3/RAB11A/PDCD6IP/EXOC5

/RBM8A/ELAVL1/CSNK2A1/LARP4/DNAJC3/TPR/C6orf52/YTHDF3/POLR2G/EIF1/SAMD4B/SRP9/SLC2A13/MAGOH/DAPK3/INPP

M1/BLM/EXOSC6/MRE11/TIMELESS/CDT1/SLX4/MCM7/VPS72/MPV17/SMARCD2/OBI1/ZNF365/NPAS2/ABL1/CHTF18/BRCA1/C
6/BAG3/FBXW11/TTC21B/SSX2IP/KIF11/NDE1/APP/SLIRP/DTNBP1

RAB5B/VPS18/VAMP2/SCARB2/S100A10/VPS25/PIP4K2A/DTNBP1/ANKFY1/TRAPPC6A/EXOC8/SEC31B/TRAPPC1/AGFG1/ATP6V

ITR1/CBFA2T2/TGFBR1/WASL

NJ10/CES1/ABCC11/IL1B/SLC38A5/EMB/SLC22A3/TNF/SLC6A7/RBP5/TNFSF11/ABCD2/SLC38A1/RBP1/PLA2G2D/PLA2G2A/NOS
NJ10/CES1/ABCC11/IL1B/SLC38A5/EMB/SLC22A3/TNF/SLC6A7/RBP5/TNFSF11/ABCD2/SLC38A1/RBP1/PLA2G2D/PLA2G2A/NOS

3/LRRC4C/MAP1A/CACNG8/GRIA4/CPEB3/CCR2/RIMS4/NTNG2/CDH11/HRH1/PLG/GHRL/STX1B/SCT/OTOF/AKAP5/ZDHHC2
3/LRRC4C/MAP1A/CACNG8/GRIA4/CPEB3/CCR2/RIMS4/NTNG2/CDH11/HRH1/PLG/GHRL/STX1B/SCT/OTOF/AKAP5/ZDHHC2

RIMS4/KCNJ5/ATP1A4/PKP2/GHRL/STX1B/TREM2/KCNA5/ABAT/KCNK12/SLC4A8/PTK2B/KCNK15/CACNA1C/UCP2/SCN1B/CA

OQ
IA1C/SCN1B/PGAM2/TACR1/TNNI2/FXYD1/SLC8A1/TNNT2/TIFAB/CXCR4/CD38/TNF/PIK3CG/RCSD1

CCR2/GDF2/CASQ1/SLC9A9/ATP1A4/TGM2/TUNAR/CDH23/SLC9C1/TTC7A/ATP2B3/STC2/SLC4A8/PTK2B/CACNA1C/CCL8/WN

P25/GPR174/RASGRF1/ARHGAP15/MYOC/RHOH/P2RY10

VIPR1/ADCY4/ADGRG4/GPR65/INSL3/ADGRG2/GPR171/ADGRL3/GNA14/PDE2A/S1PR4/S1PR1/MARCO/CXCL9/CXCL10/ADGR

GFB1/SULF2/CLDN18/TIE1/SFRP1/ALOX5/WNT5A/AREG/GATA2/DLL4/WNT7A/TACR1/JAML/MMP12/SGPP2/TGFA/KRT2/EAF2/

)3/SYT1/KMO/SV2A/SELENOM/WNT7A/LYN/TACR1/LILRB1/VSNL1/SUCNR1/LRRK2/IL1B/CHRNA6/CD38/IL1RN/INHBA/TNF/DO[illegible]

O1/ADIPOQ

RO1A/PTPN6/C3/RELN/SLC38A1/CXCL9/MMP9/CXCL10/HAMP/CEMIP/ANO9/CXCL11/P2RX5/MS4A1/CD19/ADIPOQ

TA2/JAK2/FOXC2/ITGA5/INSL3/CD40/PEAR1/SFTA3/WNT7A/LYN/TFPI/JAML/MMP12/F5/SELP/PRKCQ/DGKA/CD40LG/SCUBE1/

ᐳR4/LCK/TNF/PLEK/LAPTM5/ITK/SELE/TNFSF10/TBC1D1/RASGRP2/ARHGAP25/PTPRC/CCL5/RASGRF1/TBC1D10C/ATP2A3/C

AK2/CD40/BTC/PEAR1/LYN/TFPI/F5/SELP/PRKCQ/DGKA/CD40LG/TIFAB/SCUBE1/LCK/PLCG2/PLEK/PIK3CG/MMRN1/VAV1/PTF

A/PTPN6/PTPRC/CXCL9/CCR7/CCL5/ATP2A3/XCR1/CACNA2D2/CXCL10/CEMIP/CXCL11/FCRL3/P2RX5/MS4A1/CD19

'HY1/ATP4A/LYN/KCNJ10/FGF11/LILRB1/TLR9/FXYD1/SLC8A1/TSPAN13/CD84/NETO1/CXCR4/FXYD3/PIK3CG/ANK3/CORO1A/F

RX1/PTGS2/SCN3A/DSC2/POPDC2/ADRA2B/ADM/EDN1/ADORA2A/ITGA4/P2RY2/HTR1D/ADRA1A/KEL/DES/SOD2/SLIT2/C3AR[illegible]

ITGB2/CD84/PCDH15/IL1RN/CDHR1/MPZL2/EMB/CD6/SELE/ITGAL/CLSTN3/SELL/CLDN5/ADIPOQ

AK2/TGFB3/CD40/PIK3R6/ICAM1/NDRG4/HAVCR2/CCL21/WNT7A/SPI1/IL34/IGFBP6/TLR9/BMP2/CCL19/LRRK2/IL1B/TGFA/ACKR

1/FGF2/HGF/KLRK1/TNF/GPR18/SPN/GATA3/SIGLEC10/FGR/MMRN1/CD96/PTPN6/PTPRC/CR1/NLRC3/IL33/HAMP/SEMA3A/S

SUCNR1/PRKCQ/ITGB2/LSP1/CCL19/IL1B/SLAMF8/BIN2/ACKR3/IL12A/CXCR4/PLA2G7/CCR5/DOCK2/SEMA7A/SEMA4D/CXCL1[illegible]

SUCNR1/PRKCQ/ITGB2/LSP1/CCL19/IL1B/SLAMF8/BIN2/ACKR3/IL12A/CXCR4/PLA2G7/CCR5/DOCK2/SEMA7A/SEMA4D/CXCL1[illegible]

3AP1/RASGRP1/PTPN22/CLNK/LTF/ZBP1/KLRD1/PLCG2/KLRK1/CD180/TNF/TRAF3IP3/SPN/BTK/FGR/SCIMP/CD96/IL4I1/LILRA4

/ANK3/MMRN1/CORO1A/SELE/IL4I1/CD27/VAV1/TESPA1/PTPN6/RASAL3/FCHO1/PTPRC/CCR7/CCL5/SASH3/ALOX15/IL23A/IL2

R6/BIRC3/KLRB1/TLR4/SH2D1A/CCL5/OASL/SLAMF6/CR1/NLRC3/TLR7/AIM2/FCRL3/TLR10

3J1/OR5H2/OR2F2/RTP4/SCN11A/OR52A1/OR4C5/OR11L1/OR6B2/OR4A47/OR13C4/RHO/OR9Q2/CACNA2D4/KCNA1/OR6N2/O

.23A/IL2RG/CR1/CARD11/IL7R/IL18/RHOH/FCRL3/LIF/IKZF3/DCSTAMP/ADIPOQ

N/IL23A/PLA2G2A/IL7R/IL18/IRF8/HAMP/NOS2/PLAC8/IGHM/CCL20/CXCL13

NK/LTF/ZBP1/KLRD1/PLCG2/KLRK1/TNF/TRAF3IP3/TNFSF11/BTK/PIK3CG/SCIMP/LILRA4/VAV1/TLR6/BIRC3/C3/TLR4/CCR7/SH

CR5/CXCL11/CNR2/CCR6/CCL20/STAP1/CXCL5/CXCL13

CD84/SERPINB9/SLAMF8/CR1L/TIGIT/SLA2/UBASH3A/PTPN22/TBX21/CLNK/LTF/JAK3/KLRD1/INHBA/NCKAP1L/KLRK1/TNF/RAS

OR5T3/OR8D1/OR51E2/OR4A15/OR14J1/OR51A2/OR3A3/LPO/OR6C74/OR8J1/OR5H2/OR2F2/RTP4/SCNN1G/OR52A1/OR4C5/O

IL12A/ADAM19/RASGRP1/PTPN22/TBX21/JAK3/IL1RN/LCK/CD5/NCKAP1L/KLRK1/CD6/TNF/LAPTM5/TNFSF11/CD3E/SPN/GATA

2/OR52A1/OR4C5/OR11L1/OR6B2/OR4A47/OR13C4/OR9Q2/OR6N2/OR9G4/OR6X1/OR5C1/OR2L5/OR2B2/OR2T1/OR4K17/OR1E

R13C4/OR9Q2/OR6N2/OR9G4/P2RX2/OR6X1/OR5C1/OR2L5/OR2B2/OR2T1/OR4K17/OR1E1/OR6K2/OR8U1/OR6B1/OR4D2/TAS2

GALS2/LRRK2/IL2RA/FOXP3/IL1B/SLAMF8/NLRP1/PIK3AP1/PLA2G7/PDE2A/DNASE1L3/SEMA7A/AOAH/ZBP1/HGF/PLCG2/TNF/T

12/EBI3/LILRB1/CRTAM/TLR9/CD28/PRKCQ/CCDC88B/IL1RL1/IFIH1/CCL19/LRRK2/FOXP3/IL1B/IRF1/IL12RB1/CD84/CD40LG/NL

2/OR2F2/RTP4/OR52A1/OR4C5/OR11L1/OR6B2/OR4A47/OR13C4/OR9Q2/OR6N2/OR9G4/OR6X1/OR5C1/OR2L5/OR2B2/OR2T1/

4/OR9Q2/OR6N2/OR9G4/OR6X1/OR5C1/OR2L5/OR2B2/OR2T1/OR4K17/OR1E1/OR6K2/OR8U1/OR6B1/OR4D2/OR1L6/OR13C8/

L13

F17/JAK2/IL15/IRF7/CXCL2/IL10RA/CCL21/PADI2/LYN/SPI1/LILRB4/MMP12/IRAK2/IL17REL/EBI3/LILRB1/IL34/ACKR2/CCR4/TRAF1

L18/RHOH/TNFRSF13C/IDO1

PRC/CCR7/CCL5/SASH3/IL23A/PLA2G2A/IL2RG/CR1/CD24/CARD11/IL7R/SELL/IL18/RHOH/TNFRSF13C/IDO1

TRAF3IP3/CD247/LAPTM5/CD3E/GATA3/BTK/CD3D/FGR/SCIMP/LAT2/ITK/IGHA2/LILRA4/VAV1/THEMIS/TESPA1/PTPN6/TLR6/TI

SCIMP/LAT2/ITK/IGHA2/LILRA4/VAV1/THEMIS/TESPA1/PTPN6/TLR6/TRBC1/BIRC3/TRBC2/C3/FCHO1/PTPRC/TLR4/CCR7/SH2I

ESPA1/PTPN6/RASAL3/FCHO1/PLA2G2D/PTPRC/CCR7/CCL5/SASH3/IL23A/PLA2G2A/IL2RG/CR1/CD24/CARD11/TNFRSF9/IL7R

RD11/IL33/IL7R/IL18/HAMP/RHOH/TNFRSF13C/FCRL3/STAP1

SASH3/IL23A/IL2RG/CR1/CD24/CARD11/IL33/IL7R/IL18/HAMP/RHOH/TNFRSF13C/FCRL3/STAP1

G7/IL18/IRF8/CCR6/CD19

33/NKG7/IL18/IRF8/CCR6/CD19

D1/CD5/INHBA/NCKAP1L/KLRK1/TOX/CD2/CD6/RASSF5/LAPTM5/TNFSF11/CD3E/SPN/GATA3/BTK/FGR/RAC2/CORO1A/IL4I1/(

.23A/PLA2G2A/CR1/CD24/CARD11/TNFRSF9/IL33/IL7R/IL18/CLU/TNFRSF13C/FCRL3/CSF2RB/IKZF3/CD22/TNFRSF13B/CD79A

'RSF13C/FCRL3/IKZF3/CD22/TNFRSF13B/CD79A/CR2/MS4A1/CD19/IDO1

/IKZF3/CD22/TNFRSF13B/CD79A/CR2/MS4A1/CD19/IDO1

K1/TNF/LAPTM5/SPN/GATA3/BTK/FGR/SCIMP/CD96/RAC2/IL4I1/VAV1/PTPN6/KLRB1/WAS/C3/IL18R1/RBP4/PTPRC/TLR4/SH2I

OCS1/SPINK5/IL1B/IRF1/IL12RB1/CD1D/CD40LG/SLAMF8/DOCK2/RASGRP1/PTPN22/TBX21/JAK3/LYL1/LCK/INHBA/PLCG2/NCI

A/PLCG2/NCKAP1L/TOX/CD2/GPR18/CD3E/SPN/GATA3/BTK/CD3D/MFNG/LY9/ITK/IL4I1/CD27/VAV1/THEMIS/TESPA1/PTPN6/IT

/IGHD/BANK1

/CARD11/CD3G/IGLC7/IGHM/FCRL3/FCER2/CD22/STAP1/CD79A/BLK/CR2/MS4A1/CD19/IGHD/BANK1

.RK1/FCMR/CD2/TNF/SPN/GATA3/BTK/PIK3CG/GZMB/FGR/SCIMP/CD96/RAC2/LAT2/CORO1A/IL4I1/CD27/IGHA2/VAV1/PTPN6

.R4/IL2RB/INPP5D/SASH3/IL23A/SLAMF6/CR1/IL33/IL7R/IL18/IL21R/CLU/CFI/IGLC7/IGHM/CCR6/CSF2RB/FCER2/CXCL13/CR2/(

/CLEC6A/LYN/IL18BP/RSAD2/LTA/LILRB4/EBI3/LILRB1/BMX/TRDC/CD7/CD28/PRKCQ/GAPT/ZAP70/CD8A/C1R/IL1RL1/CCL19/TI

IL2RB/INPP5D/SASH3/IL23A/SLAMF6/CR1/IL7R/NKG7/IL18/IL21R/CLU/CFI/IGLC7/IGHM/CCR6/CSF2RB/FCER2/CR2/CD19/IGHD

UTP18/WDR18/RPUSD2/NAT10/MTREX/TRMT61B/TSR1/FBL/NOL9/IMP4/ZNF622/POP7/LYAR/FTSJ3/AATF/NOC4L/SURF6/MT[illegible]

EXOSC4/RRP36/TSR3/URB2/DDX27/PRPF3/CLNS1A/AGO1/NSA2/RRP1/SUV39H1/NOL10/ABT1/DDX10/TSR2/REXO4/SF3A3/SF[illegible]

RCB1/NVL/WDR75/RIOK3/ANG/WDR55/ERCC2/DDX52/RRP8/GTF3C4/KRR1/NOP14/ERI1/BUD23/PIH1D1/TAF1B/UTP20/TBL3/[illegible]

METTL1/AGO1/NSA2/RRP1/SUV39H1/NOL10/ABT1/DDX10/PUM1/TRUB2/TSR2/REXO4/RNF113A/INTS13/ELP6/DUS1L/DDX18/E[illegible]

10A/TPRKB/C2orf49/QRSL1

NCAPH2/PSMG2/NSL1/BECN1/BRD7/PPP2R1A/ARID2/NUMA1/ABRAXAS1/KLHL22/TUBG2/CHMP6/CHMP1A/NUP62/RHOA/RM[illegible]

1H1/CHMP2A/ANAPC1/NDEL1/RAD21/CHMP3/MIS18A/SEH1L/NCAPD2/SKA1/PUM1/KIF22/SAC3D1/SIRT2/RMDN1/TACC3/UH[illegible]

[illegible]/INCENP/ZW10/SMC2/DAPK3/MAPRE1/DYNC1LI1/HASPIN/DIS3L2/KIF11/NDE1/GEN1/FLNA/CUL9/KNTC1/CHEK1/BORA/NFIB/[illegible]

F11/RBM15B/EIF1/SNIP1/RRP1B/LSM10/SNRNP27/THOC2/SNRNP48/CCAR2/FUS/RBFOX2/MAGOH/PPP4R2/RBMX2/MTREX/T

31/CHEK2/CLASP1/TTK/PIBF1/MYBL2/CCNE2/CCSAP/INCENP/ZW10/SMC2/CENPQ/BAG6/MAPRE1/FMN2/DYNC1LI1/HASPIN/E [illegible]S/RBMX/U2AF2/CACTIN/PRPF19/DDX42/AQR/UBL5/SON/DDX41/C9orf78/SYF2/RBM23/NCBP2/PAXBP1/PPIL3/HNRNPM/RBM3[illegible]

[illegible]F2/CACTIN/PRPF19/DDX42/AQR/UBL5/SON/DDX41/C9orf78/SYF2/RBM23/NCBP2/PAXBP1/PPIL3/HNRNPM/RBM39/DYRK1A/SR[illegible]

[illegible]F2/CACTIN/PRPF19/DDX42/AQR/UBL5/SON/DDX41/C9orf78/SYF2/RBM23/NCBP2/PAXBP1/PPIL3/HNRNPM/RBM39/DYRK1A/SR[illegible]

[illegible]AC/RBM15B/EIF1/SNIP1/RRP1B/LSM10/CPSF3/SNRNP27/LSM11/THOC2/SNRNP48/PTCD2/CCAR2/RBFOX2/MAGOH/PPP4R2/[illegible]

[illegible]N1/DDX11/BRD7/ARID2/NUMA1/KLHL22/MPHOSPH8

[illegible]CHEK1/POLI/RFC5/NFIB/NFIC/MCRS1/REPIN1/DBF4/DONSON/SSRP1/GINS2/DNA2/PRIM2/DNAJC2

[illegible]/RAD51C/SGO1/CHMP2A/FIGNL1/NDEL1/CDC14B/RAD21/CHMP3/NSFL1C/SEH1L/NCAPD2/SKA1/KIF22/SIRT2/RMDN1/SPIRE[illegible]

1/BRCA2/PARP1/OOEP/DMAP1/CHEK1/ATRIP/NSMCE2/KAT5/MTA1/PIAS4/RNF8/BRD8/GINS2/DNA2/NUDT16L1/RUVBL1/KLHL

DUFA6/PET117/RCC1L/NOL3/NDUFS8/PANK2/NDUFB11/CSNK2A2/DMAC1/NDUFB2/HSPA4/CDC37/ATG10/APOOL/POLDIP2/FU

LBP1/ANAPC11/CDK1/NCAPG2/RANGRF/DCTN2/NCAPH/PRDM7/NUDC/POLDIP2/CHMP1B/ZWINT/RAD51C/SGO1/CHMP2A/F

2/UQCRC2/UQCC3/ATP5F1D/NDUFA10/UQCRQ

C1/TOP2B/BRCC3/KMT5A/OGG1/TAF10/TAF9

5/DNA2/TAOK3/MBTPS2/RB1/RAD1/TPRA1/BABAM2/PRPF19/EZH2/GNB1L/PSMG2/SYF2/TTI2/INIP/CCP110/BRD7/RBL1/PKD2/A

AXAS1/KLHL22/GIGYF2/PTEN/FBXO7/MAPK14/RFWD3/TAOK1/CCND1/ZWILCH/FZR1

3/LMAN1/LMAN2L/COG4/TEX261/GABARAPL2/EXOC2/PRKD1/KIF1C/COG1/AP1AR/RNF139/SEC23A/YIPF5/CLN3/SEC24D/RAB12

SARNP/LSM14B/PUM1/TRUB2/GFM2/CASC3/CNOT9/TCOF1/DAP/ELP6/GFM1/CAPRIN1/PABPC4/UPF3A/EIF4E2/EEF2K/TRIM7

NMT1/ANTKMT/MPHOSPH8/MEPCE/WDR6/TRMT2A/COMTD1/METTL22/TRMT13/CARNMT1/EHMT1/SPOCD1/TRMT1/CBLL

SP2/TPR/CUL4A/TRIAP1/AVEN/CRADD/CHEK2/CLASP1/TTK/LSM10/CPSF3/CCNE2/RDX/LSM11/INCENP/UBE2A/ZW10/FBXO3[illegible]

/HSPB1/ABCF1/PLD1/SARNP/LSM14B/HAGH/PUM1/TRUB2/GGT1/GFM2/CASC3/CNOT9/TCOF1/DAP/ELP6/GFM1/GSS/CAPRIN

R11/NDUFA11/CDK1/PFKM/NDUFA6/CHCHD5/G6PD/NDUFS8/PANK2/NDUFB11/AGL/ETFB/NDUFB2/FH/PRXL2C/P4HA2/CIAP

/GEN1/ATAD5/ANKRD17/KNTC1/CHEK1/MDM2/IER3/ERCC2/CUL4B/NSMCE2/BRD4/NAE1/DONSON/RRM2/ANAPC15/TAOK3/[illegible]

4/DNAJB2/FBXW11/RNF10/HSPBP1/SUMO2/FBXO4/NAGLU/GLMN/ECRG4/PSMA1/FBXO44/PSMD13/DDB1/DCAF12/CUL1/FBX

2/MLF1/TPR/CUL4A/TRIAP1/AVEN/CRADD/CHEK2/TTK/MDC1/LSM10/CPSF3/CCNE2/RDX/LSM11/INCENP/CCAR2/ZW10/FBX

N6/DET1/CTNNB1/GID8/UGGT1/FBXO2/CUL4A/ZYG11B/PSMF1/AXIN1/PSMA2/DMAC2/RNF145/FBXW8/HSP90B1/NPLOC4/WW[illegible]

[illegible]IM3/CDKN2C/DUSP3/SIRT2/ORC1/RRM1/INTS13/RPA3/BLM/SASS6/MKI67/PHB2/DLG1/PIN1/MRE11/RIOK2/EGF/LCMT1/SMAR

[illegible]F1/ANAPC5/RNF170/FBXL17/SHARPIN/ARIH2/RNF114/TRIP12/TRIM62/SHPRH/RNF168/CUL3/TRIM3/FBXO22/RNF19B/MYCB[illegible]

[illegible]/VTA1/VPS16/GABARAPL2/UBQLN2/PHF23/ATP2A2/SNX18/PRKACA/RAB5A/WDR81/BECN1/ATG16L1/ARL8B/SNX5/VPS28/USP[illegible]

XD1/FAM91A1/DLG1/DYNLL1/ATP13A2/CDT1/RIOK2/CLASP2/SNF8/COPG2/CTNNB1/ADORA2B/PDCD6/RHOT2/COPG1/MECP[illegible]

10/FBXO31/RPA2/DYNC1LI1/TELO2/CDK5RAP1/BABAM1/HASPIN/FBXO4/MCIDAS/GEN1/MUS81/BRCA2/TP53BP2/KNTC1/CHEK

O1C/BCAP31/TOMM34

FM2/WDR76/ZNF365/NPAS2/ABL1/BRCA1/KMT5C/CUL4A/PHF13/TAF6/TRIAP1/STK25/RIF1/MEAK7/USP51/SGTA/NUPR1/CCAR

5/ROCK2/DBF4/BRD4/RRM2/STXBP4/NCAPD3

P3/CNOT3/UPF1/NCBP1/JMJD4/MTG2/APP/IGF2BP2/ANG/UHMK1/IMPACT/ATF4/KHDRBS1/UPF3B

3/ELP6/SLC11A1/EXOSC2/CAPRIN1/PABPC4/UPF3A/PATL1/EIF4E2/EEF2K/TRIM71/CNOT11/CYFIP1/ZC3H10/BZW1/RBM8A/ELA

HMP4B/LAMP1/MCOLN1/ATG14/EMC6/HEXA/TRAF6/TPP1

KAP5L/UBQLN1/DUSP3/BNIP3L/GFM2/GABARAP/AIFM1/SPTBN1/SPTBN5/CARMIL1/PHB2/WIPI2/CAMSAP2/KDM4A/EIF4E2/ATF

1/LONP2/BCS1L/TRAM1/NUP35/SGTA/SRP9/EFCAB7/HSP90AA1/LAMP2/MON1A/TIMM9/TIMM29/BAG6/STAM/MVB12A/TIMM17

CY6

10/HYLS1/FLNA/YAP1/BBS4/DNAL1/KIAA0753/KIFAP3/ARF4/TRAF3IP1/KIF24/SNAP29/PCM1/ABCC4/NME5/ENKD1/PLK4/MPHOS

S4/ADCY6/PYGL

3/ARF4/TRAF3IP1/KIF24/SNAP29/PCM1/ABCC4/NME5/ENKD1/PLK4/MPHOSPH9/PCNT/CEP290/CEP350/CFAP69

DR1/ADPRHL1/MAPRE1/PRKAR1A/CENPH/HAUS3/CIRBP/CENPW/PAN3/MCIDAS/KIF11/HDAC3/DDB1/CDS2/FLNA/KNTC1/PF

AP1/CHEK1/ERCC2/NSMCE2/MCRS1/KAT5/PIAS4/RNF8/BRD8

HDF1/LARP1B/GIGYF2/CPEB1/MAPK14/EIF4ENIF1/SRSF2/FXR2/ZC3H14/METTL3/TENT4A/RBM15/FTO/TTC5/NCL/NBAS/SNW

UFV1/NDUFB3/FIGNL1/ELOVL1/DNPH1/PRKAG1/GUK1/STOML2/NDUFS4/NNT/ADCY6/SLC4A7

5E/NAT10/SECISBP2L/CDK5RAP1/SESN2/STK35/AJUBA/CIRBP/POLDIP3/CNOT3/UPF1/NCBP1/JMJD4/SCARB2/MTG2/APP/IGF

CTNNB1/SLC15A4/KMT5C/CUL4A/S100A11/PHF13/TAF6/KPNA2/MECP2/SMC6/PMS2/RIF1/PARN/XRCC5/USP51/DFFB/FUS/ERC

/0C/LAMTOR1/AP3B1/TRAPPC8/ATP6V0A4/TRAPPC12/KLHL12/AP1M1/ATP6V1F/SEC23B/SNAP29/YIPF7/ARFGAP3/TRAPPC13/U

;2

;2

/DYSF/SLC4A8/PTK2B/WNT5A/SV2B/JAK2/ACHE/SYT1/KMO/WNT7A/KCNJ10/PLCL2/TACR1/CALB2/LRRK2/IL1B/NETO1/CHRNA

/DYSF/SLC4A8/PTK2B/WNT5A/SV2B/JAK2/ACHE/SYT1/KMO/WNT7A/KCNJ10/PLCL2/TACR1/CALB2/LRRK2/IL1B/NETO1/CHRNA

CNA1I/WNT7A/KCNJ10/TACR1/FXYD1/SLC8A1/LRRK2/NETO1/HCN1/GNA14/CHRNA6/TNF/GABRB3/ANK3/RELN/MYOC/CNR2/

IT5A/SV2B/FASLG/CD40/THY1/CCL21/SV2A/ATP4A/LYN/CALB2/SLC9A7/SLC8A1/CCL19/LRRK2/SLAMF8/CCR5/LCK/FGF2/PLCG

G5/ADGRF5/CXCL11/CNR2

IL12A/JCAD/PPP1R16B/FGF2/HGF/TNF/TNFSF11/GATA3/BTK/FLT4/BCL11B/CCL5/RASGRF1/NLRC3/COL4A3/FGF1/TBX1

C2B/TNFSF11/GATA3/BTK/MCTP2/RBP4/TMEM132A/CCL5/SNCAIP/NOS2/LIF/BLK/ADIPOQ

CXCR4/LCK/FGF2/PLCG2/TNF/PLEK/PIK3CG/MMRN1/VAV1/PTPN6/WAS/TLR4/ALOX15/CARMIL2/FGF1/HBB

R1/COL4A3/ST18/AIM2/FCRL3/CCL20/CXCL13

ᴾN6/WAS/TLR4/HBB

ᴾTPN6/RELN/CXCL9/CCL5/MMP9/CXCL10/HAMP/CEMIP/ANO9/CXCL11/FCRL3/P2RX5/MS4A1/CD19

1/CDH5/OLR1/SCN4B/HTR7/FGA/THRB/FOXN4/SVEP1/HRH1/MYL2/KCNJ5/TNNI1/VEGFC/PTAFR/DLL1/PKP2/ADM2/GHRL/STA

R3/DOK3/JCAD/RASGRP1/SEMA7A/PTPN22/INHBA/FGF2/HGF/PLCG2/TNF/LAPTM5/TNFSF11/MAP4K1/PIK3CG/SCIMP/ALKAL2/

TAP1/CXCL13/IL22RA2/ADIPOQ

4/FGF2/HGF/NCKAP1L/KLRK1/ACKR4/GPR18/S1PR1/TNFSF11/SPN/PIK3CG/RAC2/CORO1A/VAV1/CXCL9/CCR7/CCL5/CXCR3/

4/FGF2/HGF/NCKAP1L/KLRK1/ACKR4/GPR18/S1PR1/TNFSF11/SPN/PIK3CG/RAC2/CORO1A/VAV1/CXCL9/CCR7/CCL5/CXCR3/

4/VAV1/PTPN6/TLR6/BIRC3/KLRB1/TLR4/SH2D1A/CCL5/IL23A/OASL/SLAMF6/CR1/NLRC3/TLR7/AIM2/FCRL3/TLR10

RG/CR1/CD24/CARD11/IL7R/MYOC/IL18/RHOH/TNFRSF13C/LIF/CXCL13

R9G4/LXN/OR6X1/HPN/OR5C1/GJA10/OR2L5/OR2B2/OR2T1/OR4K17/OR1E1/OR6K2/OR8U1/OR6B1/OR4D2/TAS2R38/OR1L6/O[illegible]

[illegible]2D1A/CCL5/IL23A/OASL/PLA2G2A/SLAMF6/NLRC3/IL33/TLR7/NKG7/IL18/AIM2/FCRL3/TLR10/STAP1/IDO1

[illegible]SF5/GPR18/LAPTM5/SPN/BTK/FGR/CD96/IL4I1/LILRA4/PTPN6/TLR6/PLA2G2D/PTPRC/TLR4/SH2D1A/INPP5D/ALOX15/TBC1D1[illegible]

R11L1/OR6B2/CALHM3/OR4A47/OR13C4/OR9Q2/OR6N2/OR9G4/P2RX2/OR6X1/OR5C1/OR2L5/OR2B2/OR2T1/OR4K17/OR1E1/[illegible]

3/ANK3/MMRN1/CORO1A/SELE/IL4I1/CD27/VAV1/TESPA1/PTPN6/RASAL3/FCHO1/PLA2G2D/PTPRC/CCR7/CCL5/SASH3/IL23A/

1/OR6K2/OR8U1/OR6B1/OR4D2/OR1L6/OR13C8/OR7G2/OR2S2/OR56A1/OR4K14/OR5D16/OR5D13/OR52B2/OR5B2/OR52L1/OR

2R38/OR1L6/OR13C8/OR7G2/OR2S2/OR56A1/KCNMB4/OR4K14/OR5D16/OR5D13/OR52B2/OR5B2/OR52L1/OR51D1/OR4D5/OR4

NFSF11/SPN/GATA3/BTK/PIK3CG/SIGLEC10/FGR/SELE/PTPN6/BIRC3/C3/PLA2G2D/PTPRC/TLR4/CCR7/CCL5/ALOX15/IL23A/P

RP1/TIGIT/IL12A/RASGRP1/SEMA7A/PTPN22/TBX21/CLNK/HGF/PLCG2/KLRK1/CD2/CD6/TNF/TRAF3IP3/LAPTM5/CD3E/SPN/G

OR4K17/OR1E1/OR6K2/OR8U1/OR6B1/OR4D2/TAS2R38/OR1L6/OR13C8/OR7G2/OR2S2/OR56A1/OR4K14/OR5D16/OR5D13/OR

R7G2/OR2S2/OR56A1/OR4K14/OR5D16/OR5D13/OR52B2/OR5B2/OR52L1/OR51D1/OR4D5/OR4E2/OR1D5/OR2AT4/OR52W1/O

/IL1RL1/IFIH1/CCL19/CD300LF/IL2RA/SOCS1/IL1B/IRF1/IL9R/IL12RB1/ACKR3/IL12A/CXCR4/CCR5/JAK3/ZBP1/IL1RN/HCLS1/INHI

RBC1/BIRC3/TRBC2/C3/FCHO1/PTPRC/TLR4/CCR7/SH2D1A/INPP5D/BLNK/OASL/CR1/CD24/NLRC3/CARD11/CD3G/TLR7/AIM

D1A/INPP5D/BLNK/OASL/CR1/NLRC3/CARD11/CD3G/TLR7/AIM2/IGLC7/IGHM/FCRL3/TLR10/CD22/STAP1/CD79A/BLK/CR2/MS4
/IL18/RHOH/TNFRSF13C/IDO1

CD27/SIT1/VAV1/TESPA1/PTPN6/RASAL3/IL15RA/FCHO1/PLA2G2D/PTPRC/TLR4/CCR7/CCL5/INPP5D/SASH3/IL23A/TBC1D10C/

/CR2/MS4A1/CD19/IDO1

D1A/SASH3/IL23A/SLAMF6/CR1/IL33/IL7R/TLR7/IL18/NOS2/FCRL3/FCER2/CD22/STAP1/CR2

KAP1L/TOX/CD2/GPR18/CD3E/SPN/GATA3/BTK/CD3D/MFNG/LY9/ITK/IL4I1/CD27/VAV1/THEMIS/TESPA1/PTPN6/ITM2A/IKZF1/IL

TM2A/IKZF1/IL18R1/IL15RA/POU2AF1/PLA2G2D/PTPRC/CCR7/BCL11B/INPP5D/BLNK/SASH3/IL23A/SLAMF6/IL2RG/CR1/CARD1

/KLRB1/WAS/C3/IL18R1/PTPRC/TLR4/SH2D1A/IL2RB/INPP5D/SASH3/IL23A/SLAMF6/CR1/IL7R/NKG7/IL18/IL21R/CLU/NOS2/CFI/

CD19/IGHD

RGC2/FOXP3/IL1B/IRF1/LAX1/IL9R/PRF1/IL12RB1/UNC13D/CD84/CD1D/CD40LG/CR1L/IL12A/SLA2/MYO1G/TBX21/JAK3/TRAC/K

G2/NMD3/NVL/WDR75/RIOK3/WDR55/ERCC2/CUL4B/DDX52/TMA16/RRP8/KRR1/NOP14/ERI1/BUD23/NIP7/PIH1D1/UTP20/TBL3

3B6/DDX18/EXOSC2/GTF2H5/DDX39B/SNRNP200/DDX31/EXOSC6/GAR1/RIOK2/DDX17/PNO1/PUF60/WDR43/MCTS1/CRNKL1/I

POLR1B/NGDN/TRIR/METTL18/NOB1/DDX11/CHD7/EXOSC7/RBM34/PELO/CAVIN1/GTPBP4/NOLC1/SPIN1/WDR36/RPUSD4

XOSC2/GTF2H5/EXOSC6/GAR1/TRIM71/RIOK2/DDX17/TRIT1/WDR43/PWP1/OSGEPL1/SRSF3/ELAC2/ZC3H10/WDR74/NOL11/W

2/ZWILCH

RF1/SASS6/MKI67/RNF4/BAZ1B/TOP3A/MRE11/CDT1/RIOK2/GOLGA2/LCMT1/CLASP2/SMARCD2/MND1/BRCA1/CSNK2A1/STAI

NSMCE2/BROX/KAT5

FIP11/CELF2/CLK2/DHX35/VIRMA/PRPF40A/STRAP/TMBIM6/TAF6L/ESS2/CIRBP/RBM41/MAGOHB/TRA2A/UPF1/NCBP1/LSM1/

CL7B/C14orf39/DIS3L2/KIF11/SMARCB1/DDB1/GEN1

9/DYRK1A/SRSF1/LUC7L2/DCPS/LUC7L/HNRNPUL1/SFPQ/SRSF2/TSSC4/YJU2/GEMIN2/WBP4/CTNNBL1/DDX39A/SF3A2/MET

SF1/LUC7L2/DCPS/LUC7L/HNRNPUL1/SFPQ/SRSF2/TSSC4/YJU2/GEMIN2/WBP4/CTNNBL1/DDX39A/SF3A2/METTL3/SNRPD3/

SF1/LUC7L2/DCPS/LUC7L/HNRNPUL1/SFPQ/SRSF2/TSSC4/YJU2/GEMIN2/WBP4/CTNNBL1/DDX39A/SF3A2/METTL3/SNRPD3/

RPUSD2/CDK11B/RBMX2/MTREX/TFIP11/CELF2/CSTF1/DHX35/VIRMA/PRPF40A/SLBP/STRAP/ESS2/CIRBP/RBM41/MAGOHB/

I/UHRF1/MKI67/PHB2/BAZ1B/FANCA/PIN1/MRE11/CDT1/SLX4/EGF/GOLGA2/LCMT1/CLASP2/MTBP/FBXO43/MND1/NME6/ORC

15/BABAM2/RAD51AP1/PRPF19/MCMDC2/SMARCAL1/XRCC1/NSD2/INIP/DDX11/BRD7/ARID2/ZCWPW1/ABRAXAS1/EYA3/IFF[illegible]

[illegible]NDC2/FUNDC1/NDUFAF6/MTCH2/ALKBH7/RTL10/TWNK/TOMM6/TTC19/SNX30/PDCD5/NDUFB3/RALA/TIMM21/TMEM223/T[illegible]

[illegible]IGNL1/NDEL1/CDC14B/RALA/RAD21/CHMP3/NSFL1C/SEH1L/NCAPD2/SKA1/KIF22/SIRT2/RMDN1/SPIRE1/UHRF1/MKI67/PHB[illegible]

ABRAXAS1/KLHL22/GIGYF2/PTEN/FBXO7

2/GCC2/CSNK1D/VAMP7/VPS54/COPB2/PPP6C/COG5/TFG/NBAS/HTT/ARCN1/PREPL/COMMD1/PITPNB/EXOC4/RAB14/MIA3/C

1/RARS2/CNOT11/IARS2/CYFIP1/MCTS1/BZW1/RBM8A/ELAVL1/YARS2/CSNK2A1/LARP4/DNAJC3/TPR/UBA52/C6orf52/YTHDF

1/FAM98B/MRM3/METTL3/SNRPD3/ATPSCKMT/PRDM15/PRDM5/BHMT/RBM15/TRMT10A/PRDM4/CAMKMT/SMARCA5

1/RPA2/DBF4B/DYNC1LI1/E2F3/CCNA2/CDC25A/BABAM1/HASPIN/BCL7B/APP/SMARCB1/DDB1/GEN1/CUL1/ATAD5/ANKRD17/

1/PABPC4/UPF3A/EIF4E2/EEF2K/TRIM71/RARS2/CNOT11/IARS2/CYFIP1/MCTS1/BZW1/RBM8A/ELAVL1/YARS2/CSNK2A1/LAR

IN1/PGK1/ACO2/NDUFV1/NDUFB3/IFNLR1/PHLDA2/COQ9/PRKAG1/STOML2/CISD1/NDUFS4/NNT/PYGL/FOXK2/SLC25A22/UQ

MBTPS2/RB1/TPRA1/BABAM2

L4/FBXO9/DAB2/PSMB4/MDM2/ARRB1/KAT5/KCTD13/PSMD11/MTA1/AMN1/WDR26/PSMD12/ANAPC15/FBXL3/UBE3A/PSMD6/

)31/RPA2/DBF4B/DYNC1LI1/TELO2/CDC25A/BABAM1/HASPIN/BCL7B/FBXO4/APP/SMARCB1/DDB1/GEN1/PAGR1/MUS81/ATAD

TR1/FEM1A/SGTA/UBE2A/TRIB1/NUPR1/CCAR2/PSMC4/AUP1/FBXO31/KLHL42/RNF122/PITHD1/DNAJC10/BAG6/GNA12/LTN1/

CD2/MTBP/FBXO43/ABL1/MBTPS1/NME6/BRCA1/ADAMTS1/CTNNB1/PTCH1/CASP2/TPR/CUL4A/TRIAP1/AVEN/CRADD/CHEK

2/TRAF6/RNF41/CDC27/BIRC2/UBE2K/KLHL20/TNFAIP3/RNF31/UBE4A/DCAF15/ARIH1/RNF26/UBE2D3/UBR5

30/TIGAR/TSC1/YOD1/FBXO7/RAB3GAP2/DNAJC16/CHMP6/WDR45B/CHMP1A/CLN3/TMEM74/MAPK3/SMURF1/HGS/RAB23/V

2/STK25/CLASP1/BICD1/SNAPIN/PIBF1/LLGL1/TRAPPC5/ARL6/MAP4/INCENP/RAB6A/ZW10/CENPQ/KIF13A/RHOT1/MAPRE1/FM

K1/MDM2/IER3/DOT1L/ATRIP/RRP8/NAE1/DONSON/INTS7/ANAPC15/DNA2/TAOK3/PTPRK/MBTPS2/RB1/RAD1/TPRA1/BABAM

R2/FUS/ERCC1/EEF1E1/PPP4R2/RPA2/BAG6/TFIP11/FMN2/BDKRB2/NACC2/TELO2/BABAM1/BCAP31/TMBIM6/PARPBP/TAF6L/

VL1/CSNK2A1/LARP4/DNAJC3/TPR/MECP2/C6orf52/YTHDF3/PARN/TSN/POLR2G/EIF1/SAMD4B/TSNAX/SRP9/FUS/MAGOH/D[illegible]

13A2/EEF2K/SH3GL3/GOLGA2/HMGA1/UFM1/CTSK/CLASP2/MCTS1/SMARCD2/MICAL1/FZD5/ULK3/SUPT16H/LIX1L/MMP11/K[illegible]

A/GNPTG/E2F3/NUP85/NUP153/TOMM5/BAG3/BCAP31/TSC2/TOMM34/TOMM70/SNUPN/NUP155/SCARB2/VPS25/SRP19/NUP1

SPH9/PCNT/CEP290

RRC2C/KIAA0753/ARHGEF10/TRAPPC12/HAUS4/CNOT6/NIP7/DOCK5/USP10/AAAS/TUBGCP4/CEP295NL/PLK4/CENPK/WASH(

1/SF1/CWC22/CDC73/HNRNPL/PCID2/FMR1/CSDE1/RBMXL1/PDE12/SRPK1/HNRNPD/PRKCD/SECISBP2/FXR1/TIA1/SRSF7/E

2BP2/ANG/UHMK1/IMPACT/ATF4/KHDRBS1/UPF3B

C1/PPP4R2/HSP90AA1/SSBP1/NAT10/RPA2/TFIP11/FMN2/DBF4B/CCNA2/CDC25A/BABAM1/PARPBP/TAF6L/BCL7B/UPF1/FBX

JNC13B/ATP6V1C1/VTA1/BLOC1S3/WASHC5

A6/CD38/TNF/SHISA8/LRRTM1/MCTP2/RELN/SLC38A1/CLSTN3/RASGRF1/SNCAIP/CACNA2D2/CNR2/ADIPOQ

A6/CD38/TNF/SHISA8/LRRTM1/MCTP2/RELN/SLC38A1/CLSTN3/RASGRF1/SNCAIP/CACNA2D2/CNR2/ADIPOQ

/P2RX5

2/PLCH2/CORO1A/PTPN6/PTPRC/CXCL9/CCR7/CCL5/ATP2A3/XCR1/TMC8/CXCL10/HAMP/HVCN1/CEMIP/CXCL11/CD19

\T1/GCH1/SPTBN4/FLI1/KCNA5/ATP2B3/TGFB1/KLF2/ABAT/XCL2/PTPRO/CACNA1C/SERPING1/PCSK5/CYSLTR1/HOPX/C2CD

/CD27/TLR6/MARCO/FLT4/PTPRC/TLR4/CCR7/CCL5/ALOX15/PLA2G2A/CD24/MYOC/SEMA3A/FGF1/FCRL3/CCL20/LIF/TBX1/BA

IL23A/XCR1/CXCL10/SEMA3A/FGF1/CXCR5/CXCL11/CNR2/CCR6/CCL20/STAP1/CXCL5/CXCL13

IL23A/XCR1/CXCL10/SEMA3A/FGF1/CXCR5/CXCL11/CNR2/CCR6/CCL20/STAP1/CXCL5/CXCL13

R13C8/OR7G2/OR2S2/OR56A1/OR4K14/OR5D16/OR5D13/OR52B2/OR5B2/OR52L1/OR51D1/OR4D5/OR4E2/OR1D5/OR2AT4/OR5

0C/PLA2G2A/CR1/NLRC3/IL33/IL7R/ADGRF5/CNR2/FCRL3/CD22/TNFRSF13B/STAP1/CR2/IDO1/ADIPOQ/BANK1

OR6K2/OR8U1/OR6B1/OR4D2/TAS2R38/OR1L6/OR13C8/OR7G2/OR2S2/OR56A1/OR4K14/OR5D16/OR5D13/OR52B2/OR5B2/OR5[illegible]

PLA2G2A/IL2RG/CR1/CD24/CARD11/IL7R/IL18/RHOH/TNFRSF13C/CXCL13/IDO1/ADIPOQ

51D1/OR4D5/OR4E2/OR1D5/OR2AT4/OR52W1/OR2C1/OR7G3/OR5A1/OR10H1/OR5K2/SLC6A3/OR5M9/OR2AG1/OR10S1/OR10[illegible]

4E2/OR1D5/OR2AT4/OR52W1/OR2C1/OR7G3/OR5A1/OR10H1/OR5K2/OR5M9/OR2AG1/OR10S1/OR10A6/OR6P1/OR4D11/OR4C[illegible]

LA2G2A/NLRC3/IL33/TLR7/MMP9/NKG7/IL18/HAMP/AIM2/STAP1/IL22RA2/IDO1/ADIPOQ

ATA3/BTK/PIK3CG/FGR/SCIMP/LY9/ITK/POU2F2/TLR6/C3/IL18R1/POU2AF1/FLT4/PTPRC/TLR4/CCR7/SASH3/IL23A/SLAMF6/CA

52B2/OR5B2/OR52L1/OR51D1/OR4D5/OR4E2/OR1D5/OR2AT4/OR52W1/OR2C1/OR7G3/OR5A1/OR10H1/OR5K2/OR5M9/OR2AG[illegible]

R2C1/OR7G3/OR5A1/OR10H1/OR5K2/OR5M9/OR2AG1/OR10S1/OR10A6/OR6P1/OR4D11/OR4C3/OR52B4/OR5AN1/OR6S1/OR51

3A/ACKR4/TNF/IL3RA/LAPTM5/TNFSF11/LILRA4/PTPN6/BIRC3/IL18R1/IL15RA/ACKR1/PTPRC/CXCL9/CCR7/IL2RB/CCL5/CXCR

2/IGLC7/IGHM/FCRL3/TLR10/FCER2/CD22/STAP1/CD79A/BLK/CR2/MS4A1/CD19/IGHD/BANK1

4A1/CD19/IGHD/BANK1

/PLA2G2A/IL2RG/CR1/CD24/CARD11/TNFRSF9/IL7R/IL18/RHOH/TNFRSF13C/FCRL3/IKZF3/CD22/TNFRSF13B/CD19/IDO1/BAN

18R1/IL15RA/POU2AF1/PLA2G2D/PTPRC/CCR7/BCL11B/INPP5D/BLNK/SASH3/IL23A/SLAMF6/IL2RG/CR1/CARD11/CD3G/TNFR

I/CD3G/TNFRSF9/IL7R/IL18/IRF8/RHOH/CCR6/FCRL3/IKZF3/CD79A/CR2/MS4A1/CD19/PAX5

IGLC7/IGHM/CCR6/CSF2RB/FCER2/STAP1/CR2/CD19/IGHD

LHL6/TRAT1/KLRD1/NCKAP1L/KLRK1/FCMR/CD6/TNF/CD247/CD3E/SPN/GATA3/BTK/LAMP3/PIK3CG/SIGLEC10/CD3D/LY9/LAT

3/NGDN/CINP/RBIS/METTL18/MDN1/NOB1/CHD7/NOA1/EXOSC7/RBM34/GTPBP4/EIF1AX/TSC1/NOL

PWP1/DHX30/WDR74/NOL11/WDR46/UTP23/TGS1/POLR2D/SNRPF/NOM1/RRP12/CUL4A/RPF2/TR

/DR46/UTP23/RRP12/RPF2/TRMT112/METTL2A/UTP25/MECP2/INTS14/NIFK/CTU2/PARN/TSN/SNIP1/RRP

RD9/TPR/PHF13/PTTG1/WRAP73/KATNB1/CHEK2/SMC6/CLASP1/TTK/TUBGCP5/PIBF1/HAUS1/MYBL2/CENPO/CC

/PUS1/SRSF12/RBM42/ALYREF/RTCB/PTBP1/SRSF9/SREK1IP1/TCERG1/GRSF1/KHDRBS1/SRSF10/UPF3B/HNRNP

TL3/SNRPD3/LSM3/HNRNPR/RBM15/NCL/HTATSF1/RBM4B/SNW1/LSM8/SF1/SNRPA/SF3B3/DHX38/SDE2/CWC22/HN

LSM3/HNRNPR/RBM15/NCL/HTATSF1/RBM4B/SNW1/LSM8/SF1/SNRPA/SF3B3/DHX38/SDE2/CWC22/HNRNPL/FMR1/C

LSM3/HNRNPR/RBM15/NCL/HTATSF1/RBM4B/SNW1/LSM8/SF1/SNRPA/SF3B3/DHX38/SDE2/CWC22/HNRNPL/FMR1/C

TRA2A/UPF1/NCBP1/RBM26/LSM1/PUS1/PAN3/SRSF12/APP/RBM42/ALYREF/CSTF2/PTBP1/SRSF9/SREK1IP1/TCE

4/TPR/PHF13/PTTG1/WRAP73/KATNB1/CHEK2/ANKLE2/CLASP1/TTK/PIBF1/MYBL2/TOM1L1/CCNE2/CCSAP/

[illegible]1/XRCC4/SFPQ/FBH1/RMI1/RFWD3/UBE2N/RMI2/TENT4A/POLQ/SMARCE1/TOPBP1/PPP4R3B/SMARCD1/DMC1/M

MEM70/STOML2/COX19/NDUFS4/LMNA/MTX1/PPP2CB/BNIP3L/SPIRE1/GABARAP/RRM1/AIFM1/APOO/UQCC3/PHB2/WIPI2/AT

[illegible]/BAZ1B/FANCA/PIN1/MRE11/CDT1/SLX4/EGF/GOLGA2/LCMT1/CLASP2/MTBP/FBXO43/MND1/NME6/ORC4/TP

:CDC22/VPS35L/SEC16A/CUL3/MPPE1/LAMP1/EXOC5/CNST/GOPC/COG7/SEC24B/NSF/NAPA/KLHL

3/POLR2G/EIF1/SAMD4B/SRP9/EEF1E1/MAGOH/GATC/TRNT1/DAPK3/INPP5E/N

KNTC1/CHEK1/MDM2/IER3/KHDRBS1/ERCC2/CUL4B/NFIB/NSMCE2/DBF4/BRD4/NAE1/DONSON/RRM2/ANAPC15/

P4/DNAJC3/TPR/UBA52/C6orf52/YTHDF3/POLR2G/EIF1/SAMD4B/SRP9/EEF1E1/M

CRC2/POR/UQCC3/PHB2/GLRX2/ATP5F1D/NDUFA10/UQCRQ

UBE3D/KLHL15/ZFAND2A

)5/ANKRD17/BRCA2/KNTC1/CHEK1/MDM2/IER3/DOT1L/ATRIP/ERCC2/CUL4B/NSMCE2/DBF4/BRD4/NAE1/DONSON/RRM

UGGT2/TRIM13/PLAA/TRAF4/BCAP31/DNAJB2/ECPAS/FBXW11/RNF185/SOCS6/RNF10/HSPBP1/SUMO2/TM

2/TTK/MDC1/LSM10/CPSF3/TOM1L1/CCNE2/RDX/LSM11/INCENP/ZW10/FBXO31/DAPK3/CDK11B/RPA2/DBF4B

PS37C/ATG4D/RALB/TMEM39A/PEX5/ATP6V1D/TOM1/ATG4C/HTT/ARFIP2/ATP6V0D2/RAB7A

N2/STAM/WASL

2/PRPF19/EZH2/GNB1L/NPR2/RNF167/PSMG2/SYF2

3CL7B/RNF185/AJUBA/OXR1/FBXO4/MSX1/USP1/PPP4C/SMARCB1/TMEM161A/DDIAS/KDM1A/ATAD5/BRCA2/

APK3/INPP5E/NAT10/SPOUT1/SECISBP2L/CDK5RAP1/SESN2/STK35/TNKS1BP1/AJUB

ATNB1/CLASP1/SNAPIN/MAP6D1/RDX/CCSAP/MAP4/DFFB/LAMP2/TRNT1/ATG2A/WDR1/STMN1/TFIP11/TGFBR1/L

88/HDAC3/SEC61G/BRCA2/FLNA/STK3/DMAP1/TOMM40L/NUP107/AP3B1/BAP1/NPEPPS/IPO8

C5/PAN2/MDN1/EDC3/PUM2/ARHGEF5/CSRP2/PRKD1/HAUS2/CCP110/NOA1/YTHDF1/NUMA1/ABRAXAS1/EIF1AX/TUB

XOSC3/EXOSC9/DHX36/SRSF4/MLH1/ELAVL4/RNASEL

O4/MCIDAS/USP1/PPP4C/SMARCB1/TMEM161A/KDM1A/NVL/ATAD5/ANKRD17/PARP1/OOEP/DMAP1/TTF1/CHEK1/I

4B/FGB/KLK1/FOXC2/SCN1B/TJP3/TACR1/TNNI2/F5/FXYD1/SUCNR1/SLC8A1/TNNT2/CXCR4/PDE2A/CD38/SH3GL2/T

ANK1

[illegible]2W1/OR2C1/OR7G3/OR5A1/OR10H1/OR5K2/EYS/OR5M9/OR2AG1/OR10S1/OR10A6/OR6P1/OR4D11/OR

2L1/OR51D1/OR4D5/OR4E2/OR1D5/OR2AT4/OR52W1/OR2C1/OR7G3/OR5A1/OR10H1/OR5K2/SLC6A3/

A6/OR6P1/B2M/OR4D11/OR4C3/OR52B4/OR5AN1/OR6S1/OR51F1/OR52M1/OR52H1/OR8B2/OR7D2/OR

3/OR52B4/OR5AN1/OR6S1/OR51F1/OR52M1/OR52H1/OR8B2/OR7D2/OR1N1/OR5L1/OR6C70/OR2AJ1/C

RD11/IL33/TLR7/LTB/IL18/IRF8/CLU/CLEC9

1/OR10S1/OR10A6/OR6P1/OR4D11/OR4C3/OR52B4/OR5AN1/OR6S1/OR51F1/OR52M1/OR52H1/OR8B2

F1/OR52M1/OR52H1/OR8B2/OR7D2/OR1N1/OR5L1/OR6C70/OR2AJ1/OR5B12/OR2D3/OR12D2/OR51T1

:3/OASL/IL2RG/CD24/XCR1

K1

SF9/IL7R/IL18/IRF8/RHOH/CCR6/FCRL3/IKZ

2/ITK/IL4I1/CD27/CLEC4C/SIT1/IGHA2/THEMIS/PTPN6/TRBC

P5
